# How to account for the uncertainty from standard toxicity tests in species sensitivity distributions: an example in non-target plants

**DOI:** 10.1101/2020.07.02.183863

**Authors:** Sandrine Charles, Dan Wu, Virginie Ducrot

## Abstract

This research aims to account for the uncertainty on 50% effective rates (ER_50_) in species sensitivity distribution (SSD) analyses and to study how including this uncertainty may influence the 5% Hazard Rate (HR_5_) estimation. We explored various endpoints (survival, emergence, shoot dry weight) for non-target plants from seven standard greenhouse studies that used different experimental approaches (vegetative vigour vs. seedling emergence) and applied seven herbicides at different growth stages. Firstly for each endpoint of each study, a three-parameter log-logistic model was fitted to experimental toxicity test data for each species under a Bayesian framework to get a posterior probability distribution for ER_50_. Then in order to account for the uncertainty on the ER_50_, we explored two censoring criteria to censor ER_50_ taking the ER_50_ distribution and the range of tested rates into account. Based on dose-response fitting results and censoring criteria, we considered input ER_50_ values SSD analyses in three ways (only point estimates chosen as ER_50_ medians, interval-censored ER_50_ based on their 95% credible interval and censored ER_50_ according to one of the two criteria), by fitting a log-normal distribution under a frequentist framework to get the three corresponding HR_5_ estimates. We observed that SSD fitted reasonably well when there were at least six distinct ER_50_ values. By comparing the three SSD curves and the three HR_5_ estimates, we found that propagating the uncertainty from ER_50_ and including censored data into the SSD analysis often leads to smaller point estimates of HR_5_, which is more conservative in a risk assessment context. In addition, we recommend not to focus solely on the point estimate of the HR_5_, but also to look at the precision of this estimate as depicted by the 95% confidence interval.

## Introduction

Today, Species Sensitivity Distributions (SSD) are established as a key tool for the environmental risk assessment (ERA) of chemicals [1]. They provide a reliable assessment of the range of sensitivities within a plant or animal community of interest and thereby allow to estimate indicators such as the 5% hazard concentration/rate (HC_5_/HR_5_) that is the hazardous concentration/rate prone to affect 5% of the species within the community. An estimation of the HC_5_/HR_5_ can be obtained from the fit of a probability distribution on a collection of toxicity values, such as 50% effective concentrations/rates (ER_50_/ER_50_), thus requiring performing a statistically robust analysis. Toxicity values are usually derived from a regression model fitted on toxicity test data observed at several treatment levels at a target time point. This fit provides toxicity values as point estimates, but an uncertainty can also be associated to them, either through a confidence interval (under a frequentist framework) or a credible interval (under a Bayesian framework). Nevertheless, this uncertainty is rarely accounted for in HR_5_ estimates afterwards, what motivated our study.

The SSD method is largely used in the field of non-target terrestrial plant (NTTP) studies, for the purpose of assessing the risk of plant protection products to NTTP [2]. NTTP are defined as non-crop plants located outside the treatment area according to the Guidance Document (GD) on Terrestrial Ecotoxicology [2]. In the case of NTTP studies, treatment levels or exposure concentrations are rather called “tested rates”, corresponding to application rates in field. Subsequently, we will use the notations HR_5_, ER_50_ hereafter.

For the SSD analyses, up to 10 species are usually studied for the ERA of a pesticide. The used tested rates are selected prior to the experiments, sometimes the same for some of or all the chosen species. The highest tested rate usually corresponds to the highest authorised application rate of the herbicide in the field, which ensures the realism of the ecotoxicological evaluation towards agricultural practices. Iowever, this highest tested rate might be too low to elicit large toxic effects (i.e., close to 100% effect, as it is classically done when setting the highest dose for a dose-response analysis for animals) especially for fungicides and insecticides. This specific point will be discussed in our paper. Unbounded right-censored ER_50_ values (namely ER_50_ greater than the highest tested rate) can occur when the range of tested rates does not really match the observed sensitivity of a plant species or when this species is not affected at the highest tested rate intended according to good agricultural practices. Note that such unbound ER_50_ values may have been produced using a validated standard experimental procedure, so that there is no reason to question them. Additionally, the GD on Terrestrial Ecotoxicology does not provide any advice on how to deal with unbound ER_50_ values or with the uncertainty associated with the ER_50_ estimates when performing SSD analyses [2]. As a consequence, the common practice is first to ignore the uncertainty by considering point estimates only (usually the mean estimate), and second either to discard unbound ER_50_ values from the analyses or to substitute them with arbitrary values (*e.g*., the highest tested rate), even if rarely done in practice. Nevertheless, performing in such a way is a clear loss of valuable information with some drawbacks. Ignoring uncertainty prevents to account for low (i.e., in the lower tail of their probability distribution) or high (i.e., in the upper tail of their probability distribution) ER_50_ values that are statistically probable, thus leading to potentially biased HR_5_ estimates (*i.e*., either over- or under-estimated values). Discarding unbound ER_50_ values may derive in (i) a range of remaining ER_50_ values that may not cover the full range of sensitivities as originally displayed in the set of the chosen species (the most sensitive or the most tolerant species may for example be excluded, thus producing biased HR_5_ estimates, either over- or under-estimated); (ii) unbound ER_50_ values can occur for many species, so that, after discarding them, the input data set might not be sufficient to allow an SSD analysis to be performed. This latter issue is of great concern for risk assessment, since the SSD analysis is currently the only higher tier option prescribed by the GD on Terrestrial Ecotoxicology and widely accepted by authorities. Thus, being unable to finalise an SSD analysis may prevent refining the risk assessment of some chemical substances. The GD on Terrestrial Ecotoxicology indeed considers that the SSD analysis is more suitable than a tier-1 approach (based on a single endpoint and a single species) to achieve the environmental protection goal because it takes into account the available data on the sensitivity of several species [2]. Moreover, substituting unbounded ER_50_ with arbitrary values would be a fairly arbitrary way that does not make much sense and leads to the possibility to produce biased HR_5_ estimates subjectively.

Within this context, based on seven NTTP case studies, we aimed to revisit SSD analyses by accounting for both the uncertainty on ER_50_ values (referred to as interval-censored values hereafter) together with the inclusion of censored values, in particular right-censored values, which happen to be commonly observed from toxicity tests in practice. Indeed, left-censored values are rare because the tested rate range as imposed by the standard protocols is better adapted to assess effects for the more sensitive species. We also tried to quantify how both types of censored values may influence the final estimate of the HR_5_.

## Materials and methods

### Materials

Laboratory experiment data sets on NTTP were available for seven studies on products with various herbicidal mode of action (Table 1). Each study provides data from two toxicity tests: seedling emergence (SE) according to OECD guideline 208 [3] and vegetative vigour (VV) according to OECD guideline 227 [4]. For each study, 10 species (thereafter named using their EPPO code [5]; see S1 Table for common names of species) were exposed to a range of five tested rates of a product plus a control (i.e., absence of product), which were applied either to the soil surface (SE test) or directly to the plants (VV test). Besides, in study 4, extra experiments at lower tested rates were conducted for two of the species (CUMSA and LYPES) in the VV test, and for study 7, extra experiments at lower tested rates for two of the species (ALLCE and BEAVA) were carried out in the SE test.

**Table 1.** Brief description of the seven studies.

| Study | Product<sup>1</sup> | Tested species (EPPO code)<sup>2</sup> | Tested rate unit |
| --- | --- | --- | --- |
| study 1 | product 1 | ALLCE AVESA BEAVA BRSNW CUMSA<br>GLXMA HELAN LYPES TRZAW ZEAMA | ml product/ha |
| study 2 | product 2 | ALLCE AVESA BEAVA BRSNW CUMSA<br>GLXMA HELAN LOLPE LYPES ZEAMA | g a.s./ha |
| study 3 | product 3 | ALLCE BEAVA BRSNW CUMSA FAGES<br>GLXMA LOLPE LYPES TRZAW ZEAMA | ml product/ha |
| study 4 | product 4 | ALLCE AVESA BEAVA BRSNW CUMSA<br>GLXMA HELAN LYPES TRZAW ZEAMA | ml product/ha |
| study 5 | product 5 | ALLCE AVESA BEAVA BRSNW CUMSA<br>GLXMA HELAN LOLPE LYPES ZEAMA | g a.s./ha |
| study 6 | product 6 | ALLCE AVESA BEAVA BRSNW CUMSA<br>GLXMA HELAN LYPES TRZAW ZEAMA | g product/ha |
| study 7 | product 7 | ALLCE AVESA BEAVA BRSNW CUMSA<br>GLXMA HELAN LYPES TRZAW ZEAMA | g product/ha |

<sup>1</sup> See S2 Table for formulations of active substances of the seven products.
<sup>2</sup> EPPO: European and mediterranean Plant Protection Organization; see S1 Table for corresponding species and common names to the EPPO code.

During experiments, plants were observed for 21 days after day 0. Day 0 is defined as the day at which 50% of the control seedlings have emerged for SE tests and as the day of application for VV tests. During the 21-day observation period, seedling emergence, seedling survival and visual injury in each replicate were followed weekly (at days 0, 7, 14 and 21) in SE tests, while plant survival and visual injuries were followed weekly in VV tests, also in each replicate. At the end of the experiments (in both SE and VV tests), shoots were cut-off and dried up, then the shoot dry weight was measured in each replicate. For each study, five endpoints at day 21 were thus available: emergence, survival and shoot dry weight for SE tests, survival and shoot dry weight for VV tests.

### Methods

To assess the effects of the studied herbicides on NTTP, we first analysed the effects of the seven products on the five endpoints (that is a total of 7 × 5 × 10 data sets) by fitting a dose-response model to experimental toxicity test data thus getting ER_50_ estimates for each endpoint of each of 10 species. The modelling process was carried out under a Bayesian framework, which ensures to get a posterior probability distribution for the ER_50_ which can then be used as a basis to quantify the uncertainty on the ER_50_. Then, these ER_50_ values, also considering their uncertainty, were used as inputs for the SSD analyses leading to the HR_5_ estimates.

#### Dose-response model

For SE tests, observed data at day 21 for replicate i can be described as 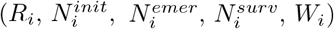, where *R_i_* is the tested rate, 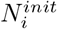 the number of sown seeds, 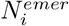 the number of emerged seedlings, 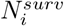 the number of surviving seedlings and *W_i_* the average shoot dry weight of surviving seedlings. For VV tests, observed data at day 21 for replicate i can be described as 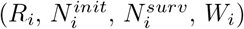, where *R_i_* is the tested rate, 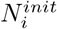 the number of initial plants, 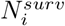 the number of surviving plants and *W_i_* the average shoot dry weight of surviving plants.

The number of emerged seedlings (SE test) and the number of surviving seedlings/plants (SE and VV tests) follow a binomial distribution, with an emergence probability (resp. a survival probability) as a function of the tested rate (see Eq (1), (2) and (3)):

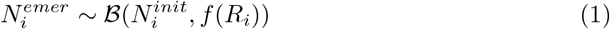

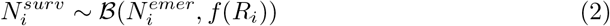

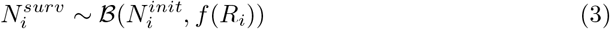

Assuming that *W_i_* is normally distributed with mean *μ_i_* and standard deviation *σ*, with *μ_i_* defined as a function of the tested rate, we get:

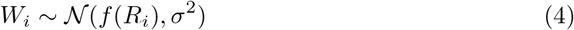

In Eq (1) to (4), *f* was chosen as three-parameter log-logistic function:

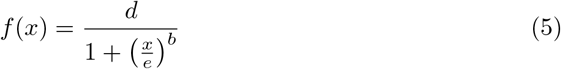

Parameters *b, d* and *e* are positive. Parameter *b* is a shape parameter translating the intensity of the effect, d corresponds to the endpoint in control data (i.e., in absence of product) and *e* corresponds to the ER_50_. Within the Bayesian framework, we have to specify a prior distribution for model parameters b, d, e (and *σ* in case of modelling shoot dry weight data). The prior distributions are given in Table 2.

**Table 2.** Specification of prior distributions for model parameters.

| Emergence/survival |  | Shoot dry weight |  |
| --- | --- | --- | --- |
| Parameter | Prior distribution | Parameter | Prior distribution |
| $\log_{10}b$ | $\mathcal{U}(-2, 2)$ | $\log_{10}b$ | $\mathcal{U}(-2, 2)$ |
| $d$ | $\mathcal{U}(0, 1)$ | $d$ | $\mathcal{U}(0, dmax)^1$ |
| $\log_{10}e$ | $\mathcal{N}(\mu, \sigma)^2$ | $\log_{10}e$ | $\mathcal{N}(\mu, \sigma)^2$ |
| | | $\sigma$ | $\mathcal{U}(0, dmax/2)^1$ |

<sup>1</sup> $dmax$ equals twice the highest observed shoot dry weight for the species under consideration. The observation with the highest observed shoot dry weight is excluded from the data set before running inference.
<sup>2</sup> $\mu = \frac{\log_{10}(maxR) + \log_{10}(minR)}{2}$ and $\sigma = \frac{\log_{10}(maxR) - \log_{10}(minR)}{2}$ , where $minR$ and $maxR$ are the lowest and the highest tested rates, respectively.

##### Estimation of parameters

Model computations were performed in R [6] with JAGS using Gibbs sampling via Markov Chain Monte Carlo (MCMC) simulation [7]. The R-package “morse” [8] was used to analyse emergence and survival data. In package “morse”, if no inhibition of plant emergence (or no survival) is observed in control groups, parameter d is automatically set to 1 by default. Hence a two-parameter log-logistic model is fitted to the data. Emergence and survival data can partically be analysed with the MOSAIC platform [9]. A modus operandi is provided in S1 Appendix. A specific R-code based on the R-package ‘rjags’ was built to fit shoot dry weight data. This code is made freely available through an R-shiny web application (https://mosaic.univ-lyon1.fr/growth), for reproducibility of the results for shoot dry weight data.

Three chains were run firstly for an intialization phase of 3000 iterations and a burn-in phase of 5000 iterations, then Ratery and Lewis’s Diagnostic was used to set the necessary thining and the number of iterations to reach a given level of precision in posterior samples. These posterior samples allows to get a joint posterior probability distribution as well as marginal posterior probability distributions for all model parameters.

#### Censoring ER_50_ estimates to account for the uncertainty

The output of interest from the previous dose-response analyses consists of the posterior probability distribution of the ER_50_ (Fig 1A) allowing to quantify the uncertainty on the ER_50_ estimation summarized as a 95% credible interval (CI95), representing the range of values within which the ER_50_ has 95% of chance to be found. Hence, we considered the use of CI95 of ER_50_ estimates as inputs of SSD analyses instead of point estimates (median/mean values) as a good way to account for the uncertainty on the ER_50_ estimates into subsequent analyses. But the following questions then arise: should we always consider the bounds of CI95 as the uncertainty limits of the ER_50_ and use this interval as an ER_50_ input value for the SSD analysis? Is the ER_50_ estimate always precise enough to be used as it is in the SSD compared to the range of tested rates? How to account for the precision of the ER_50_ estimate regarding the range of tested rates? Is there any situation for which the estimated ER_50_ should be considered as right-censored?

**Fig 1.**
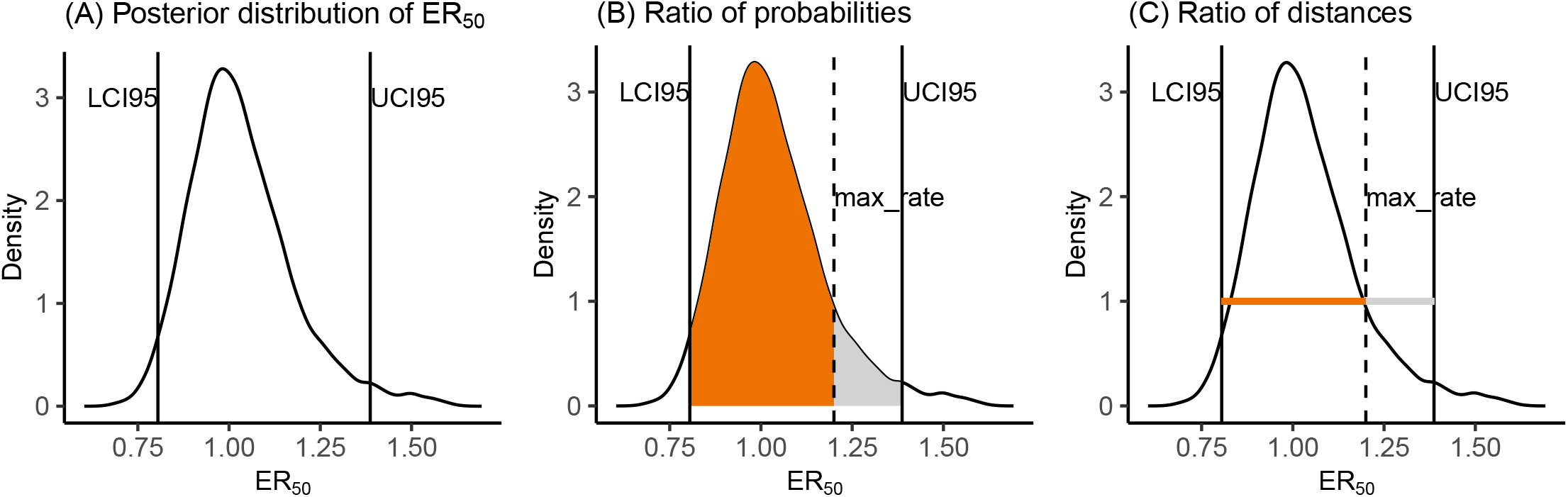
Example of posterior probability distribution of ER_50_ (A) and calculation of both censoring criteria (B-C).

To ensure agronomic realism, the common practice in standard toxicity tests with NTTP is to use the maximal field application rate as the maximum tested rate in the experimental design. However, some species are not affected or only slightly affected at the highest intended application rate: the application rates that would be needed to create high effects in the plants are unknown, thus leading to unbounded values of ER_50_ estimates (greater than the highest tested rate). Such ER_50_ estimates will usually be not precise (having a large CI95) which may suggest to rather consider them as a right-censored value. That’s why we carefully considered the relevance of the ER_50_ estimates (quantified through its CI95) regarding the range of tested rates, in particular the highest tested rate (*max_rate*). In order to decide on the most appropriate mathematical option for right-censoring the ER_50_, we propose two criteria based on overlapping ratios between [LCI95; *max_rate*] and [LCI95; UCI95], where LCI95/UCI95 are the lower/upper bound of the CI95, respectively:

1. A first criterion based on a ratio of probabilities (denoted C_1_, Eq (6)) defined as the ratio of the probability that the ER_50_ lies within [LCI95; *max_rate*] over the probability that the ER_50_ lies within [LCI95; UCI95]; as illustrated on Fig 1B, criterion C_1_ is calculated as the ratio of the orange surface divided by the (orange + grey) surface.

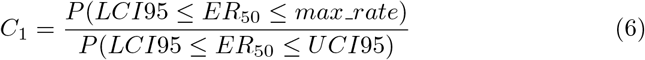
2. A second criterion based on a ratio of distances (denoted C_2_, Eq (7)) defined as the ratio of the distance *max_rate* - LCI95 (if *max_rate* LCI95, then the distance is set to 0) over the extend of the CI95; as illustrated on Fig 1C, criterion C_2_ is calculated as the ratio of the orange segment divided by the (orange + grey) segment.

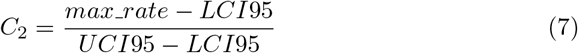

##### Decision

Once the criterion is calculated, we need a decision threshold (denoted T) to right-censor or not the ER_50_. If the ratio is greater than T, we keep an interval-censored ER_50_ corresponding to the whole CI95; otherwise, we consider a right-censored ER_50_ with a lower bound being the minimum between LCI95 and *max_rate* (Eq (8)):

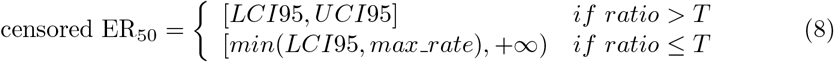

#### SSD analyses

Our final objective is to explore the influence of considering the uncertainty on ER_50_ in SSD analyses and specifically its impact on HR_5_ estimates. Given the way we have taken the uncertainy on ER_50_ into account (see above), this means studying how interval- and/or right-censored ER_50_ input values impact the SSD analysis and the HR_5_ estimation. Thus SSD analyses were carried out based on ER_50_ values coming from the seven studied firstly analysed with a dose-response model as previously described. For each experimental study and each endpoint, based on dose-response fitting results, we considered input ER_50_ values for an SSD analysis in the different following ways:

i. only point estimates (chosen as the medians of probability distributions of the ER_50_ estimates).
ii. interval-censored ER_50_ based on their CI95, and use of these intervals as such in a mathematically sound way;
iii. censored ER_50_ according to criterion 1 with a decision threshold T = 0.5 (denoted C1T0.5), and use of these censored ER_50_ as such in a mathematically sound way.

SSD analyses were run by fitting a log-normal probability distribution to ER_50_ input values under a frequentist framework based on the R-package ‘fitdistrplus’ [10]. This R-package allows the user to deal with censored data in a mathematically sound way. An alternative way is to use the web platform MOSAIC and its SSD module https://mosaic.univ-lyon1.fr/ssd [11].

## Results

### Dose-response analyses

All results on dose-response analyses are displayed in files “report_xxx.pdf” in S1 Archive for each case study, each species and each endpoint (five files per case study). Under a Bayesian framework, whatever the data set, the species and the endpoint, we always succeeded in fitting a dose-response curve and getting a posterior probability distribution on the ER_50_. For certain endpoints in certain studies, we got a well-shaped sigmoidal dose-response curve with a median estimate of the ER_50_ within the range of tested rates. Nevertheless, in cases where the herbicides did not elicit a strong effect on the studied species, we got a flat dose-reponse curve with a high median estimate of the ER_50_, in particular for the survival endpoint of VV tests.

Fig 2 illustrates an example of a dose-response curve along with some goodness-of-fit criteria. The data we used for this example is the shoot dry weight of the VV test of study 1 for species BEAVA. The median fitted dose-response curve in Fig 2A is represented by a solid orange line associated with its CI95 in grey band; it describes the shoot dry weight of the sugar beet as a function of the product tested rate. The goodness-of-fit for the fitted model can be checked using posterior predictive check (PPC) plot illustrated in Fig 2B. The PPC plot shows the observed shoot dry weight values against their corresponding shoot dry weight predictions (black dots), along with their CI95 (vertical segments, green if the CI95 contains the observed value and red otherwise). The model is considered to be well fitted if around 95% of black dots are within CI95. Please note that for the emergence and survival datasets, the previous statement is not necessarily expected because observations are pooled per tested rate. The precision of the model parameter estimation can be visualized in Fig 2C by comparing the posterior distribution (orange surface) to the prior one (grey surface) for each parameter and in Fig 2D, we can check for correlations between parameters. A narrower posterior distribution compared to the prior one for each parameter and low correlations between parameters are expected to ensure the goodness-of-fit of the model; that is the case in this example.

**Fig 2.**
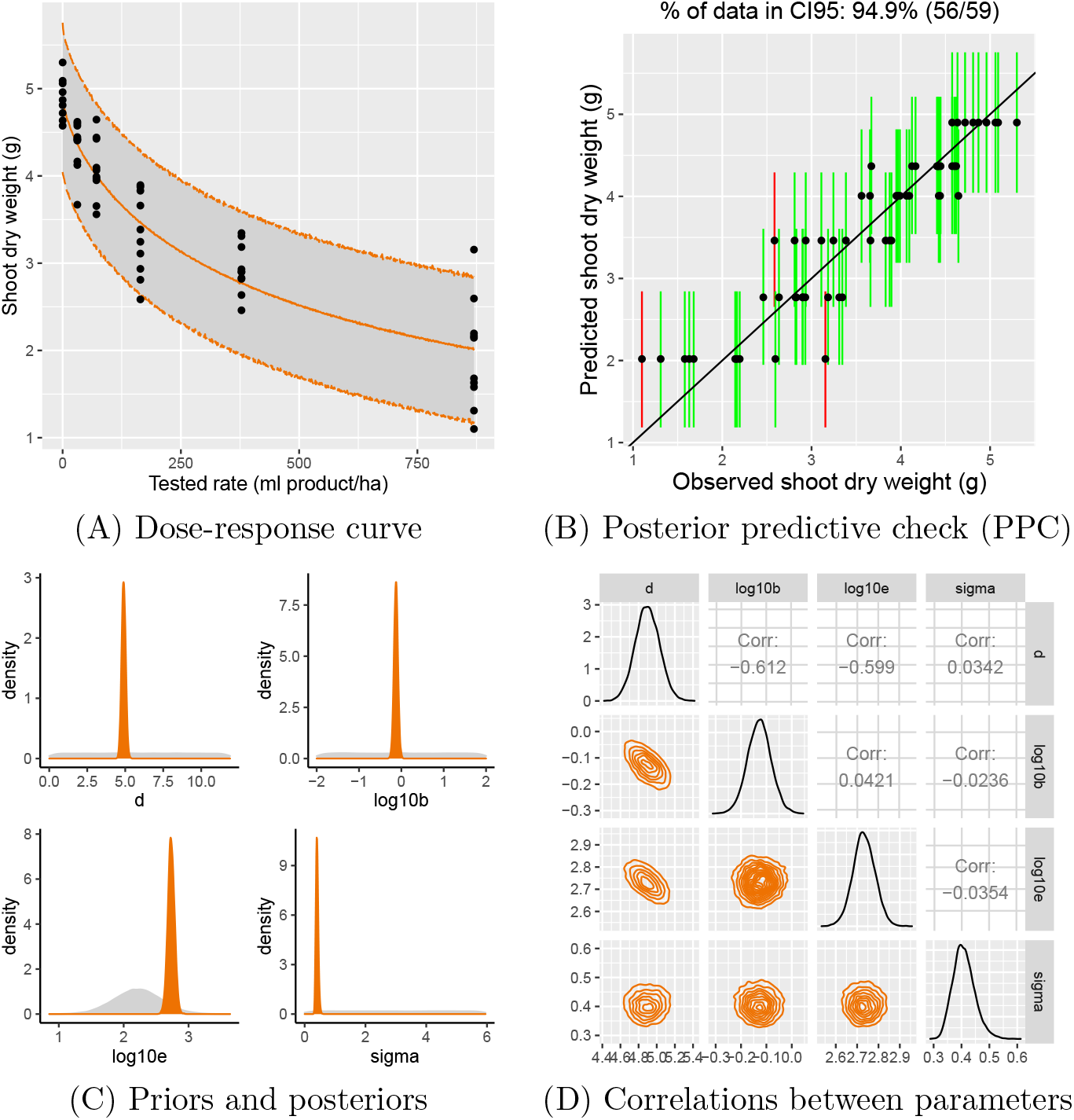
Example of a dose-response curve (A), posterior predictive check (B), prior and posterior distributions of parameters (C) and correlations between parameters (D).

### Censoring on ER_50_ estimates

The censoring decision for an ER_50_ depends on both the criterion (C_1_ or C_2_) and the decision threshold T. To study the influence of the criterion and the decision threshold on censoring decisions, we tried seven T values: 0, 0.2, 0.4, 0.5, 0.6, 0.8, 1.0, with each criterion. Fig 3 provides an example of censored ER_50_ values obtained according to both criteria and the seven decision thresholds. The data we used for this example is the shoot dry weight of the VV test of study 1 for species ALLCE.

**Fig 3.**
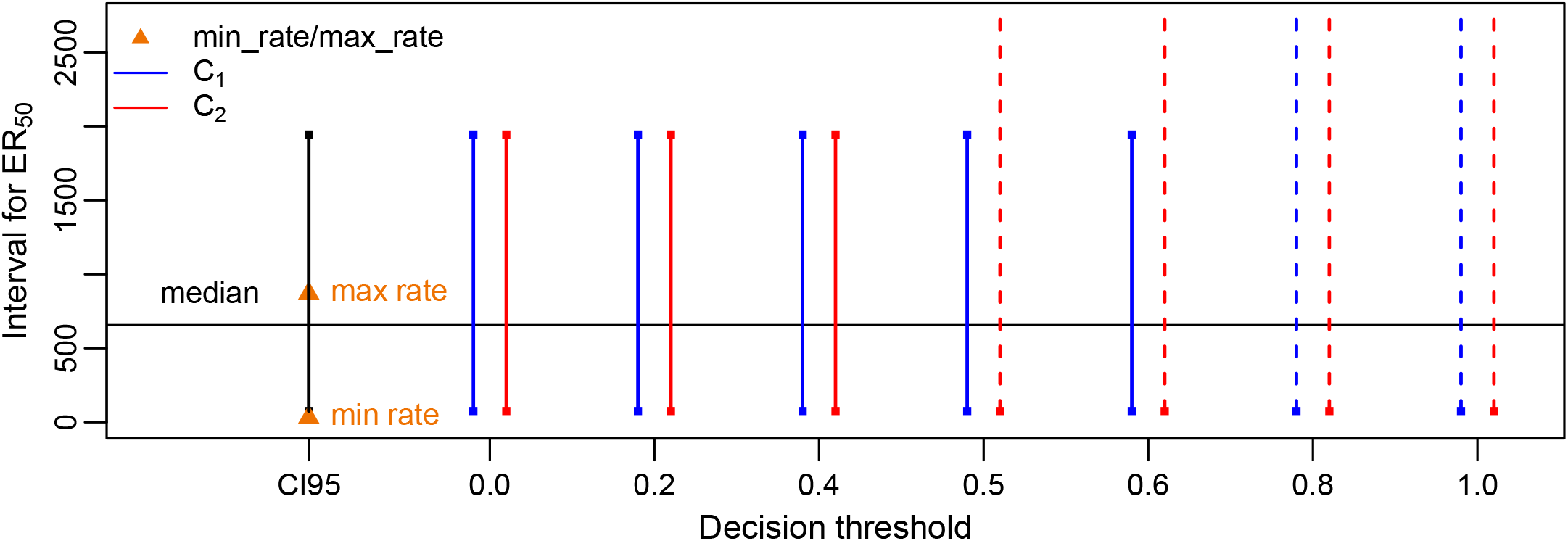
Censored ER_50_ according to both criteria and the seven decision thresholds for the shoot dry weight endpoint of the VV test of study 1 - species ALLCE. The two orange triangles stand for the lowest and the highest tested rates. Vertical segments of three different colors (black, blue, red) represent the CI95 of ER_50_, the censored ER_50_ according to C_1_ and the censored ER_50_ according to C_2_, respectively; solid vertical segments are for bounded intervals while dotted vertical segments stand for right-unbounded intervals; the black horizontal line represents the median of the ER_50_ estimate.

In this example, the ER_50_ is either interval-censored or right-censored depending on the criterion and the T value. Moreover, censored ER_50_ values vary slightly according to the criterion and the decision threshold. Most of the time, criteria C_1_ and C_2_ lead to the same censoring decision for our seven case studies, thus criterion C_1_ (based on the whole probability distribution of the ER_50_) was finally preferred. See files

“ER_50__censoring.pdf” in S1 Archive (seven files in total) for results on other species and other endpoints. Regarding the decision threshold T, in the following cases, the seven decision thresholds led to the same censoring decision:

- case 1: when the CI95 of the ER_50_ is utterly within the range of tested rates, an ER_50_ interval-censored by its CI95 is always obtained;
- case 2: when the CI95 of the ER_50_ is utterly out of the range of tested rates, a right-censored ER_50_ [*max_rate*, +∞) is always obtained.

Consequently, the decision threshold influences the censoring decision when there is an overlap between the CI95 of the ER_50_ and the range of tested rates. In this case, the higher T is, the more often we will decide to right-censor the ER_50_. Hence, in certain cases, a too high T value may generate a lot of right-censored ER_50_ values and lead to consider some ER_50_ estimates right-censored while we would have rather prefered to use their CI95 to quantify their uncertainty. On the other hand, a too low T value may almost always lead to decide to use an interval-censored ER_50_ with its CI95, even in cases where most of the possible values for the ER_50_ estimate within the support of its posterior probability distribution are greater than the highest tested rate, while in such a case, we would have rather considered to right-censor it regarding the range of the tested rates. Therefore, we have a preference for T = 0.5, as a neutral value. Hence, for the subsequent SSD analyses, we will consider only censored ER_50_ values according to C_1_ and T = 0.5 (C1T0.5).

### SSD and HR_5_

Three ways of handling ER_50_ values in SSD analyses were studied and compared for the seven case studies. For each case study, all results on SSD and HR_5_ are displayed in files “SSD_analyses.pdf” in S1 Archive (seven files in total). In total, we did SSD analyses on 105 data sets (7 studies × 5 endpoints × 3 types of ER_50_). We had almost no convergence problem for parameter estimation, except for the survival endpoint of the VV test for studies 2 and 5. We had almost no convergence problem for parameter estimation, except for the survival and shoot dry weight endpoints of the VV test for studies 2 and 5. In the case of study 2 for the survival endpoint, fitting a log-normal distribution to the data set with censored ER_50_ values according to C1T0.5 failed because the 10 censored ER_50_ values were in fact equal to the same interval [*max_rate*, +∞). In other cases, convergence failed because all censored ER_50_ values were too close from each other.

Fig 4 illustrates an example of three SSD analyses based only on medians of ER_50_ (A), ER_50_ interval-censored by their CI95 (B) and ER_50_ censored according to C1T0.5 (C). The obtained HR_5_ results are denoted by HR_5,1_, HR_5,2_ and HR_5,3_, respectively. The data for this example is the shoot dry weight of the VV test of study 1. In this example, the three SSD curves fitted well to ER_50_ values. The SSD curve in Fig 4C has a larger 95% confidence interval (CoI95) than the ones in Fig 4A and Fig 4B. Estimated HR_5,3_ was smaller than HR_5,1_ and HR_5,2_ with a larger CoI95.

**Fig 4.**
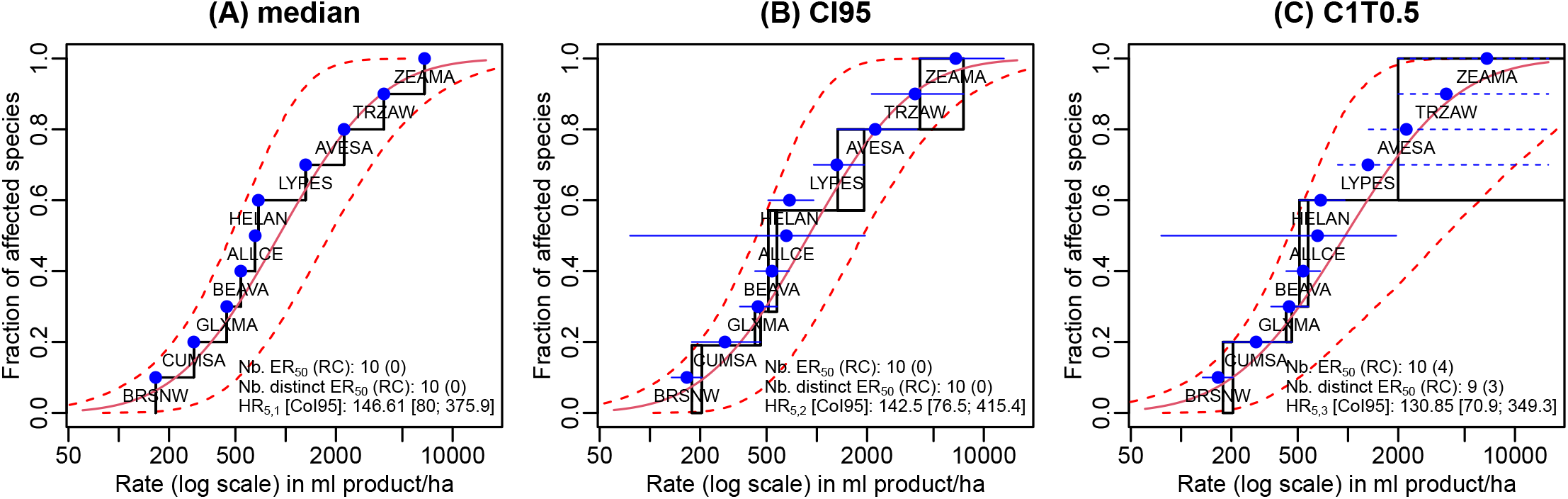
Example of three SSD analyses based respectively on ER_50_ medians, CI95 of ER_50_ and censored ER_50_ according to C1T0.5 for the shoot dry weight endpoint of the VV test of study 1. Blue points represent fractions of affected species (EPPO code) ordered by the median of their ER_50_. Solid blue horizontal segments stand for interval-censored ER_50_ by their CI95 and dotted ones stand for right-censored ER_50_. Solid red curves represent fitted SSD curves and dotted ones represent 95% confidence interval (CoI95) of the fitted SSD curves. Meaning of legends: Nb. ER_50_ (RC) stands for the number of ER_50_ (the number of right-censored ER_50_); Nb. distinct ER_50_ (RC) stands for the number of distinct ER_50_ (the number of distinct right-censored ER_50_); HR_5_ [CoI95] stands for the estimated HR_5_ with its CoI95.

#### Summary from the seven case studies

Results on HR_5_ based on the three ways of handling ER_50_ values for the seven data sets are given in Table 3 and corresponding SSD curves are given in files “SSD_analyses.pdf” in S1 Archive.

**Table 3.** Comparison of results on HR_5_ based on different handling criteria for input ER_50_ values.

| Study | Endpoint | Median |  | CI95 |  | C1T0.5 |  |  |
| --- | --- | --- | --- | --- | --- | --- | --- | --- |
| | | Nb.(RC)<sup>1</sup> | $HR_{5,1}[CoI95]^2$ | Nb.(RC)<sup>1</sup> | $HR_{5,2}[CoI95]^2$ | Nb.(RC)<sup>1</sup> | $HR_{5,3}[CoI95]^2$ | CV<sup>3</sup> |
| 1 | Emergence (SE) | 10 (0) | 832.1 [694.6; 1165] | 10 (0) | 1581 [1245; 2211] | 5 (4) | 2260 [NA; NA] | NA |
| 1 | Survival (SE) | 10 (0) | 177.7 [98.08; 460.6] | 10 (0) | 172.2 [93; 465.5] | 8 (2) | 146.4 [78.27; 412.6] | 0.6200 |
| 1 | Shoot dry weight (SE) | 10 (0) | 126.4 [84.87; 257.1] | 10 (0) | 125.6 [84.2; 259.5] | 9 (1) | 120.5 [80.6; 253.8] | 0.3216 |
| 1 | Survival (VV) | 10 (0) | 787 [703.1; 1069] | 10 (0) | 1184 [922.6; 2458] | 3 (3) | 8697 [NA; NA] | NA |
| 1 | Shoot dry weight (VV) | 10 (0) | 146.6 [79.96; 375.9] | 10 (0) | 142.5 [76.49; 415.4] | 9 (3) | 130.9 [70.93; 349.3] | 0.5326 |
| 2 | Emergence (SE) | 10 (0) | 41.82 [20.37; 151.6] | 10 (0) | 68.16 [32.16; 436.9] | 3 (3) | 721.9 [NA; NA] | NA |
| 2 | Survival (SE) | 10 (0) | 18.25 [6.061; 165] | 10 (0) | 18.89 [5.698; 115] | 5 (3) | 9.66 [2.666; 53700] | 2.8780 |
| 2 | Shoot dry weight (SE) | 10 (0) | 16.77 [5.313; 76.45] | 10 (0) | 17.62 [5.376; 90.29] | 8 (5) | 12.17 [2.489; 6972] | 5.2380 |
| 2 | Survival (VV) | 10 (0) | 169.5 [169; 271.5] | 10 (0) | 486.1 [NA; NA] | 1 (1) | NA [NA; NA] | NA |
| 2 | Shoot dry weight (VV) | 10 (0) | 193.3 [159.3; 259.4] | 10 (0) | 198.7 [173.5; 501.6] | 2 (1) | NA [NA; NA] | NA |
| 3 | Emergence (SE) | 10 (0) | 1495 [1343; 2199] | 10 (0) | 3049 [NA; NA] | 2 (2) | 6172 [NA; NA] | NA |
| 3 | Survival (SE) | 10 (0) | 985.2 [712; 1773] | 10 (0) | 937.6 [718.9; 2487] | 4 (1) | 902.4 [707.7; 60040] | 3.6700 |
| 3 | Shoot dry weight (SE) | 10 (0) | 278.5 [135; 936.6] | 10 (0) | 257 [129.6; 858.5] | 7 (1) | 220.8 [105.5; 899.9] | 0.6879 |
| 3 | Survival (VV) | 10 (0) | 998.9 [745.5; 1848] | 10 (0) | 973.2 [756.5; 2929] | 4 (1) | 970.3 [752.5; 3160] | 0.4557 |
| 3 | Shoot dry weight (VV) | 10 (0) | 156.5 [88.91; 410.6] | 10 (0) | 158.4 [90.34; 419] | 10 (3) | 135 [71.15; 360.8] | 0.4794 |
| 4 | Emergence (SE) | 10 (0) | 40.25 [5.791; 402.3] | 10 (0) | 50.42 [7.072; 644.8] | 6 (6) | 304600 [NA; NA] | NA |
| 4 | Survival (SE) | 10 (0) | 23.15 [4.837; 180.5] | 10 (0) | 25.9 [5.794; 199.2] | 10 (6) | 48.58 [26.2; 238.3] | 5.4630 |
| 4 | Shoot dry weight (SE) | 10 (0) | 12.3 [4.498; 41.74] | 10 (0) | 12.95 [3.311; 44.09] | 10 (1) | 25.25 [16.96; 53.34] | 0.3277 |
| 4 | Survival (VV) | 10 (0) | 71.47 [38.53; 188.6] | 10 (0) | 70 [35.79; 213.8] | 9 (4) | 57.05 [26.1; 243.4] | 3.2630 |
| 4 | Shoot dry weight (VV) | 10 (0) | 3.91 [2.018; 12.52] | 10 (0) | 3.944 [2.062; 12.3] | 10 (1) | 3.799 [1.681; 11.74] | 0.5550 |
| 5 | Emergence (SE) | 10 (0) | 0.3523 [0.1793; 1.282] | 10 (0) | 0.3731 [0.1895; 1.332] | 9 (3) | 0.3297 [0.1527; 1.594] | 70.6900 |
| 5 | Survival (SE) | 10 (0) | 0.3586 [0.1488; 1.138] | 10 (0) | 0.3836 [0.1551; 1.343] | 10 (3) | 0.3415 [0.1174; 1.317] | 0.6793 |
| 5 | Shoot dry weight (SE) | 10 (0) | 0.3437 [0.1509; 1.207] | 10 (0) | 0.3655 [0.1665; 1.297] | 10 (1) | 0.3405 [0.1427; 1.241] | 0.6585 |
| 5 | Survival (VV) | 10 (0) | 17.91 [13.86; 24.83] | 10 (0) | NA [NA; NA] | 3 (2) | 21.06 [21.06; 28.58] | 0.1296 |
| 5 | Shoot dry weight (VV) | 10 (0) | 12.24 [7.933; 21.78] | 10 (0) | 15.22 [10.38; 49.8] | 4 (2) | NA [NA; NA] | NA |
| 6 | Emergence (SE) | 10 (0) | 115.2 [54.57; 327.4] | 10 (0) | 113.2 [53.84; 574.3] | 3 (2) | 86.47 [47.41; 3535] | 1.2790 |
| 6 | Survival (SE) | 10 (0) | 29.9 [13; 121.2] | 10 (0) | 29.99 [12.3; 135.9] | 6 (2) | 21.32 [8.864; 133.9] | 2.1450 |
| 6 | Shoot dry weight (SE) | 10 (0) | 7.416 [4.924; 16.11] | 10 (0) | 7.47 [5.008; 16.77] | 10 (2) | 6.499 [3.96; 14.16] | 0.3548 |
| 6 | Survival (VV) | 10 (0) | 22.5 [14.43; 43.68] | 10 (0) | 22.53 [13.19; 50.85] | 6 (2) | 16.88 [9.297; 55.46] | 0.5118 |
| 6 | Shoot dry weight (VV) | 10 (0) | 3.525 [2.454; 7.914] | 10 (0) | 3.553 [2.449; 8.065] | 10 (1) | 3.487 [2.414; 8.764] | 0.4082 |
| 7 | Emergence (SE) | 10 (0) | 73.88 [61.31; 103.6] | 10 (0) | 151.7 [131.5; 235.4] | 3 (3) | 302.4 [NA; NA] | NA |
| 7 | Survival (SE) | 10 (0) | 7.758 [1.734; 67.84] | 10 (0) | 7.658 [1.659; 76.36] | 7 (2) | 5.275 [0.9925; 80.1] | 1.7270 |
| 7 | Shoot dry weight (SE) | 10 (0) | 1.062 [0.2098; 6.958] | 10 (0) | 1.553 [0.3054; 6.965] | 10 (1) | 1.401 [0.2398; 6.885] | 0.8095 |
| 7 | Survival (VV) | 10 (0) | 6.341 [3.258; 22.57] | 10 (0) | 6.516 [3.28; 25.68] | 7 (3) | 5.612 [2.314; 35.4] | 4.6280 |
| 7 | Shoot dry weight (VV) | 10 (0) | 1.448 [0.7201; 4.423] | 10 (0) | 1.402 [0.7252; 4.51] | 10 (2) | 1.152 [0.496; 3.481] | 0.5719 |

NA stands for Not Available; it may appear either when there is a problem of convergence, or when the proximity or the equality of $ER_{50}$ values leads to always bootstrapping the same set of $ER_{50}$ values thus providing equal lower and upper bounds of the CoI95. Lines coloured in gray stand for $HR_{5,3}$ poorly estimated or not-estimated based on less than six distinct $ER_{50}$ , or for only right-censored $ER_{50}$ as inputs for SSD analyses.
<sup>1</sup> Number of distinct $ER_{50}$ (number of distinct right-censored $ER_{50}$ );
<sup>2</sup> Estimated $HR_5$ [95% confidence interval];
<sup>3</sup> Coefficient of variation for $HR_5$ .

According to SSD curves, SSD were poorly fitted when there were less than six distinct ER_50_ among the 10 input ER_50_ values, most of which being right-censored (e.g., the emergence endpoint for studies 1, 2, 3, 6 and 7; the survival endpoint of the SE test for studies 2 and 3; the survival endpoint of the VV test for studies 1, 3 and 5; the shoot dry weight endpoint of the VV test for study 5). In such cases, we often found that HR_5_ estimates were imprecise with a large CoI95. The HR_5,3_ estimates taking the right censoring into account were often greater than the HR_5,1_ and HR_5,2_ estimates based on only medians of ER_50_ or based on interval-censored ER_50_ by their CI95. Please note that we performed these SSD analyses anyway to obtain HR_5_ estimates and to illustrate that HR_5_ values obtained for data sets where less than six species show clear adverse effects are not precise. Therefore, it would make no sense to fit an SSD in such cases for routine analyses.

According to SSD curves, SSD fitted reasonably well when there were at least six distinct ER_50_ values as inputs, only some of which being right-censored (e.g., the emergence endpoint for study 5; the survival endpoint of the SE test for studies 1, 4, 5, 6 and 7; the shoot dry weight endpoint of the SE test for all seven studies; the survival endpoint of the VV test for studies 4, 6 and 7; the shoot dry weight endpoint of the VV test for studies 1, 3, 4, 6 and 7). We almost always observed that the HR_5,3_ were lower than the HR_5,1_ and HR_5,2_, except for the survival endpoint of the SE test for study 4, where we had a HR_5,3_ twice greater than the HR_5,1_ and HR_5,2_; for the three endpoints of the SE test for study 5, HR_5,1_, HR_5,2_ and HR_5,3_ were very close and for the shoot dry weight endpoint of the VV test for study 7, the HR_5,3_ was a little greater than the HR_5,1_ but less than HR_5,2_.

Concerning the influence of including right-censored data or not on the HR_5_ estimate, based on SSD curves for our seven case studies, we found that for cases with an HR_5,3_ lower than the HR_5,1_ and HR_5,2_ estimates, the right-censored ER_50_ values were rather obtained for less sensitive species (i.e., species having higher median ER_50_ values). This result was found frequently among the seven case studies and most of the time in studies for which there were few right-censored ER_50_ values for measured endpoints; this reflects that such a situation will be the case in most of the cases encountered when analysing SSD data for NTTP. This comes from the fact that the tested rate range (as imposed by the standard protocols) is better adapted to assess effects on the more sensitive species. On the other hand, we found that for certain cases where the HR_5,3_ was greater than the HR_5,1_ and HR_5,2_ estimates, the right-censored ER_50_ values were not only obtained for less sensitive species but also for highly sensitive species (i.e., species having lower median ER_50_ values). This happened often in studies for which there were lots of right-censored ER_50_ values, as the application rates were not adapted for most of the species for the measured endpoints. In a risk assessment context, this situation would typically lead to new experiments with higher tested rates.

## Discussion

### Dose-response modelling

#### Model choice

There are a vast variety of models in common use to describe a dose-response relationship for ecotoxicity test data, such as probit, log-logistic, Weibull, etc. [12–14]. The log-logistic models have been widely used in weed science and they have been recommended as a standard herbicide dose-response [15]. The log-logistic models are by far the most commonly used model for describing toxicity test data [16]. The log-logistic models can be used to properly analyze not only continuous data but also quantal data. Hence, for our NTTP data, we used log-logistic models always with the same deterministic part to analyze emergence, survival and shoot dry weight data, thus facilitating comparisons.

Hence a three-parameter log-logistic model was chosen to analyze not only emergence and survival data, but also for shoot dry weight data. Although a four-parameter log-logistic model could have been tested and may be chosen for shoot dry weight data, we prefered to use the three-parameter one, since it’s reasonable to fix the lower asymptote (parameter c) at 0 considering that at really high application rates all plants can die, even though the actual data are not fully supporting this assumption for few rare cases [16]. Morever, for any of our data sets, the addition of one extra parameter did not significantly improve the model fitting (results not shown).

#### Choice of priors

A quasi-non-informative uniform prior distribution was chosen for the logarithm of parameter *b* within the interval [-2,2], in order to cover a wide variety of dose-response shapes. In certain cases, we had an extremely flat dose-response curve (no effect of any of the tested rates was observed on the endpoint) so that the posterior of parameter *b* was as wide as the prior, even if we enlarged the support prior interval. In addition, in these cases, the imprecise estimation of parameter *b* did not influence our conclusions on the ER_50_ estimates, since the ER_50_ values were considered as right-censored with their lower bound being the highest tested rate.

For parameter *e*, we used the same prior as the one used in the R-package “morse” [8,17]: a normal distribution was chosen for the logarithm of parameter *e*, with specific mean and standard deviation (presented in section method) computed from the experimental design. This choice implies that parameter *e* has a probability slightly greater than 95% to lie within the range of tested rates.

A uniform prior distribution was chosen for parameter *d* within the interval [0, *dmax*], For the emergence and survival data sets, *dmax* equals 1 representing a 100% probability of emergence or a 100% probability of survival. For the shoot dry weight data sets, *dmax* should ideally be chosen according to expert knowledge and equal to the highest expected shoot dry weight regarding the species and environmental conditions (in the experiment) under consideration. Nevertheless, for pratical convience, *dmax* was chosen as twice the highest observed shoot dry weight for the species under consideration, and then the observation with the highest observed value was discarded from the dose-response analyses.

Concerning prior distributions for the variance parameter of the Gaussian distribution (shoot dry weight data only), there are commonly used prior specifications (e.g., an inverse gamma, an inverse chi-square distribution on variance parameter, a uniform distribution on standard devation parameter). We finally assigned a uniform prior distribution to standard deviation parameter σ within the interval 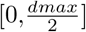.

### Censoring of ER_50_ estimates

We chose to use criterion C_1_ based on the ratio of probabilities and a T value equal to 0.5 to censor ER_50_ estimates. Indeed, criteria C_1_ and C_2_ led to very close censoring decisions for ER_50_ estimates, but criterion C_1_ was preferred since it is based on the whole probability distribution of the ER_50_. However, if criterion C_2_ had been chosen for censoring ER_50_ estimates, there would be very few changes for SSD analyses and HR_5_ estimates. Regarding decision threshold T, there is no rule set in stone for its choice. We have a preference for T = 0.5, as a neutral value. This medium value may avoid considering very imprecise ER_50_ estimates as interval-censored by their CI95 and may also avoid considering enough precise ER_50_ estimates as right-censored. Based on the 350 data sets we analysed, the same results and conclusions are almost always reached for a choice of a T value between 0.4 and 0.6. We propose to always use the intermediate value of 0.5 for the sake of simplicity and comparability. We recommend to avoid using T values below 0.4 or above 0.6, for the reasons mentioned above.

#### Uncertainty on the ER_50_ estimates

There are some limitations in the way we considered the uncertainty on the ER_50_ estimate. Here, this uncertainty was simply summarized by either an interval-censored ER_50_ with its CI95 or by a right-censored ER_50_ accounting for the range of tested rates. This method does not use all available information on ER_50_ (i.e., the full posterior distribution). It is conceivable that the uncertainty on the ER_50_ estimates could be considered in other better ways, allowing to take account of the full posterior distribution of ER_50_. Further research would be needed to explore this possibility.

### SSD analyses

Compared to the traditional deterministic approach that relies on the most sensitive individual toxicity data, the probalistic SSD method has numerous advantages [18]. As always in statistics, SSD can be built with either parametric or non-parametric methods. Some parametric distributions have already been proposed for SSD, such as log-logistic [19], log-normal [20,21], Burr Type III [22], Weibull distributions, etc.. The common use of the parametric approach for SSD is due to its mathematical simplicity and because it requires less data points compared to non-parametric approaches. Log-normal and log-logistic distributions are the most commonly used for SSD. The present paper has used a log-normal distribution to fit ER_50_ values without testing the normality of the logarithm of tested rates, since it is not our purpose to find the best fit to toxicity values, but rather to study the influence of accounting for the uncertainty on HR_5_ estimates.

Concerning minimum data requirements for fitting an SSD, a minimum of five to ten species is deemed acceptable for regulatory purposes in aquatic ecotoxicity [18] and in the context of environmental risk assessment, a minimum of six species is required [2]. Ten data points were suggested by Wheeler et al. [23] to generate reliable estimate upon which regulatory decisions may be based. In the present paper, ten species were tested for the seven case studies, allowing to collect ten ER_50_ values (generally distinct) for SSD analyses. However, for certain endpoints of certain studies, when taking into account right-censored ER_50_ values in SSD analyses, the number of distinct ER_50_ values for SSD diminished, since some of right-censored values were in fact equal to the same interval [*max_rate*, +∞), with *max_rate* equal for all the species. Consequently, we had some cases where SSD were poorly fitted with less than six distinct ER_50_ values, some of which being right-censored. When this happens, it is better not to consider SSD results and their corresponding estimated HR_5_ values, and if possible to add new experiments with higher application rates. Based on our seven cases studies, at least six distinct ER_50_ values (i.e., six distinct intervals, whatever the ER_50_ point value) appears as the minimum requirement to reasonably fit an SSD and obtain a relevant 95% confidence interval on the final HR_5_ estimate.

#### Influence of right-censored data on HR_5_ estimates

For endpoints for which the SSD was fitted on at least six distinct ER_50_ inputs, most of the time, HR_5_ estimates taking the right-censoring into account were lower than HR_5_ estimates based on medians of ER_50_ or interval-censored ER_50_ by their CI95. The results were consistent with a simulation study conducted by Green et al. [12], which demonstrated that the mathematically sound way of using censored data tends to underestimate HR_5_ compared to the true HR_5_ resulting from a data set without censored values, with greater underestimation associated with greater percentage of censoring. In addition, we found that when right-censored ER_50_ values were obtained for the less sensitive species, which is the most common case in practice, the HR_5_ values obtained by including the right-censoring were most of the time smaller than those obtained by handling censored data as non-censored data.

On a general point of view, based on our seven case studies for the five endpoints we analysed, the influence of including right-censored data on the HR_5_ estimate depends on the right-censored ER_50_ values being obtained rather on more sensitive species or on less sensitive species. In addition, we can say that, if right-censored data spread in a random way among the chosen species, the HR_5_ obtained by considering right-censored ER_50_ values can be both greater or smaller than the HR_5_ obtained by handling censored data as non-censored ones.

#### Sensitivity of endpoints

In the results section, we have not mentioned the sensitivity of endpoints. However from Table 3, we found that the shoot dry weight endpoint from SE or VV tests appeared almost always as the most sensitive endpoint with the lowest estimated HR_50_ value, except for study 5 for which HR_5_ results for three endpoints (emergence, survival and shoot dry weight of the SE test) were very close. The shoot dry weight from the VV test appeared more often as the most sensitive one (four times out of the seven case studies) than the one from the SE test. In addition, for the shoot dry weight endpoint, we had rarely right-censored ER_50_ values according to C1T0.5. Therefore, it would be recommended to always collect and analyse shoot dry weight data in order to assess risk of herbicide on NTTP by using SSD analyses based on censored ER_50_ inputs.

#### Experimental Design

In certain studies, the tested rates were not specifically adapted to the sensitivity of some species. Hence, some species were not affected or slightly affected at the highest intended application rate, leading to right-censored ER_50_ values. These right-censored values may affect the estimation of HR_5_, for example making the estimate less precise. If the precision of the HR_5_ is not considered as sufficient (i.e., with a high value of the CV), then conducting new experiments with higher application rates may help to refine the estimation of the HR_5_.

## Conclusion

The method we proposed in this paper is applicable to any taxon in ecotoxicology. The results presented here are based on a total 350 data sets consisting of seven studies, each with five endpoints (survival, emergence, shoot dry weight) for 10 non-target plants from standard greenhouse experiments that used different experimental designs (vegetative vigour vs. seedling emergence) and applied herbicides at different growth stages. The Bayesian approach allowed estimating ER_50_ values and 95% credibility intervals for all data sets, even when the dose-response curve did not reach a strong effect for the highest tested rate. Combined with a statistically sound approach for inclusion of censored ER_50_ estimates in SSD computing, we maximized the use of existing species data when building SSD, thus avoiding discarding right and/or left-censored data that may be obtained from lab studies for less or more sensitive species. Our overarching study confirmed that at least six distinct ER_50_ values (i.e., six distinct intervals, whatever the ER_50_ point value) are required as input to the SSD analysis to ensure obtaining a reliable estimate of the HR_5_. Our paper proposes a statistically sound method for propagating the uncertainty of ER_50_ estimates considered as interval-censored values towards the HR_5_ estimates. This method delivers both point estimates and bootstrap 95% confidence intervals of HR_5_. It illustrates that propagating the uncertainty from ER_50_ and including censored data as inputs for SSD analyses may change both the point estimate and the 95% confidence interval. The extend of the change depends on the characteristics of the ER_50_ input values (*e.g*., whether censored data were obtained for less/more sensitive species or were randomly spread among the tested species) and on the chosen criteria for handling the uncertainty of ER_50_ values. Consequently, when comparing and interpreting the final results, we recommend not to focus solely on the point estimate of the HR_5_, but also to look at the precision of this estimate as depicted by the 95% confidence interval. A small confidence interval stands for a precise estimate of the HR_5_, and thus a low uncertainty. This information integrates the differences in sensitivity and the uncertainty of the ER_50_ estimates across a range of species all the way down to the HR_5_ estimation: therefore, it is particularly valuable for an informed use of the HR_5_ value in the context of environmental risk assessment.

## Supporting information

Appendix-modus-operandi

Archive with reports on studies

## Acknowledgements

The authors thank the European Crop Protection Association (ECPA) for the financial support of this work.

## Supporting information

**S1 Table. Corresponding species and common names to the EPPO code.**

**S2 Table. Formulation of active substances in seven products.**

**S1 Appendix. Reproduction of results via MOSAIC.**

**S1 Archive. Zip file containing all supplementary results.** It is a zip file containing 7 folders (one folder per case study). In each folder, it contains 5 files “report_xxx.pdf” for detail results of dose-response analyses, one file corresponding to does-response analyses per endpoint. It also contains one file “ER50censoring.pdf” for censored ER_50_ and one file “SSD_analyses.pdf” for results of SSD analyses.

