## Appendix-modus-operandi for "How to account for the uncertainty from standard toxicity tests in species sensitivity distributions: an example in non-target plants"

Sandrine Charles Dan Wu Virginie Ducrot

29 June 2020

Contact:

### S1 Appendix: reproduction of results via MOSAIC

This document gives the modus operandi to reproduce the results in the paper with the MOSAIC platform (MOdeling and StAtistical tools for ecotoxiCology): https://mosaic.univ-lyon1.fr/. Several modules of MOSAIC must be used successively, and multiple times for one of them.

#### First step: dose-response analyses

In case of emergence and survival data, please use the "surv" module of MOSAIC (menu 'surv' -> 'standard': https://mosaic.univ-lyon1.fr/survival). It allows to get a posterior distribution of $ER_{50}$ from a dose-response fitting (file named 'mcmc.txt', soon available). For emergence data sets, please rename column "Nemer" into "Nsurv" when using the "surv" module of MOSAIC. Then use the following function in R to get a censored $ER_{50}$ value according to C1T0.5.

```
# distER50: distribution of ER50 (sample of ER50 values)
# max_rate: highest tested rate
censorC1T0.5 <- function(distER50, max_rate) {
  LCI95 <- quantile(distER50, probs = 0.025)
  UCI95 <- quantile(distER50, probs = 0.975)
  ratio <- sum(distER50 >= LCI95 & distER50 <= max_rate)/length(distER50)/0.95
  left <- ifelse(ratio > 0.5, LCI95, min(LCI95, max_rate))
  right <- ifelse(ratio > 0.5, UCI95, NA)
  return(data.frame(left, right))
}
mcmc <- read.table("mcmc.txt", header = T, sep = "\t")
result <- censorC1T0.5(10^mcmc[, "log10e"], max_rate) # max_rate: highest tested rate
```

Exception: if no inhibition of plant emergence (or no survival) is observed for any tested rate, an "internal error" may be displayed. In such a case, you must use R directly and the R-package 'Morse'. The corresponding R-script can be copied and pasted from MOSAIC (bottom of result page) when running on another data set not showing this internal error. You will just have to change the input data.

In case of shoot-dry-weight data, please use the $MOSAIC_{growth}$ application https://mosaic.univ-lyon1.fr/growth. The $MOSAIC_{growth}$ application makes possible to analyse multiple data sets simultaneously and to get censored $ER_x$ values according to C1T0.5. These censored $ER_x$ values can be downloaded as a txt file and used for further SSD analyses.

#### Second step: SSD analyses

Use "SSD" module of MOSAIC to run SSD analyses: https://mosaic.univ-lyon1.fr/ssd
