## Supplementary material for "How to account for the uncertainty from standard toxicity tests in species sensitivity distributions: an example in non-target plants": Archive with reports on studies: ER50_censoring.pdf

### $ER_{50}$ censoring after dose-response analyses

Study 1

25 June 2020

Contact:

---

This document provides the $ER_{50}$ censoring after dose-response analyses for study 1, including four parts:

- Raw data description (test item rates);
- Summary on $ER_{50}$ from a Bayesian dose-response fit;
- Censoring on $ER_{50}$ according to two criteria;
  - Criterion 1 ($C_1$) based on the ratio of the probablities, that is the ratio of the probability that the $ER_{50}$ lies within the range of the LCI95 and the maximum tested rate over the probability that the $ER_{50}$ lies in the CI95;
  - Criterion 2 ($C_2$) based on the ratio of distances, that is the ratio of the distance between the LCI95 of $ER_{50}$ and the maximum tested rate over the extend of the CI95 of $ER_{50}$;
- Graphical presentation of censored $ER_{50}$ per species per endpoint.

##### Meaning of symbols used in the document

- min_rate: minimum tested rate;
- max_rate: maximum tested rate;
- $ER_{50}$: 50% effective rate;
- CI95: 95% credible interval;
- $ER_{50}$_median: median of $ER_{50}$;
- $ER_{50}$_LCI95: lower bound of the CI95 of $ER_{50}$;
- $ER_{50}$_UCI95: upper bound of the CI95 of $ER_{50}$;
- $C_1$: criterion based on the ratio of probabilities;
- $C_2$: criterion based on the ratio of distances;
- T: decision threshold;
- Censored $ER_{50}$ [lower bound; upper bound]:
  - C1T0.0: censored $ER_{50}$ according to $C_1$ with T = 0.0;
  - C1T0.2: censored $ER_{50}$ according to $C_1$ with T = 0.2;

- C1T0.4: censored $ER_{50}$ according to $C_1$ with T = 0.4;
- C1T0.5: censored $ER_{50}$ according to $C_1$ with T = 0.5;
- C1T0.6: censored $ER_{50}$ according to $C_1$ with T = 0.6;
- C1T0.8: censored $ER_{50}$ according to $C_1$ with T = 0.8;
- C1T1.0: censored $ER_{50}$ according to $C_1$ with T = 1.0;
- C2T0.0: censored $ER_{50}$ according to $C_2$ with T = 0.0;
- C2T0.2: censored $ER_{50}$ according to $C_2$ with T = 0.2;
- C2T0.4: censored $ER_{50}$ according to $C_2$ with T = 0.4;
- C2T0.5: censored $ER_{50}$ according to $C_2$ with T = 0.5;
- C2T0.6: censored $ER_{50}$ according to $C_2$ with T = 0.6;
- C2T0.8: censored $ER_{50}$ according to $C_2$ with T = 0.8;
- C2T1.0: censored $ER_{50}$ according to $C_2$ with T = 1.0.

### Seeding emergence (SE) test of study 1

##### Raw data description (test item rates) for the SE test

```
Number of species:  10
EPPO code:  ALLCE AVESA BEAVA BRSNW CUMSA GLXMA HELAN LYPES TRZAW ZEAMA
```

Table 1: Correspondance between species and common names according to the EPPO code

| EPPO code | Species name | Common name |
|---|---|---|
| ALLCE | *Allium cepa* | Onion |
| AVESA | *Avena sativa* | Oat |
| BEAVA | *Beta vulgaris* | Sugar beet |
| BRSNW | *Brassica napus winter* | Oilseed rape |
| CUMSA | *Cucumis sativus* | Cucumber |
| GLXMA | *Glycine max* | Soybean |
| HELAN | *Helianthus annuus* | Sunflower |
| LYPES | *Solanum lycopersicum* | Tomato |
| TRZAW | *Triticum aestivum winter* | Winter wheat |
| ZEAMA | *Zea mays subs. Amylacea* | Corn |

Table 2: Tested rates per species (ml product/ha) for the SE test of study 1

| ALLCE | AVESA | BEAVA | BRSNW | CUMSA | GLXMA | HELAN | LYPES | TRZAW | ZEAMA |
|---|---|---|---|---|---|---|---|---|---|
| 0.00 | 0.00 | 0.00 | 0.00 | 0.00 | 0.00 | 0.00 | 0.00 | 0.00 | 0.00 |
| 31.07 | 71.47 | 31.07 | 31.07 | 31.07 | 31.07 | 31.07 | 31.07 | 71.47 | 71.47 |
| 71.47 | 164.38 | 71.47 | 71.47 | 71.47 | 71.47 | 71.47 | 71.47 | 164.38 | 164.38 |
| 164.38 | 378.07 | 164.38 | 164.38 | 164.38 | 164.38 | 164.38 | 164.38 | 378.07 | 378.07 |
| 378.07 | 869.57 | 378.07 | 378.07 | 378.07 | 378.07 | 378.07 | 378.07 | 869.57 | 869.57 |
| 869.57 | 2000.00 | 869.57 | 869.57 | 869.57 | 869.57 | 869.57 | 869.57 | 2000.00 | 2000.00 |

#### SE test - emergence endpoint

Table 3: Minimum and maximum tested rates, median, quantile 2.5% and quantile 97.5% of the $ER_{50}$ posterior distribution - emergence endpoint of the SE test of study 1

| species | min_rate | max_rate | $ER_{50}$_median | $ER_{50}$_LCI95 | $ER_{50}$_UCI95 |
|---|---|---|---|---|---|
| ALLCE | 31.07 | 869.57 | 1420.196 | 692.4289 | 3756.931 |
| AVESA | 71.47 | 2000.00 | 1506.596 | 1053.5823 | 2425.402 |
| BEAVA | 31.07 | 869.57 | 1203.154 | 908.4504 | 2777.003 |
| BRSNW | 31.07 | 869.57 | 1262.485 | 932.6021 | 2937.444 |
| CUMSA | 31.07 | 869.57 | 2674.779 | 1416.8116 | 6583.219 |
| GLXMA | 31.07 | 869.57 | 1270.655 | 933.6909 | 3001.201 |
| HELAN | 31.07 | 869.57 | 2543.192 | 1395.4354 | 6113.227 |
| LYPES | 31.07 | 869.57 | 1246.794 | 924.3375 | 2886.373 |
| TRZAW | 71.47 | 2000.00 | 2093.930 | 1879.2971 | 2909.464 |
| ZEAMA | 71.47 | 2000.00 | 5813.628 | 3111.8055 | 14090.879 |

Table 4: Censored $ER_{50}$ according to criterion 1 and the seven decision thresholds - emergence endpoint of the SE test of study 1

| species | C1T0.0 | C1T0.2 | C1T0.4 | C1T0.5 | C1T0.6 | C1T0.8 | C1T1.0 |
|---|---|---|---|---|---|---|---|
| ALLCE | [692.43; 3756.93] | [692.43; Inf) | [692.43; Inf) | [692.43; Inf) | [692.43; Inf) | [692.43; Inf) | [692.43; Inf) |
| AVESA | [1053.58; 2425.4] | [1053.58; 2425.4] | [1053.58; 2425.4] | [1053.58; 2425.4] | [1053.58; 2425.4] | [1053.58; 2425.4] | [1053.58; Inf) |
| BEAVA | [869.57; Inf) | [869.57; Inf) | [869.57; Inf) | [869.57; Inf) | [869.57; Inf) | [869.57; Inf) | [869.57; Inf) |
| BRSNW | [869.57; Inf) | [869.57; Inf) | [869.57; Inf) | [869.57; Inf) | [869.57; Inf) | [869.57; Inf) | [869.57; Inf) |
| CUMSA | [869.57; Inf) | [869.57; Inf) | [869.57; Inf) | [869.57; Inf) | [869.57; Inf) | [869.57; Inf) | [869.57; Inf) |
| GLXMA | [869.57; Inf) | [869.57; Inf) | [869.57; Inf) | [869.57; Inf) | [869.57; Inf) | [869.57; Inf) | [869.57; Inf) |
| HELAN | [869.57; Inf) | [869.57; Inf) | [869.57; Inf) | [869.57; Inf) | [869.57; Inf) | [869.57; Inf) | [869.57; Inf) |
| LYPES | [869.57; Inf) | [869.57; Inf) | [869.57; Inf) | [869.57; Inf) | [869.57; Inf) | [869.57; Inf) | [869.57; Inf) |
| TRZAW | [1879.3; 2909.46] | [1879.3; Inf) | [1879.3; Inf) | [1879.3; Inf) | [1879.3; Inf) | [1879.3; Inf) | [1879.3; Inf) |
| ZEAMA | [2000; Inf) | [2000; Inf) | [2000; Inf) | [2000; Inf) | [2000; Inf) | [2000; Inf) | [2000; Inf) |

Table 5: Censored $ER_{50}$ according to criterion 2 and the seven decision thresholds - emergence endpoint of the SE test of study 1

| species | C2T0.0 | C2T0.2 | C2T0.4 | C2T0.5 | C2T0.6 | C2T0.8 | C2T1.0 |
|---|---|---|---|---|---|---|---|
| ALLCE | [692.43; 3756.93] | [692.43; Inf) | [692.43; Inf) | [692.43; Inf) | [692.43; Inf) | [692.43; Inf) | [692.43; Inf) |
| AVESA | [1053.58; 2425.4] | [1053.58; 2425.4] | [1053.58; 2425.4] | [1053.58; 2425.4] | [1053.58; 2425.4] | [1053.58; Inf) | [1053.58; Inf) |
| BEAVA | [869.57; Inf) | [869.57; Inf) | [869.57; Inf) | [869.57; Inf) | [869.57; Inf) | [869.57; Inf) | [869.57; Inf) |
| BRSNW | [869.57; Inf) | [869.57; Inf) | [869.57; Inf) | [869.57; Inf) | [869.57; Inf) | [869.57; Inf) | [869.57; Inf) |
| CUMSA | [869.57; Inf) | [869.57; Inf) | [869.57; Inf) | [869.57; Inf) | [869.57; Inf) | [869.57; Inf) | [869.57; Inf) |
| GLXMA | [869.57; Inf) | [869.57; Inf) | [869.57; Inf) | [869.57; Inf) | [869.57; Inf) | [869.57; Inf) | [869.57; Inf) |
| HELAN | [869.57; Inf) | [869.57; Inf) | [869.57; Inf) | [869.57; Inf) | [869.57; Inf) | [869.57; Inf) | [869.57; Inf) |
| LYPES | [869.57; Inf) | [869.57; Inf) | [869.57; Inf) | [869.57; Inf) | [869.57; Inf) | [869.57; Inf) | [869.57; Inf) |
| TRZAW | [1879.3; 2909.46] | [1879.3; Inf) | [1879.3; Inf) | [1879.3; Inf) | [1879.3; Inf) | [1879.3; Inf) | [1879.3; Inf) |
| ZEAMA | [2000; Inf) | [2000; Inf) | [2000; Inf) | [2000; Inf) | [2000; Inf) | [2000; Inf) | [2000; Inf) |

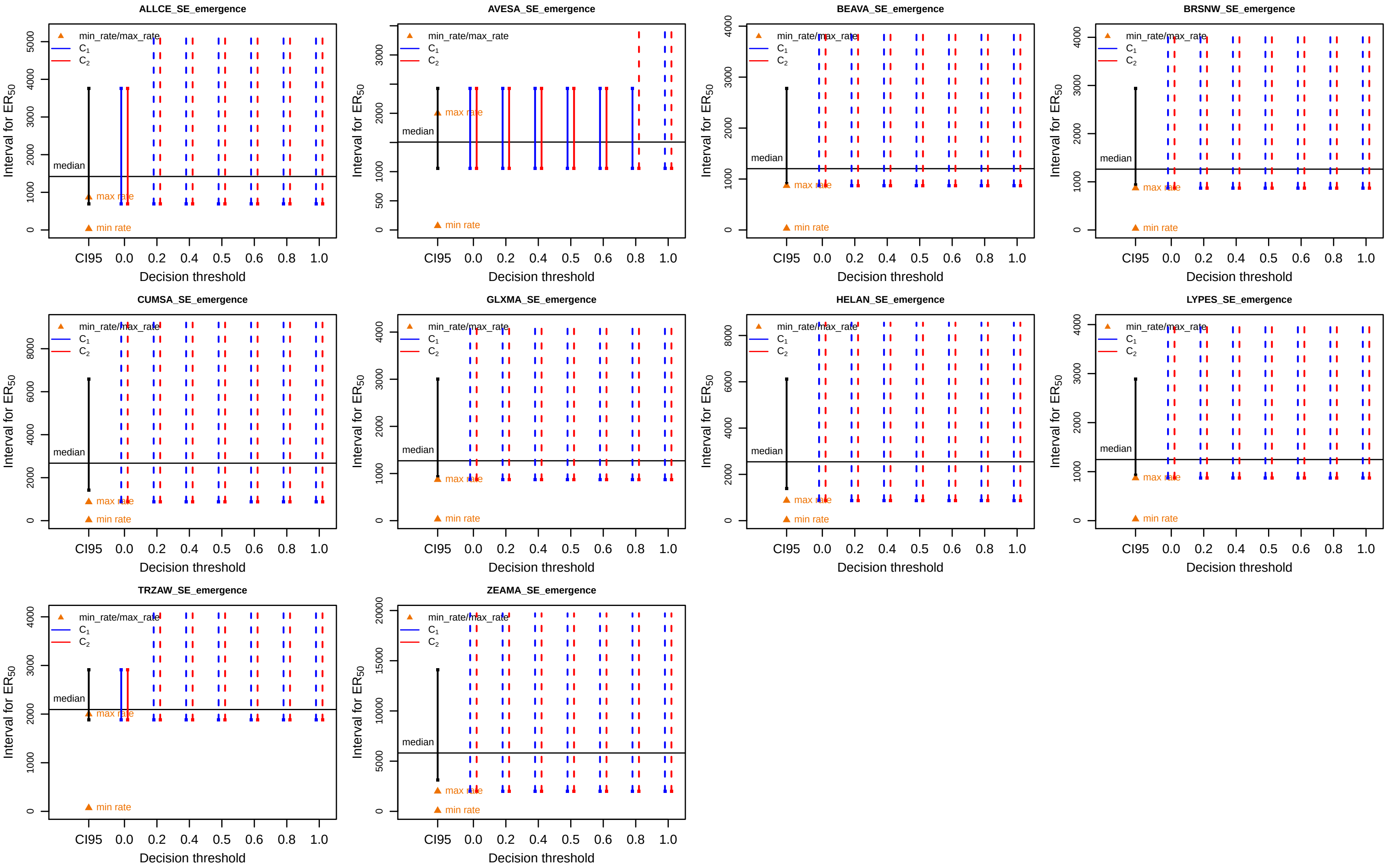


Figure 1: CI95 of $ER_{50}$ and censored $ER_{50}$ according to both criteria and the seven decision thresholds for all species - emergence endpoint of the SE test of study 1. The two orange triangles stand for the lowest and the highest tested rates. Vertical segments of three different colors (black, blue, red) represent the CI95 of $ER_{50}$, the censored $ER_{50}$ according to $C_1$ and the censored $ER_{50}$ according to $C_2$, respectively; solid vertical segments are for bounded intervals while dotted vertical segments stand for right-unbounded intervals; the black horizontal line represents the median of the $ER_{50}$.

#### SE test - survival endpoint

Table 6: Minimum and maximum tested rates, median, quantile 2.5% and quantile 97.5% of the $ER_{50}$ posterior distribution - survival endpoint of the SE test of study 1

| species | min_rate | max_rate | $ER_{50}$_median | $ER_{50}$_LCI95 | $ER_{50}$_UCI95 |
|---|---|---|---|---|---|
| ALLCE | 31.07 | 869.57 | 163.0892 | 128.5010 | 206.3729 |
| AVESA | 71.47 | 2000.00 | 264.4067 | 201.1559 | 352.9132 |
| BEAVA | 31.07 | 869.57 | 368.2296 | 300.3761 | 448.5027 |
| BRSNW | 31.07 | 869.57 | 539.8981 | 444.0571 | 661.1070 |
| CUMSA | 31.07 | 869.57 | 555.7471 | 430.3477 | 754.0101 |
| GLXMA | 31.07 | 869.57 | 1270.8044 | 931.2418 | 2845.2630 |
| HELAN | 31.07 | 869.57 | 1269.0585 | 930.7878 | 2936.6626 |
| LYPES | 31.07 | 869.57 | 1108.0447 | 901.0288 | 2353.7261 |
| TRZAW | 71.47 | 2000.00 | 1938.4264 | 1369.9977 | 3381.9892 |
| ZEAMA | 71.47 | 2000.00 | 2794.5687 | 2085.5955 | 6353.9845 |

Table 7: Censored $ER_{50}$ according to criterion 1 and the seven decision thresholds - survival endpoint of the SE test of study 1

| species | C1T0.0 | C1T0.2 | C1T0.4 | C1T0.5 | C1T0.6 | C1T0.8 | C1T1.0 |
|---|---|---|---|---|---|---|---|
| ALLCE | [128.5; 206.37] | [128.5; 206.37] | [128.5; 206.37] | [128.5; 206.37] | [128.5; 206.37] | [128.5; 206.37] | [128.5; 206.37] |
| AVESA | [201.16; 352.91] | [201.16; 352.91] | [201.16; 352.91] | [201.16; 352.91] | [201.16; 352.91] | [201.16; 352.91] | [201.16; 352.91] |
| BEAVA | [300.38; 448.5] | [300.38; 448.5] | [300.38; 448.5] | [300.38; 448.5] | [300.38; 448.5] | [300.38; 448.5] | [300.38; 448.5] |
| BRSNW | [444.06; 661.11] | [444.06; 661.11] | [444.06; 661.11] | [444.06; 661.11] | [444.06; 661.11] | [444.06; 661.11] | [444.06; 661.11] |
| CUMSA | [430.35; 754.01] | [430.35; 754.01] | [430.35; 754.01] | [430.35; 754.01] | [430.35; 754.01] | [430.35; 754.01] | [430.35; 754.01] |
| GLXMA | [869.57; Inf) | [869.57; Inf) | [869.57; Inf) | [869.57; Inf) | [869.57; Inf) | [869.57; Inf) | [869.57; Inf) |
| HELAN | [869.57; Inf) | [869.57; Inf) | [869.57; Inf) | [869.57; Inf) | [869.57; Inf) | [869.57; Inf) | [869.57; Inf) |
| LYPES | [869.57; Inf) | [869.57; Inf) | [869.57; Inf) | [869.57; Inf) | [869.57; Inf) | [869.57; Inf) | [869.57; Inf) |
| TRZAW | [1370; 3381.99] | [1370; 3381.99] | [1370; 3381.99] | [1370; 3381.99] | [1370; Inf) | [1370; Inf) | [1370; Inf) |
| ZEAMA | [2000; Inf) | [2000; Inf) | [2000; Inf) | [2000; Inf) | [2000; Inf) | [2000; Inf) | [2000; Inf) |

Table 8: Censored $ER_{50}$ according to criterion 2 and the seven decision thresholds - survival endpoint of the SE test of study 1

| species | C2T0.0 | C2T0.2 | C2T0.4 | C2T0.5 | C2T0.6 | C2T0.8 | C2T1.0 |
|---|---|---|---|---|---|---|---|
| ALLCE | [128.5; 206.37] | [128.5; 206.37] | [128.5; 206.37] | [128.5; 206.37] | [128.5; 206.37] | [128.5; 206.37] | [128.5; 206.37] |
| AVESA | [201.16; 352.91] | [201.16; 352.91] | [201.16; 352.91] | [201.16; 352.91] | [201.16; 352.91] | [201.16; 352.91] | [201.16; 352.91] |
| BEAVA | [300.38; 448.5] | [300.38; 448.5] | [300.38; 448.5] | [300.38; 448.5] | [300.38; 448.5] | [300.38; 448.5] | [300.38; 448.5] |
| BRSNW | [444.06; 661.11] | [444.06; 661.11] | [444.06; 661.11] | [444.06; 661.11] | [444.06; 661.11] | [444.06; 661.11] | [444.06; 661.11] |
| CUMSA | [430.35; 754.01] | [430.35; 754.01] | [430.35; 754.01] | [430.35; 754.01] | [430.35; 754.01] | [430.35; 754.01] | [430.35; 754.01] |
| GLXMA | [869.57; Inf) | [869.57; Inf) | [869.57; Inf) | [869.57; Inf) | [869.57; Inf) | [869.57; Inf) | [869.57; Inf) |
| HELAN | [869.57; Inf) | [869.57; Inf) | [869.57; Inf) | [869.57; Inf) | [869.57; Inf) | [869.57; Inf) | [869.57; Inf) |
| LYPES | [869.57; Inf) | [869.57; Inf) | [869.57; Inf) | [869.57; Inf) | [869.57; Inf) | [869.57; Inf) | [869.57; Inf) |
| TRZAW | [1370; 3381.99] | [1370; 3381.99] | [1370; Inf) | [1370; Inf) | [1370; Inf) | [1370; Inf) | [1370; Inf) |
| ZEAMA | [2000; Inf) | [2000; Inf) | [2000; Inf) | [2000; Inf) | [2000; Inf) | [2000; Inf) | [2000; Inf) |

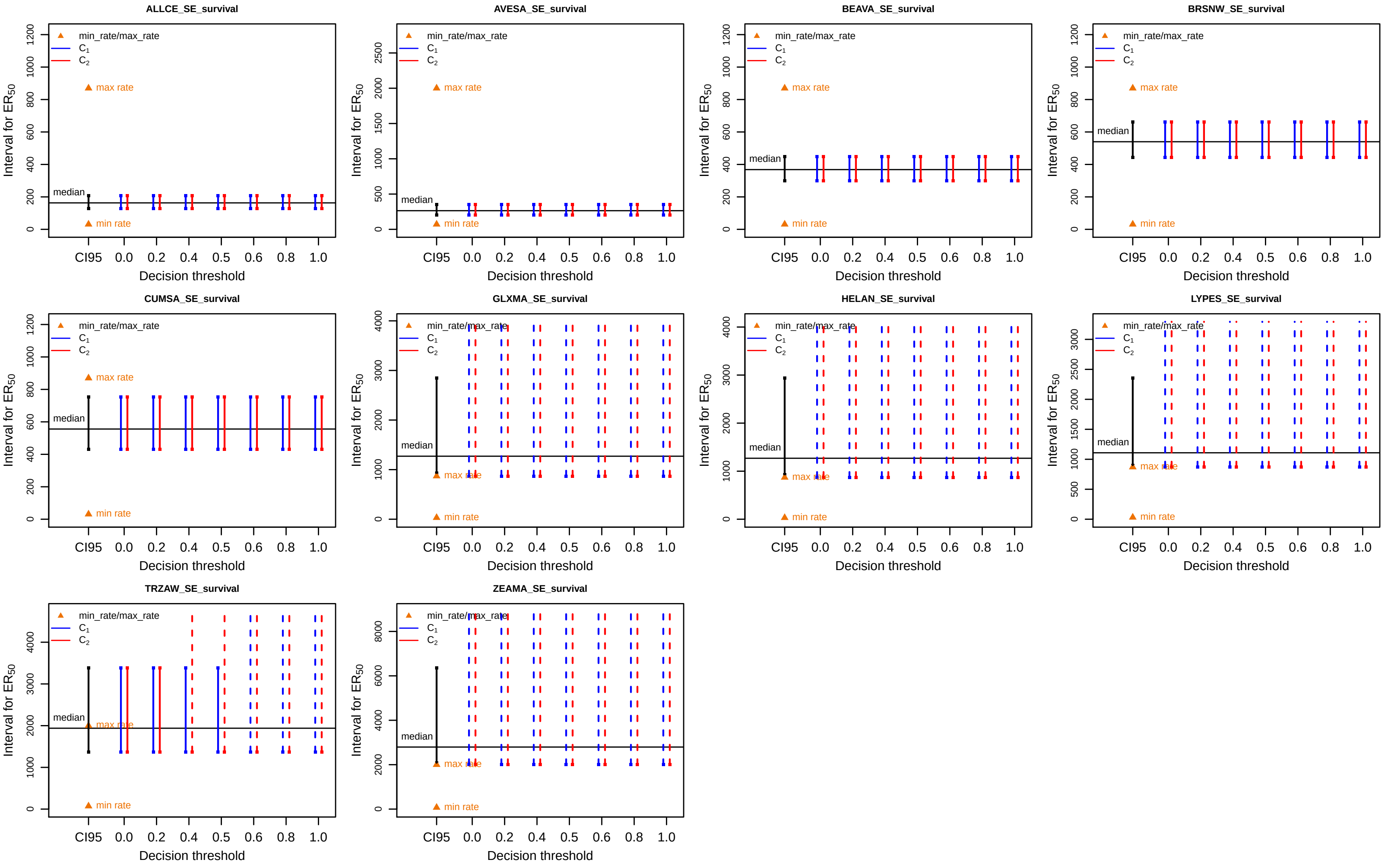


Figure 2: CI95 of $ER_{50}$ and censored $ER_{50}$ according to both criteria and the seven decision thresholds for all species - survival endpoint of the SE test of study 1. The two orange triangles stand for the lowest and the highest tested rates. Vertical segments of three different colors (black, blue, red) represent the CI95 of $ER_{50}$, the censored $ER_{50}$ according to $C_1$ and the censored $ER_{50}$ according to $C_2$, respectively; solid vertical segments are for bounded intervals while dotted vertical segments stand for right-unbounded intervals; the black horizontal line represents the median of the $ER_{50}$.

#### SE test - shoot dry weight endpoint

Table 9: Minimum and maximum tested rates, median, quantile 2.5% and quantile 97.5% of the $ER_{50}$ posterior distribution - shoot dry weight endpoint of the SE test of study 1

| species | min_rate | max_rate | $ER_{50}$_median | $ER_{50}$_LCI95 | $ER_{50}$_UCI95 |
|---|---|---|---|---|---|
| ALLCE | 31.07 | 378.07 | 155.8689 | 106.6271 | 222.6088 |
| AVESA | 71.47 | 869.57 | 183.3126 | 142.1188 | 244.2603 |
| BEAVA | 31.07 | 378.07 | 283.2377 | 230.0457 | 351.8027 |
| BRSNW | 31.07 | 869.57 | 321.0205 | 272.8069 | 372.1631 |
| CUMSA | 31.07 | 869.57 | 275.3910 | 236.3495 | 321.3627 |
| GLXMA | 31.07 | 869.57 | 1215.5481 | 913.7874 | 2761.6941 |
| HELAN | 31.07 | 869.57 | 1209.7758 | 911.7498 | 2903.6297 |
| LYPES | 31.07 | 869.57 | 597.0532 | 507.2770 | 701.1788 |
| TRZAW | 71.47 | 2000.00 | 670.0644 | 532.6734 | 841.1321 |
| ZEAMA | 71.47 | 2000.00 | 1992.5124 | 1312.3398 | 3350.1970 |

Table 10: Censored $ER_{50}$ according to criterion 1 and the seven decision thresholds - shoot dry weight endpoint of the SE test of study 1

| species | C1T0.0 | C1T0.2 | C1T0.4 | C1T0.5 | C1T0.6 | C1T0.8 | C1T1.0 |
|---|---|---|---|---|---|---|---|
| ALLCE | [106.63; 222.61] | [106.63; 222.61] | [106.63; 222.61] | [106.63; 222.61] | [106.63; 222.61] | [106.63; 222.61] | [106.63; 222.61] |
| AVESA | [142.12; 244.26] | [142.12; 244.26] | [142.12; 244.26] | [142.12; 244.26] | [142.12; 244.26] | [142.12; 244.26] | [142.12; 244.26] |
| BEAVA | [230.05; 351.8] | [230.05; 351.8] | [230.05; 351.8] | [230.05; 351.8] | [230.05; 351.8] | [230.05; 351.8] | [230.05; 351.8] |
| BRSNW | [272.81; 372.16] | [272.81; 372.16] | [272.81; 372.16] | [272.81; 372.16] | [272.81; 372.16] | [272.81; 372.16] | [272.81; 372.16] |
| CUMSA | [236.35; 321.36] | [236.35; 321.36] | [236.35; 321.36] | [236.35; 321.36] | [236.35; 321.36] | [236.35; 321.36] | [236.35; 321.36] |
| GLXMA | [869.57; Inf) | [869.57; Inf) | [869.57; Inf) | [869.57; Inf) | [869.57; Inf) | [869.57; Inf) | [869.57; Inf) |
| HELAN | [869.57; Inf) | [869.57; Inf) | [869.57; Inf) | [869.57; Inf) | [869.57; Inf) | [869.57; Inf) | [869.57; Inf) |
| LYPES | [507.28; 701.18] | [507.28; 701.18] | [507.28; 701.18] | [507.28; 701.18] | [507.28; 701.18] | [507.28; 701.18] | [507.28; 701.18] |
| TRZAW | [532.67; 841.13] | [532.67; 841.13] | [532.67; 841.13] | [532.67; 841.13] | [532.67; 841.13] | [532.67; 841.13] | [532.67; 841.13] |
| ZEAMA | [1312.34; 3350.2] | [1312.34; 3350.2] | [1312.34; 3350.2] | [1312.34; 3350.2] | [1312.34; Inf) | [1312.34; Inf) | [1312.34; Inf) |

Table 11: Censored $ER_{50}$ according to criterion 2 and the seven decision thresholds - shoot dry weight endpoint of the SE test of study 1

| species | C2T0.0 | C2T0.2 | C2T0.4 | C2T0.5 | C2T0.6 | C2T0.8 | C2T1.0 |
|---|---|---|---|---|---|---|---|
| ALLCE | [106.63; 222.61] | [106.63; 222.61] | [106.63; 222.61] | [106.63; 222.61] | [106.63; 222.61] | [106.63; 222.61] | [106.63; 222.61] |
| AVESA | [142.12; 244.26] | [142.12; 244.26] | [142.12; 244.26] | [142.12; 244.26] | [142.12; 244.26] | [142.12; 244.26] | [142.12; 244.26] |
| BEAVA | [230.05; 351.8] | [230.05; 351.8] | [230.05; 351.8] | [230.05; 351.8] | [230.05; 351.8] | [230.05; 351.8] | [230.05; 351.8] |
| BRSNW | [272.81; 372.16] | [272.81; 372.16] | [272.81; 372.16] | [272.81; 372.16] | [272.81; 372.16] | [272.81; 372.16] | [272.81; 372.16] |
| CUMSA | [236.35; 321.36] | [236.35; 321.36] | [236.35; 321.36] | [236.35; 321.36] | [236.35; 321.36] | [236.35; 321.36] | [236.35; 321.36] |
| GLXMA | [869.57; Inf) | [869.57; Inf) | [869.57; Inf) | [869.57; Inf) | [869.57; Inf) | [869.57; Inf) | [869.57; Inf) |
| HELAN | [869.57; Inf) | [869.57; Inf) | [869.57; Inf) | [869.57; Inf) | [869.57; Inf) | [869.57; Inf) | [869.57; Inf) |
| LYPES | [507.28; 701.18] | [507.28; 701.18] | [507.28; 701.18] | [507.28; 701.18] | [507.28; 701.18] | [507.28; 701.18] | [507.28; 701.18] |
| TRZAW | [532.67; 841.13] | [532.67; 841.13] | [532.67; 841.13] | [532.67; 841.13] | [532.67; 841.13] | [532.67; 841.13] | [532.67; 841.13] |
| ZEAMA | [1312.34; 3350.2] | [1312.34; 3350.2] | [1312.34; Inf) | [1312.34; Inf) | [1312.34; Inf) | [1312.34; Inf) | [1312.34; Inf) |

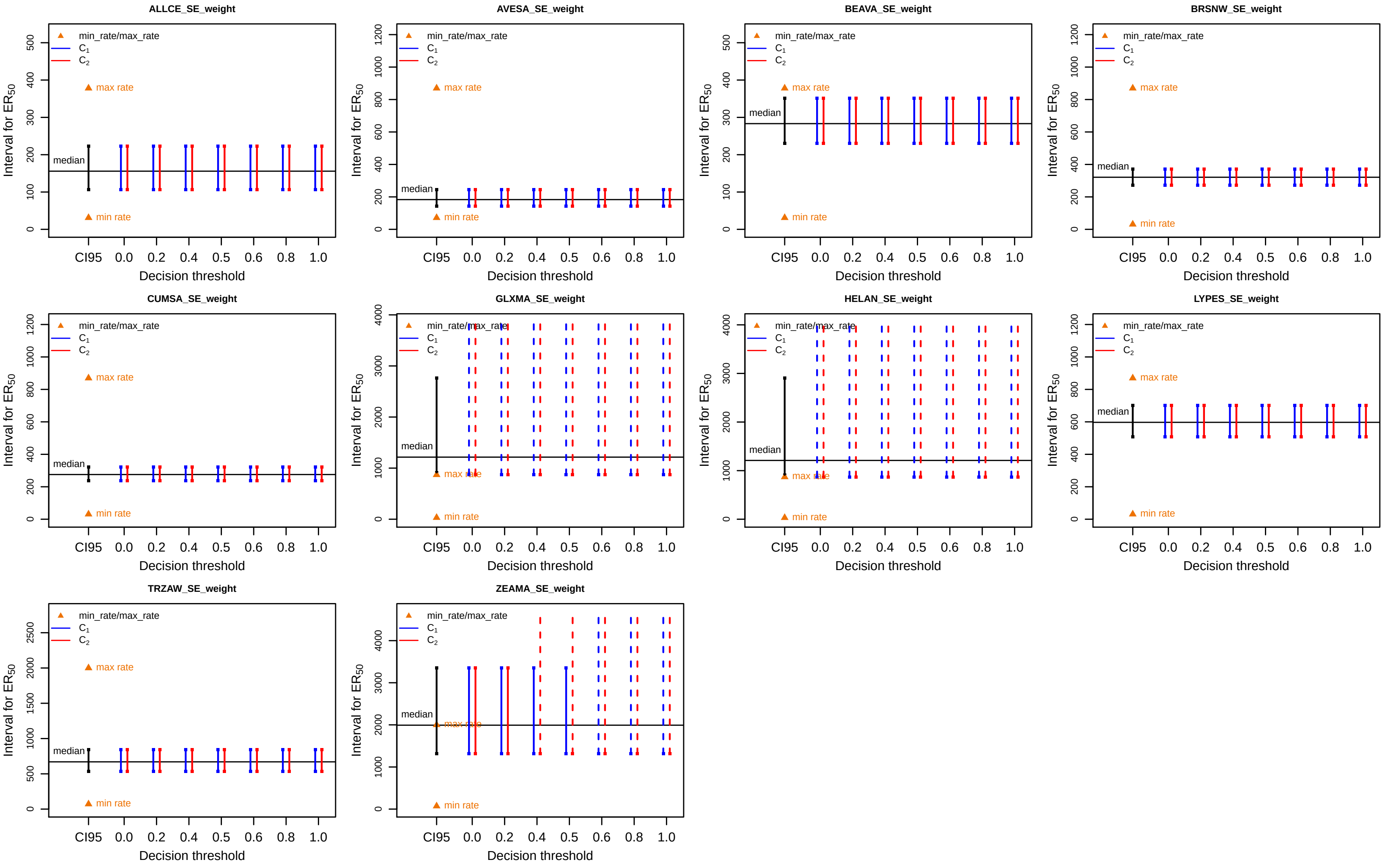


Figure 3: CI95 of $ER_{50}$ and censored $ER_{50}$ according to both criteria and the seven decision thresholds for all species - shoot dry weight endpoint of the SE test of study 1. The two orange triangles stand for the lowest and the highest tested rates. Vertical segments of three different colors (black, blue, red) represent the CI95 of $ER_{50}$, the censored $ER_{50}$ according to $C_1$ and the censored $ER_{50}$ according to $C_2$, respectively; solid vertical segments are for bounded intervals while dotted vertical segments stand for right-unbounded intervals; the black horizontal line represents the median of the $ER_{50}$.

### Vegatative vigor (VV) test of study 1

##### Raw data description (test item rates) for VV test

```
Number of species:  10
EPPO code:  ALLCE AVESA BEAVA BRSNW CUMSA GLXMA HELAN LYPES TRZAW ZEAMA
```

Table 12: Corresponding species and common names to the EPPO code

| EPPO code | Species name | Common name |
|---|---|---|
| ALLCE | *Allium cepa* | Onion |
| AVESA | *Avena sativa* | Oat |
| BEAVA | *Beta vulgaris* | Sugar beet |
| BRSNW | *Brassica napus winter* | Oilseed rape |
| CUMSA | *Cucumis sativus* | Cucumber |
| GLXMA | *Glycine max* | Soybean |
| HELAN | *Helianthus annuus* | Sunflower |
| LYPES | *Solanum lycopersicum* | Tomato |
| TRZAW | *Triticum aestivum winter* | Winter wheat |
| ZEAMA | *Zea mays subs. Amylacea* | Corn |

Table 13: Tested rates per species (ml product/ha) for the VV test of study 1

| ALLCE | AVESA | BEAVA | BRSNW | CUMSA | GLXMA | HELAN | LYPES | TRZAW | ZEAMA |
|---|---|---|---|---|---|---|---|---|---|
| 0.00 | 0.00 | 0.00 | 0.00 | 0.00 | 0.00 | 0.00 | 0.00 | 0.00 | 0.00 |
| 31.07 | 71.47 | 31.07 | 31.07 | 31.07 | 31.07 | 31.07 | 31.07 | 71.47 | 71.47 |
| 71.47 | 164.38 | 71.47 | 71.47 | 71.47 | 71.47 | 71.47 | 71.47 | 164.38 | 164.38 |
| 164.38 | 378.07 | 164.38 | 164.38 | 164.38 | 164.38 | 164.38 | 164.38 | 378.07 | 378.07 |
| 378.07 | 869.57 | 378.07 | 378.07 | 378.07 | 378.07 | 378.07 | 378.07 | 869.57 | 869.57 |
| 869.57 | 2000.00 | 869.57 | 869.57 | 869.57 | 869.57 | 869.57 | 869.57 | 2000.00 | 2000.00 |

#### VV test - survival endpoint

Table 14: Minimum and maximum tested rates, median, quantile 2.5% and quantile 97.5% of the $ER_{50}$ posterior distribution - survival endpoint of the VV test of study 1

| species | min_rate | max_rate | $ER_{50}$_median | $ER_{50}$_LCI95 | $ER_{50}$_UCI95 |
|---|---|---|---|---|---|
| ALLCE | 31.07 | 869.57 | 1268.5992 | 931.0119 | 2901.631 |
| AVESA | 71.47 | 2000.00 | 2920.8134 | 2147.1338 | 6425.718 |
| BEAVA | 31.07 | 869.57 | 1267.0010 | 933.1524 | 2950.169 |
| BRSNW | 31.07 | 869.57 | 956.0488 | 862.9716 | 1475.258 |
| CUMSA | 31.07 | 869.57 | 1113.0925 | 901.9258 | 2351.503 |
| GLXMA | 31.07 | 869.57 | 1272.9332 | 934.5864 | 2917.581 |
| HELAN | 31.07 | 869.57 | 1271.9763 | 934.9826 | 2842.151 |
| LYPES | 31.07 | 869.57 | 1268.7299 | 933.6295 | 2893.262 |
| TRZAW | 71.47 | 2000.00 | 2930.0900 | 2147.2277 | 6668.731 |
| ZEAMA | 71.47 | 2000.00 | 2901.5010 | 2144.8387 | 6692.480 |

Table 15: Censored $ER_{50}$ according to criterion 1 and the seven decision thresholds - survival endpoint of the VV test of study 1

| species | C1T0.0 | C1T0.2 | C1T0.4 | C1T0.5 | C1T0.6 | C1T0.8 | C1T1.0 |
|---|---|---|---|---|---|---|---|
| ALLCE | [869.57; Inf) | [869.57; Inf) | [869.57; Inf) | [869.57; Inf) | [869.57; Inf) | [869.57; Inf) | [869.57; Inf) |
| AVESA | [2000; Inf) | [2000; Inf) | [2000; Inf) | [2000; Inf) | [2000; Inf) | [2000; Inf) | [2000; Inf) |
| BEAVA | [869.57; Inf) | [869.57; Inf) | [869.57; Inf) | [869.57; Inf) | [869.57; Inf) | [869.57; Inf) | [869.57; Inf) |
| BRSNW | [862.97; 1475.26] | [862.97; Inf) | [862.97; Inf) | [862.97; Inf) | [862.97; Inf) | [862.97; Inf) | [862.97; Inf) |
| CUMSA | [869.57; Inf) | [869.57; Inf) | [869.57; Inf) | [869.57; Inf) | [869.57; Inf) | [869.57; Inf) | [869.57; Inf) |
| GLXMA | [869.57; Inf) | [869.57; Inf) | [869.57; Inf) | [869.57; Inf) | [869.57; Inf) | [869.57; Inf) | [869.57; Inf) |
| HELAN | [869.57; Inf) | [869.57; Inf) | [869.57; Inf) | [869.57; Inf) | [869.57; Inf) | [869.57; Inf) | [869.57; Inf) |
| LYPES | [869.57; Inf) | [869.57; Inf) | [869.57; Inf) | [869.57; Inf) | [869.57; Inf) | [869.57; Inf) | [869.57; Inf) |
| TRZAW | [2000; Inf) | [2000; Inf) | [2000; Inf) | [2000; Inf) | [2000; Inf) | [2000; Inf) | [2000; Inf) |
| ZEAMA | [2000; Inf) | [2000; Inf) | [2000; Inf) | [2000; Inf) | [2000; Inf) | [2000; Inf) | [2000; Inf) |

Table 16: Censored $ER_{50}$ according to criterion 2 and the seven decision thresholds - survival endpoint of the VV test of study 1

| species | C2T0.0 | C2T0.2 | C2T0.4 | C2T0.5 | C2T0.6 | C2T0.8 | C2T1.0 |
|---|---|---|---|---|---|---|---|
| ALLCE | [869.57; Inf) | [869.57; Inf) | [869.57; Inf) | [869.57; Inf) | [869.57; Inf) | [869.57; Inf) | [869.57; Inf) |
| AVESA | [2000; Inf) | [2000; Inf) | [2000; Inf) | [2000; Inf) | [2000; Inf) | [2000; Inf) | [2000; Inf) |
| BEAVA | [869.57; Inf) | [869.57; Inf) | [869.57; Inf) | [869.57; Inf) | [869.57; Inf) | [869.57; Inf) | [869.57; Inf) |
| BRSNW | [862.97; 1475.26] | [862.97; Inf) | [862.97; Inf) | [862.97; Inf) | [862.97; Inf) | [862.97; Inf) | [862.97; Inf) |
| CUMSA | [869.57; Inf) | [869.57; Inf) | [869.57; Inf) | [869.57; Inf) | [869.57; Inf) | [869.57; Inf) | [869.57; Inf) |
| GLXMA | [869.57; Inf) | [869.57; Inf) | [869.57; Inf) | [869.57; Inf) | [869.57; Inf) | [869.57; Inf) | [869.57; Inf) |
| HELAN | [869.57; Inf) | [869.57; Inf) | [869.57; Inf) | [869.57; Inf) | [869.57; Inf) | [869.57; Inf) | [869.57; Inf) |
| LYPES | [869.57; Inf) | [869.57; Inf) | [869.57; Inf) | [869.57; Inf) | [869.57; Inf) | [869.57; Inf) | [869.57; Inf) |
| TRZAW | [2000; Inf) | [2000; Inf) | [2000; Inf) | [2000; Inf) | [2000; Inf) | [2000; Inf) | [2000; Inf) |
| ZEAMA | [2000; Inf) | [2000; Inf) | [2000; Inf) | [2000; Inf) | [2000; Inf) | [2000; Inf) | [2000; Inf) |

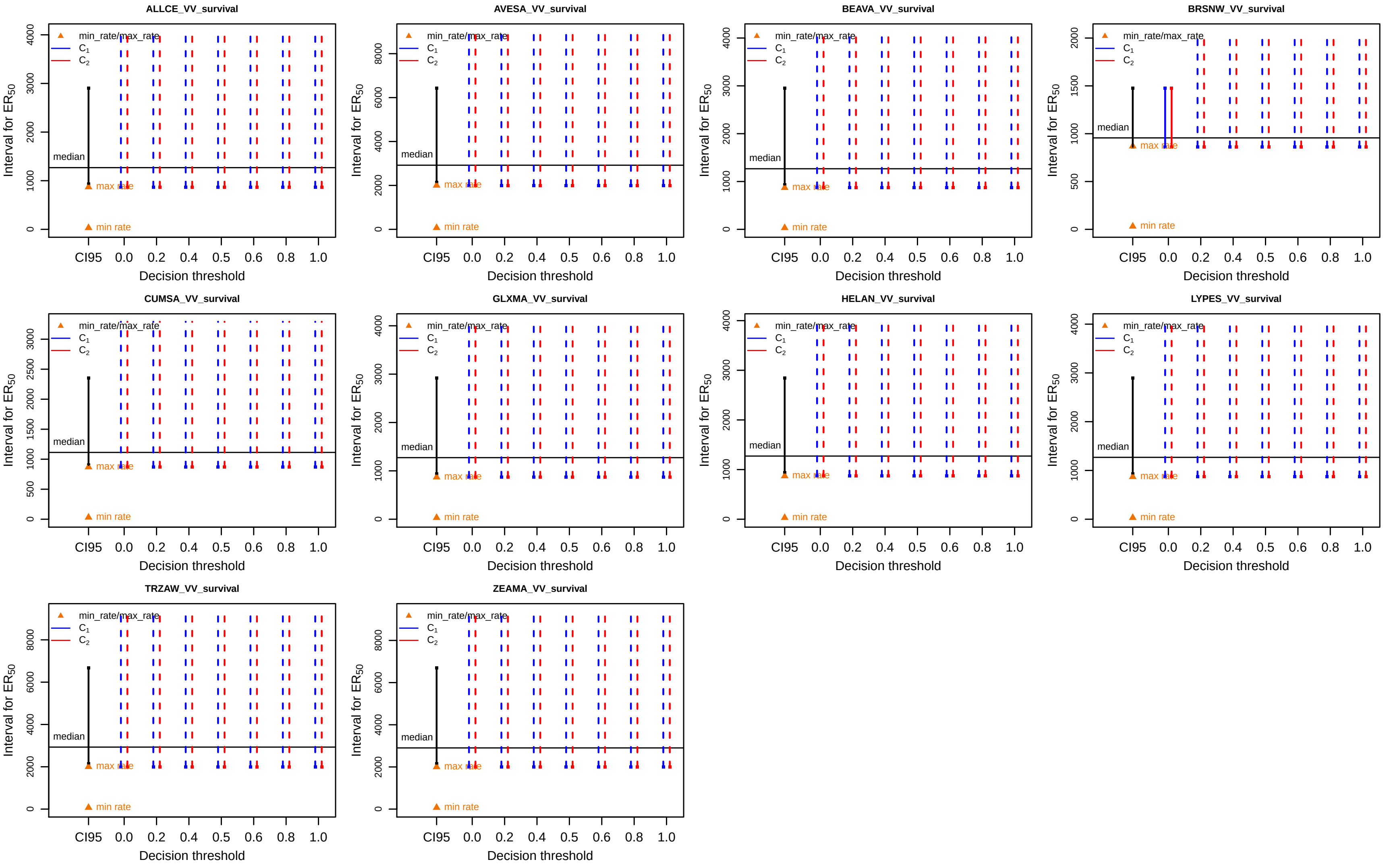


Figure 4: CI95 of $ER_{50}$ and censored $ER_{50}$ according to both criteria and the seven decision thresholds for all species - survival endpoint of the VV test of study 1. The two orange triangles stand for the lowest and the highest tested rates. Vertical segments of three different colors (black, blue, red) represent the CI95 of $ER_{50}$, the censored $ER_{50}$ according to $C_1$ and the censored $ER_{50}$ according to $C_2$, respectively; solid vertical segments are for bounded intervals while dotted vertical segments stand for right-unbounded intervals; the black horizontal line represents the median of the $ER_{50}$.

#### VV test - shoot dry weight endpoint

Table 17: Minimum and maximum tested rates, median, quantile 2.5% and quantile 97.5% of the $ER_{50}$ posterior distribution - shoot dry weight endpoint of the VV test of study 1

| species | min_rate | max_rate | $ER_{50}$_median | $ER_{50}$_LCI95 | $ER_{50}$_UCI95 |
|---|---|---|---|---|---|
| ALLCE | 31.07 | 869.57 | 656.8173 | 76.04318 | 1945.3813 |
| AVESA | 71.47 | 2000.00 | 2235.4933 | 1330.00639 | 3967.5197 |
| BEAVA | 31.07 | 869.57 | 538.5002 | 426.64664 | 681.0293 |
| BRSNW | 31.07 | 869.57 | 166.3894 | 134.76212 | 204.4930 |
| CUMSA | 31.07 | 869.57 | 281.4183 | 177.97705 | 460.0662 |
| GLXMA | 31.07 | 869.57 | 443.8683 | 346.20412 | 577.4326 |
| HELAN | 31.07 | 869.57 | 685.7885 | 510.86219 | 955.9358 |
| LYPES | 31.07 | 869.57 | 1316.7886 | 958.54475 | 1924.9520 |
| TRZAW | 71.47 | 2000.00 | 3874.8076 | 2129.18665 | 7560.3578 |
| ZEAMA | 71.47 | 2000.00 | 6788.7400 | 4142.47676 | 13203.0575 |

Table 18: Censored $ER_{50}$ according to criterion 1 and the seven decision thresholds - shoot dry weight endpoint of the VV test of study 1

| species | C1T0.0 | C1T0.2 | C1T0.4 | C1T0.5 | C1T0.6 | C1T0.8 | C1T1.0 |
|---|---|---|---|---|---|---|---|
| ALLCE | [76.04; 1945.38] | [76.04; 1945.38] | [76.04; 1945.38] | [76.04; 1945.38] | [76.04; 1945.38] | [76.04; Inf) | [76.04; Inf) |
| AVESA | [1330.01; 3967.52] | [1330.01; 3967.52] | [1330.01; Inf) | [1330.01; Inf) | [1330.01; Inf) | [1330.01; Inf) | [1330.01; Inf) |
| BEAVA | [426.65; 681.03] | [426.65; 681.03] | [426.65; 681.03] | [426.65; 681.03] | [426.65; 681.03] | [426.65; 681.03] | [426.65; 681.03] |
| BRSNW | [134.76; 204.49] | [134.76; 204.49] | [134.76; 204.49] | [134.76; 204.49] | [134.76; 204.49] | [134.76; 204.49] | [134.76; 204.49] |
| CUMSA | [177.98; 460.07] | [177.98; 460.07] | [177.98; 460.07] | [177.98; 460.07] | [177.98; 460.07] | [177.98; 460.07] | [177.98; 460.07] |
| GLXMA | [346.2; 577.43] | [346.2; 577.43] | [346.2; 577.43] | [346.2; 577.43] | [346.2; 577.43] | [346.2; 577.43] | [346.2; 577.43] |
| HELAN | [510.86; 955.94] | [510.86; 955.94] | [510.86; 955.94] | [510.86; 955.94] | [510.86; 955.94] | [510.86; 955.94] | [510.86; Inf) |
| LYPES | [869.57; Inf) | [869.57; Inf) | [869.57; Inf) | [869.57; Inf) | [869.57; Inf) | [869.57; Inf) | [869.57; Inf) |
| TRZAW | [2000; Inf) | [2000; Inf) | [2000; Inf) | [2000; Inf) | [2000; Inf) | [2000; Inf) | [2000; Inf) |
| ZEAMA | [2000; Inf) | [2000; Inf) | [2000; Inf) | [2000; Inf) | [2000; Inf) | [2000; Inf) | [2000; Inf) |

Table 19: Censored $ER_{50}$ according to criterion 2 and the seven decision thresholds - shoot dry weight endpoint of the VV test of study 1

| species | C2T0.0 | C2T0.2 | C2T0.4 | C2T0.5 | C2T0.6 | C2T0.8 | C2T1.0 |
|---|---|---|---|---|---|---|---|
| ALLCE | [76.04; 1945.38] | [76.04; 1945.38] | [76.04; 1945.38] | [76.04; Inf) | [76.04; Inf) | [76.04; Inf) | [76.04; Inf) |
| AVESA | [1330.01; 3967.52] | [1330.01; 3967.52] | [1330.01; Inf) | [1330.01; Inf) | [1330.01; Inf) | [1330.01; Inf) | [1330.01; Inf) |
| BEAVA | [426.65; 681.03] | [426.65; 681.03] | [426.65; 681.03] | [426.65; 681.03] | [426.65; 681.03] | [426.65; 681.03] | [426.65; 681.03] |
| BRSNW | [134.76; 204.49] | [134.76; 204.49] | [134.76; 204.49] | [134.76; 204.49] | [134.76; 204.49] | [134.76; 204.49] | [134.76; 204.49] |
| CUMSA | [177.98; 460.07] | [177.98; 460.07] | [177.98; 460.07] | [177.98; 460.07] | [177.98; 460.07] | [177.98; 460.07] | [177.98; 460.07] |
| GLXMA | [346.2; 577.43] | [346.2; 577.43] | [346.2; 577.43] | [346.2; 577.43] | [346.2; 577.43] | [346.2; 577.43] | [346.2; 577.43] |
| HELAN | [510.86; 955.94] | [510.86; 955.94] | [510.86; 955.94] | [510.86; 955.94] | [510.86; 955.94] | [510.86; 955.94] | [510.86; Inf) |
| LYPES | [869.57; Inf) | [869.57; Inf) | [869.57; Inf) | [869.57; Inf) | [869.57; Inf) | [869.57; Inf) | [869.57; Inf) |
| TRZAW | [2000; Inf) | [2000; Inf) | [2000; Inf) | [2000; Inf) | [2000; Inf) | [2000; Inf) | [2000; Inf) |
| ZEAMA | [2000; Inf) | [2000; Inf) | [2000; Inf) | [2000; Inf) | [2000; Inf) | [2000; Inf) | [2000; Inf) |

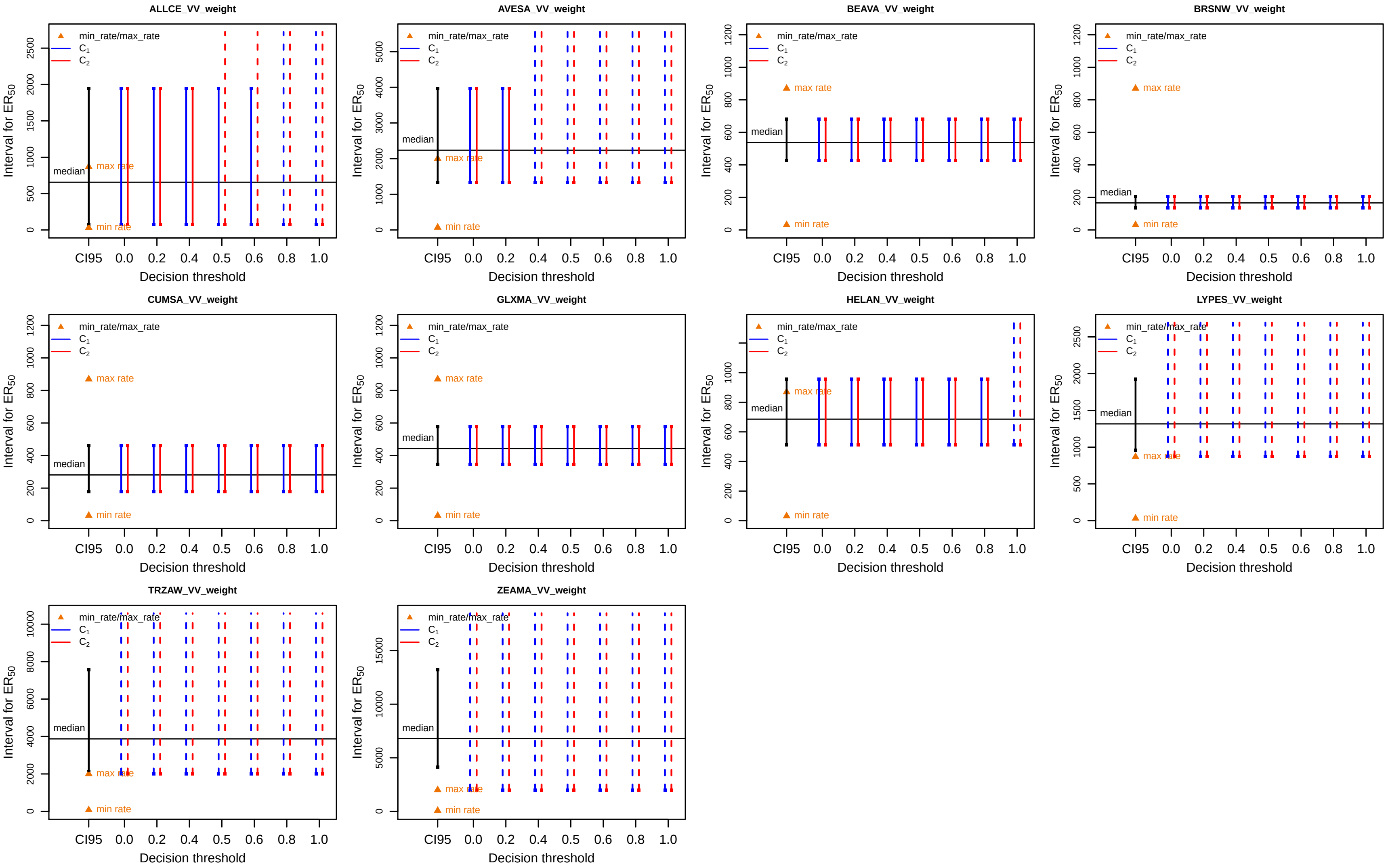


Figure 5: CI95 of $ER_{50}$ and censored $ER_{50}$ according to both criteria and the seven decision thresholds for all species - shoot dry weight endpoint of the VV test of study 1. The two orange triangles stand for the lowest and the highest tested rates. Vertical segments of three different colors (black, blue, red) represent the CI95 of $ER_{50}$, the censored $ER_{50}$ according to $C_1$ and the censored $ER_{50}$ according to $C_2$, respectively; solid vertical segments are for bounded intervals while dotted vertical segments stand for right-unbounded intervals; the black horizontal line represents the median of the $ER_{50}$.
