## Supplementary material for "How to account for the uncertainty from standard toxicity tests in species sensitivity distributions: an example in non-target plants": Archive with reports on studies: report_SE_survival.pdf

### Dose-response analyses

Study 1

Seedling Emergence test - survival endpoint

25 June 2020

Contact:

---

This is a report which provides results on all performed dose-response analyses for the survival endpoint of the Seedling Emergence test for study 1.

---

#### Data set: ALLCE_SE_survival

Table 1: Summary of parameter estimates (parameter d is set to 1) for ALLCE_SE_survival data set

| Parameter | median | Q2.5 | Q97.5 |
|---|---|---|---|
| b | 3.408 | 2.141 | 5.439 |
| e | 163.089 | 128.501 | 206.373 |

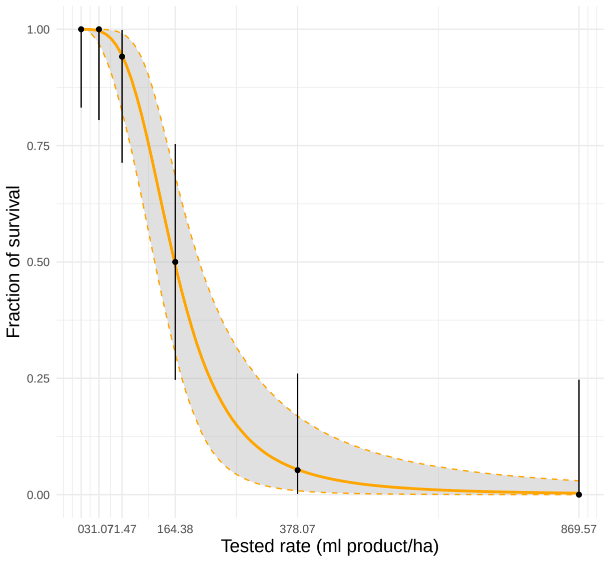

(a) Dose-response curve

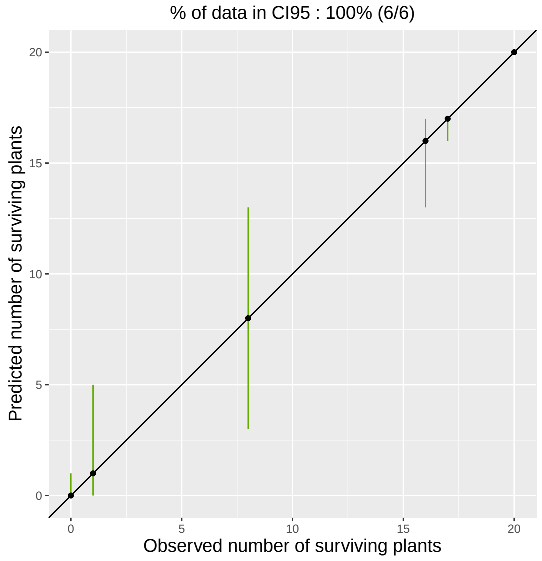

(b) Posterior predictive check (PPC)

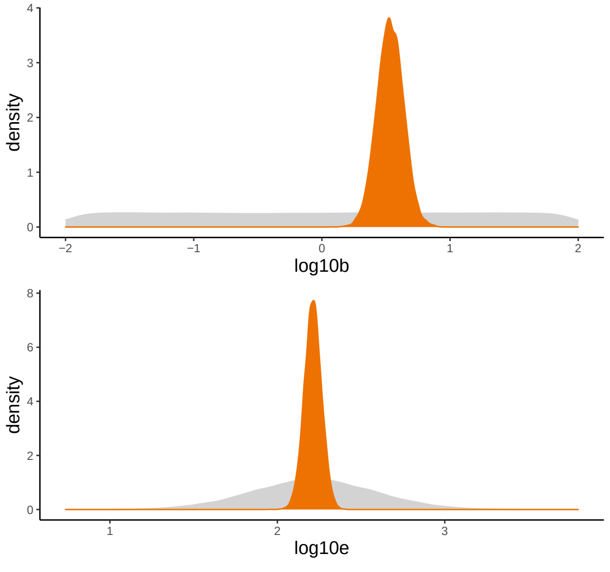

(c) Priors and posteriors

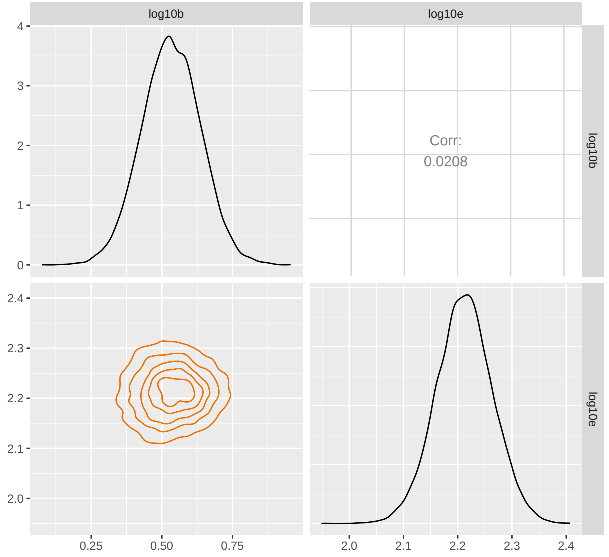

(d) Correlations between parameters

Figure 1: Dose-response curve (a), PPC (b), prior and posterior distributions (c) and correlations between parameters (d).

#### Data set: AVESA_SE_survival

Table 2: Summary of parameter estimates (parameter d is set to 1) for AVESA_SE_survival data set

| Parameter | median | Q2.5 | Q97.5 |
|---|---|---|---|
| b | 2.420 | 1.507 | 3.621 |
| e | 264.407 | 201.156 | 352.913 |

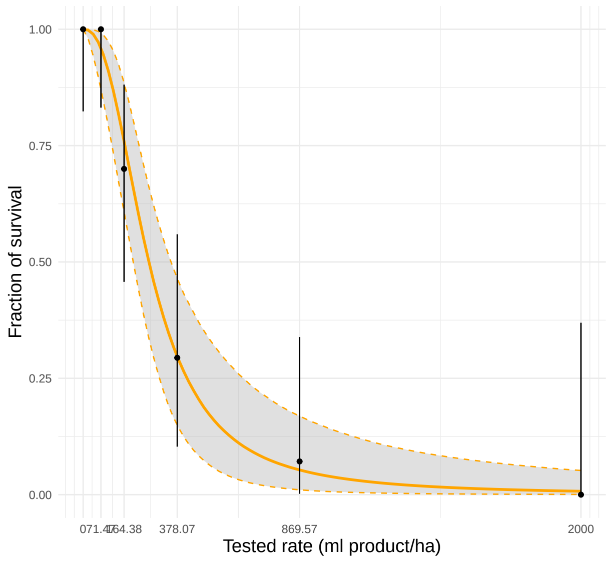

(a) Dose-response curve

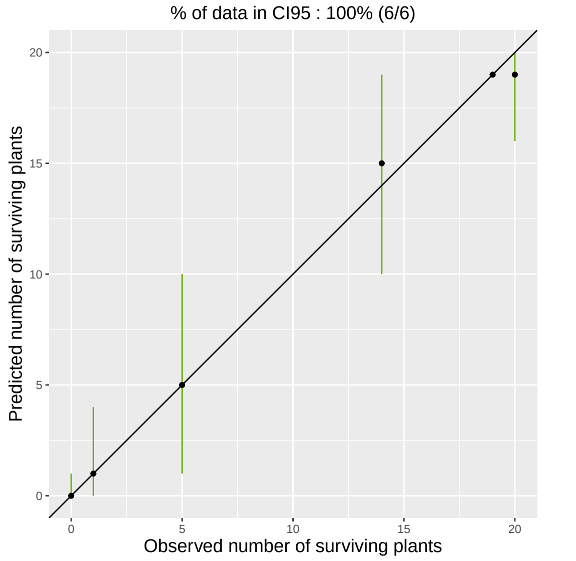

(b) Posterior predictive check (PPC)

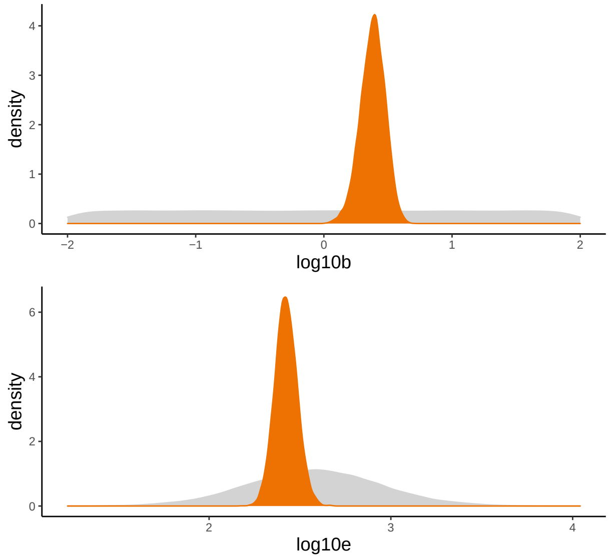

(c) Priors and posteriors

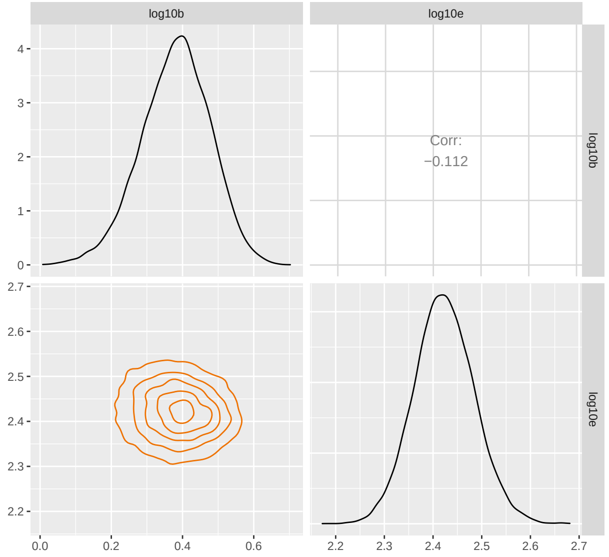

(d) Correlations between parameters

Figure 2: Dose-response curve (a), PPC (b), prior and posterior distributions (c) and correlations between parameters (d).

#### Data set: BEAVA_SE_survival

Table 3: Summary of parameter estimates (parameter d is set to 1) for BEAVA_SE_survival data set

| Parameter | median | Q2.5 | Q97.5 |
|---|---|---|---|
| b | 4.175 | 2.493 | 7.365 |
| e | 368.230 | 300.376 | 448.503 |

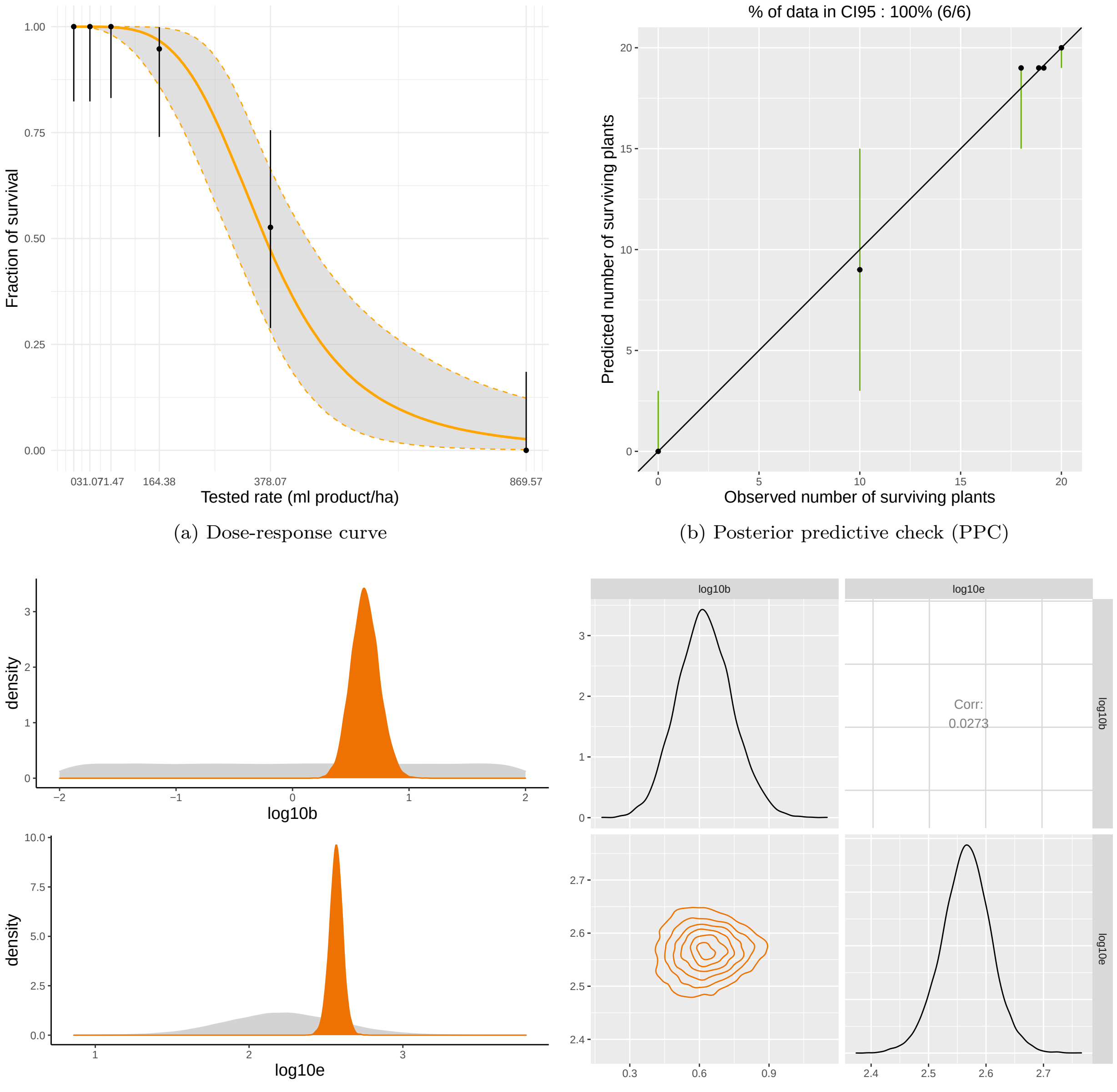

(a) Dose-response curve

(b) Posterior predictive check (PPC)

(c) Priors and posteriors

(d) Correlations between parameters

Figure 3: Dose-response curve (a), PPC (b), prior and posterior distributions (c) and correlations between parameters (d).

#### Data set: BRSNW_SE_survival

Table 4: Summary of parameter estimates (parameter d is set to 1) for BRSNW_SE_survival data set

| Parameter | median | Q2.5 | Q97.5 |
|---|---|---|---|
| b | 4.695 | 2.797 | 7.286 |
| e | 539.898 | 444.057 | 661.107 |

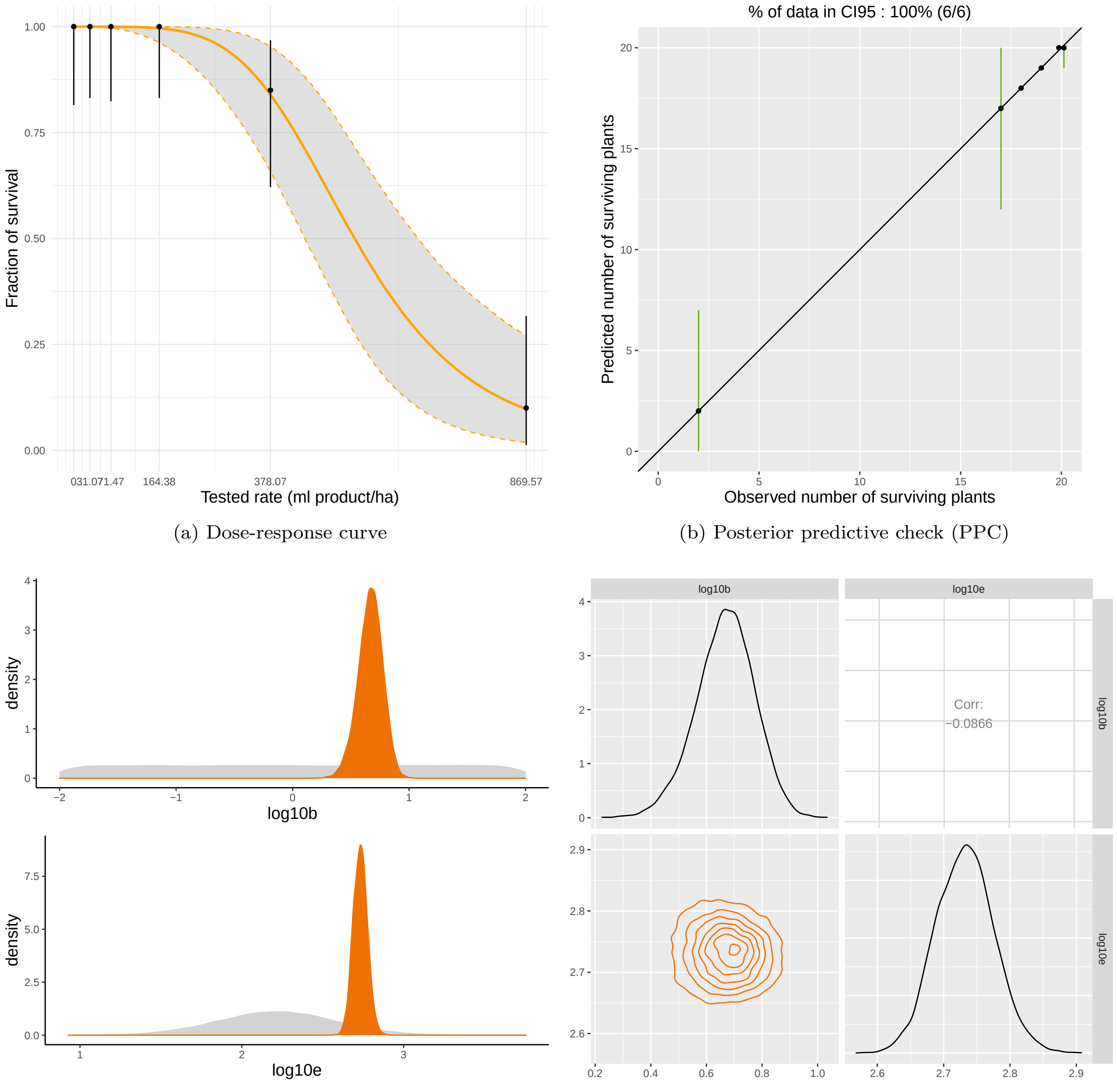

(a) Dose-response curve

(b) Posterior predictive check (PPC)

(c) Priors and posteriors

(d) Correlations between parameters

Figure 4: Dose-response curve (a), PPC (b), prior and posterior distributions (c) and correlations between parameters (d).

### Data set: CUMSA_SE_survival

Table 5: Summary of parameter estimates (parameter d is set to 1) for CUMSA_SE_survival data set

| Parameter | median | Q2.5 | Q97.5 |
|---|---|---|---|
| b | 2.482 | 1.507 | 3.814 |
| e | 555.747 | 430.348 | 754.010 |

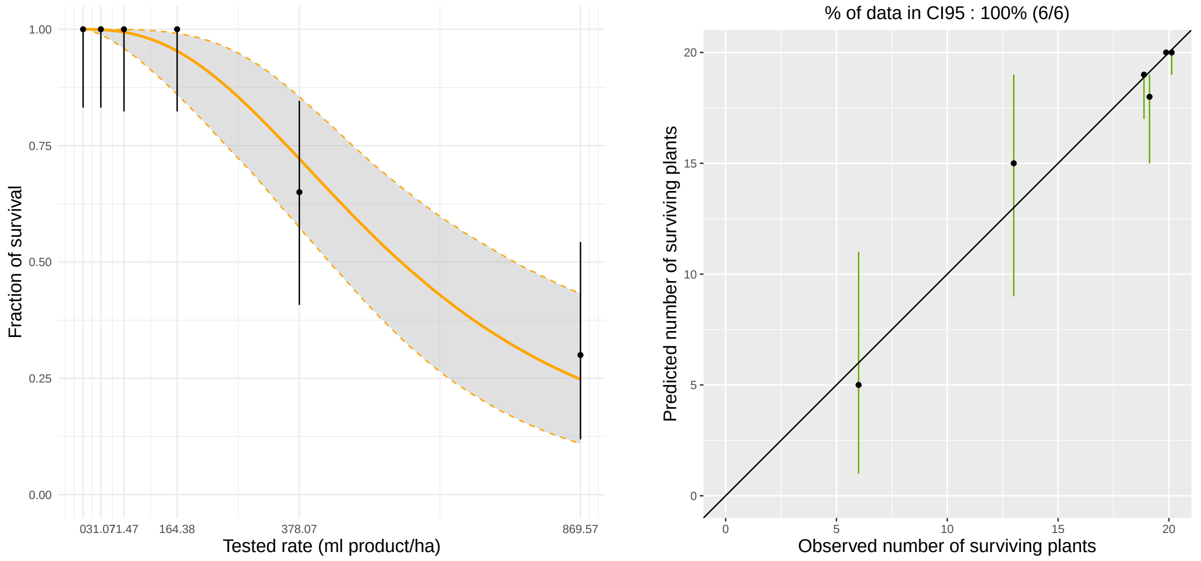

(a) Dose-response curve

(b) Posterior predictive check (PPC)

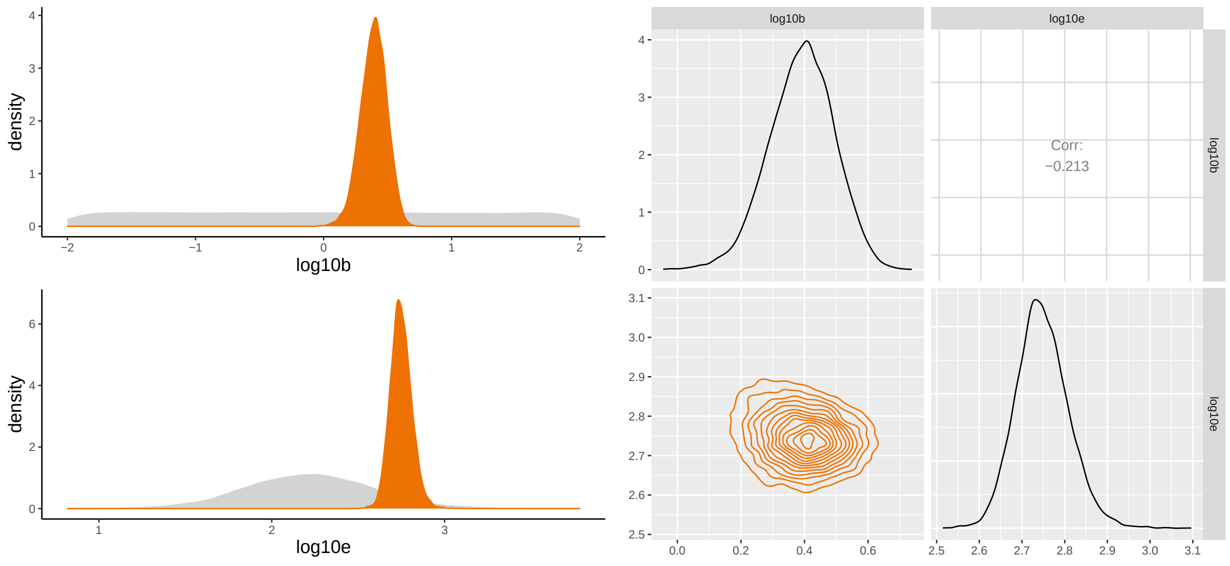

(c) Priors and posteriors

(d) Correlations between parameters

Figure 5: Dose-response curve (a), PPC (b), prior and posterior distributions (c) and correlations between parameters (d).

#### Data set: GLXMA_SE_survival

Table 6: Summary of parameter estimates (parameter d is set to 1) for GLXMA_SE_survival data set

| Parameter | median | Q2.5 | Q97.5 |
|---|---|---|---|
| b | 34.323 | 4.846 | 95.138 |
| e | 1270.804 | 931.242 | 2845.263 |

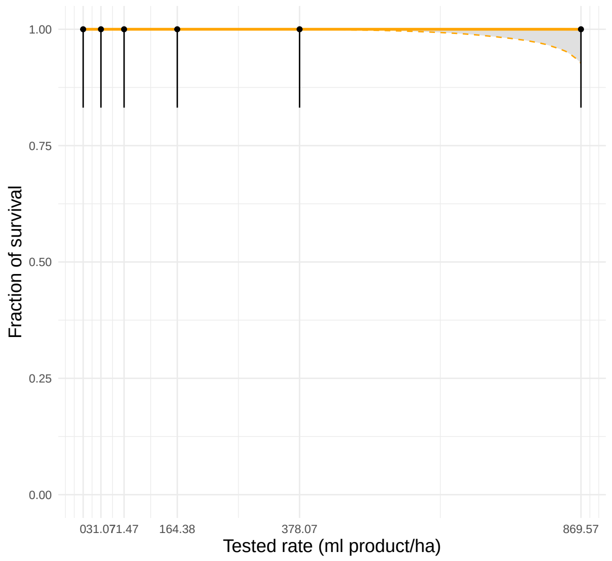

(a) Dose-response curve

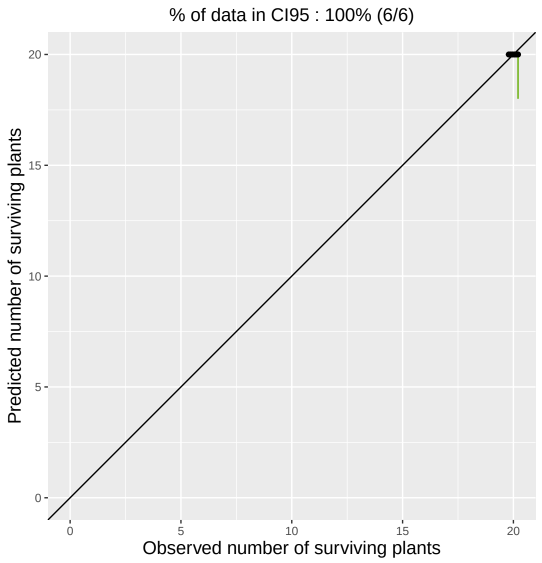

(b) Posterior predictive check (PPC)

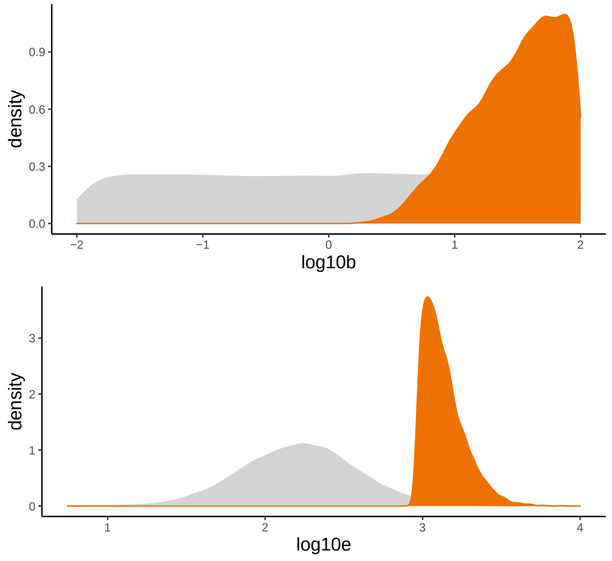

(c) Priors and posteriors

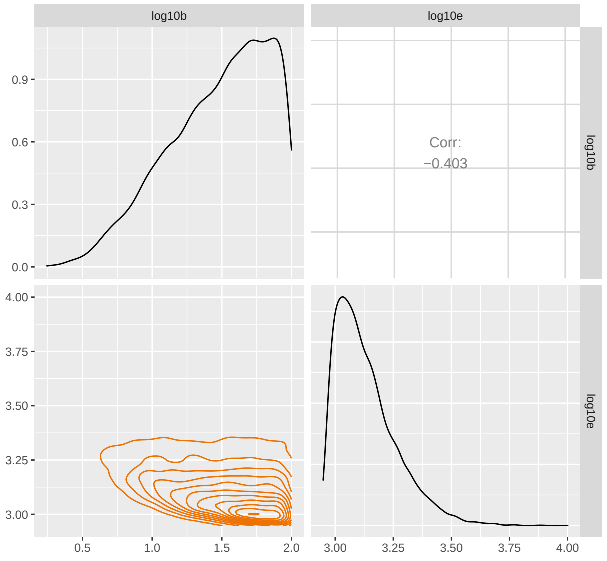

(d) Correlations between parameters

Figure 6: Dose-response curve (a), PPC (b), prior and posterior distributions (c) and correlations between parameters (d).

#### Data set: HELAN_SE_survival

Table 7: Summary of parameter estimates (parameter d is set to 1) for HELAN_SE_survival data set

| Parameter | median | Q2.5 | Q97.5 |
|---|---|---|---|
| b | 33.830 | 5.064 | 95.658 |
| e | 1269.059 | 930.788 | 2936.663 |

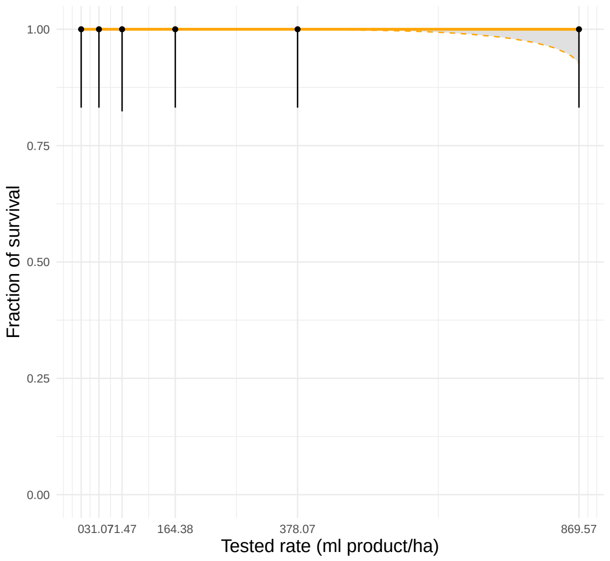

(a) Dose-response curve

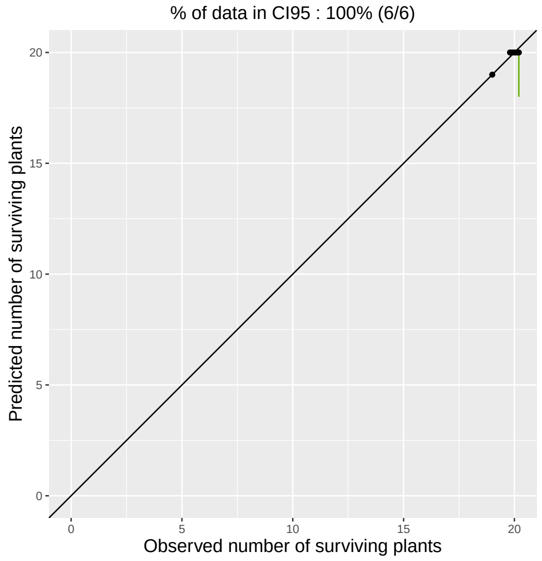

(b) Posterior predictive check (PPC)

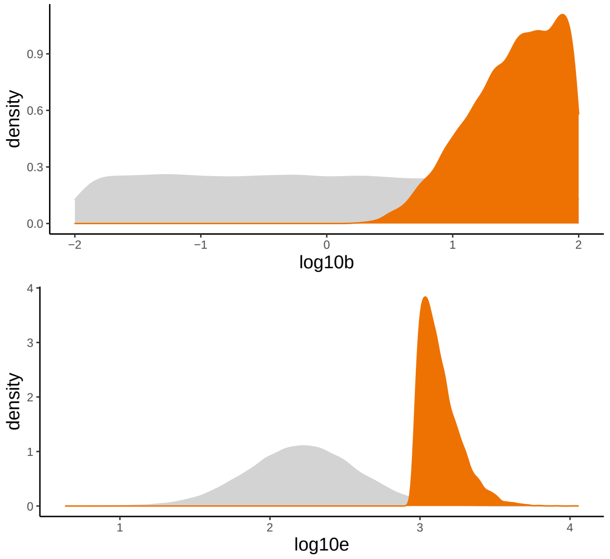

(c) Priors and posteriors

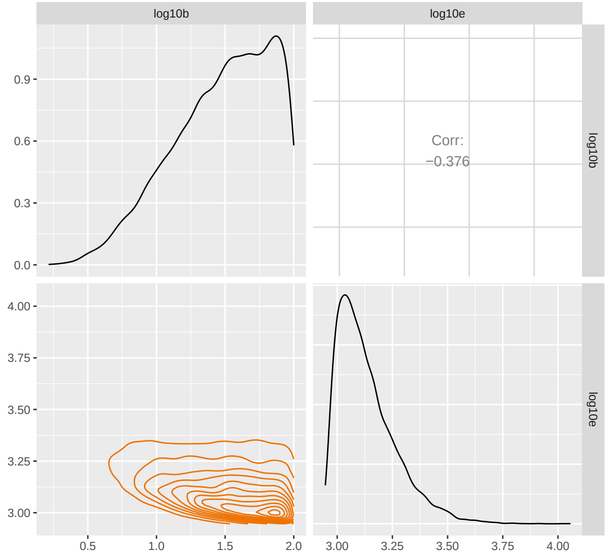

(d) Correlations between parameters

Figure 7: Dose-response curve (a), PPC (b), prior and posterior distributions (c) and correlations between parameters (d).

#### Data set: LYPES_SE_survival

Table 8: Summary of parameter estimates (parameter d is set to 1) for LYPES_SE_survival data set

| Parameter | median | Q2.5 | Q97.5 |
|---|---|---|---|
| b | 11.791 | 2.548 | 77.385 |
| e | 1108.045 | 901.029 | 2353.726 |

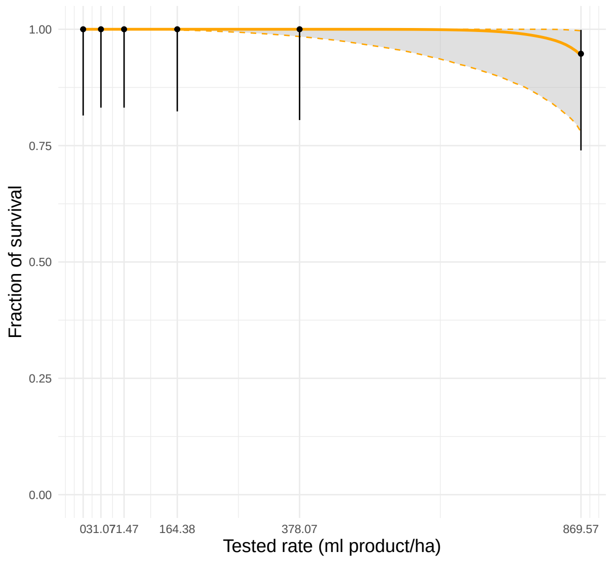

(a) Dose-response curve

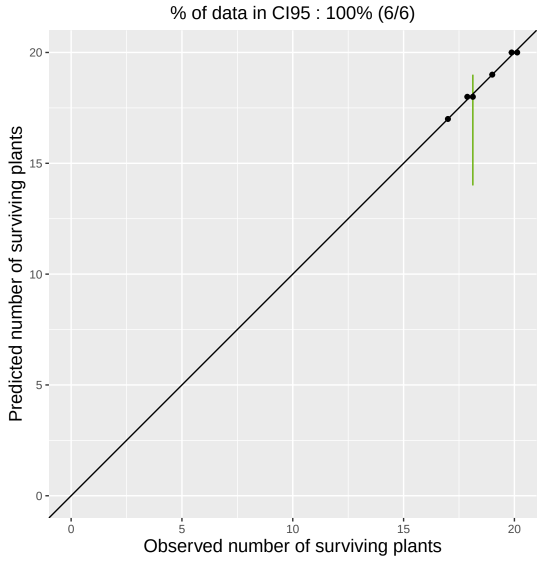

(b) Posterior predictive check (PPC)

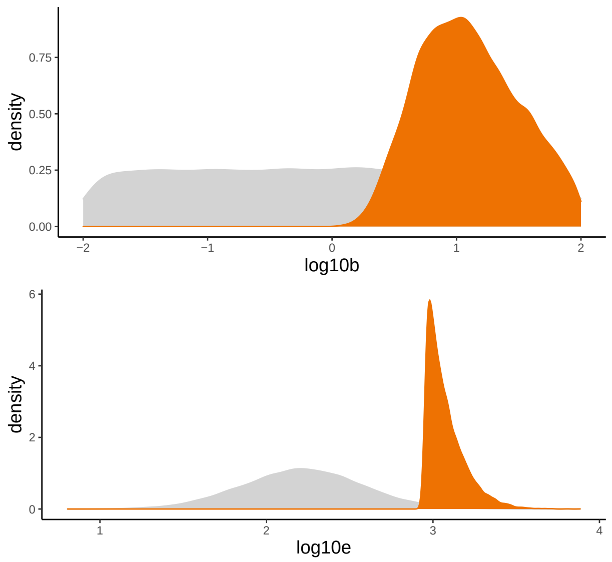

(c) Priors and posteriors

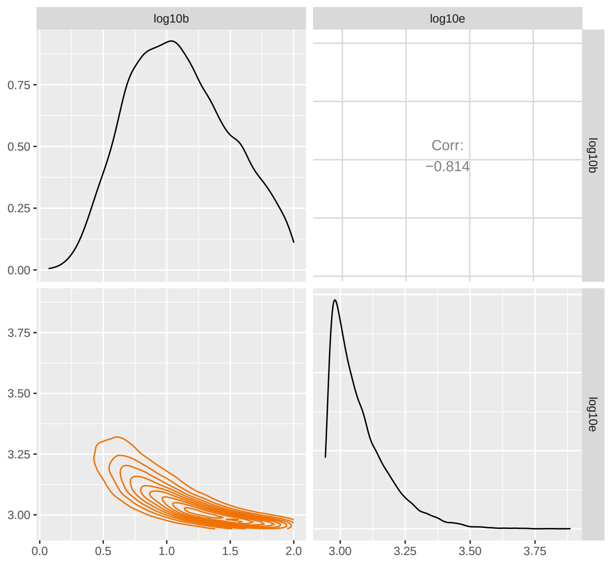

(d) Correlations between parameters

Figure 8: Dose-response curve (a), PPC (b), prior and posterior distributions (c) and correlations between parameters (d).

#### Data set: TRZAW_SE_survival

Table 9: Summary of parameter estimates (parameter d is set to 1) for TRZAW_SE_survival data set

| Parameter | median | Q2.5 | Q97.5 |
|---|---|---|---|
| b | 2.172 | 1.212 | 3.659 |
| e | 1938.426 | 1369.998 | 3381.989 |

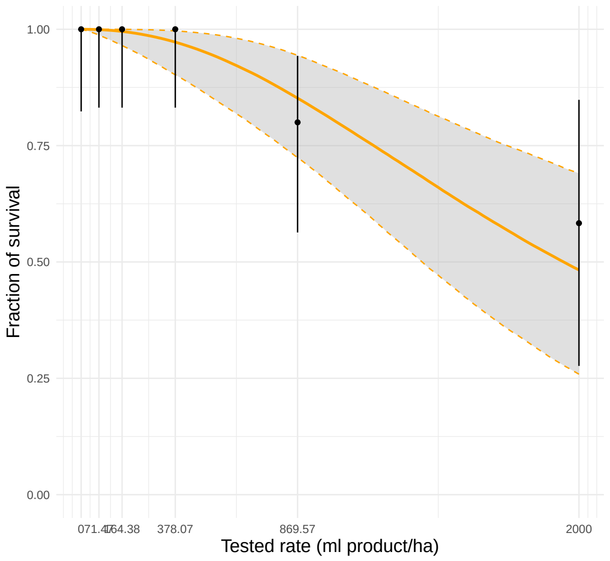

(a) Dose-response curve

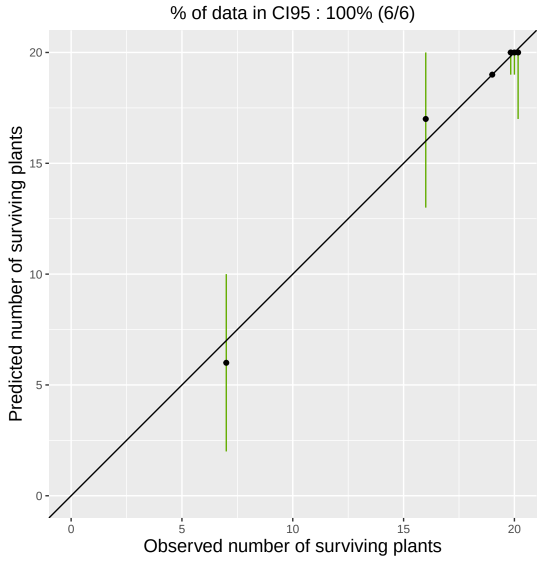

(b) Posterior predictive check (PPC)

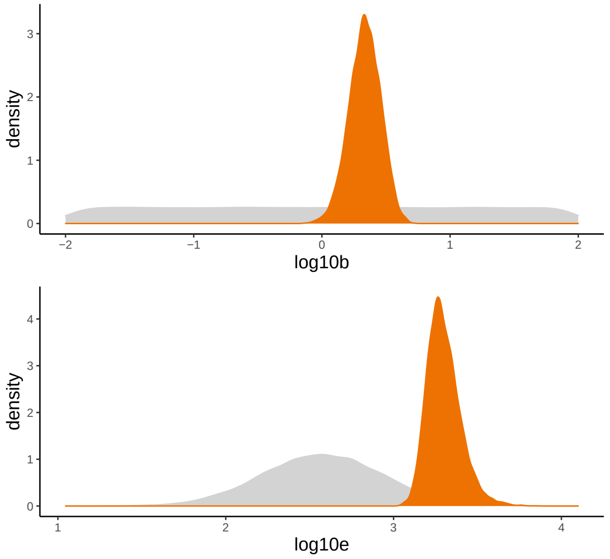

(c) Priors and posteriors

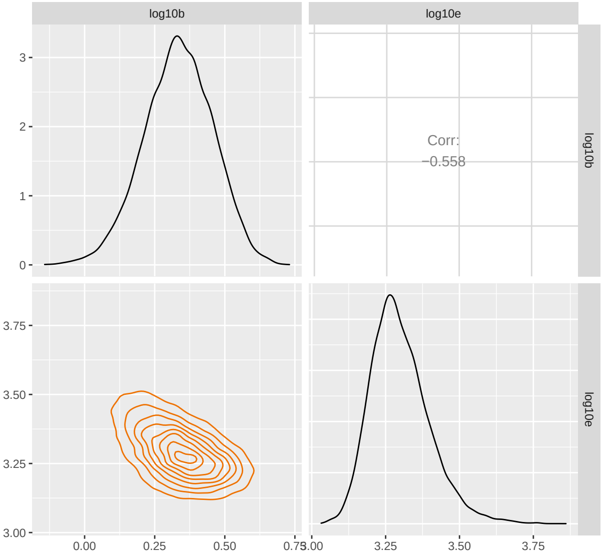

(d) Correlations between parameters

Figure 9: Dose-response curve (a), PPC (b), prior and posterior distributions (c) and correlations between parameters (d).

#### Data set: ZEAMA_SE_survival

Table 10: Summary of parameter estimates for ZEAMA_SE_survival data set

| Parameter | median | Q2.5 | Q97.5 |
|---|---|---|---|
| b | 15.267 | 1.823 | 90.674 |
| d | 0.960 | 0.911 | 0.991 |
| e | 2794.569 | 2085.595 | 6353.985 |

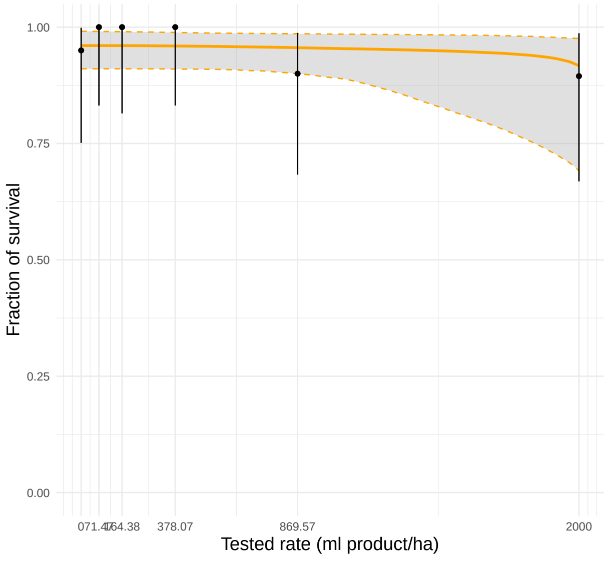

(a) Dose-response curve

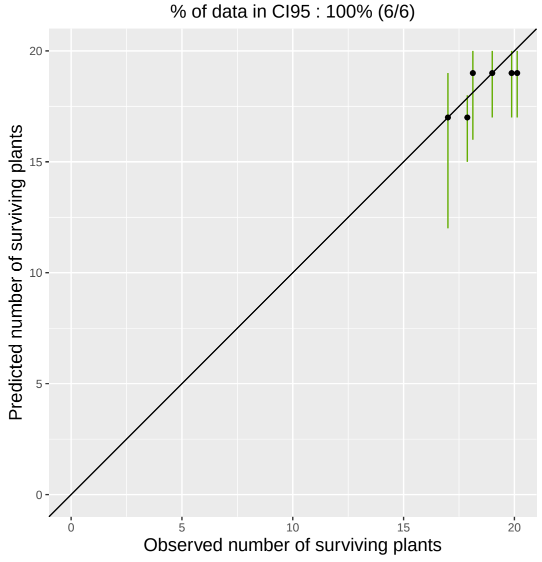

(b) Posterior predictive check (PPC)

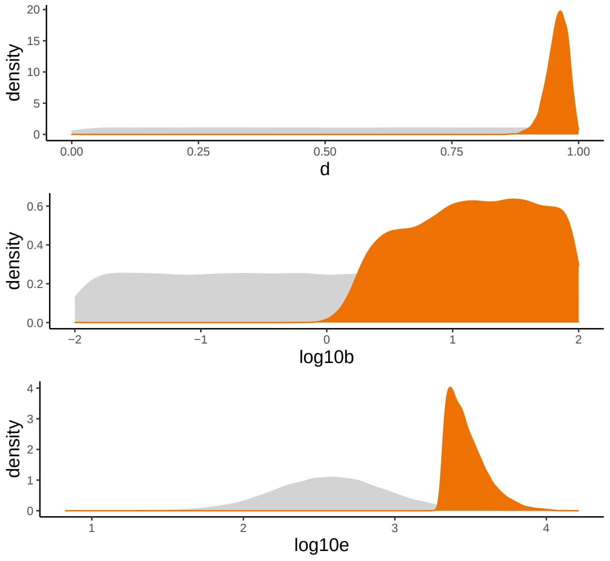

(c) Priors and posteriors

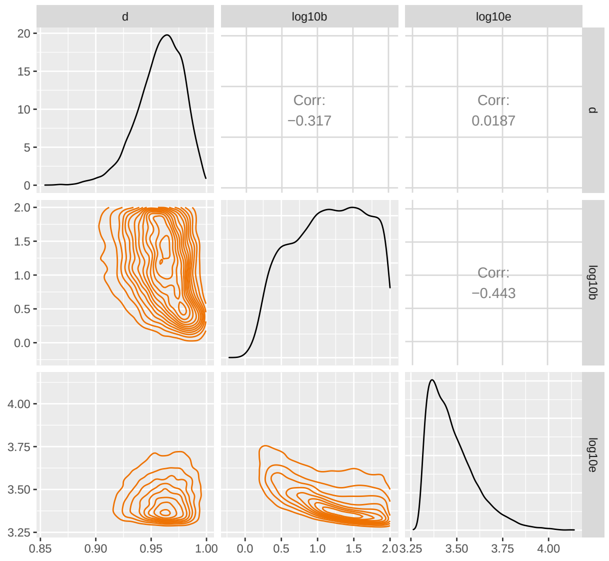

(d) Correlations between parameters

Figure 10: Dose-response curve (a), PPC (b), prior and posterior distributions (c) and correlations between parameters (d).
