## Supplementary material for "How to account for the uncertainty from standard toxicity tests in species sensitivity distributions: an example in non-target plants": Archive with reports on studies: report_VV_survival.pdf

### Dose-response analyses

Study 1

Vegetative Vigour test - survival endpoint

25 June 2020

Contact:

---

This is a report which provides results on all performed dose-response analyses for the survival endpoint of the Vegetative Vigour test for study 1.

---

#### Data set: ALLCE_VV_survival

Table 1: Summary of parameter estimates (parameter d is set to 1) for ALLCE_VV_survival data set

| Parameter | median | Q2.5 | Q97.5 |
|---|---|---|---|
| b | 33.329 | 4.842 | 95.573 |
| e | 1268.599 | 931.012 | 2901.631 |

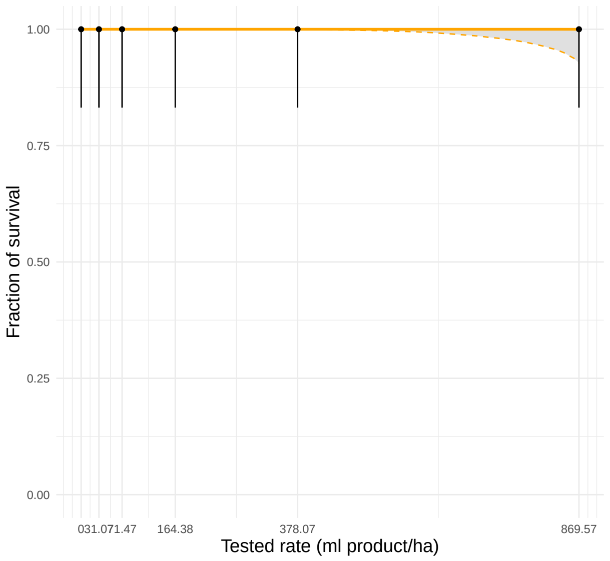

(a) Dose-response curve

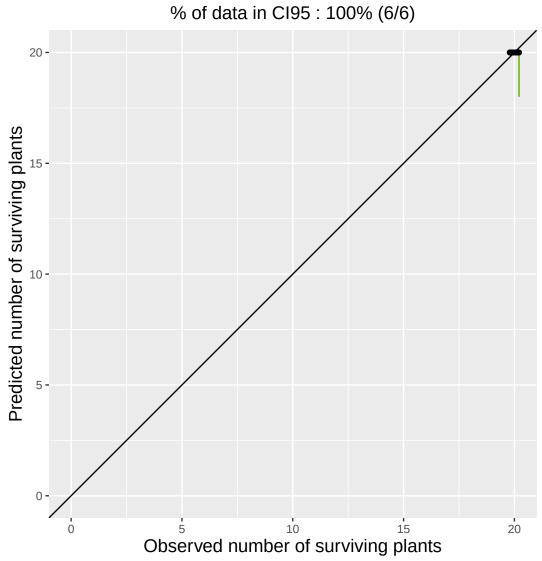

(b) Posterior predictive check (PPC)

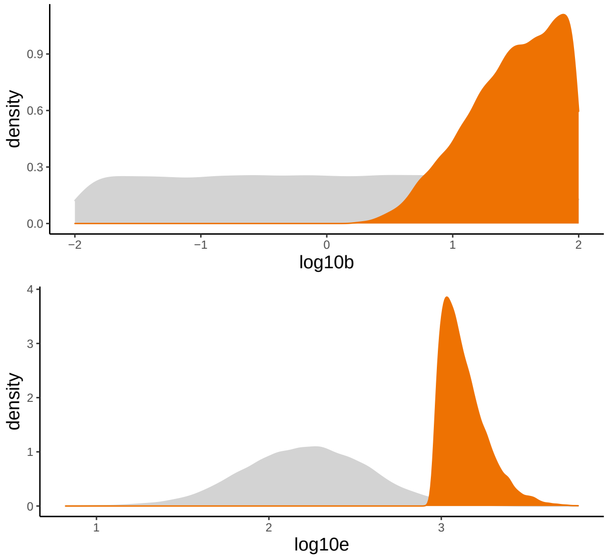

(c) Priors and posteriors

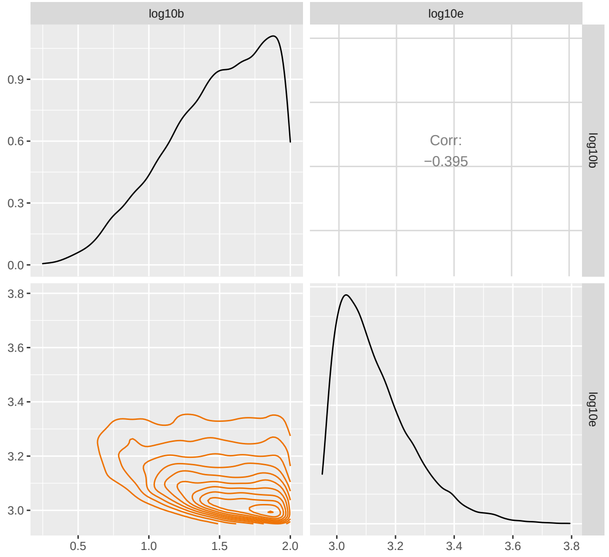

(d) Correlations between parameters

Figure 1: Dose-response curve (a), PPC (b), prior and posterior distributions (c) and correlations between parameters (d).

#### Data set: AVESA_VV_survival

Table 2: Summary of parameter estimates (parameter d is set to 1) for AVESA_VV_survival data set

| Parameter | median | Q2.5 | Q97.5 |
|---|---|---|---|
| b | 34.022 | 5.052 | 95.518 |
| e | 2920.813 | 2147.134 | 6425.718 |

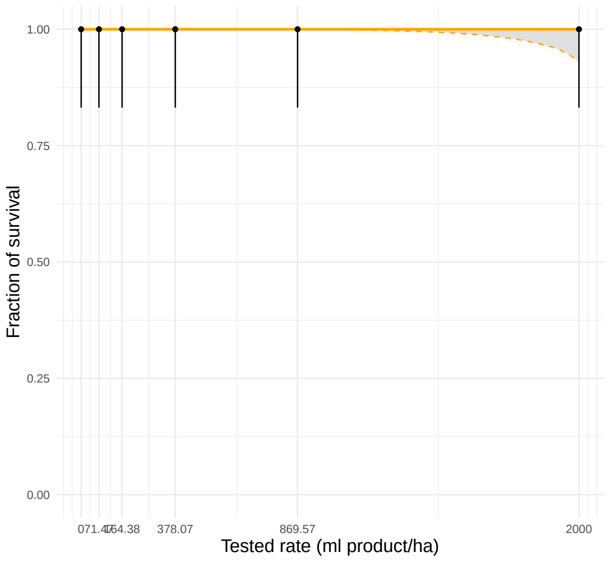

(a) Dose-response curve

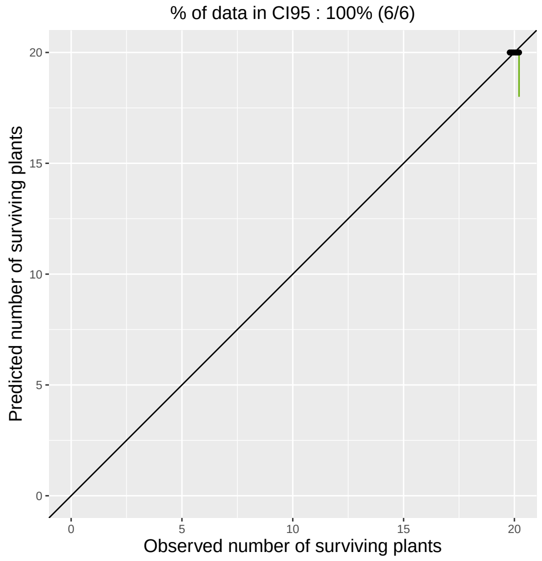

(b) Posterior predictive check (PPC)

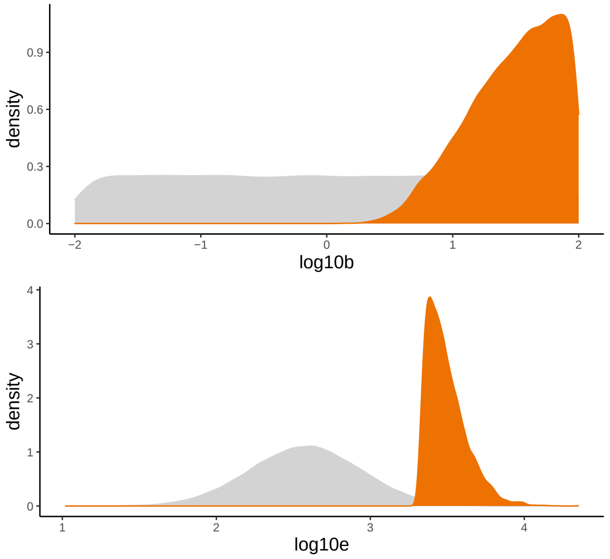

(c) Priors and posteriors

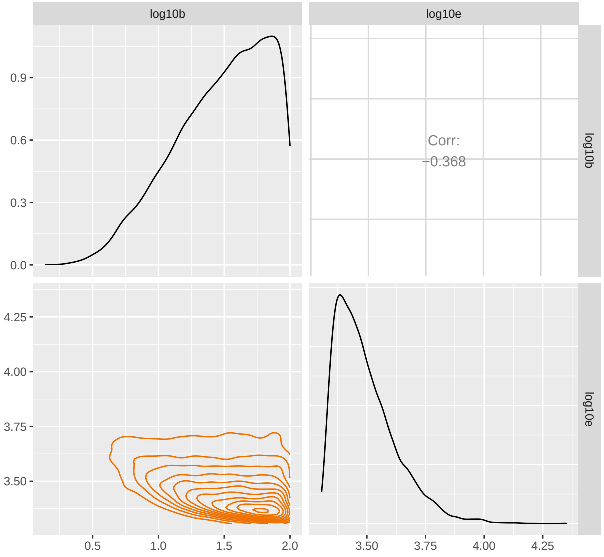

(d) Correlations between parameters

Figure 2: Dose-response curve (a), PPC (b), prior and posterior distributions (c) and correlations between parameters (d).

#### Data set: BEAVA_VV_survival

Table 3: Summary of parameter estimates (parameter d is set to 1) for BEAVA_VV_survival data set

| Parameter | median | Q2.5 | Q97.5 |
|---|---|---|---|
| b | 34.465 | 4.823 | 95.069 |
| e | 1267.001 | 933.152 | 2950.169 |

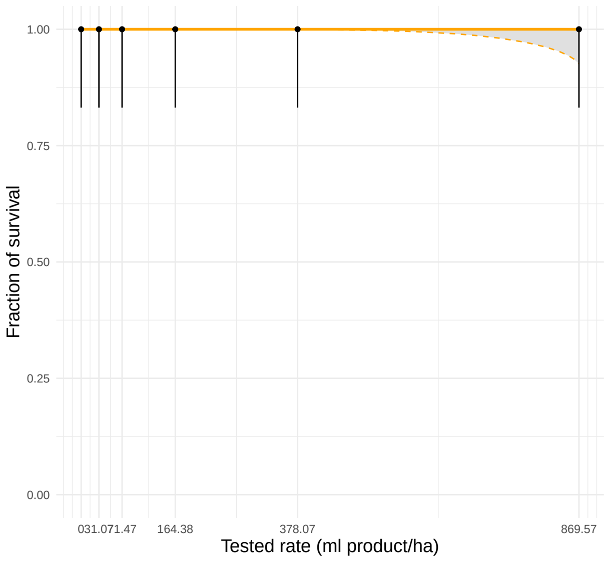

(a) Dose-response curve

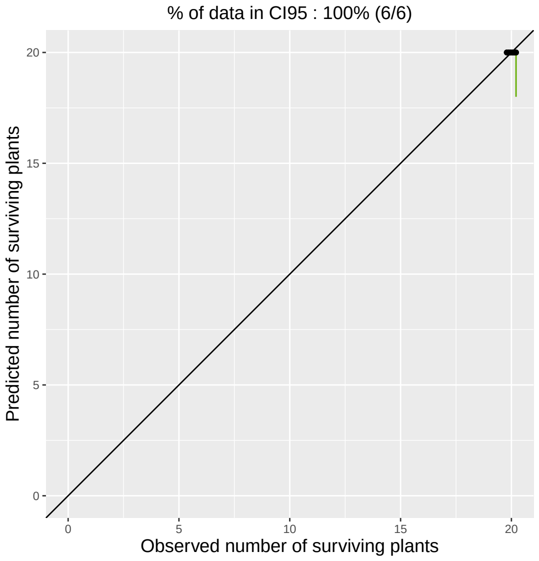

(b) Posterior predictive check (PPC)

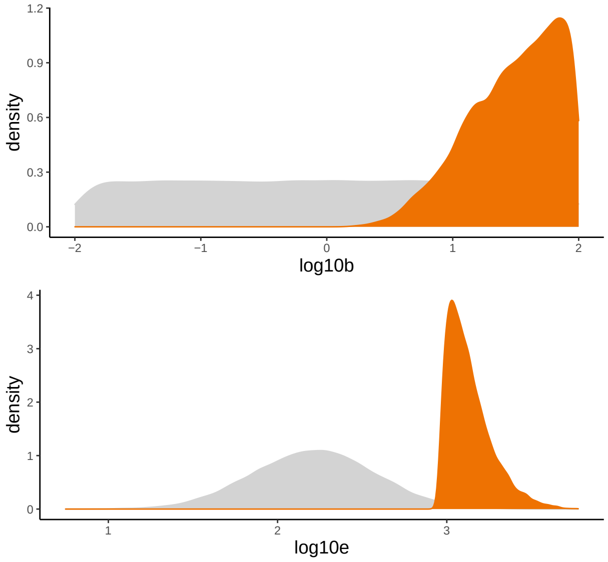

(c) Priors and posteriors

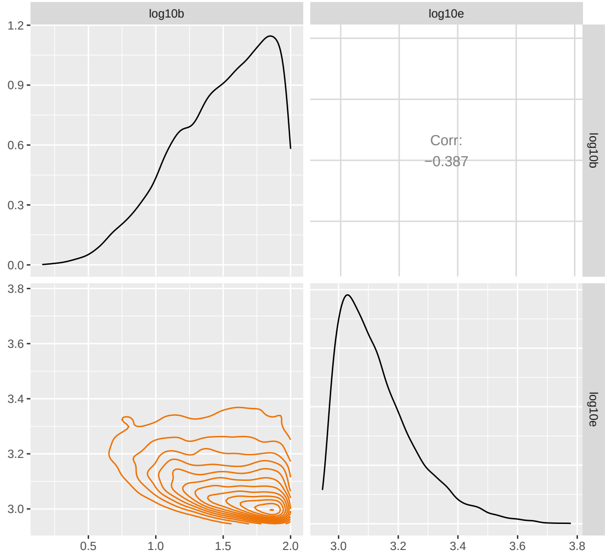

(d) Correlations between parameters

Figure 3: Dose-response curve (a), PPC (b), prior and posterior distributions (c) and correlations between parameters (d).

### Data set: BRSNW_VV_survival

Table 4: Summary of parameter estimates (parameter d is set to 1) for BRSNW_VV_survival data set

| Parameter | median | Q2.5 | Q97.5 |
|---|---|---|---|
| b | 7.288 | 2.213 | 62.937 |
| e | 956.049 | 862.972 | 1475.258 |

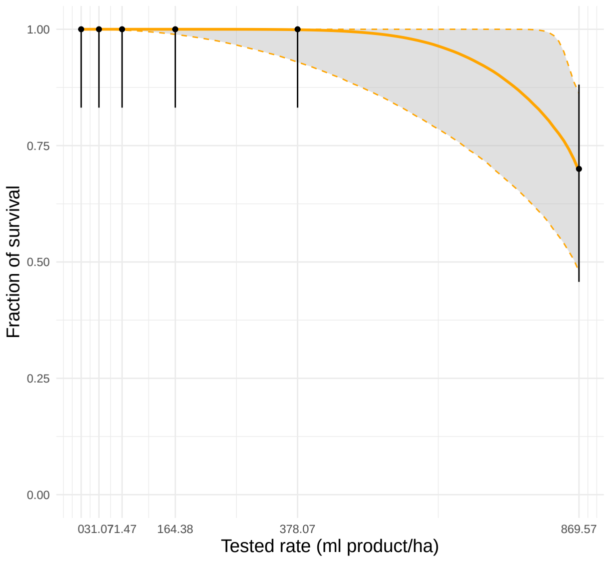

(a) Dose-response curve

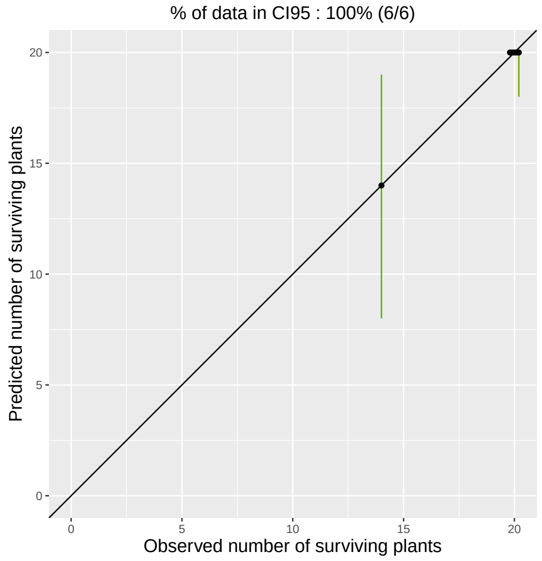

(b) Posterior predictive check (PPC)

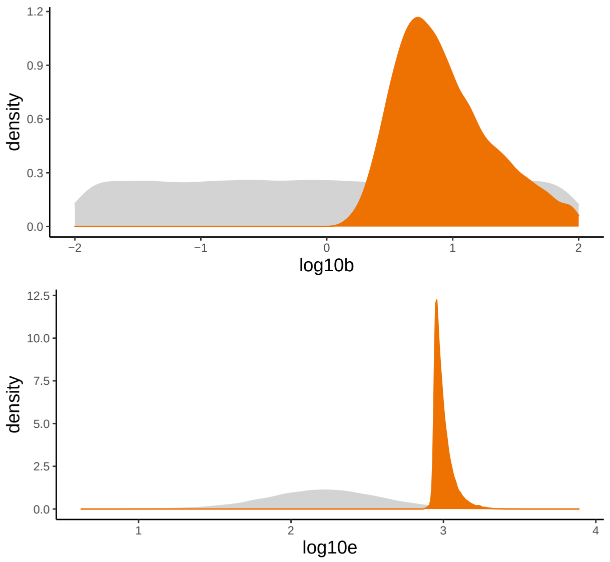

(c) Priors and posteriors

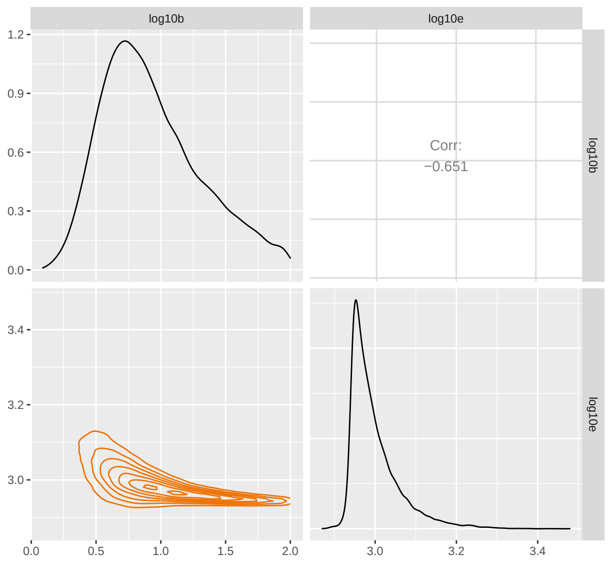

(d) Correlations between parameters

Figure 4: Dose-response curve (a), PPC (b), prior and posterior distributions (c) and correlations between parameters (d).

#### Data set: CUMSA_VV_survival

Table 5: Summary of parameter estimates (parameter d is set to 1) for CUMSA_VV_survival data set

| Parameter | median | Q2.5 | Q97.5 |
|---|---|---|---|
| b | 11.819 | 2.716 | 78.563 |
| e | 1113.093 | 901.926 | 2351.503 |

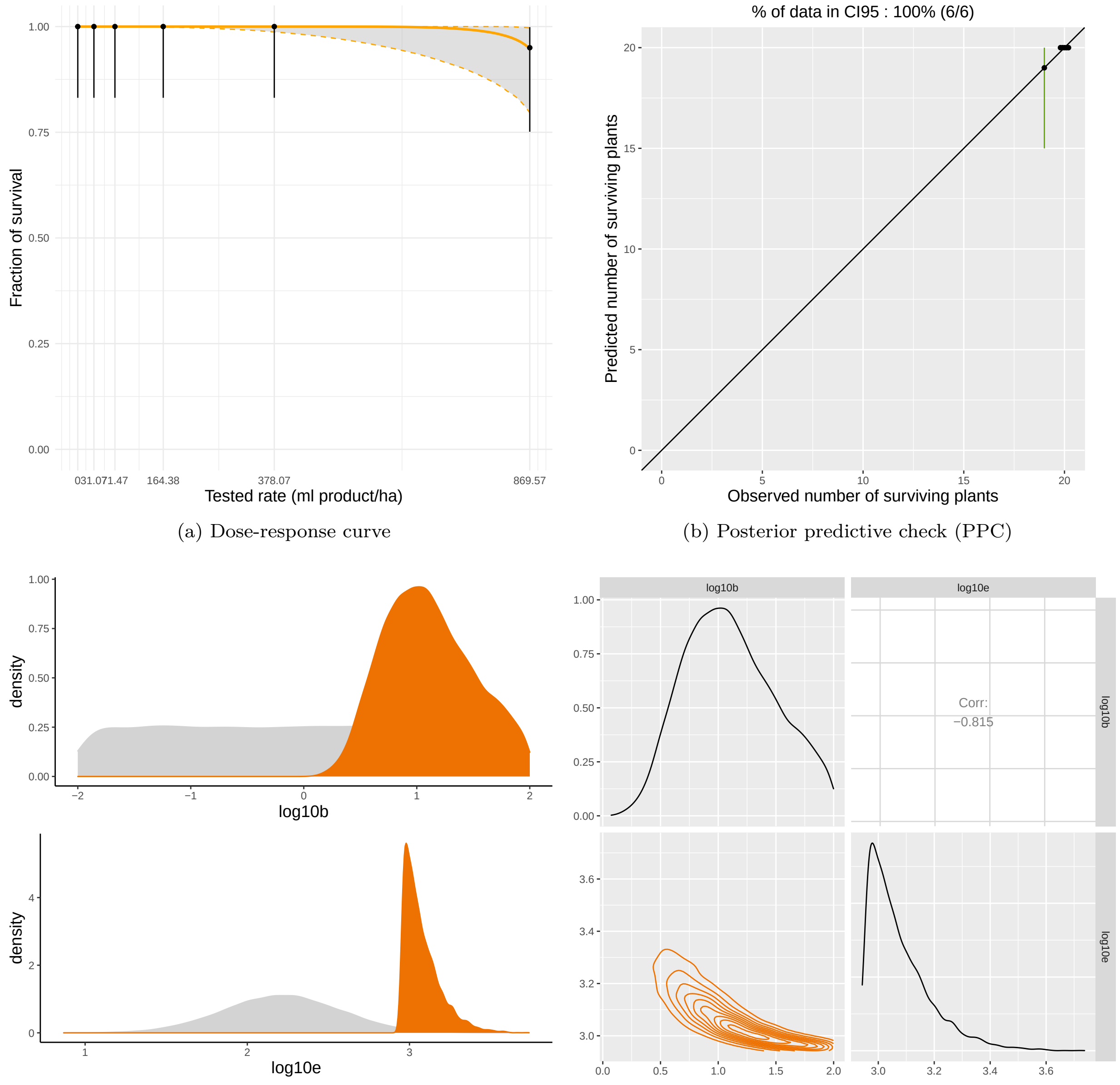

(a) Dose-response curve

(b) Posterior predictive check (PPC)

(c) Priors and posteriors

(d) Correlations between parameters

Figure 5: Dose-response curve (a), PPC (b), prior and posterior distributions (c) and correlations between parameters (d).

### Data set: GLXMA_VV_survival

Table 6: Summary of parameter estimates (parameter d is set to 1) for GLXMA_VV_survival data set

| Parameter | median | Q2.5 | Q97.5 |
|---|---|---|---|
| b | 33.407 | 4.837 | 94.562 |
| e | 1272.933 | 934.586 | 2917.581 |

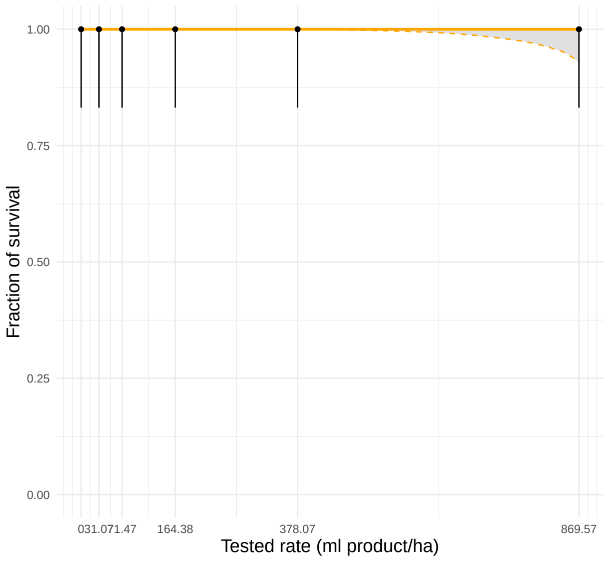

(a) Dose-response curve

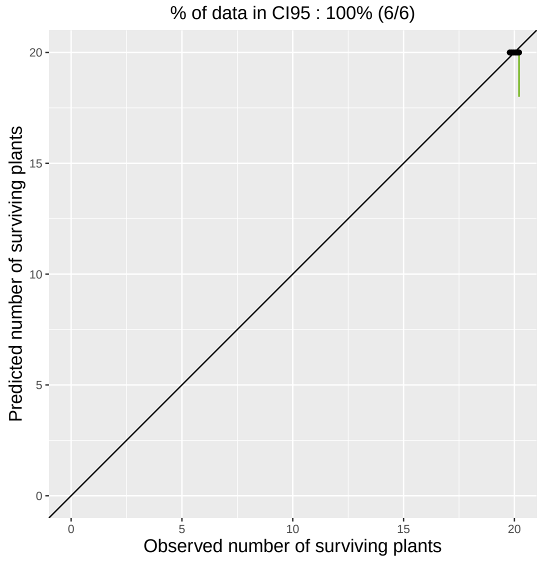

(b) Posterior predictive check (PPC)

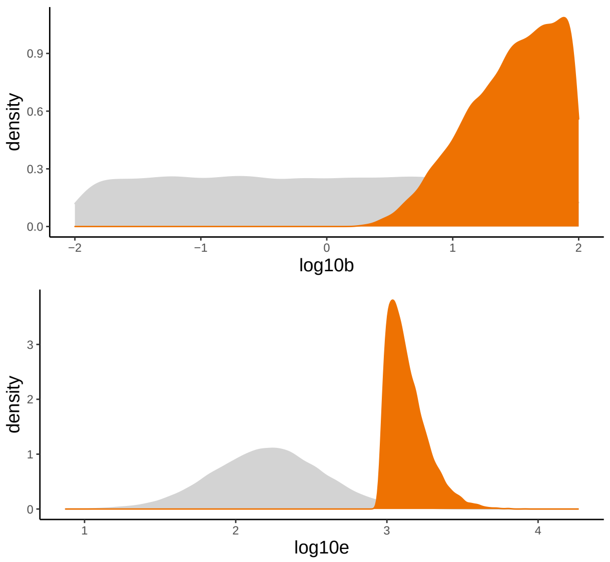

(c) Priors and posteriors

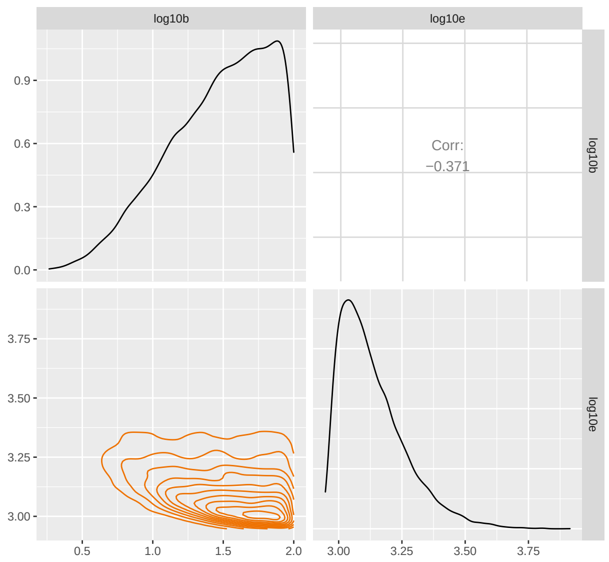

(d) Correlations between parameters

Figure 6: Dose-response curve (a), PPC (b), prior and posterior distributions (c) and correlations between parameters (d).

#### Data set: HELAN_VV_survival

Table 7: Summary of parameter estimates (parameter d is set to 1) for HELAN_VV_survival data set

| Parameter | median | Q2.5 | Q97.5 |
|---|---|---|---|
| b | 33.614 | 4.788 | 94.815 |
| e | 1271.976 | 934.983 | 2842.151 |

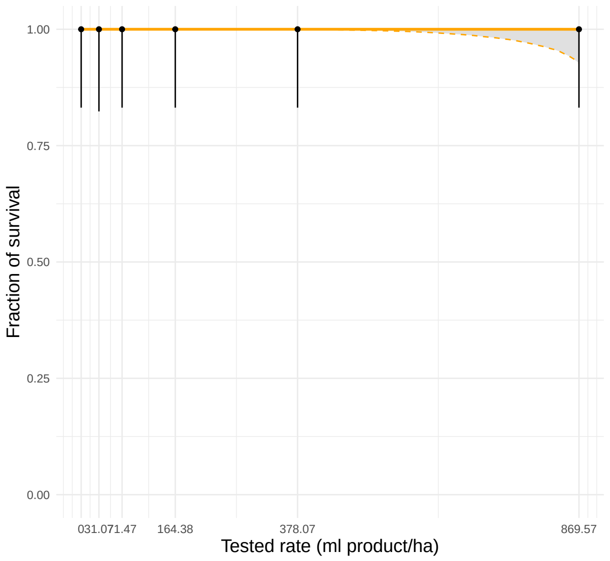

(a) Dose-response curve

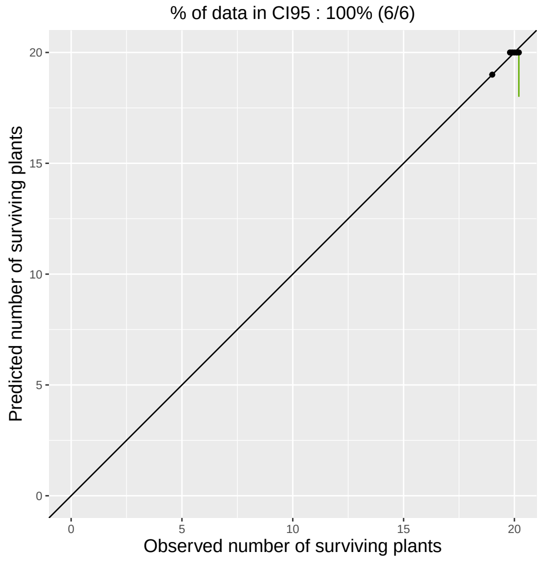

(b) Posterior predictive check (PPC)

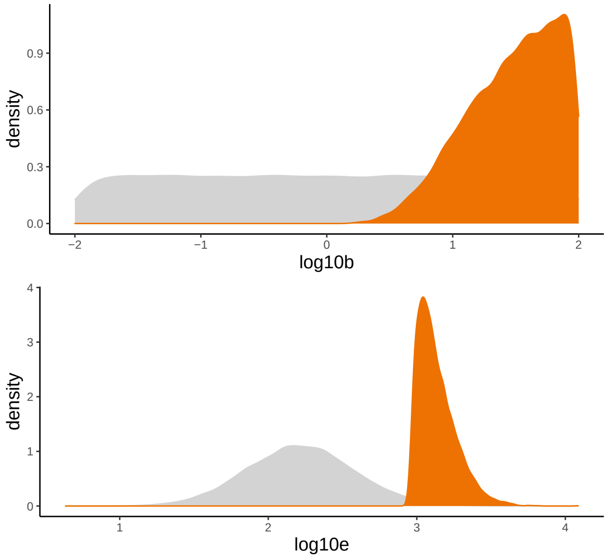

(c) Priors and posteriors

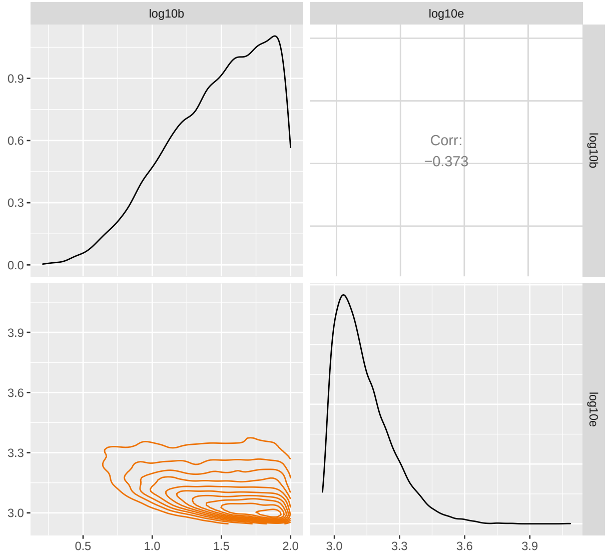

(d) Correlations between parameters

Figure 7: Dose-response curve (a), PPC (b), prior and posterior distributions (c) and correlations between parameters (d).

#### Data set: LYPES_VV_survival

Table 8: Summary of parameter estimates (parameter d is set to 1) for LYPES_VV_survival data set

| Parameter | median | Q2.5 | Q97.5 |
|---|---|---|---|
| b | 34.464 | 4.868 | 94.787 |
| e | 1268.730 | 933.630 | 2893.262 |

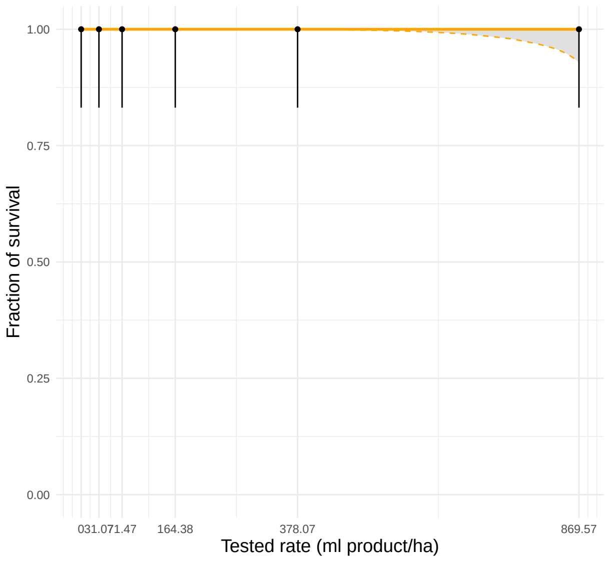

(a) Dose-response curve

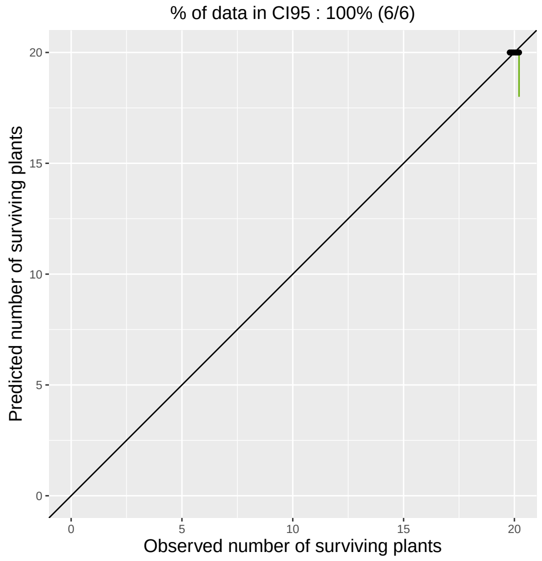

(b) Posterior predictive check (PPC)

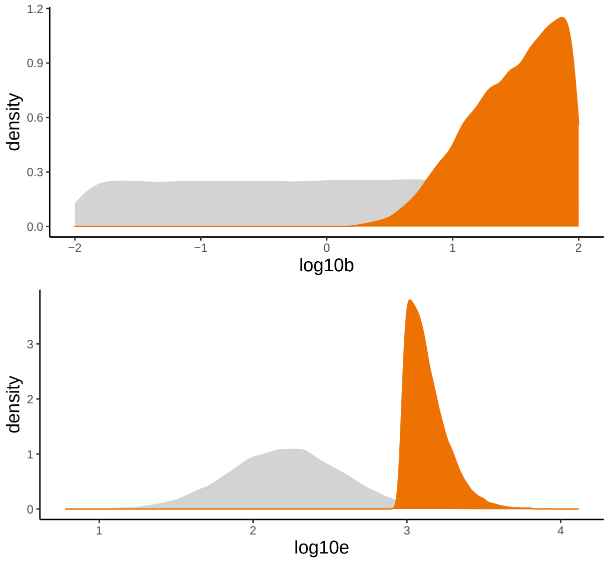

(c) Priors and posteriors

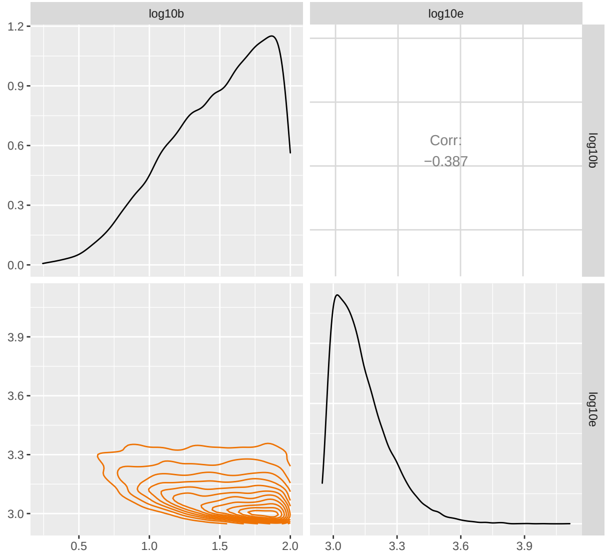

(d) Correlations between parameters

Figure 8: Dose-response curve (a), PPC (b), prior and posterior distributions (c) and correlations between parameters (d).

#### Data set: TRZAW_VV_survival

Table 9: Summary of parameter estimates (parameter d is set to 1) for TRZAW_VV_survival data set

| Parameter | median | Q2.5 | Q97.5 |
|---|---|---|---|
| b | 33.757 | 4.746 | 95.143 |
| e | 2930.090 | 2147.228 | 6668.731 |

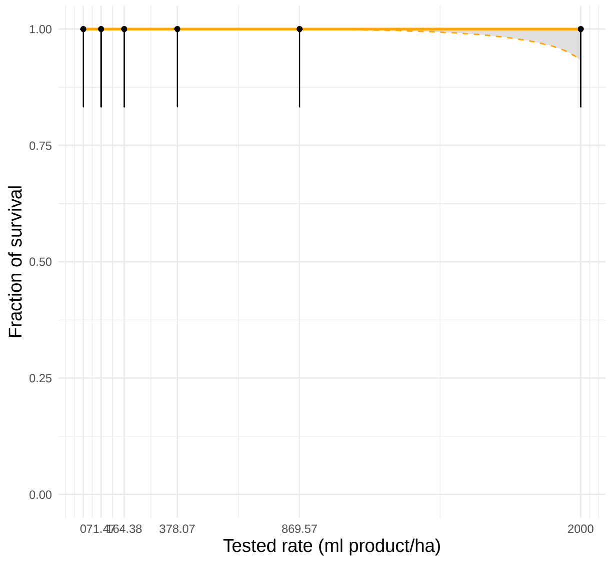

(a) Dose-response curve

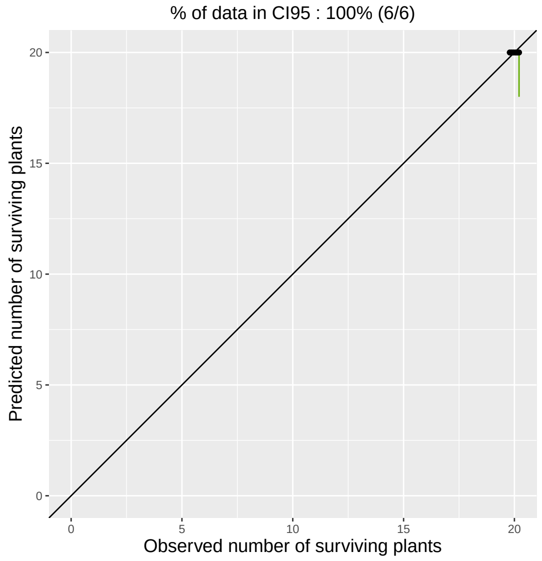

(b) Posterior predictive check (PPC)

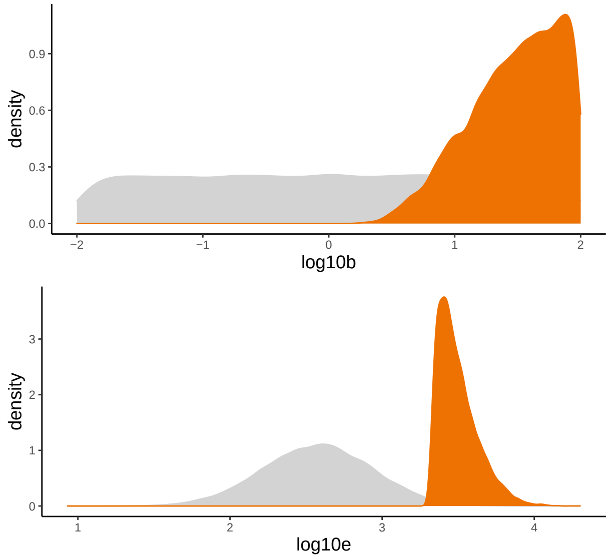

(c) Priors and posteriors

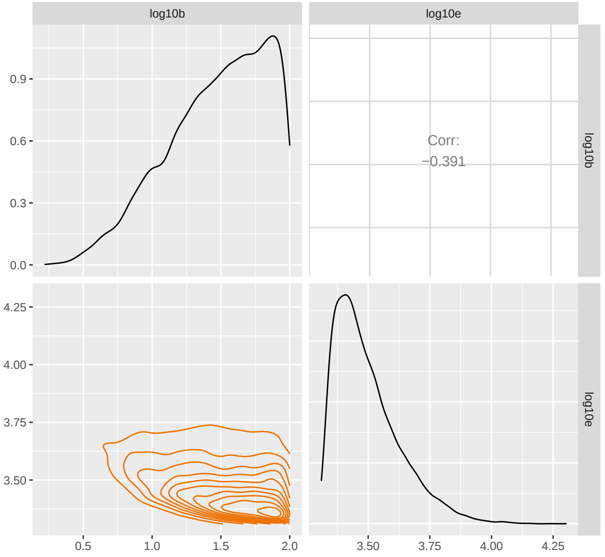

(d) Correlations between parameters

Figure 9: Dose-response curve (a), PPC (b), prior and posterior distributions (c) and correlations between parameters (d).

#### Data set: ZEAMA_VV_survival

Table 10: Summary of parameter estimates (parameter d is set to 1) for ZEAMA_VV_survival data set

| Parameter | median | Q2.5 | Q97.5 |
|---|---|---|---|
| b | 34.276 | 4.920 | 95.269 |
| e | 2901.501 | 2144.839 | 6692.480 |

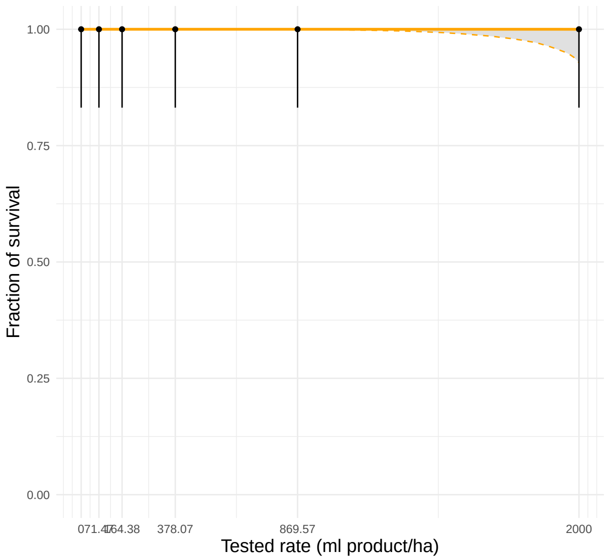

(a) Dose-response curve

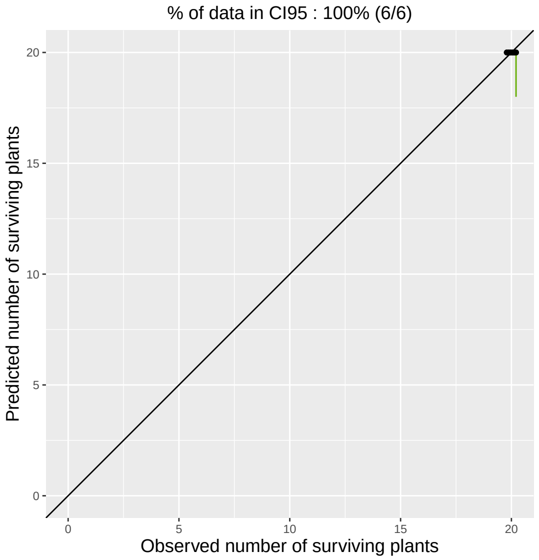

(b) Posterior predictive check (PPC)

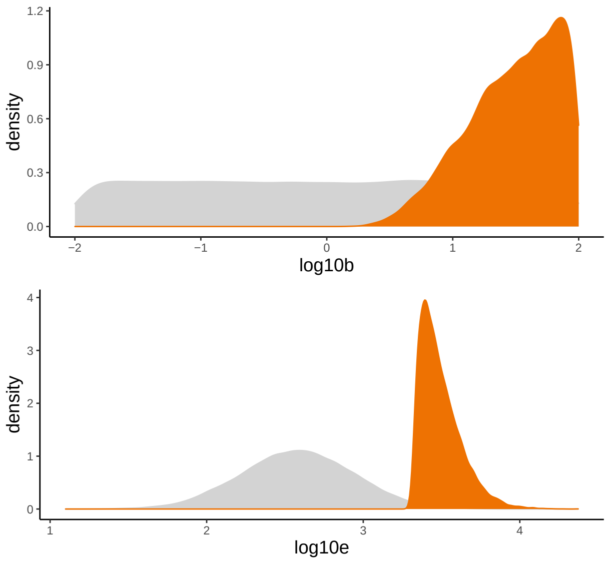

(c) Priors and posteriors

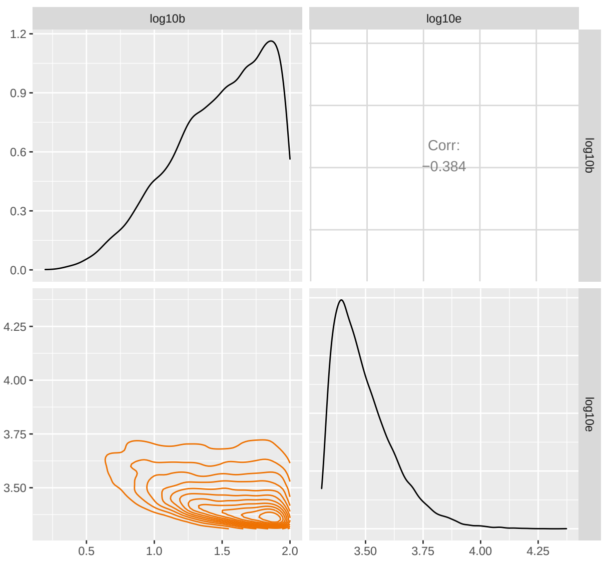

(d) Correlations between parameters

Figure 10: Dose-response curve (a), PPC (b), prior and posterior distributions (c) and correlations between parameters (d).
