## Supplementary material for "How to account for the uncertainty from standard toxicity tests in species sensitivity distributions: an example in non-target plants": Archive with reports on studies: report_VV_weight.pdf

### Dose-response analysis

Study 1

Vegetative Vigour test - shoot dry VV_weight endpoint

25 June 2020

Contact:

---

This is a report which provides results on all performed dose-response analyses for the shoot dry VV_weight endpoint of the Vegetative Vigour test for study 1.

---

#### Data set: ALLCE_VV_weight

Table 1: Summary of parameter estimates for ALLCE_VV_weight data set

| Parameter | median | Q2.5 | Q97.5 |
|---|---|---|---|
| b | 0.379 | 0.014 | 32.599 |
| d | 0.306 | 0.211 | 0.387 |
| e | 656.817 | 76.043 | 1945.381 |
| sigma | 0.082 | 0.062 | 0.114 |

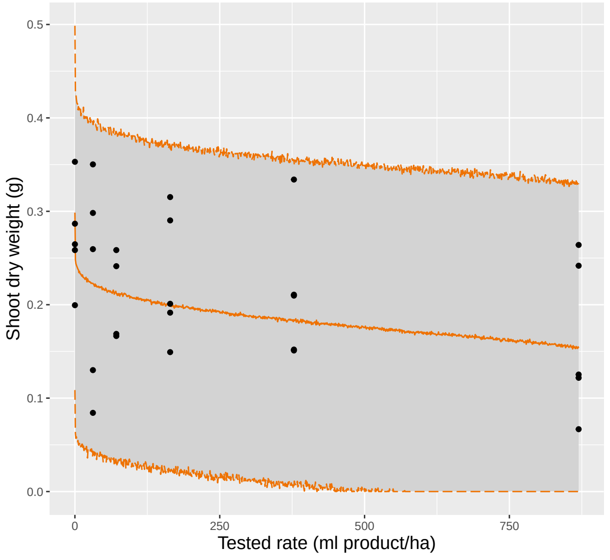

(a) Dose-response curve

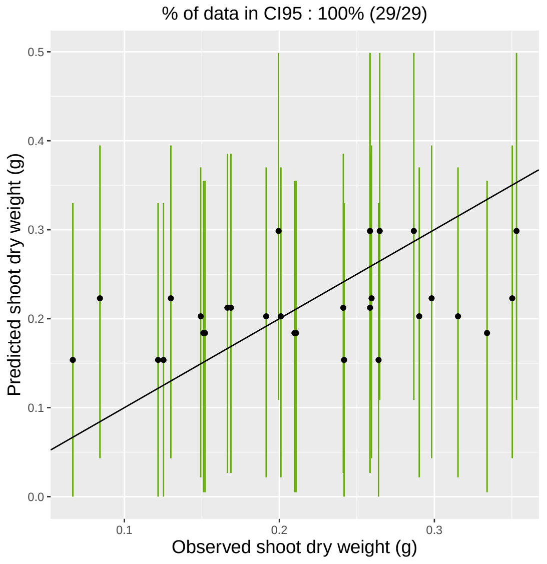

(b) Posterior predictive check (PPC)

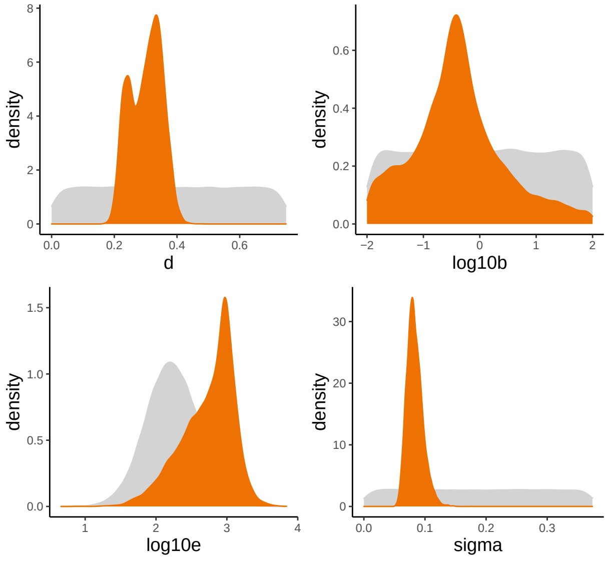

(c) Priors and posteriors

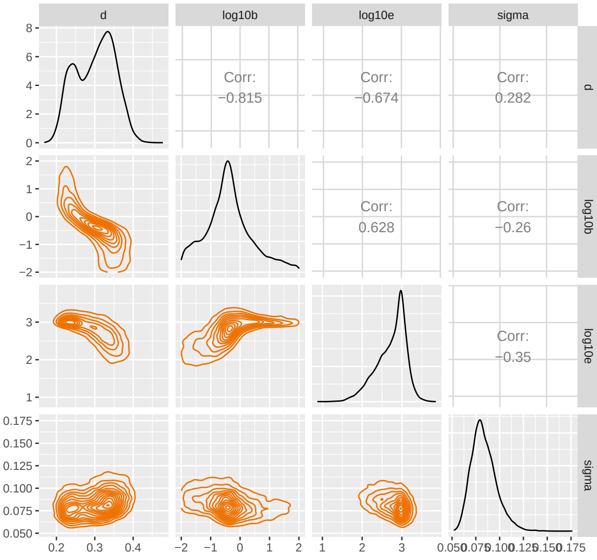

(d) Correlations between parameters

Figure 1: Dose-response curve (a), PPC (b), prior and posterior distributions (c) and correlations between parameters (d).

#### Data set: AVESA_VV_weight

Table 2: Summary of parameter estimates for AVESA_VV_weight data set

| Parameter | median | Q2.5 | Q97.5 |
|---|---:|---:|---:|
| b | 0.585 | 0.380 | 0.950 |
| d | 1.992 | 1.804 | 2.199 |
| e | 2235.493 | 1330.006 | 3967.520 |
| sigma | 0.196 | 0.152 | 0.270 |

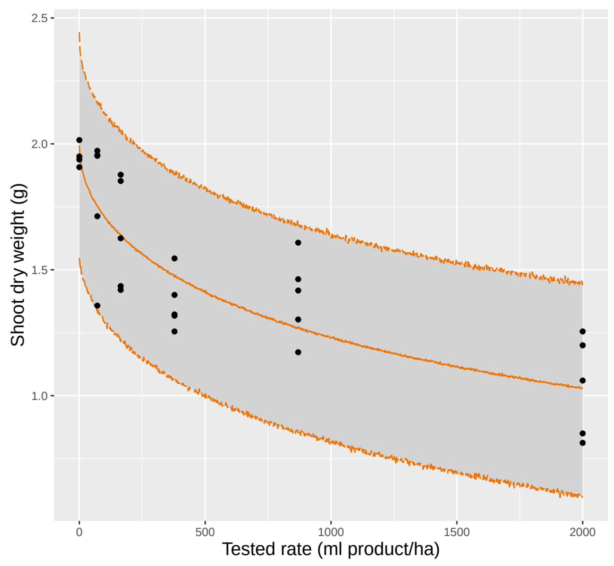

(a) Dose-response curve

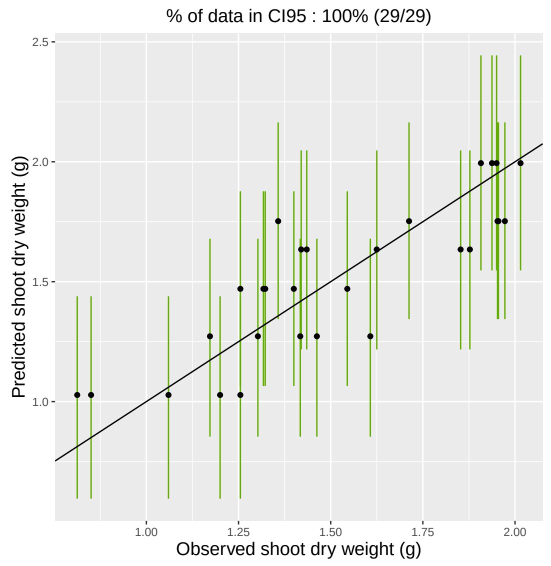

(b) Posterior predictive check (PPC)

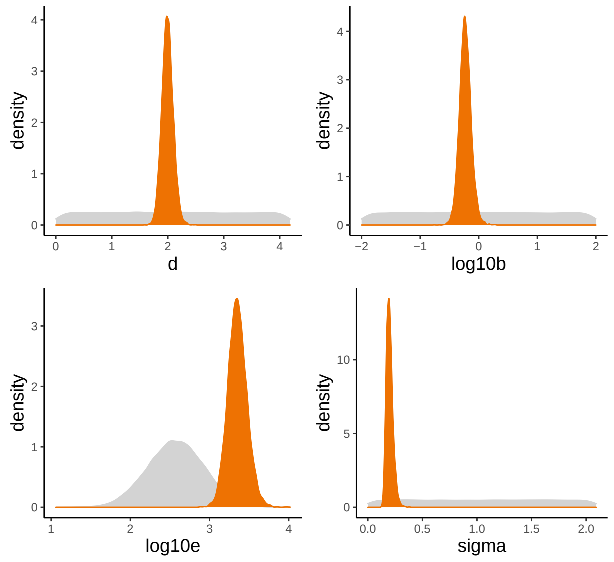

(c) Priors and posteriors

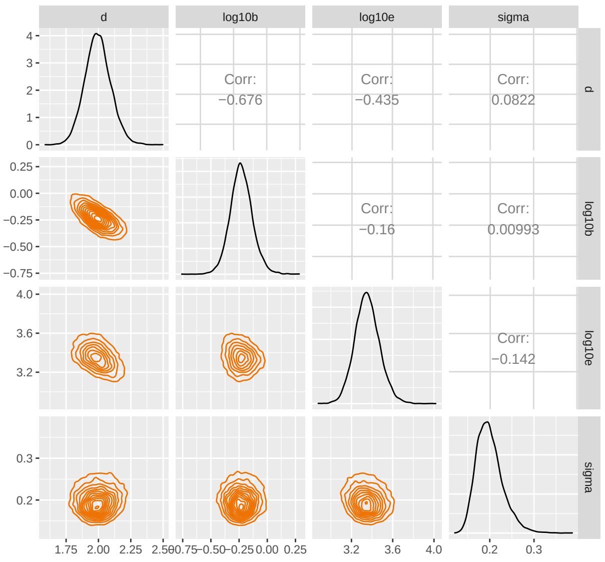

(d) Correlations between parameters

Figure 2: Dose-response curve (a), PPC (b), prior and posterior distributions (c) and correlations between parameters (d).

### Data set: BEAVA_VV_weight

Table 3: Summary of parameter estimates for BEAVA_VV_weight data set

| Parameter | median | Q2.5 | Q97.5 |
|---|---|---|---|
| b | 0.745 | 0.602 | 0.919 |
| d | 4.896 | 4.638 | 5.159 |
| e | 538.500 | 426.647 | 681.029 |
| sigma | 0.406 | 0.341 | 0.494 |

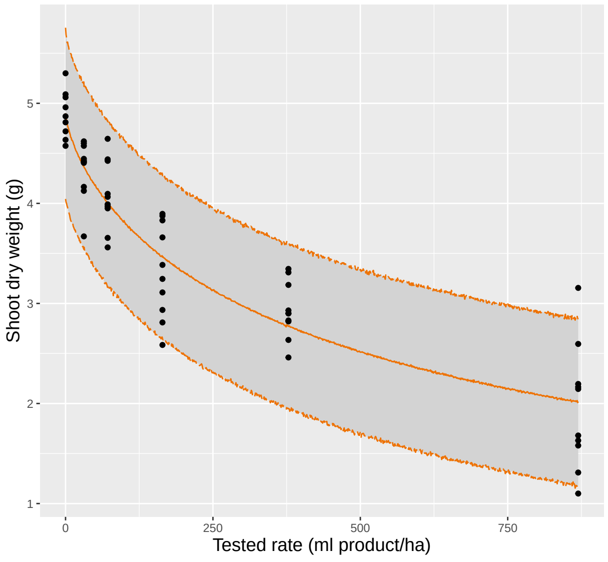

(a) Dose-response curve

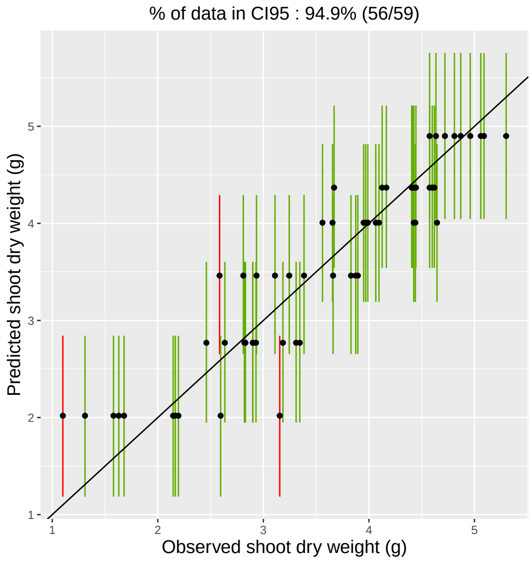

(b) Posterior predictive check (PPC)

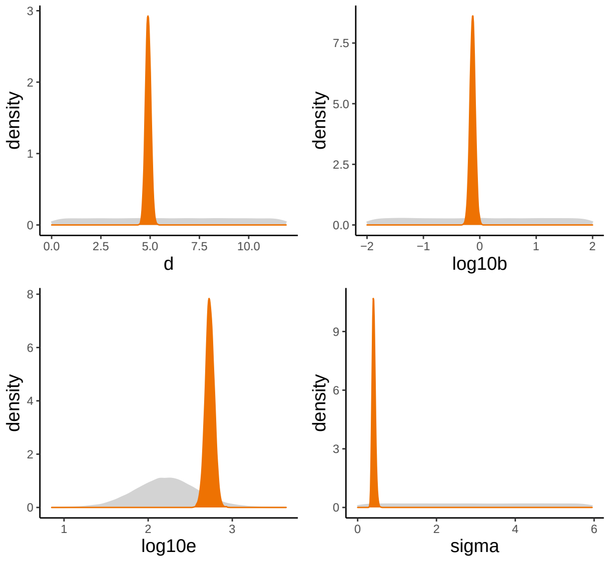

(c) Priors and posteriors

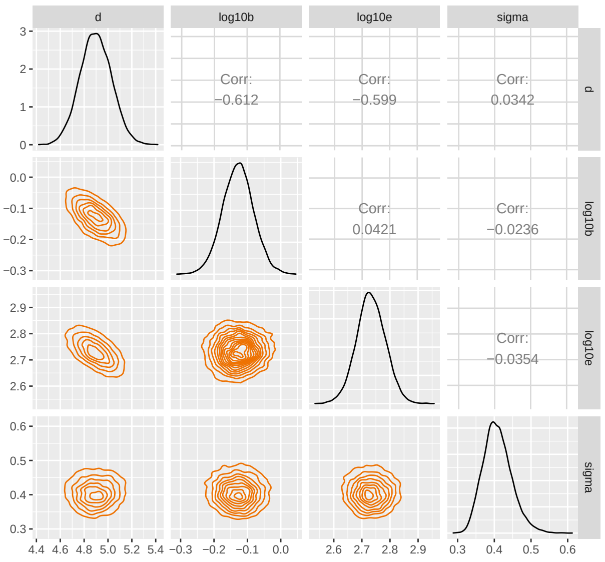

(d) Correlations between parameters

Figure 3: Dose-response curve (a), PPC (b), prior and posterior distributions (c) and correlations between parameters (d).

#### Data set: BRSNW_VV_weight

Table 4: Summary of parameter estimates for BRSNW_VV_weight data set

| Parameter | median | Q2.5 | Q97.5 |
|---|---|---|---|
| b | 1.061 | 0.881 | 1.279 |
| d | 8.883 | 8.329 | 9.466 |
| e | 166.389 | 134.762 | 204.493 |
| sigma | 0.897 | 0.752 | 1.097 |

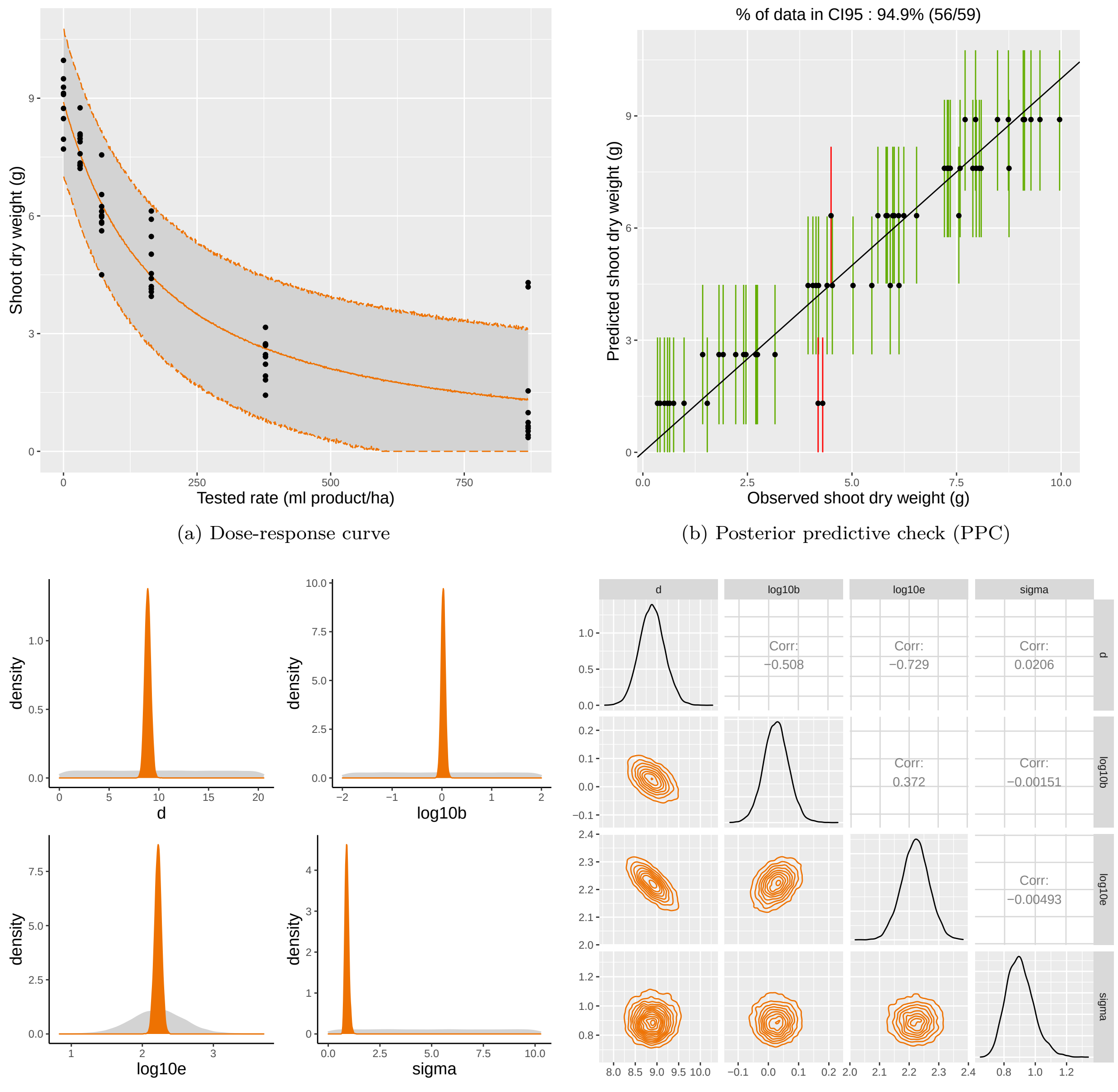

(a) Dose-response curve

(b) Posterior predictive check (PPC)

(c) Priors and posteriors

(d) Correlations between parameters

Figure 4: Dose-response curve (a), PPC (b), prior and posterior distributions (c) and correlations between parameters (d).

#### Data set: CUMSA_VV_weight

Table 5: Summary of parameter estimates for CUMSA_VV_weight data set

| Parameter | median | Q2.5 | Q97.5 |
|---|---|---|---|
| b | 0.485 | 0.347 | 0.644 |
| d | 16.043 | 14.784 | 17.311 |
| e | 281.418 | 177.977 | 460.066 |
| sigma | 1.977 | 1.654 | 2.419 |

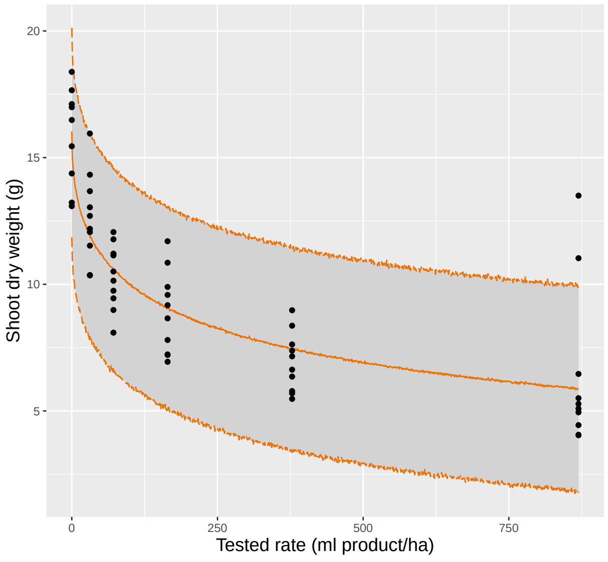

(a) Dose-response curve

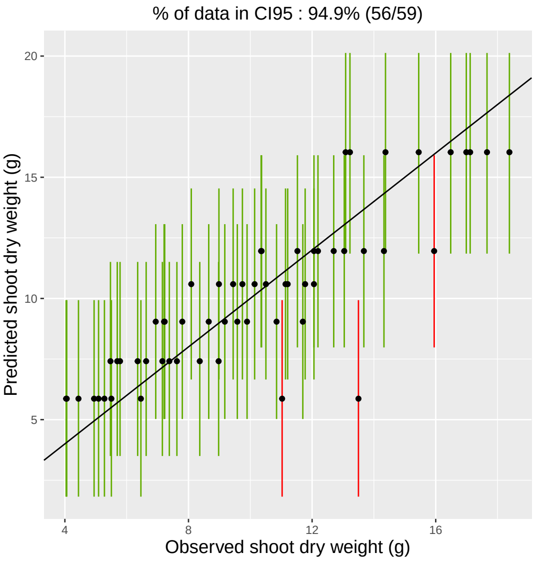

(b) Posterior predictive check (PPC)

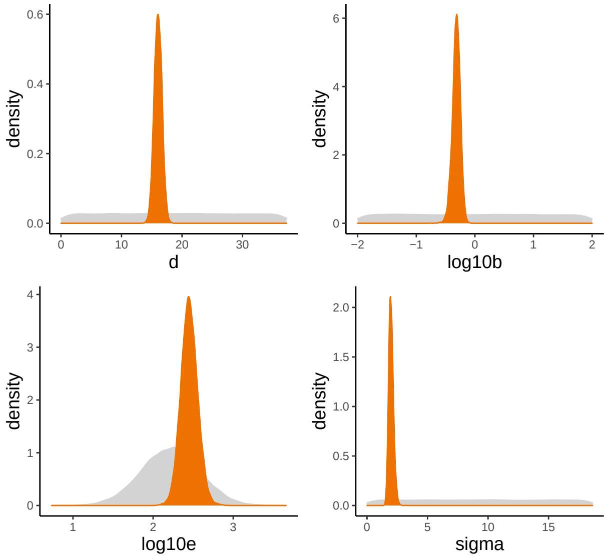

(c) Priors and posteriors

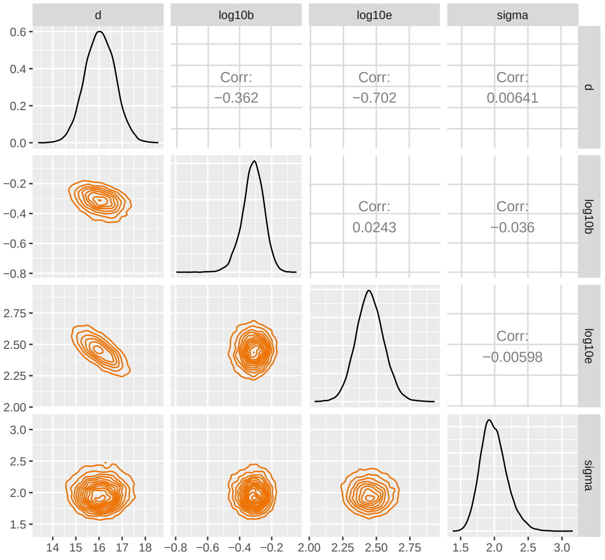

(d) Correlations between parameters

Figure 5: Dose-response curve (a), PPC (b), prior and posterior distributions (c) and correlations between parameters (d).

#### Data set: GLXMA_VV_weight

Table 6: Summary of parameter estimates for GLXMA_VV_weight data set

| Parameter | median | Q2.5 | Q97.5 |
|---|---|---|---|
| b | 0.676 | 0.557 | 0.818 |
| d | 5.563 | 5.278 | 5.863 |
| e | 443.868 | 346.204 | 577.433 |
| sigma | 0.461 | 0.385 | 0.563 |

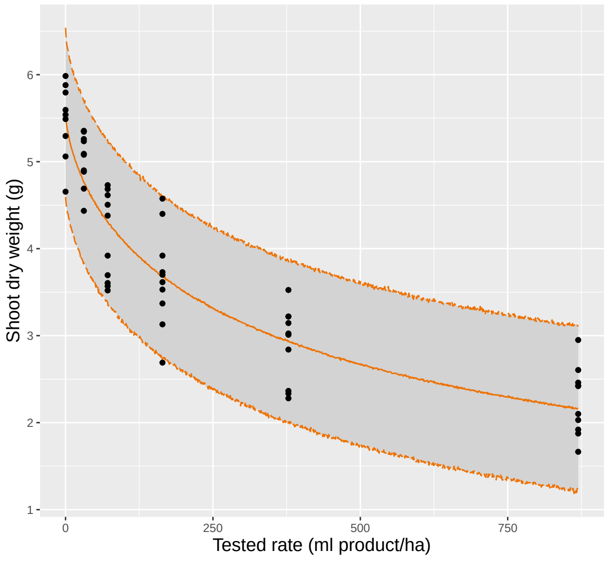

(a) Dose-response curve

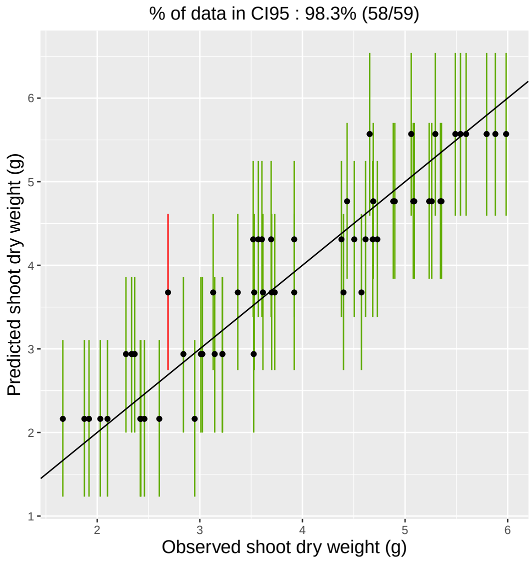

(b) Posterior predictive check (PPC)

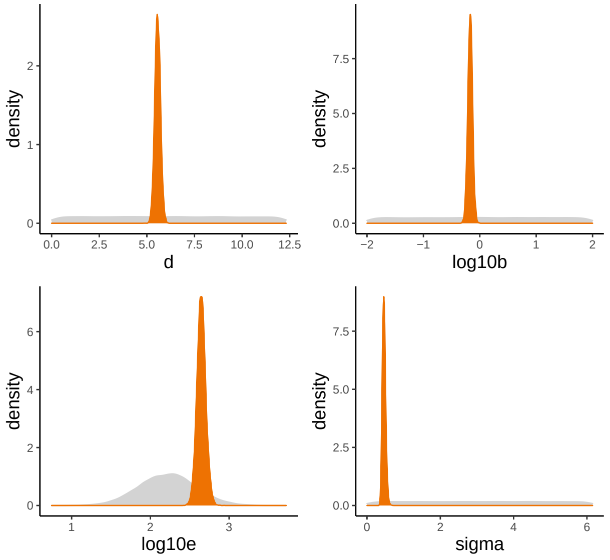

(c) Priors and posteriors

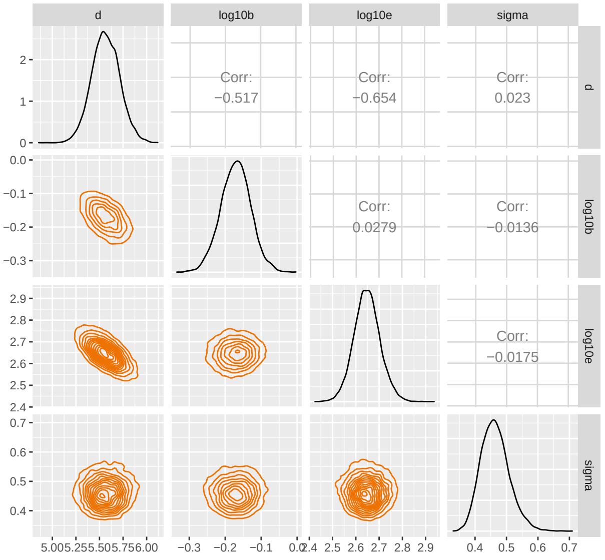

(d) Correlations between parameters

Figure 6: Dose-response curve (a), PPC (b), prior and posterior distributions (c) and correlations between parameters (d).

#### Data set: HELAN_VV_weight

Table 7: Summary of parameter estimates for HELAN_VV_weight data set

| Parameter | median | Q2.5 | Q97.5 |
|---|---|---|---|
| b | 0.629 | 0.498 | 0.785 |
| d | 3.464 | 3.284 | 3.655 |
| e | 685.788 | 510.862 | 955.936 |
| sigma | 0.310 | 0.260 | 0.380 |

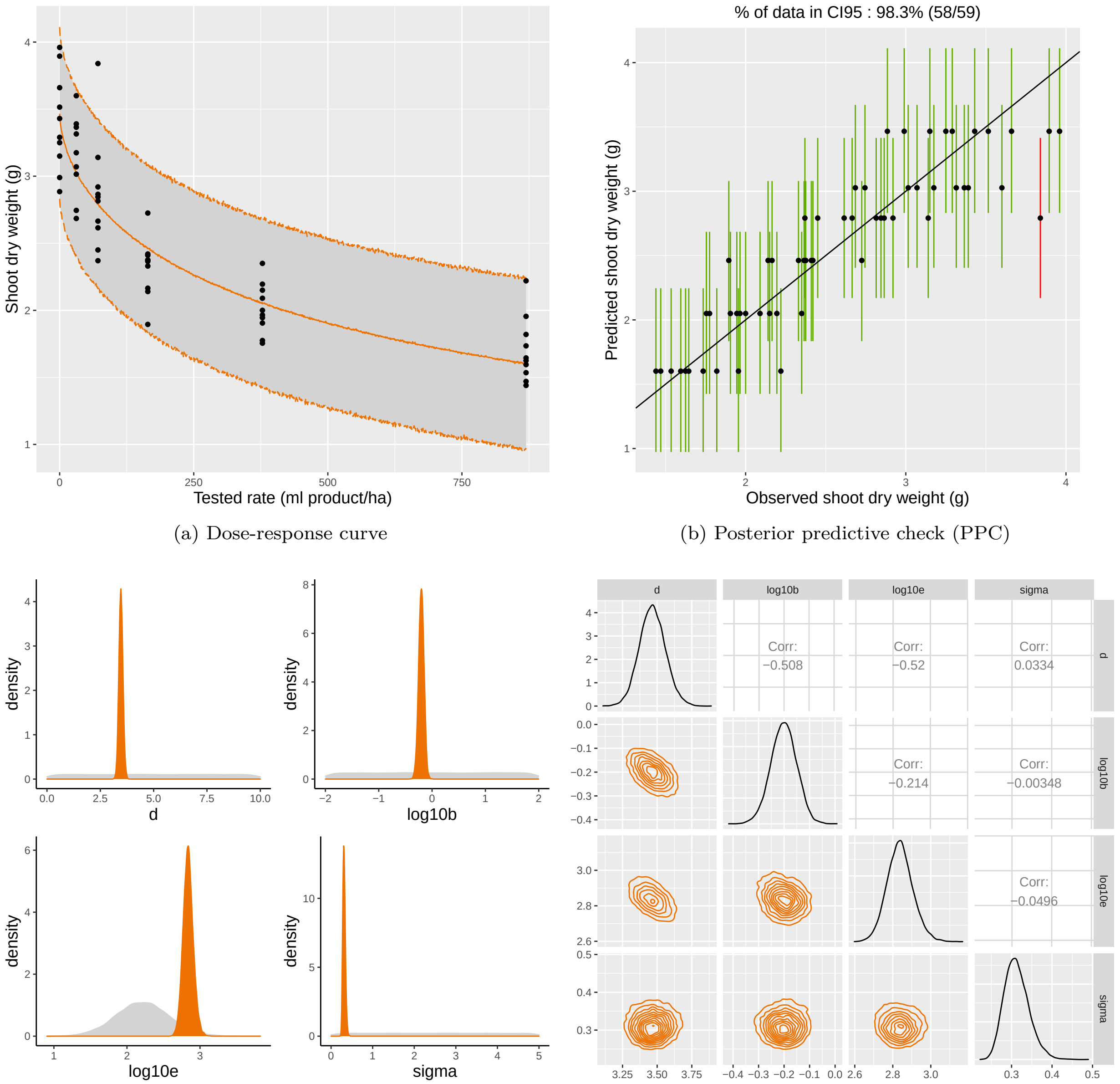

(a) Dose-response curve

(b) Posterior predictive check (PPC)

(c) Priors and posteriors

(d) Correlations between parameters

Figure 7: Dose-response curve (a), PPC (b), prior and posterior distributions (c) and correlations between parameters (d).

#### Data set: LYPES_VV_weight

Table 8: Summary of parameter estimates for LYPES_VV_weight data set

| Parameter | median | Q2.5 | Q97.5 |
|---|---:|---:|---:|
| b | 0.628 | 0.492 | 0.809 |
| d | 12.221 | 11.629 | 12.825 |
| e | 1316.789 | 958.545 | 1924.952 |
| sigma | 0.985 | 0.827 | 1.194 |

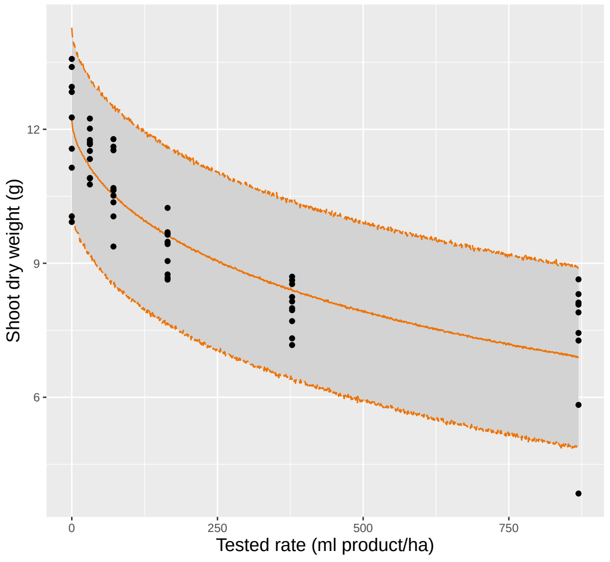

(a) Dose-response curve

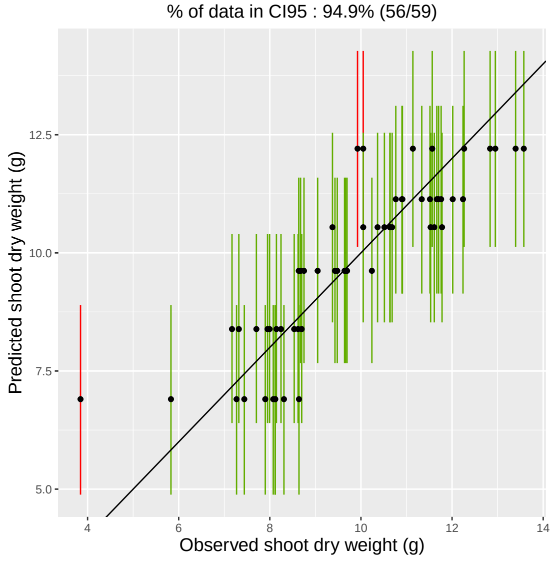

(b) Posterior predictive check (PPC)

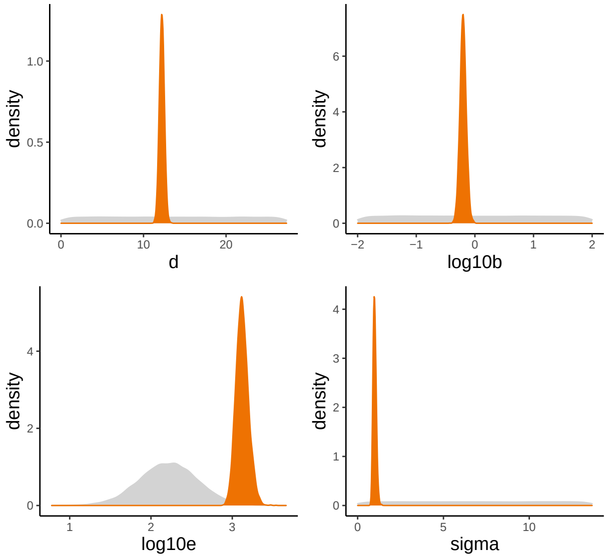

(c) Priors and posteriors

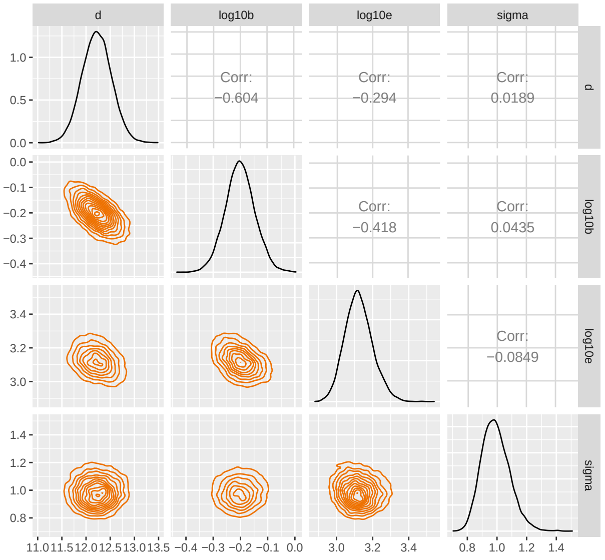

(d) Correlations between parameters

Figure 8: Dose-response curve (a), PPC (b), prior and posterior distributions (c) and correlations between parameters (d).

#### Data set: TRZAW_VV_weight

Table 9: Summary of parameter estimates for TRZAW_VV_weight data set

| Parameter | median | Q2.5 | Q97.5 |
|---|---|---|---|
| b | 1.565 | 0.804 | 24.256 |
| d | 1.143 | 1.079 | 1.218 |
| e | 3874.808 | 2129.187 | 7560.358 |
| sigma | 0.110 | 0.085 | 0.151 |

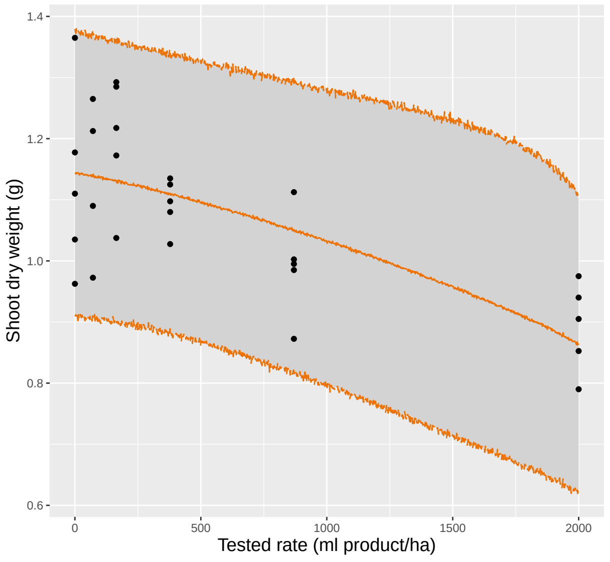

(a) Dose-response curve

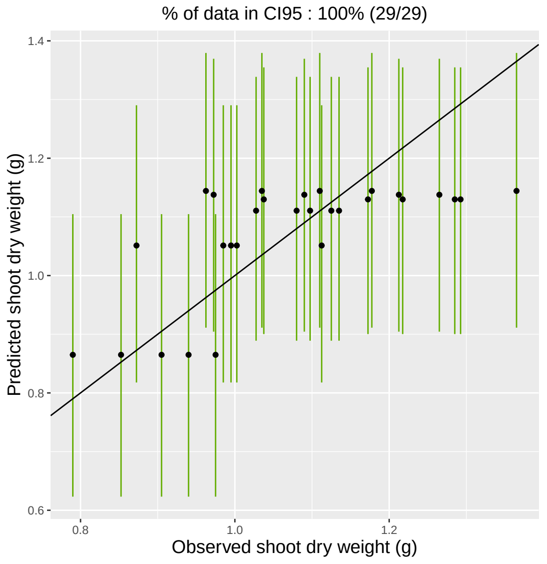

(b) Posterior predictive check (PPC)

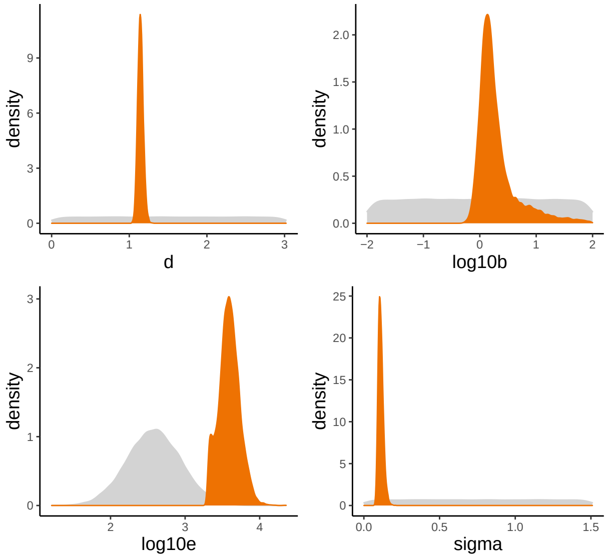

(c) Priors and posteriors

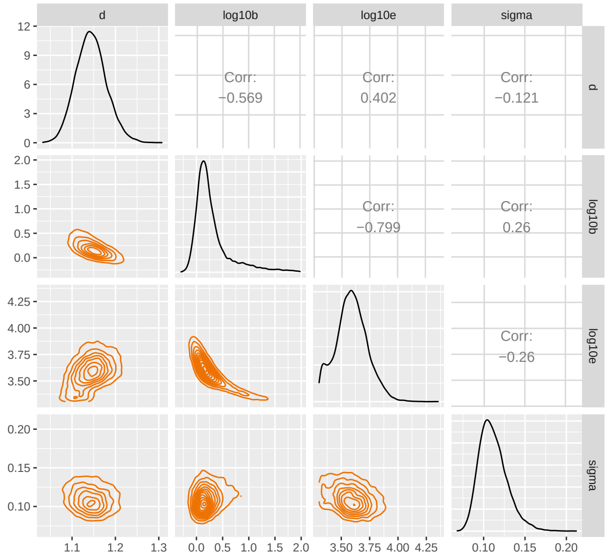

(d) Correlations between parameters

Figure 9: Dose-response curve (a), PPC (b), prior and posterior distributions (c) and correlations between parameters (d).

#### Data set: ZEAMA_VV_weight

Table 10: Summary of parameter estimates for ZEAMA_VV_weight data set

| Parameter | median | Q2.5 | Q97.5 |
|---|---|---|---|
| b | 0.732 | 0.492 | 1.208 |
| d | 8.628 | 8.195 | 9.109 |
| e | 6788.740 | 4142.477 | 13203.057 |
| sigma | 0.774 | 0.650 | 0.953 |

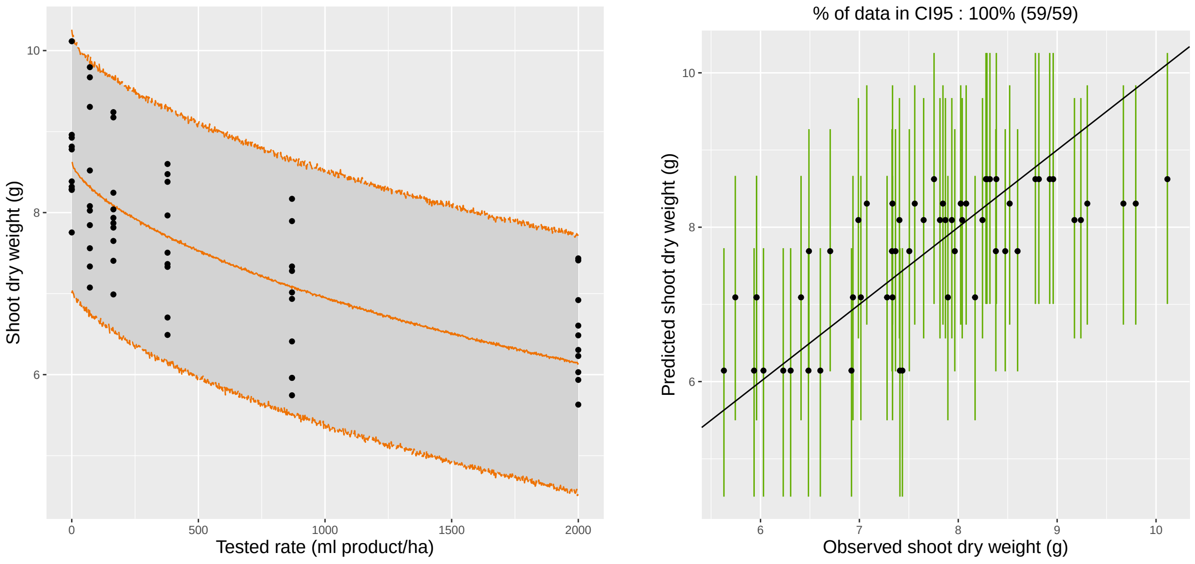

(a) Dose-response curve

(b) Posterior predictive check (PPC)

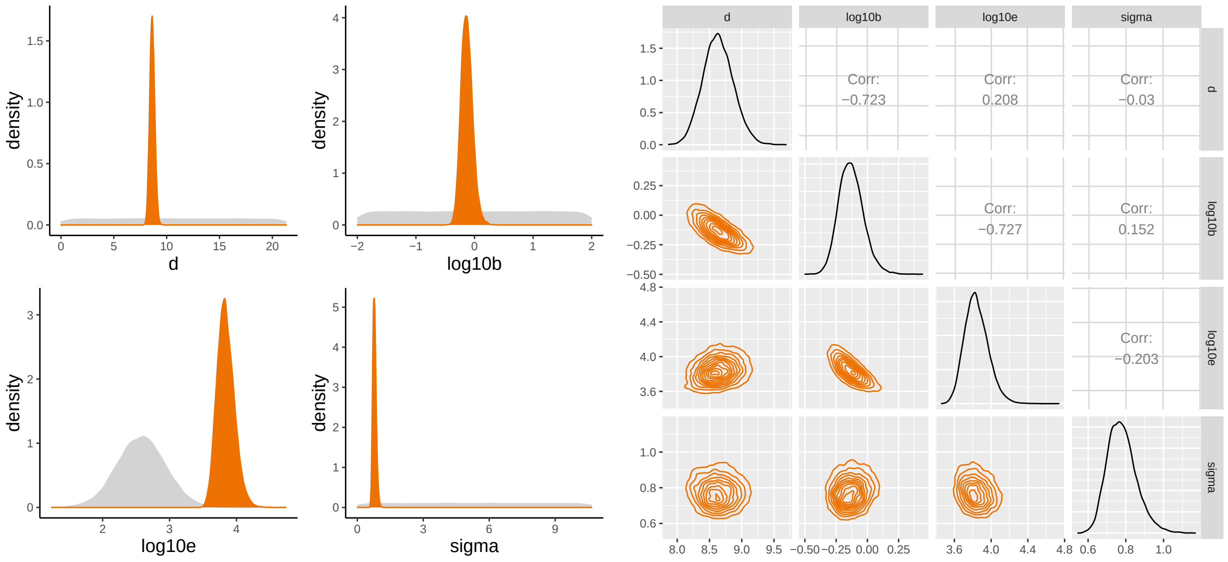

(c) Priors and posteriors

(d) Correlations between parameters

Figure 10: Dose-response curve (a), PPC (b), prior and posterior distributions (c) and correlations between parameters (d).
