## Supplementary material for "How to account for the uncertainty from standard toxicity tests in species sensitivity distributions: an example in non-target plants": Archive with reports on studies: SSD_analyses.pdf

### Species sensitivity distribution (SSD) analyses by fitting a log-normal distribution

#### Study 1

25 June 2020

Contact:

---

The document provides results on species sensitivity distribution (SSD) analyses for study 1. There are five endpoints for study 1: emergence, survival, shoot dry weight of the seedling emergence (SE) test, and survival and shoot dry weight of vegetative vigour (VV) test.

We used SSD to describe the toxicity variation of herbicides among a set of non-target terrestrial plants (NTTP). Based on this SSD, the hazardous rate at which five percent of the species are affected ($HR_5$) can be estimated. This document aims to compare SSD and $HR_5$ based on 50% effective rates ($ER_{50}$) generated in three different ways. We considered:

1. Point estimates (medians of posterior distributions of $ER_{50}$) as inputs for SSD analyses;
2. Interval-censored $ER_{50}$ by 95% credible interval (CI95);
3. censored $ER_{50}$ according to criterion 1 with a decision threshold T = 0.5

For each endpoint, a total of three SSD are produced:

1. SSD based on medians of posterior distributions of $ER_{50}$ (figure title: median);
2. SSD based on CI95 of $ER_{50}$ (figure title: CI95);
3. SSD based on censored $ER_{50}$ according to criterion 1 with T = 0.5 (figure title: C1T0.5).

**Meaning of symbols used in Tables 1-5:**

- nb_$ER_{50}$ (RC): number of $ER_{50}$ (number of right-censored $ER_{50}$);
- nb_dist$ER_{50}$ (RC): number of distinct $ER_{50}$ (number of distinct right-censored $ER_{50}$);
- $HR_5$_est: $HR_5$ mean estimate;
- $HR_5$_median: median of $HR_5$;
- $HR_5$_LCoI95: lower bound of the 95% confidence interval (CoI95) of $HR_5$;
- $HR_5$_UCoI95: upper bound of the CoI95 of $HR_5$.

**Meaning of legends, lines or curves in Figures 1-5:**

- Nb. $ER_{50}$ (RC): number of $ER_{50}$ (number of right-censored $ER_{50}$);
- Nb. distinct $ER_{50}$ (RC): number of distinct $ER_{50}$ (number of distinct right-censored $ER_{50}$);
- $HR_5$ [CoI95]: $HR_5$ mean estimate [CoI95 of $HR_5$];
- Blue points: fractions of affected species (EPPO code) ordered by the median of $ER_{50}$;
- Solid blue horizontal segment: interval-censored $ER_{50}$ by the CI95 of the $ER_{50}$;
- Dotted blue horizontal segment: right-censored $ER_{50}$;
- Solid red curve: fitted SSD curve;
- Dotted red curves: CoI95 of the fitted SSD curve.

### $HR_5$ results and SSD curves

#### $HR_5$ and SSD for the emergence endpiont of the SE test of study 1

Table 1: Number of (distinct) $ER_{50}$ (right-censored) and $HR_5$ estimate for the emergence endpoint of the SE test of study 1 (tested rate unit: ml product/ha)

| criterion | nb_$ER_{50}$ (RC) | nb_dist$ER_{50}$ (RC) | $HR_5$_est | $HR_5$_median | $HR_5$_LCoI95 | $HR_5$_UCoI95 | CV |
|---|---|---|---|---|---|---|---|
| median | 10 (0) | 10 (0) | 832.125 | 898.580 | 694.639 | 1164.750 | 0.142 |
| CI95 | 10 (0) | 10 (0) | 1581.358 | 1659.924 | 1245.310 | 2211.102 | 0.164 |
| C1T0.5 | 10 (9) | 5 (4) | 2260.459 | 2260.459 | 2260.459 | 2260.459 | 0.000 |

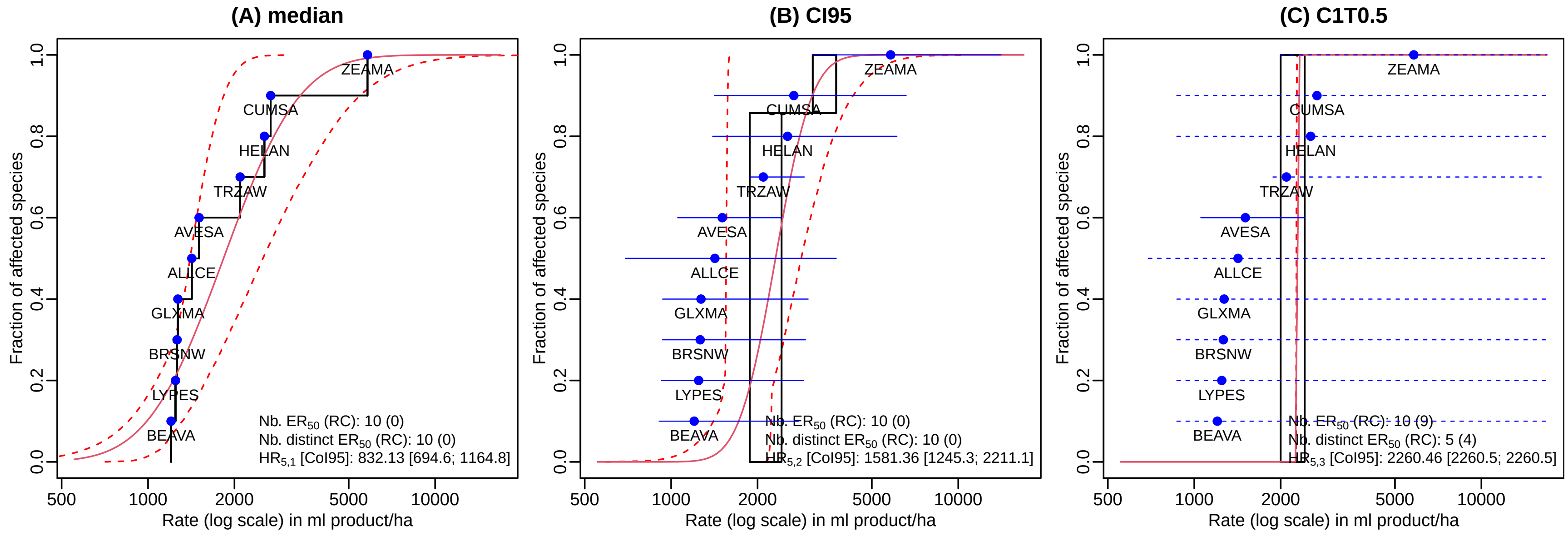


Figure 1: Example of three SSD based on respectively $ER_{50}$ medians (A), CI95 of $ER_{50}$ (B) and censored $ER_{50}$ according to C1T0.5 (C) for the emergnece endpoint of the SE test of study 1.

#### $HR_5$ and SSD for the survival endpiont of the SE test of study 1

Table 2: Number of (distinct) $ER_{50}$ (right-censored) and $HR_5$ estimate for the survival endpoint of the SE test of study 1 (tested rate unit: ml product/ha)

| criterion | nb_$ER_{50}$ (RC) | nb_dist$ER_{50}$ (RC) | $HR_5$_est | $HR_5$_median | $HR_5$_LCoI95 | $HR_5$_UCoI95 | CV |
|---|---|---|---|---|---|---|---|
| median | 10 (0) | 10 (0) | 177.685 | 188.283 | 98.078 | 460.631 | 0.451 |
| CI95 | 10 (0) | 10 (0) | 172.208 | 185.712 | 93.005 | 465.532 | 0.512 |
| C1T0.5 | 10 (4) | 8 (2) | 146.369 | 158.137 | 78.272 | 412.595 | 0.620 |

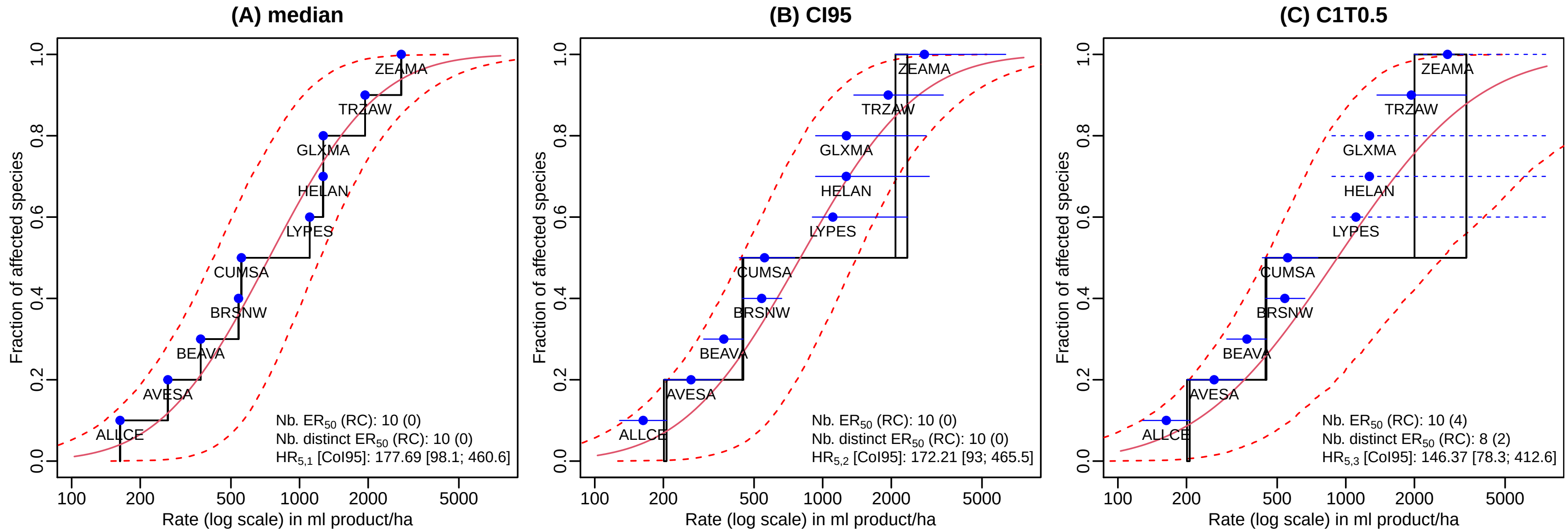


Figure 2: Example of three SSD based on respectively $ER_{50}$ medians (A), CI95 of $ER_{50}$ (B) and censored $ER_{50}$ according to C1T0.5 (C) for the survival endpoint of the SE test of study 1.

#### $HR_5$ and SSD for the shoot dry weight endpiont of the SE test of study 1

Table 3: Number of (distinct) $ER_{50}$ (right-censored) and $HR_5$ estimate for the shoot dry weight endpoint of the SE test of study 1 (tested rate unit: ml product/ha)

| criterion | nb_$ER_{50}$ (RC) | nb_dist$ER_{50}$ (RC) | $HR_5$_est | $HR_5$_median | $HR_5$_LCoI95 | $HR_5$_UCoI95 | CV |
|---|---|---|---|---|---|---|---|
| median | 10 (0) | 10 (0) | 126.388 | 133.673 | 84.871 | 257.074 | 0.311 |
| CI95 | 10 (0) | 10 (0) | 125.604 | 134.848 | 84.195 | 259.481 | 0.331 |
| C1T0.5 | 10 (2) | 9 (1) | 120.480 | 128.796 | 80.598 | 253.838 | 0.322 |

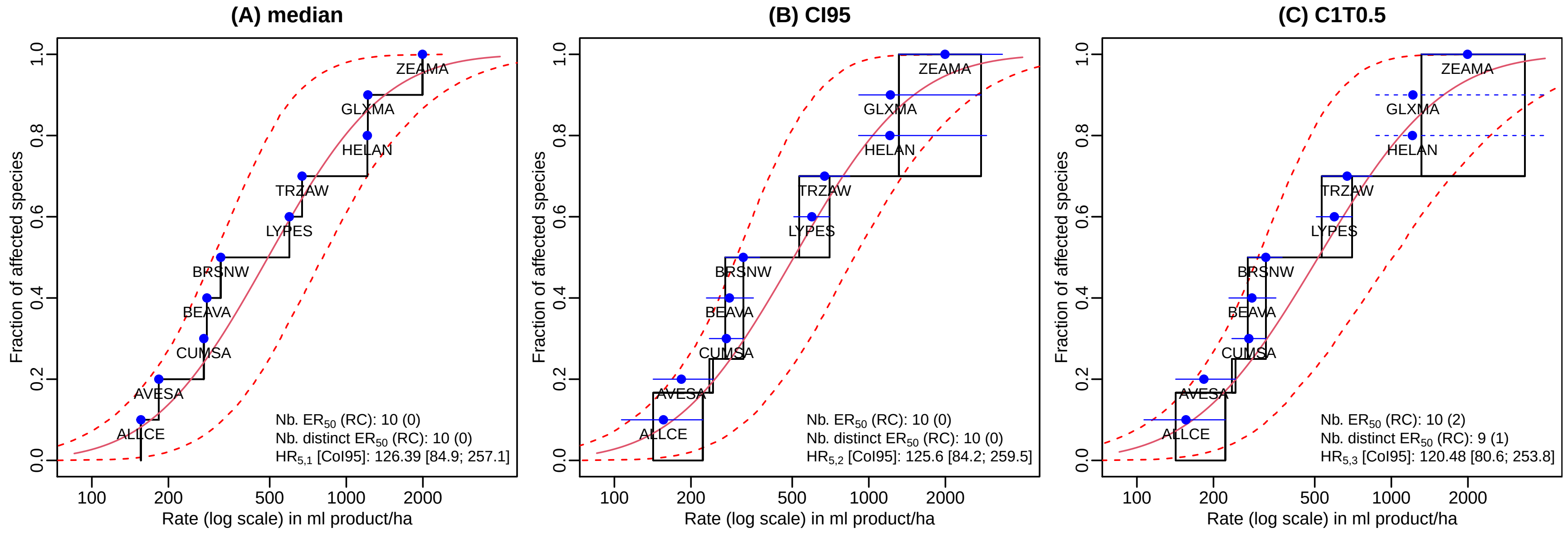


Figure 3: Example of three SSD based on respectively $ER_{50}$ medians (A), CI95 of $ER_{50}$ (B) and censored $ER_{50}$ according to C1T0.5 (C) for the shoot dry weight endpoint of the SE test of study 1.

#### $HR_5$ and SSD for the survival endpiont of the VV test of study 1

Table 4: Number of (distinct) $ER_{50}$ (right-censored) and $HR_5$ estimate for the survival endpoint of the VV test of study 1 (tested rate unit: ml product/ha)

| criterion | nb_$ER_{50}$ (RC) | nb_dist$ER_{50}$ (RC) | $HR_5$_est | $HR_5$_median | $HR_5$_LCoI95 | $HR_5$_UCoI95 | CV |
|---|---|---|---|---|---|---|---|
| median | 10 (0) | 10 (0) | 787.045 | 812.425 | 703.078 | 1068.519 | 0.101 |
| CI95 | 10 (0) | 10 (0) | 1183.658 | 1161.124 | 922.577 | 2458.266 | 0.376 |
| C1T0.5 | 10 (10) | 3 (3) | 8696.715 | 8696.715 | 8696.715 | 8696.715 | 0.000 |

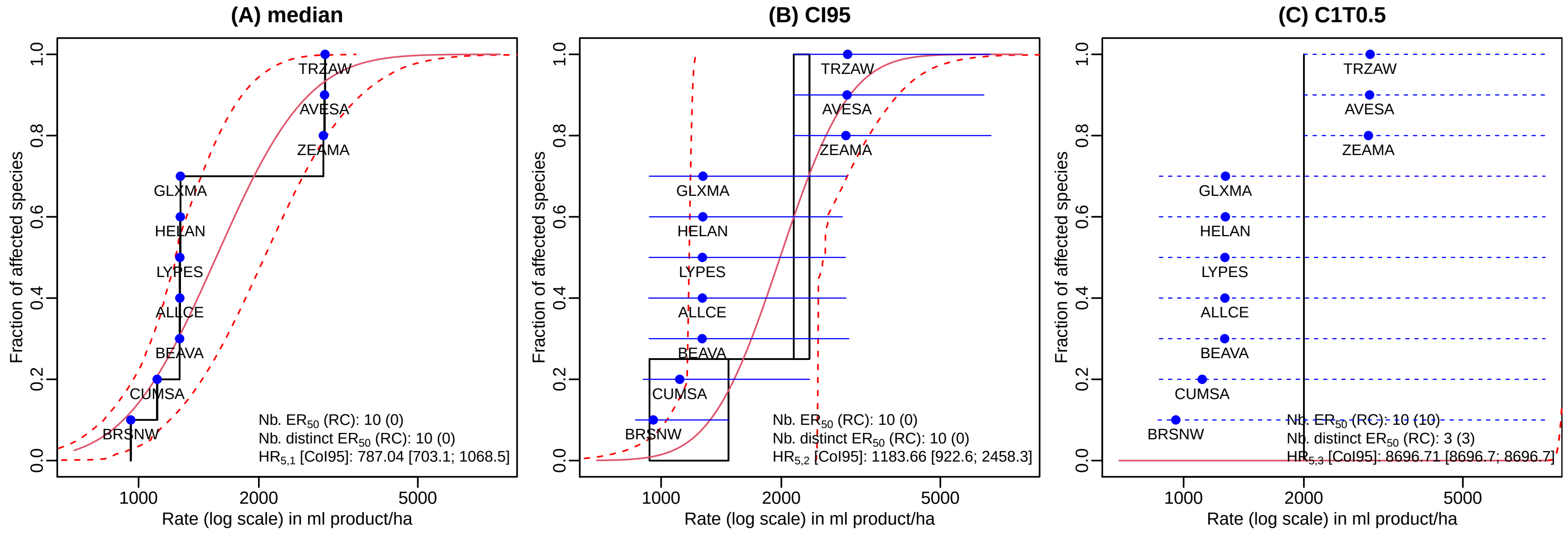


Figure 4: Example of three SSD based on respectively $ER_{50}$ medians (A), CI95 of $ER_{50}$ (B) and censored $ER_{50}$ according to C1T0.5 (C) for the survival endpoint of the VV test of study 1.

#### $HR_5$ and SSD for the shoot dry weight endpiont of the VV test of study 1

Table 5: Number of (distinct) $ER_{50}$ (right-censored) and $HR_5$ estimate for the shoot dry weight endpoint of the VV test of study 1 (tested rate unit: ml product/ha)

| criterion | nb_$ER_{50}$ (RC) | nb_dist$ER_{50}$ (RC) | $HR_5$_est | $HR_5$_median | $HR_5$_LCoI95 | $HR_5$_UCoI95 | CV |
|---|---|---|---|---|---|---|---|
| median | 10 (0) | 10 (0) | 146.606 | 162.827 | 79.961 | 375.887 | 0.436 |
| CI95 | 10 (0) | 10 (0) | 142.501 | 160.413 | 76.488 | 415.390 | 0.512 |
| C1T0.5 | 10 (4) | 9 (3) | 130.853 | 144.164 | 70.932 | 349.320 | 0.533 |

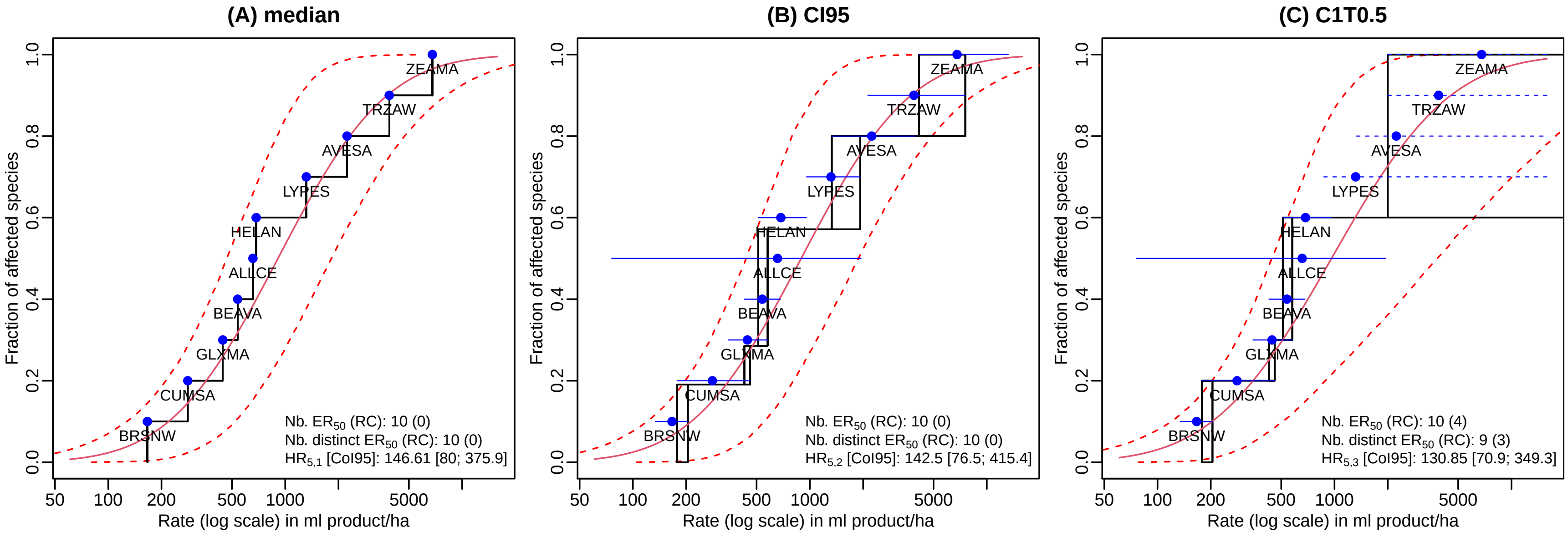


Figure 5: Example of three SSD based on respectively $ER_{50}$ medians (A), CI95 of $ER_{50}$ (B) and censored $ER_{50}$ according to C1T0.5 (C) for the shoot dry weight endpoint of the VV test of study 1.
