## Supplementary material for "How to account for the uncertainty from standard toxicity tests in species sensitivity distributions: an example in non-target plants": Archive with reports on studies: report_SE_emergence.pdf

### Dose-response analyses

Study 2

Seedling Emergence test - emergence endpoint

25 June 2020

Contact:

---

This is a report which provides results on all performed dose-response analyses for the emergence endpoint of the Seedling Emergence test for study 2.

---

### Data set: ALLCE_SE_emergence

Table 1: Summary of parameter estimates for ALLCE_SE_emergence data set

| Parameter | median | Q2.5 | Q97.5 |
|---|---|---|---|
| b | 18.855 | 2.070 | 92.244 |
| d | 0.883 | 0.835 | 0.924 |
| e | 41.298 | 30.148 | 105.057 |

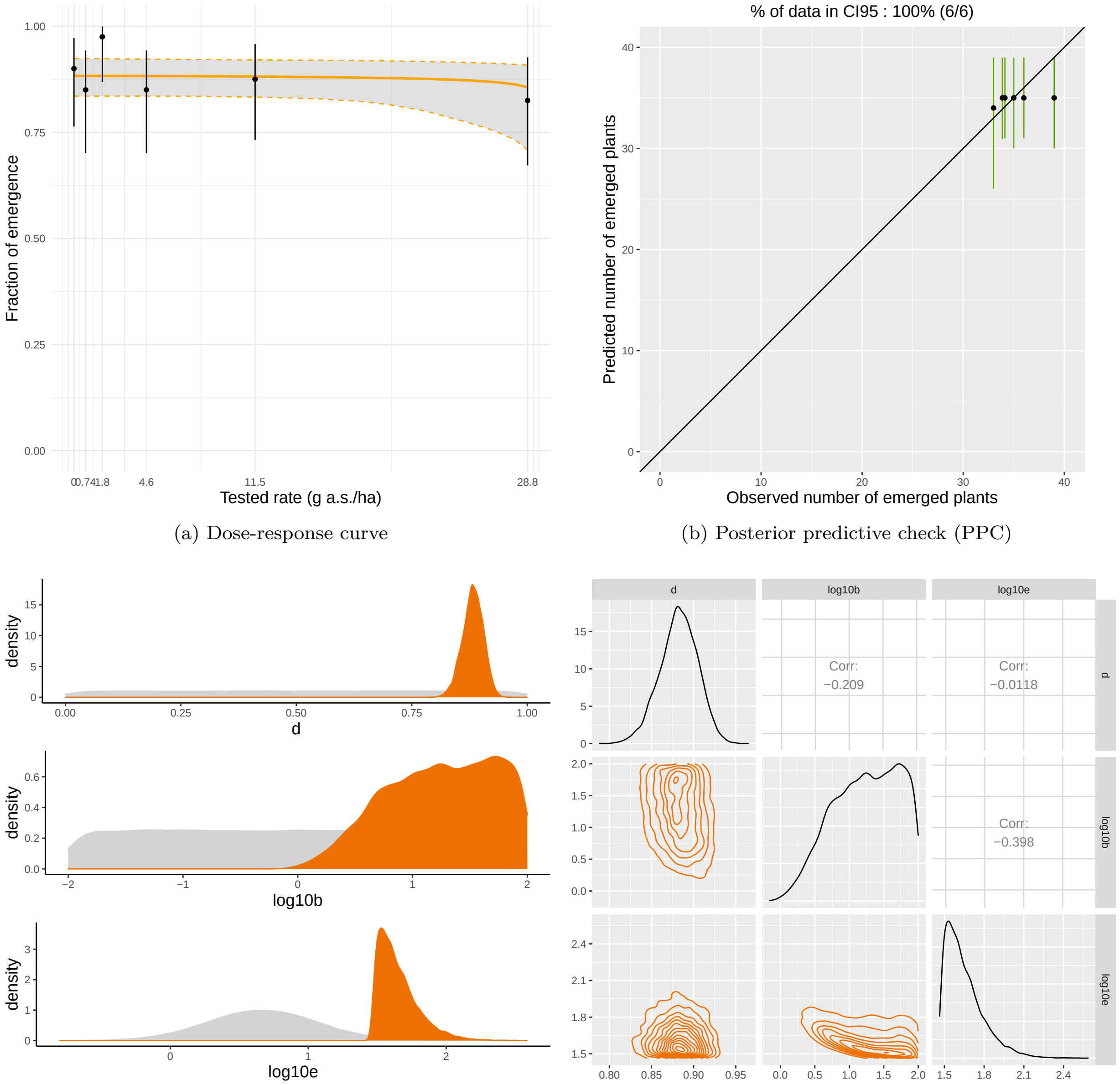

(a) Dose-response curve

(b) Posterior predictive check (PPC)

(c) Priors and posteriors

(d) Correlations between parameters

Figure 1: Dose-response curve (a), PPC (b), prior and posterior distributions (c) and correlations between parameters (d).

#### Data set: AVESA_SE_emergence

Table 2: Summary of parameter estimates for AVESA_SE_emergence data set

| Parameter | median | Q2.5 | Q97.5 |
|---|---|---|---|
| b | 32.577 | 4.714 | 94.702 |
| d | 0.969 | 0.942 | 0.986 |
| e | 268.672 | 193.990 | 661.026 |

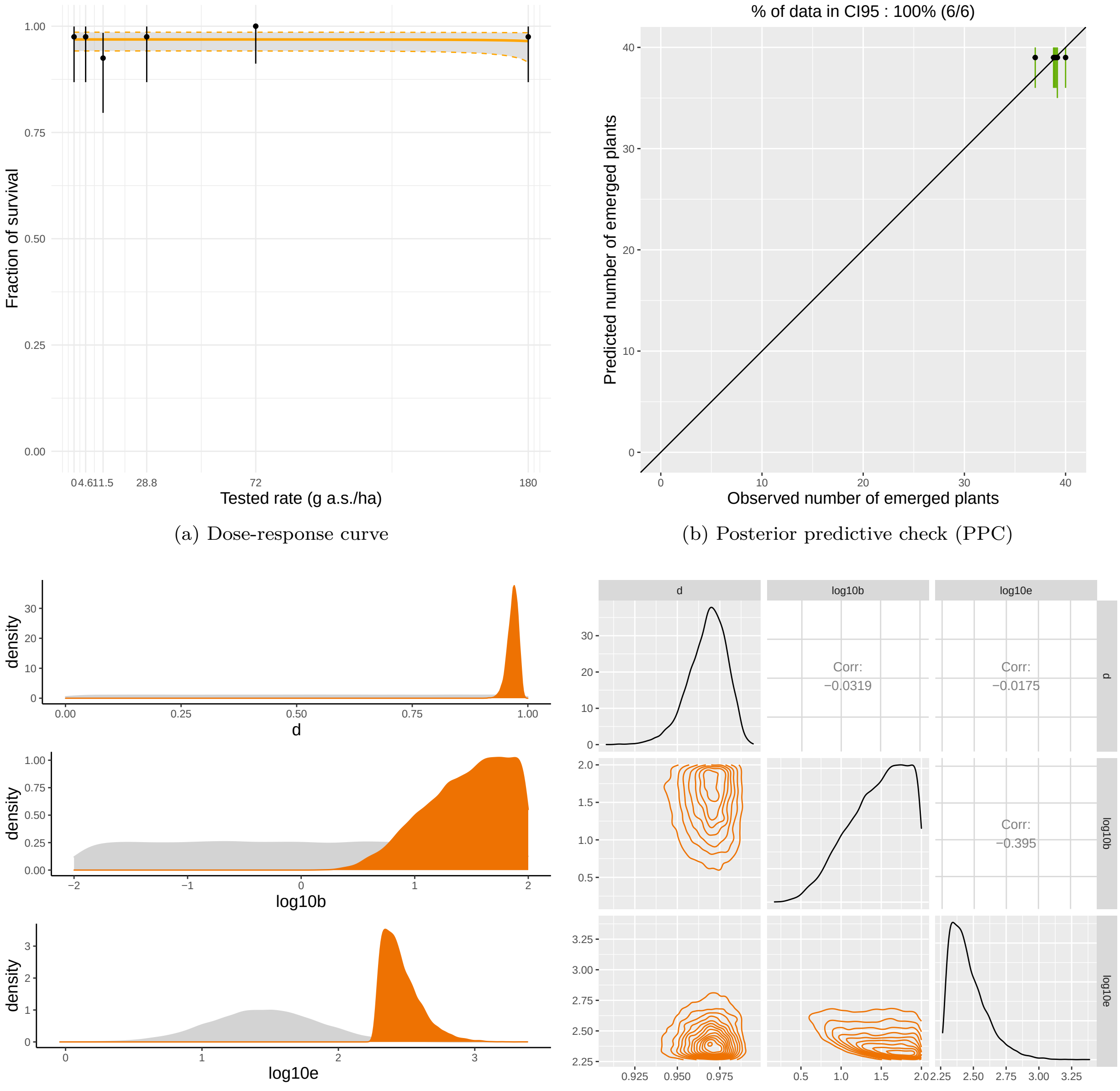

(a) Dose-response curve

(b) Posterior predictive check (PPC)

(c) Priors and posteriors

(d) Correlations between parameters

Figure 2: Dose-response curve (a), PPC (b), prior and posterior distributions (c) and correlations between parameters (d).

#### Data set: BEAVA_SE_emergence

Table 3: Summary of parameter estimates for BEAVA_SE_emergence data set

| Parameter | median | Q2.5 | Q97.5 |
|---|---|---|---|
| b | 29.112 | 3.458 | 94.438 |
| d | 0.861 | 0.816 | 0.902 |
| e | 42.933 | 30.786 | 107.747 |

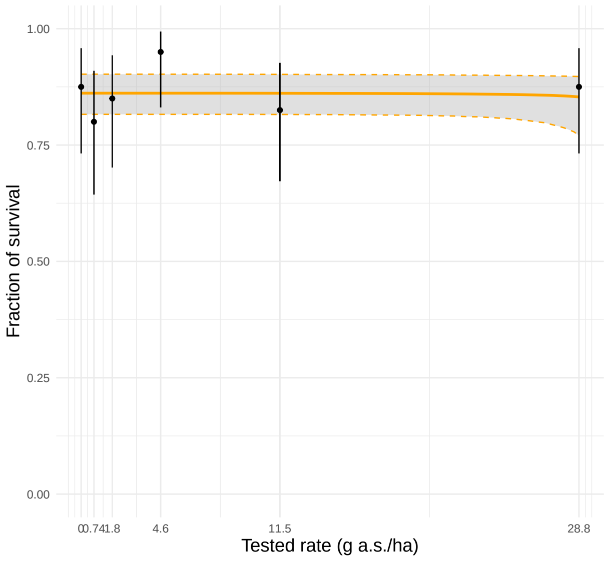

(a) Dose-response curve

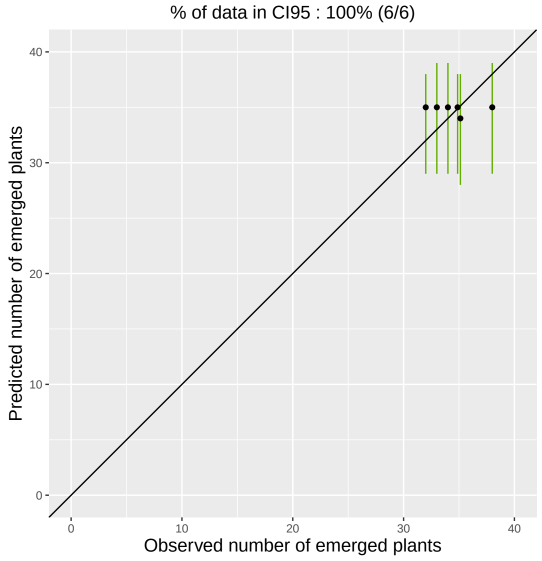

(b) Posterior predictive check (PPC)

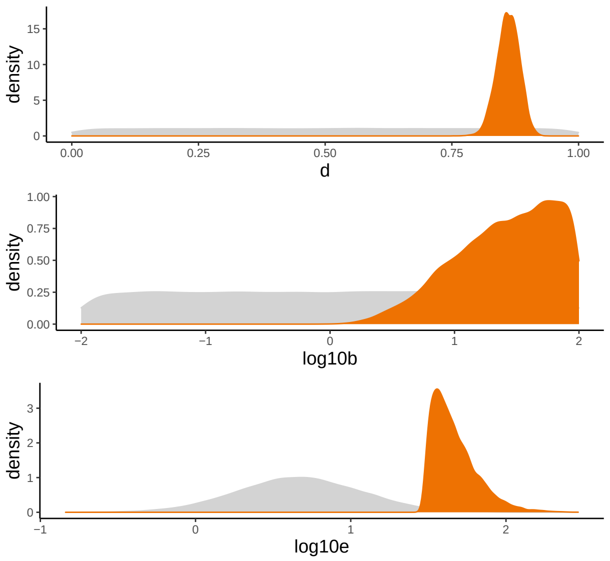

(c) Priors and posteriors

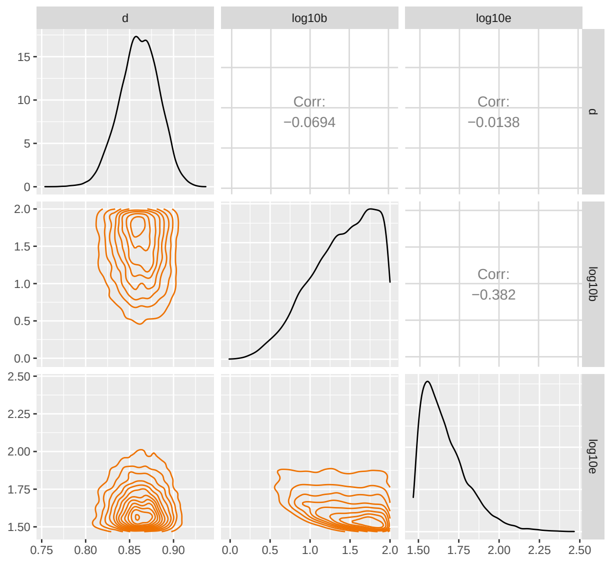

(d) Correlations between parameters

Figure 3: Dose-response curve (a), PPC (b), prior and posterior distributions (c) and correlations between parameters (d).

#### Data set: BRSNW_SE_emergence

Table 4: Summary of parameter estimates for BRSNW_SE_emergence data set

| Parameter | median | Q2.5 | Q97.5 |
|---|---|---|---|
| b | 29.073 | 3.879 | 94.291 |
| d | 0.928 | 0.891 | 0.956 |
| e | 268.648 | 192.273 | 673.321 |

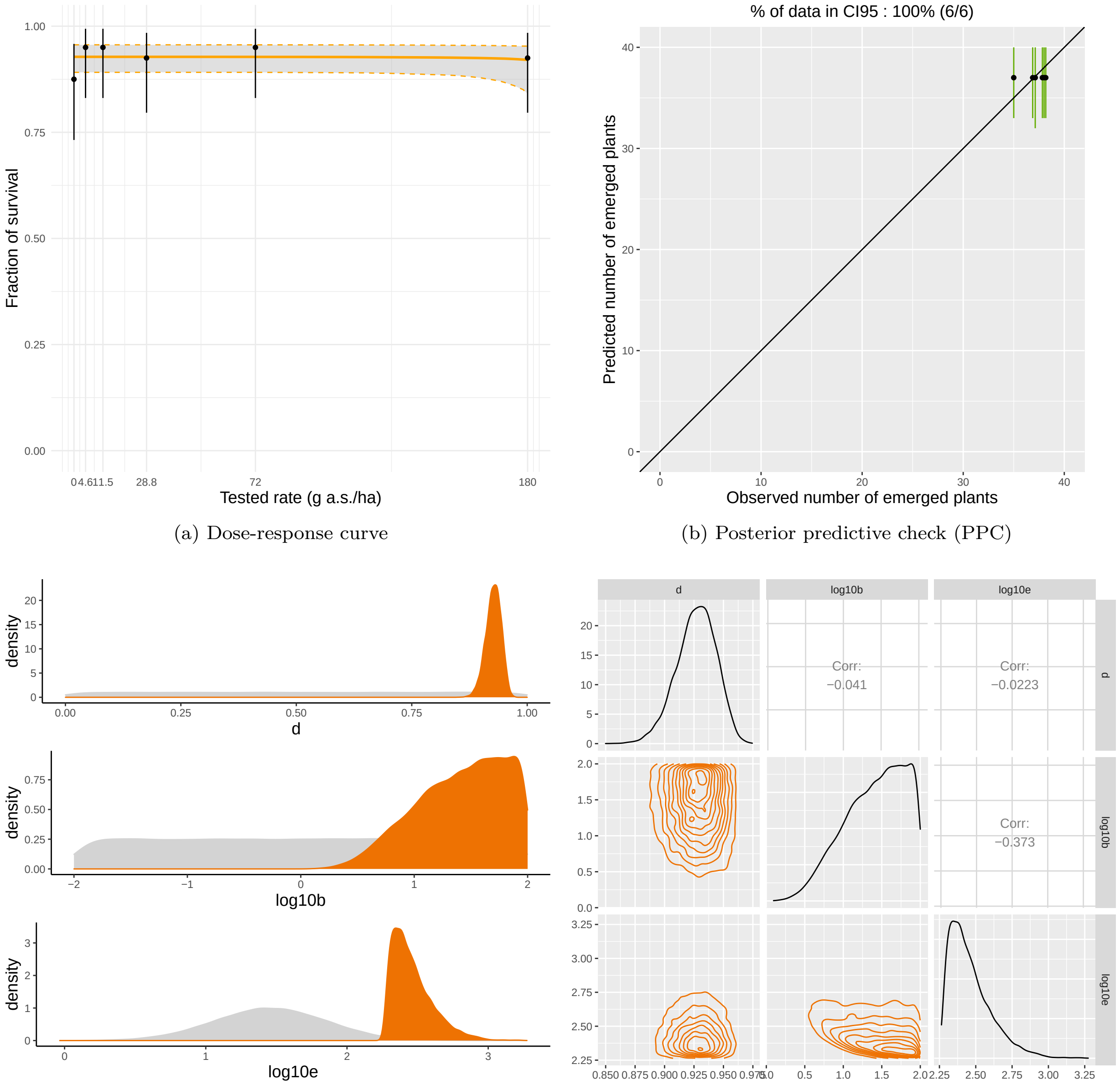

(a) Dose-response curve

(b) Posterior predictive check (PPC)

(c) Priors and posteriors

(d) Correlations between parameters

Figure 4: Dose-response curve (a), PPC (b), prior and posterior distributions (c) and correlations between parameters (d).

#### Data set: CUMSA_SE_emergence

Table 5: Summary of parameter estimates for CUMSA_SE_emergence data set

| Parameter | median | Q2.5 | Q97.5 |
|---|---|---|---|
| b | 26.365 | 3.297 | 93.365 |
| d | 0.920 | 0.882 | 0.951 |
| e | 262.856 | 190.861 | 656.414 |

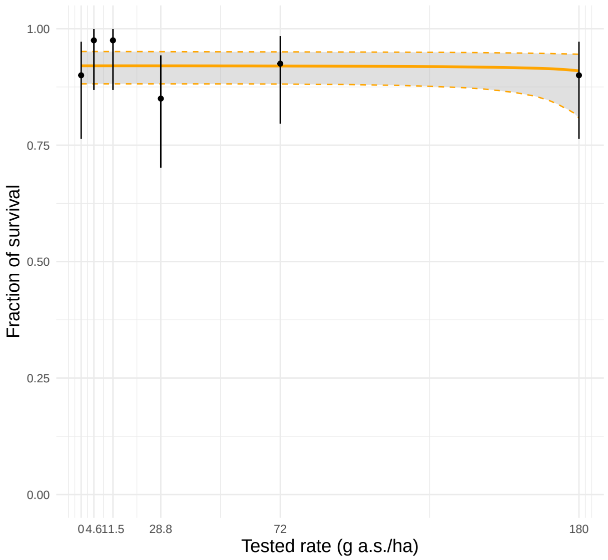

(a) Dose-response curve

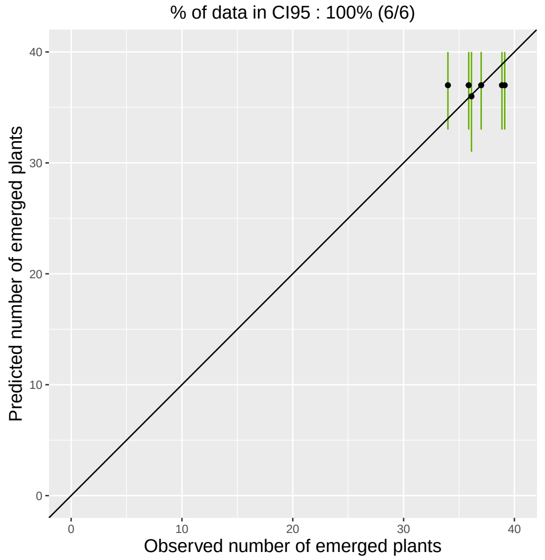

(b) Posterior predictive check (PPC)

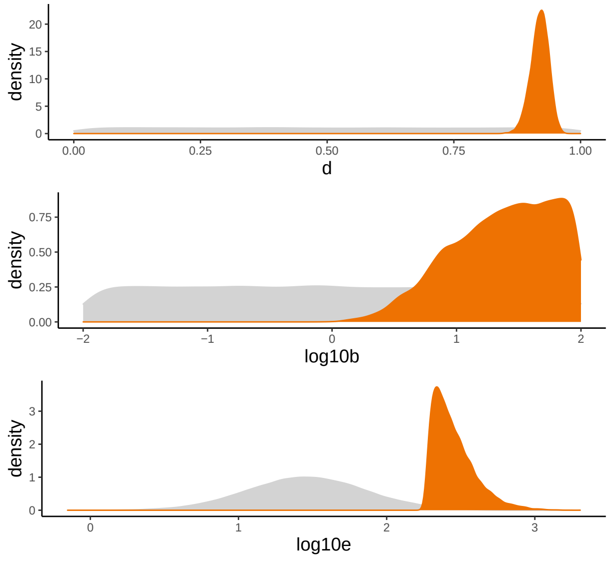

(c) Priors and posteriors

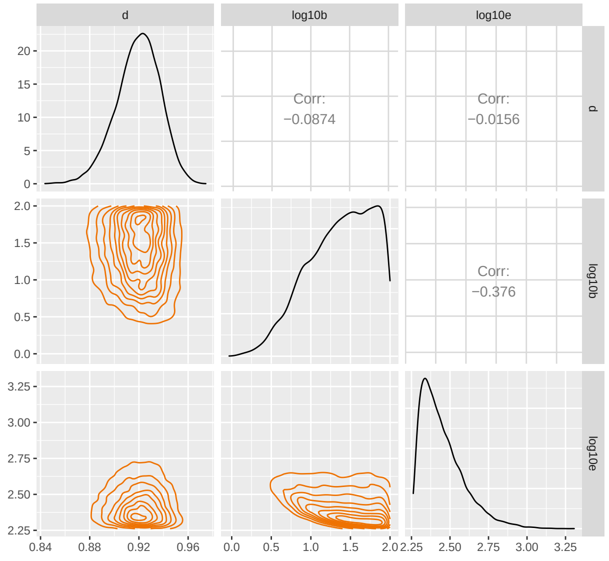

(d) Correlations between parameters

Figure 5: Dose-response curve (a), PPC (b), prior and posterior distributions (c) and correlations between parameters (d).

### Data set: GLXMA_SE_emergence

Table 6: Summary of parameter estimates for GLXMA_SE_emergence data set

| Parameter | median | Q2.5 | Q97.5 |
|---|---|---|---|
| b | 21.972 | 2.247 | 92.993 |
| d | 0.918 | 0.878 | 0.951 |
| e | 262.148 | 189.786 | 661.344 |

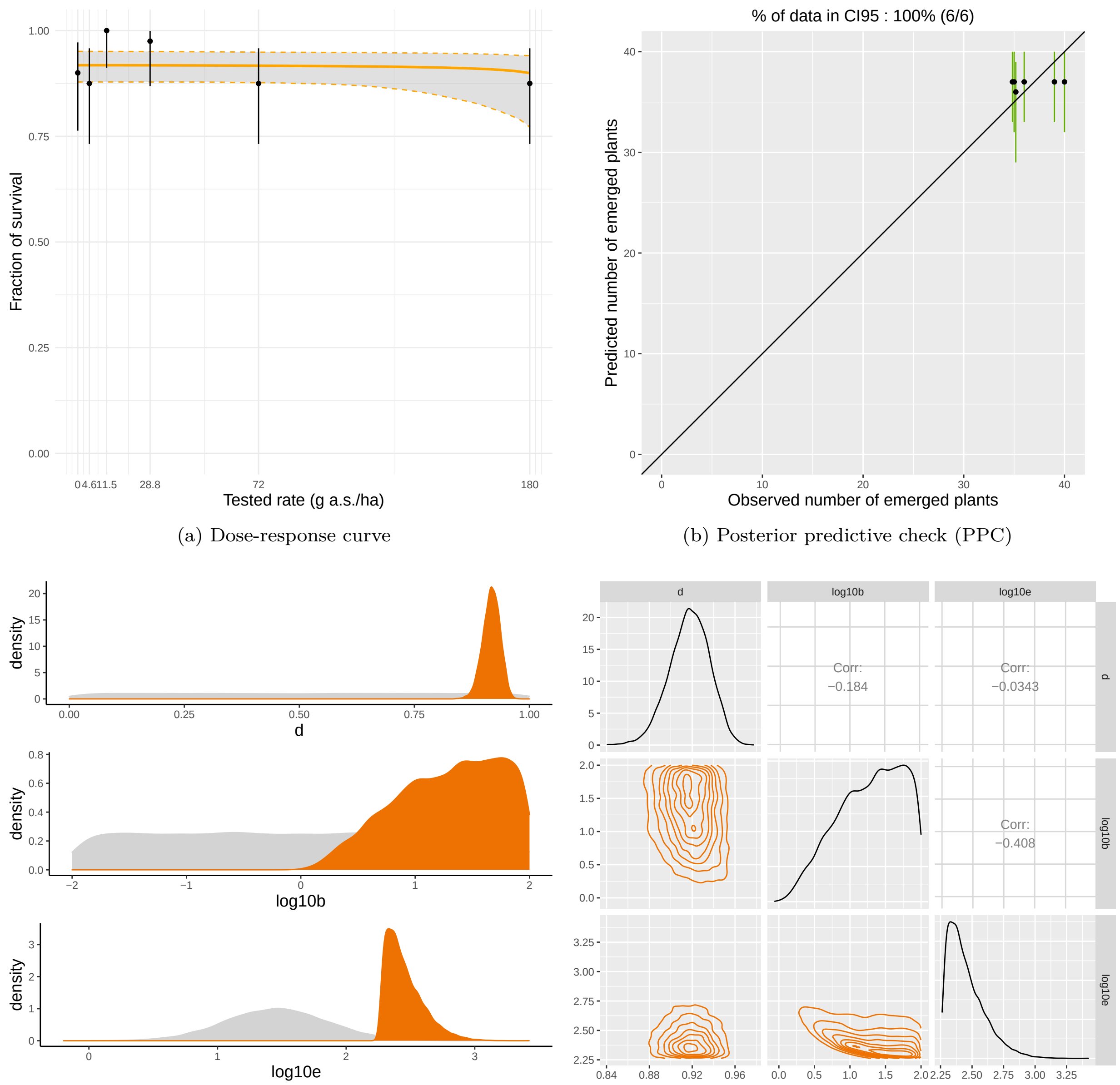

(a) Dose-response curve

(b) Posterior predictive check (PPC)

(c) Priors and posteriors

(d) Correlations between parameters

Figure 6: Dose-response curve (a), PPC (b), prior and posterior distributions (c) and correlations between parameters (d).

#### Data set: HELAN_SE_emergence

Table 7: Summary of parameter estimates for HELAN_SE_emergence data set

| Parameter | median | Q2.5 | Q97.5 |
|---|---|---|---|
| b | 35.784 | 5.436 | 94.713 |
| d | 0.960 | 0.931 | 0.980 |
| e | 274.842 | 195.480 | 668.667 |

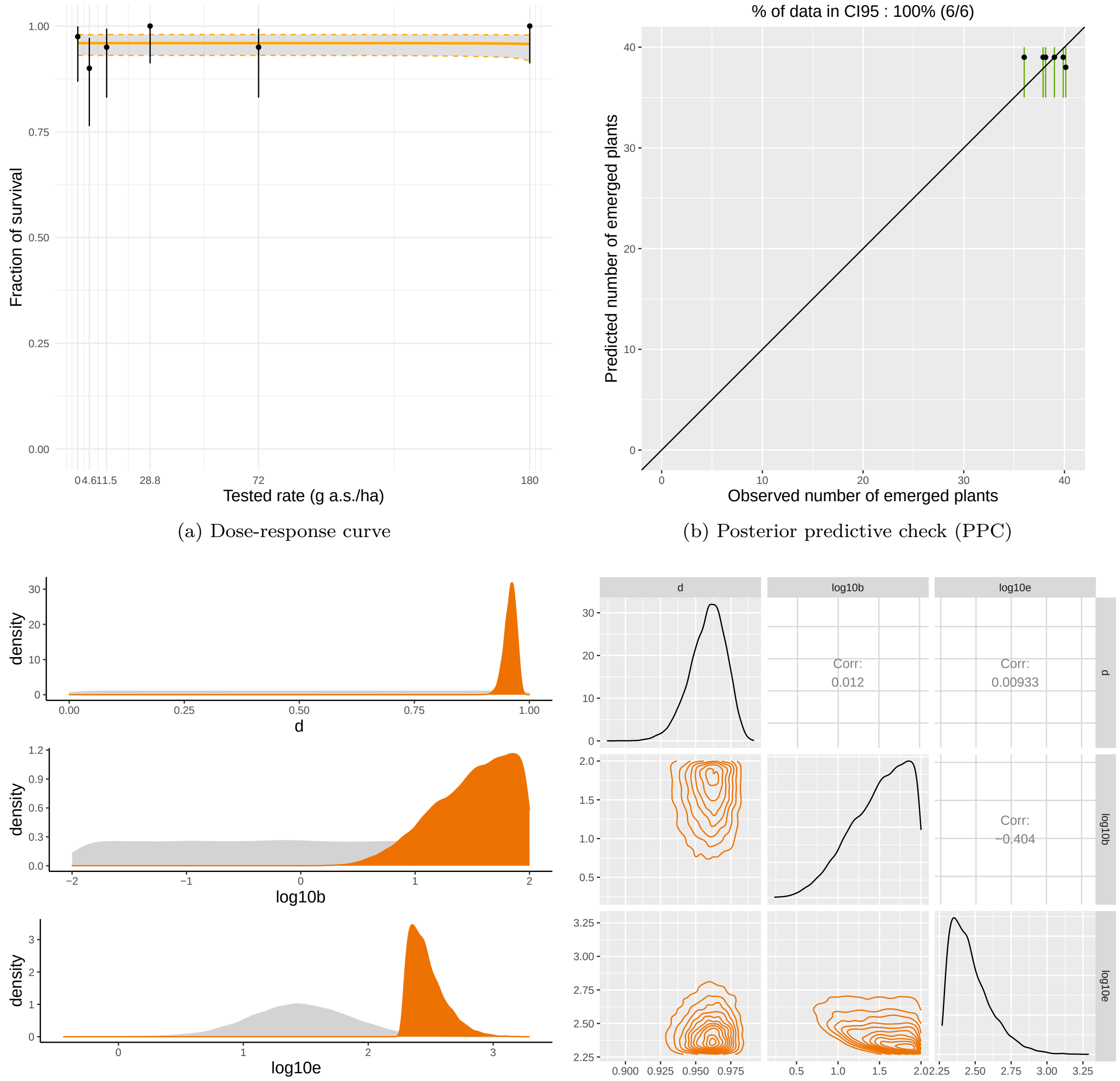

(a) Dose-response curve

(b) Posterior predictive check (PPC)

(c) Priors and posteriors

(d) Correlations between parameters

Figure 7: Dose-response curve (a), PPC (b), prior and posterior distributions (c) and correlations between parameters (d).

#### Data set: LOLPE_SE_emergence

Table 8: Summary of parameter estimates for LOLPE_SE_emergence data set

| Parameter | median | Q2.5 | Q97.5 |
|---|---|---|---|
| b | 31.536 | 4.408 | 94.940 |
| d | 0.890 | 0.847 | 0.925 |
| e | 107.631 | 77.598 | 267.661 |

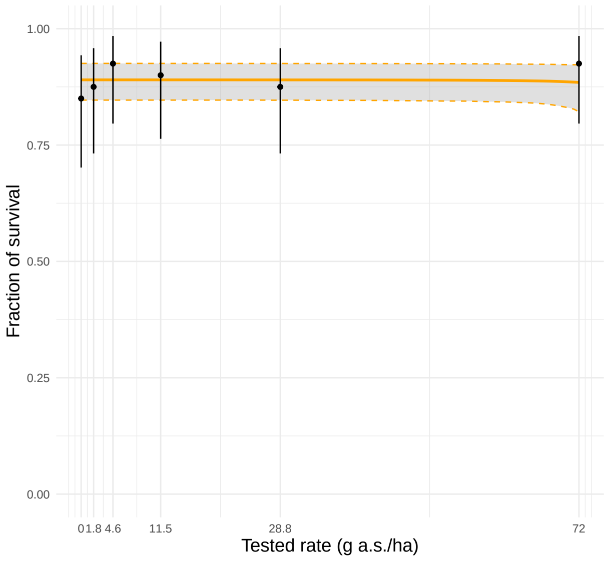

(a) Dose-response curve

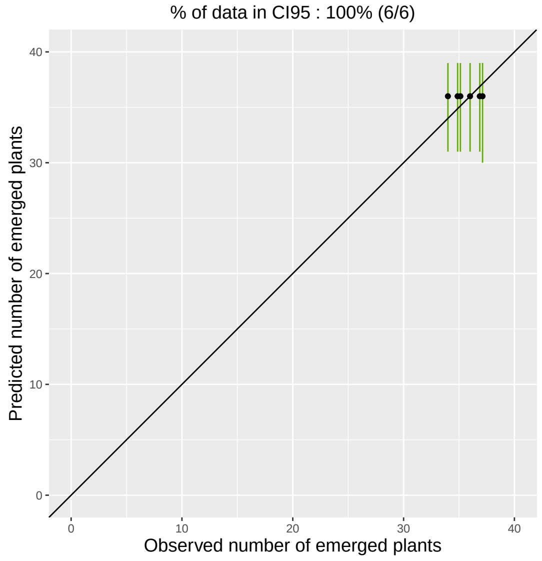

(b) Posterior predictive check (PPC)

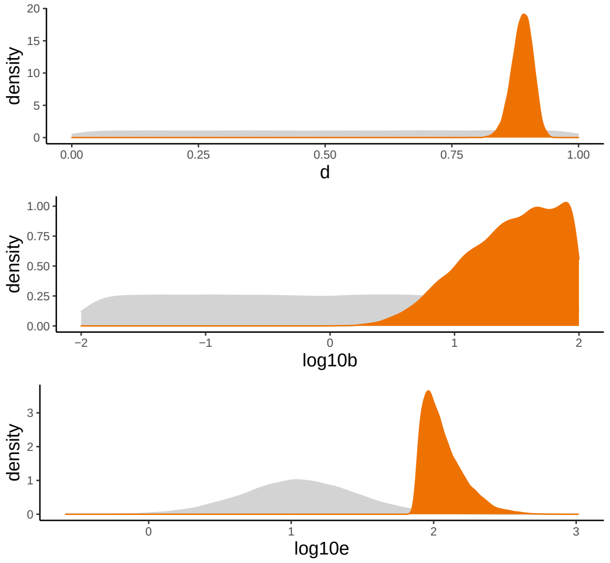

(c) Priors and posteriors

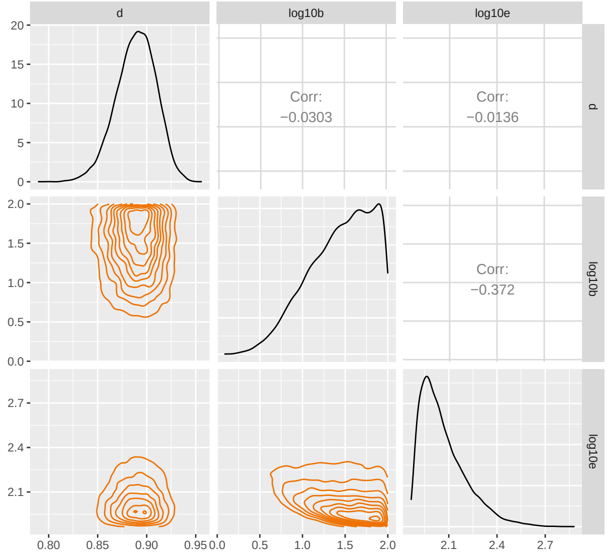

(d) Correlations between parameters

Figure 8: Dose-response curve (a), PPC (b), prior and posterior distributions (c) and correlations between parameters (d).

#### Data set: LYPES_SE_emergence

Table 9: Summary of parameter estimates (parameter d is set to 1) for LYPES_SE_emergence data set

| Parameter | median | Q2.5 | Q97.5 |
|---|---|---|---|
| b | 2.326 | 1.318 | 4.176 |
| e | 543.278 | 311.511 | 1338.246 |

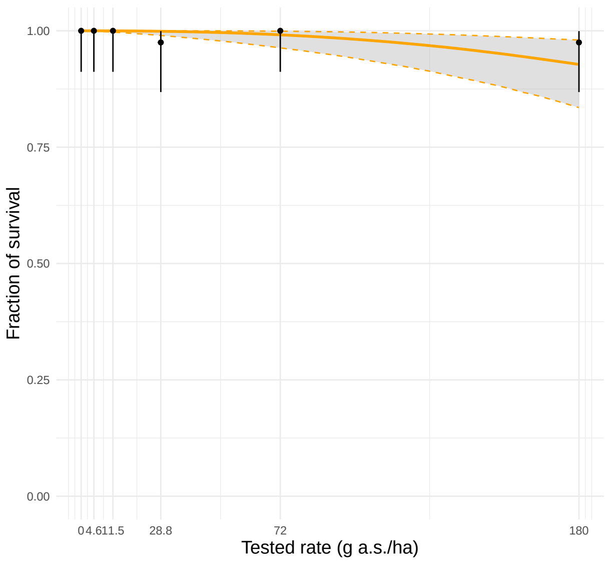

(a) Dose-response curve

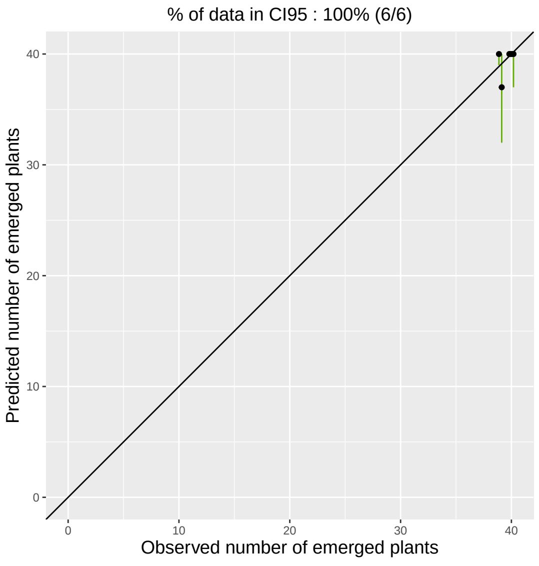

(b) Posterior predictive check (PPC)

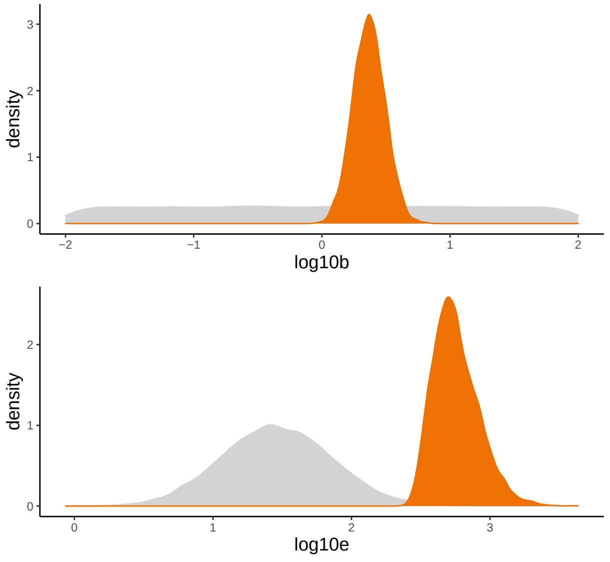

(c) Priors and posteriors

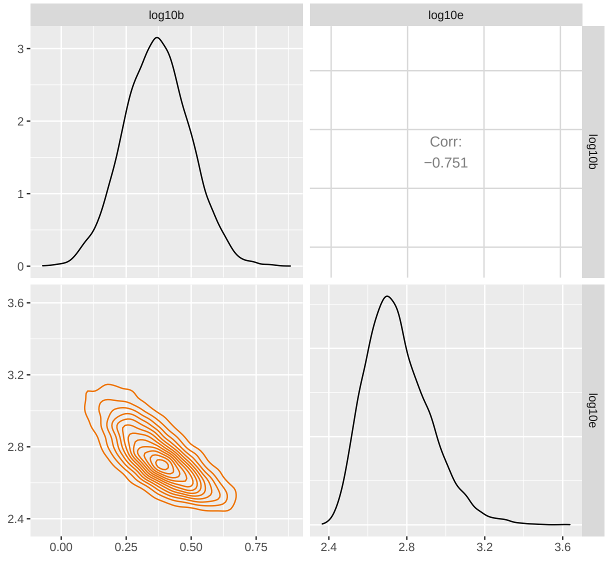

(d) Correlations between parameters

Figure 9: Dose-response curve (a), PPC (b), prior and posterior distributions (c) and correlations between parameters (d).

#### Data set: ZEAMA_SE_emergence

Table 10: Summary of parameter estimates (parameter d is set to 1) for ZEAMA_SE_emergence data set

| Parameter | median | Q2.5 | Q97.5 |
|---|---|---|---|
| b | 1.039 | 0.721 | 1.498 |
| e | 1096.913 | 516.603 | 2938.759 |

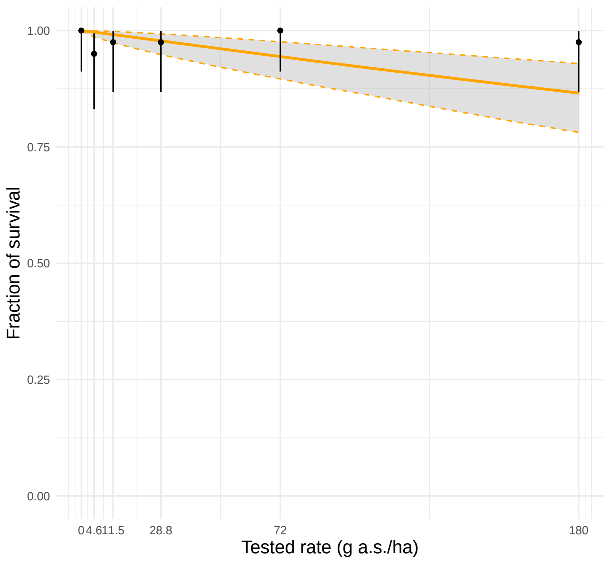

(a) Dose-response curve

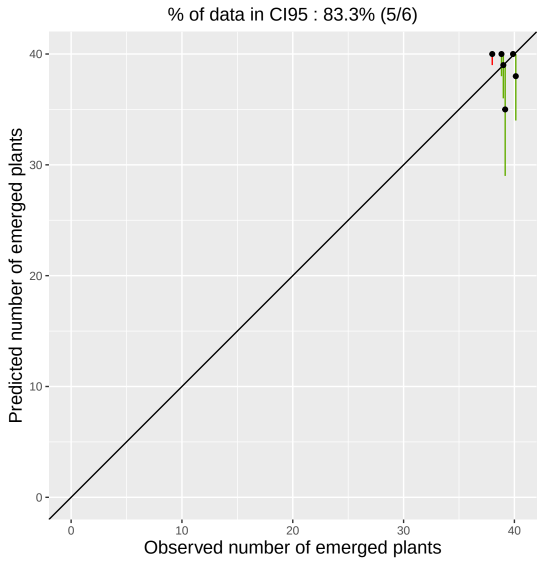

(b) Posterior predictive check (PPC)

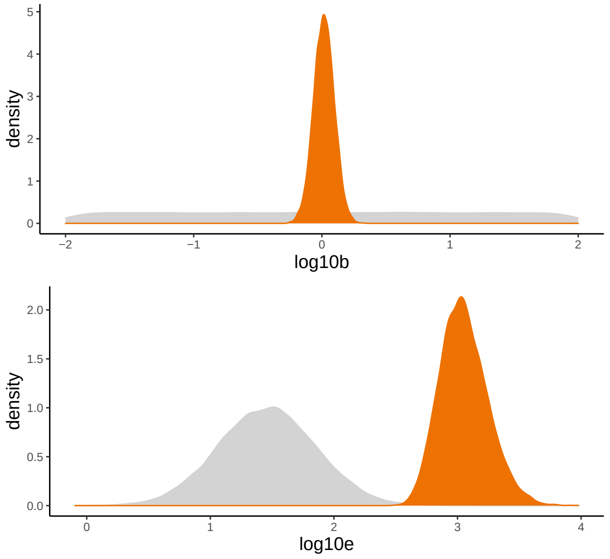

(c) Priors and posteriors

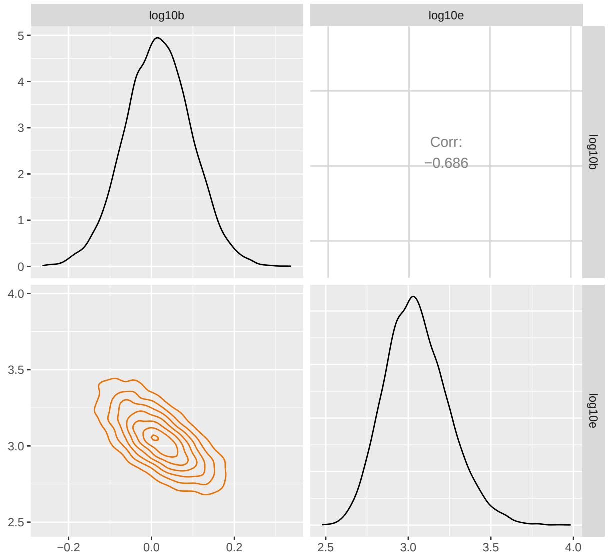

(d) Correlations between parameters

Figure 10: Dose-response curve (a), PPC (b), prior and posterior distributions (c) and correlations between parameters (d).
