## Supplementary material for "How to account for the uncertainty from standard toxicity tests in species sensitivity distributions: an example in non-target plants": Archive with reports on studies: report_SE_weight.pdf

### Dose-response analysis

Study 2

Seedling Emergence test - shoot dry SE_weight endpoint

25 June 2020

Contact:

---

This is a report which provides results on all performed dose-response analyses for the shoot dry SE_weight endpoint of the Seedling Emergence test for study 2.

---

#### Data set: ALLCE_SE_weight

Table 1: Summary of parameter estimates for ALLCE_SE_weight data set

| Parameter | median | Q2.5 | Q97.5 |
|---|---|---|---|
| b | 1.964 | 1.055 | 4.073 |
| d | 0.045 | 0.041 | 0.050 |
| e | 7.954 | 5.785 | 10.746 |
| sigma | 0.010 | 0.009 | 0.013 |

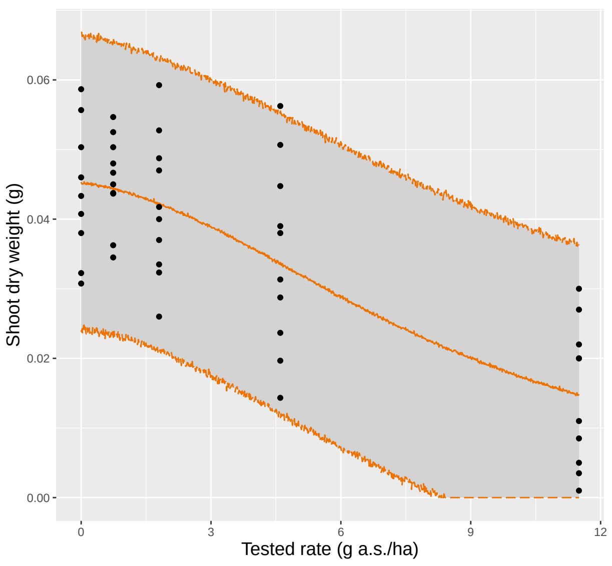

(a) Dose-response curve

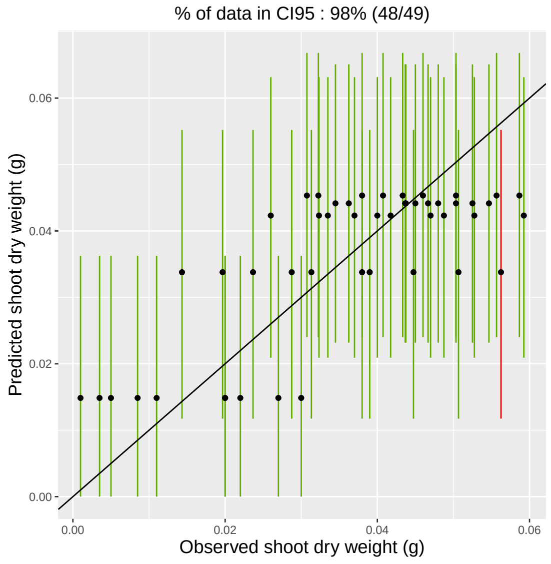

(b) Posterior predictive check (PPC)

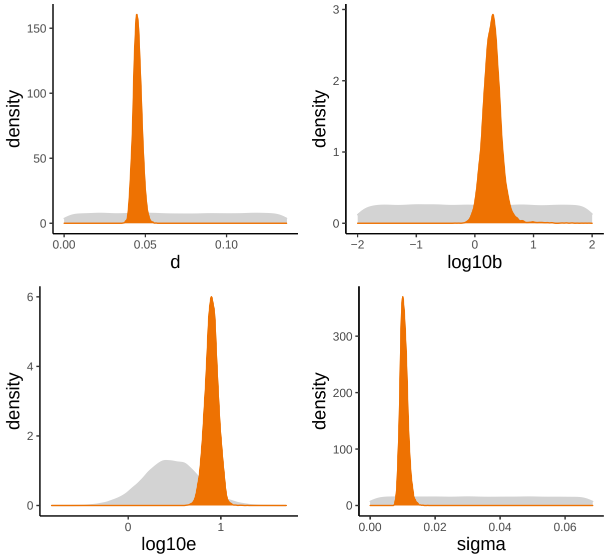

(c) Priors and posteriors

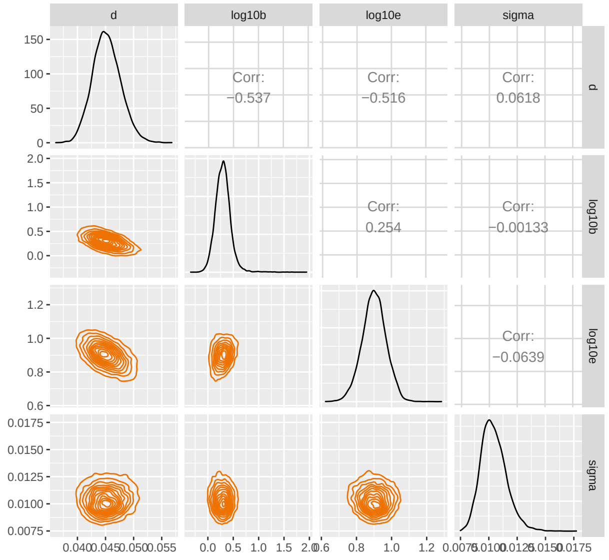

(d) Correlations between parameters

Figure 1: Dose-response curve (a), PPC (b), prior and posterior distributions (c) and correlations between parameters (d).

#### Data set: AVESA_SE_weight

Table 2: Summary of parameter estimates for AVESA_SE_weight data set

| Parameter | median | Q2.5 | Q97.5 |
|---|---|---|---|
| b | 0.971 | 0.548 | 1.707 |
| d | 0.599 | 0.552 | 0.660 |
| e | 217.238 | 147.926 | 374.312 |
| sigma | 0.102 | 0.086 | 0.125 |

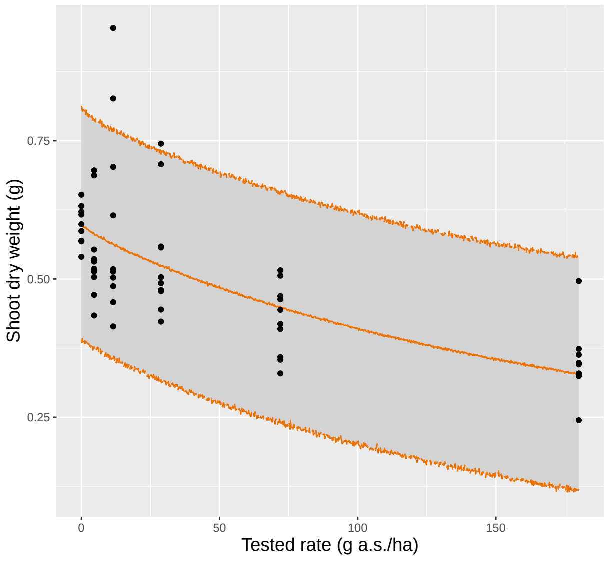

(a) Dose-response curve

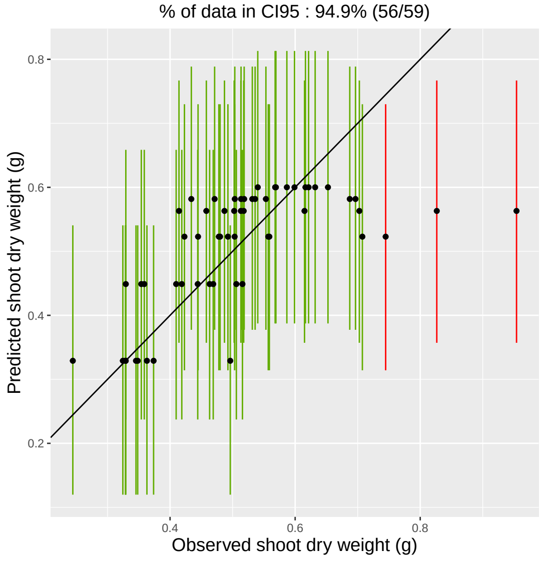

(b) Posterior predictive check (PPC)

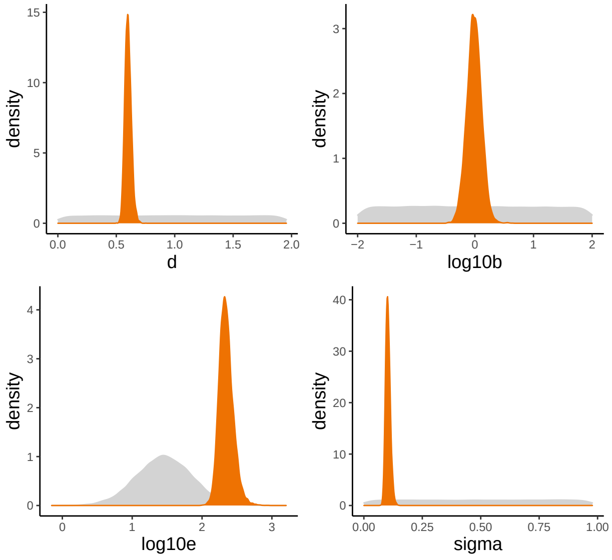

(c) Priors and posteriors

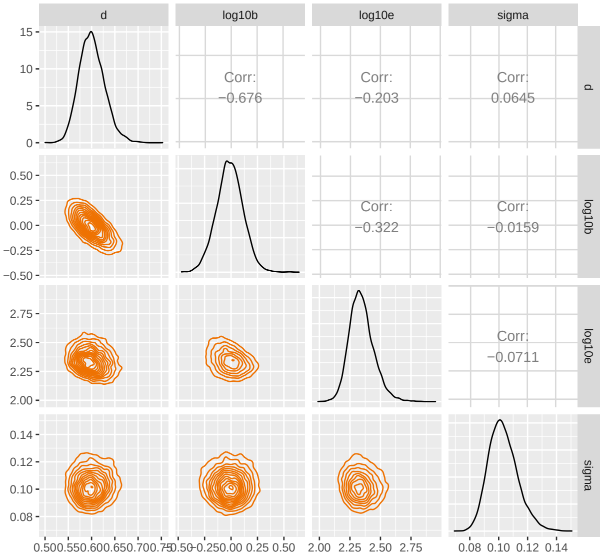

(d) Correlations between parameters

Figure 2: Dose-response curve (a), PPC (b), prior and posterior distributions (c) and correlations between parameters (d).

#### Data set: BEAVA_SE_weight

Table 3: Summary of parameter estimates for BEAVA_SE_weight data set

| Parameter | median | Q2.5 | Q97.5 |
|---|---|---|---|
| b | 5.248 | 1.150 | 64.725 |
| d | 1.014 | 0.957 | 1.073 |
| e | 38.374 | 29.507 | 88.484 |
| sigma | 0.256 | 0.226 | 0.293 |

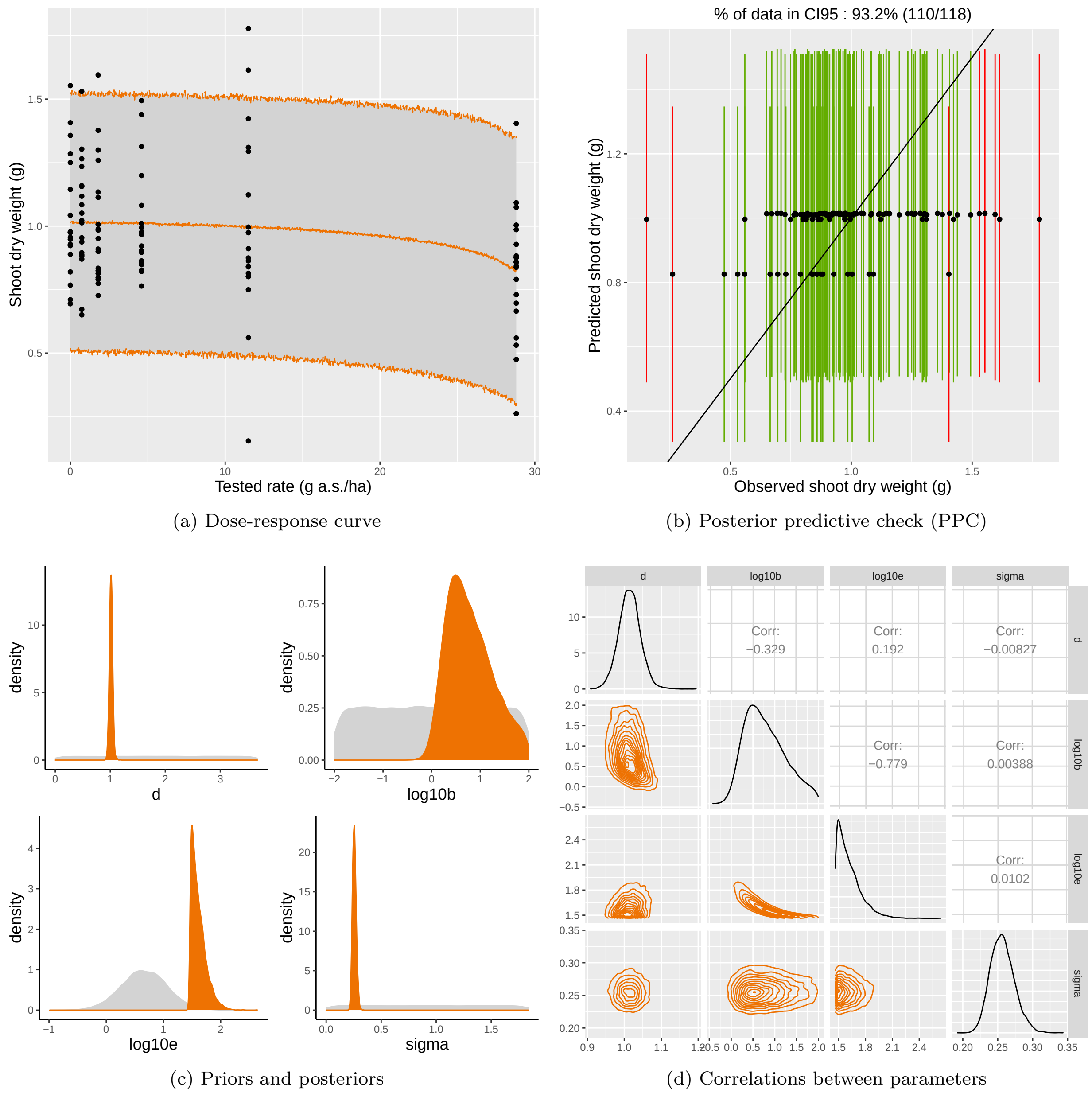

(a) Dose-response curve

(b) Posterior predictive check (PPC)

(c) Priors and posteriors

(d) Correlations between parameters

Figure 3: Dose-response curve (a), PPC (b), prior and posterior distributions (c) and correlations between parameters (d).

#### Data set: BRSNW_SE_weight

Table 4: Summary of parameter estimates for BRSNW_SE_weight data set

| Parameter | median | Q2.5 | Q97.5 |
|---|---|---|---|
| b | 0.965 | 0.586 | 1.770 |
| d | 1.321 | 1.220 | 1.443 |
| e | 208.450 | 143.197 | 357.522 |
| sigma | 0.314 | 0.276 | 0.358 |

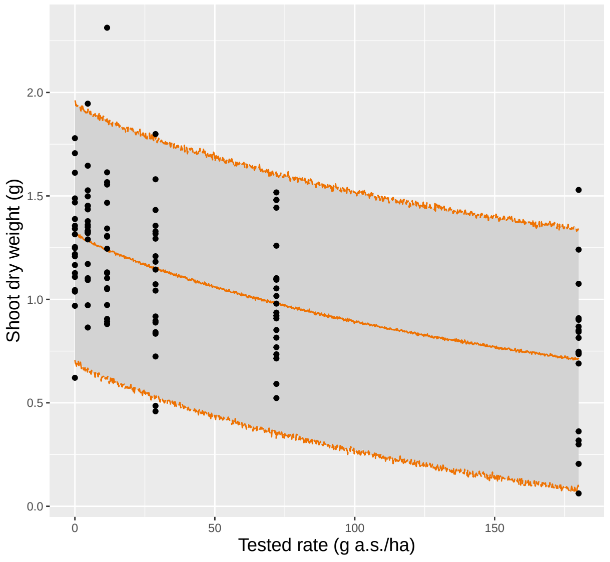

(a) Dose-response curve

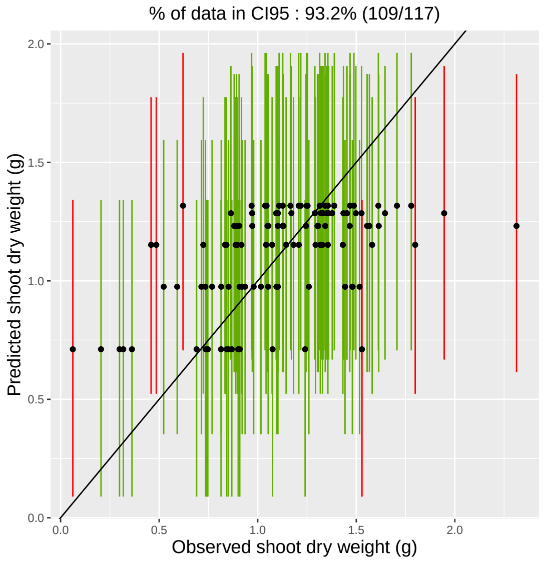

(b) Posterior predictive check (PPC)

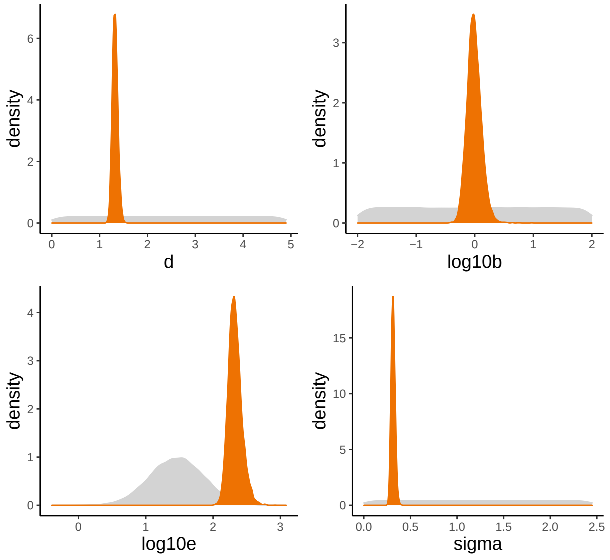

(c) Priors and posteriors

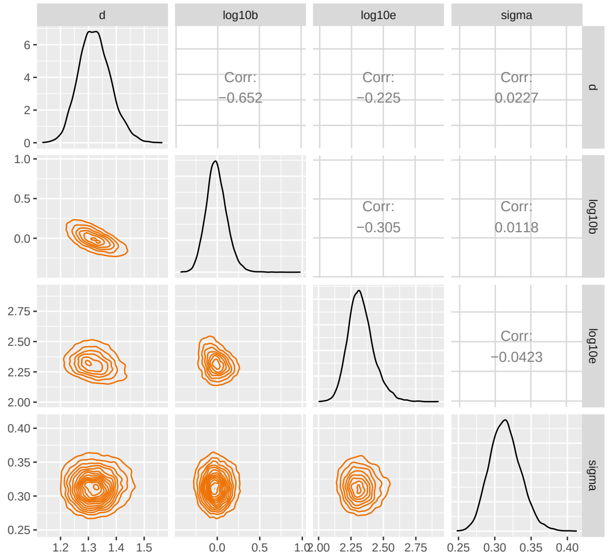

(d) Correlations between parameters

Figure 4: Dose-response curve (a), PPC (b), prior and posterior distributions (c) and correlations between parameters (d).

### Data set: CUMSA_SE_weight

Table 5: Summary of parameter estimates for CUMSA_SE_weight data set

| Parameter | median | Q2.5 | Q97.5 |
|---|---|---|---|
| b | 0.904 | 0.628 | 1.315 |
| d | 2.871 | 2.614 | 3.137 |
| e | 51.999 | 36.541 | 75.689 |
| sigma | 0.654 | 0.576 | 0.750 |

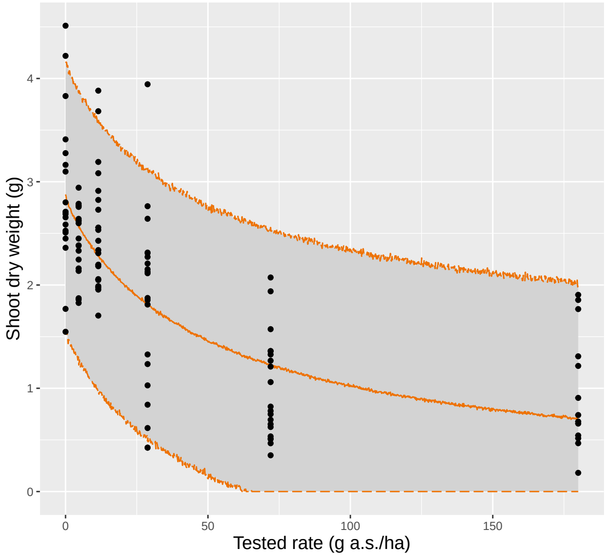

(a) Dose-response curve

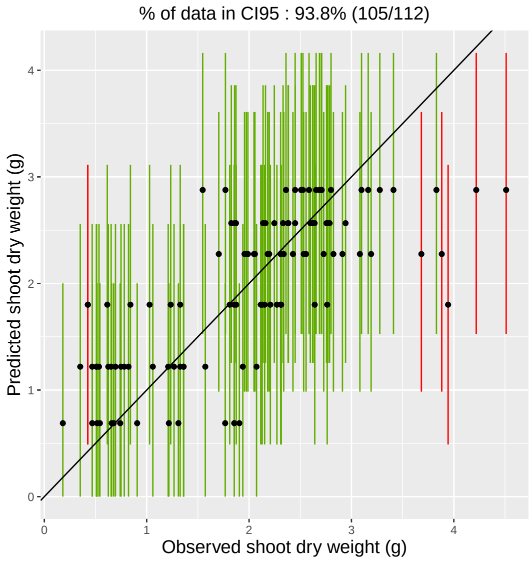

(b) Posterior predictive check (PPC)

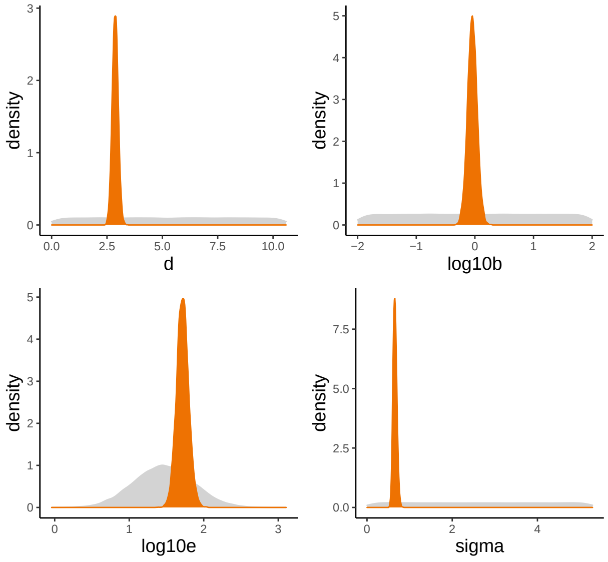

(c) Priors and posteriors

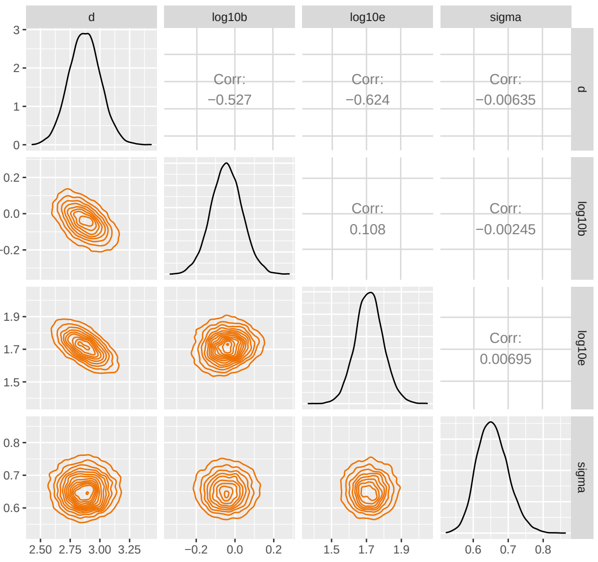

(d) Correlations between parameters

Figure 5: Dose-response curve (a), PPC (b), prior and posterior distributions (c) and correlations between parameters (d).

#### Data set: GLXMA_SE_weight

Table 6: Summary of parameter estimates for GLXMA_SE_weight data set

| Parameter | median | Q2.5 | Q97.5 |
|---|---|---|---|
| b | 25.033 | 3.336 | 93.096 |
| d | 2.197 | 2.116 | 2.284 |
| e | 259.458 | 189.967 | 632.591 |
| sigma | 0.437 | 0.386 | 0.501 |

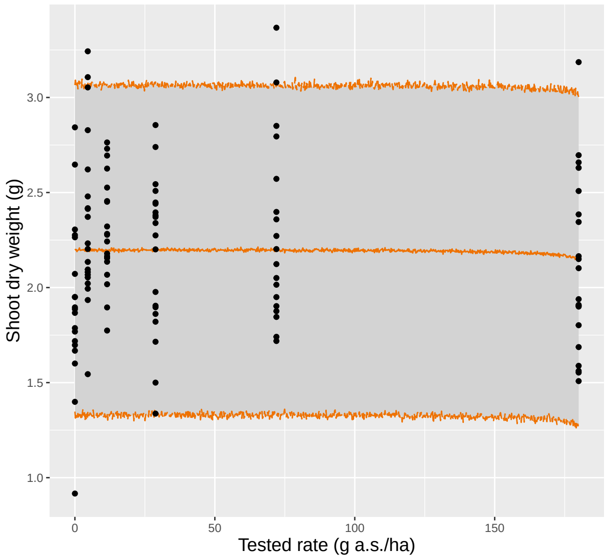

(a) Dose-response curve

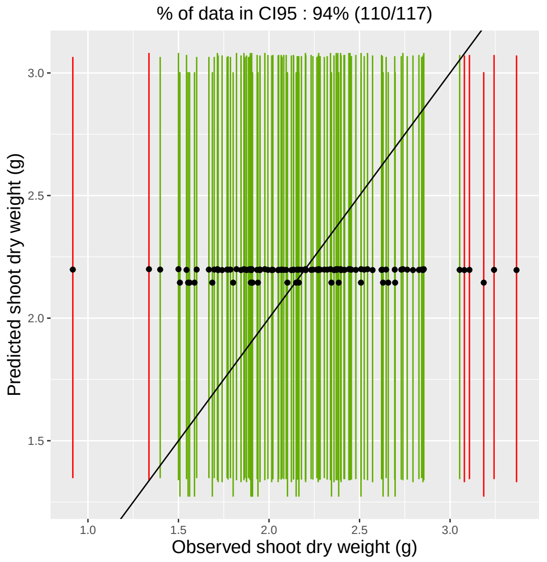

(b) Posterior predictive check (PPC)

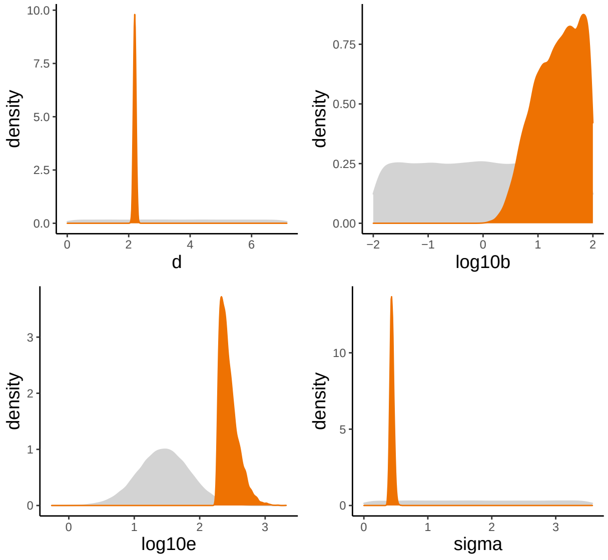

(c) Priors and posteriors

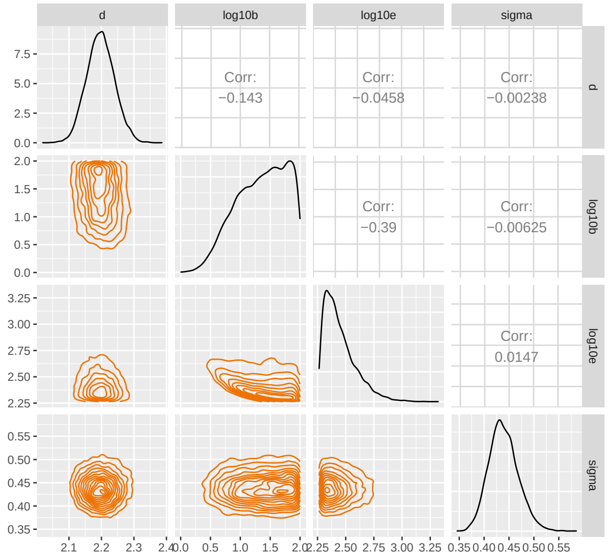

(d) Correlations between parameters

Figure 6: Dose-response curve (a), PPC (b), prior and posterior distributions (c) and correlations between parameters (d).

#### Data set: HELAN_SE_weight

Table 7: Summary of parameter estimates for HELAN_SE_weight data set

| Parameter | median | Q2.5 | Q97.5 |
|---|---|---|---|
| b | 33.124 | 4.954 | 94.387 |
| d | 0.551 | 0.530 | 0.570 |
| e | 272.888 | 195.420 | 674.027 |
| sigma | 0.109 | 0.097 | 0.125 |

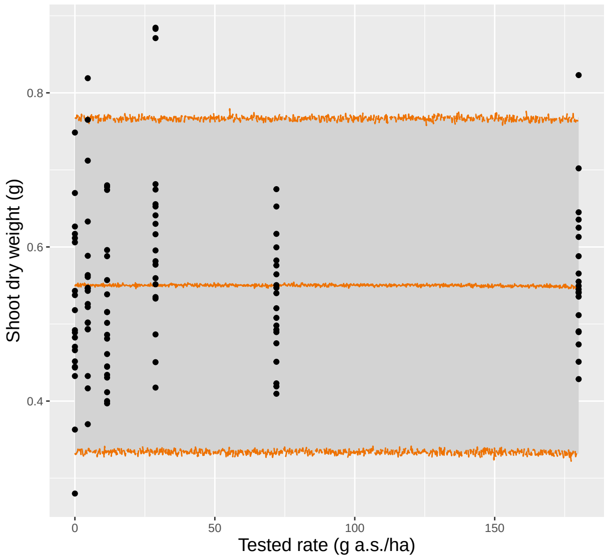

(a) Dose-response curve

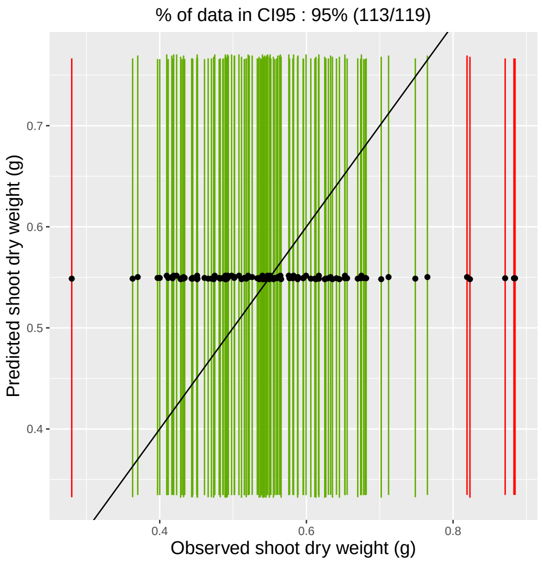

(b) Posterior predictive check (PPC)

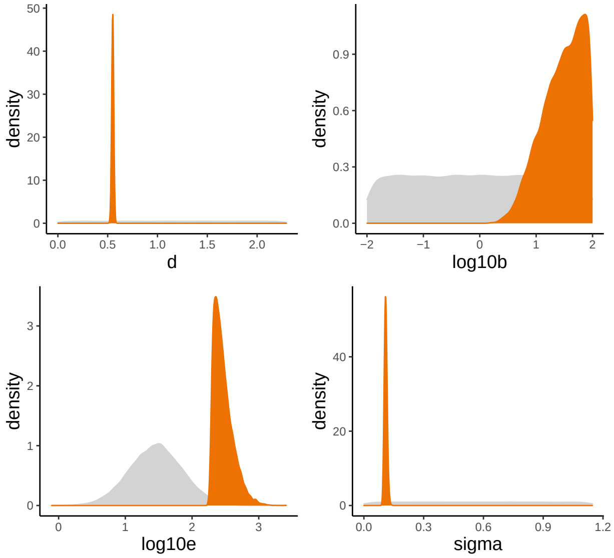

(c) Priors and posteriors

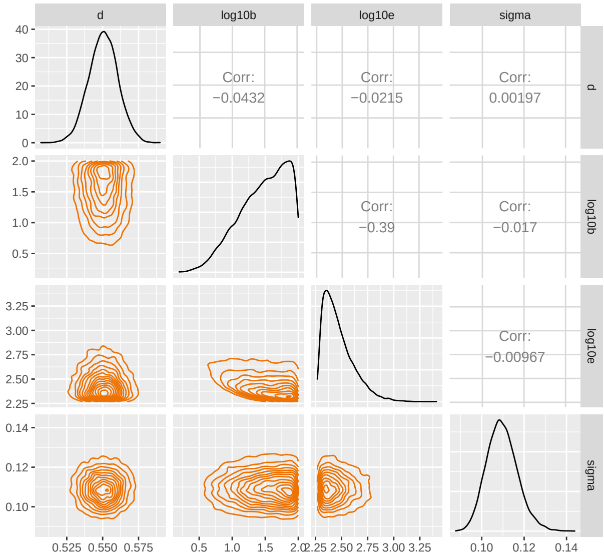

(d) Correlations between parameters

Figure 7: Dose-response curve (a), PPC (b), prior and posterior distributions (c) and correlations between parameters (d).

#### Data set: LOLPE_SE_weight

Table 8: Summary of parameter estimates for LOLPE_SE_weight data set

| Parameter | median | Q2.5 | Q97.5 |
|---|---|---|---|
| b | 4.431 | 1.384 | 56.887 |
| d | 0.104 | 0.094 | 0.114 |
| e | 74.133 | 51.281 | 137.118 |
| sigma | 0.031 | 0.026 | 0.039 |

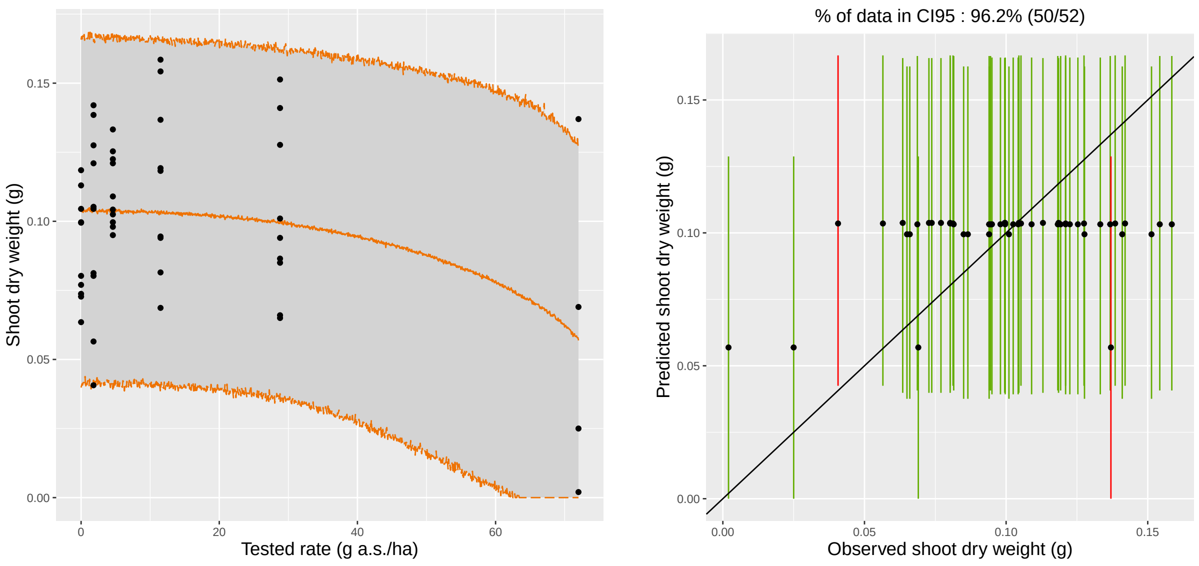

(a) Dose-response curve

(b) Posterior predictive check (PPC)

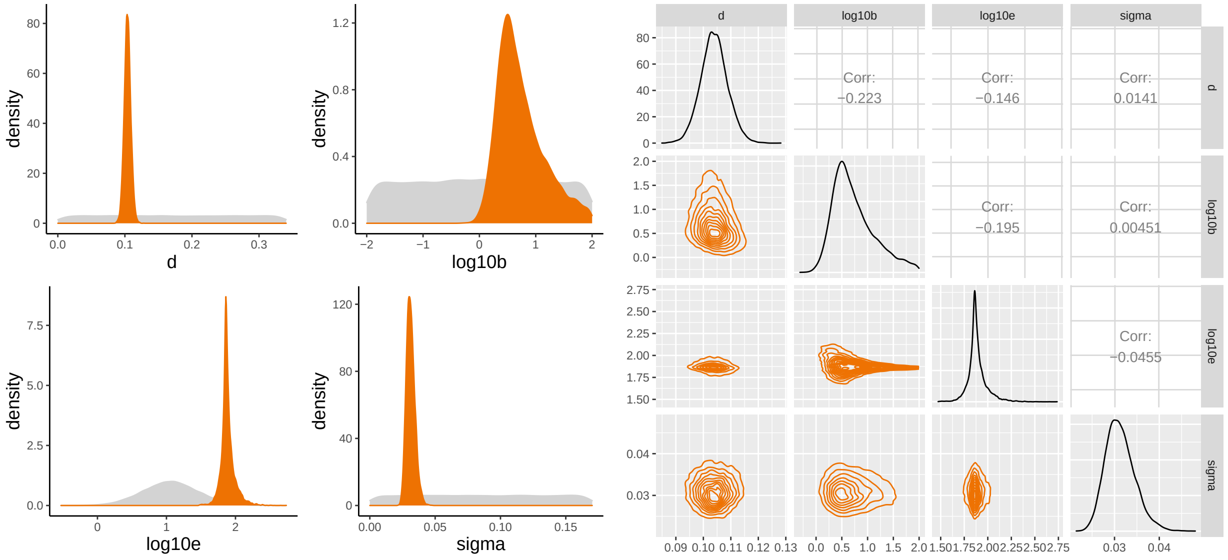

(c) Priors and posteriors

(d) Correlations between parameters

Figure 8: Dose-response curve (a), PPC (b), prior and posterior distributions (c) and correlations between parameters (d).

#### Data set: LYPES_SE_weight

Table 9: Summary of parameter estimates for LYPES_SE_weight data set

| Parameter | median | Q2.5 | Q97.5 |
|---|---:|---:|---:|
| b | 0.889 | 0.620 | 1.321 |
| d | 1.286 | 1.187 | 1.396 |
| e | 103.170 | 74.600 | 144.647 |
| sigma | 0.258 | 0.228 | 0.296 |

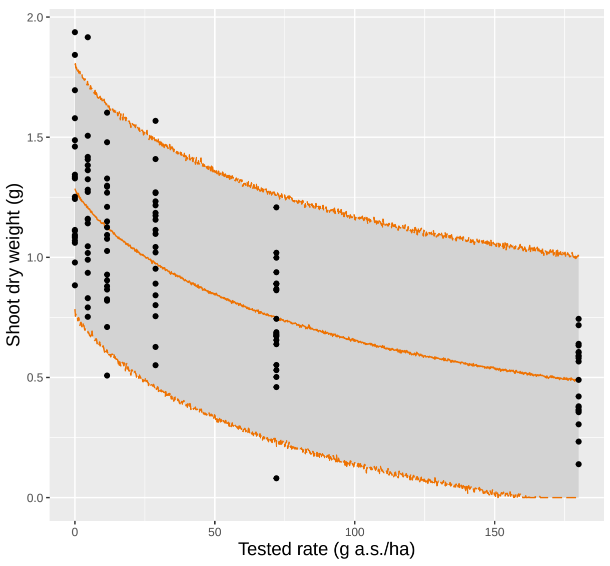

(a) Dose-response curve

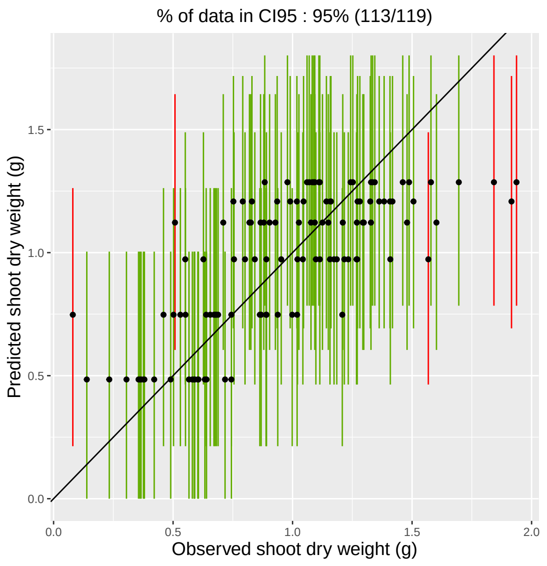

(b) Posterior predictive check (PPC)

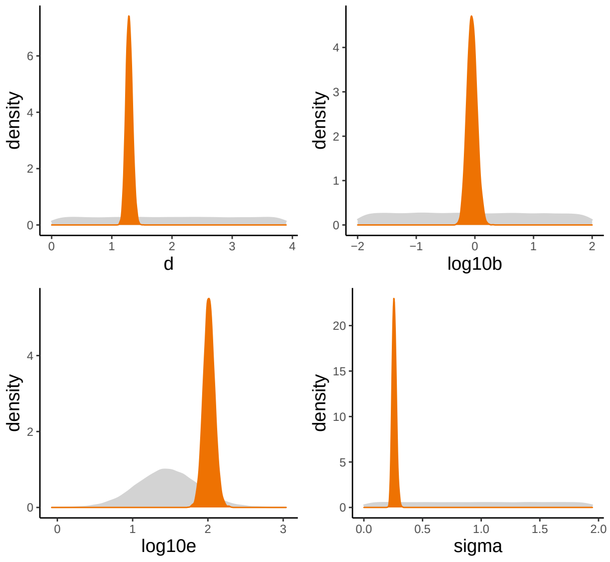

(c) Priors and posteriors

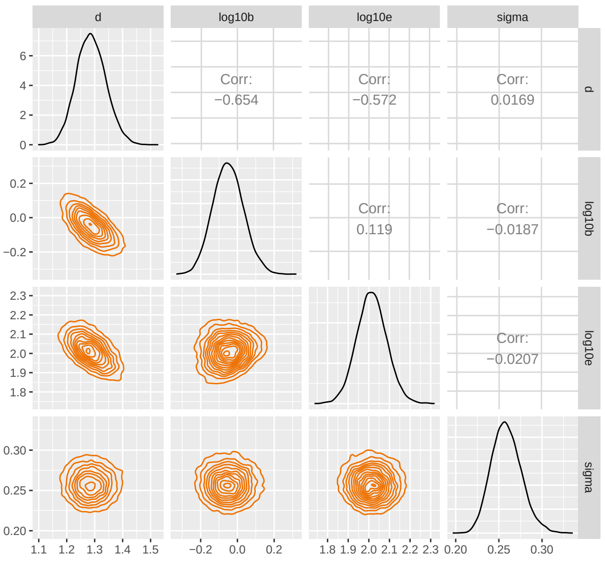

(d) Correlations between parameters

Figure 9: Dose-response curve (a), PPC (b), prior and posterior distributions (c) and correlations between parameters (d).

#### Data set: ZEAMA_SE_weight

Table 10: Summary of parameter estimates for ZEAMA_SE_weight data set

| Parameter | median | Q2.5 | Q97.5 |
|---|---|---|---|
| b | 17.004 | 2.717 | 91.516 |
| d | 3.783 | 3.674 | 3.895 |
| e | 252.786 | 188.393 | 622.364 |
| sigma | 0.554 | 0.489 | 0.633 |

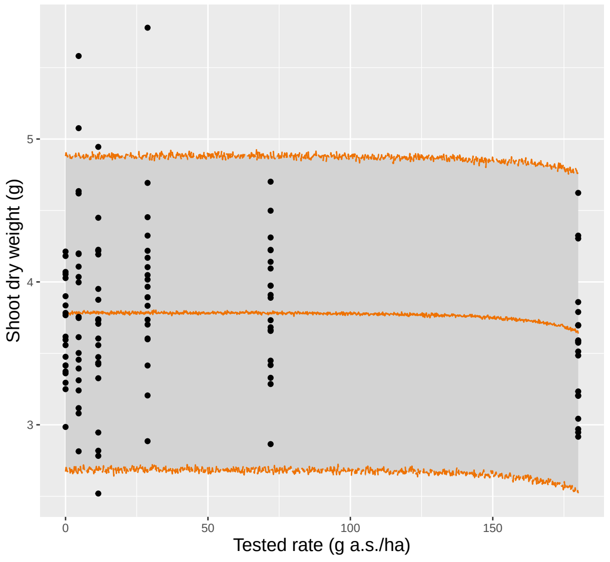

(a) Dose-response curve

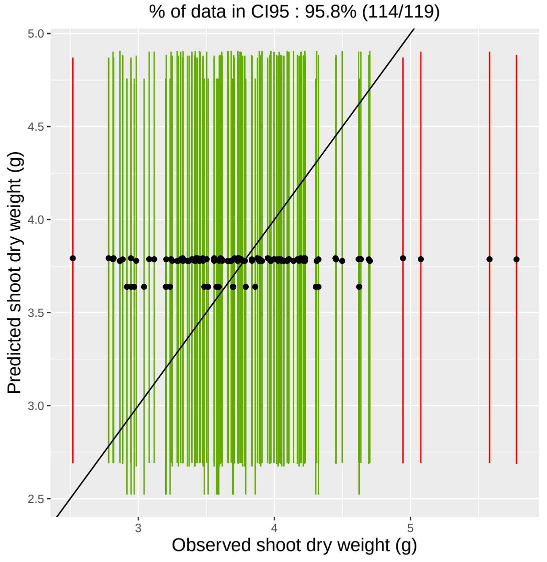

(b) Posterior predictive check (PPC)

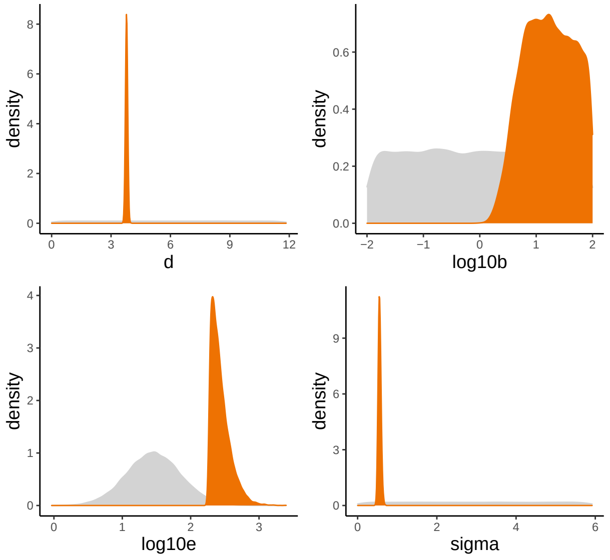

(c) Priors and posteriors

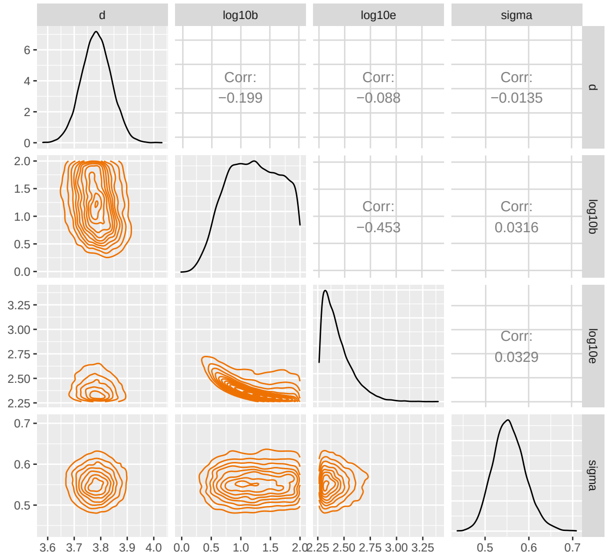

(d) Correlations between parameters

Figure 10: Dose-response curve (a), PPC (b), prior and posterior distributions (c) and correlations between parameters (d).
