## Supplementary material for "How to account for the uncertainty from standard toxicity tests in species sensitivity distributions: an example in non-target plants": Archive with reports on studies: report_VV_weight.pdf

### Dose-response analysis

Study 2

Vegetative Vigour test - shoot dry VV_weight endpoint

25 June 2020

Contact:

---

This is a report which provides results on all performed dose-response analyses for the shoot dry VV_weight endpoint of the Vegetative Vigour test for study 2.

---

#### Data set: ALLCE_VV_weight

Table 1: Summary of parameter estimates for ALLCE_VV_weight data set

| Parameter | median | Q2.5 | Q97.5 |
|---|---|---|---|
| b | 27.377 | 3.206 | 94.450 |
| d | 0.069 | 0.064 | 0.073 |
| e | 267.731 | 191.643 | 702.042 |
| sigma | 0.015 | 0.012 | 0.019 |

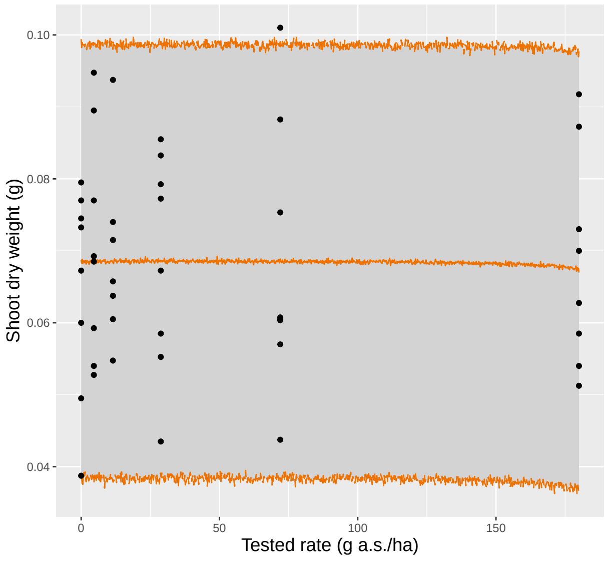

(a) Dose-response curve

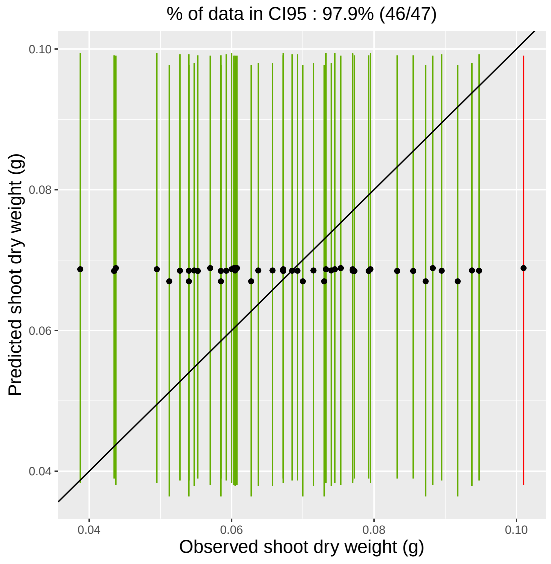

(b) Posterior predictive check (PPC)

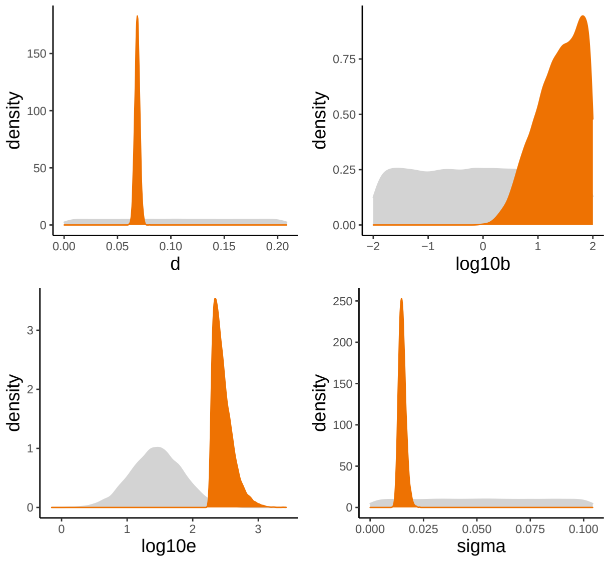

(c) Priors and posteriors

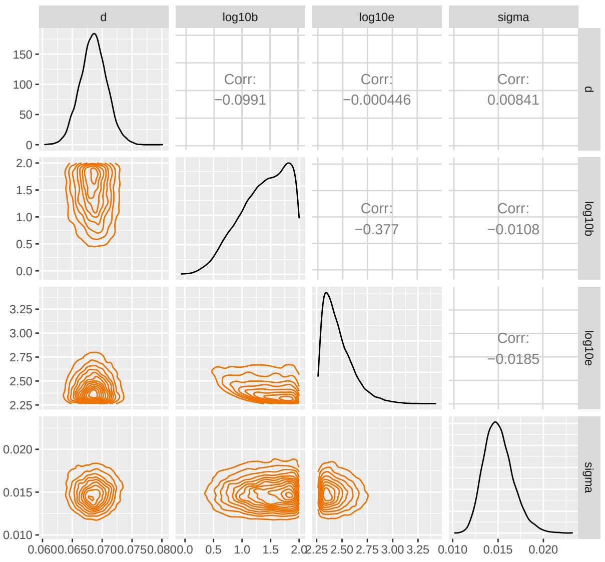

(d) Correlations between parameters

Figure 1: Dose-response curve (a), PPC (b), prior and posterior distributions (c) and correlations between parameters (d).

#### Data set: AVESA_VV_weight

Table 2: Summary of parameter estimates for AVESA_VV_weight data set

| Parameter | median | Q2.5 | Q97.5 |
|---|---|---|---|
| b | 36.267 | 5.761 | 95.815 |
| d | 2.781 | 2.685 | 2.881 |
| e | 275.835 | 195.680 | 719.176 |
| sigma | 0.329 | 0.271 | 0.413 |

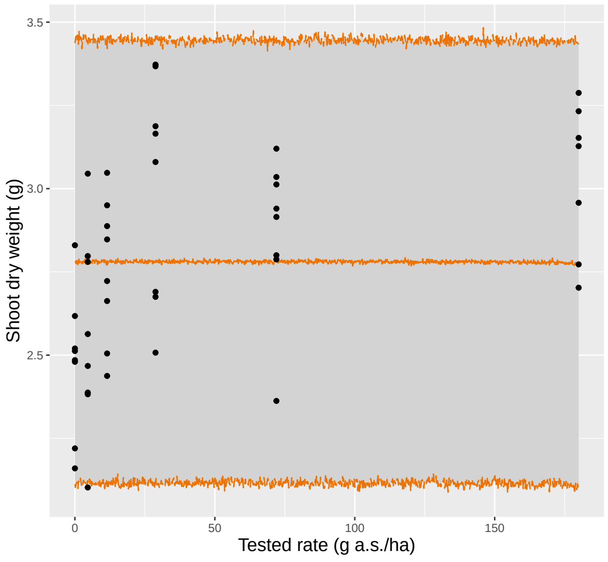

(a) Dose-response curve

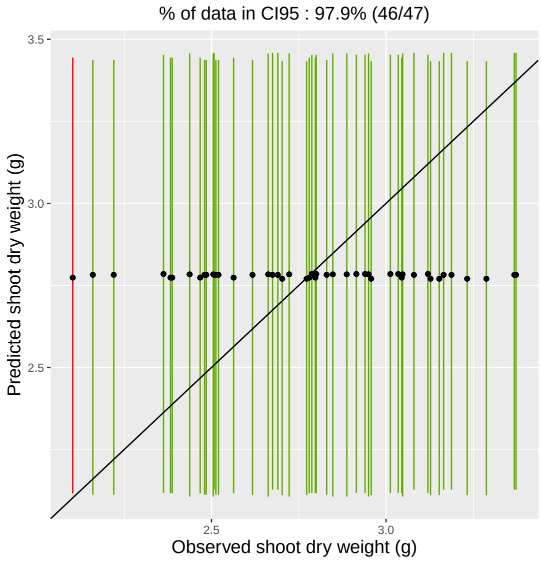

(b) Posterior predictive check (PPC)

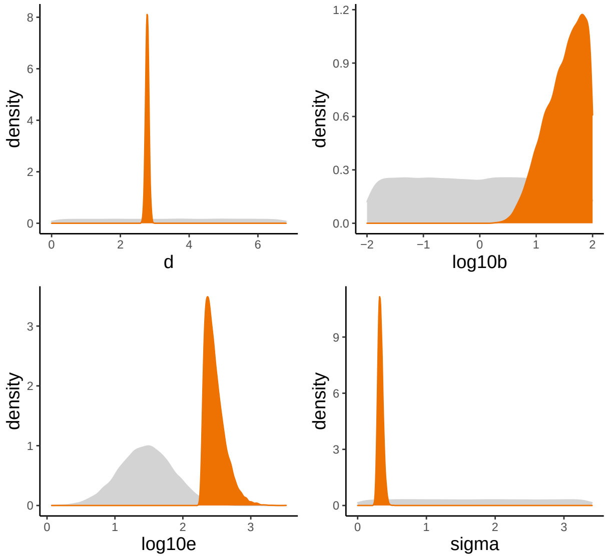

(c) Priors and posteriors

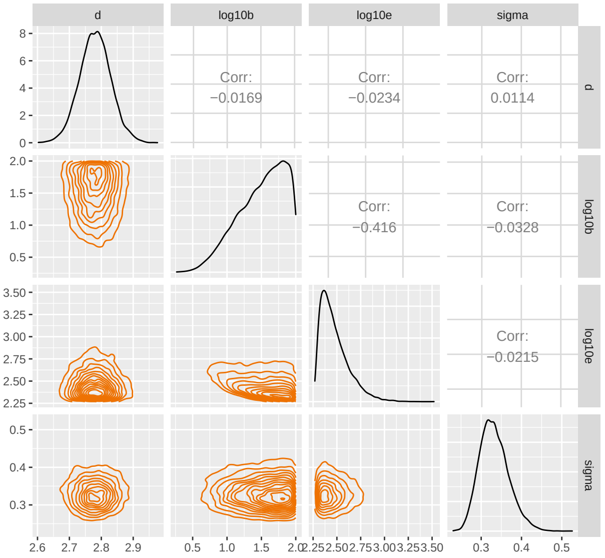

(d) Correlations between parameters

Figure 2: Dose-response curve (a), PPC (b), prior and posterior distributions (c) and correlations between parameters (d).

#### Data set: BEAVA_VV_weight

Table 3: Summary of parameter estimates for BEAVA_VV_weight data set

| Parameter | median | Q2.5 | Q97.5 |
|---|---|---|---|
| b | 4.309 | 1.092 | 54.930 |
| d | 4.464 | 4.361 | 4.600 |
| e | 260.558 | 185.483 | 711.614 |
| sigma | 0.433 | 0.377 | 0.506 |

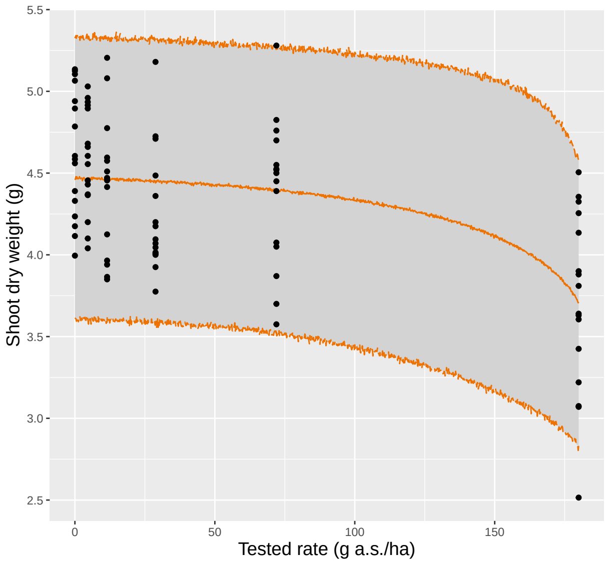

(a) Dose-response curve

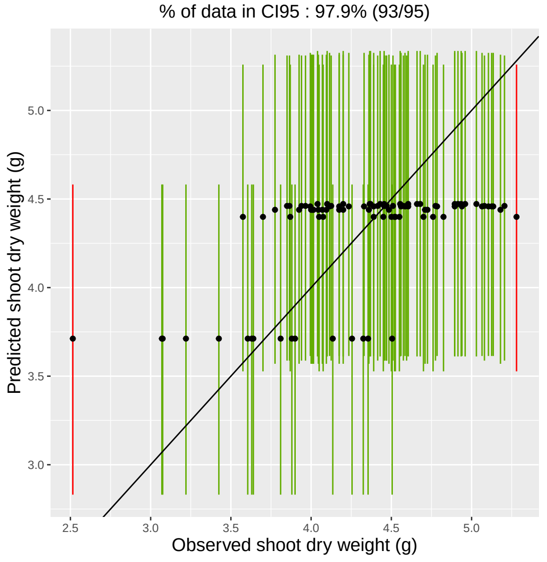

(b) Posterior predictive check (PPC)

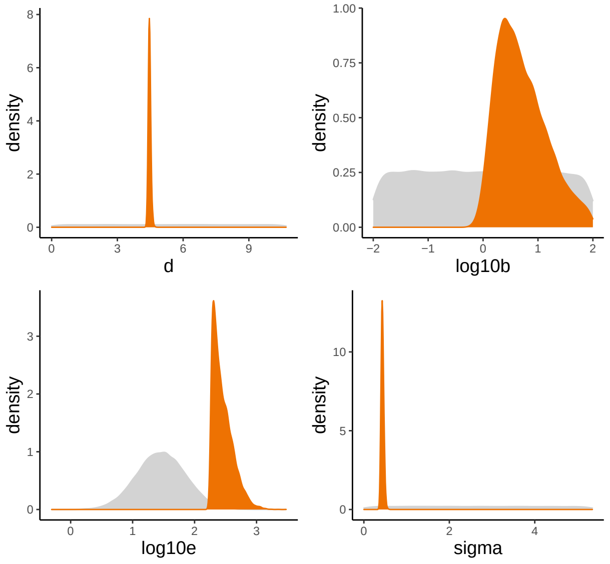

(c) Priors and posteriors

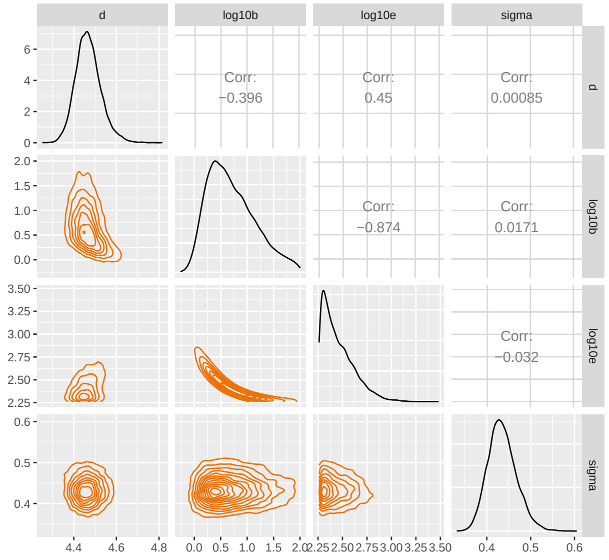

(d) Correlations between parameters

Figure 3: Dose-response curve (a), PPC (b), prior and posterior distributions (c) and correlations between parameters (d).

#### Data set: BRSNW_VV_weight

Table 4: Summary of parameter estimates for BRSNW_VV_weight data set

| Parameter | median | Q2.5 | Q97.5 |
|---|---|---|---|
| b | 0.828 | 0.566 | 1.256 |
| d | 6.786 | 6.440 | 7.187 |
| e | 373.208 | 257.356 | 643.918 |
| sigma | 0.906 | 0.787 | 1.056 |

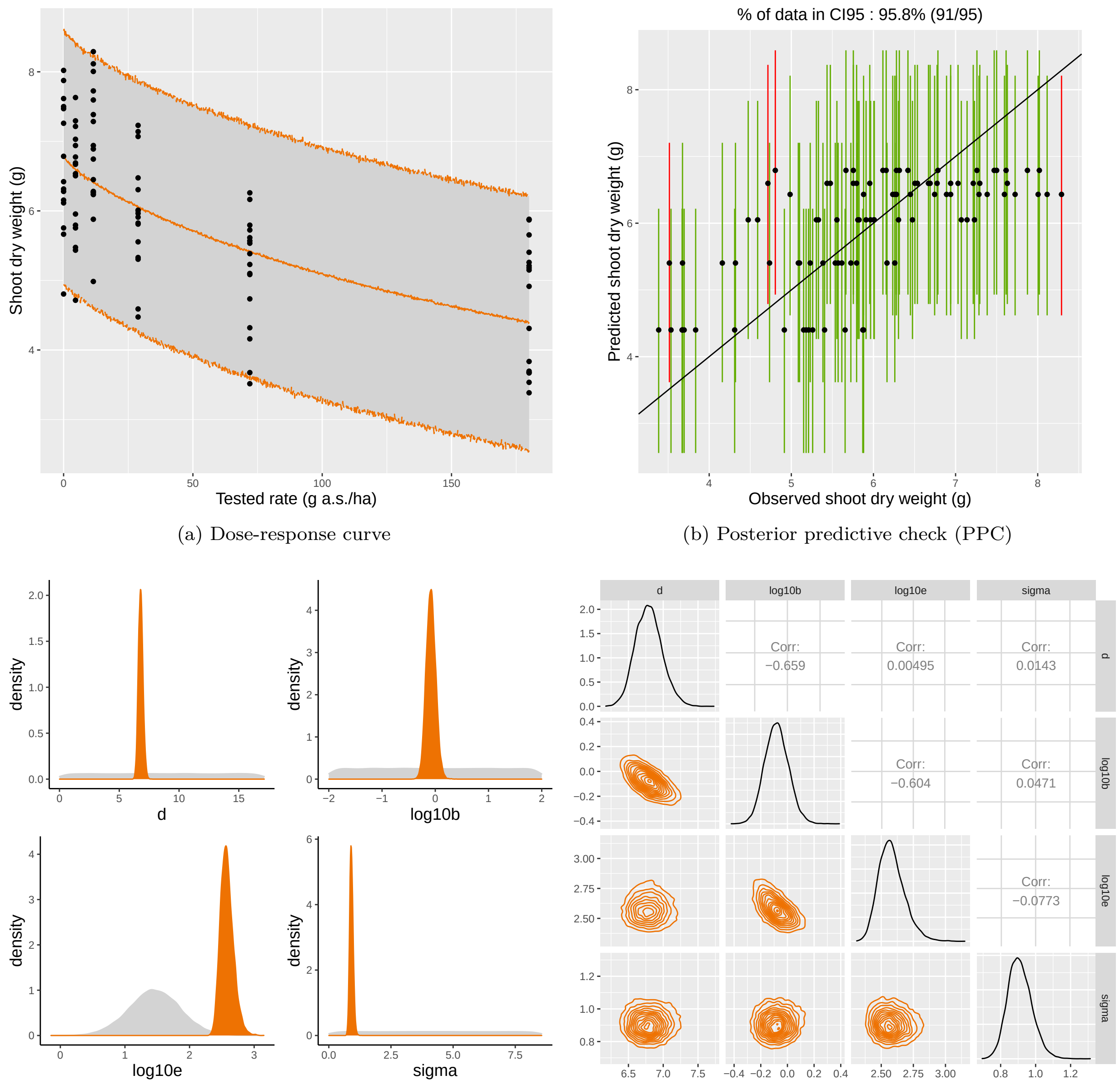

(a) Dose-response curve

(b) Posterior predictive check (PPC)

(c) Priors and posteriors

(d) Correlations between parameters

Figure 4: Dose-response curve (a), PPC (b), prior and posterior distributions (c) and correlations between parameters (d).

#### Data set: CUMSA_VV_weight

Table 5: Summary of parameter estimates for CUMSA_VV_weight data set

| Parameter | median | Q2.5 | Q97.5 |
|---|---|---|---|
| b | 0.509 | 0.380 | 0.686 |
| d | 14.056 | 13.292 | 14.866 |
| e | 422.594 | 267.973 | 776.848 |
| sigma | 1.609 | 1.398 | 1.879 |

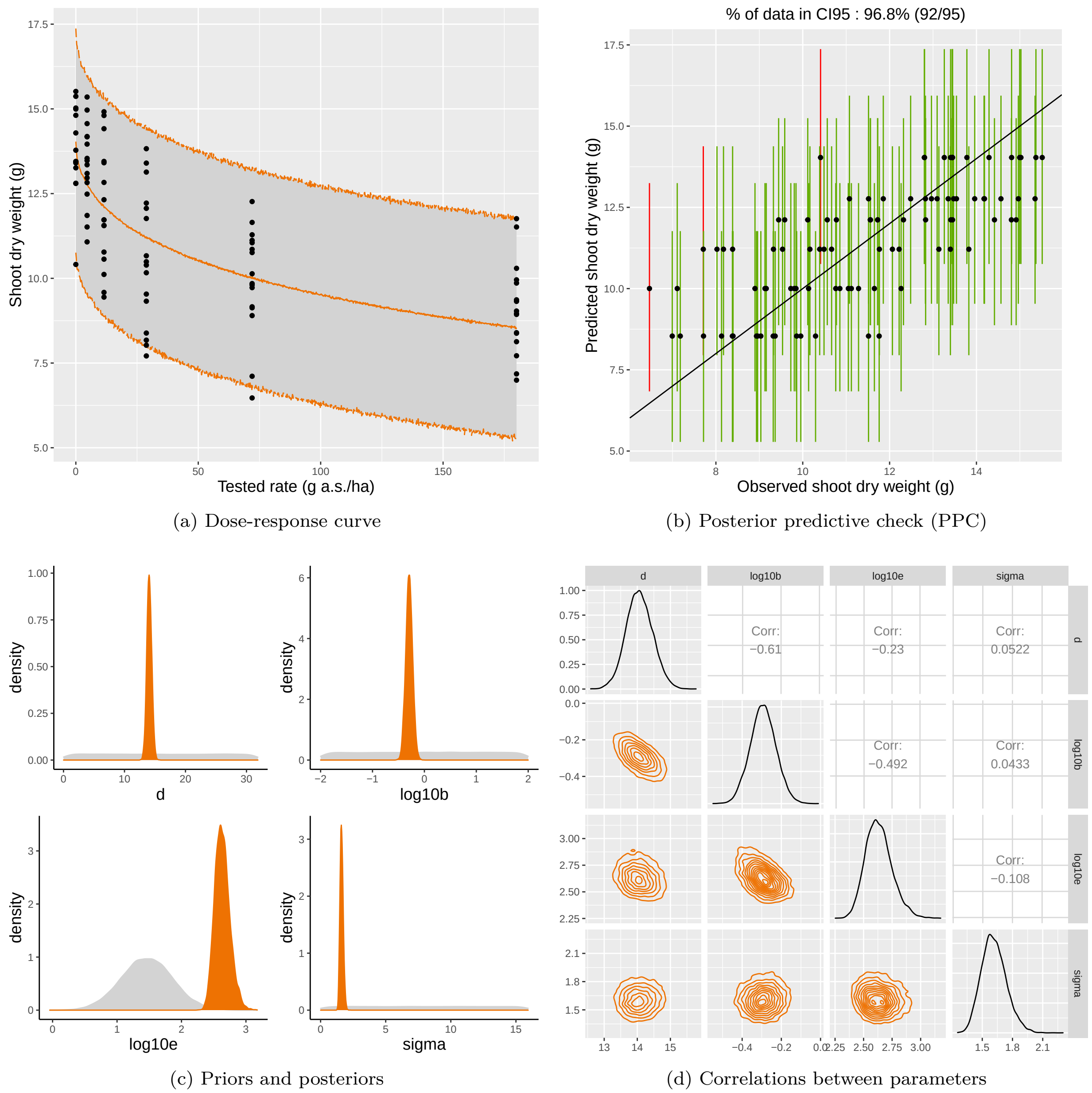

(a) Dose-response curve

(b) Posterior predictive check (PPC)

(c) Priors and posteriors

(d) Correlations between parameters

Figure 5: Dose-response curve (a), PPC (b), prior and posterior distributions (c) and correlations between parameters (d).

#### Data set: GLXMA_VV_weight

Table 6: Summary of parameter estimates for GLXMA_VV_weight data set

| Parameter | median | Q2.5 | Q97.5 |
|---|---|---|---|
| b | 31.159 | 3.593 | 94.501 |
| d | 5.780 | 5.645 | 5.923 |
| e | 270.189 | 193.516 | 720.077 |
| sigma | 0.658 | 0.576 | 0.765 |

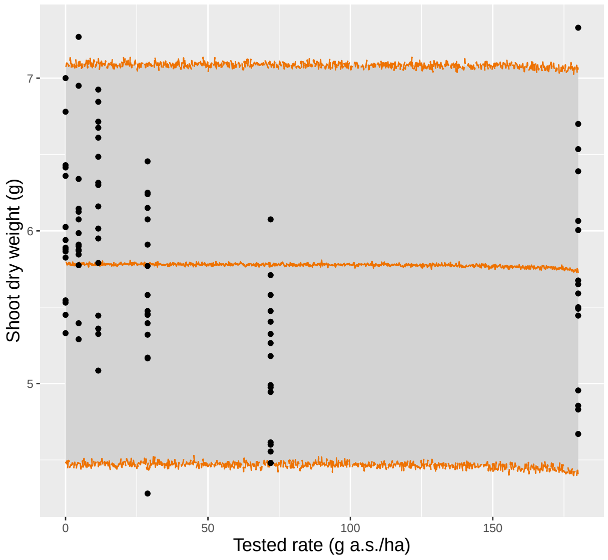

(a) Dose-response curve

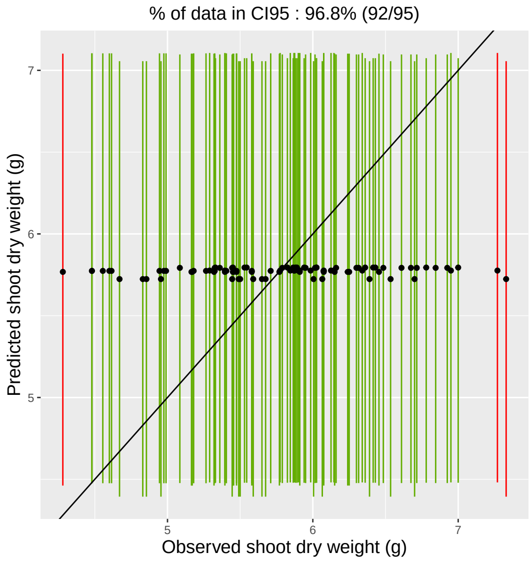

(b) Posterior predictive check (PPC)

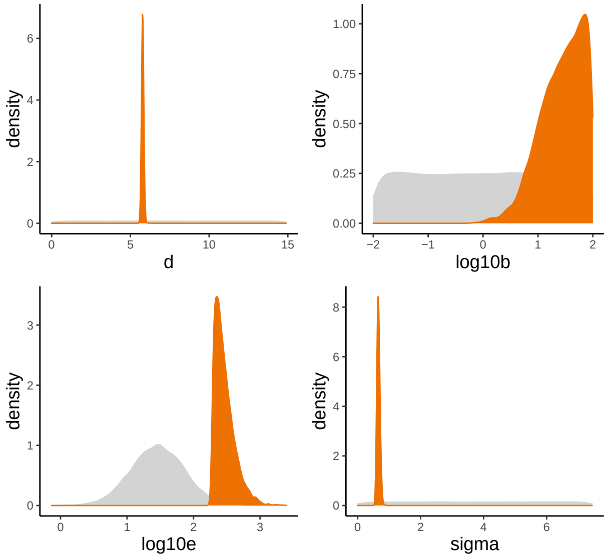

(c) Priors and posteriors

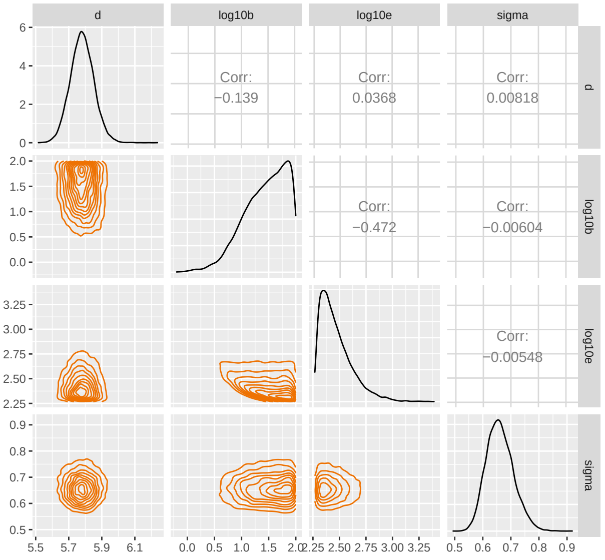

(d) Correlations between parameters

Figure 6: Dose-response curve (a), PPC (b), prior and posterior distributions (c) and correlations between parameters (d).

#### Data set: HELAN_VV_weight

Table 7: Summary of parameter estimates for HELAN_VV_weight data set

| Parameter | median | Q2.5 | Q97.5 |
|---|---|---|---|
| b | 21.936 | 1.979 | 92.641 |
| d | 2.759 | 2.690 | 2.837 |
| e | 265.261 | 190.155 | 816.097 |
| sigma | 0.310 | 0.270 | 0.361 |

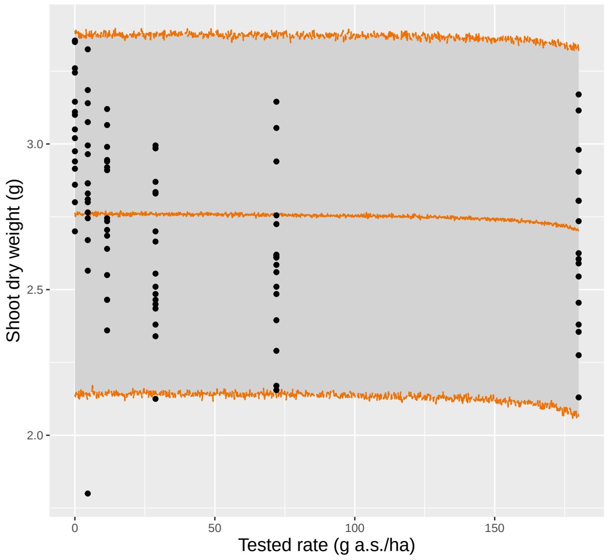

(a) Dose-response curve

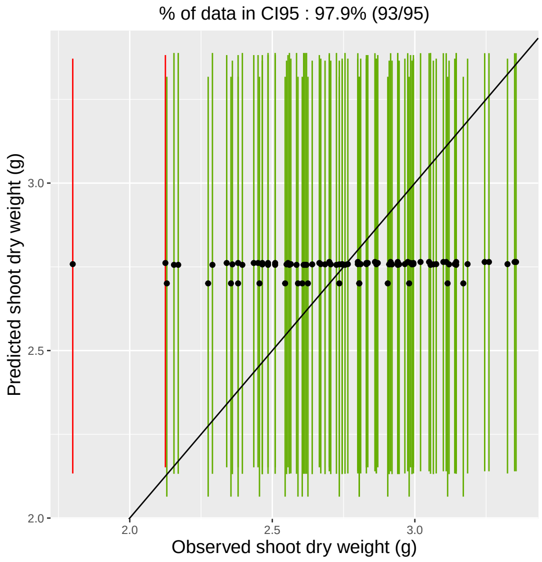

(b) Posterior predictive check (PPC)

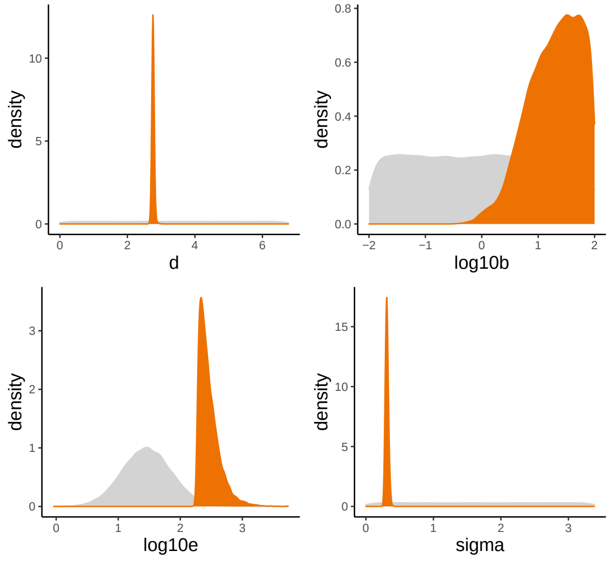

(c) Priors and posteriors

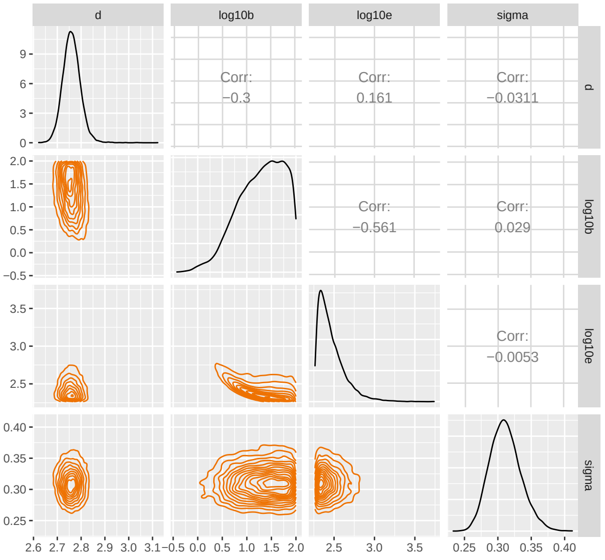

(d) Correlations between parameters

Figure 7: Dose-response curve (a), PPC (b), prior and posterior distributions (c) and correlations between parameters (d).

#### Data set: LOLPE_VV_weight

Table 8: Summary of parameter estimates for LOLPE_VV_weight data set

| Parameter | median | Q2.5 | Q97.5 |
|---|---|---|---|
| b | 30.217 | 4.185 | 94.400 |
| d | 0.253 | 0.239 | 0.267 |
| e | 266.256 | 192.719 | 665.884 |
| sigma | 0.049 | 0.041 | 0.062 |

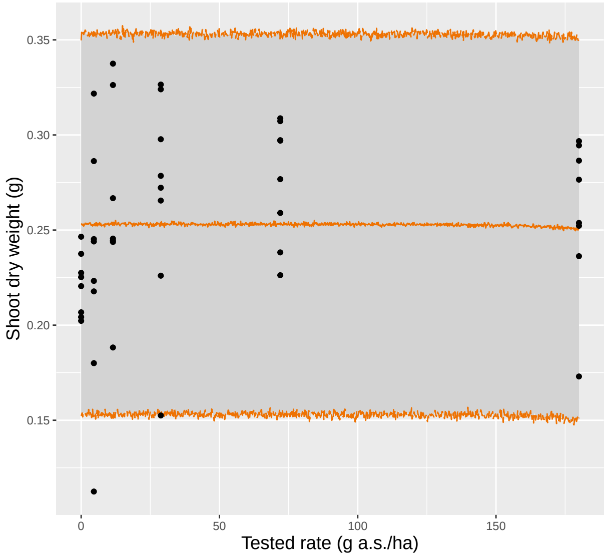

(a) Dose-response curve

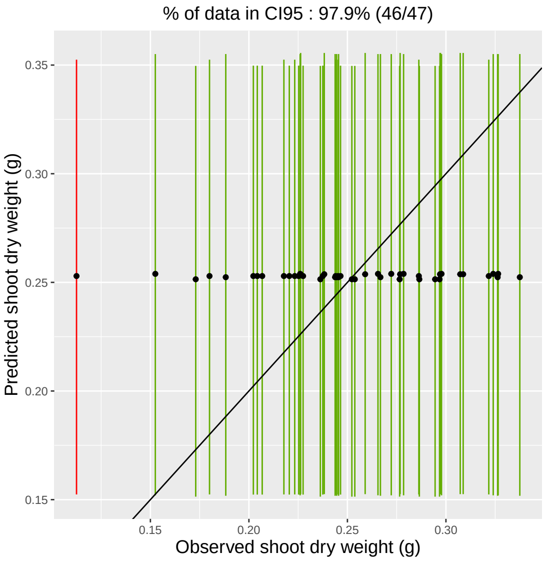

(b) Posterior predictive check (PPC)

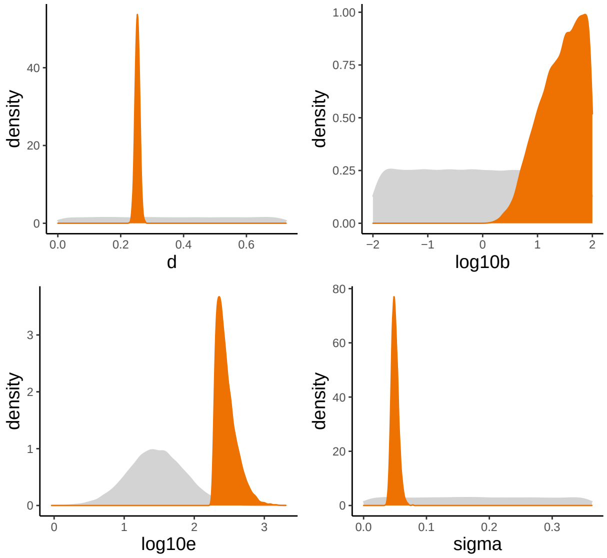

(c) Priors and posteriors

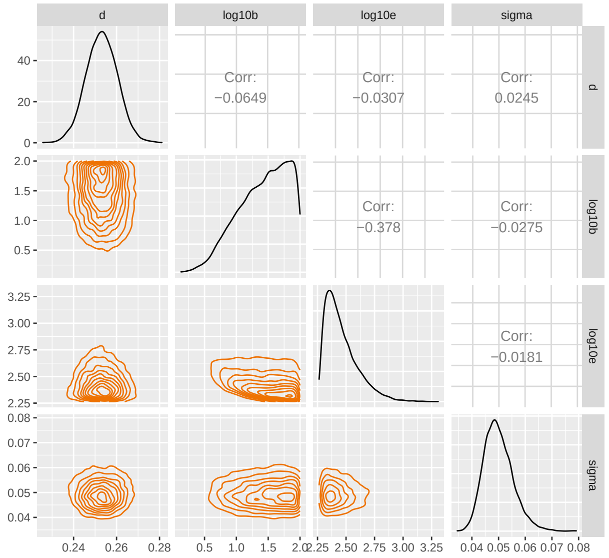

(d) Correlations between parameters

Figure 8: Dose-response curve (a), PPC (b), prior and posterior distributions (c) and correlations between parameters (d).

#### Data set: LYPES_VV_weight

Table 9: Summary of parameter estimates for LYPES_VV_weight data set

| Parameter | median | Q2.5 | Q97.5 |
|---|---|---|---|
| b | 0.812 | 0.651 | 1.020 |
| d | 6.851 | 6.553 | 7.163 |
| e | 176.247 | 141.137 | 229.857 |
| sigma | 0.732 | 0.639 | 0.852 |

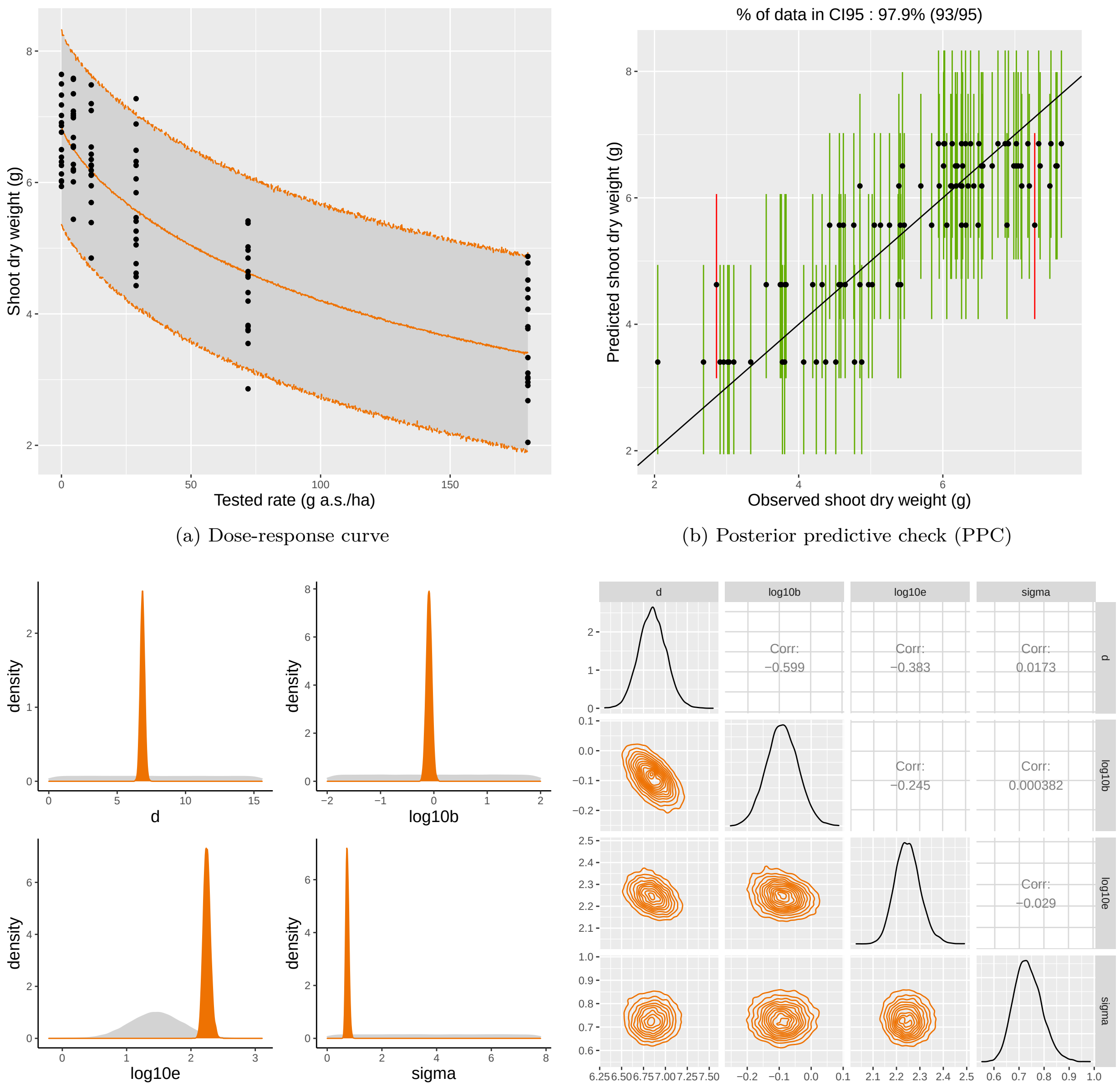

(a) Dose-response curve

(b) Posterior predictive check (PPC)

(c) Priors and posteriors

(d) Correlations between parameters

Figure 9: Dose-response curve (a), PPC (b), prior and posterior distributions (c) and correlations between parameters (d).

#### Data set: ZEAMA_VV_weight

Table 10: Summary of parameter estimates for ZEAMA_VV_weight data set

| Parameter | median | Q2.5 | Q97.5 |
|---|---|---|---|
| b | 22.991 | 1.864 | 92.923 |
| d | 7.636 | 7.365 | 7.947 |
| e | 264.931 | 190.651 | 729.949 |
| sigma | 1.298 | 1.131 | 1.510 |

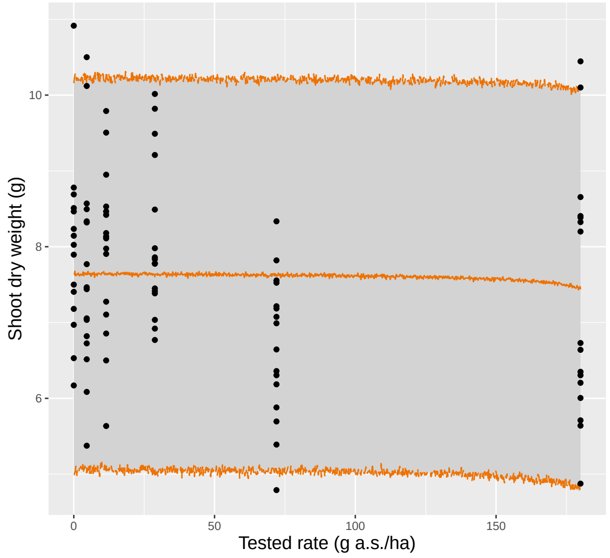

(a) Dose-response curve

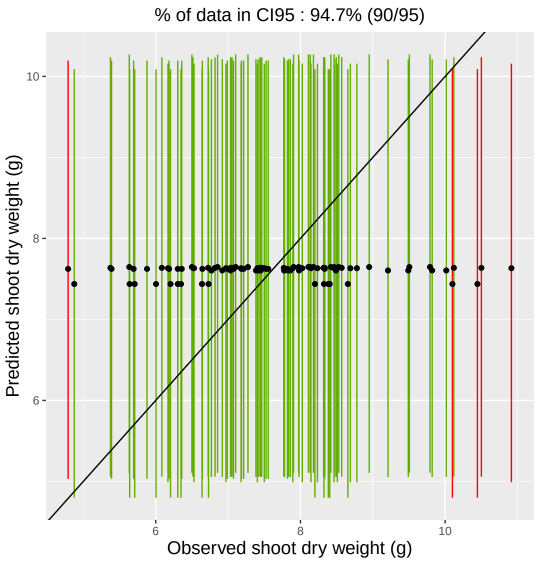

(b) Posterior predictive check (PPC)

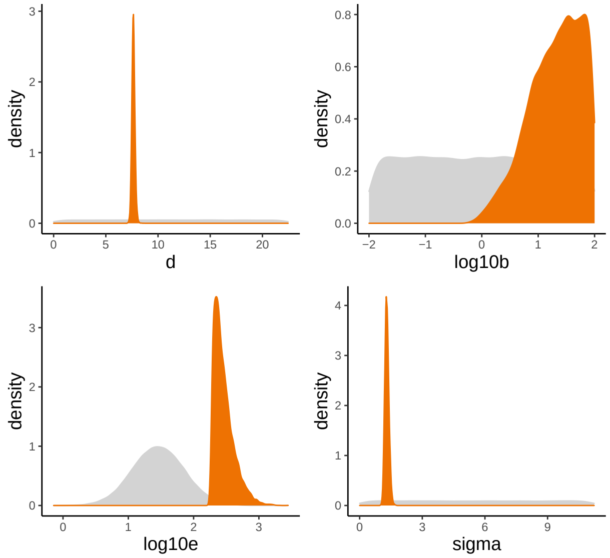

(c) Priors and posteriors

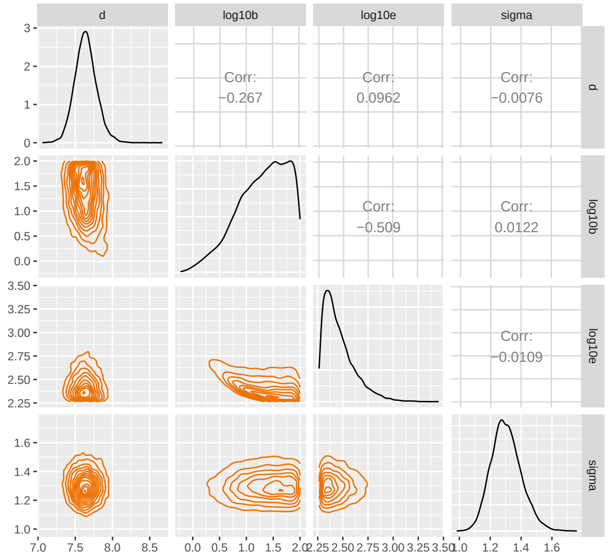

(d) Correlations between parameters

Figure 10: Dose-response curve (a), PPC (b), prior and posterior distributions (c) and correlations between parameters (d).
