## Supplementary material for "How to account for the uncertainty from standard toxicity tests in species sensitivity distributions: an example in non-target plants": Archive with reports on studies: report_SE_emergence.pdf

### Dose-response analyses

Study 3

Seedling Emergence test - emergence endpoint

25 June 2020

Contact:

---

This is a report which provides results on all performed dose-response analyses for the emergence endpoint of the Seedling Emergence test for study 3.

---

#### Data set: ALLCE_SE_emergence

Table 1: Summary of parameter estimates for ALLCE_SE_emergence data set

| Parameter | median | Q2.5 | Q97.5 |
|---|---:|---:|---:|
| b | 6.208 | 0.404 | 75.086 |
| d | 0.793 | 0.704 | 0.936 |
| e | 1742.724 | 1386.898 | 4285.237 |

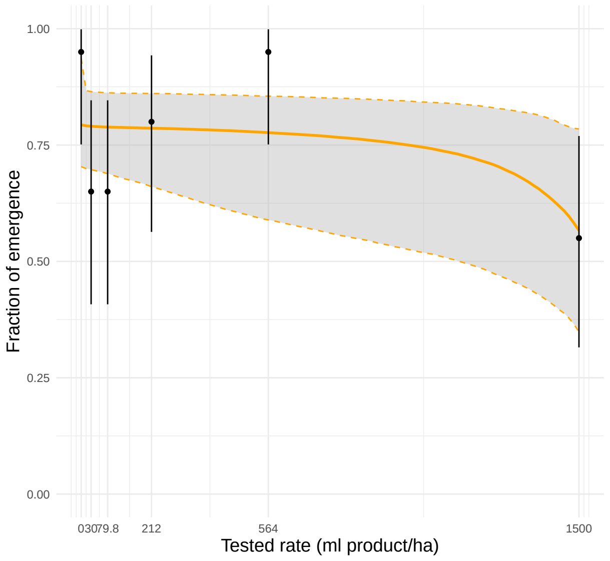

(a) Dose-response curve

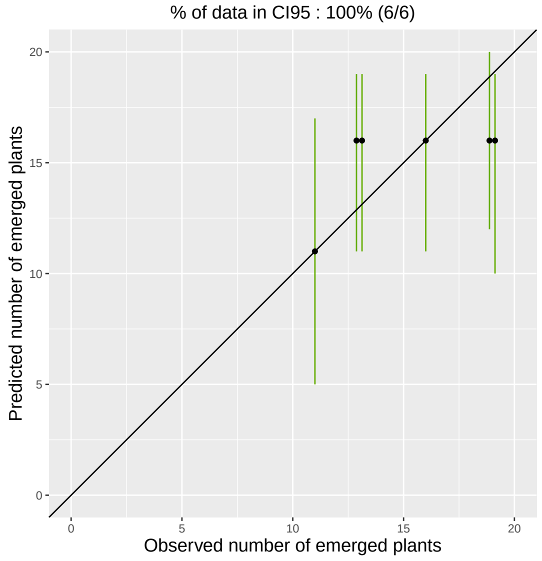

(b) Posterior predictive check (PPC)

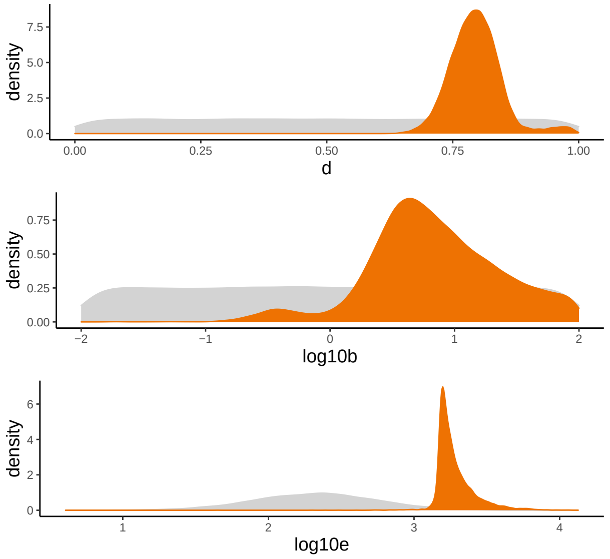

(c) Priors and posteriors

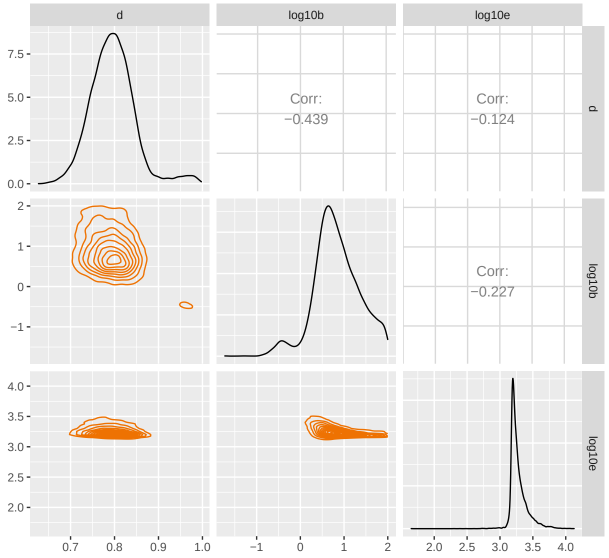

(d) Correlations between parameters

Figure 1: Dose-response curve (a), PPC (b), prior and posterior distributions (c) and correlations between parameters (d).

#### Data set: BEAVA_SE_emergence

Table 2: Summary of parameter estimates for BEAVA_SE_emergence data set

| Parameter | median | Q2.5 | Q97.5 |
|---|---:|---:|---:|
| b | 31.797 | 4.206 | 94.386 |
| d | 0.962 | 0.918 | 0.986 |
| e | 2314.562 | 1618.871 | 6004.500 |

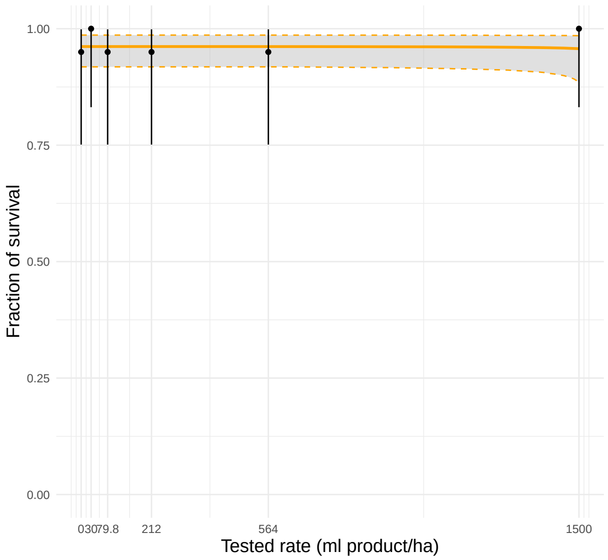

(a) Dose-response curve

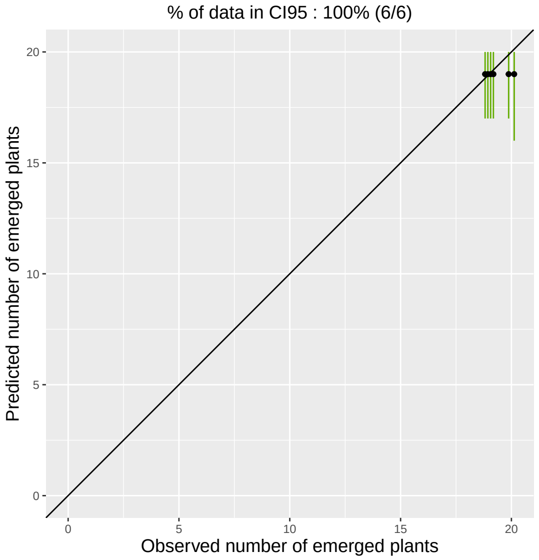

(b) Posterior predictive check (PPC)

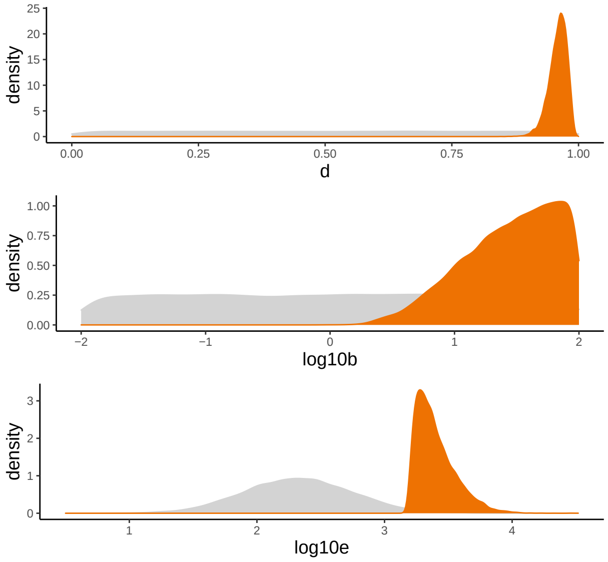

(c) Priors and posteriors

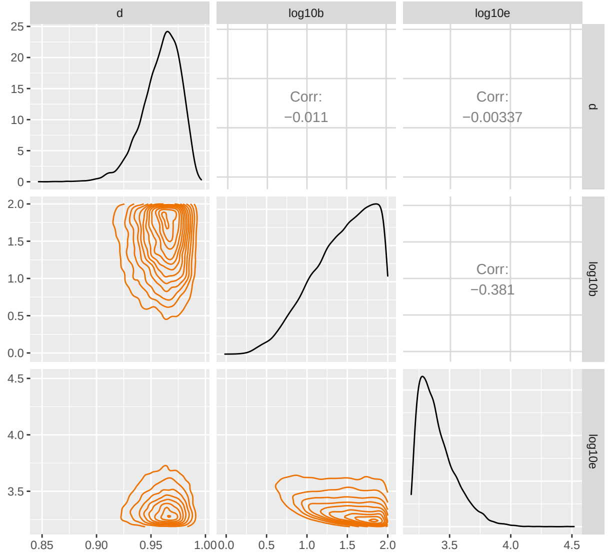

(d) Correlations between parameters

Figure 2: Dose-response curve (a), PPC (b), prior and posterior distributions (c) and correlations between parameters (d).

#### Data set: BRSNW_SE_emergence

Table 3: Summary of parameter estimates for BRSNW_SE_emergence data set

| Parameter | median | Q2.5 | Q97.5 |
|---|---|---|---|
| b | 19.929 | 1.788 | 92.748 |
| d | 0.942 | 0.887 | 0.978 |
| e | 2218.620 | 1583.885 | 5925.315 |

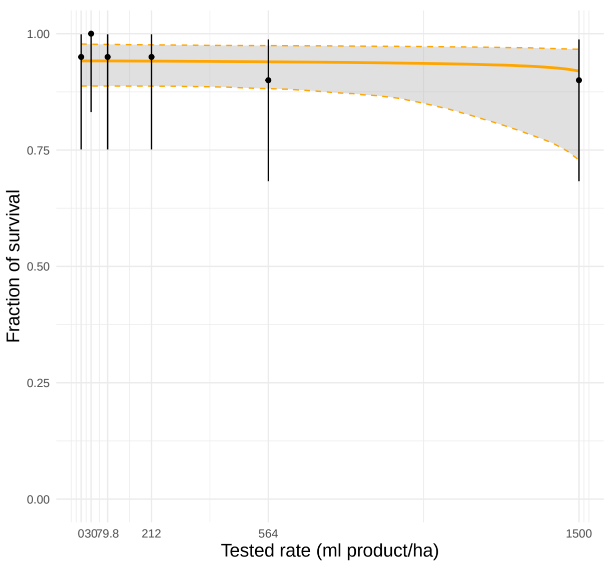

(a) Dose-response curve

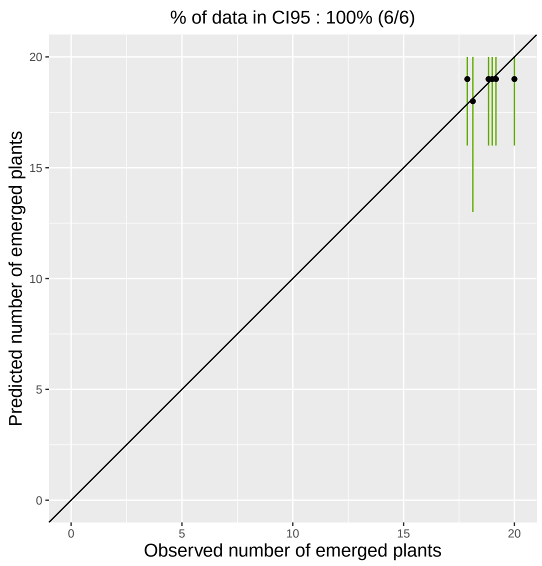

(b) Posterior predictive check (PPC)

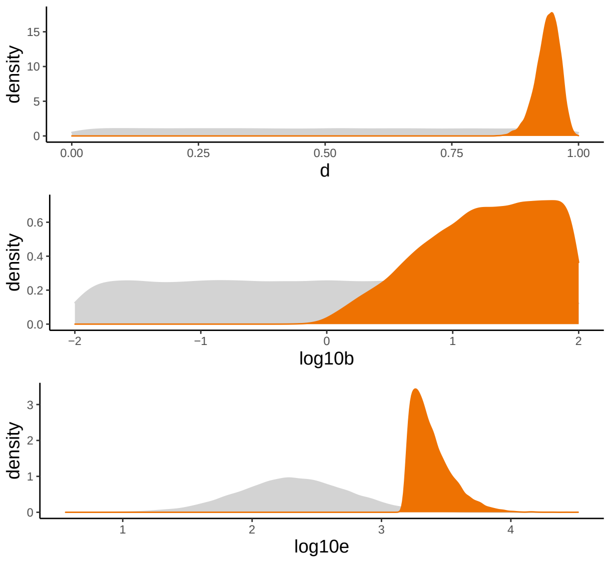

(c) Priors and posteriors

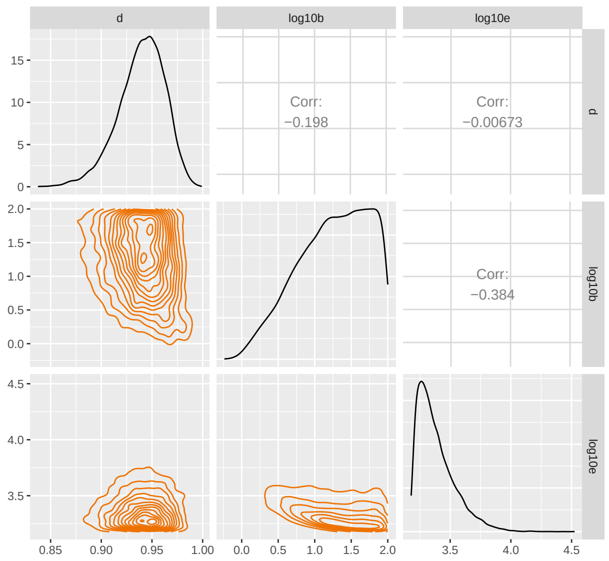

(d) Correlations between parameters

Figure 3: Dose-response curve (a), PPC (b), prior and posterior distributions (c) and correlations between parameters (d).

### Data set: CUMSA_SE_emergence

Table 4: Summary of parameter estimates for CUMSA_SE_emergence data set

| Parameter | median | Q2.5 | Q97.5 |
|---|---|---|---|
| b | 28.851 | 3.342 | 94.485 |
| d | 0.905 | 0.845 | 0.948 |
| e | 2289.370 | 1605.898 | 6129.611 |

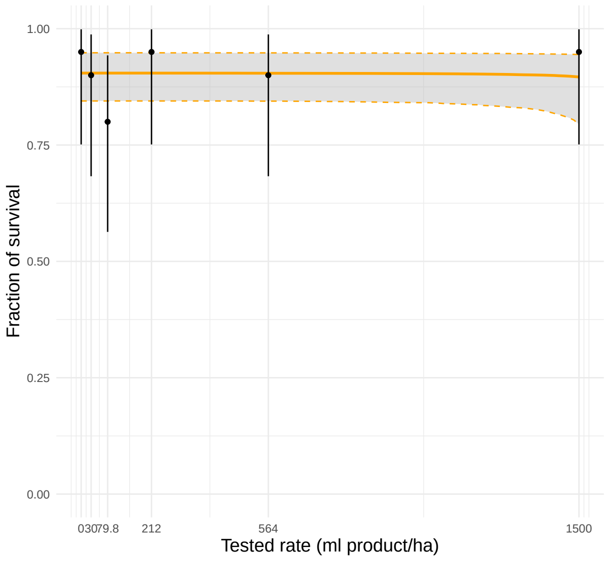

(a) Dose-response curve

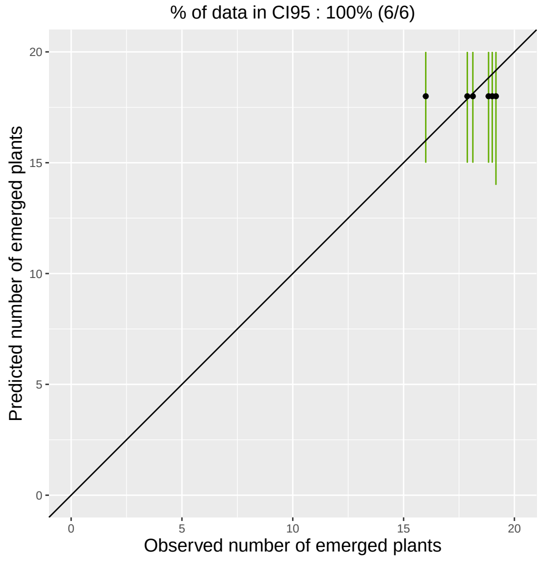

(b) Posterior predictive check (PPC)

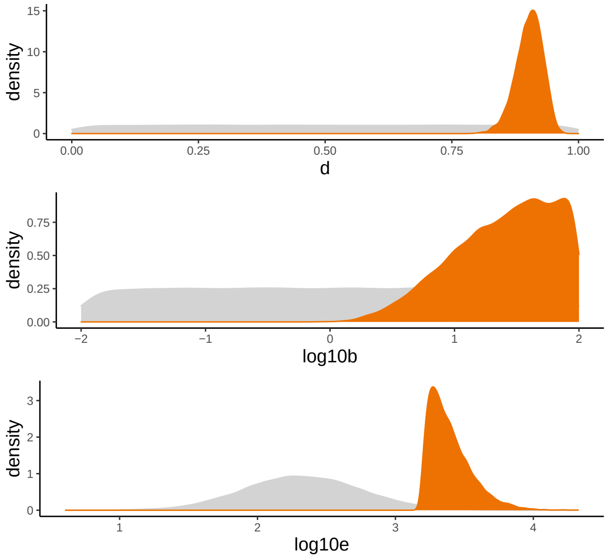

(c) Priors and posteriors

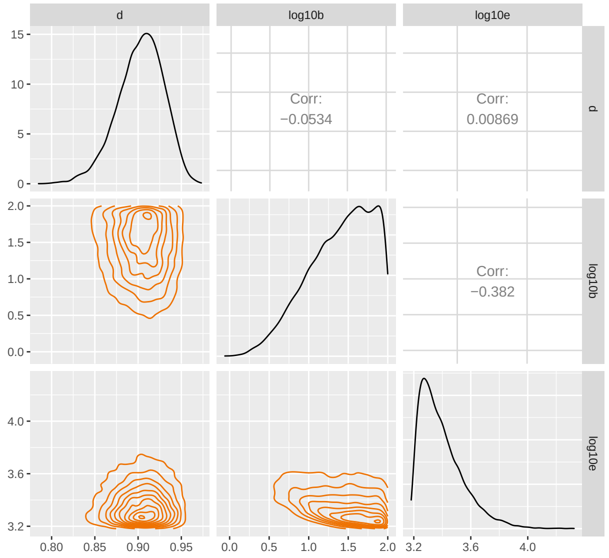

(d) Correlations between parameters

Figure 4: Dose-response curve (a), PPC (b), prior and posterior distributions (c) and correlations between parameters (d).

#### Data set: FAGES_SE_emergence

Table 5: Summary of parameter estimates for FAGES_SE_emergence data set

| Parameter | median | Q2.5 | Q97.5 |
|---|---|---|---|
| b | 27.233 | 2.658 | 94.217 |
| d | 0.864 | 0.798 | 0.918 |
| e | 2269.224 | 1600.636 | 6043.838 |

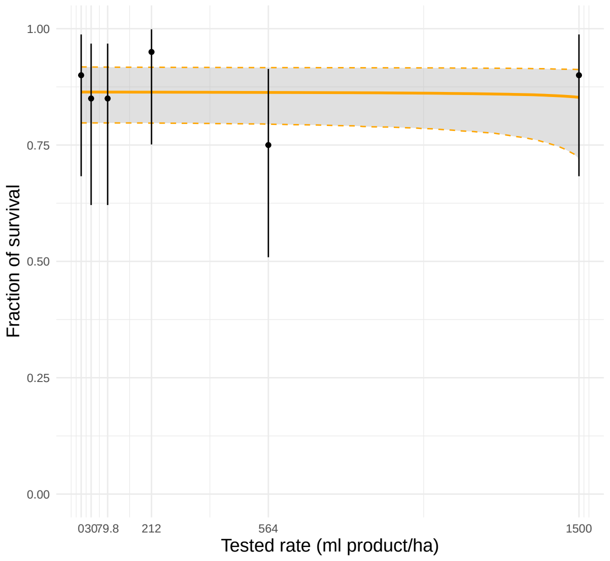

(a) Dose-response curve

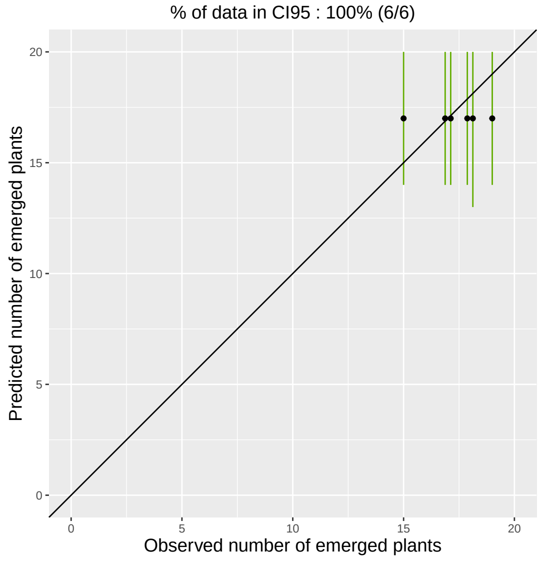

(b) Posterior predictive check (PPC)

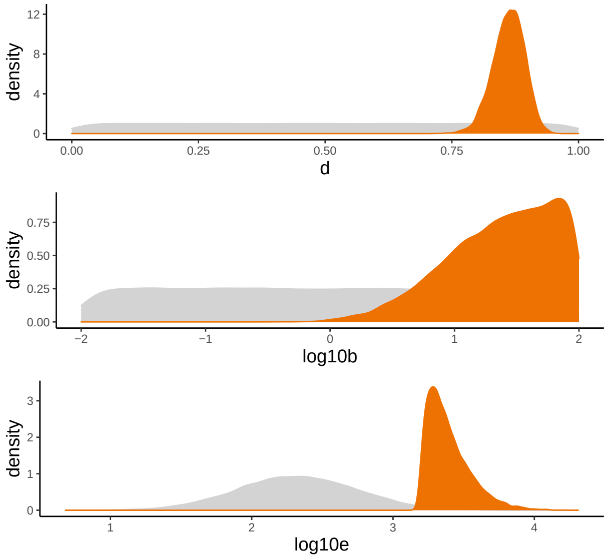

(c) Priors and posteriors

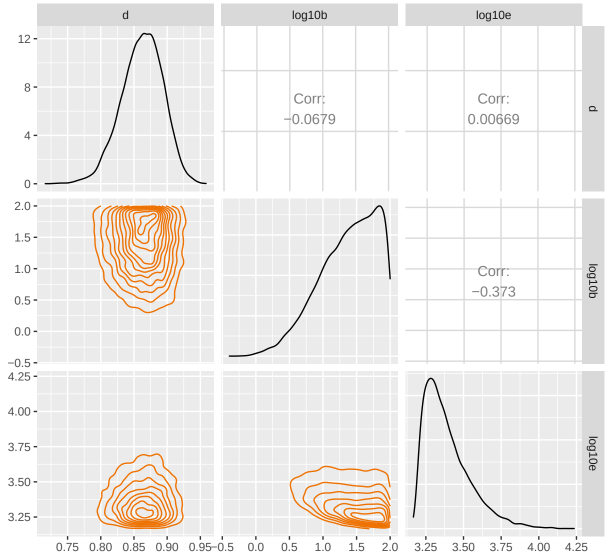

(d) Correlations between parameters

Figure 5: Dose-response curve (a), PPC (b), prior and posterior distributions (c) and correlations between parameters (d).

### Data set: GLXMA_SE_emergence

Table 6: Summary of parameter estimates (parameter d is set to 1) for GLXMA_SE_emergence data set

| Parameter | median | Q2.5 | Q97.5 |
|---|---|---|---|
| b | 1.035 | 0.65 | 1.636 |
| e | 5292.275 | 2475.91 | 15515.520 |

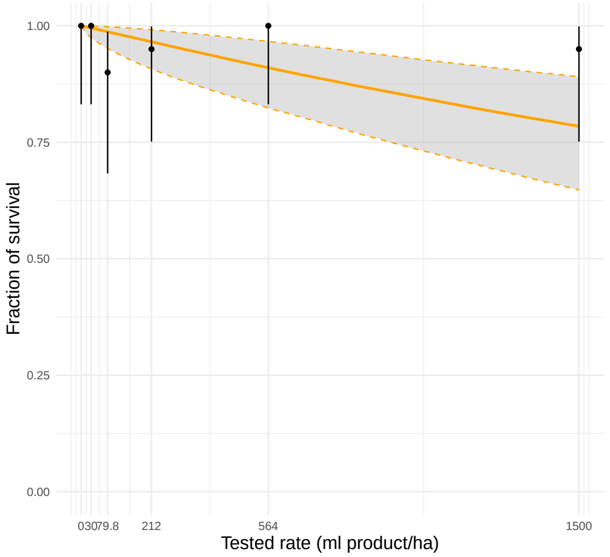

(a) Dose-response curve

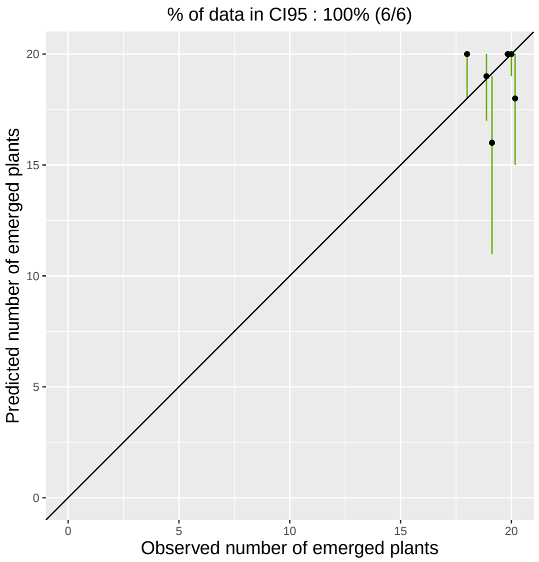

(b) Posterior predictive check (PPC)

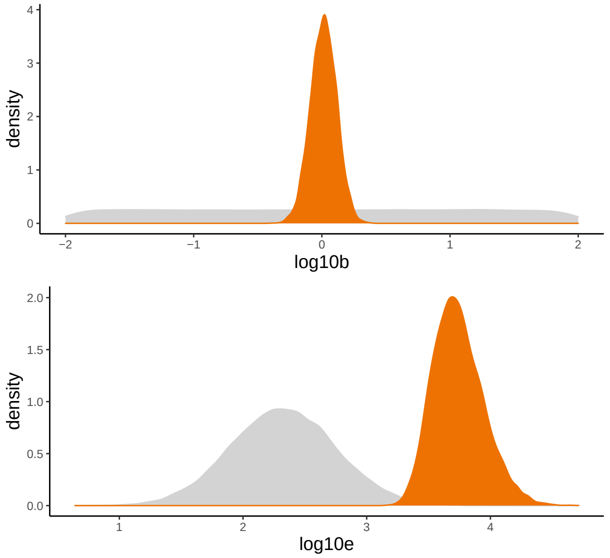

(c) Priors and posteriors

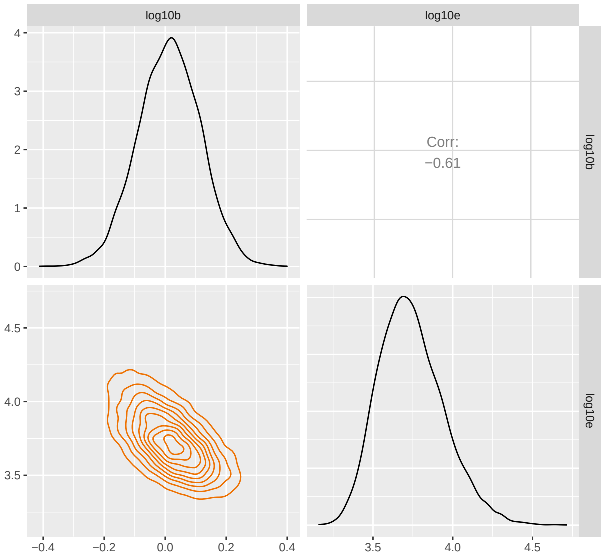

(d) Correlations between parameters

Figure 6: Dose-response curve (a), PPC (b), prior and posterior distributions (c) and correlations between parameters (d).

#### Data set: LOLPE_SE_emergence

Table 7: Summary of parameter estimates for LOLPE_SE_emergence data set

| Parameter | median | Q2.5 | Q97.5 |
|---|---:|---:|---:|
| b | 24.548 | 2.401 | 93.117 |
| d | 0.849 | 0.779 | 0.906 |
| e | 2250.291 | 1589.317 | 5903.808 |

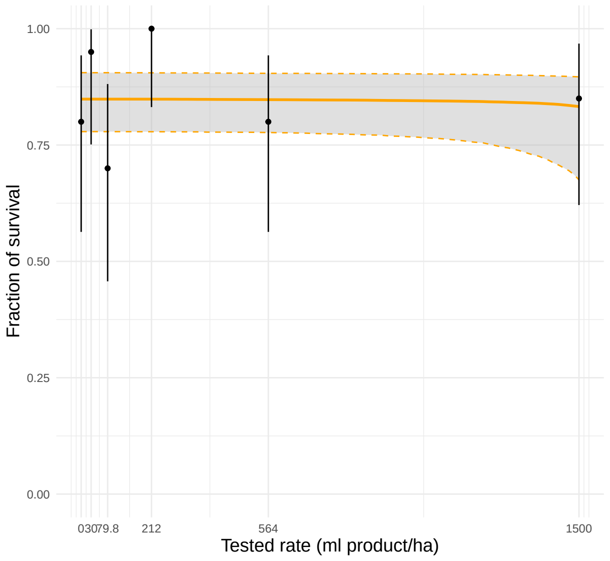

(a) Dose-response curve

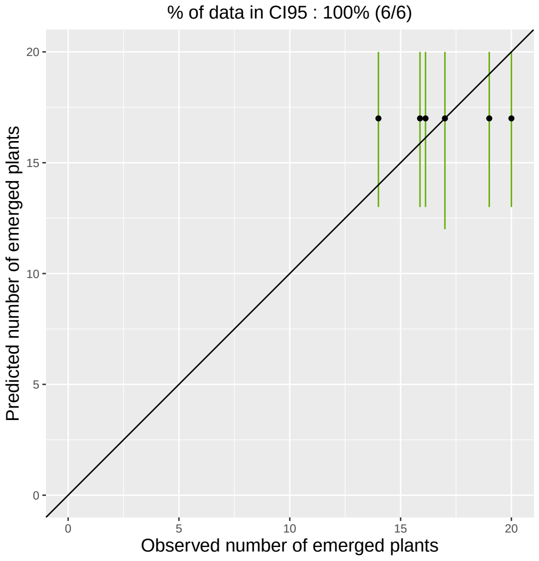

(b) Posterior predictive check (PPC)

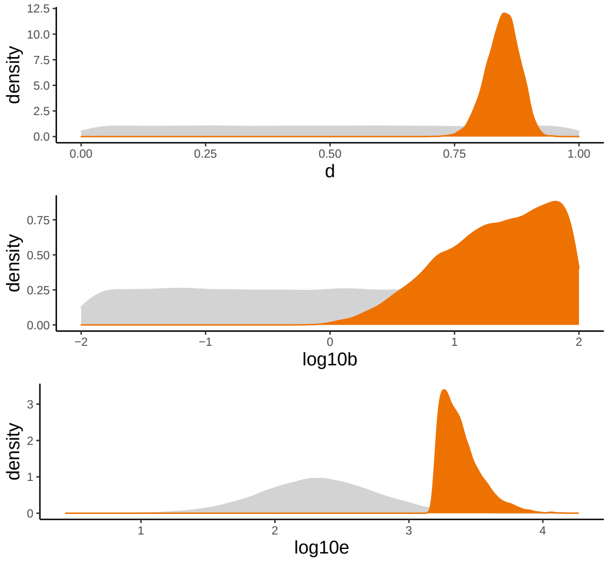

(c) Priors and posteriors

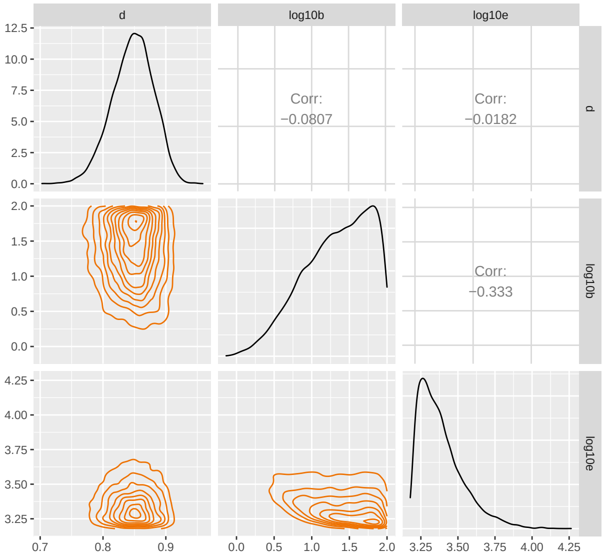

(d) Correlations between parameters

Figure 7: Dose-response curve (a), PPC (b), prior and posterior distributions (c) and correlations between parameters (d).

#### Data set: LYPES_SE_emergence

Table 8: Summary of parameter estimates for LYPES_SE_emergence data set

| Parameter | median | Q2.5 | Q97.5 |
|---|---|---|---|
| b | 22.348 | 2.289 | 93.094 |
| d | 0.932 | 0.876 | 0.970 |
| e | 2229.507 | 1585.341 | 5793.903 |

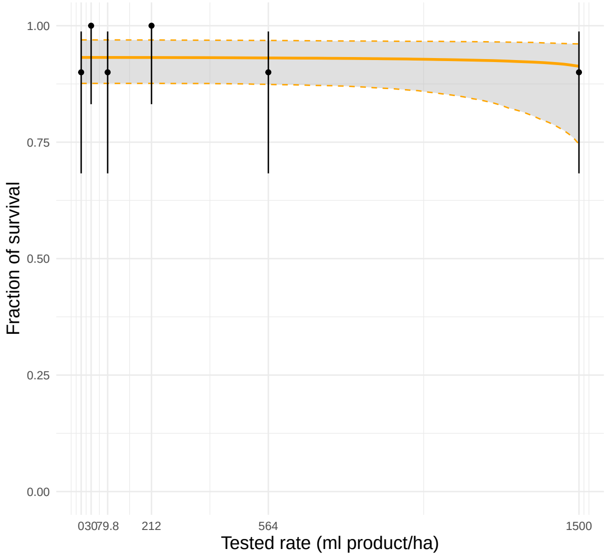

(a) Dose-response curve

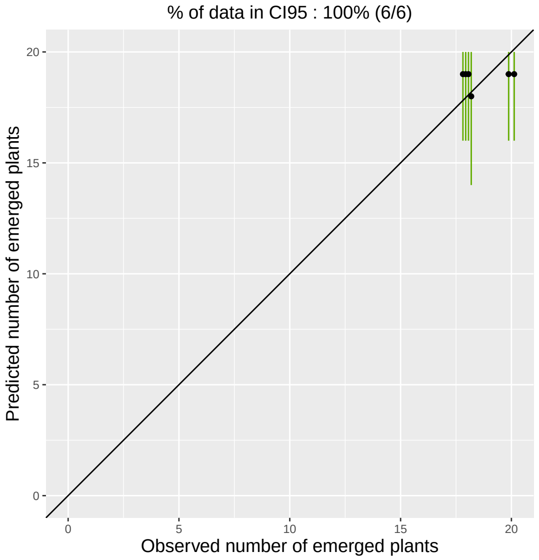

(b) Posterior predictive check (PPC)

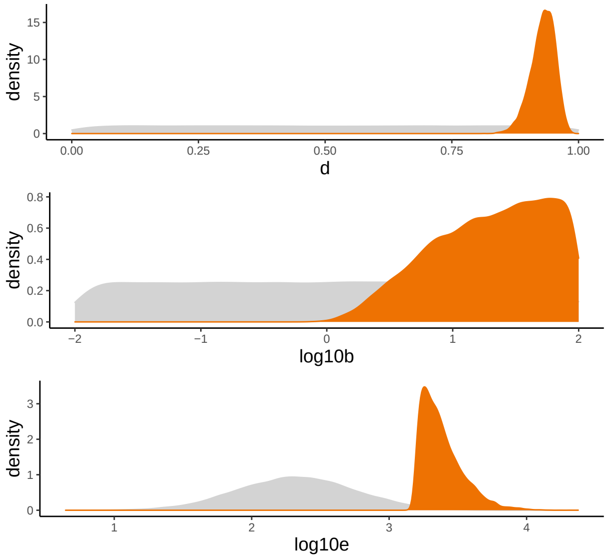

(c) Priors and posteriors

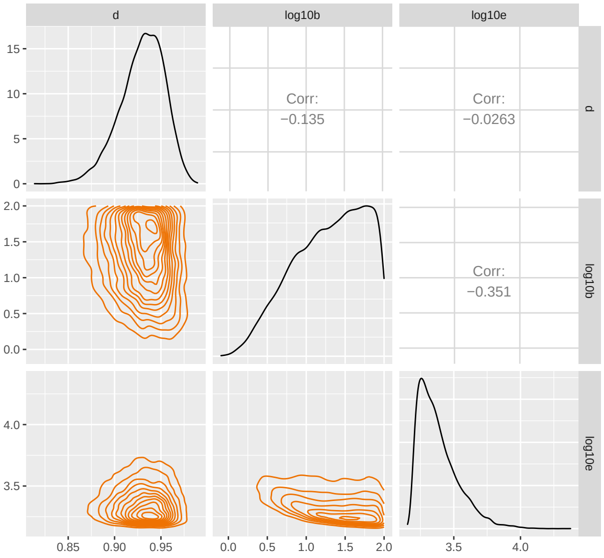

(d) Correlations between parameters

Figure 8: Dose-response curve (a), PPC (b), prior and posterior distributions (c) and correlations between parameters (d).

#### Data set: TRZAW_SE_emergence

Table 9: Summary of parameter estimates for TRZAW_SE_emergence data set

| Parameter | median | Q2.5 | Q97.5 |
|---|---:|---:|---:|
| b | 25.690 | 3.213 | 93.378 |
| d | 0.963 | 0.918 | 0.988 |
| e | 2257.608 | 1595.494 | 6090.512 |

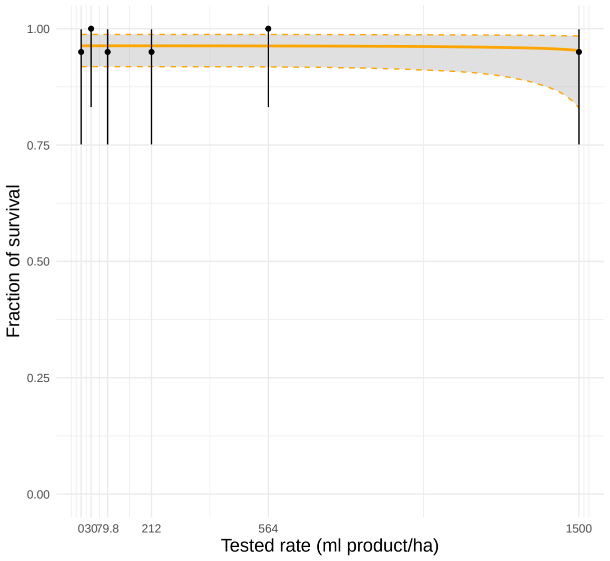

(a) Dose-response curve

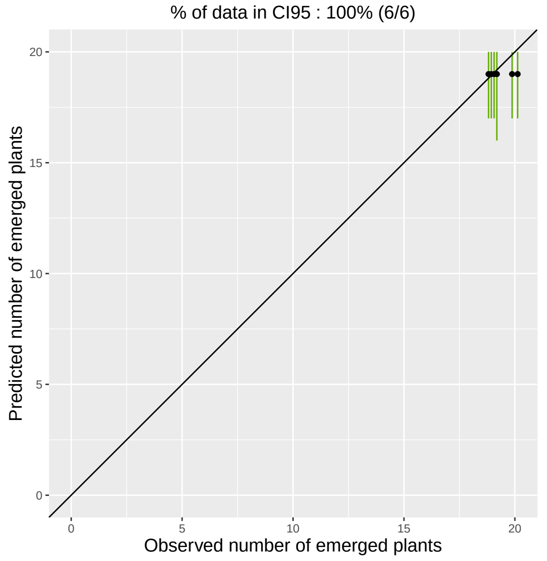

(b) Posterior predictive check (PPC)

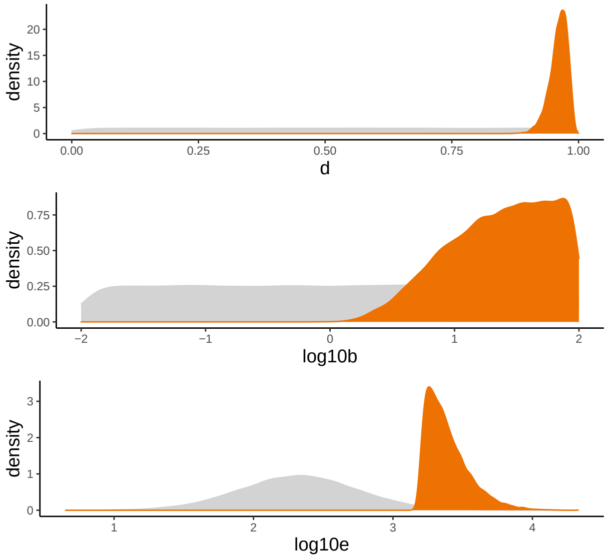

(c) Priors and posteriors

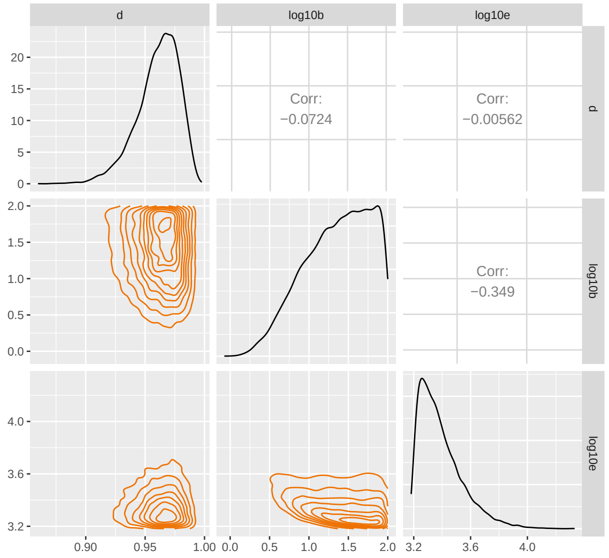

(d) Correlations between parameters

Figure 9: Dose-response curve (a), PPC (b), prior and posterior distributions (c) and correlations between parameters (d).

#### Data set: ZEAMA_SE_emergence

Table 10: Summary of parameter estimates (parameter d is set to 1) for ZEAMA_SE_emergence data set

| Parameter | median | Q2.5 | Q97.5 |
|---|---:|---:|---:|
| b | 1.400 | 0.842 | 2.366 |
| e | 4389.401 | 2291.687 | 11578.225 |

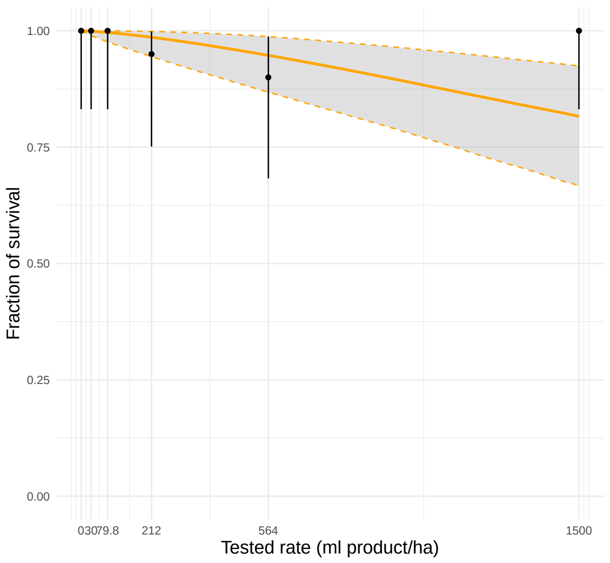

(a) Dose-response curve

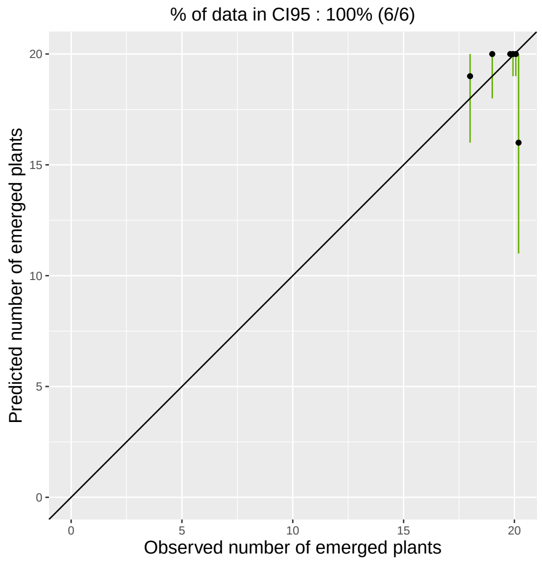

(b) Posterior predictive check (PPC)

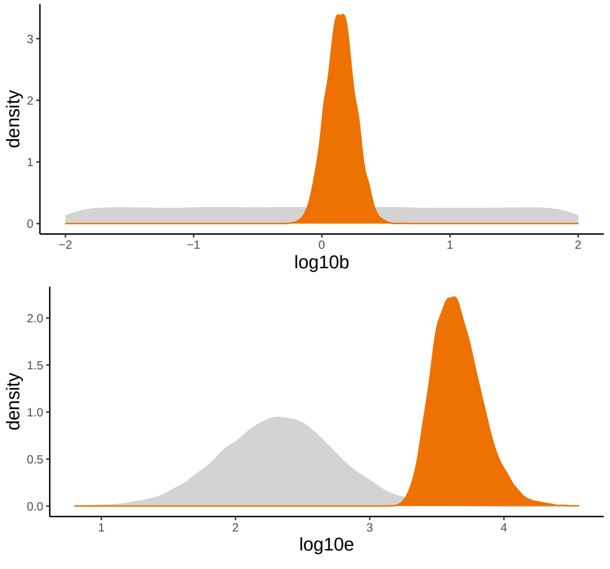

(c) Priors and posteriors

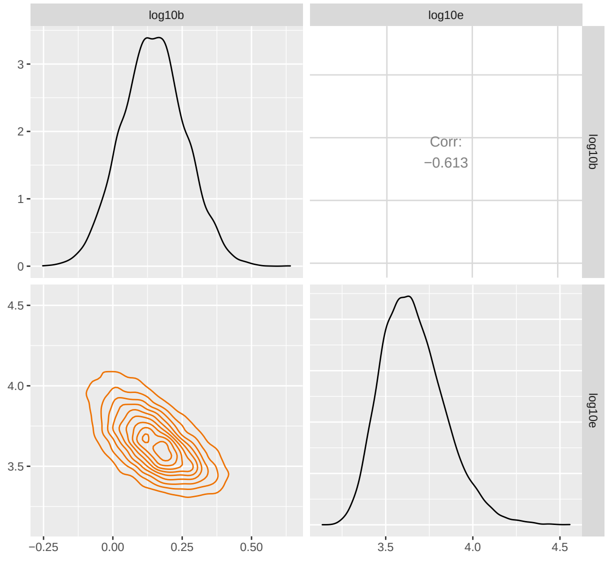

(d) Correlations between parameters

Figure 10: Dose-response curve (a), PPC (b), prior and posterior distributions (c) and correlations between parameters (d).
