## Supplementary material for "How to account for the uncertainty from standard toxicity tests in species sensitivity distributions: an example in non-target plants": Archive with reports on studies: report_SE_survival.pdf

### Dose-response analyses

Study 3

Seedling Emergence test - survival endpoint

25 June 2020

Contact:

---

This is a report which provides results on all performed dose-response analyses for the survival endpoint of the Seedling Emergence test for study 3.

---

#### Data set: ALLCE_SE_survival

Table 1: Summary of parameter estimates (parameter d is set to 1) for ALLCE_SE_survival data set

| Parameter | median | Q2.5 | Q97.5 |
|---|---|---|---|
| b | 29.526 | 3.697 | 95.007 |
| e | 2272.116 | 1603.563 | 6155.935 |

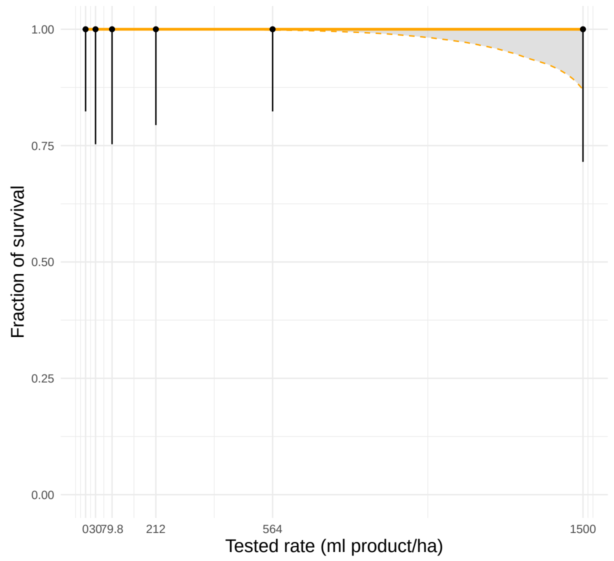

(a) Dose-response curve

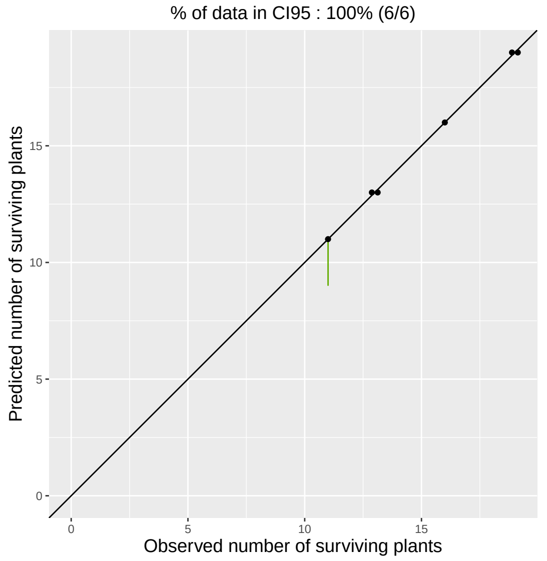

(b) Posterior predictive check (PPC)

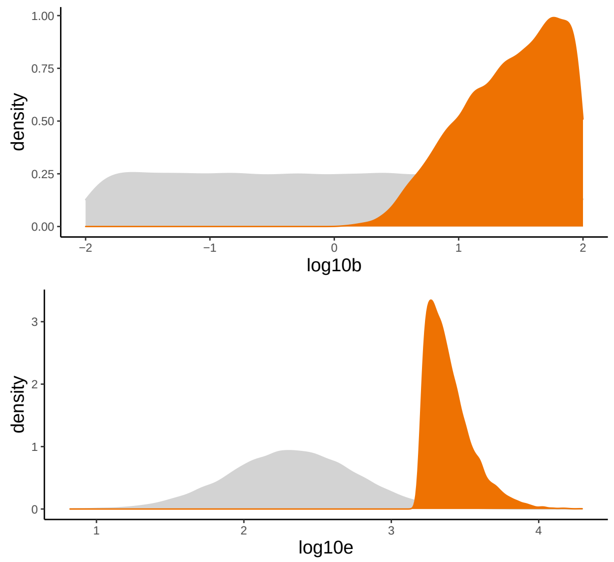

(c) Priors and posteriors

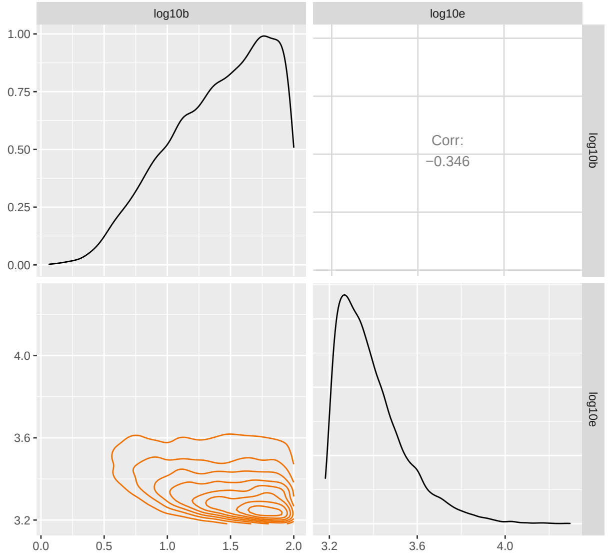

(d) Correlations between parameters

Figure 1: Dose-response curve (a), PPC (b), prior and posterior distributions (c) and correlations between parameters (d).

#### Data set: BEAVA_SE_survival

Table 2: Summary of parameter estimates (parameter d is set to 1) for BEAVA_SE_survival data set

| Parameter | median | Q2.5 | Q97.5 |
|---|---|---|---|
| b | 31.338 | 4.181 | 95.024 |
| e | 2311.868 | 1620.401 | 5999.744 |

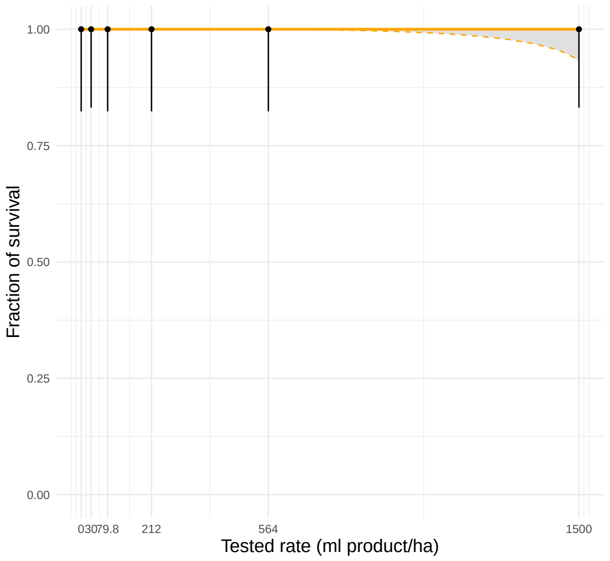

(a) Dose-response curve

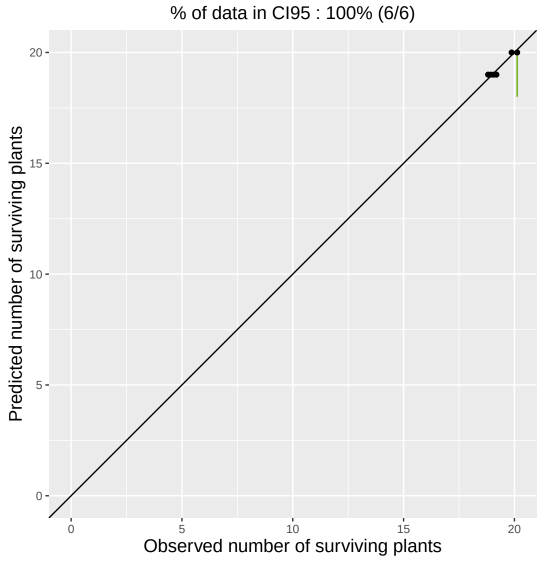

(b) Posterior predictive check (PPC)

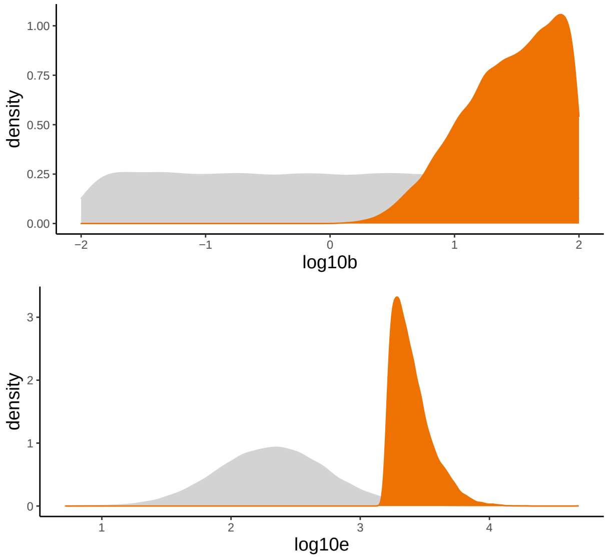

(c) Priors and posteriors

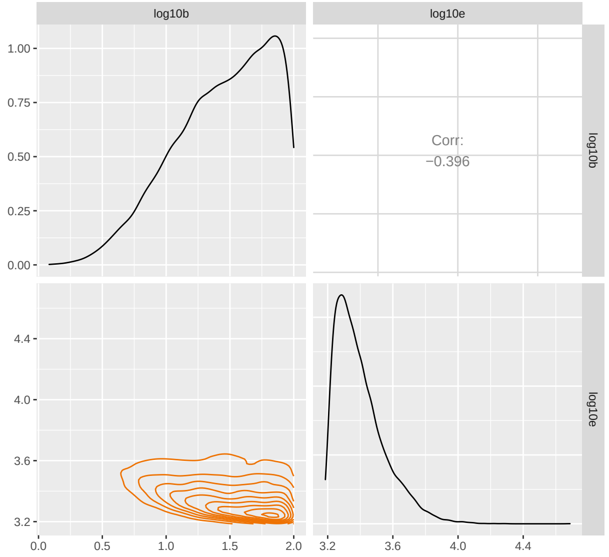

(d) Correlations between parameters

Figure 2: Dose-response curve (a), PPC (b), prior and posterior distributions (c) and correlations between parameters (d).

#### Data set: BRSNW_SE_survival

Table 3: Summary of parameter estimates (parameter d is set to 1) for BRSNW_SE_survival data set

| Parameter | median | Q2.5 | Q97.5 |
|---|---|---|---|
| b | 7.022 | 2.656 | 57.657 |
| e | 1377.016 | 1073.671 | 1557.188 |

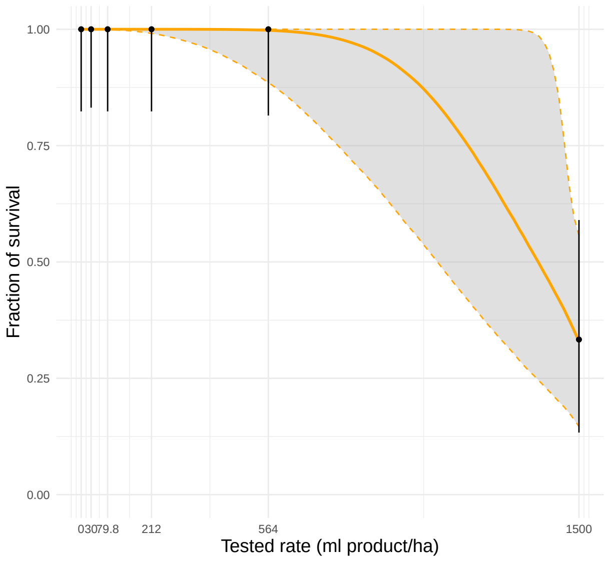

(a) Dose-response curve

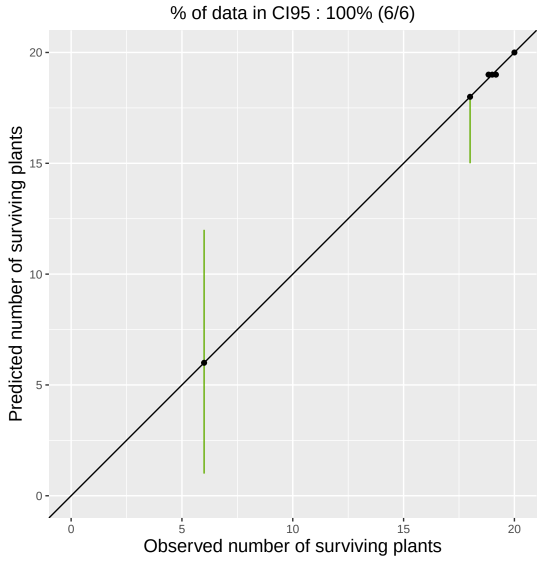

(b) Posterior predictive check (PPC)

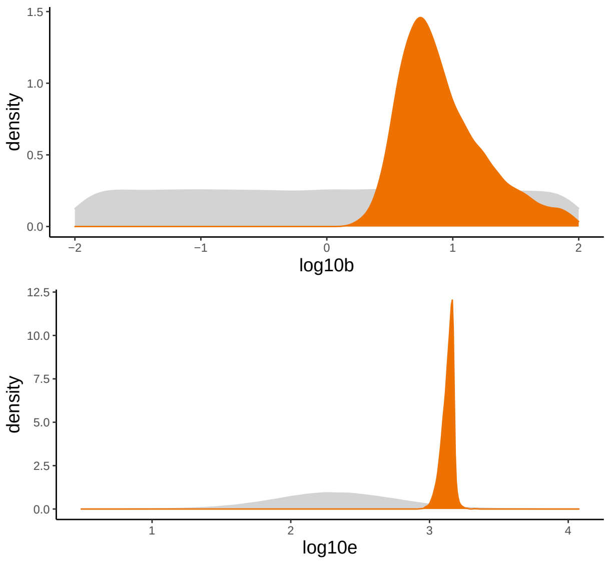

(c) Priors and posteriors

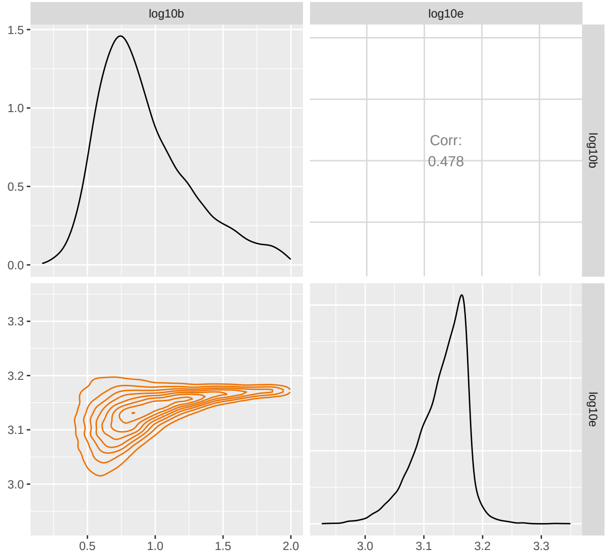

(d) Correlations between parameters

Figure 3: Dose-response curve (a), PPC (b), prior and posterior distributions (c) and correlations between parameters (d).

#### Data set: CUMSA_SE_survival

Table 4: Summary of parameter estimates (parameter d is set to 1) for CUMSA_SE_survival data set

| Parameter | median | Q2.5 | Q97.5 |
|---|---|---|---|
| b | 4.430 | 2.516 | 7.529 |
| e | 1024.014 | 796.974 | 1279.489 |

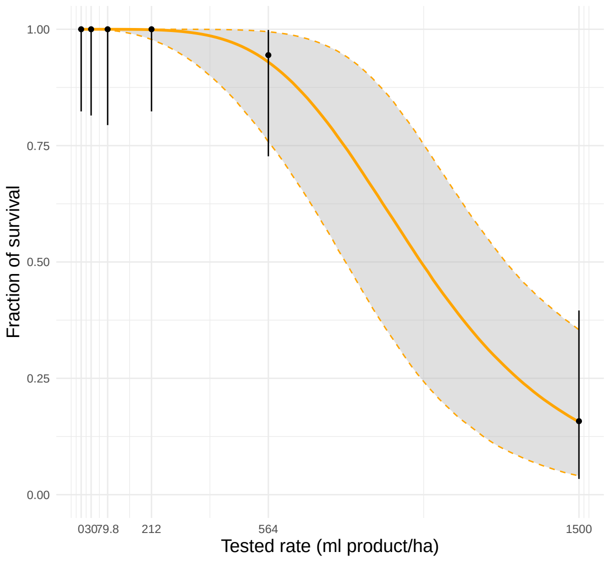

(a) Dose-response curve

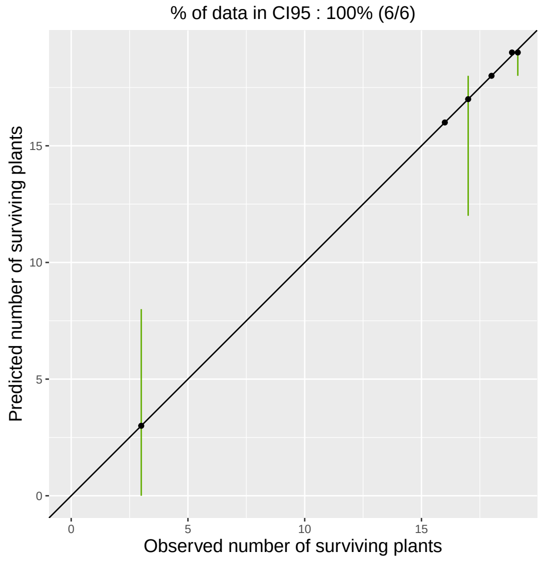

(b) Posterior predictive check (PPC)

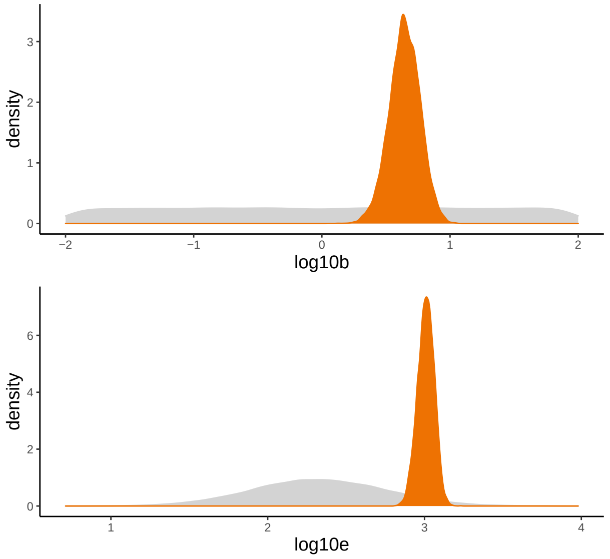

(c) Priors and posteriors

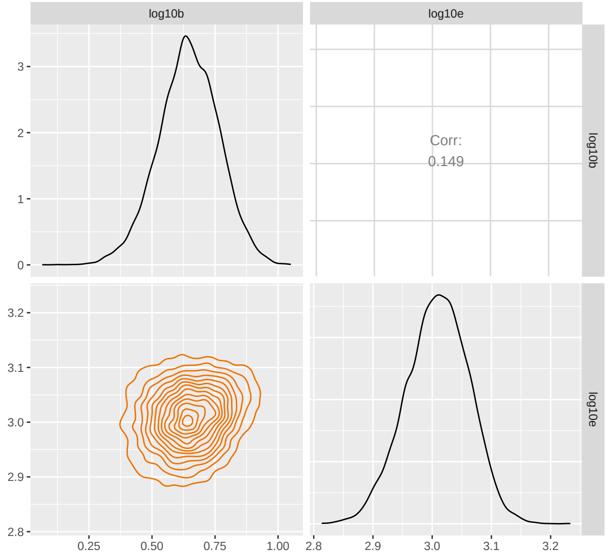

(d) Correlations between parameters

Figure 4: Dose-response curve (a), PPC (b), prior and posterior distributions (c) and correlations between parameters (d).

#### Data set: FAGES_SE_survival

Table 5: Summary of parameter estimates (parameter d is set to 1) for FAGES_SE_survival data set

| Parameter | median | Q2.5 | Q97.5 |
|---|---|---|---|
| b | 6.562 | 1.734 | 61.971 |
| e | 1870.314 | 1532.043 | 3884.872 |

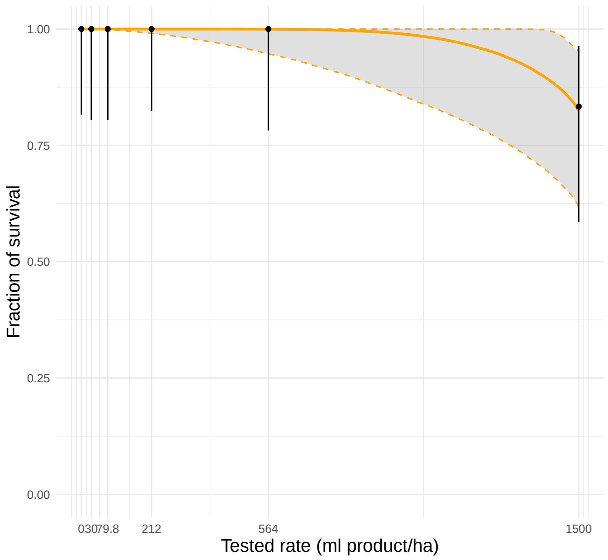

(a) Dose-response curve

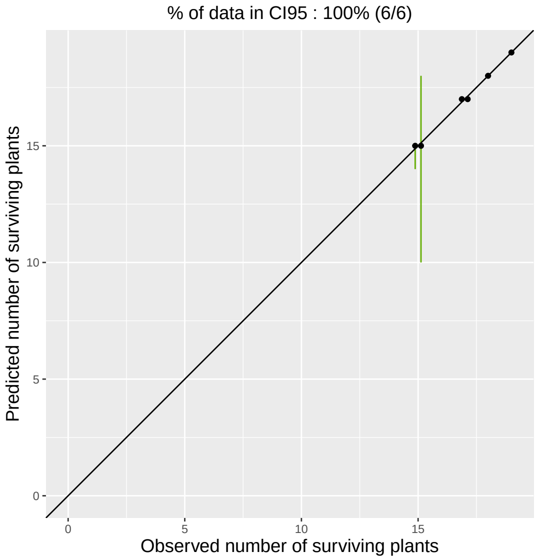

(b) Posterior predictive check (PPC)

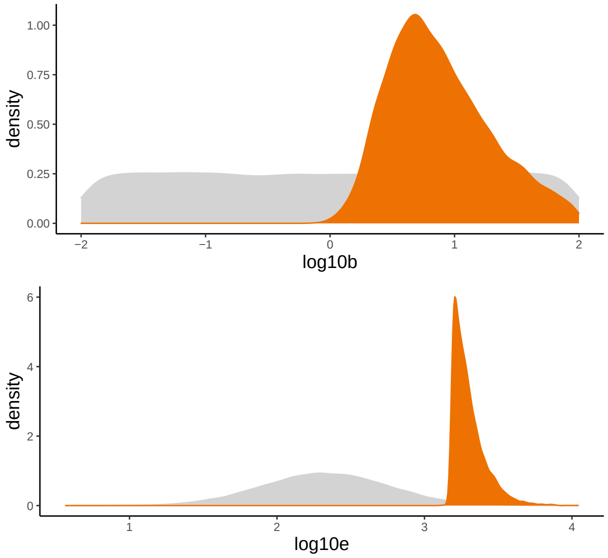

(c) Priors and posteriors

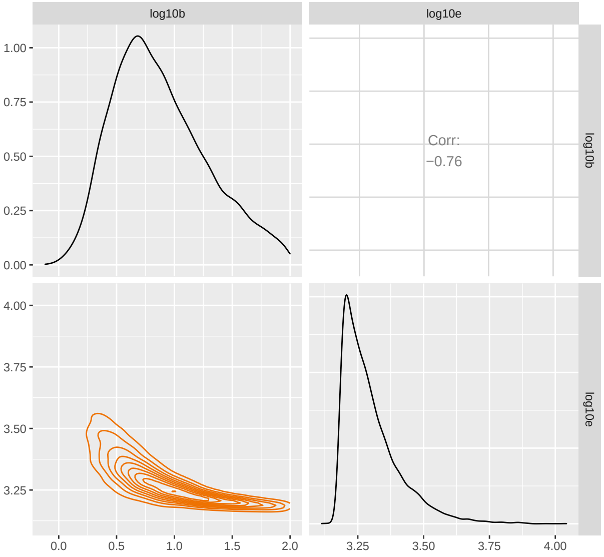

(d) Correlations between parameters

Figure 5: Dose-response curve (a), PPC (b), prior and posterior distributions (c) and correlations between parameters (d).

#### Data set: GLXMA_SE_survival

Table 6: Summary of parameter estimates (parameter d is set to 1) for GLXMA_SE_survival data set

| Parameter | median | Q2.5 | Q97.5 |
|---|---:|---:|---:|
| b | 31.926 | 4.190 | 95.159 |
| e | 2318.128 | 1618.988 | 6272.430 |

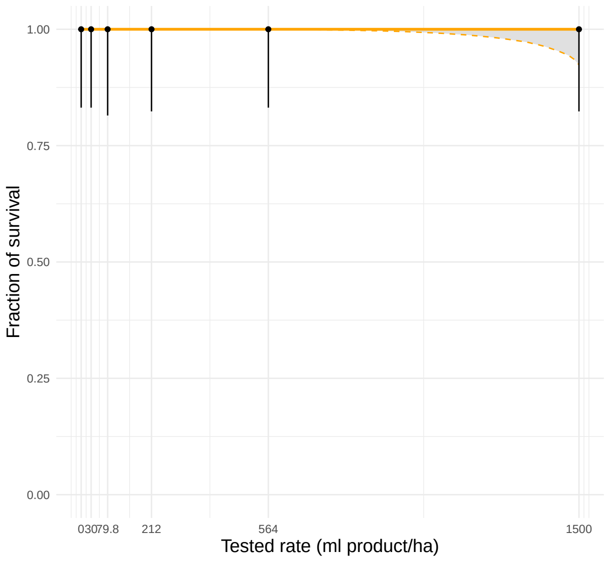

(a) Dose-response curve

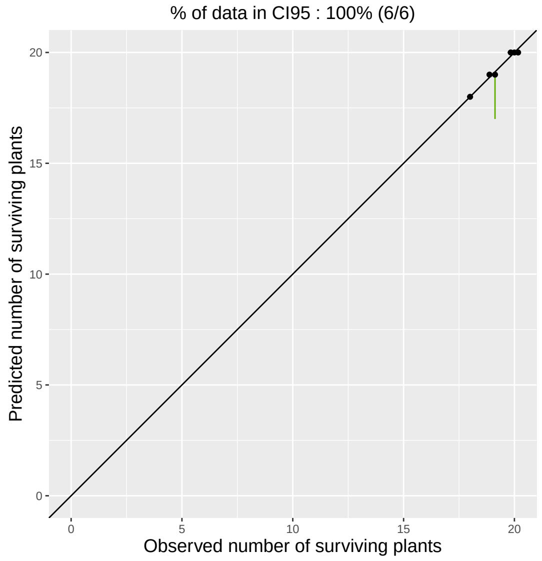

(b) Posterior predictive check (PPC)

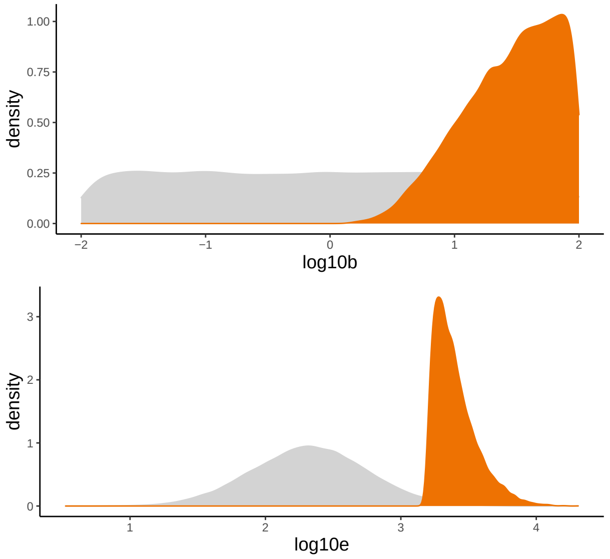

(c) Priors and posteriors

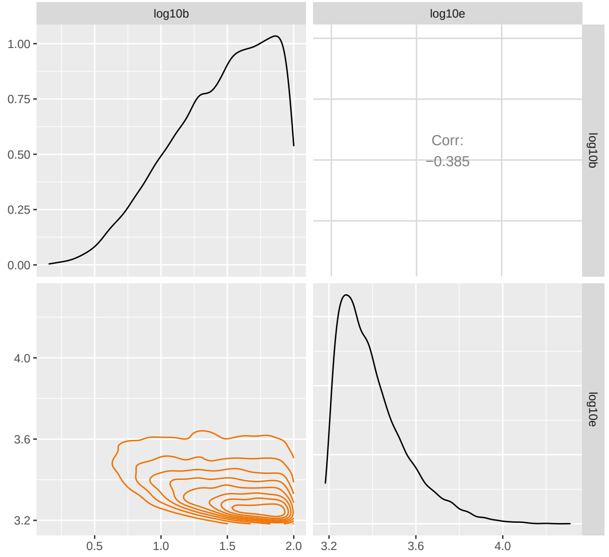

(d) Correlations between parameters

Figure 6: Dose-response curve (a), PPC (b), prior and posterior distributions (c) and correlations between parameters (d).

#### Data set: LOLPE_SE_survival

Table 7: Summary of parameter estimates (parameter d is set to 1) for LOLPE_SE_survival data set

| Parameter | median | Q2.5 | Q97.5 |
|---|---|---|---|
| b | 1.810 | 0.947 | 3.380 |
| e | 2461.879 | 1537.680 | 5801.728 |

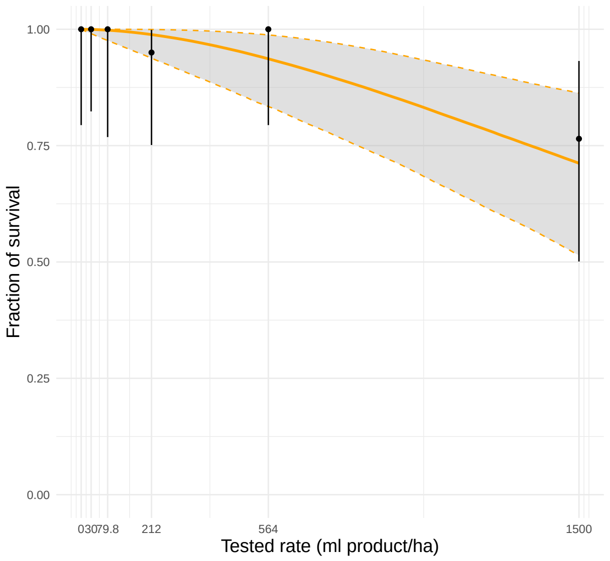

(a) Dose-response curve

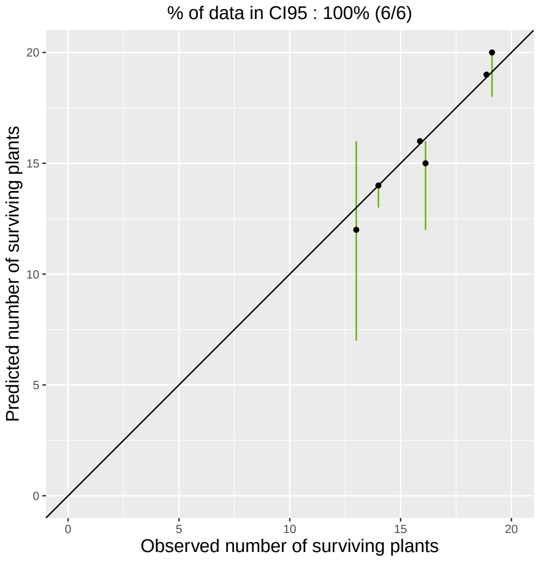

(b) Posterior predictive check (PPC)

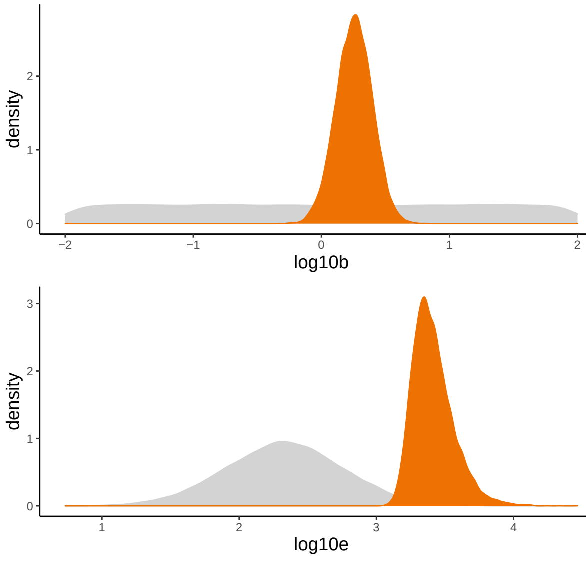

(c) Priors and posteriors

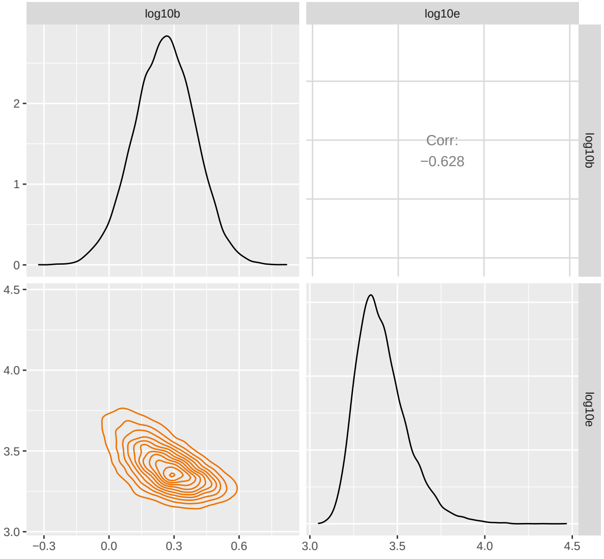

(d) Correlations between parameters

Figure 7: Dose-response curve (a), PPC (b), prior and posterior distributions (c) and correlations between parameters (d).

#### Data set: LYPES_SE_survival

Table 8: Summary of parameter estimates (parameter d is set to 1) for LYPES_SE_survival data set

| Parameter | median | Q2.5 | Q97.5 |
|---|---|---|---|
| b | 6.617 | 1.742 | 65.892 |
| e | 1857.573 | 1530.865 | 3901.110 |

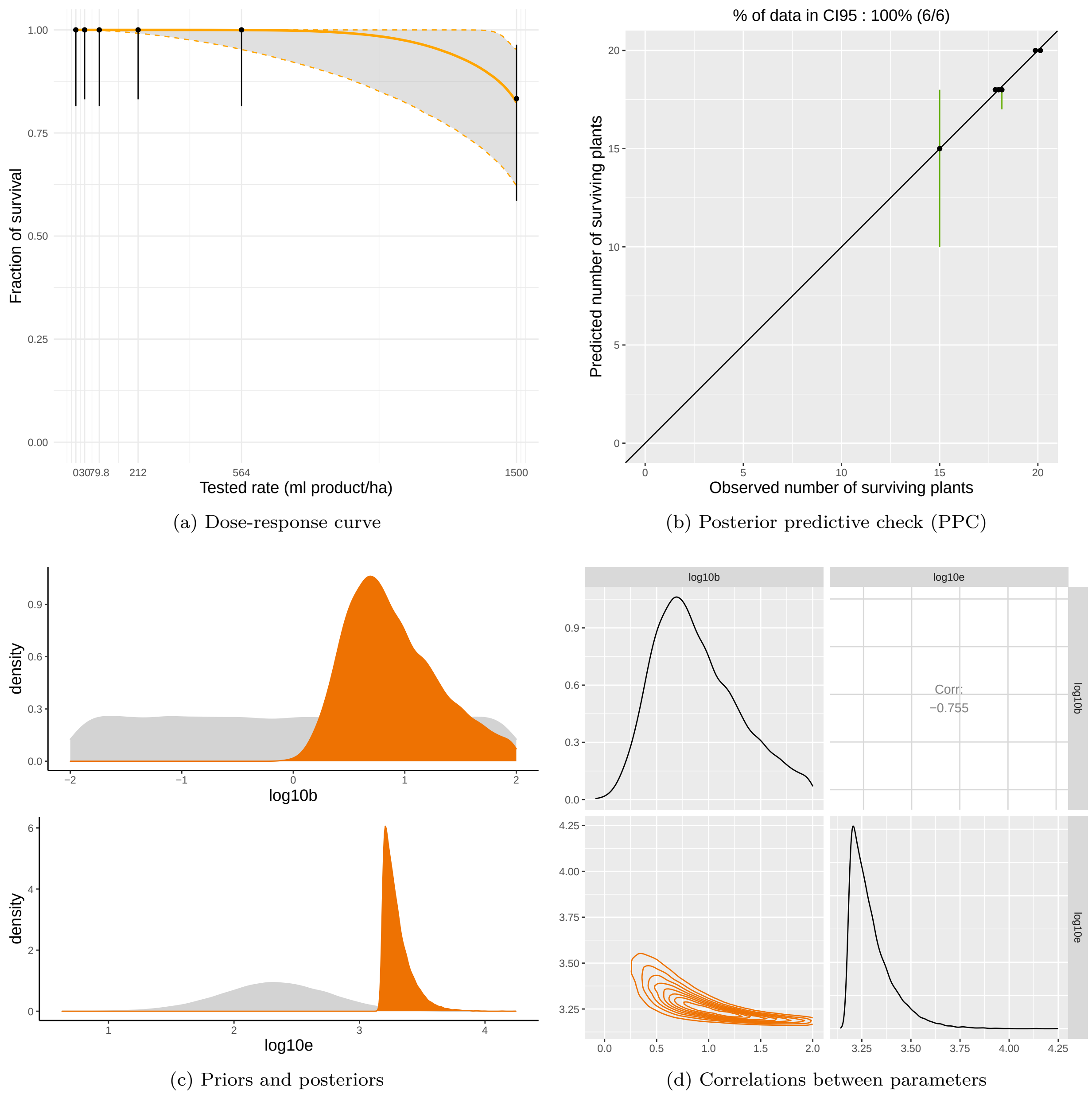

(a) Dose-response curve

(b) Posterior predictive check (PPC)

(c) Priors and posteriors

(d) Correlations between parameters

Figure 8: Dose-response curve (a), PPC (b), prior and posterior distributions (c) and correlations between parameters (d).

#### Data set: TRZAW_SE_survival

Table 9: Summary of parameter estimates (parameter d is set to 1) for TRZAW_SE_survival data set

| Parameter | median | Q2.5 | Q97.5 |
|---|---|---|---|
| b | 3.196 | 1.929 | 4.976 |
| e | 869.590 | 675.933 | 1143.524 |

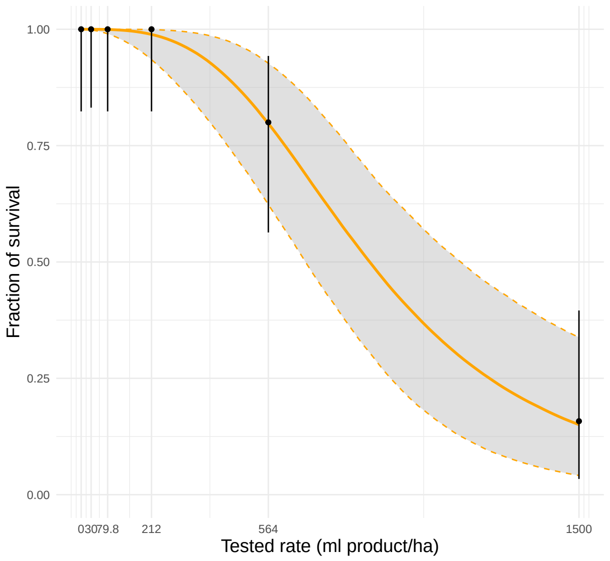

(a) Dose-response curve

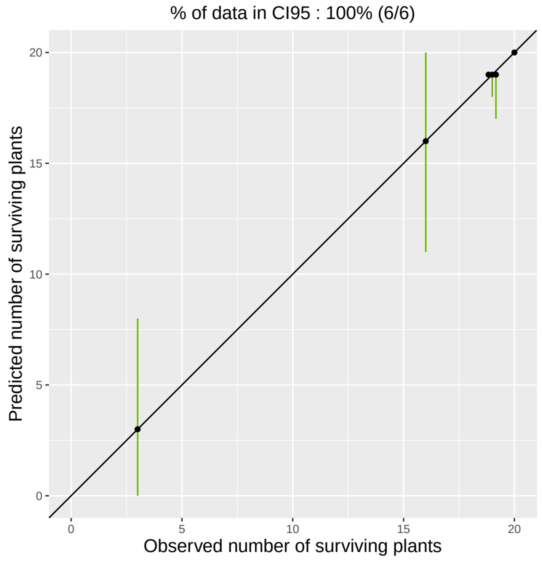

(b) Posterior predictive check (PPC)

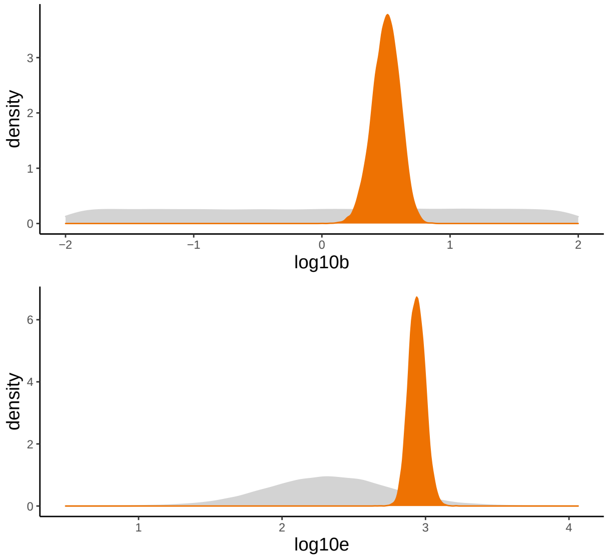

(c) Priors and posteriors

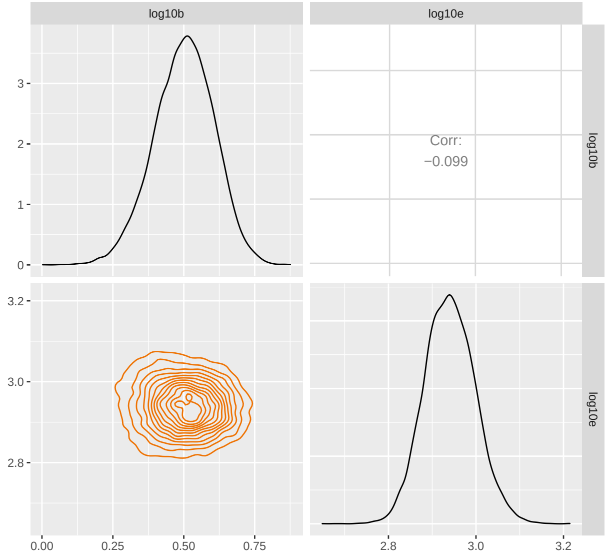

(d) Correlations between parameters

Figure 9: Dose-response curve (a), PPC (b), prior and posterior distributions (c) and correlations between parameters (d).

#### Data set: ZEAMA_SE_survival

Table 10: Summary of parameter estimates (parameter d is set to 1) for ZEAMA_SE_survival data set

| Parameter | median | Q2.5 | Q97.5 |
|---|---|---|---|
| b | 32.203 | 4.456 | 94.979 |
| e | 2303.701 | 1622.054 | 5941.655 |

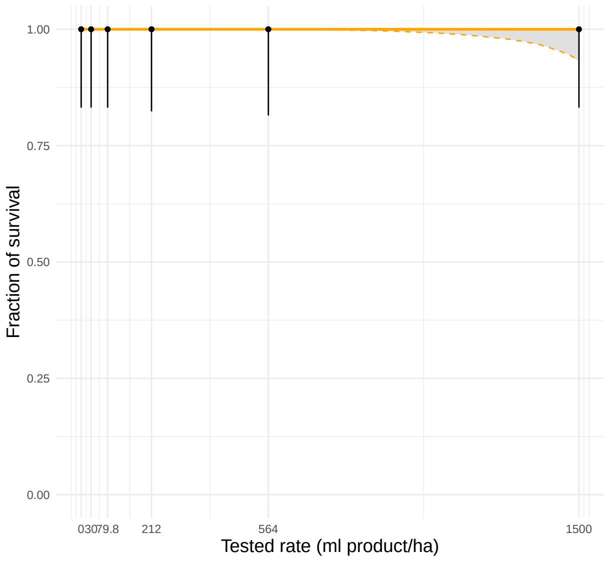

(a) Dose-response curve

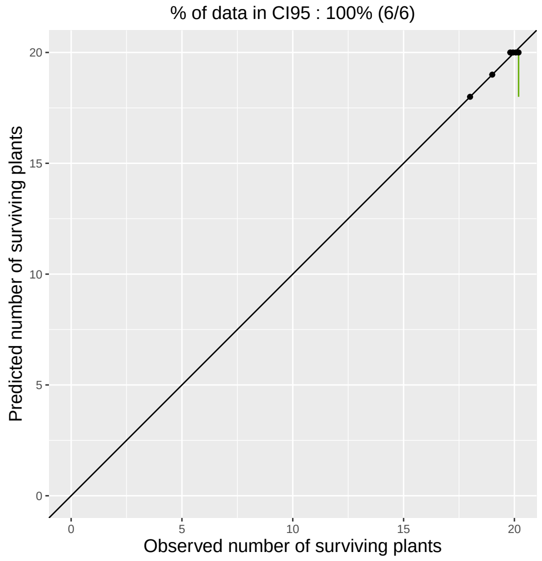

(b) Posterior predictive check (PPC)

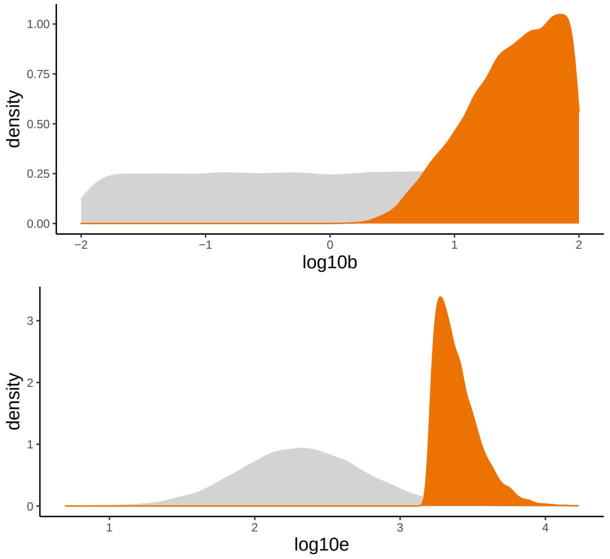

(c) Priors and posteriors

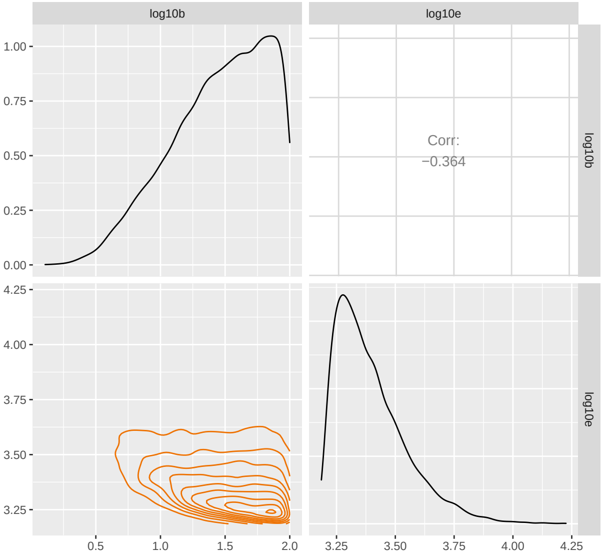

(d) Correlations between parameters

Figure 10: Dose-response curve (a), PPC (b), prior and posterior distributions (c) and correlations between parameters (d).
