## Supplementary material for "How to account for the uncertainty from standard toxicity tests in species sensitivity distributions: an example in non-target plants": Archive with reports on studies: report_SE_weight.pdf

### Dose-response analysis

Study 3

Seedling Emergence test - shoot dry SE_weight endpoint

25 June 2020

Contact:

---

This is a report which provides results on all performed dose-response analyses for the shoot dry SE_weight endpoint of the Seedling Emergence test for study 3.

---

#### Data set: ALLCE_SE_weight

Table 1: Summary of parameter estimates for ALLCE_SE_weight data set

| Parameter | median | Q2.5 | Q97.5 |
|---|---|---|---|
| b | 8.798 | 1.261 | 85.575 |
| d | 0.032 | 0.029 | 0.036 |
| e | 1996.753 | 1532.316 | 4942.222 |
| sigma | 0.008 | 0.006 | 0.011 |

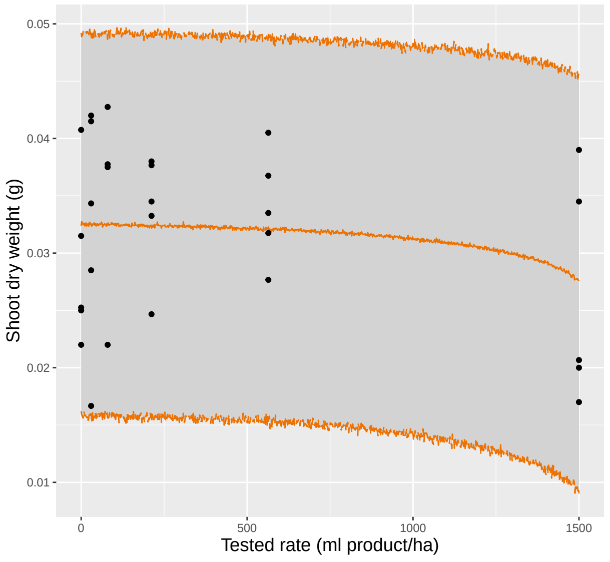

(a) Dose-response curve

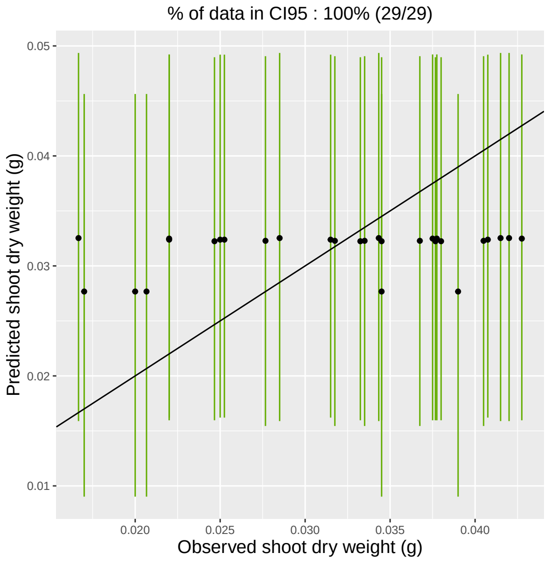

(b) Posterior predictive check (PPC)

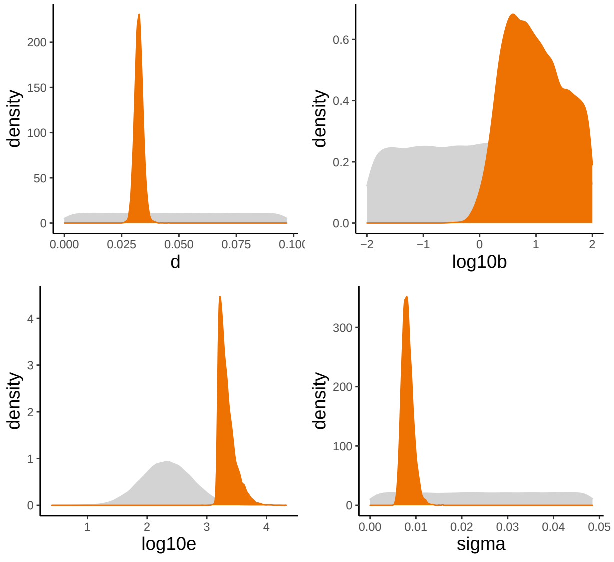

(c) Priors and posteriors

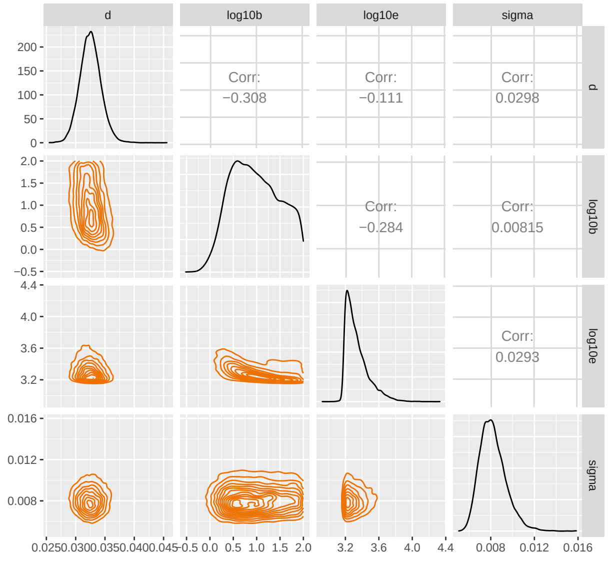

(d) Correlations between parameters

Figure 1: Dose-response curve (a), PPC (b), prior and posterior distributions (c) and correlations between parameters (d).

#### Data set: BEAVA_SE_weight

Table 2: Summary of parameter estimates for BEAVA_SE_weight data set

| Parameter | median | Q2.5 | Q97.5 |
|---|---:|---:|---:|
| b | 8.275 | 1.597 | 82.410 |
| d | 1.247 | 1.181 | 1.312 |
| e | 2034.080 | 1544.751 | 4912.110 |
| sigma | 0.221 | 0.186 | 0.270 |

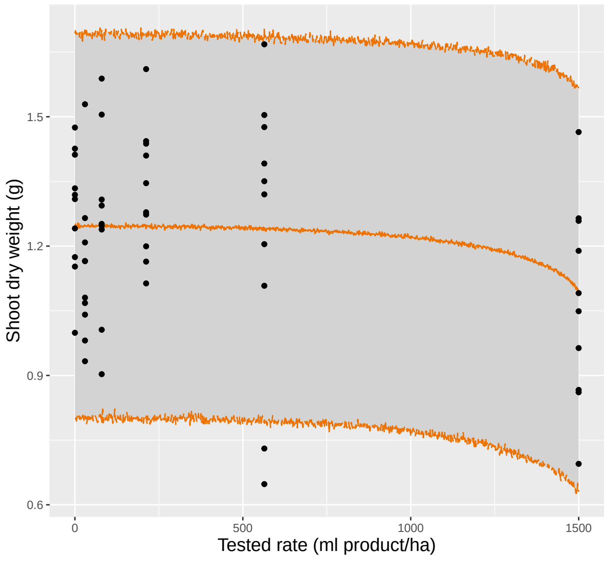

(a) Dose-response curve

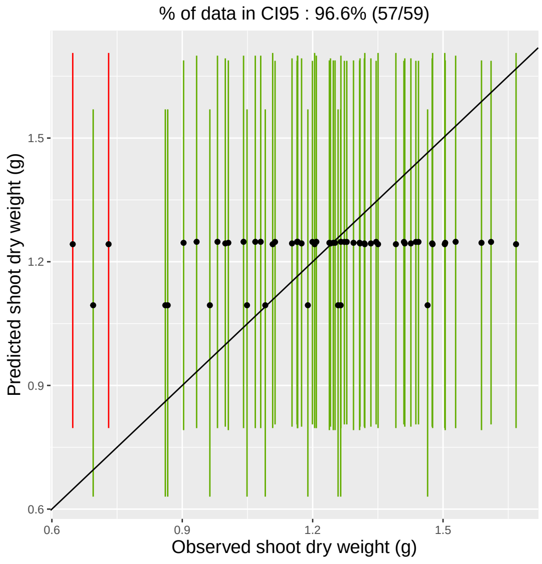

(b) Posterior predictive check (PPC)

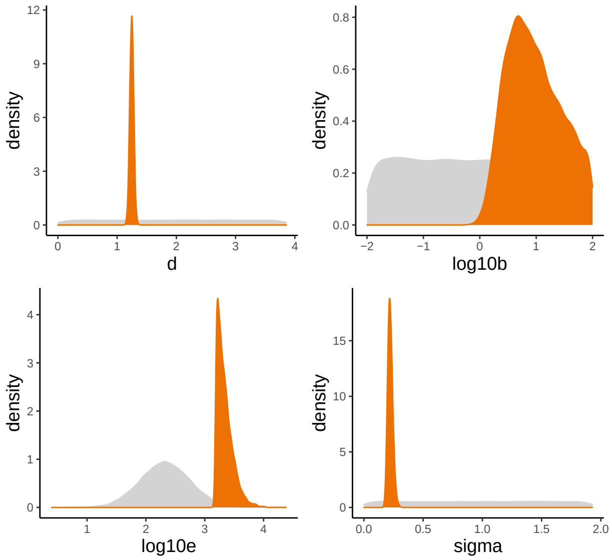

(c) Priors and posteriors

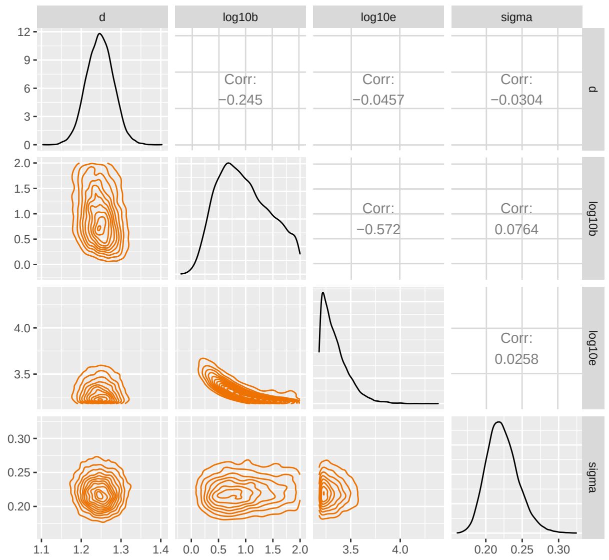

(d) Correlations between parameters

Figure 2: Dose-response curve (a), PPC (b), prior and posterior distributions (c) and correlations between parameters (d).

#### Data set: BRSNW_SE_weight

Table 3: Summary of parameter estimates for BRSNW_SE_weight data set

| Parameter | median | Q2.5 | Q97.5 |
|---|---|---|---|
| b | 2.850 | 1.699 | 15.829 |
| d | 1.871 | 1.761 | 1.989 |
| e | 839.927 | 606.825 | 1093.059 |
| sigma | 0.333 | 0.276 | 0.412 |

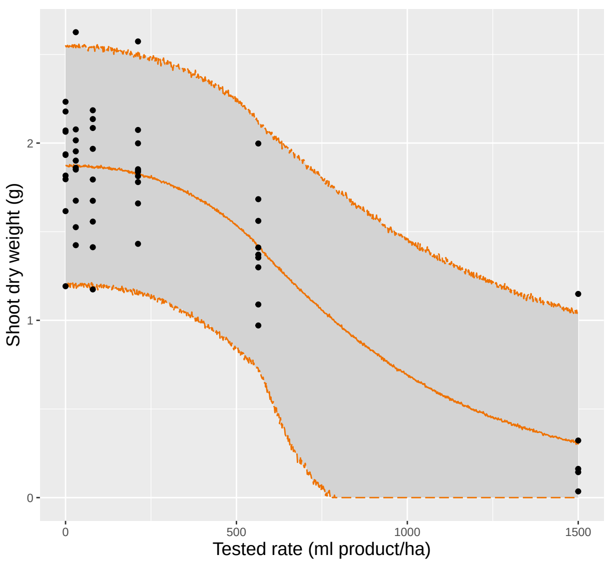

(a) Dose-response curve

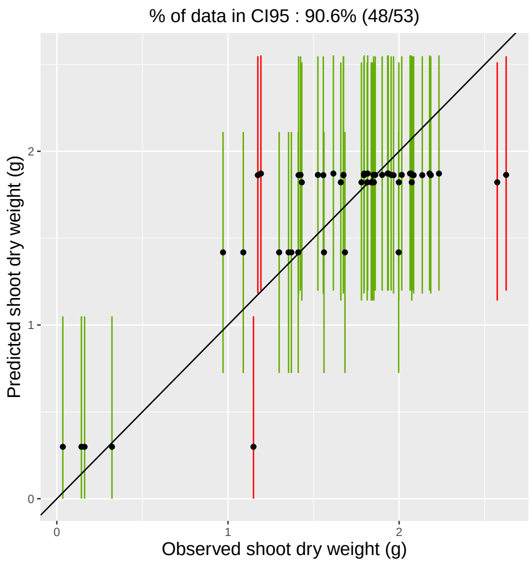

(b) Posterior predictive check (PPC)

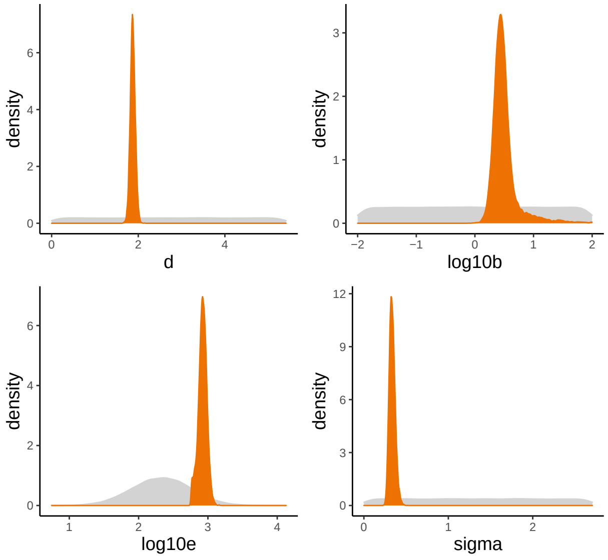

(c) Priors and posteriors

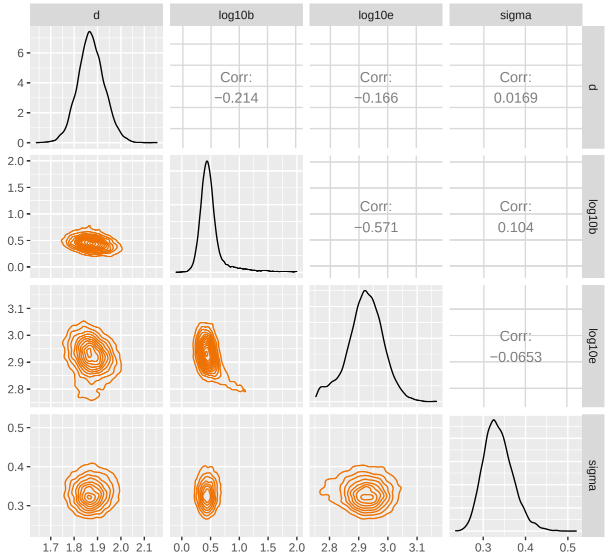

(d) Correlations between parameters

Figure 3: Dose-response curve (a), PPC (b), prior and posterior distributions (c) and correlations between parameters (d).

#### Data set: CUMSA_SE_weight

Table 4: Summary of parameter estimates for CUMSA_SE_weight data set

| Parameter | median | Q2.5 | Q97.5 |
|---|---|---|---|
| b | 2.608 | 1.850 | 4.451 |
| d | 3.713 | 3.490 | 3.942 |
| e | 284.863 | 238.887 | 337.332 |
| sigma | 0.540 | 0.446 | 0.673 |

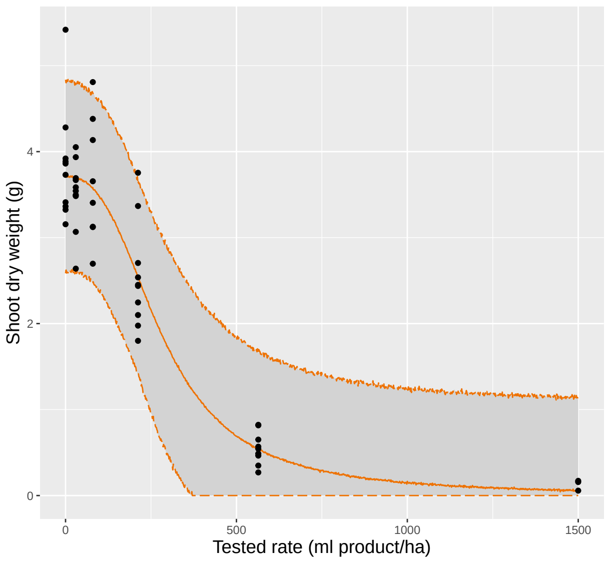

(a) Dose-response curve

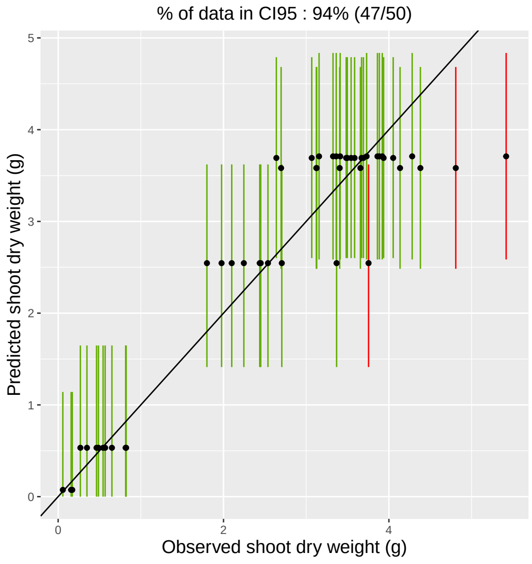

(b) Posterior predictive check (PPC)

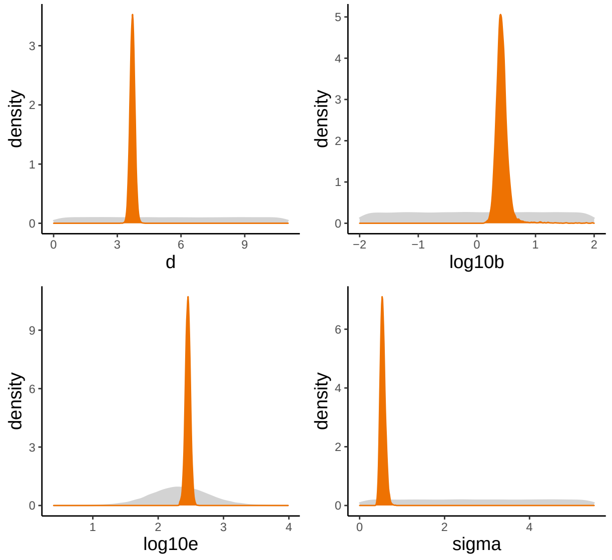

(c) Priors and posteriors

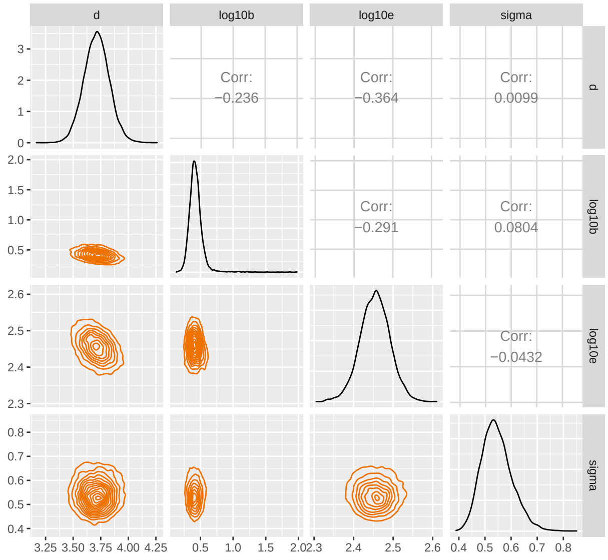

(d) Correlations between parameters

Figure 4: Dose-response curve (a), PPC (b), prior and posterior distributions (c) and correlations between parameters (d).

#### Data set: FAGES_SE_weight

Table 5: Summary of parameter estimates for FAGES_SE_weight data set

| Parameter | median | Q2.5 | Q97.5 |
|---|---|---|---|
| b | 1.146 | 0.498 | 3.124 |
| d | 1.727 | 1.541 | 1.994 |
| e | 1248.414 | 730.313 | 2214.897 |
| sigma | 0.435 | 0.365 | 0.533 |

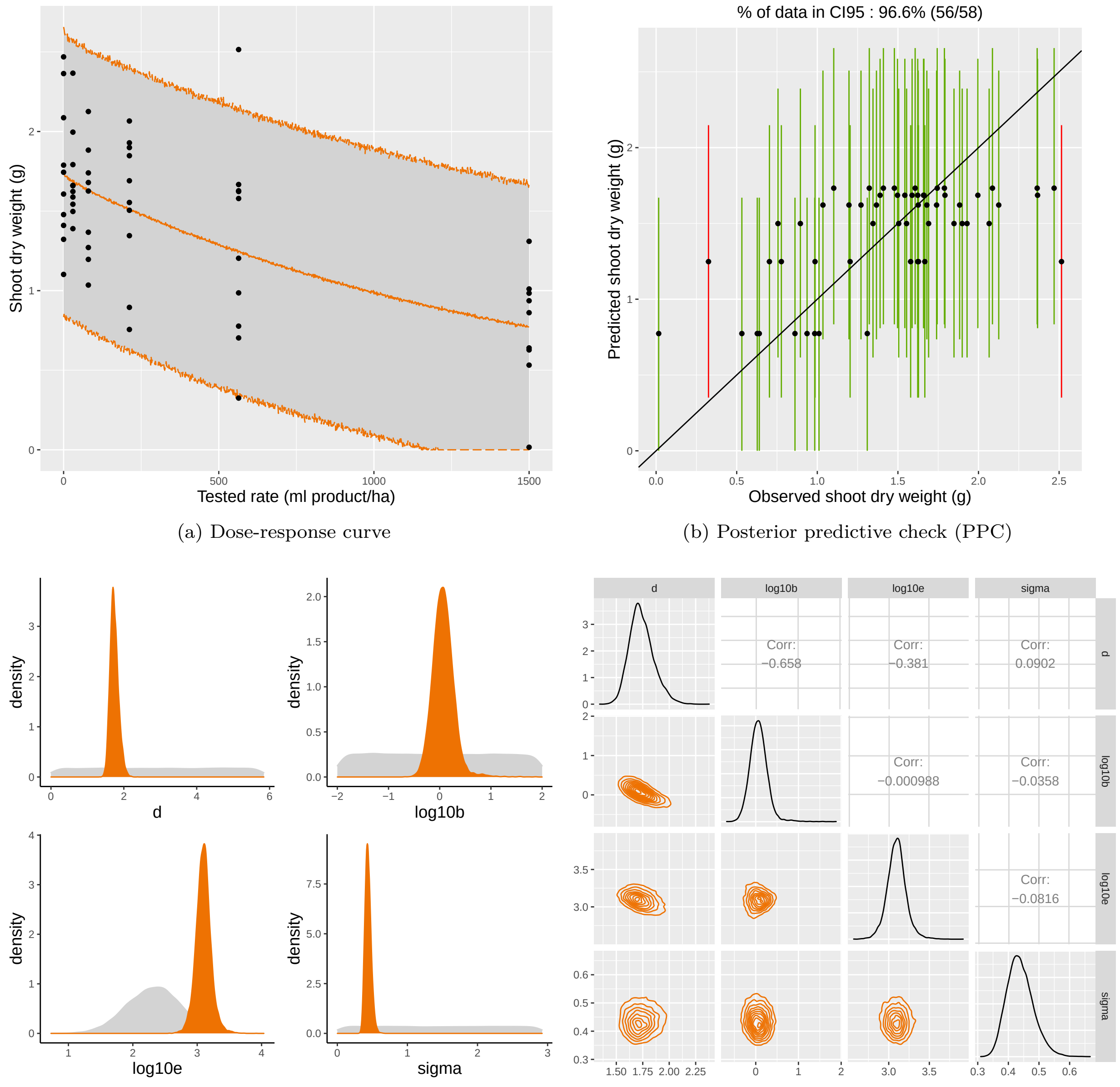

(a) Dose-response curve

(b) Posterior predictive check (PPC)

(c) Priors and posteriors

(d) Correlations between parameters

Figure 5: Dose-response curve (a), PPC (b), prior and posterior distributions (c) and correlations between parameters (d).

#### Data set: GLXMA_SE_weight

Table 6: Summary of parameter estimates for GLXMA_SE_weight data set

| Parameter | median | Q2.5 | Q97.5 |
|---|---:|---:|---:|
| b | 5.882 | 1.844 | 57.119 |
| d | 1.443 | 1.378 | 1.509 |
| e | 1742.039 | 1523.044 | 2651.235 |
| sigma | 0.225 | 0.188 | 0.274 |

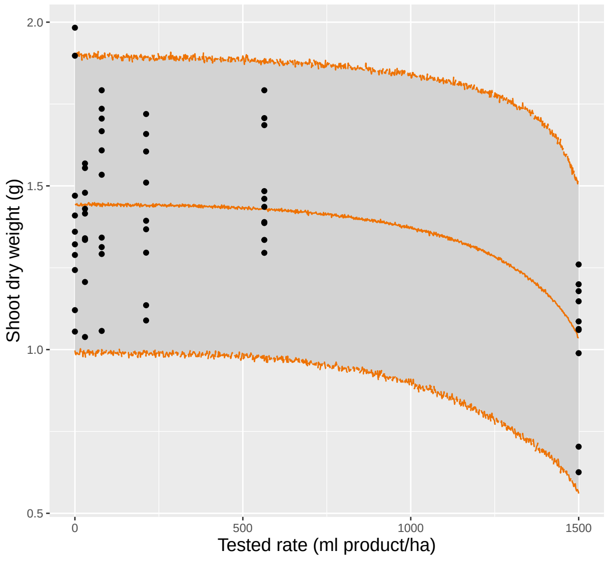

(a) Dose-response curve

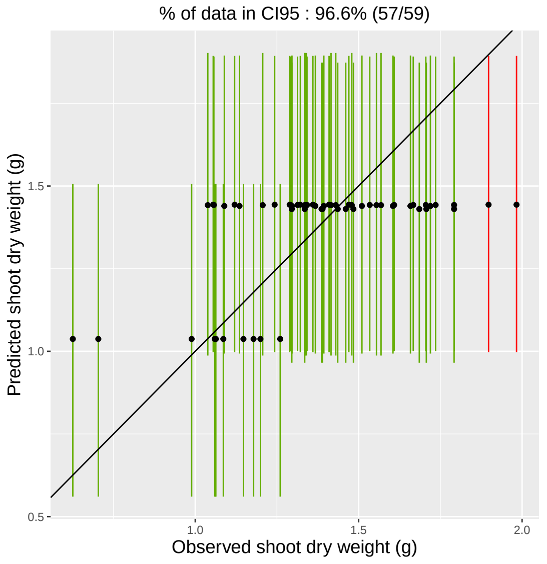

(b) Posterior predictive check (PPC)

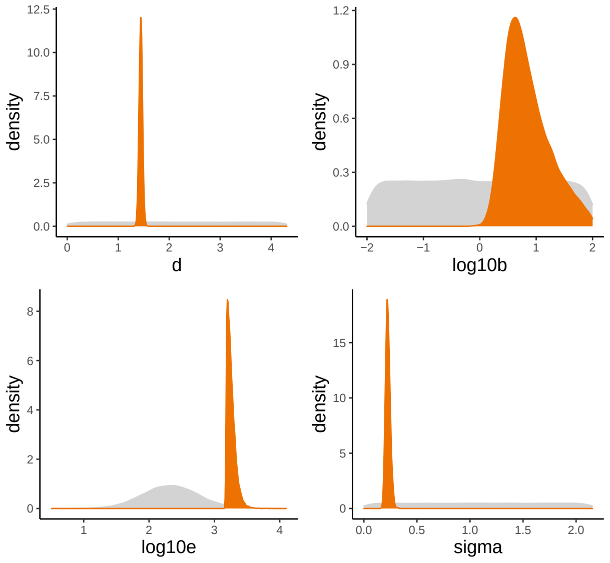

(c) Priors and posteriors

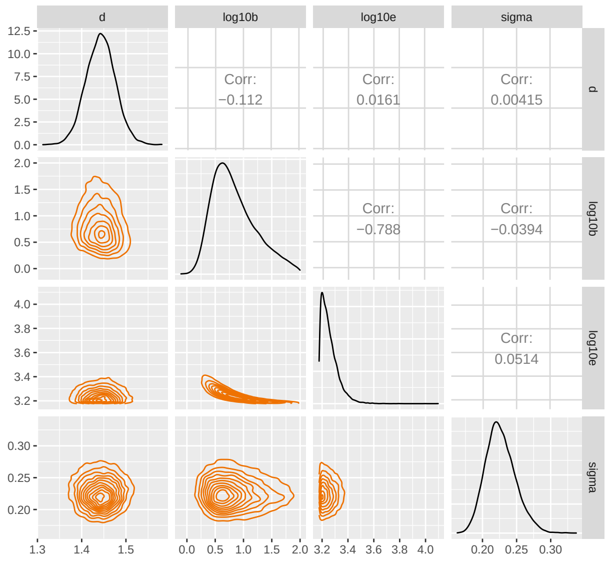

(d) Correlations between parameters

Figure 6: Dose-response curve (a), PPC (b), prior and posterior distributions (c) and correlations between parameters (d).

#### Data set: LOLPE_SE_weight

Table 7: Summary of parameter estimates for LOLPE_SE_weight data set

| Parameter | median | Q2.5 | Q97.5 |
|---|---|---|---|
| b | 3.865 | 1.016 | 41.831 |
| d | 0.119 | 0.107 | 0.134 |
| e | 1420.284 | 959.080 | 1924.471 |
| sigma | 0.027 | 0.021 | 0.038 |

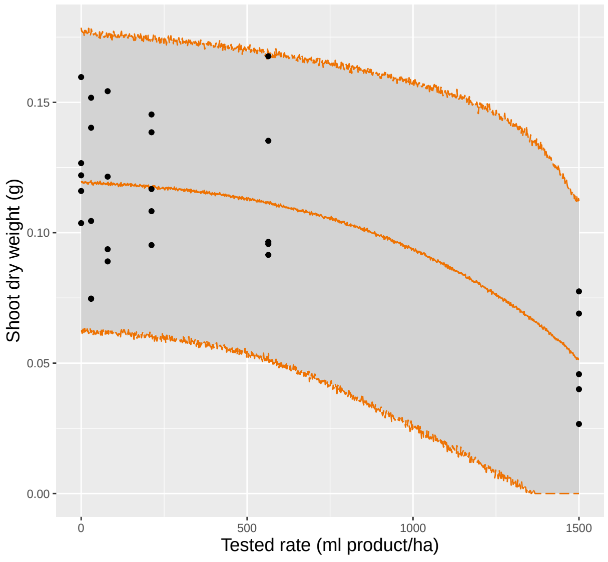

(a) Dose-response curve

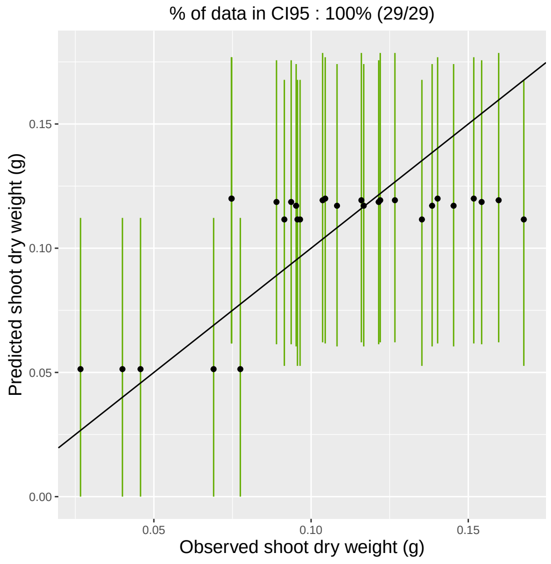

(b) Posterior predictive check (PPC)

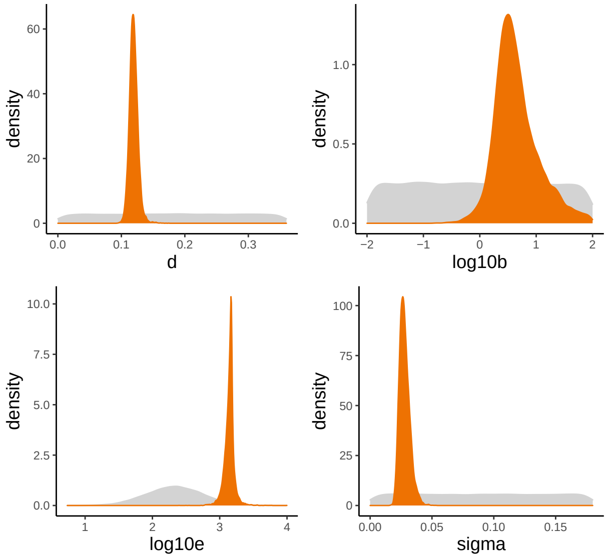

(c) Priors and posteriors

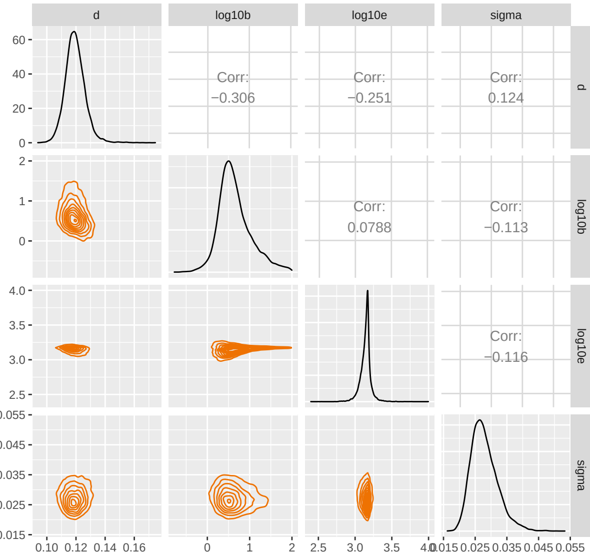

(d) Correlations between parameters

Figure 7: Dose-response curve (a), PPC (b), prior and posterior distributions (c) and correlations between parameters (d).

#### Data set: LYPES_SE_weight

Table 8: Summary of parameter estimates for LYPES_SE_weight data set

| Parameter | median | Q2.5 | Q97.5 |
|---|---|---|---|
| b | 4.450 | 2.794 | 46.781 |
| d | 1.332 | 1.258 | 1.404 |
| e | 772.613 | 583.507 | 970.832 |
| sigma | 0.219 | 0.183 | 0.269 |

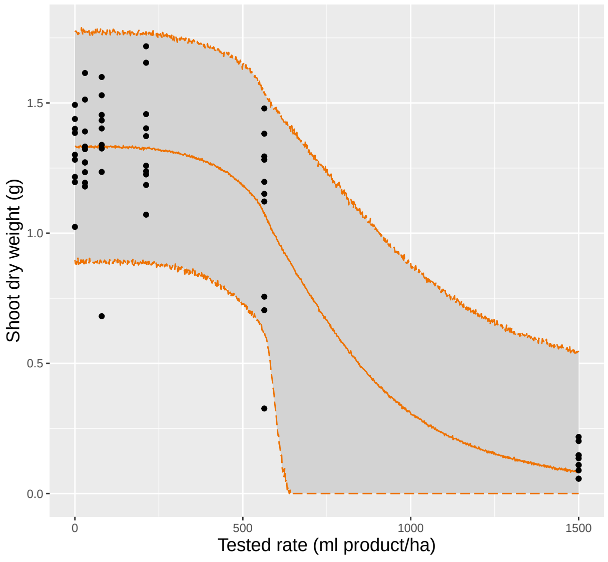

(a) Dose-response curve

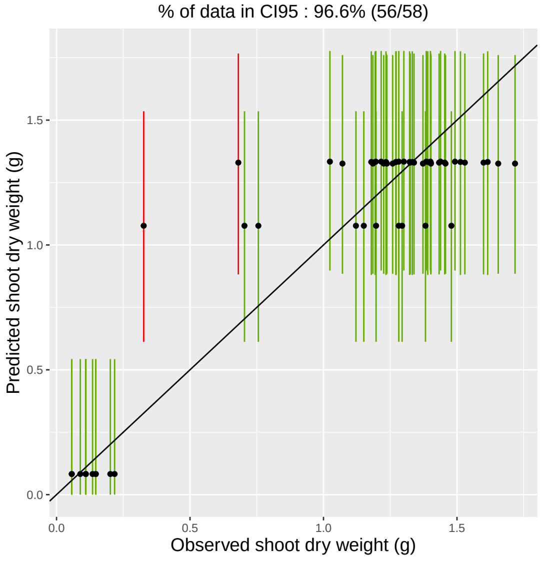

(b) Posterior predictive check (PPC)

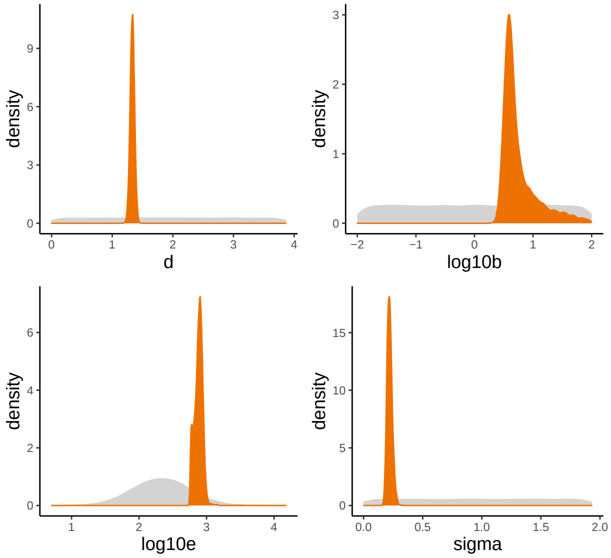

(c) Priors and posteriors

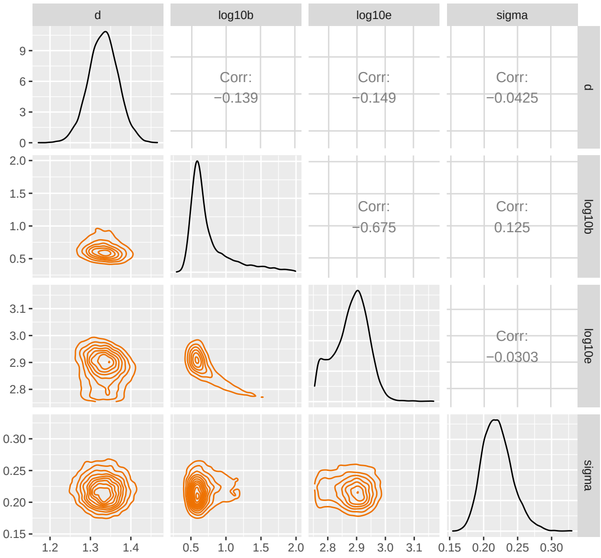

(d) Correlations between parameters

Figure 8: Dose-response curve (a), PPC (b), prior and posterior distributions (c) and correlations between parameters (d).

#### Data set: TRZAW_SE_weight

Table 9: Summary of parameter estimates for TRZAW_SE_weight data set

| Parameter | median | Q2.5 | Q97.5 |
|---|---:|---:|---:|
| b | 3.944 | 2.421 | 37.643 |
| d | 0.442 | 0.421 | 0.463 |
| e | 213.878 | 198.568 | 234.769 |
| sigma | 0.036 | 0.028 | 0.049 |

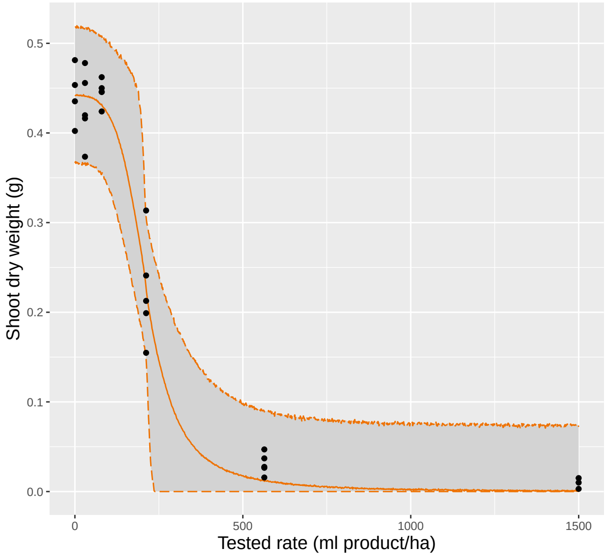

(a) Dose-response curve

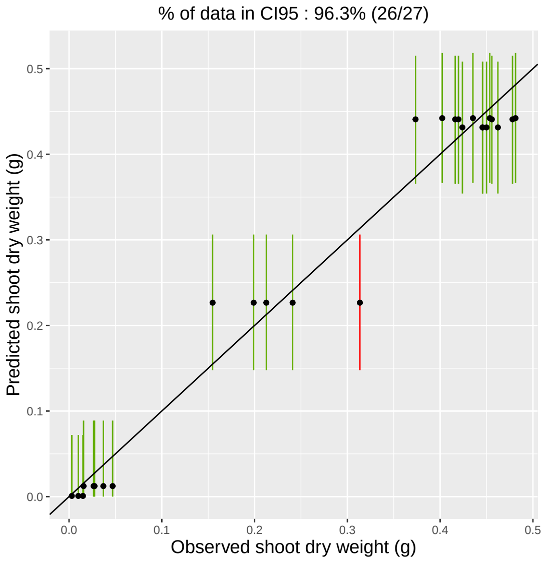

(b) Posterior predictive check (PPC)

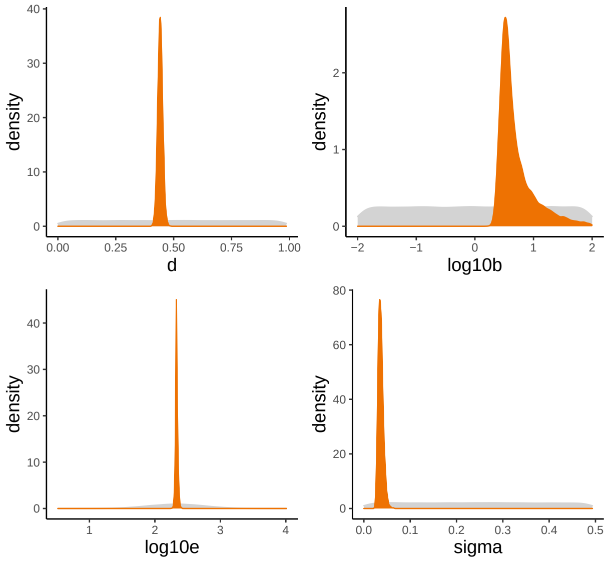

(c) Priors and posteriors

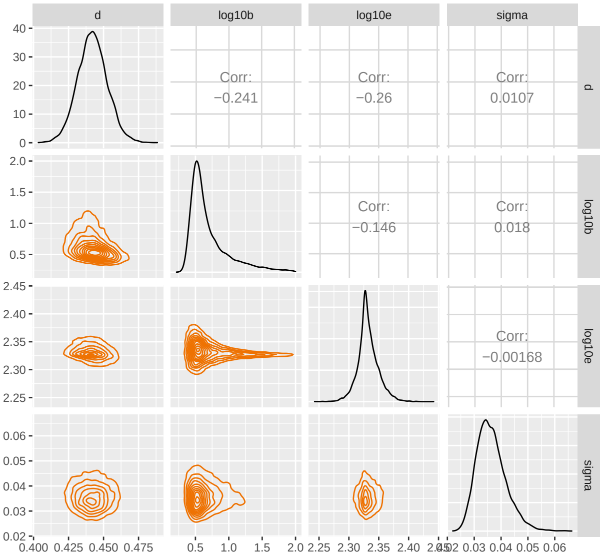

(d) Correlations between parameters

Figure 9: Dose-response curve (a), PPC (b), prior and posterior distributions (c) and correlations between parameters (d).

### Data set: ZEAMA_SE_weight

Table 10: Summary of parameter estimates for ZEAMA_SE_weight data set

| Parameter | median | Q2.5 | Q97.5 |
|---|---|---|---|
| b | 19.384 | 2.734 | 91.540 |
| d | 4.123 | 3.953 | 4.298 |
| e | 2149.669 | 1573.526 | 5601.577 |
| sigma | 0.621 | 0.524 | 0.755 |

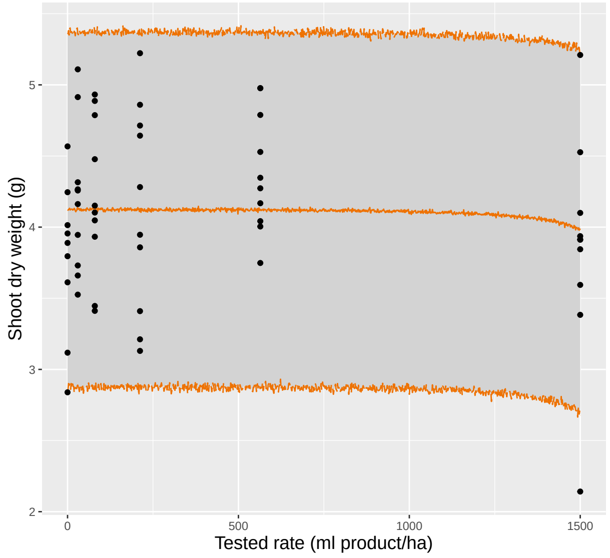

(a) Dose-response curve

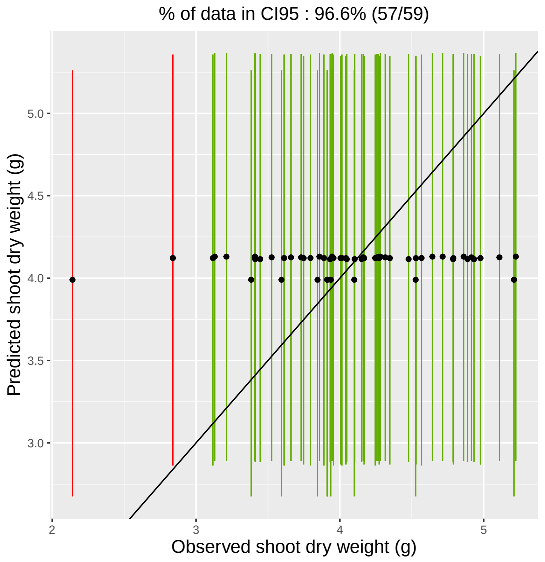

(b) Posterior predictive check (PPC)

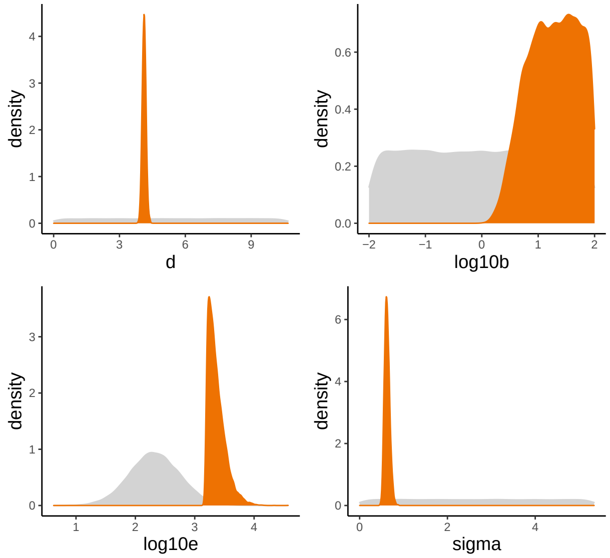

(c) Priors and posteriors

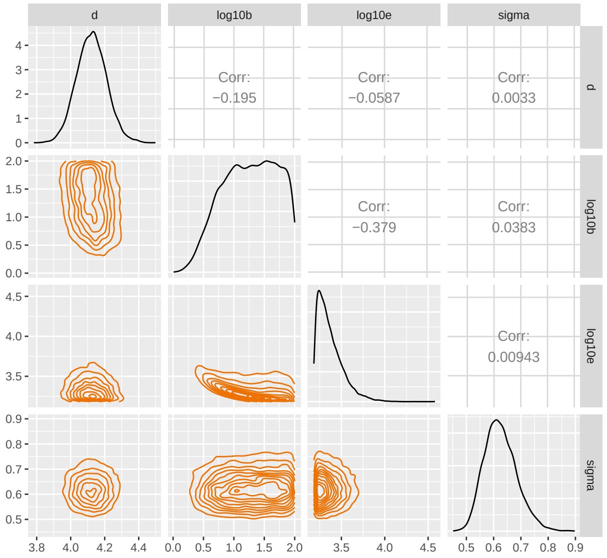

(d) Correlations between parameters

Figure 10: Dose-response curve (a), PPC (b), prior and posterior distributions (c) and correlations between parameters (d).
