## Supplementary material for "How to account for the uncertainty from standard toxicity tests in species sensitivity distributions: an example in non-target plants": Archive with reports on studies: report_VV_weight.pdf

### Dose-response analysis

Study 3

Vegetative Vigour test - shoot dry VV_weight endpoint

25 June 2020

Contact:

---

This is a report which provides results on all performed dose-response analyses for the shoot dry VV_weight endpoint of the Vegetative Vigour test for study 3.

---

#### Data set: ALLCE_VV_weight

Table 1: Summary of parameter estimates for ALLCE_VV_weight data set

| Parameter | median | Q2.5 | Q97.5 |
|---|---|---|---|
| b | 3.275 | 1.145 | 42.386 |
| d | 0.318 | 0.288 | 0.350 |
| e | 1558.400 | 1196.420 | 2439.766 |
| sigma | 0.068 | 0.053 | 0.092 |

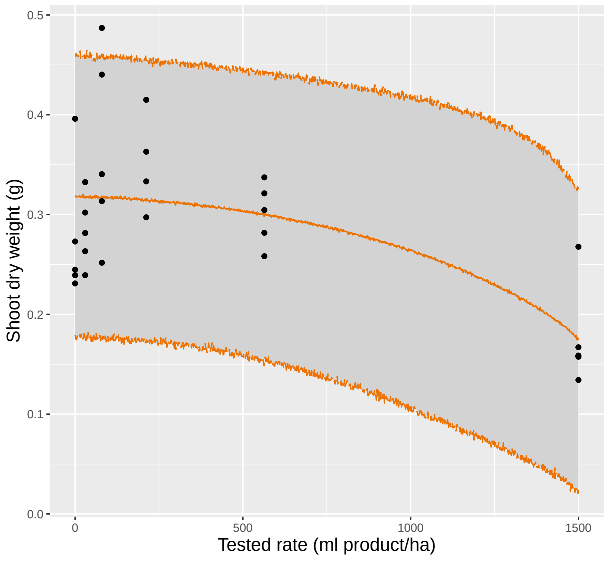

(a) Dose-response curve

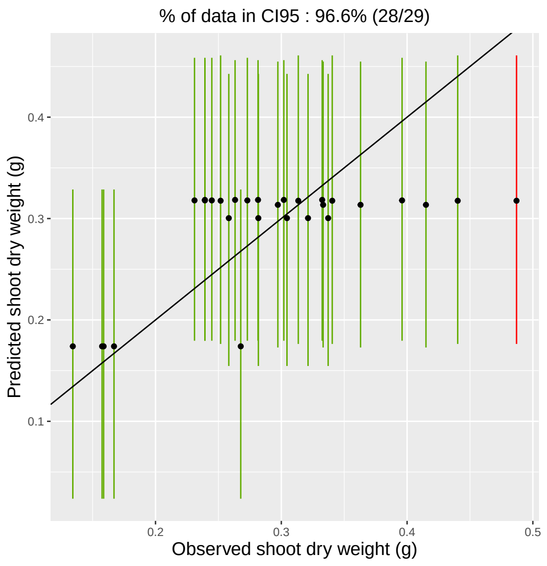

(b) Posterior predictive check (PPC)

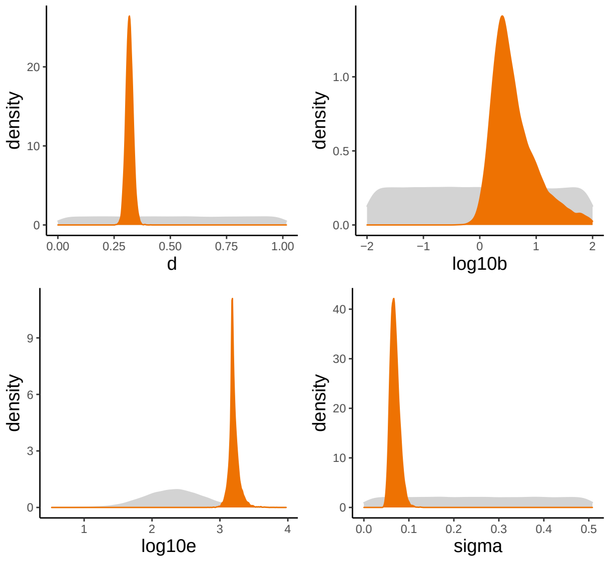

(c) Priors and posteriors

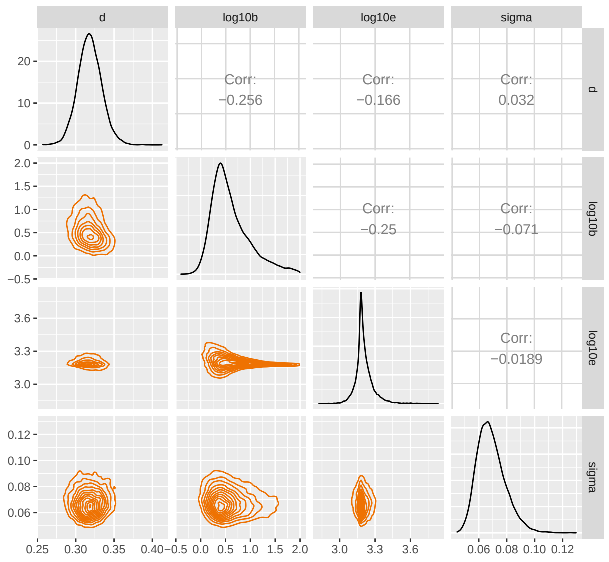

(d) Correlations between parameters

Figure 1: Dose-response curve (a), PPC (b), prior and posterior distributions (c) and correlations between parameters (d).

#### Data set: BEAVA_VV_weight

Table 2: Summary of parameter estimates for BEAVA_VV_weight data set

| Parameter | median | Q2.5 | Q97.5 |
|---|---|---|---|
| b | 1.413 | 0.718 | 10.140 |
| d | 4.086 | 3.871 | 4.352 |
| e | 2377.972 | 1609.346 | 4221.291 |
| sigma | 0.522 | 0.437 | 0.638 |

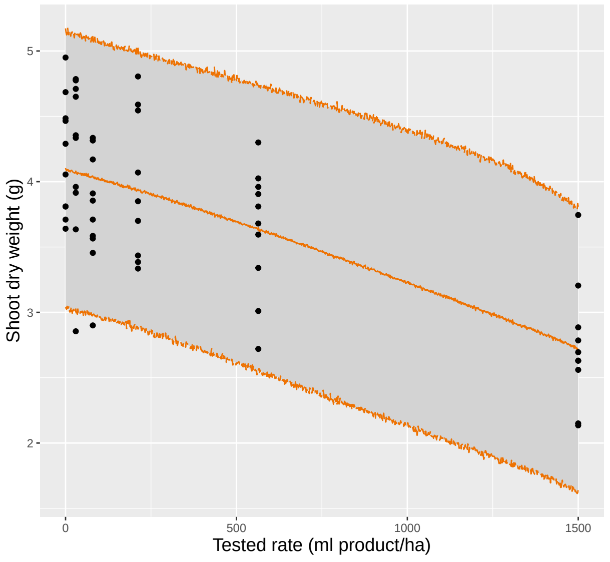

(a) Dose-response curve

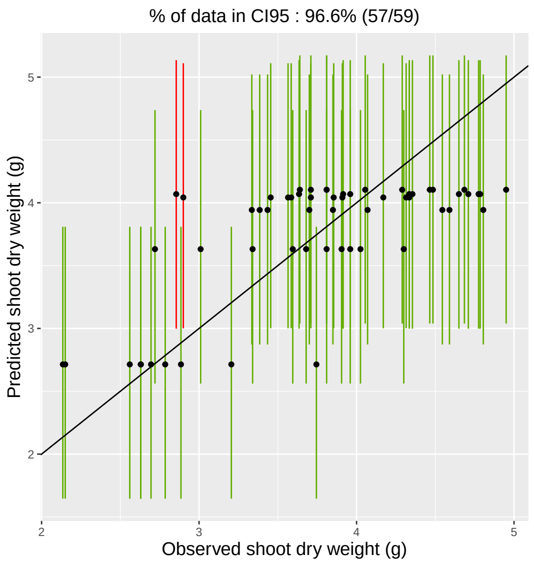

(b) Posterior predictive check (PPC)

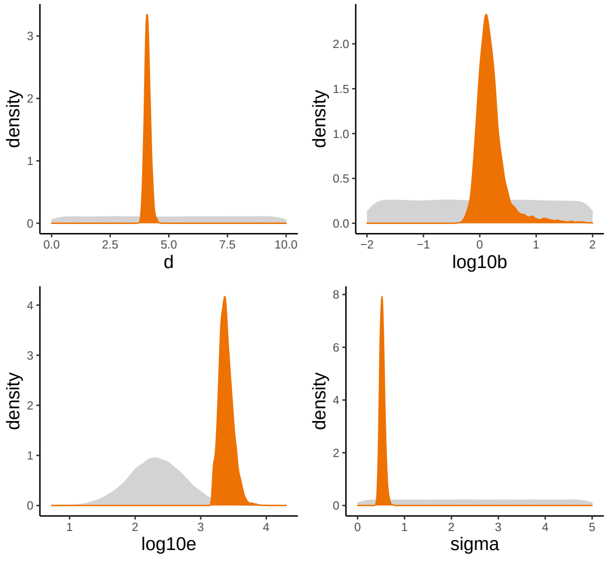

(c) Priors and posteriors

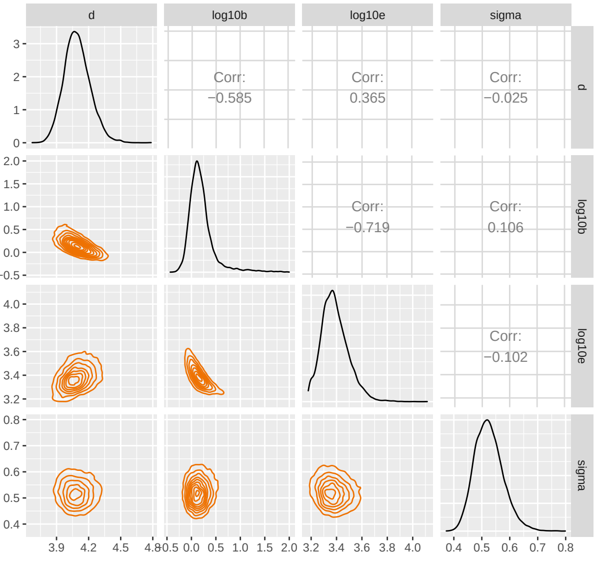

(d) Correlations between parameters

Figure 2: Dose-response curve (a), PPC (b), prior and posterior distributions (c) and correlations between parameters (d).

#### Data set: BRSNW_VV_weight

Table 3: Summary of parameter estimates for BRSNW_VV_weight data set

| Parameter | median | Q2.5 | Q97.5 |
|---|---|---|---|
| b | 1.605 | 1.214 | 2.123 |
| d | 8.226 | 7.817 | 8.658 |
| e | 838.599 | 699.395 | 1011.098 |
| sigma | 1.061 | 0.889 | 1.302 |

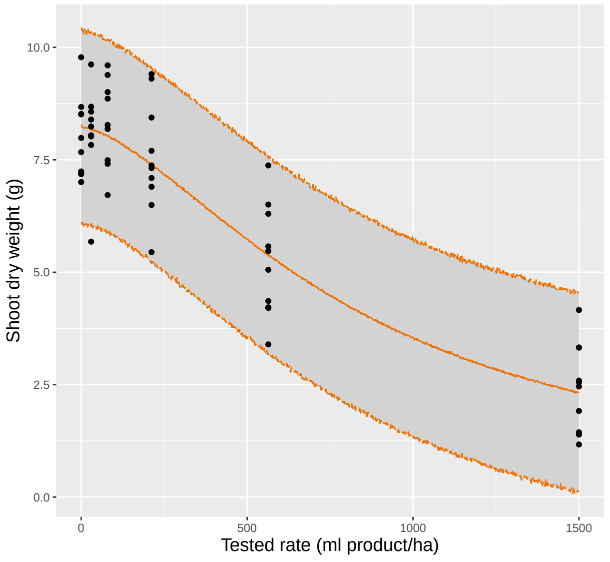

(a) Dose-response curve

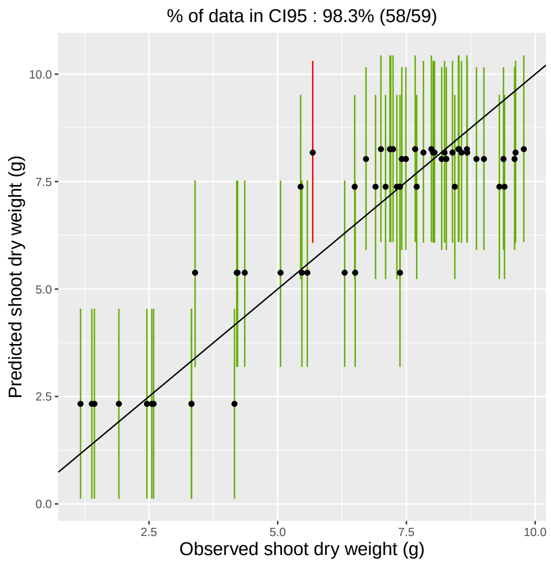

(b) Posterior predictive check (PPC)

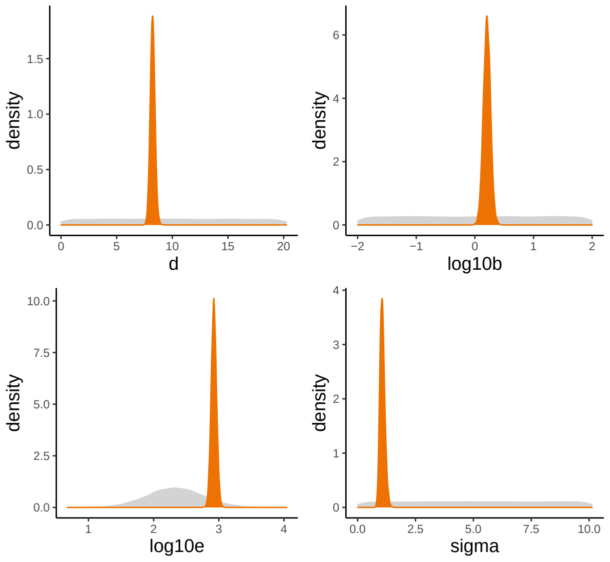

(c) Priors and posteriors

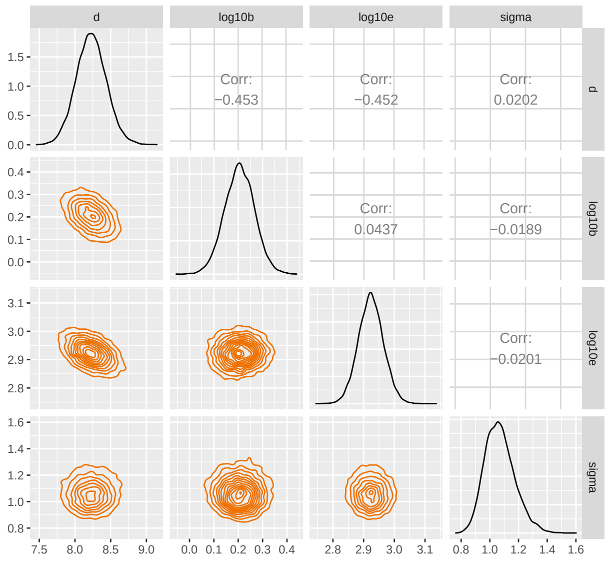

(d) Correlations between parameters

Figure 3: Dose-response curve (a), PPC (b), prior and posterior distributions (c) and correlations between parameters (d).

### Data set: CUMSA_VV_weight

Table 4: Summary of parameter estimates for CUMSA_VV_weight data set

| Parameter | median | Q2.5 | Q97.5 |
|---|---|---|---|
| b | 1.245 | 0.979 | 1.620 |
| d | 11.063 | 10.387 | 11.798 |
| e | 302.975 | 245.543 | 372.040 |
| sigma | 1.277 | 1.060 | 1.581 |

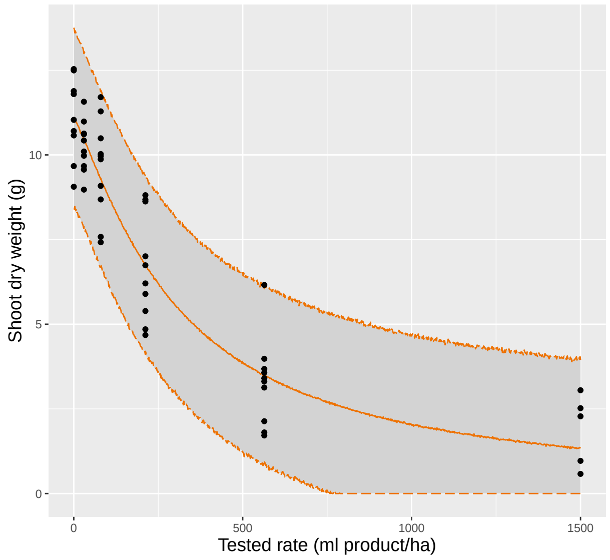

(a) Dose-response curve

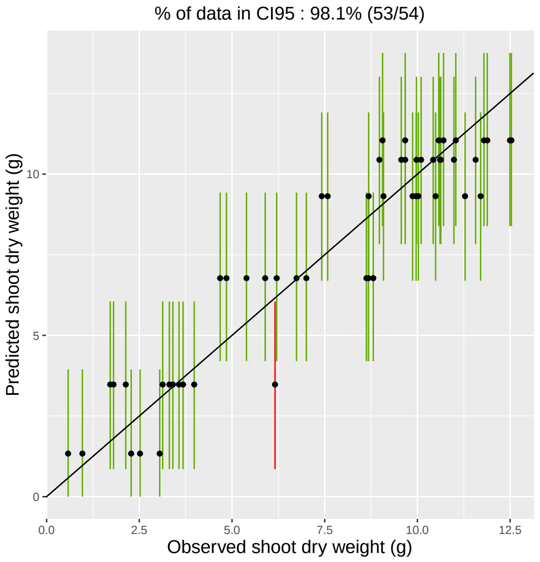

(b) Posterior predictive check (PPC)

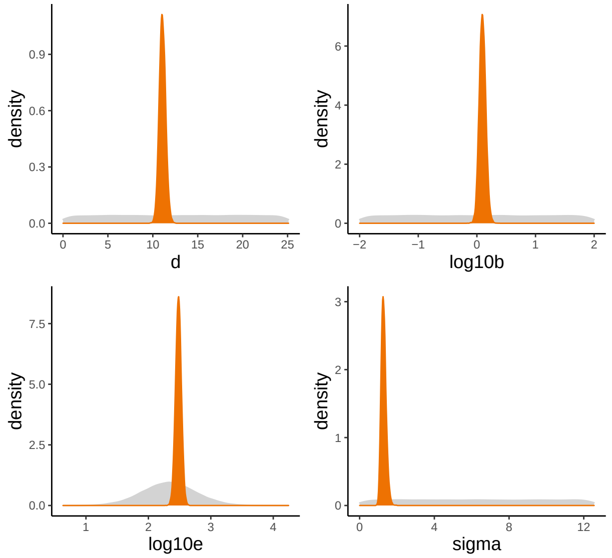

(c) Priors and posteriors

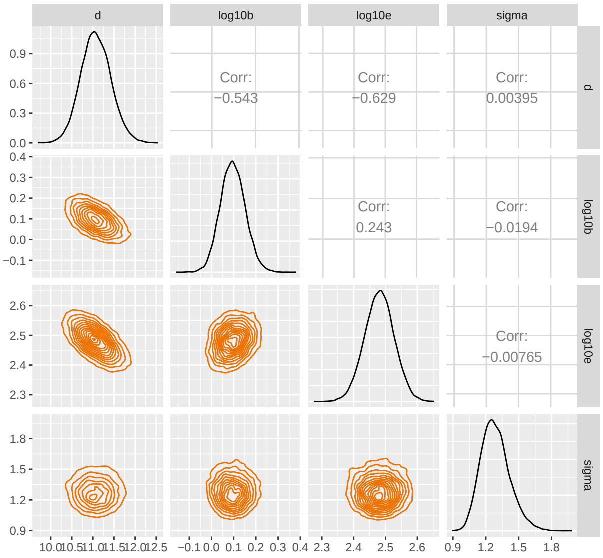

(d) Correlations between parameters

Figure 4: Dose-response curve (a), PPC (b), prior and posterior distributions (c) and correlations between parameters (d).

#### Data set: FAGES_VV_weight

Table 5: Summary of parameter estimates for FAGES_VV_weight data set

| Parameter | median | Q2.5 | Q97.5 |
|---|---|---|---|
| b | 1.353 | 1.034 | 1.831 |
| d | 6.613 | 6.129 | 7.129 |
| e | 262.085 | 205.337 | 333.376 |
| sigma | 0.926 | 0.772 | 1.141 |

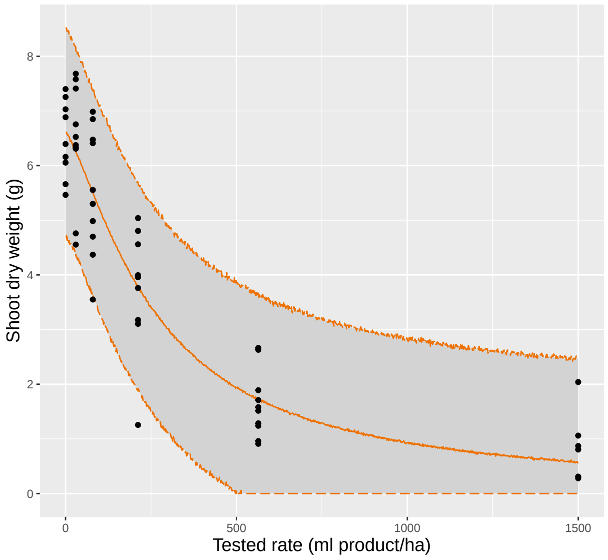

(a) Dose-response curve

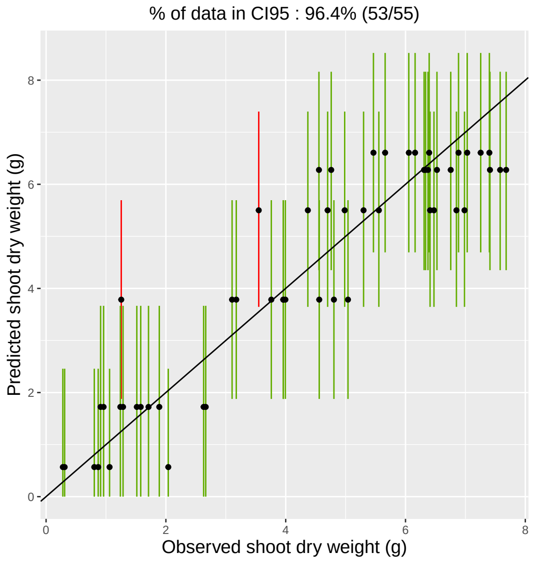

(b) Posterior predictive check (PPC)

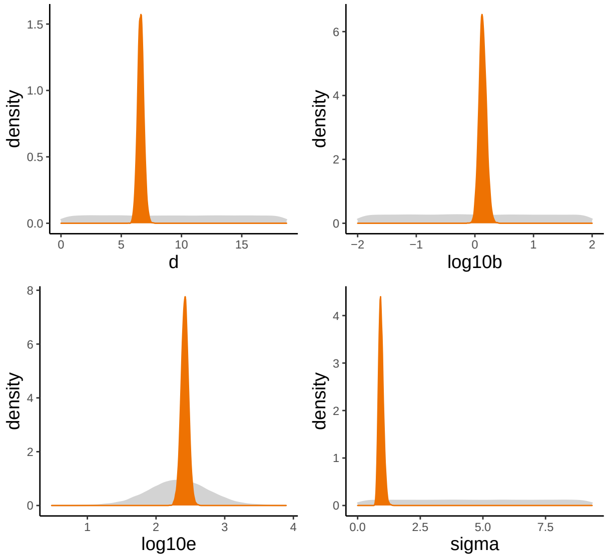

(c) Priors and posteriors

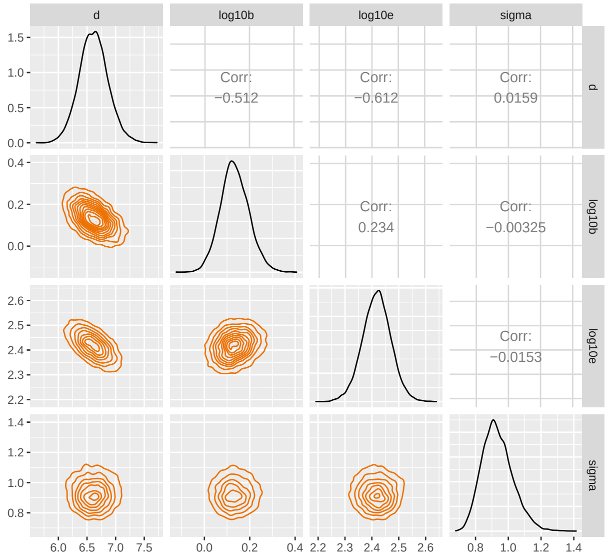

(d) Correlations between parameters

Figure 5: Dose-response curve (a), PPC (b), prior and posterior distributions (c) and correlations between parameters (d).

#### Data set: GLXMA_VV_weight

Table 6: Summary of parameter estimates for GLXMA_VV_weight data set

| Parameter | median | Q2.5 | Q97.5 |
|---|---|---|---|
| b | 1.395 | 1.090 | 1.773 |
| d | 5.744 | 5.576 | 5.924 |
| e | 1590.354 | 1396.262 | 1865.475 |
| sigma | 0.433 | 0.362 | 0.528 |

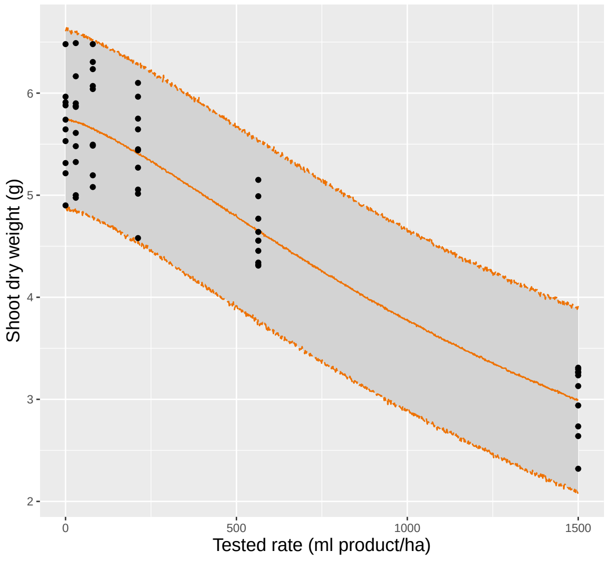

(a) Dose-response curve

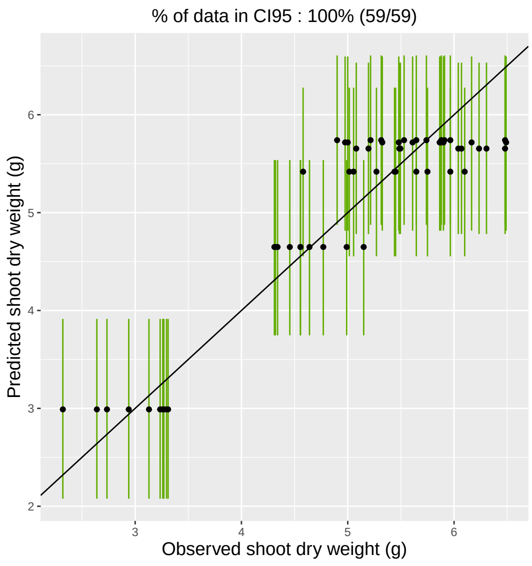

(b) Posterior predictive check (PPC)

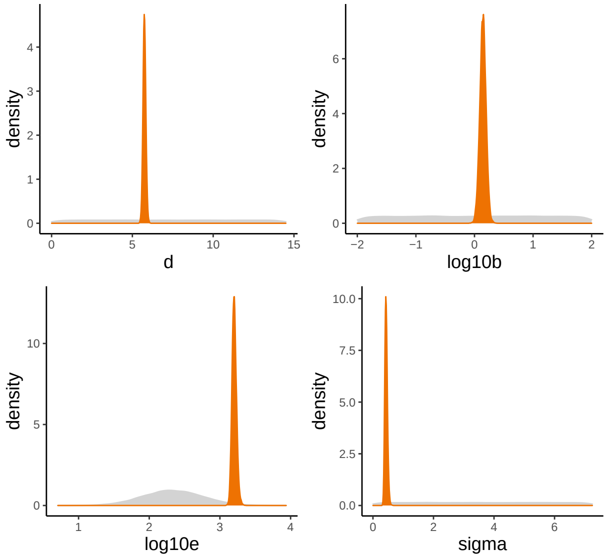

(c) Priors and posteriors

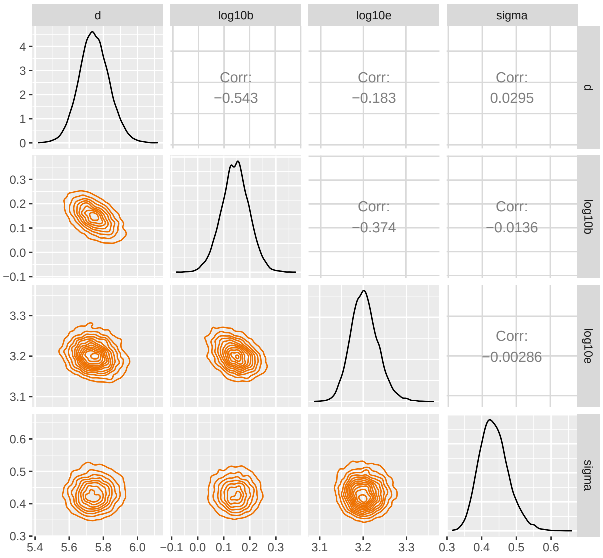

(d) Correlations between parameters

Figure 6: Dose-response curve (a), PPC (b), prior and posterior distributions (c) and correlations between parameters (d).

#### Data set: LOLPE_VV_weight

Table 7: Summary of parameter estimates for LOLPE_VV_weight data set

| Parameter | median | Q2.5 | Q97.5 |
|---|---|---|---|
| b | 1.124 | 0.726 | 1.715 |
| d | 0.473 | 0.439 | 0.515 |
| e | 1038.131 | 759.704 | 1420.947 |
| sigma | 0.051 | 0.040 | 0.070 |

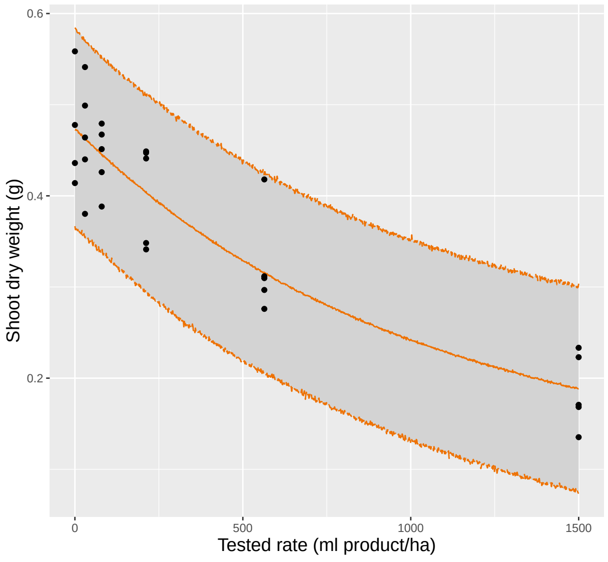

(a) Dose-response curve

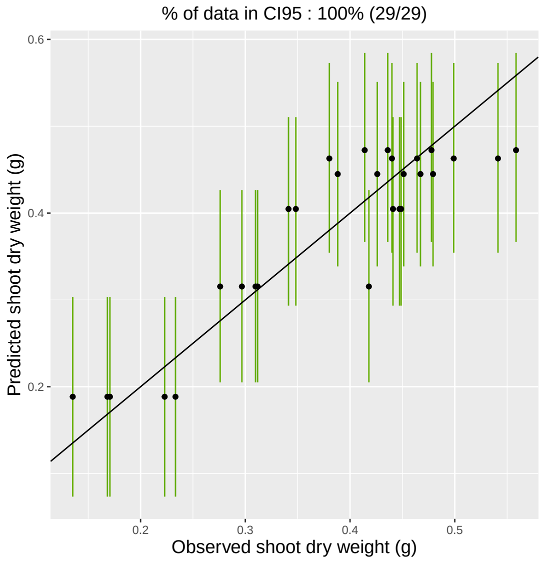

(b) Posterior predictive check (PPC)

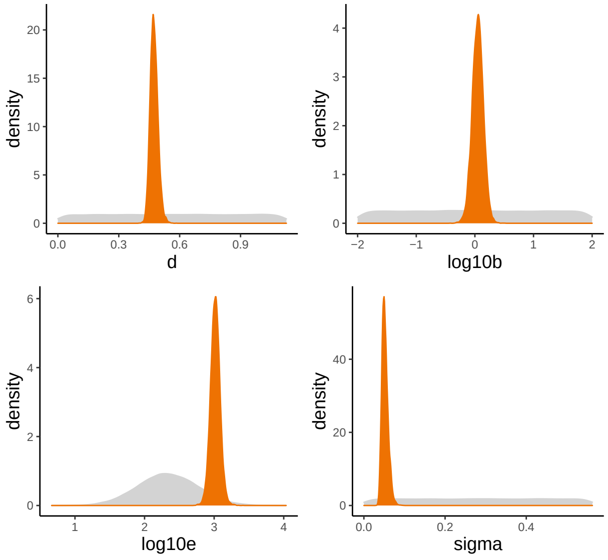

(c) Priors and posteriors

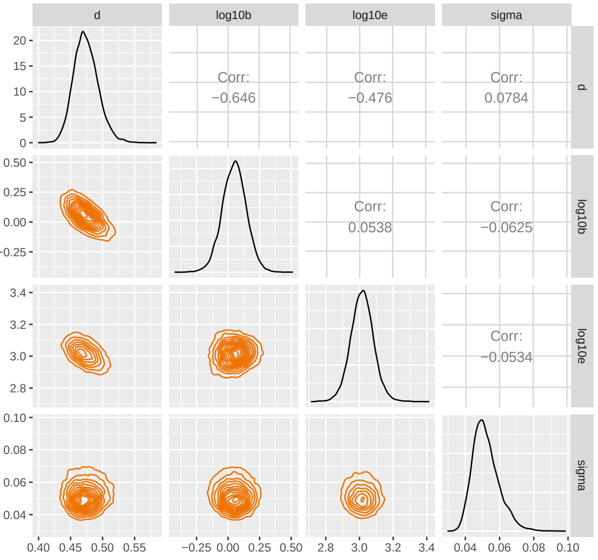

(d) Correlations between parameters

Figure 7: Dose-response curve (a), PPC (b), prior and posterior distributions (c) and correlations between parameters (d).

#### Data set: LYPES_VV_weight

Table 8: Summary of parameter estimates for LYPES_VV_weight data set

| Parameter | median | Q2.5 | Q97.5 |
|---|---|---|---|
| b | 1.248 | 0.922 | 1.712 |
| d | 3.135 | 2.852 | 3.435 |
| e | 147.991 | 109.404 | 199.499 |
| sigma | 0.508 | 0.418 | 0.633 |

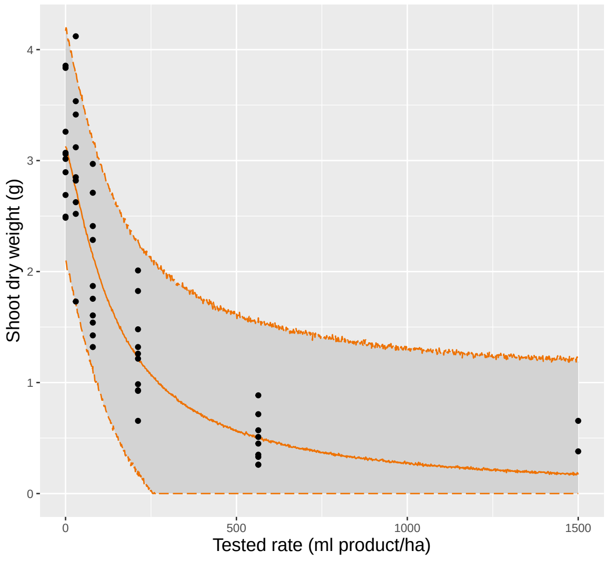

(a) Dose-response curve

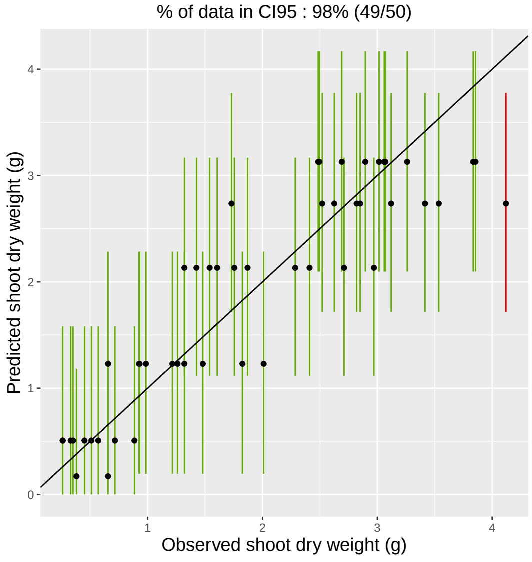

(b) Posterior predictive check (PPC)

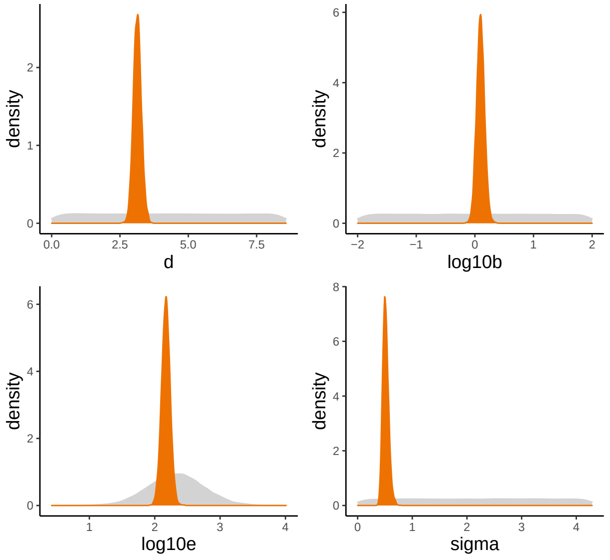

(c) Priors and posteriors

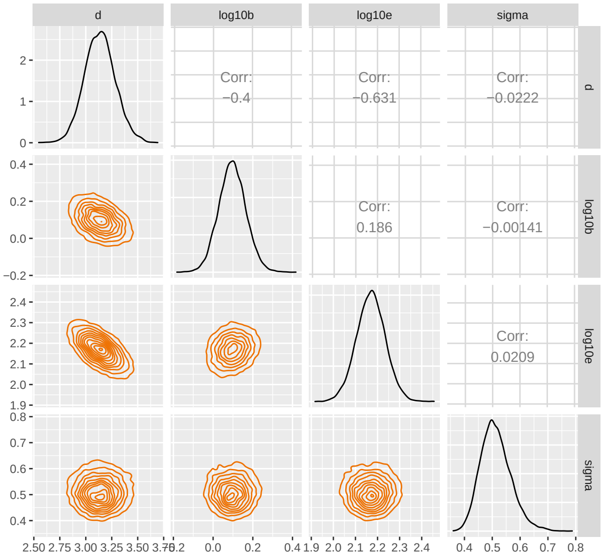

(d) Correlations between parameters

Figure 8: Dose-response curve (a), PPC (b), prior and posterior distributions (c) and correlations between parameters (d).

### Data set: TRZAW_VV_weight

Table 9: Summary of parameter estimates for TRZAW_VV_weight data set

| Parameter | median | Q2.5 | Q97.5 |
|---|---|---|---|
| b | 1.846 | 1.431 | 2.423 |
| d | 0.922 | 0.867 | 0.980 |
| e | 342.308 | 285.442 | 409.994 |
| sigma | 0.087 | 0.067 | 0.118 |

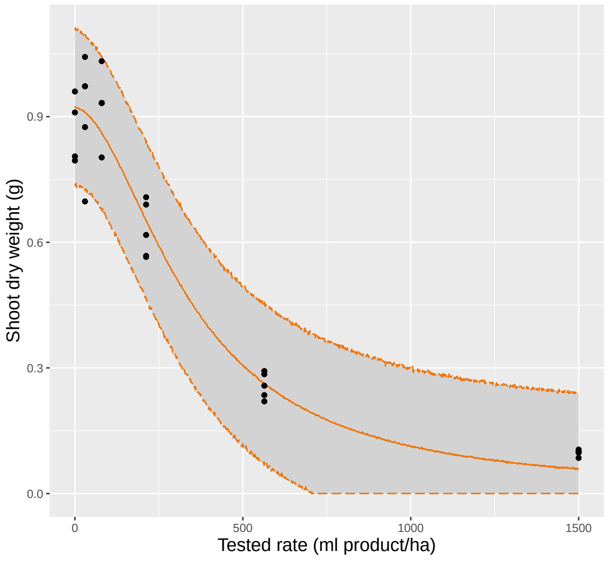

(a) Dose-response curve

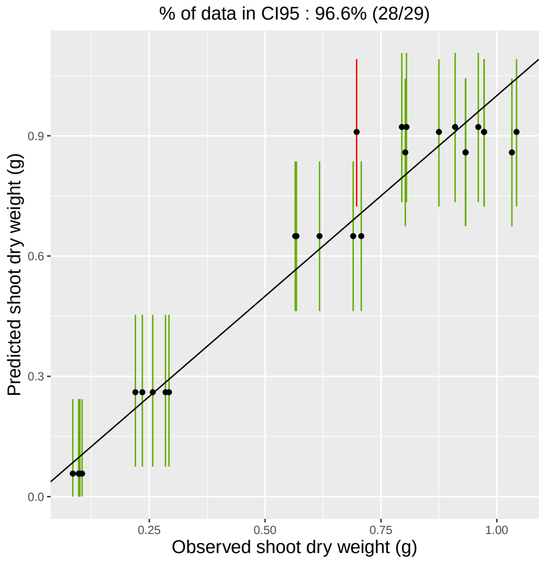

(b) Posterior predictive check (PPC)

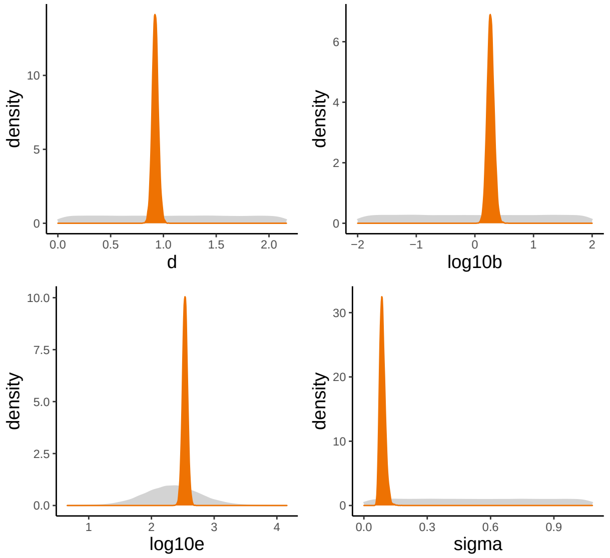

(c) Priors and posteriors

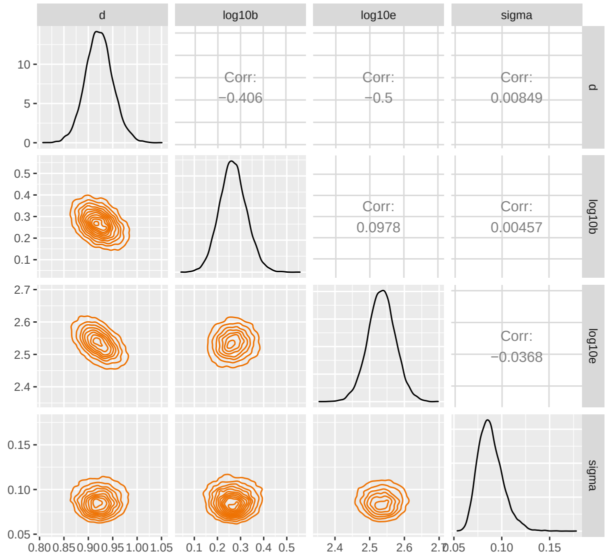

(d) Correlations between parameters

Figure 9: Dose-response curve (a), PPC (b), prior and posterior distributions (c) and correlations between parameters (d).

### Data set: ZEAMA_VV_weight

Table 10: Summary of parameter estimates for ZEAMA_VV_weight data set

| Parameter | median | Q2.5 | Q97.5 |
|---|---|---|---|
| b | 1.484 | 1.188 | 1.846 |
| d | 9.111 | 8.739 | 9.514 |
| e | 682.828 | 586.529 | 792.433 |
| sigma | 0.904 | 0.758 | 1.111 |

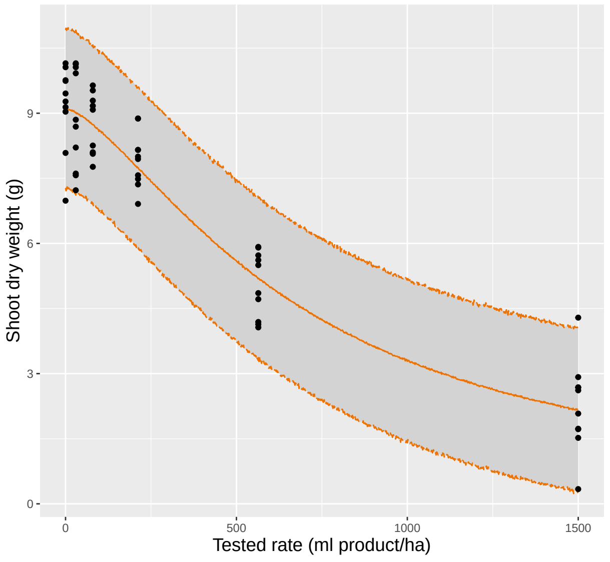

(a) Dose-response curve

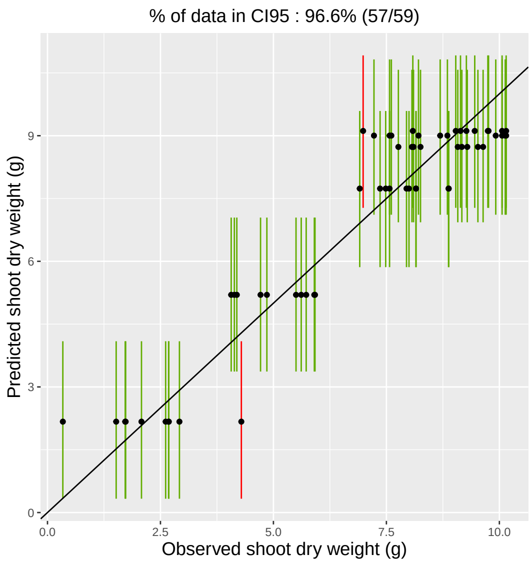

(b) Posterior predictive check (PPC)

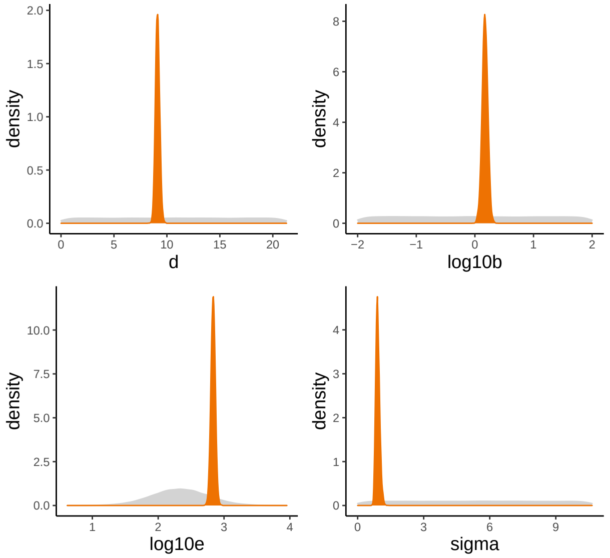

(c) Priors and posteriors

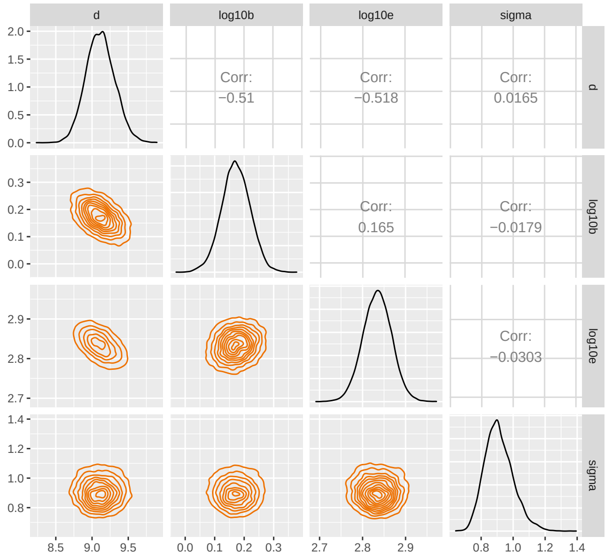

(d) Correlations between parameters

Figure 10: Dose-response curve (a), PPC (b), prior and posterior distributions (c) and correlations between parameters (d).
