## Supplementary material for "How to account for the uncertainty from standard toxicity tests in species sensitivity distributions: an example in non-target plants": Archive with reports on studies: report_SE_emergence.pdf

### Dose-response analyses

Study 4

Seedling Emergence test - emergence endpoint

25 June 2020

Contact:

---

This is a report which provides results on all performed dose-response analyses for the emergence endpoint of the Seedling Emergence test for study 4.

---

#### Data set: ALLCE_SE_emergence

Table 1: Summary of parameter estimates for ALLCE_SE_emergence data set

| Parameter | median | Q2.5 | Q97.5 |
|---|---|---|---|
| b | 28.304 | 3.314 | 94.325 |
| d | 0.741 | 0.649 | 0.823 |
| e | 8.737 | 7.038 | 16.469 |

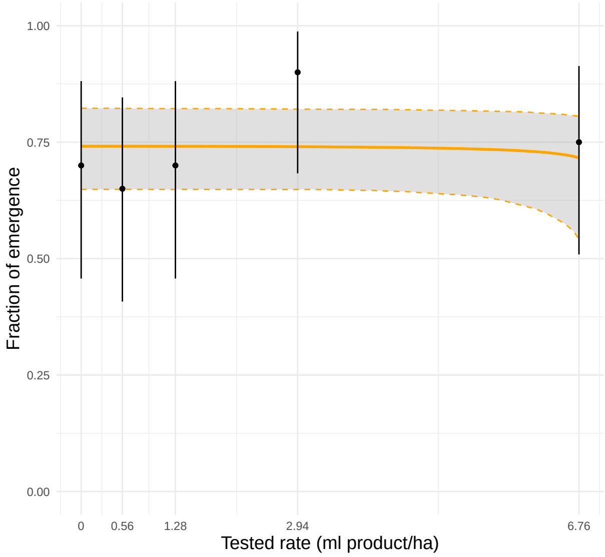

(a) Dose-response curve

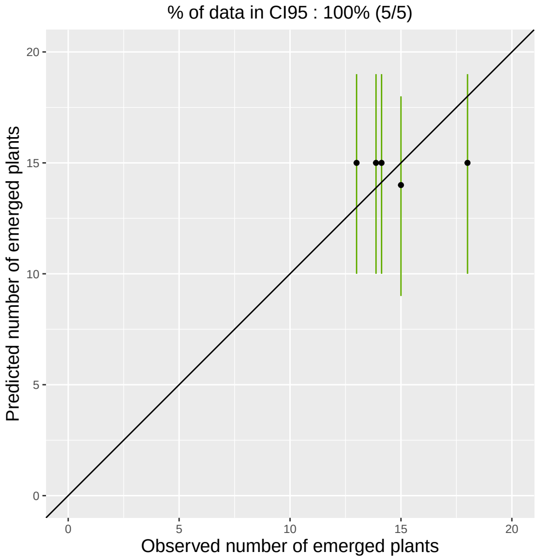

(b) Posterior predictive check (PPC)

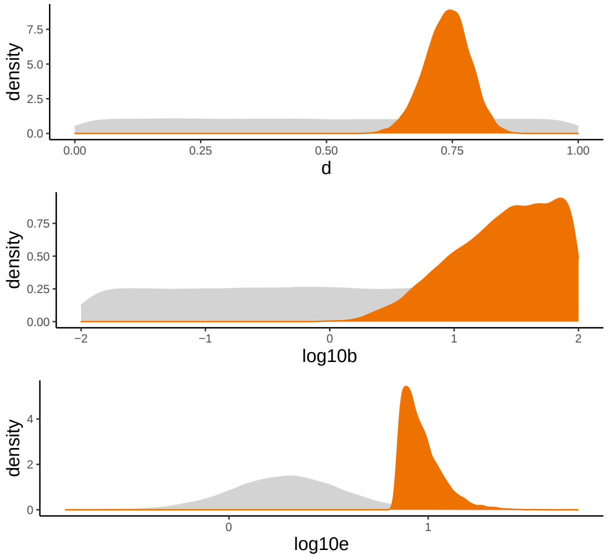

(c) Priors and posteriors

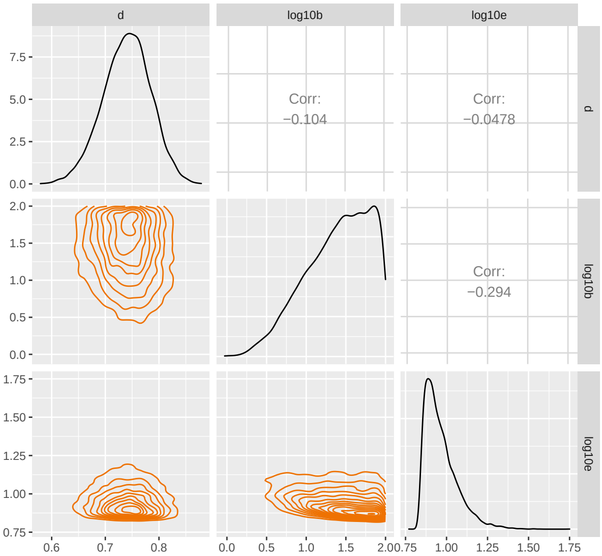

(d) Correlations between parameters

Figure 1: Dose-response curve (a), PPC (b), prior and posterior distributions (c) and correlations between parameters (d).

### Data set: AVESA_SE_emergence

Table 2: Summary of parameter estimates for AVESA_SE_emergence data set

| Parameter | median | Q2.5 | Q97.5 |
|---|---|---|---|
| b | 26.782 | 3.558 | 93.486 |
| d | 0.972 | 0.931 | 0.993 |
| e | 1405.380 | 1051.828 | 3219.676 |

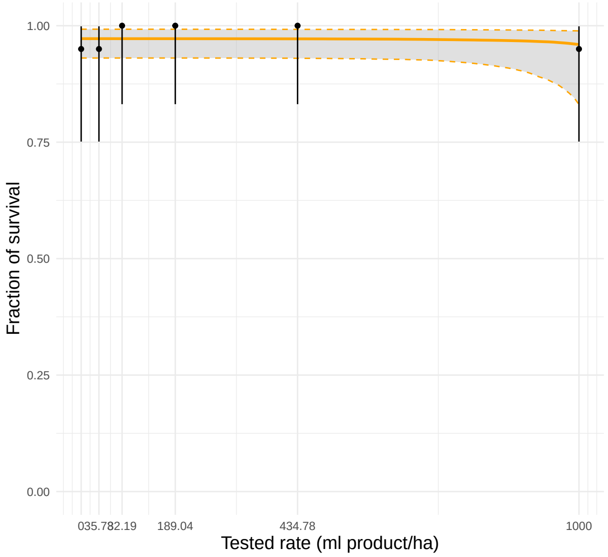

(a) Dose-response curve

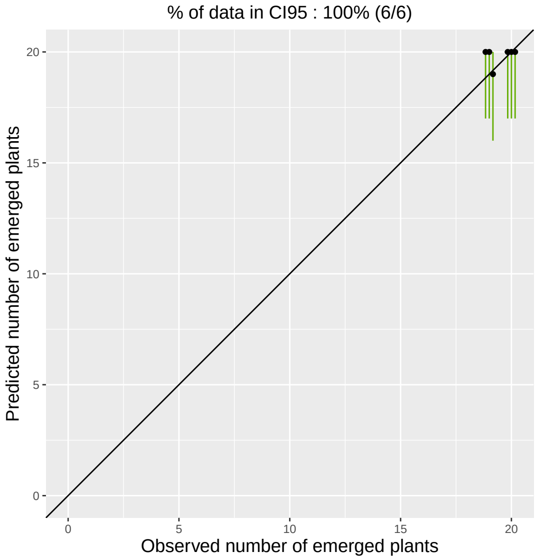

(b) Posterior predictive check (PPC)

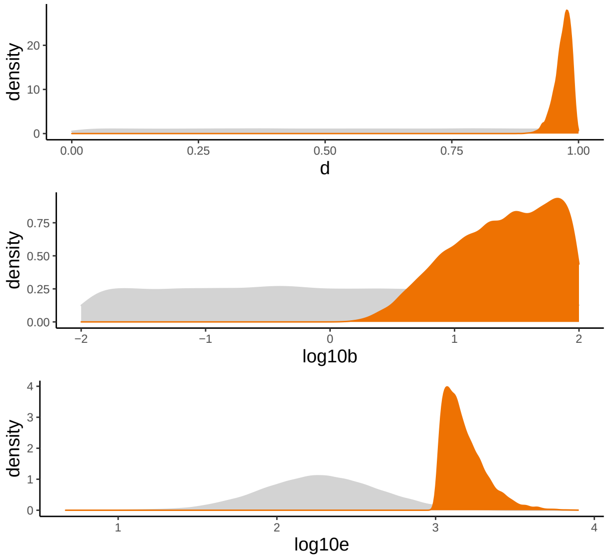

(c) Priors and posteriors

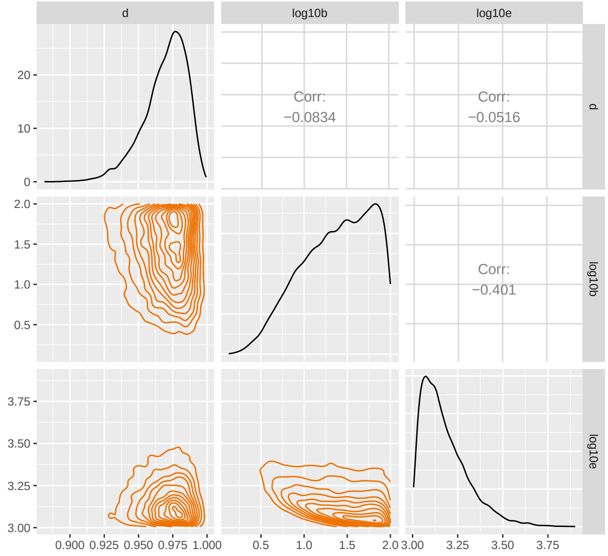

(d) Correlations between parameters

Figure 2: Dose-response curve (a), PPC (b), prior and posterior distributions (c) and correlations between parameters (d).

#### Data set: BEAVA_SE_emergence

Table 3: Summary of parameter estimates for BEAVA_SE_emergence data set

| Parameter | median | Q2.5 | Q97.5 |
|---|---|---|---|
| b | 5.403 | 1.157 | 77.886 |
| d | 0.918 | 0.847 | 0.972 |
| e | 246.873 | 192.773 | 535.733 |

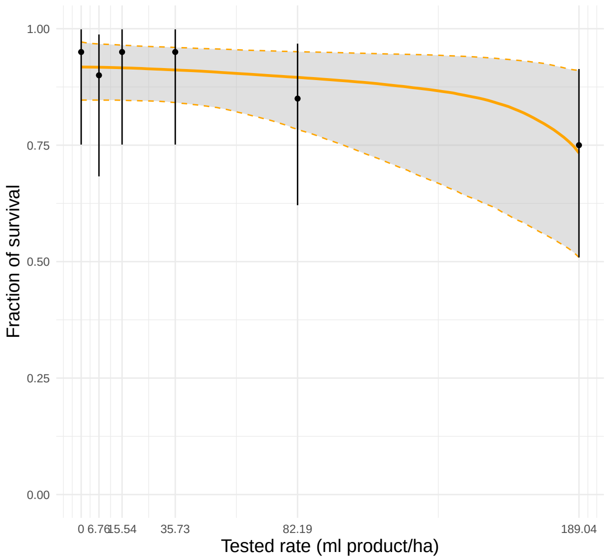

(a) Dose-response curve

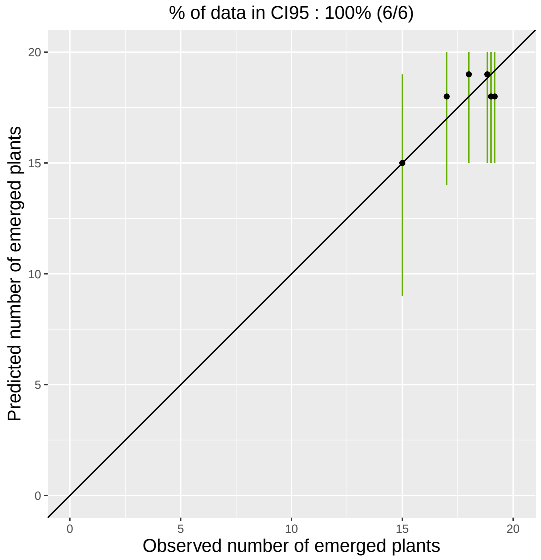

(b) Posterior predictive check (PPC)

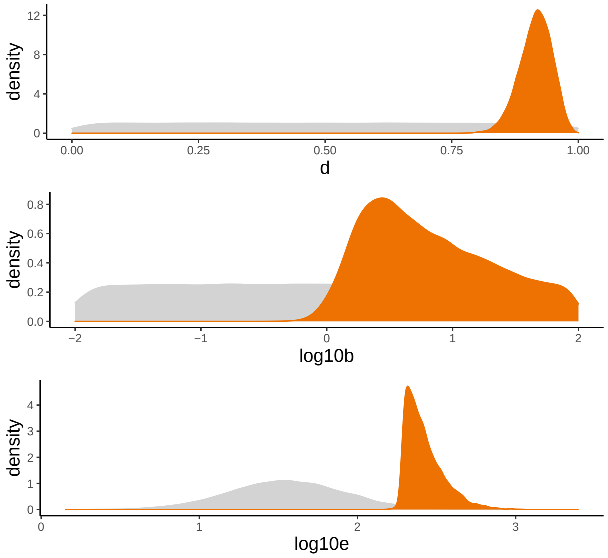

(c) Priors and posteriors

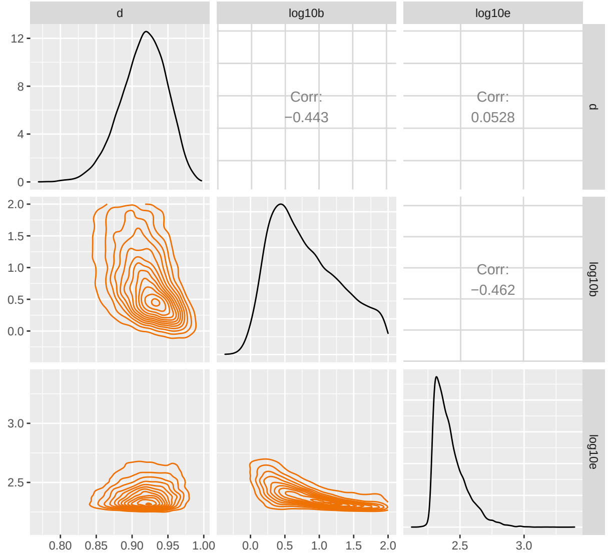

(d) Correlations between parameters

Figure 3: Dose-response curve (a), PPC (b), prior and posterior distributions (c) and correlations between parameters (d).

#### Data set: BRSNW_SE_emergence

Table 4: Summary of parameter estimates (parameter d is set to 1) for BRSNW_SE_emergence data set

| Parameter | median | Q2.5 | Q97.5 |
|---|---|---|---|
| b | 0.669 | 0.395 | 1.043 |
| e | 287.261 | 143.556 | 745.987 |

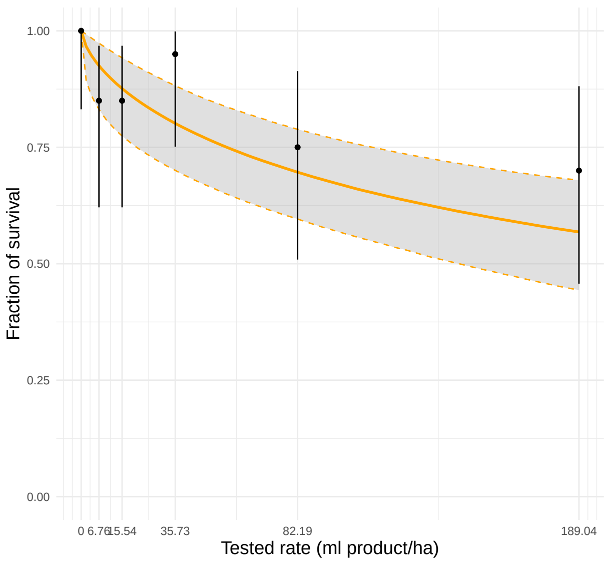

(a) Dose-response curve

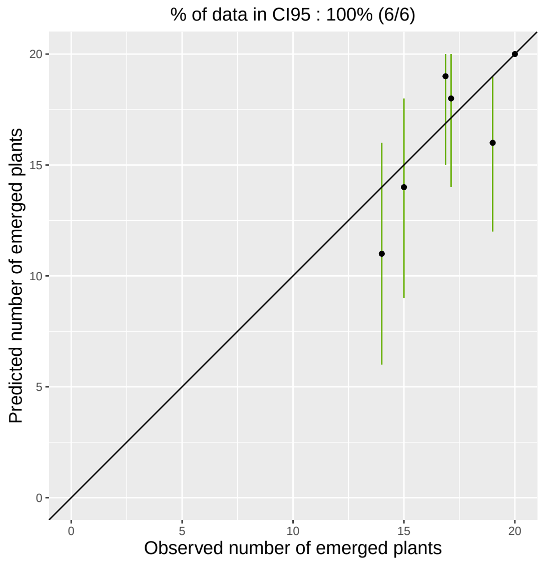

(b) Posterior predictive check (PPC)

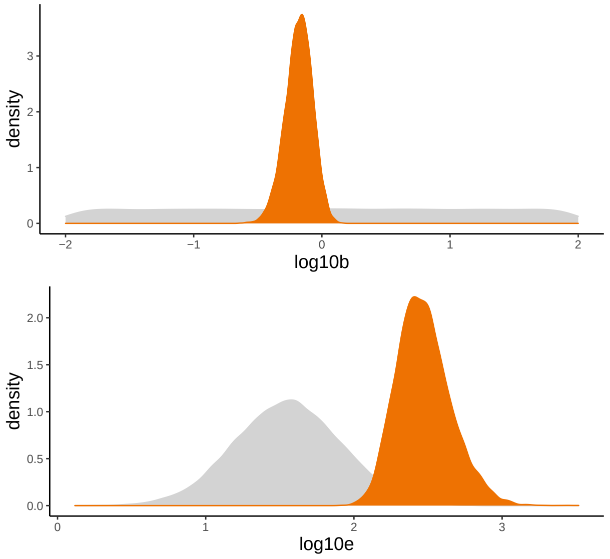

(c) Priors and posteriors

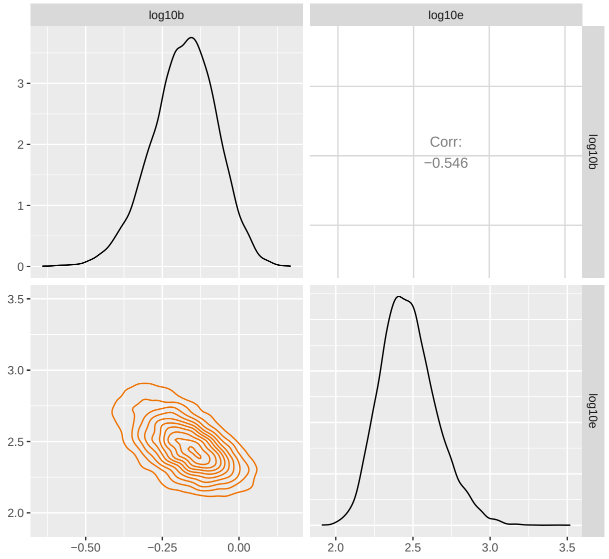

(d) Correlations between parameters

Figure 4: Dose-response curve (a), PPC (b), prior and posterior distributions (c) and correlations between parameters (d).

#### Data set: CUMSA_SE_emergence

Table 5: Summary of parameter estimates (parameter d is set to 1) for CUMSA_SE_emergence data set

| Parameter | median | Q2.5 | Q97.5 |
|---|---|---|---|
| b | 1.295 | 0.815 | 2.063 |
| e | 1344.514 | 705.783 | 3354.320 |

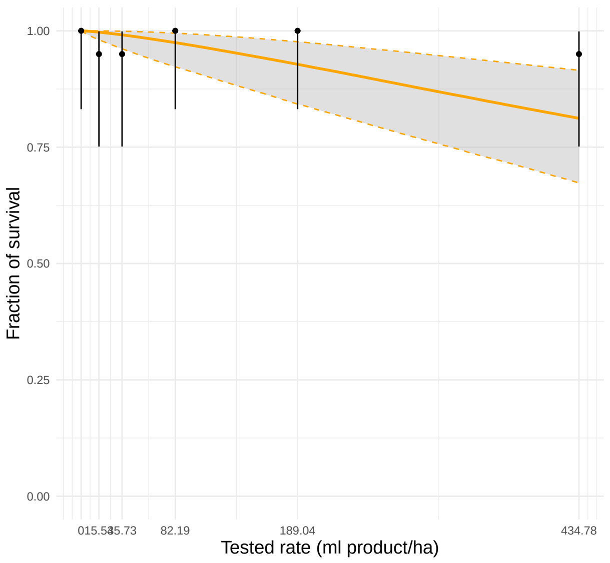

(a) Dose-response curve

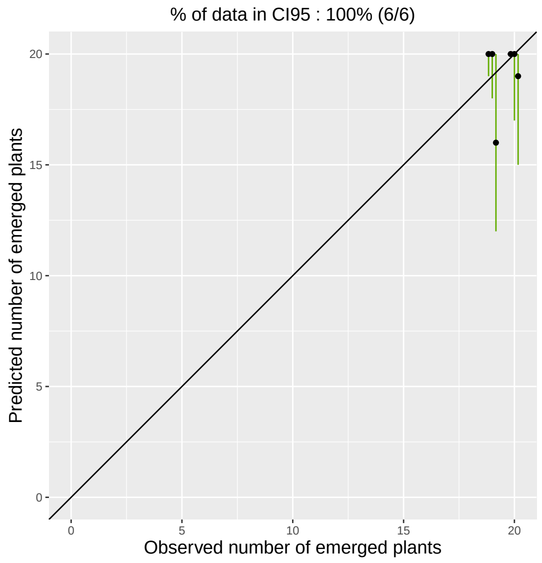

(b) Posterior predictive check (PPC)

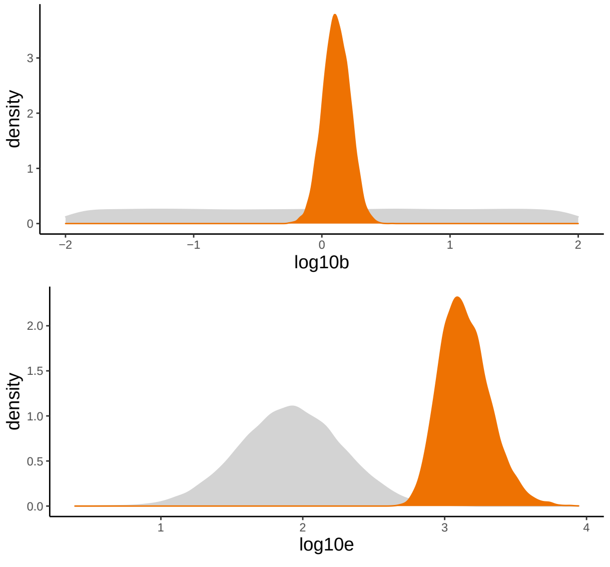

(c) Priors and posteriors

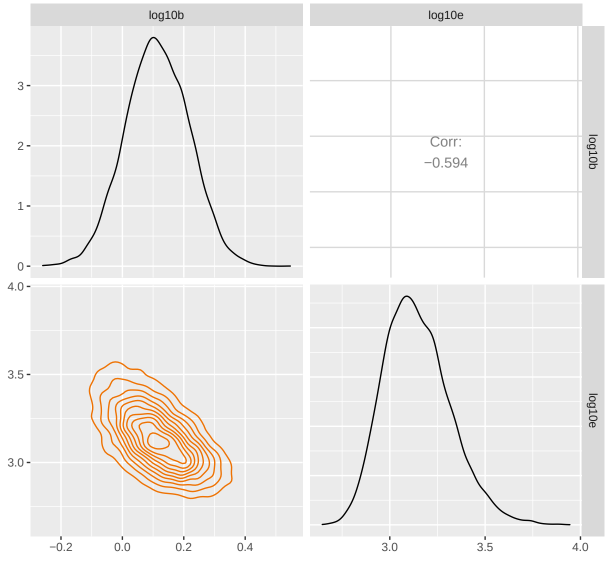

(d) Correlations between parameters

Figure 5: Dose-response curve (a), PPC (b), prior and posterior distributions (c) and correlations between parameters (d).

### Data set: GLXMA_SE_emergence

Table 6: Summary of parameter estimates for GLXMA_SE_emergence data set

| Parameter | median | Q2.5 | Q97.5 |
|---|---|---|---|
| b | 28.030 | 3.726 | 94.461 |
| d | 0.963 | 0.919 | 0.988 |
| e | 619.582 | 459.193 | 1410.823 |

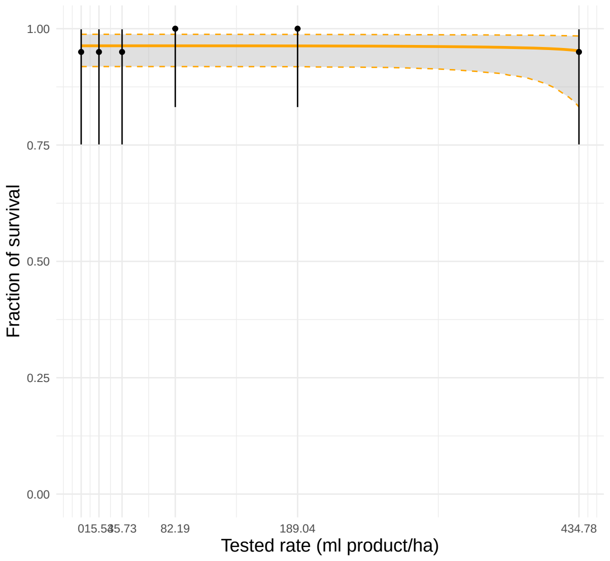

(a) Dose-response curve

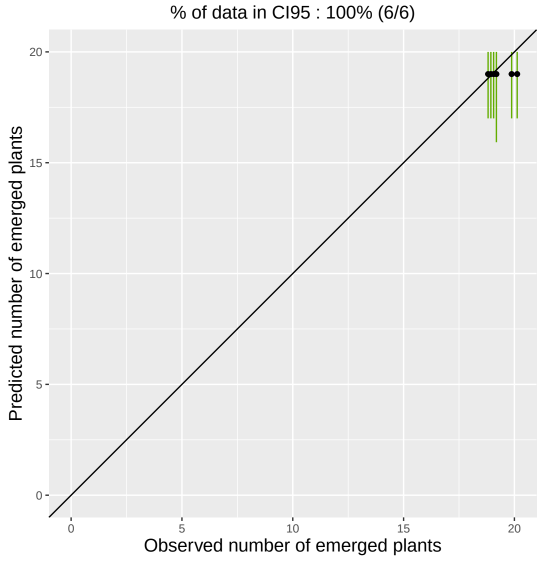

(b) Posterior predictive check (PPC)

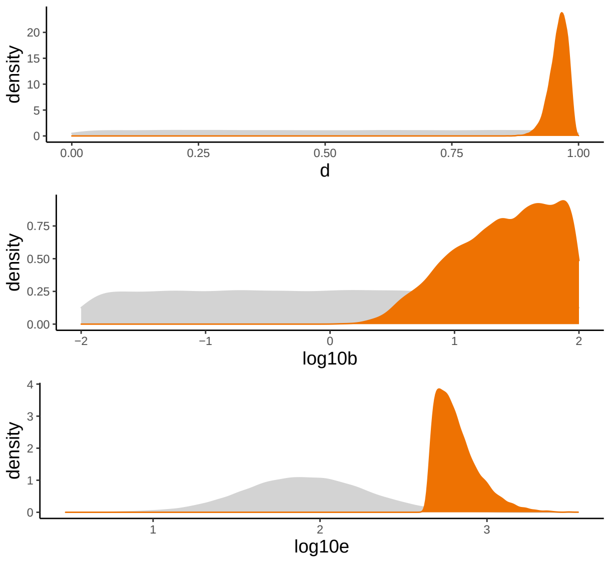

(c) Priors and posteriors

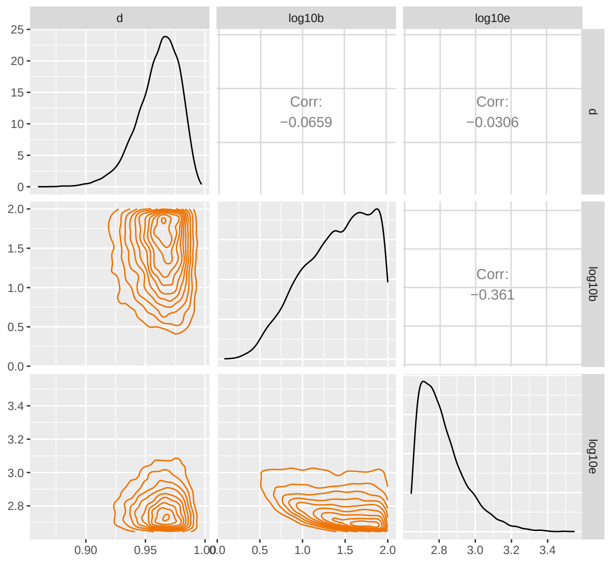

(d) Correlations between parameters

Figure 6: Dose-response curve (a), PPC (b), prior and posterior distributions (c) and correlations between parameters (d).

#### Data set: HELAN_SE_emergence

Table 7: Summary of parameter estimates (parameter d is set to 1) for HELAN_SE_emergence data set

| Parameter | median | Q2.5 | Q97.5 |
|---|---|---|---|
| b | 1.302 | 0.791 | 2.103 |
| e | 670.562 | 407.864 | 1441.417 |

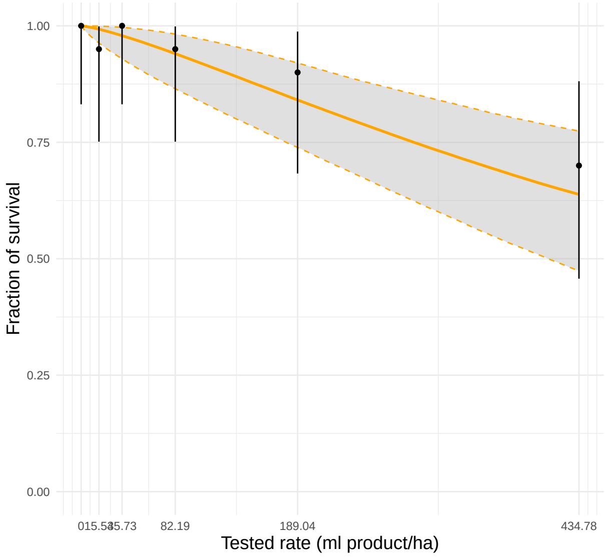

(a) Dose-response curve

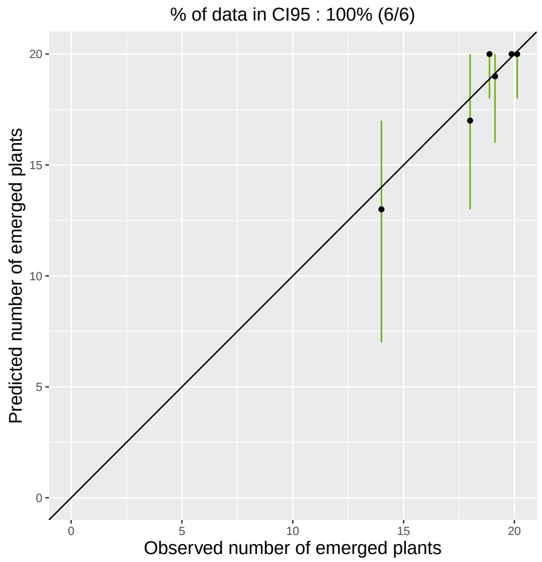

(b) Posterior predictive check (PPC)

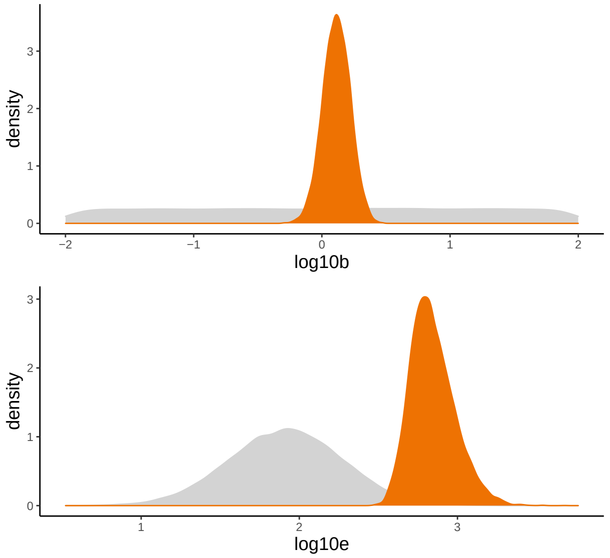

(c) Priors and posteriors

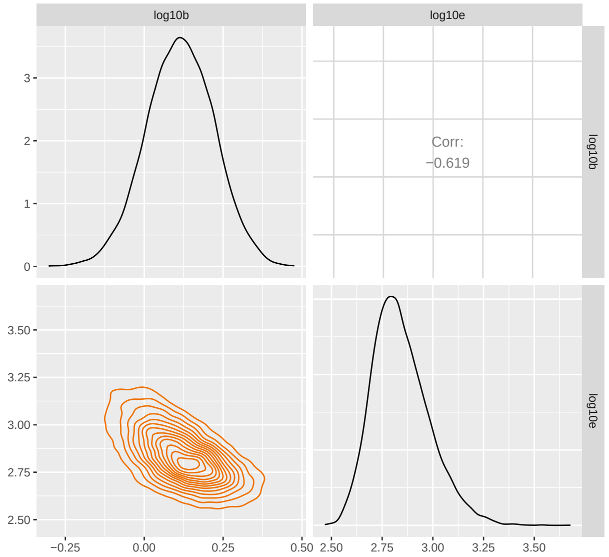

(d) Correlations between parameters

Figure 7: Dose-response curve (a), PPC (b), prior and posterior distributions (c) and correlations between parameters (d).

#### Data set: LYPES_SE_emergence

Table 8: Summary of parameter estimates for LYPES_SE_emergence data set

| Parameter | median | Q2.5 | Q97.5 |
|---|---|---|---|
| b | 6.959 | 1.423 | 75.728 |
| d | 0.893 | 0.820 | 0.948 |
| e | 532.938 | 439.202 | 1124.719 |

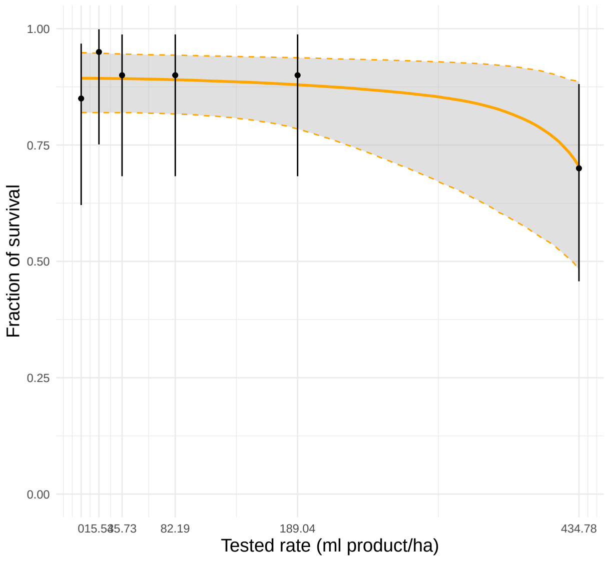

(a) Dose-response curve

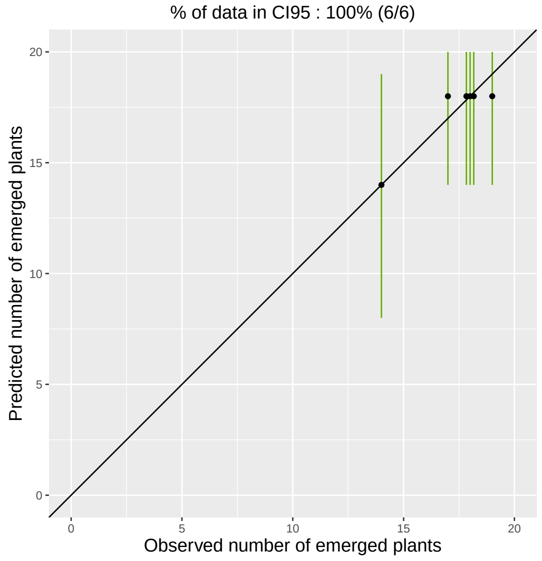

(b) Posterior predictive check (PPC)

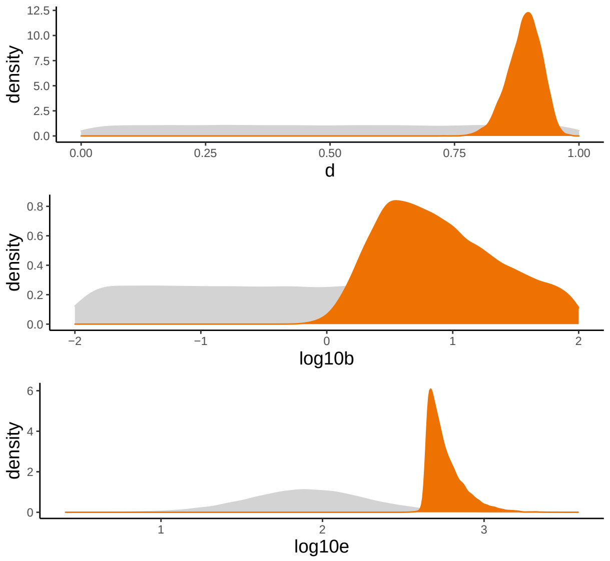

(c) Priors and posteriors

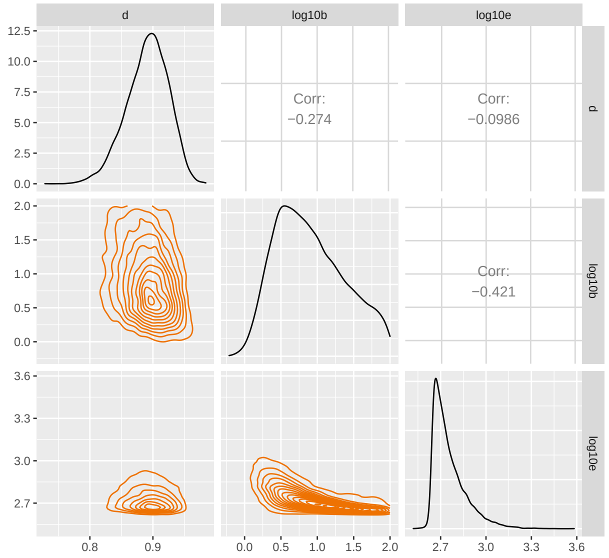

(d) Correlations between parameters

Figure 8: Dose-response curve (a), PPC (b), prior and posterior distributions (c) and correlations between parameters (d).

#### Data set: TRZAW_SE_emergence

Table 9: Summary of parameter estimates for TRZAW_SE_emergence data set

| Parameter | median | Q2.5 | Q97.5 |
|---|---:|---:|---:|
| b | 30.012 | 3.919 | 95.029 |
| d | 0.947 | 0.898 | 0.977 |
| e | 1420.400 | 1061.659 | 3185.006 |

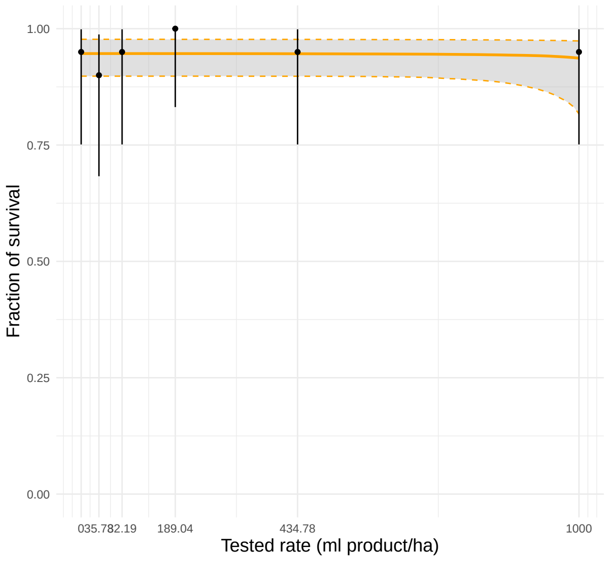

(a) Dose-response curve

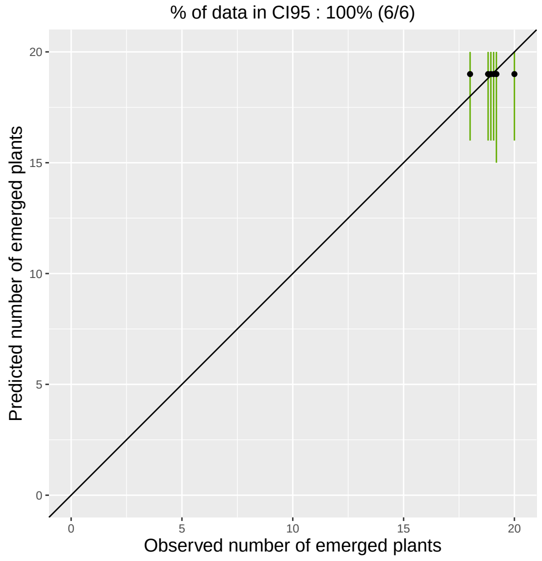

(b) Posterior predictive check (PPC)

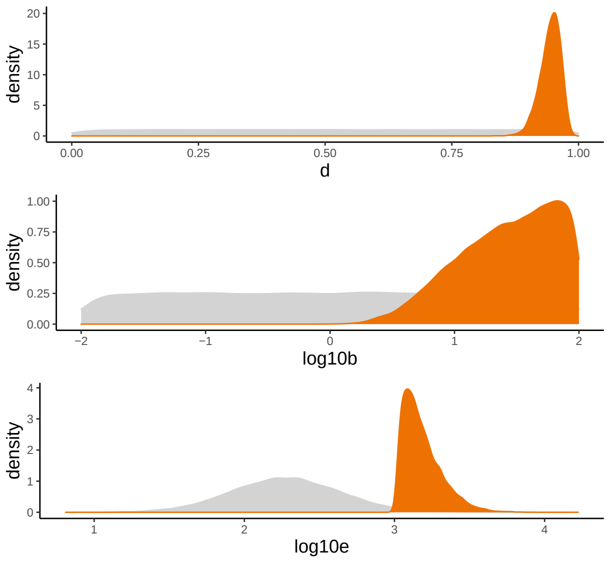

(c) Priors and posteriors

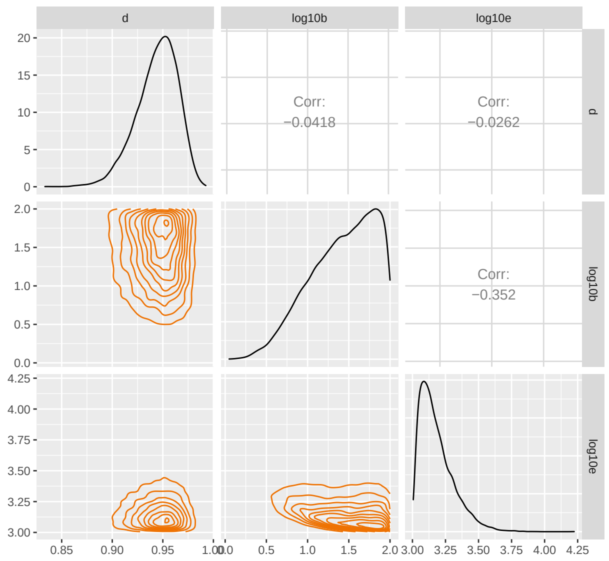

(d) Correlations between parameters

Figure 9: Dose-response curve (a), PPC (b), prior and posterior distributions (c) and correlations between parameters (d).

#### Data set: ZEAMA_SE_emergence

Table 10: Summary of parameter estimates (parameter d is set to 1) for ZEAMA_SE_emergence data set

| Parameter | median | Q2.5 | Q97.5 |
|---|---|---|---|
| b | 1.518 | 0.932 | 2.524 |
| e | 3051.785 | 1639.983 | 7638.387 |

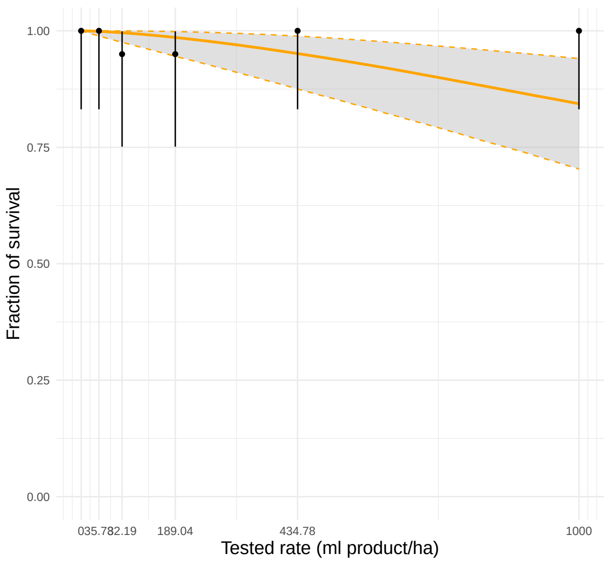

(a) Dose-response curve

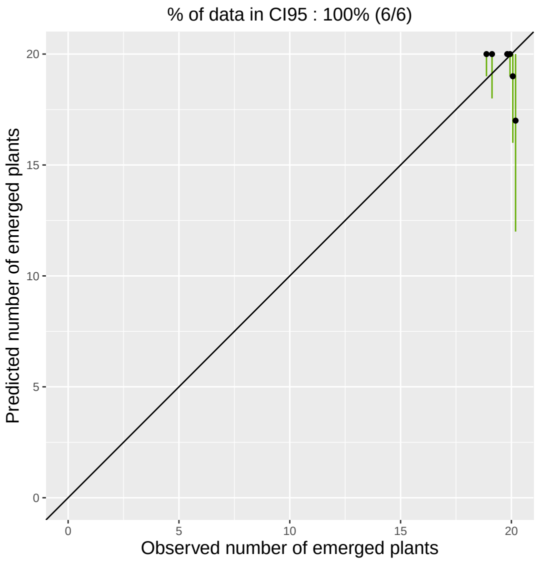

(b) Posterior predictive check (PPC)

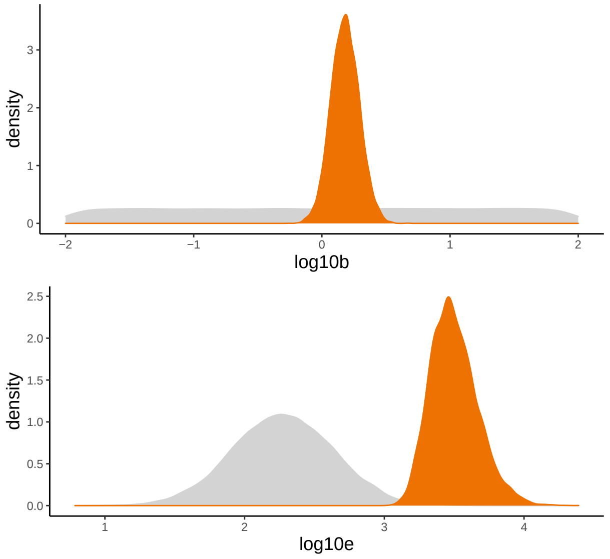

(c) Priors and posteriors

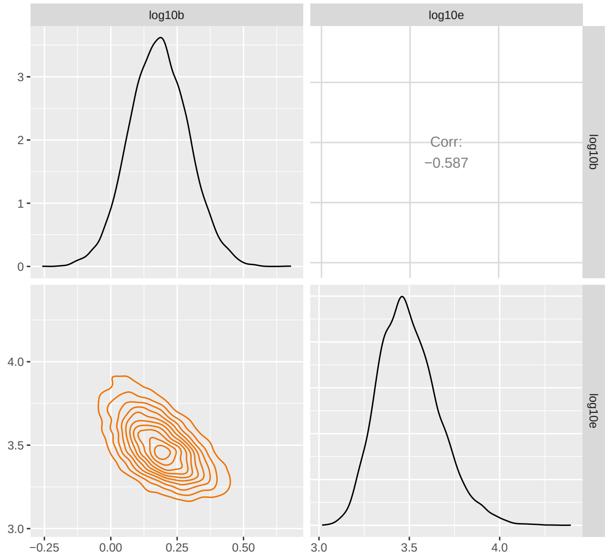

(d) Correlations between parameters

Figure 10: Dose-response curve (a), PPC (b), prior and posterior distributions (c) and correlations between parameters (d).
