## Supplementary material for "How to account for the uncertainty from standard toxicity tests in species sensitivity distributions: an example in non-target plants": Archive with reports on studies: report_SE_survival.pdf

### Dose-response analyses

Study 4

Seedling Emergence test - survival endpoint

25 June 2020

Contact:

---

This is a report which provides results on all performed dose-response analyses for the survival endpoint of the Seedling Emergence test for study 4.

---

#### Data set: ALLCE_SE_survival

Table 1: Summary of parameter estimates for ALLCE_SE_survival data set

| Parameter | median | Q2.5 | Q97.5 |
|---|---|---|---|
| b | 26.419 | 2.900 | 94.184 |
| d | 0.890 | 0.805 | 0.951 |
| e | 8.767 | 7.024 | 16.151 |

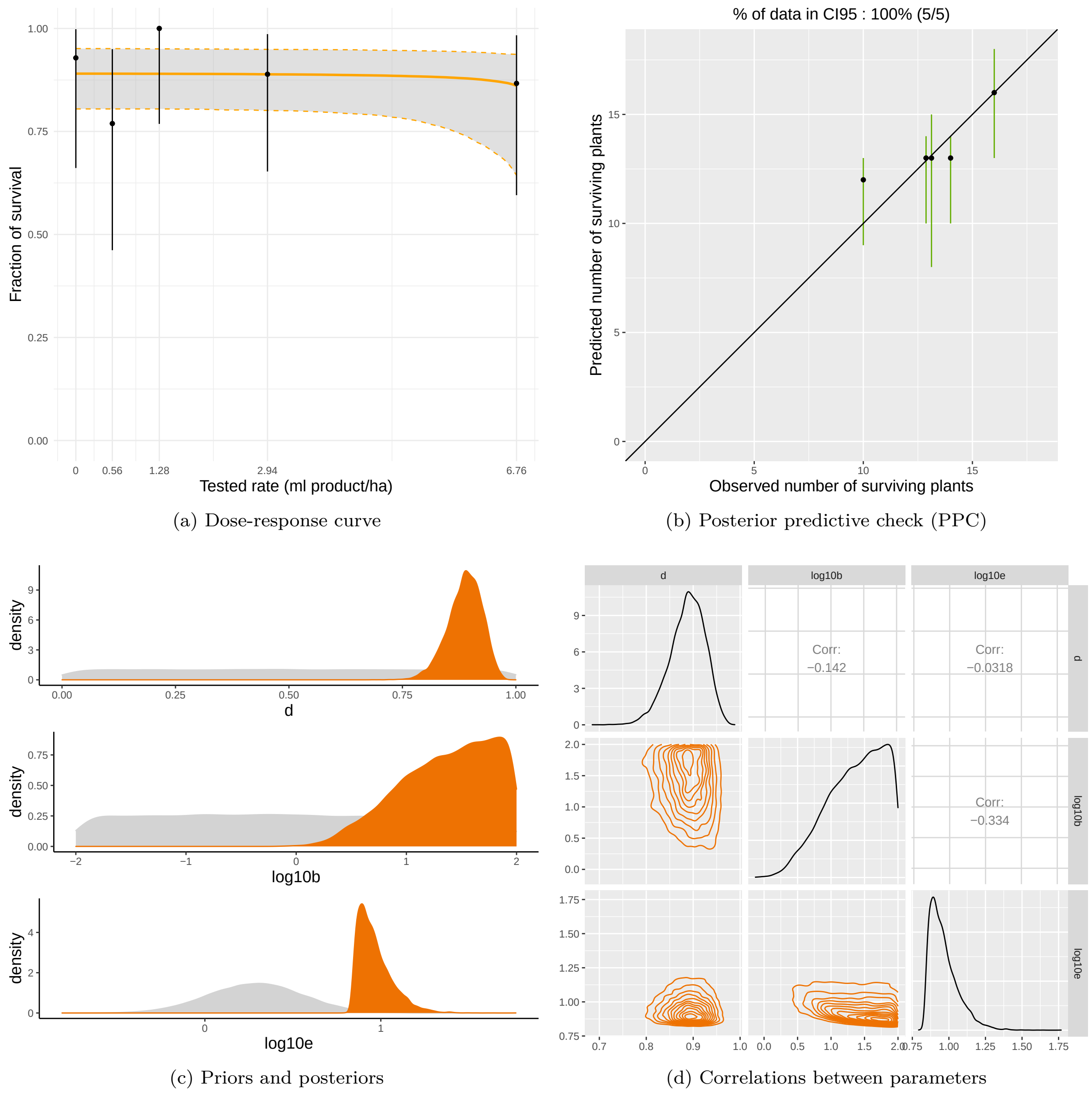

(a) Dose-response curve

(b) Posterior predictive check (PPC)

(c) Priors and posteriors

(d) Correlations between parameters

Figure 1: Dose-response curve (a), PPC (b), prior and posterior distributions (c) and correlations between parameters (d).

#### Data set: AVESA_SE_survival

Table 2: Summary of parameter estimates (parameter d is set to 1) for AVESA_SE_survival data set

| Parameter | median | Q2.5 | Q97.5 |
|---|---:|---:|---:|
| b | 7.270 | 2.357 | 60.412 |
| e | 1029.937 | 908.490 | 1393.721 |

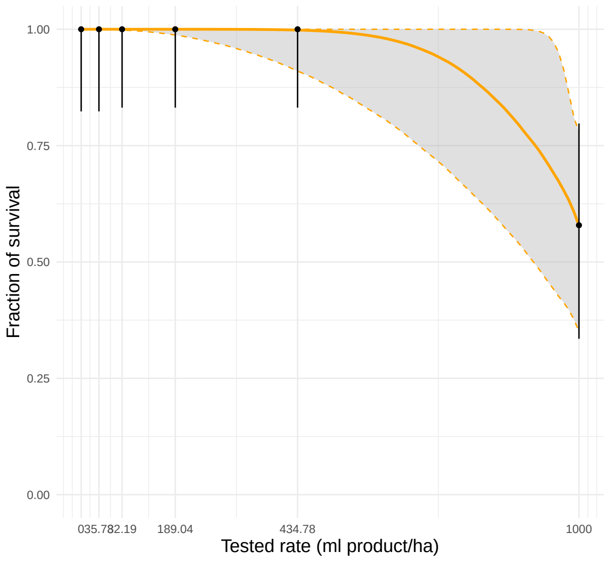

(a) Dose-response curve

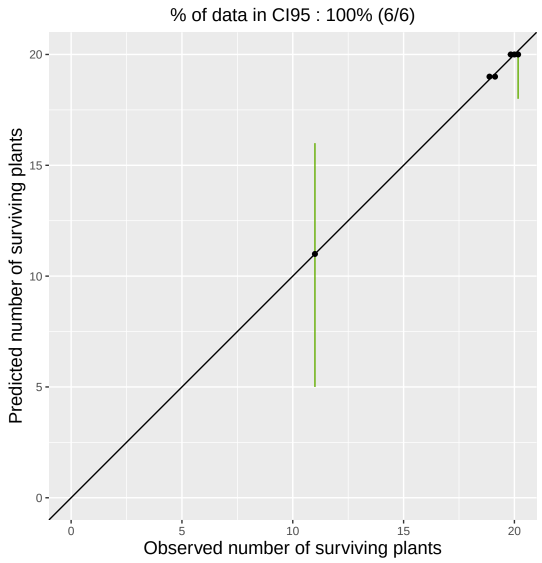

(b) Posterior predictive check (PPC)

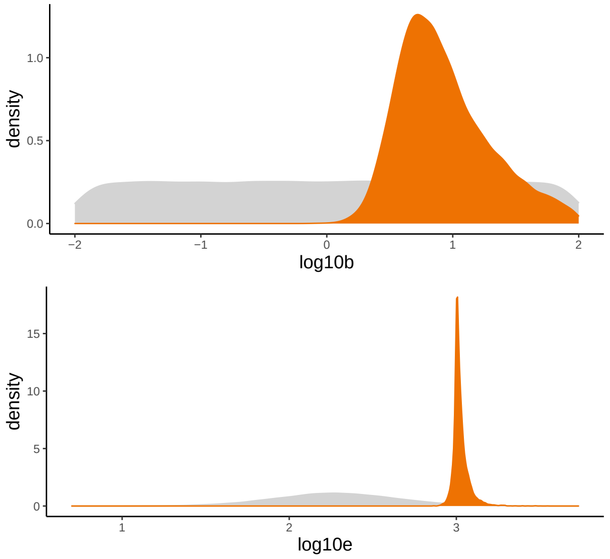

(c) Priors and posteriors

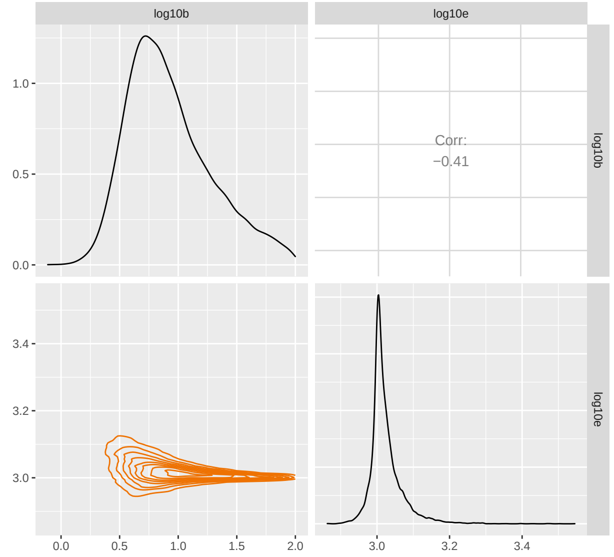

(d) Correlations between parameters

Figure 2: Dose-response curve (a), PPC (b), prior and posterior distributions (c) and correlations between parameters (d).

#### Data set: BEAVA_SE_survival

Table 3: Summary of parameter estimates (parameter d is set to 1) for BEAVA_SE_survival data set

| Parameter | median | Q2.5 | Q97.5 |
|---|---|---|---|
| b | 1.745 | 1.067 | 2.630 |
| e | 91.198 | 65.274 | 137.672 |

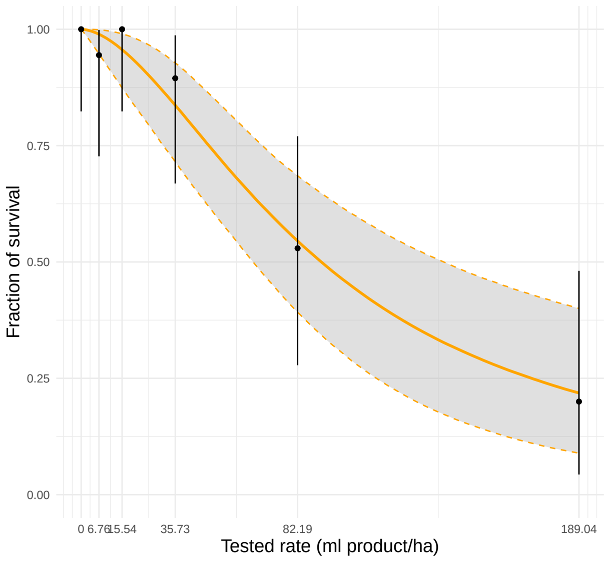

(a) Dose-response curve

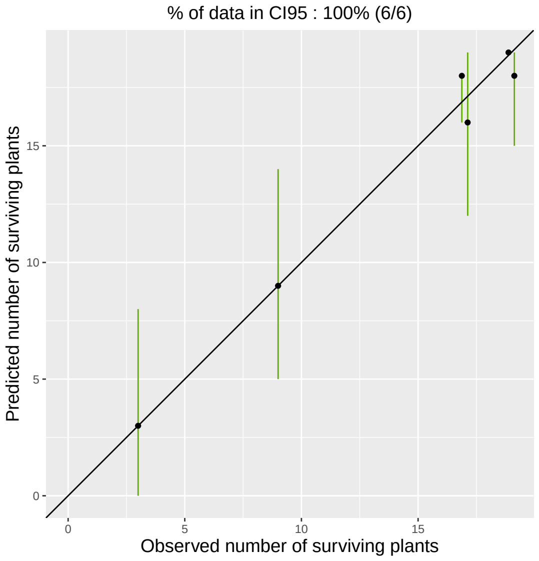

(b) Posterior predictive check (PPC)

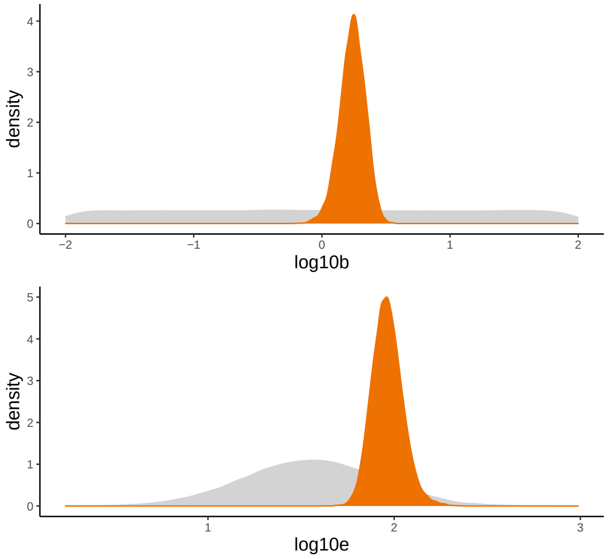

(c) Priors and posteriors

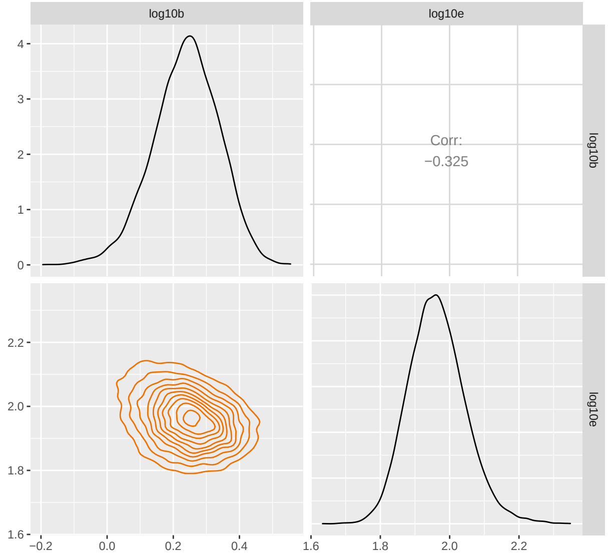

(d) Correlations between parameters

Figure 3: Dose-response curve (a), PPC (b), prior and posterior distributions (c) and correlations between parameters (d).

#### Data set: BRSNW_SE_survival

Table 4: Summary of parameter estimates (parameter d is set to 1) for BRSNW_SE_survival data set

| Parameter | median | Q2.5 | Q97.5 |
|---|---|---|---|
| b | 2.465 | 1.501 | 3.825 |
| e | 86.435 | 65.437 | 119.943 |

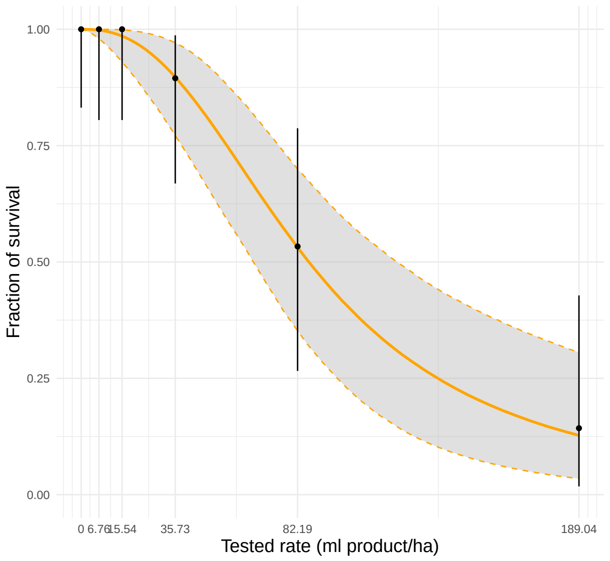

(a) Dose-response curve

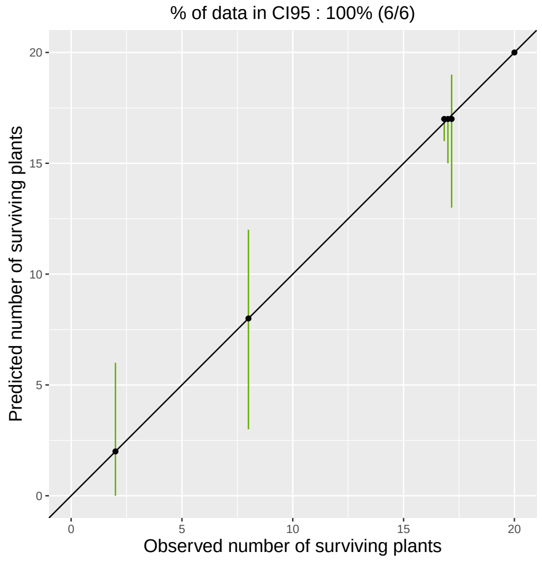

(b) Posterior predictive check (PPC)

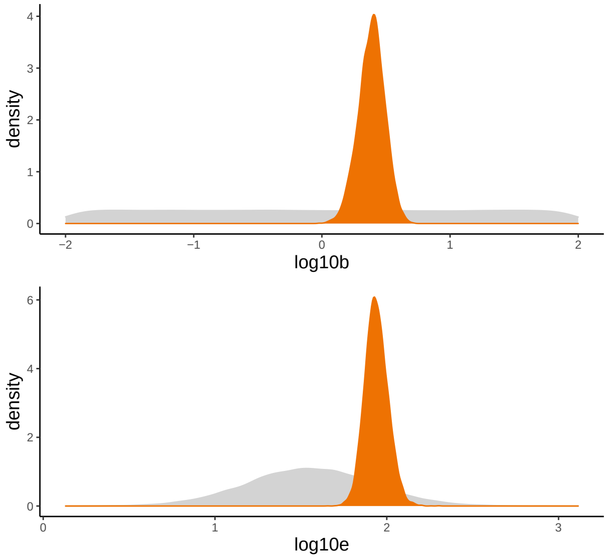

(c) Priors and posteriors

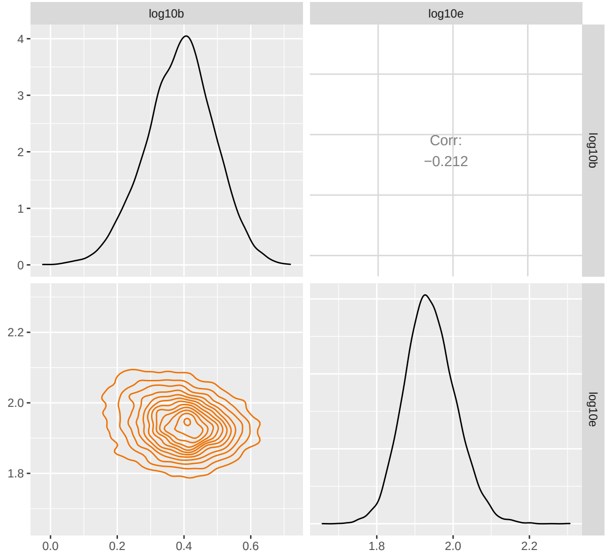

(d) Correlations between parameters

Figure 4: Dose-response curve (a), PPC (b), prior and posterior distributions (c) and correlations between parameters (d).

#### Data set: CUMSA_SE_survival

Table 5: Summary of parameter estimates (parameter d is set to 1) for CUMSA_SE_survival data set

| Parameter | median | Q2.5 | Q97.5 |
|---|---|---|---|
| b | 1.277 | 0.788 | 2.042 |
| e | 1180.950 | 628.530 | 2904.809 |

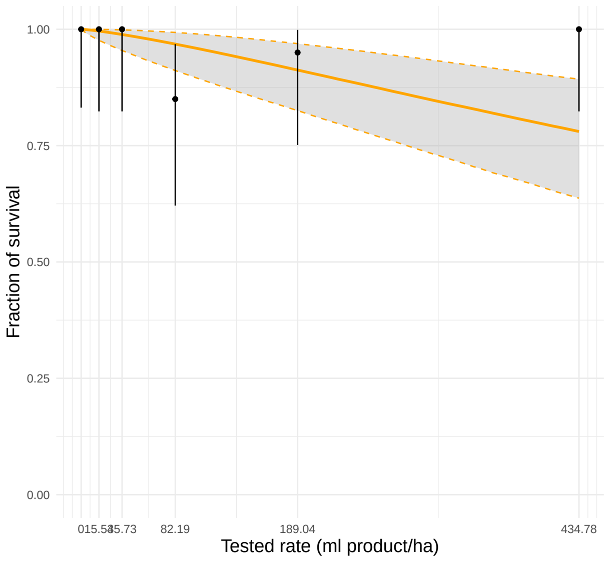

(a) Dose-response curve

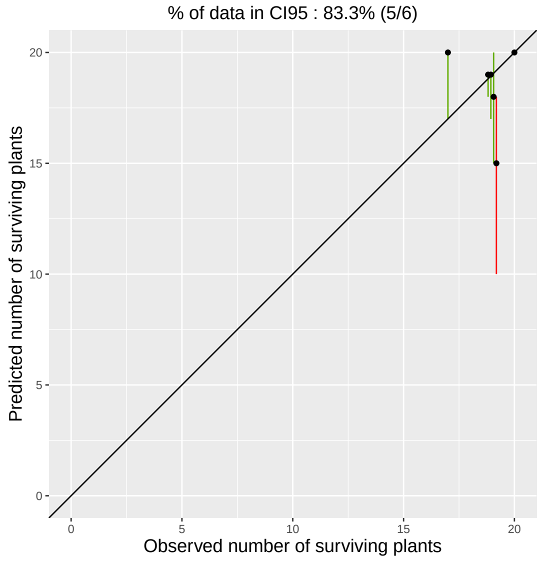

(b) Posterior predictive check (PPC)

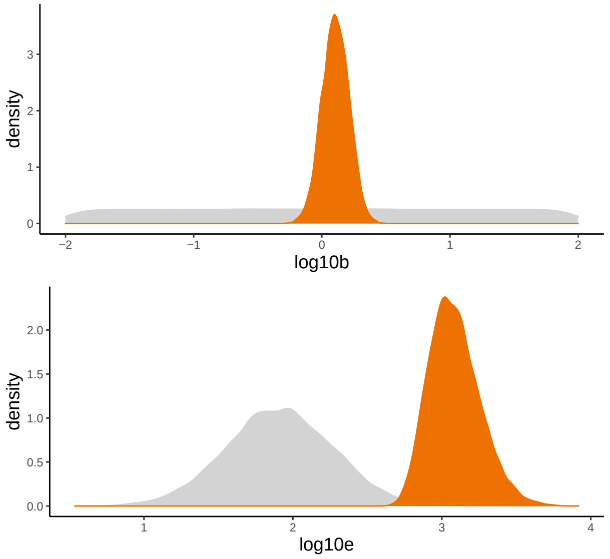

(c) Priors and posteriors

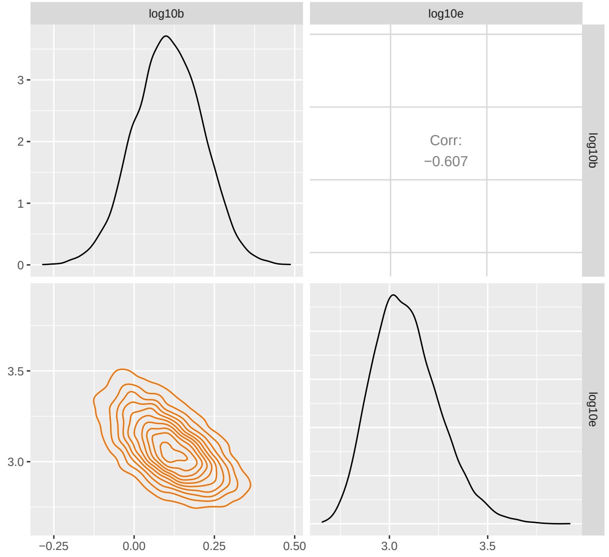

(d) Correlations between parameters

Figure 5: Dose-response curve (a), PPC (b), prior and posterior distributions (c) and correlations between parameters (d).

#### Data set: GLXMA_SE_survival

Table 6: Summary of parameter estimates (parameter d is set to 1) for GLXMA_SE_survival data set

| Parameter | median | Q2.5 | Q97.5 |
|---|---|---|---|
| b | 3.064 | 1.534 | 6.142 |
| e | 498.272 | 385.882 | 797.662 |

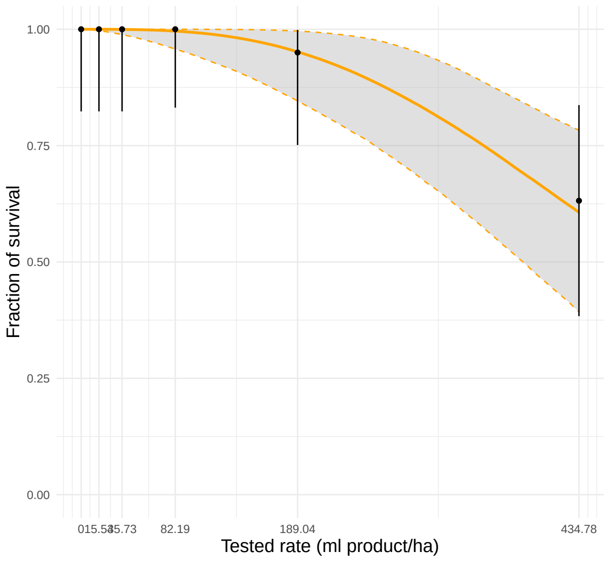

(a) Dose-response curve

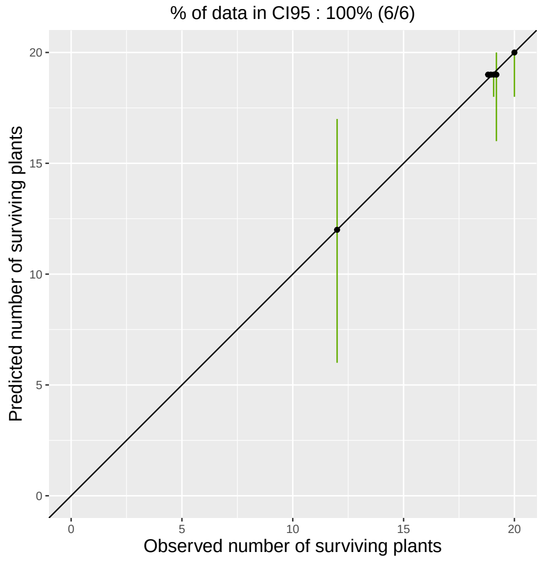

(b) Posterior predictive check (PPC)

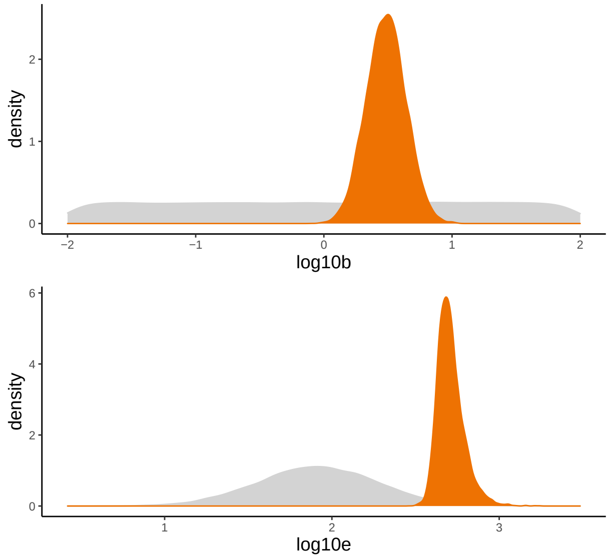

(c) Priors and posteriors

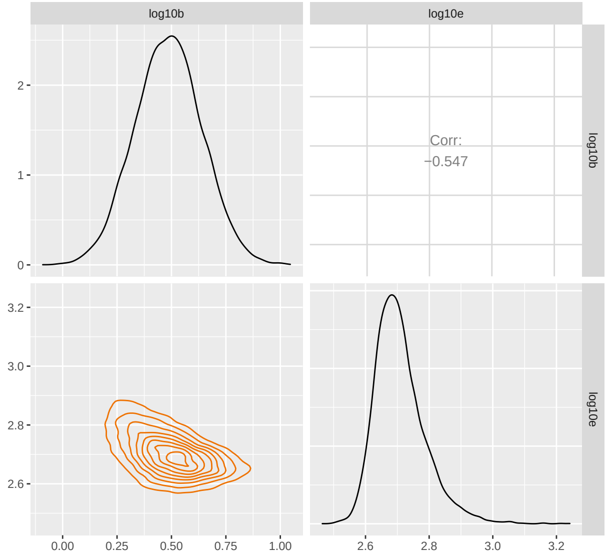

(d) Correlations between parameters

Figure 6: Dose-response curve (a), PPC (b), prior and posterior distributions (c) and correlations between parameters (d).

#### Data set: HELAN_SE_survival

Table 7: Summary of parameter estimates (parameter d is set to 1) for HELAN_SE_survival data set

| Parameter | median | Q2.5 | Q97.5 |
|---|---|---|---|
| b | 2.792 | 1.716 | 4.396 |
| e | 201.733 | 156.997 | 268.471 |

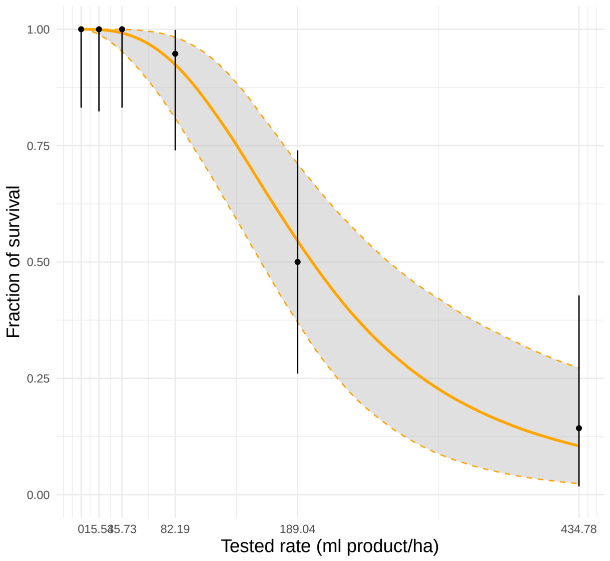

(a) Dose-response curve

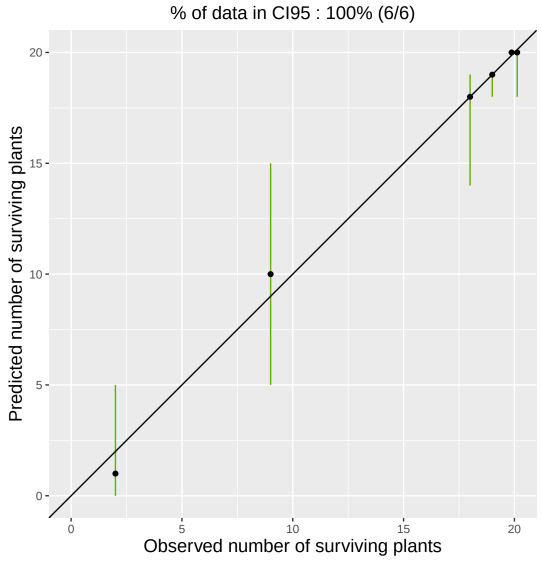

(b) Posterior predictive check (PPC)

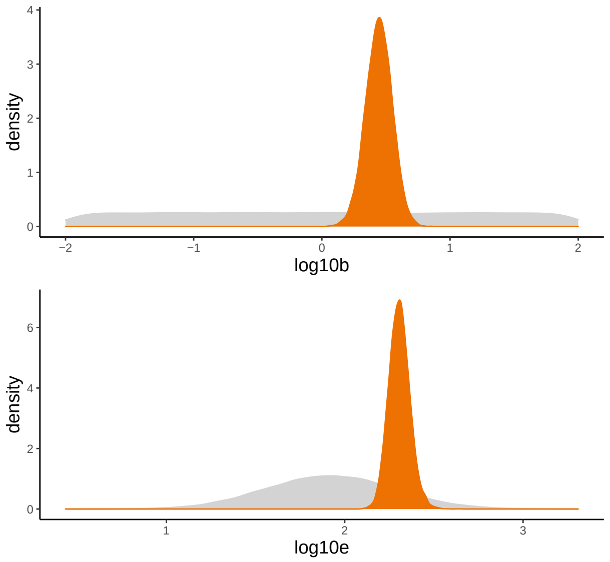

(c) Priors and posteriors

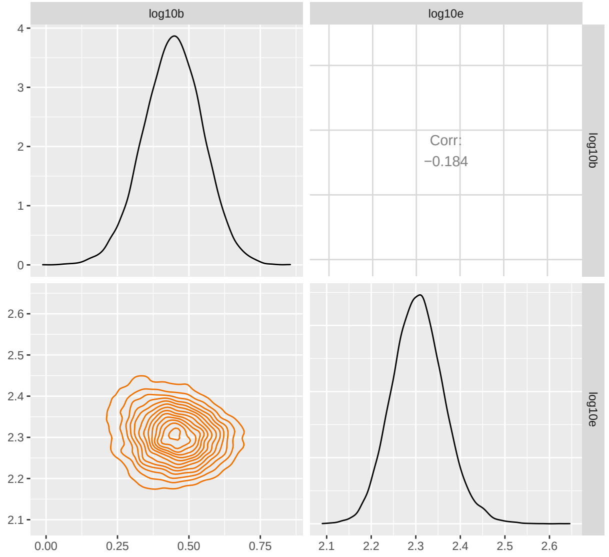

(d) Correlations between parameters

Figure 7: Dose-response curve (a), PPC (b), prior and posterior distributions (c) and correlations between parameters (d).

#### Data set: LYPES_SE_survival

Table 8: Summary of parameter estimates (parameter d is set to 1) for LYPES_SE_survival data set

| Parameter | median | Q2.5 | Q97.5 |
|---|---|---|---|
| b | 1.814 | 1.056 | 2.866 |
| e | 281.458 | 197.889 | 459.516 |

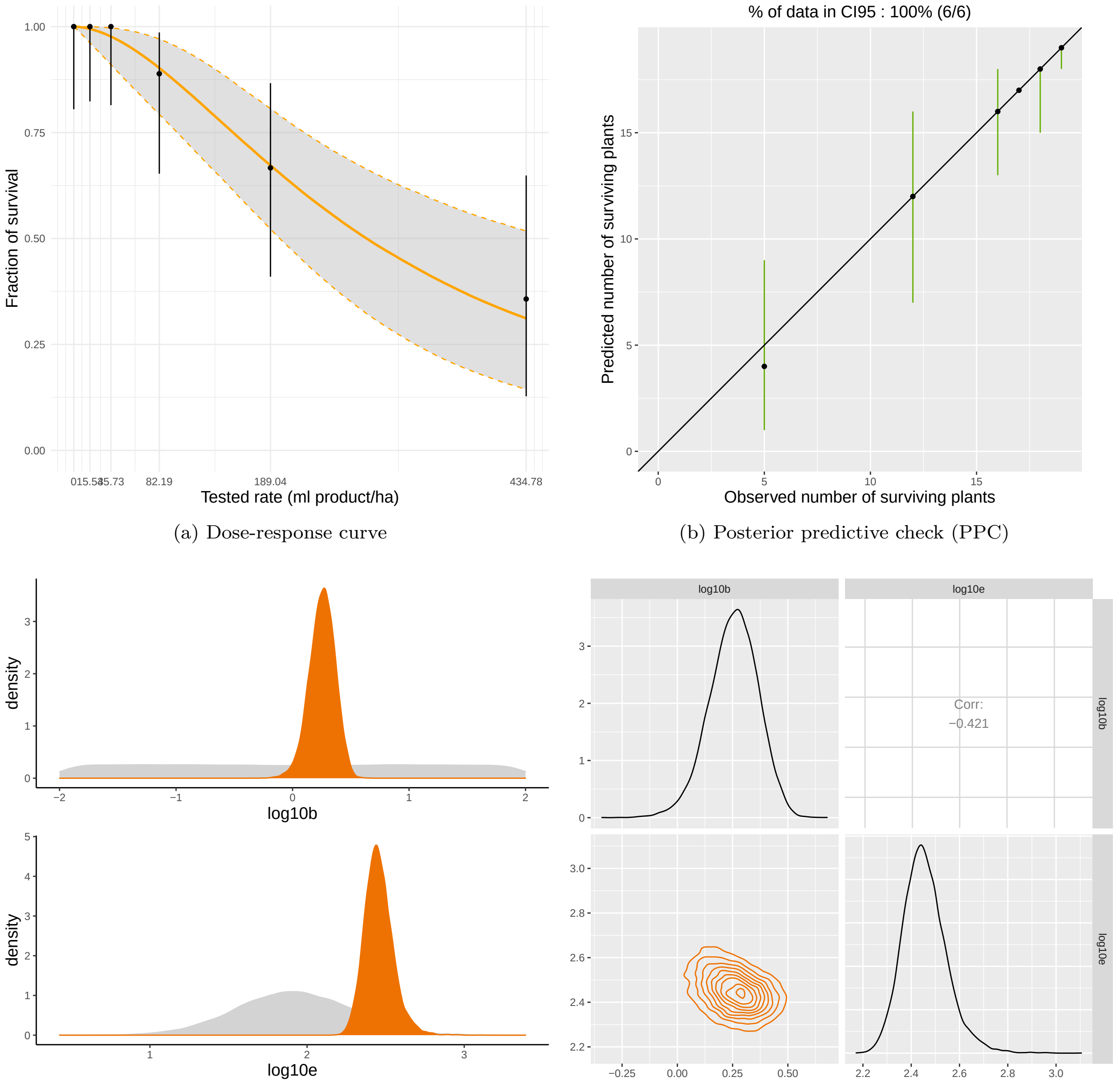

(a) Dose-response curve

(b) Posterior predictive check (PPC)

(c) Priors and posteriors

(d) Correlations between parameters

Figure 8: Dose-response curve (a), PPC (b), prior and posterior distributions (c) and correlations between parameters (d).

#### Data set: TRZAW_SE_survival

Table 9: Summary of parameter estimates (parameter d is set to 1) for TRZAW_SE_survival data set

| Parameter | median | Q2.5 | Q97.5 |
|---|---|---|---|
| b | 33.857 | 4.806 | 94.821 |
| e | 1461.708 | 1072.712 | 3253.785 |

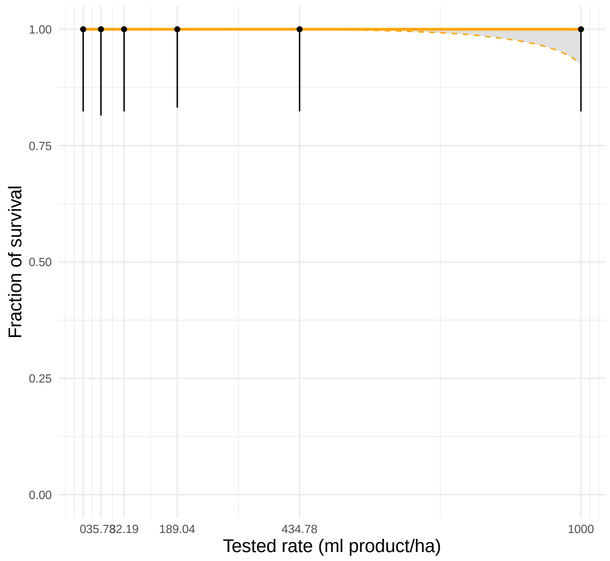

(a) Dose-response curve

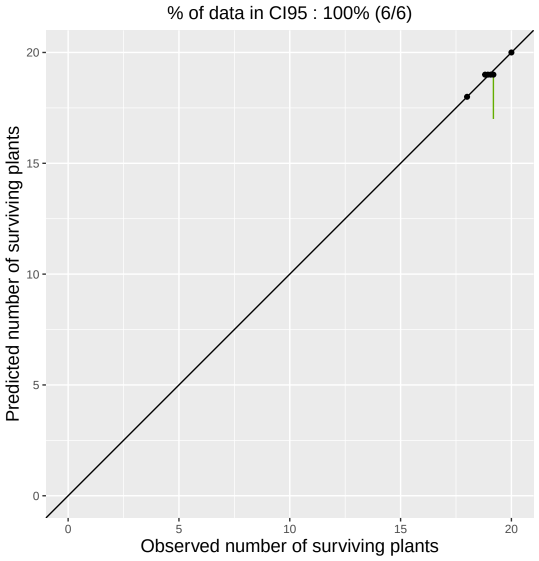

(b) Posterior predictive check (PPC)

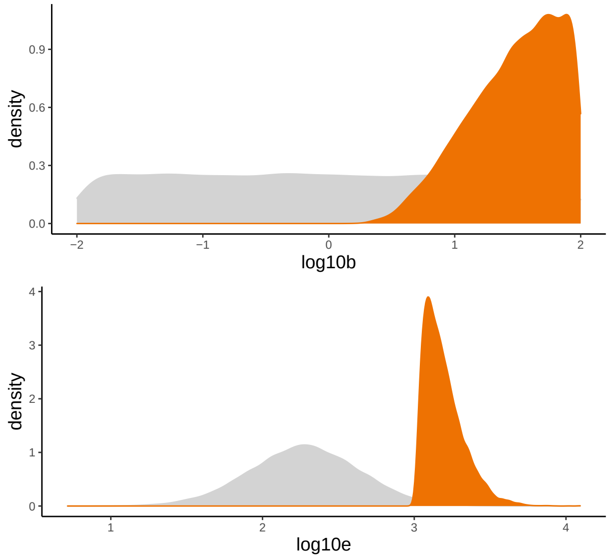

(c) Priors and posteriors

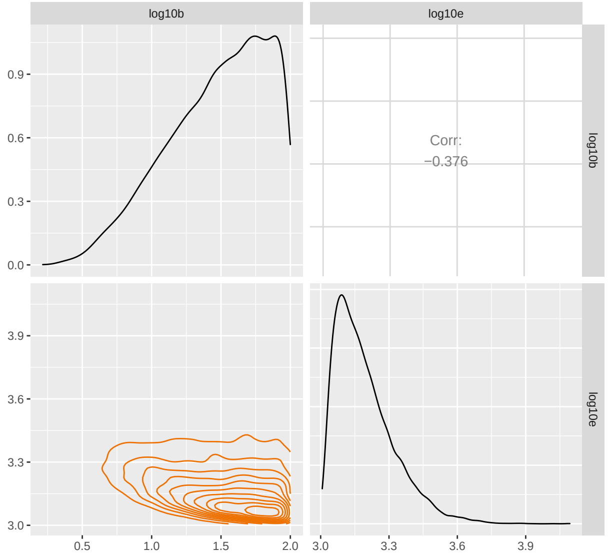

(d) Correlations between parameters

Figure 9: Dose-response curve (a), PPC (b), prior and posterior distributions (c) and correlations between parameters (d).

### Data set: ZEAMA_SE_survival

Table 10: Summary of parameter estimates (parameter d is set to 1) for ZEAMA_SE_survival data set

| Parameter | median | Q2.5 | Q97.5 |
|---|---|---|---|
| b | 7.225 | 2.180 | 59.800 |
| e | 1100.903 | 989.903 | 1676.113 |

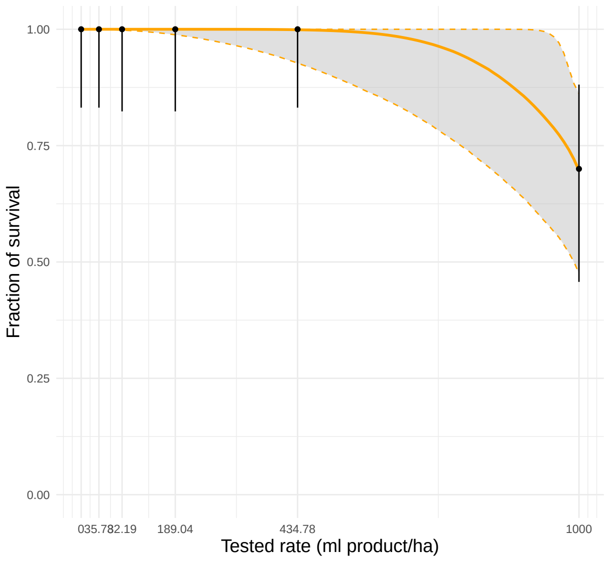

(a) Dose-response curve

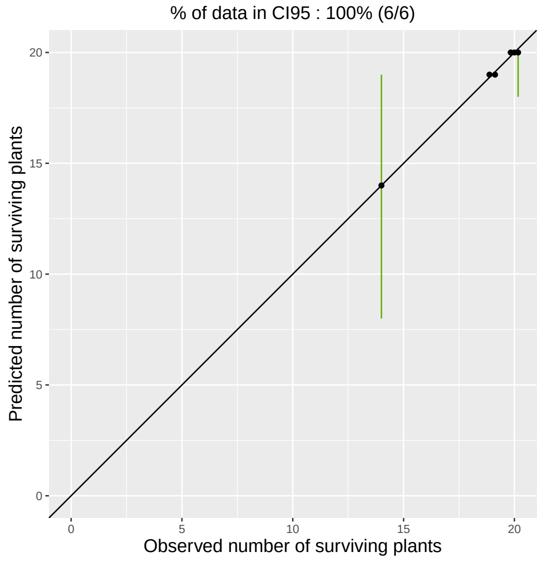

(b) Posterior predictive check (PPC)

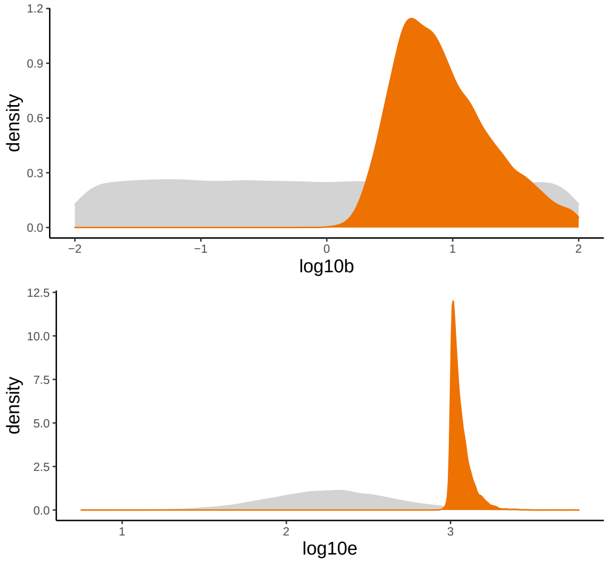

(c) Priors and posteriors

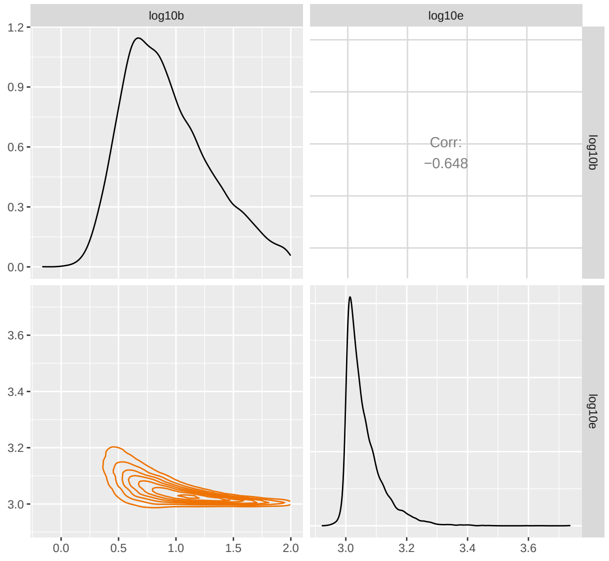

(d) Correlations between parameters

Figure 10: Dose-response curve (a), PPC (b), prior and posterior distributions (c) and correlations between parameters (d).
