## Supplementary material for "How to account for the uncertainty from standard toxicity tests in species sensitivity distributions: an example in non-target plants": Archive with reports on studies: report_SE_weight.pdf

### Dose-response analysis

Study 4

Seedling Emergence test - shoot dry SE_weight endpoint

25 June 2020

Contact:

---

This is a report which provides results on all performed dose-response analyses for the shoot dry SE_weight endpoint of the Seedling Emergence test for study 4.

---

#### Data set: ALLCE_SE_weight

Table 1: Summary of parameter estimates for ALLCE_SE_weight data set

| Parameter | median | Q2.5 | Q97.5 |
|---|---|---|---|
| b | 12.432 | 0.016 | 90.220 |
| d | 0.021 | 0.017 | 0.034 |
| e | 8.029 | 1.189 | 15.364 |
| sigma | 0.008 | 0.006 | 0.013 |

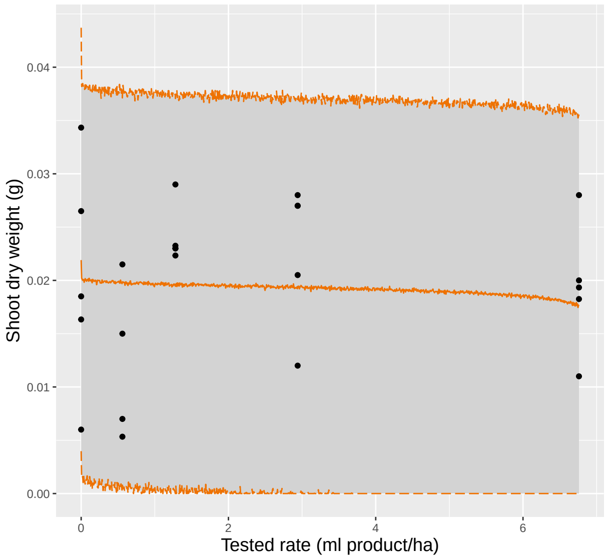

(a) Dose-response curve

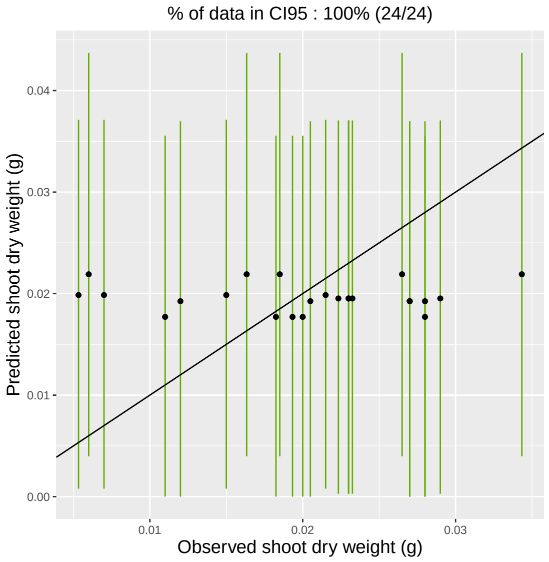

(b) Posterior predictive check (PPC)

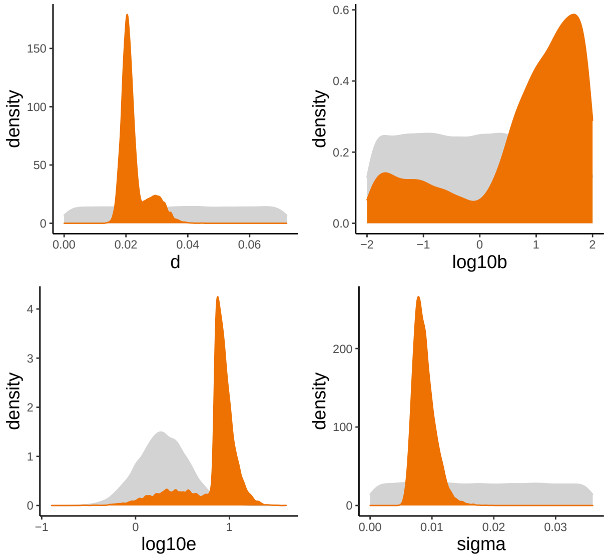

(c) Priors and posteriors

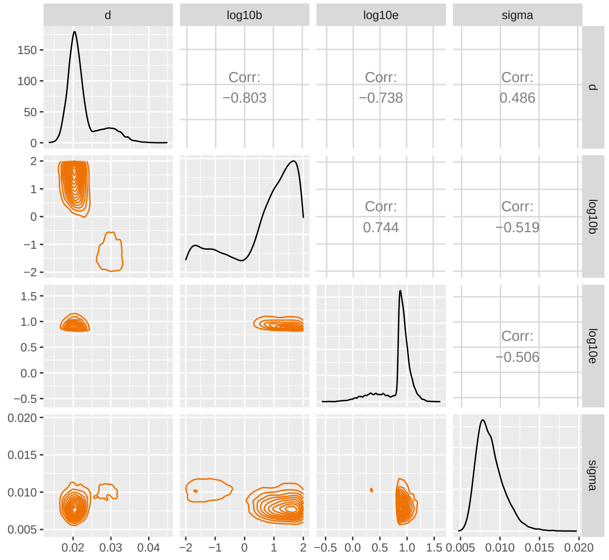

(d) Correlations between parameters

Figure 1: Dose-response curve (a), PPC (b), prior and posterior distributions (c) and correlations between parameters (d).

#### Data set: AVESA_SE_weight

Table 2: Summary of parameter estimates for AVESA_SE_weight data set

| Parameter | median | Q2.5 | Q97.5 |
|---|---:|---:|---:|
| b | 1.710 | 1.377 | 2.164 |
| d | 0.752 | 0.698 | 0.807 |
| e | 110.671 | 93.192 | 130.310 |
| sigma | 0.059 | 0.045 | 0.080 |

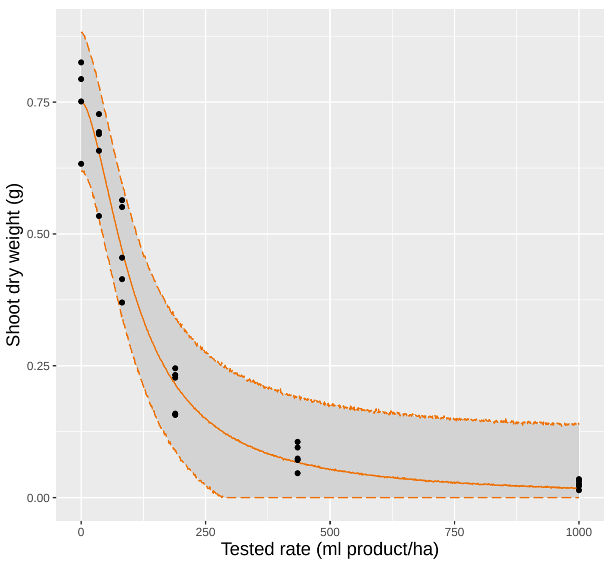

(a) Dose-response curve

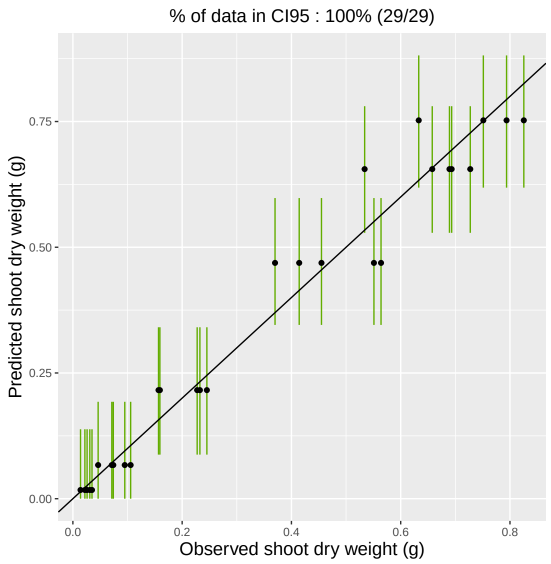

(b) Posterior predictive check (PPC)

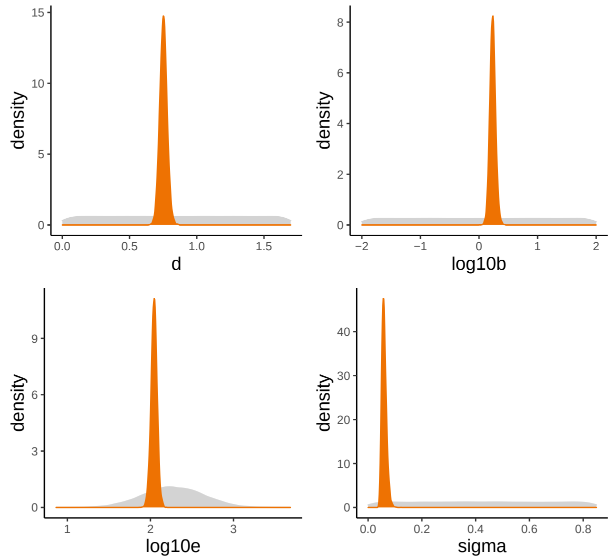

(c) Priors and posteriors

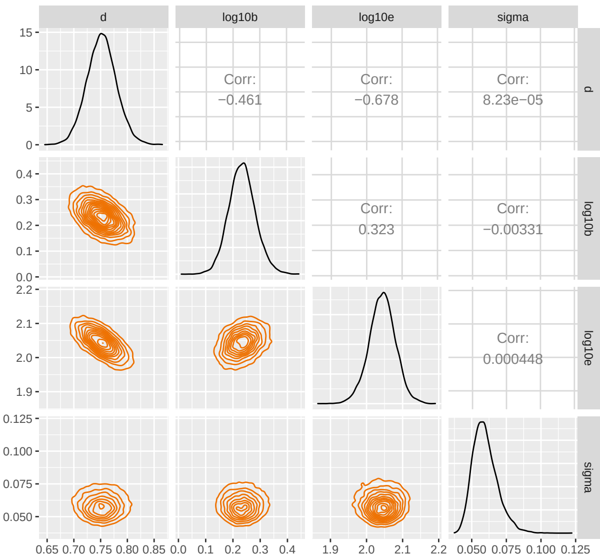

(d) Correlations between parameters

Figure 2: Dose-response curve (a), PPC (b), prior and posterior distributions (c) and correlations between parameters (d).

### Data set: BEAVA_SE_weight

Table 3: Summary of parameter estimates for BEAVA_SE_weight data set

| Parameter | median | Q2.5 | Q97.5 |
|---|---|---|---|
| b | 1.305 | 0.911 | 1.955 |
| d | 1.735 | 1.590 | 1.897 |
| e | 54.289 | 41.534 | 70.856 |
| sigma | 0.257 | 0.212 | 0.321 |

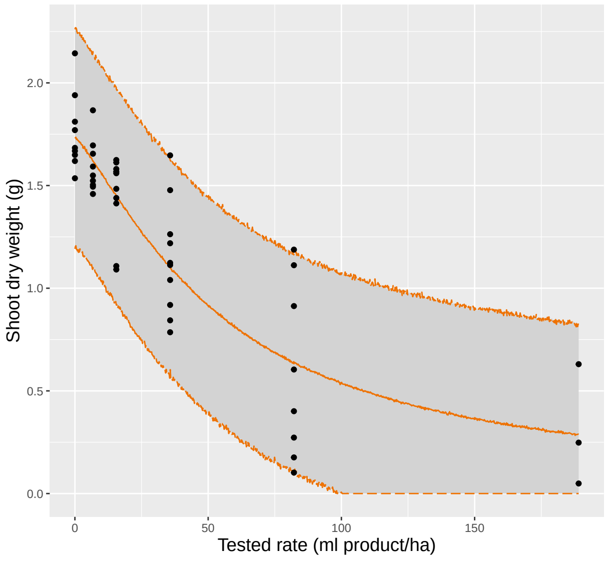

(a) Dose-response curve

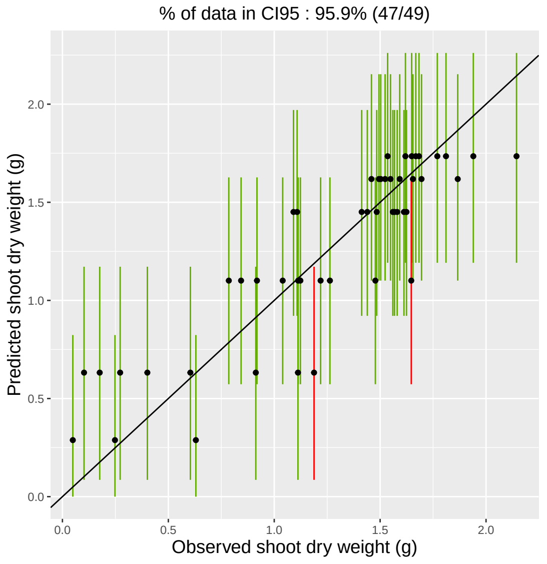

(b) Posterior predictive check (PPC)

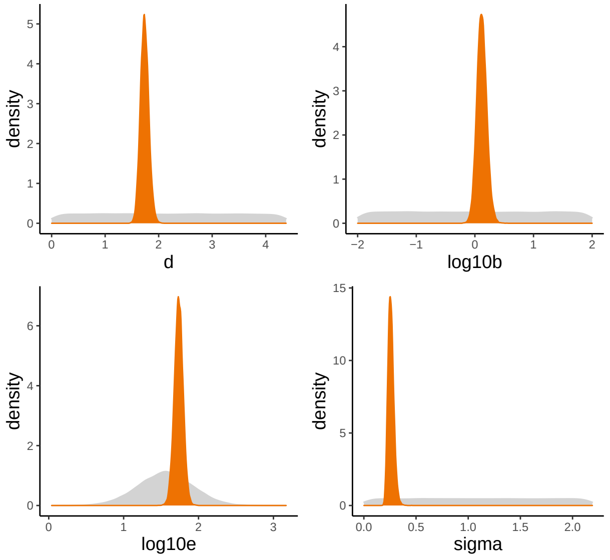

(c) Priors and posteriors

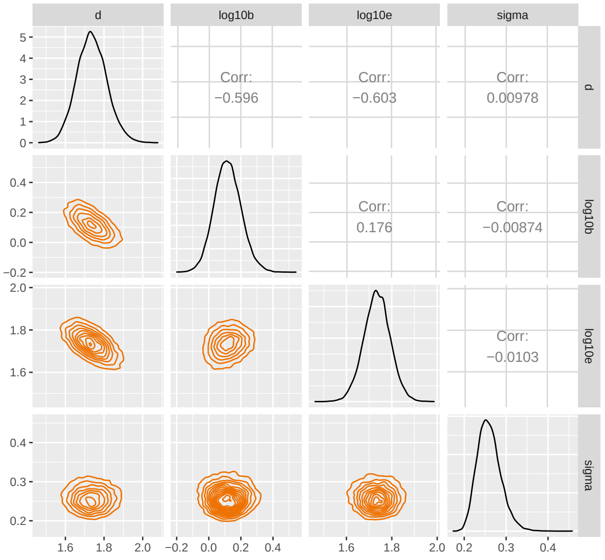

(d) Correlations between parameters

Figure 3: Dose-response curve (a), PPC (b), prior and posterior distributions (c) and correlations between parameters (d).

#### Data set: BRSNW_SE_weight

Table 4: Summary of parameter estimates for BRSNW_SE_weight data set

| Parameter | median | Q2.5 | Q97.5 |
|---|---|---|---|
| b | 2.081 | 1.277 | 3.965 |
| d | 3.167 | 2.857 | 3.504 |
| e | 44.903 | 34.536 | 61.399 |
| sigma | 0.711 | 0.583 | 0.885 |

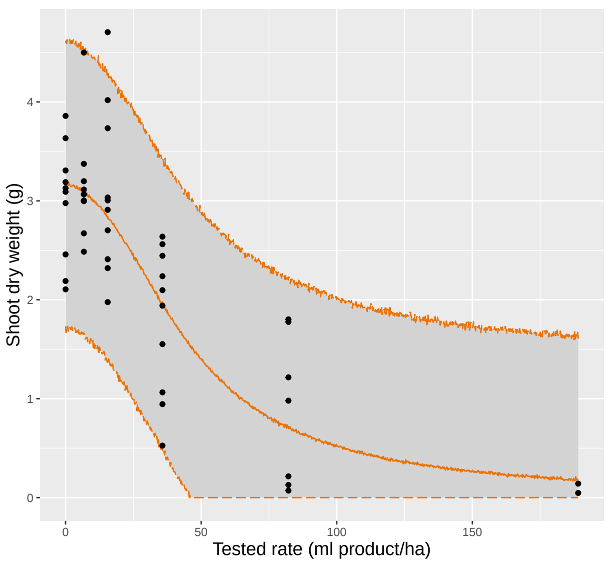

(a) Dose-response curve

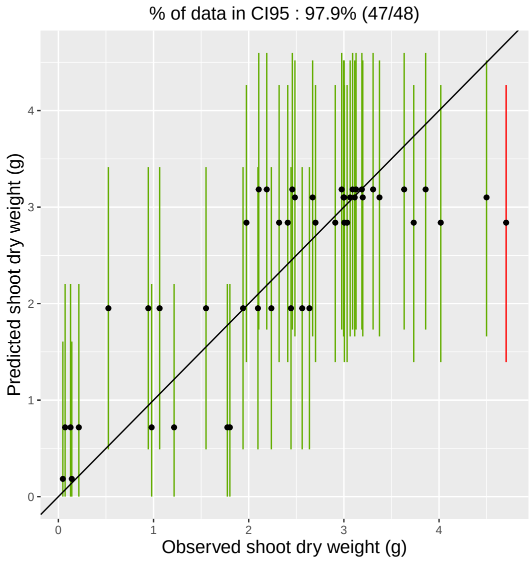

(b) Posterior predictive check (PPC)

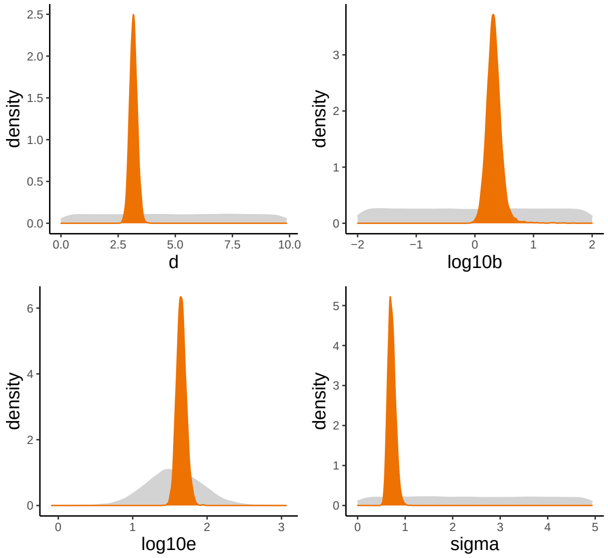

(c) Priors and posteriors

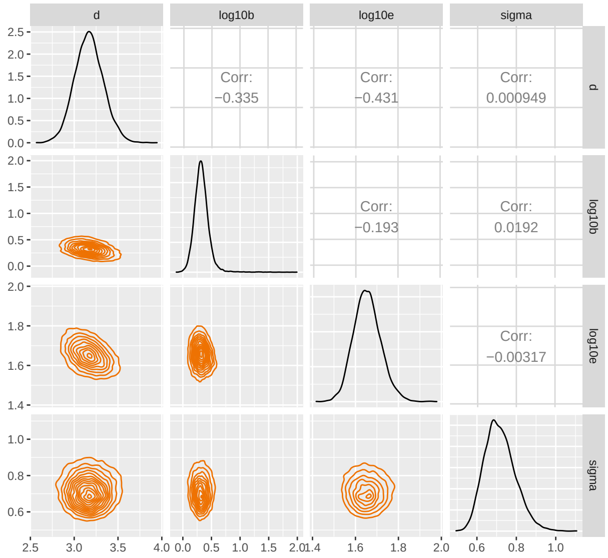

(d) Correlations between parameters

Figure 4: Dose-response curve (a), PPC (b), prior and posterior distributions (c) and correlations between parameters (d).

### Data set: CUMSA_SE_weight

Table 5: Summary of parameter estimates for CUMSA_SE_weight data set

| Parameter | median | Q2.5 | Q97.5 |
|---|---|---|---|
| b | 2.231 | 1.562 | 3.395 |
| d | 4.413 | 4.062 | 4.796 |
| e | 98.977 | 80.277 | 120.276 |
| sigma | 0.751 | 0.631 | 0.915 |

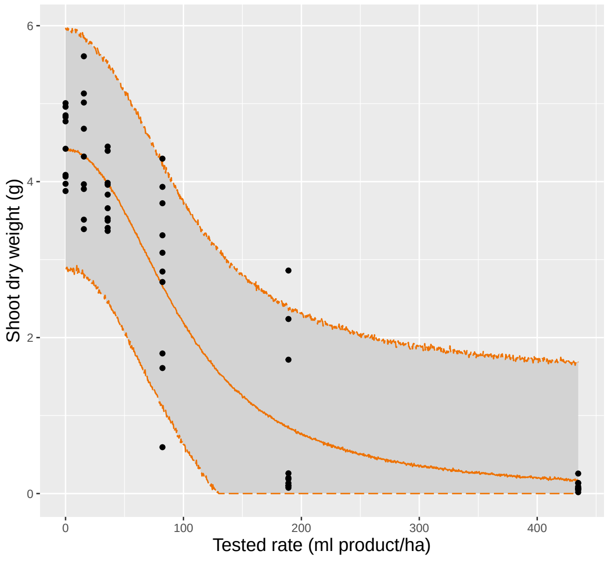

(a) Dose-response curve

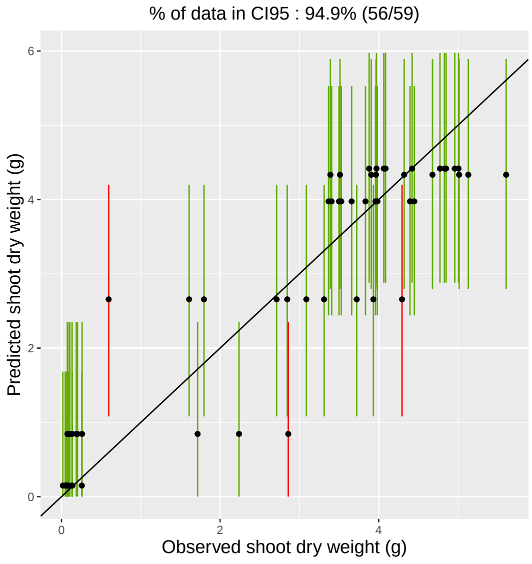

(b) Posterior predictive check (PPC)

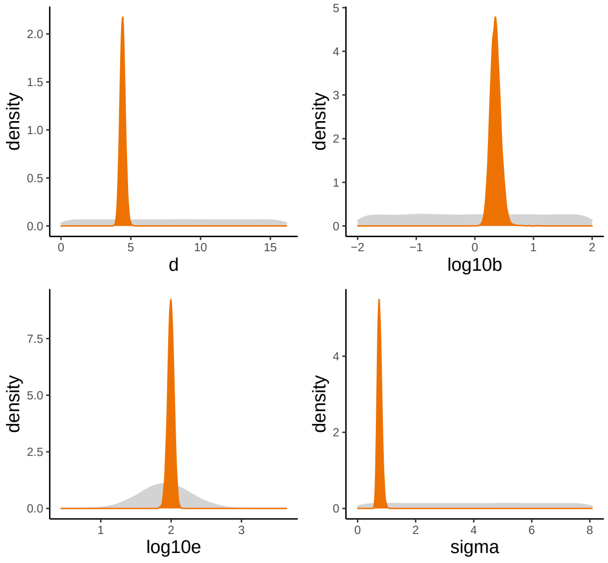

(c) Priors and posteriors

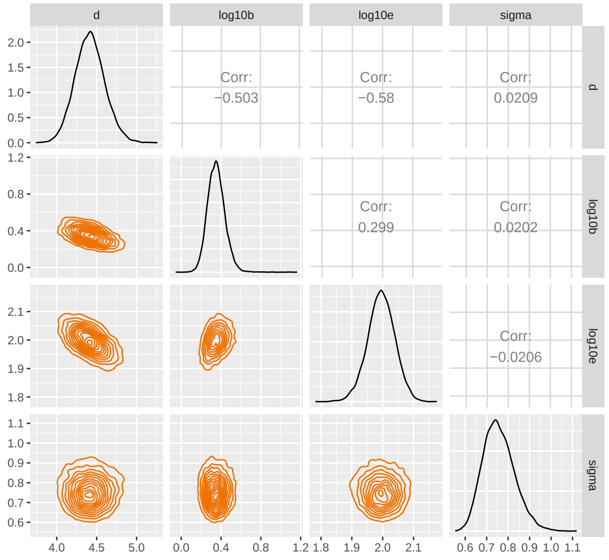

(d) Correlations between parameters

Figure 5: Dose-response curve (a), PPC (b), prior and posterior distributions (c) and correlations between parameters (d).

#### Data set: GLXMA_SE_weight

Table 6: Summary of parameter estimates for GLXMA_SE_weight data set

| Parameter | median | Q2.5 | Q97.5 |
|---|---|---|---|
| b | 1.263 | 0.900 | 1.798 |
| d | 1.696 | 1.572 | 1.840 |
| e | 186.609 | 143.242 | 237.514 |
| sigma | 0.247 | 0.206 | 0.303 |

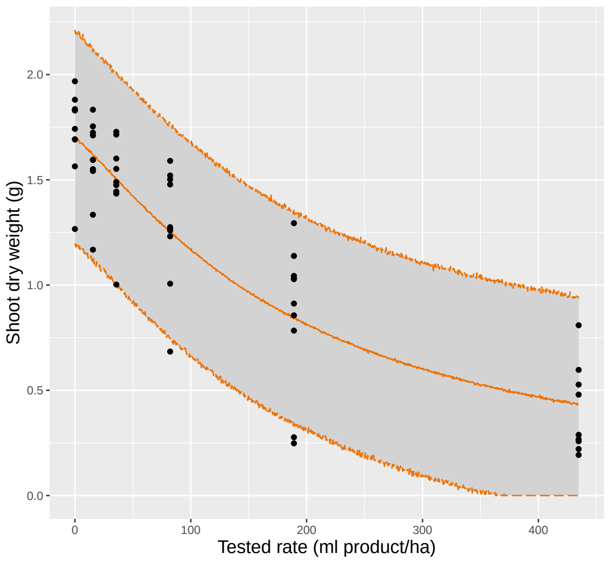

(a) Dose-response curve

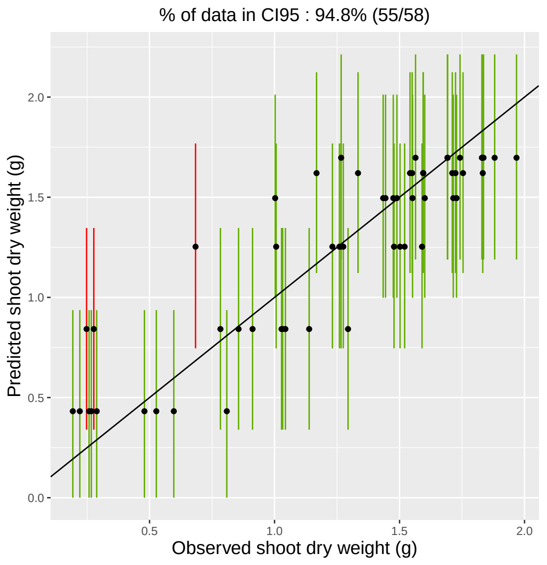

(b) Posterior predictive check (PPC)

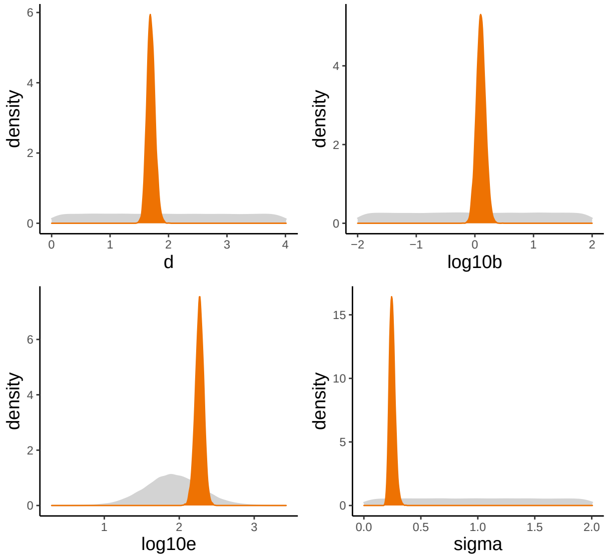

(c) Priors and posteriors

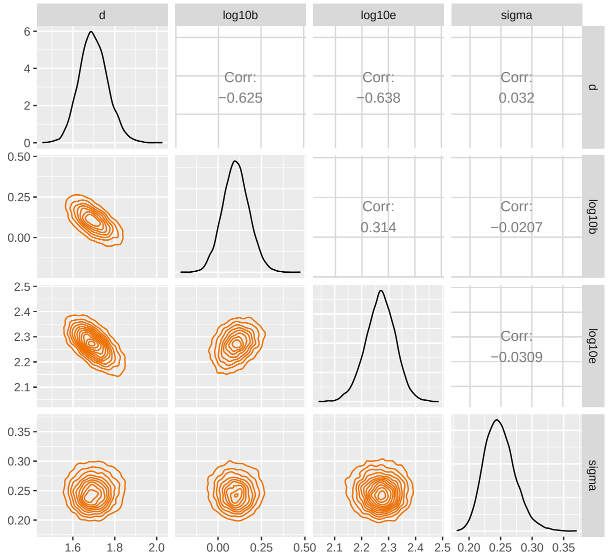

(d) Correlations between parameters

Figure 6: Dose-response curve (a), PPC (b), prior and posterior distributions (c) and correlations between parameters (d).

#### Data set: HELAN_SE_weight

Table 7: Summary of parameter estimates for HELAN_SE_weight data set

| Parameter | median | Q2.5 | Q97.5 |
|---|---|---|---|
| b | 1.901 | 1.377 | 2.807 |
| d | 1.043 | 0.958 | 1.135 |
| e | 69.388 | 56.477 | 84.429 |
| sigma | 0.154 | 0.127 | 0.194 |

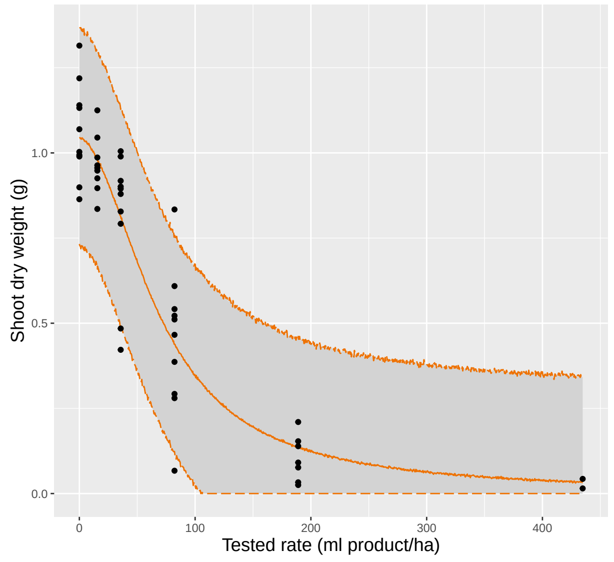

(a) Dose-response curve

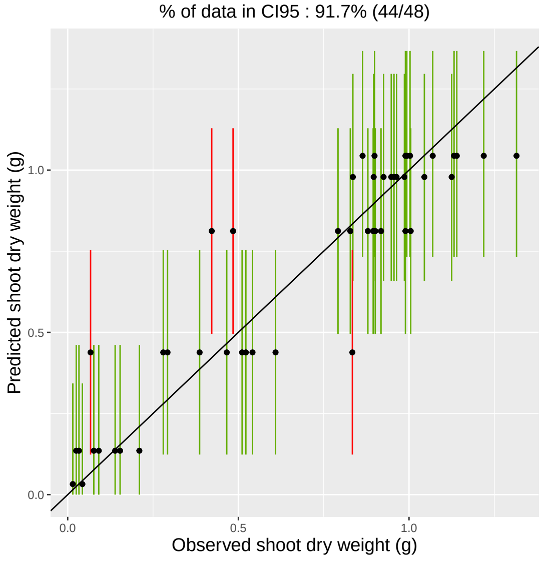

(b) Posterior predictive check (PPC)

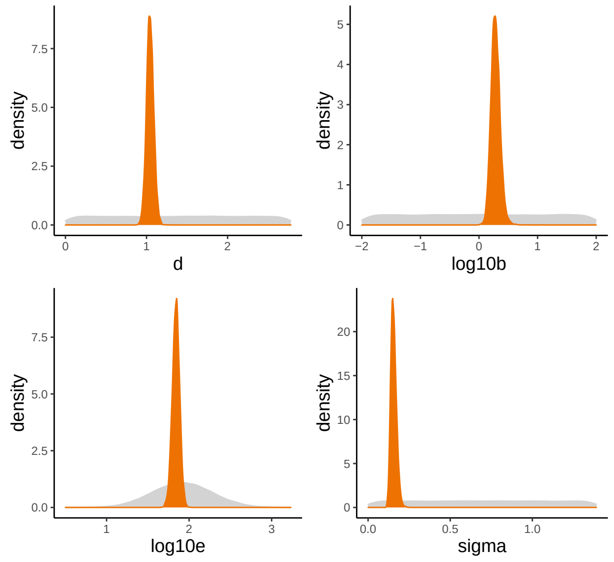

(c) Priors and posteriors

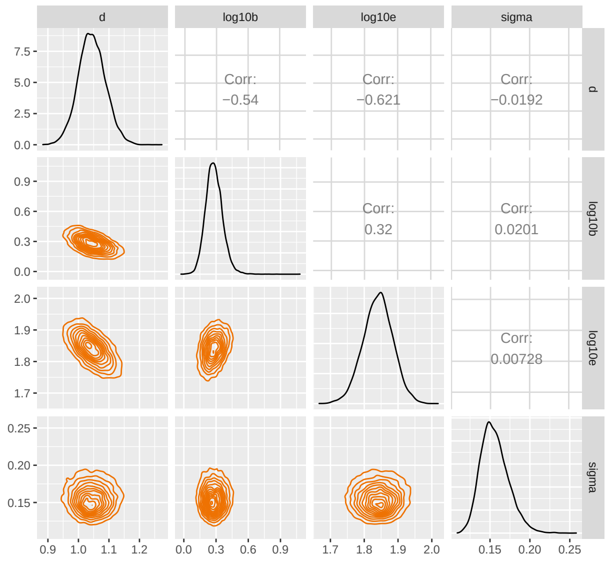

(d) Correlations between parameters

Figure 7: Dose-response curve (a), PPC (b), prior and posterior distributions (c) and correlations between parameters (d).

#### Data set: LYPES_SE_weight

Table 8: Summary of parameter estimates for LYPES_SE_weight data set

| Parameter | median | Q2.5 | Q97.5 |
|---|---|---|---|
| b | 3.121 | 1.856 | 18.591 |
| d | 1.205 | 1.076 | 1.348 |
| e | 42.107 | 35.353 | 52.337 |
| sigma | 0.240 | 0.199 | 0.298 |

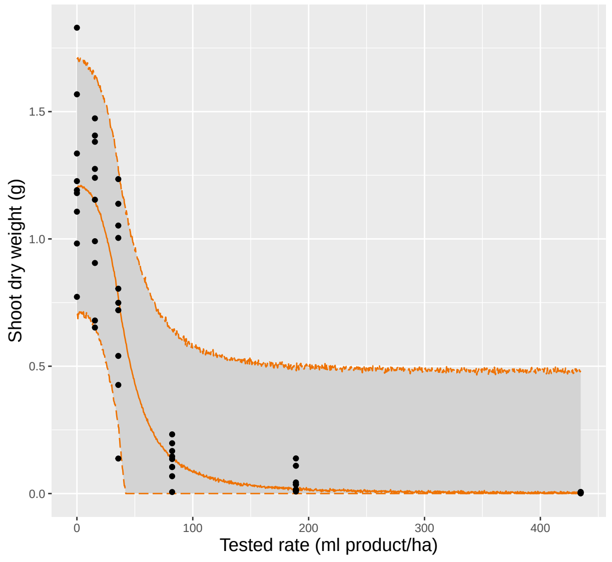

(a) Dose-response curve

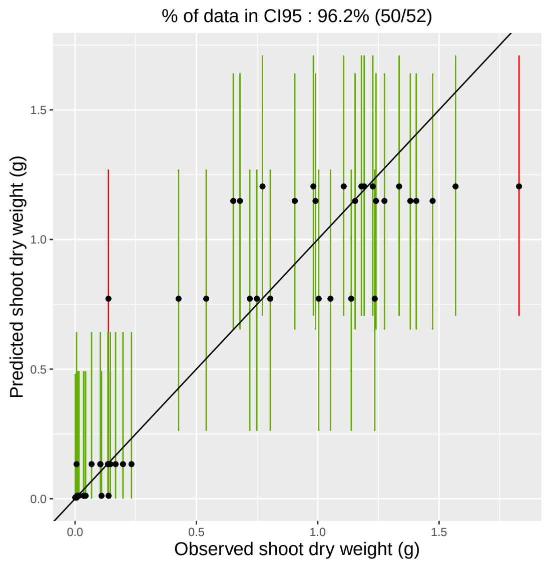

(b) Posterior predictive check (PPC)

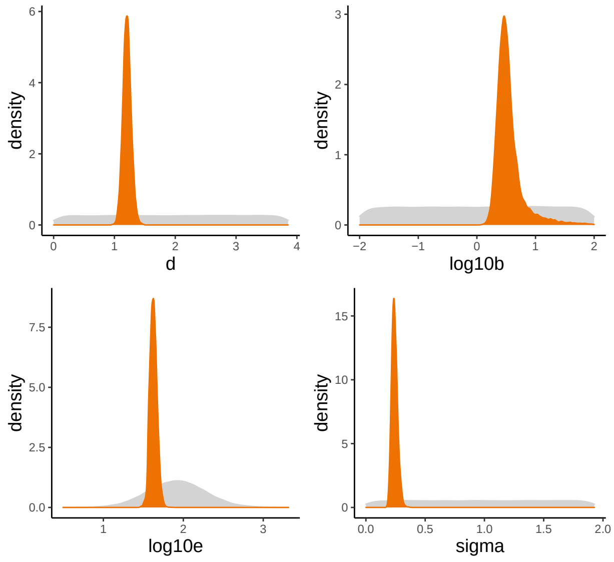

(c) Priors and posteriors

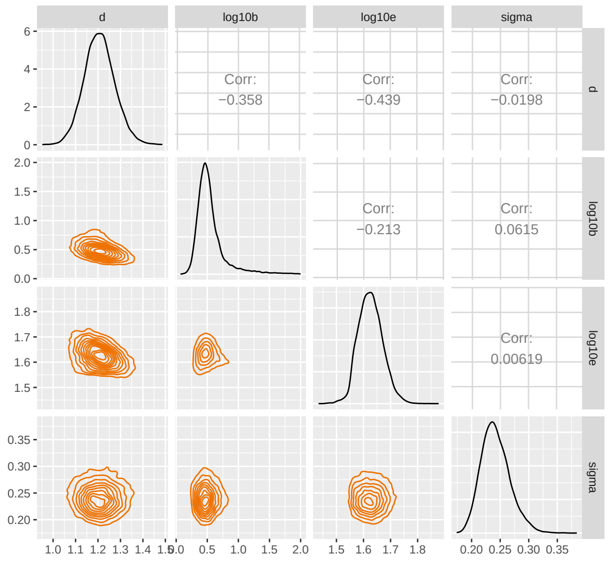

(d) Correlations between parameters

Figure 8: Dose-response curve (a), PPC (b), prior and posterior distributions (c) and correlations between parameters (d).

#### Data set: TRZAW_SE_weight

Table 9: Summary of parameter estimates for TRZAW_SE_weight data set

| Parameter | median | Q2.5 | Q97.5 |
|---|---|---|---|
| b | 0.846 | 0.643 | 1.114 |
| d | 0.610 | 0.575 | 0.647 |
| e | 880.597 | 672.858 | 1195.803 |
| sigma | 0.045 | 0.034 | 0.061 |

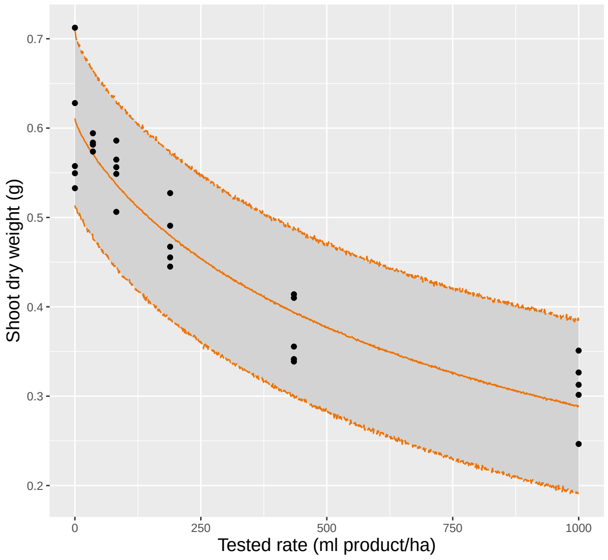

(a) Dose-response curve

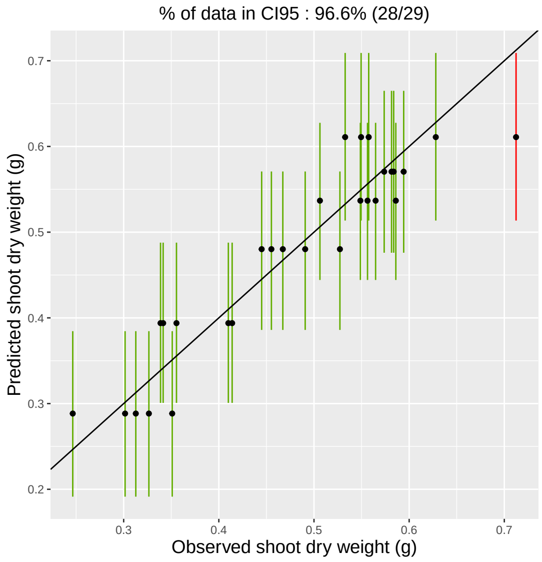

(b) Posterior predictive check (PPC)

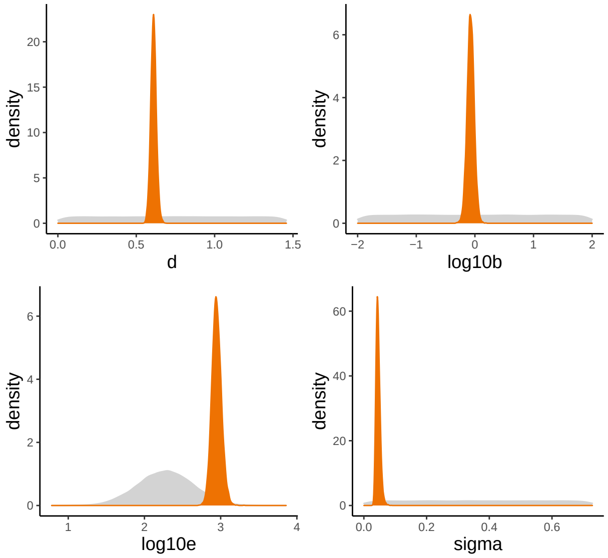

(c) Priors and posteriors

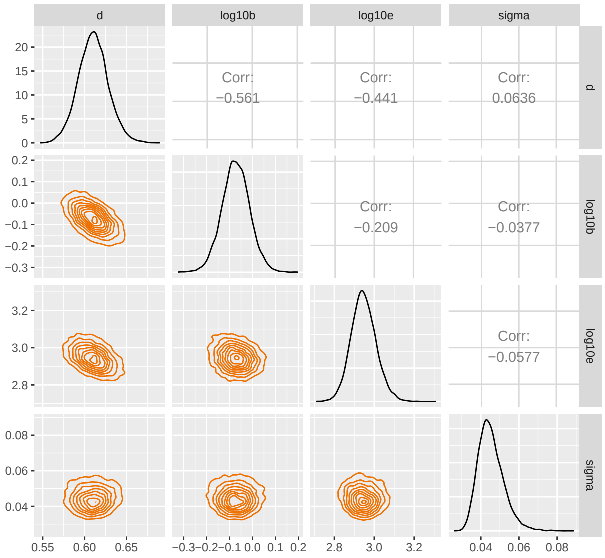

(d) Correlations between parameters

Figure 9: Dose-response curve (a), PPC (b), prior and posterior distributions (c) and correlations between parameters (d).

#### Data set: ZEAMA_SE_weight

Table 10: Summary of parameter estimates for ZEAMA_SE_weight data set

| Parameter | median | Q2.5 | Q97.5 |
|---|---|---|---|
| b | 1.103 | 0.792 | 1.557 |
| d | 5.416 | 4.937 | 5.954 |
| e | 302.849 | 219.225 | 417.390 |
| sigma | 0.906 | 0.757 | 1.115 |

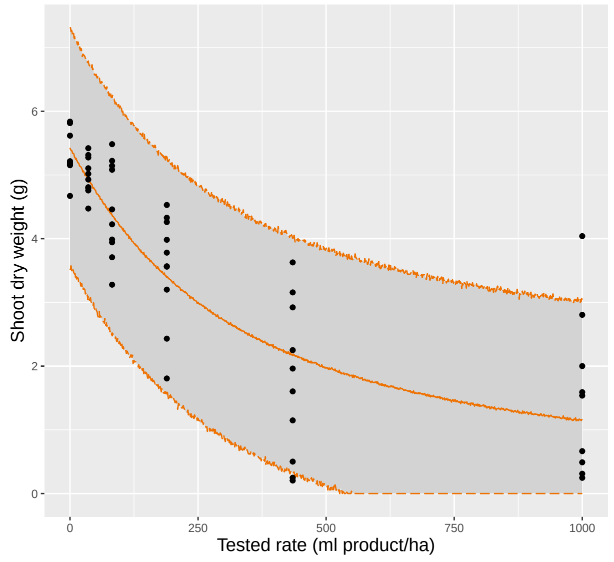

(a) Dose-response curve

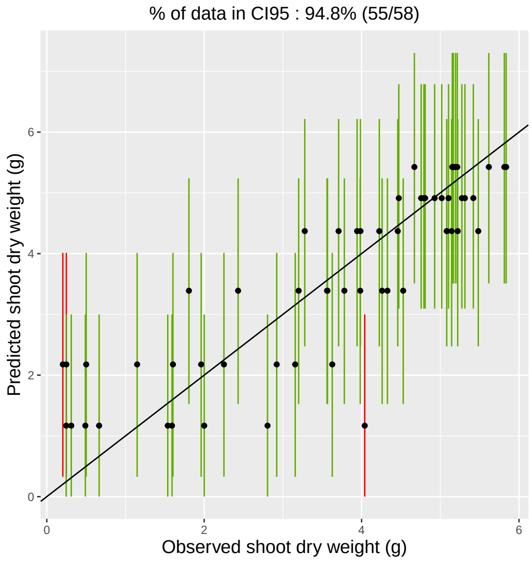

(b) Posterior predictive check (PPC)

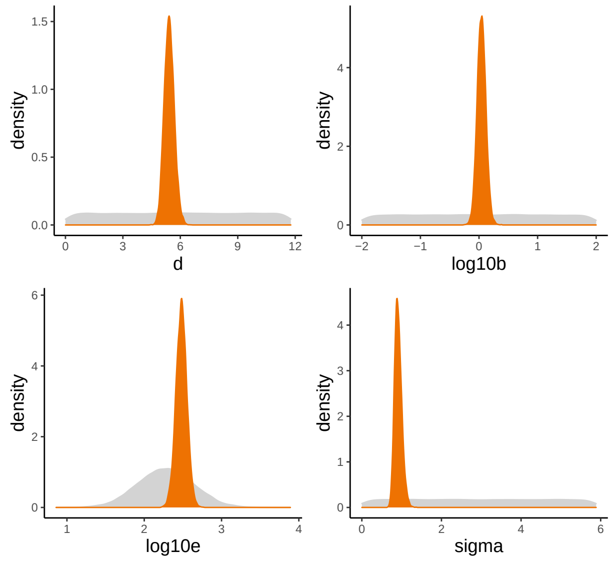

(c) Priors and posteriors

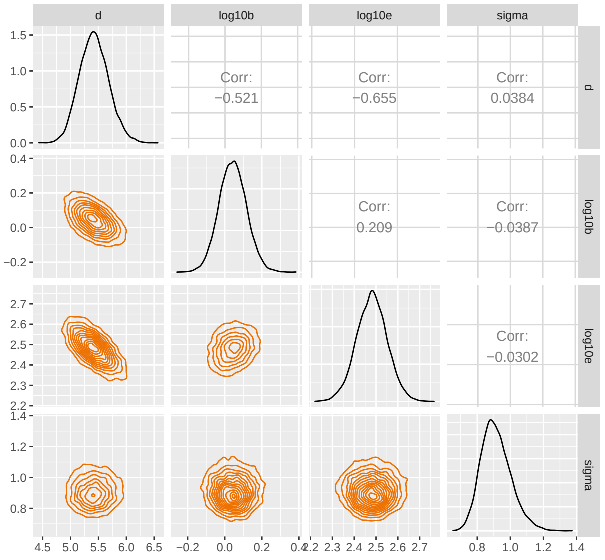

(d) Correlations between parameters

Figure 10: Dose-response curve (a), PPC (b), prior and posterior distributions (c) and correlations between parameters (d).
