## Supplementary material for "How to account for the uncertainty from standard toxicity tests in species sensitivity distributions: an example in non-target plants": Archive with reports on studies: report_VV_survival.pdf

### Dose-response analyses

Study 4

Vegetative Vigour test - survival endpoint

25 June 2020

Contact:

---

This is a report which provides results on all performed dose-response analyses for the survival endpoint of the Vegetative Vigour test for study 4.

---

#### Data set: ALLCE_VV_survival

Table 1: Summary of parameter estimates (parameter d is set to 1) for ALLCE_VV_survival data set

| Parameter | median | Q2.5 | Q97.5 |
|---|---|---|---|
| b | 1.030 | 0.614 | 1.590 |
| e | 459.047 | 278.567 | 980.247 |

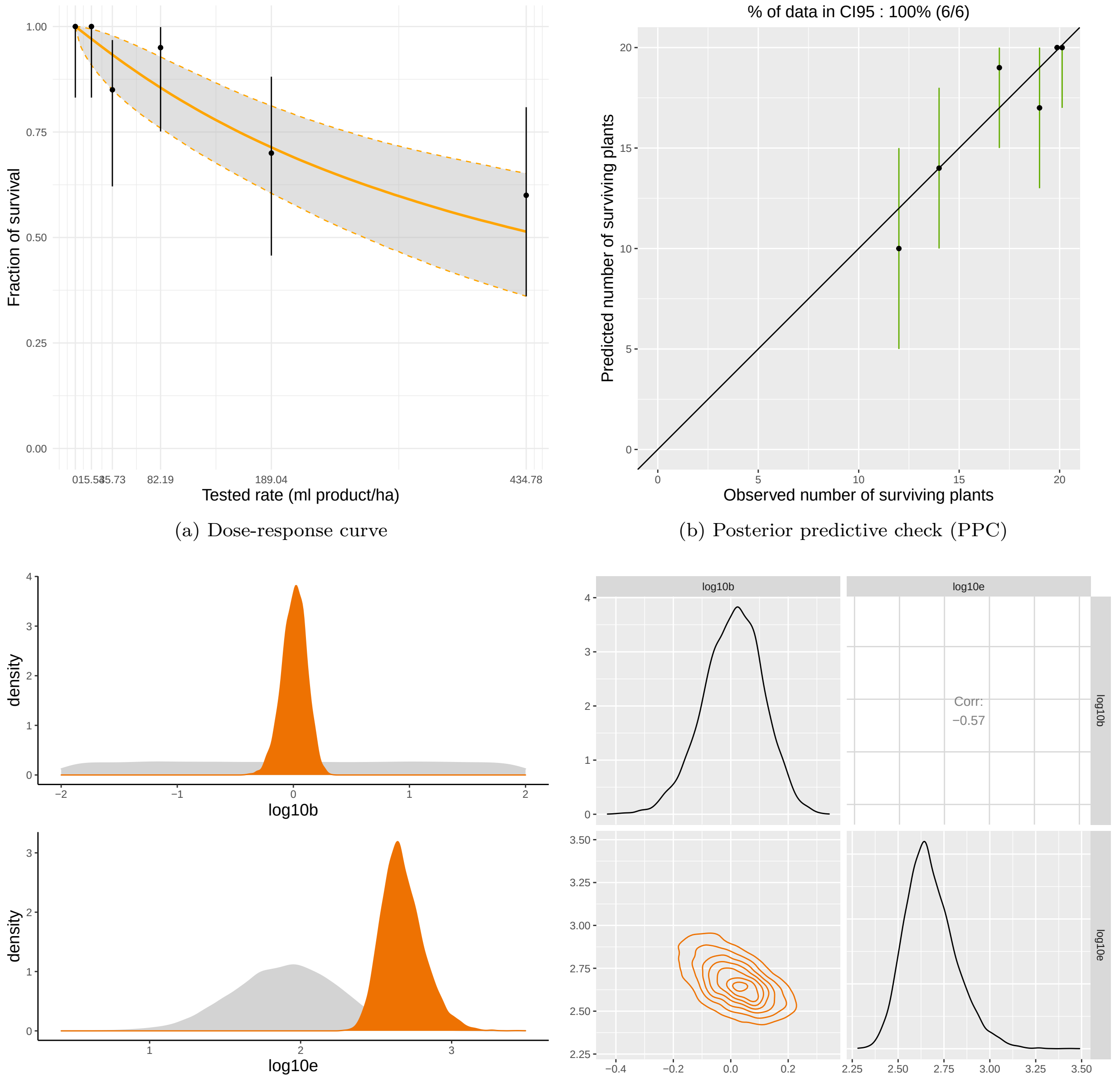

(a) Dose-response curve

(b) Posterior predictive check (PPC)

(c) Priors and posteriors

(d) Correlations between parameters

Figure 1: Dose-response curve (a), PPC (b), prior and posterior distributions (c) and correlations between parameters (d).

#### Data set: AVESA_VV_survival

Table 2: Summary of parameter estimates (parameter d is set to 1) for AVESA_VV_survival data set

| Parameter | median | Q2.5 | Q97.5 |
|---|---|---|---|
| b | 9.655 | 3.732 | 68.712 |
| e | 446.877 | 406.814 | 524.025 |

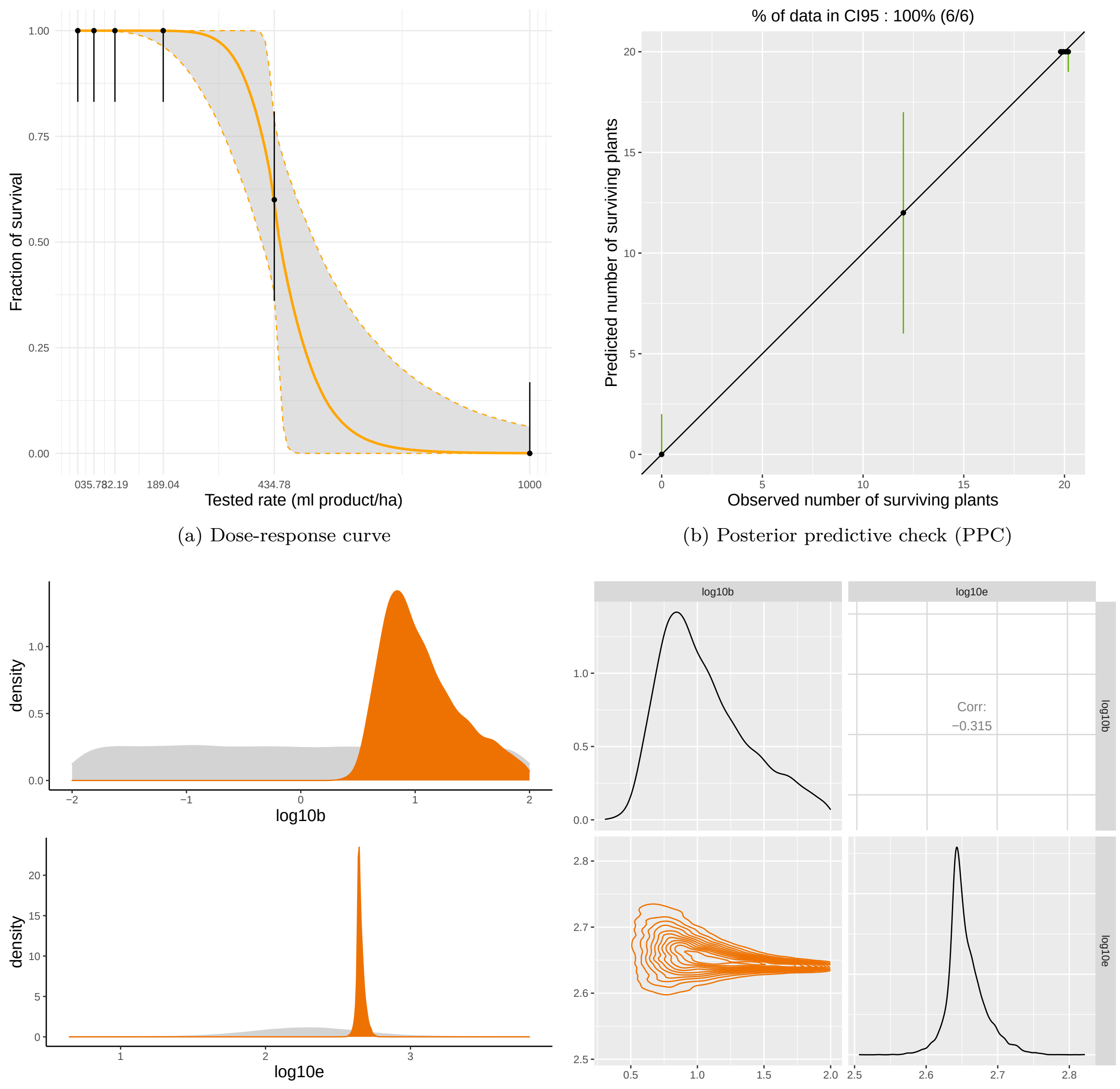

(a) Dose-response curve

(b) Posterior predictive check (PPC)

(c) Priors and posteriors

(d) Correlations between parameters

Figure 2: Dose-response curve (a), PPC (b), prior and posterior distributions (c) and correlations between parameters (d).

#### Data set: BEAVA_VV_survival

Table 3: Summary of parameter estimates (parameter d is set to 1) for BEAVA_VV_survival data set

| Parameter | median | Q2.5 | Q97.5 |
|---|---|---|---|
| b | 1.177 | 0.702 | 1.757 |
| e | 122.099 | 79.748 | 215.885 |

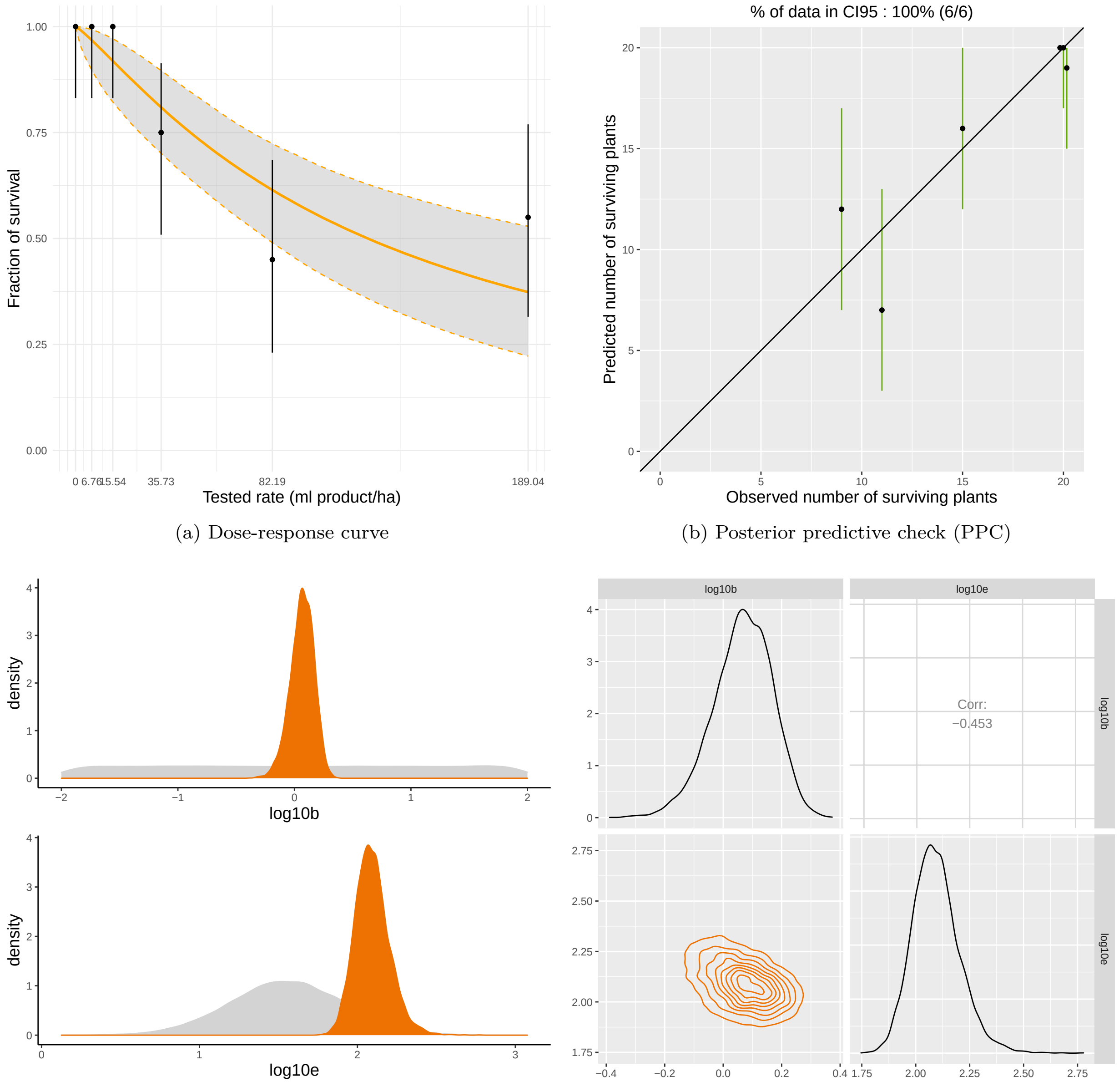

(a) Dose-response curve

(b) Posterior predictive check (PPC)

(c) Priors and posteriors

(d) Correlations between parameters

Figure 3: Dose-response curve (a), PPC (b), prior and posterior distributions (c) and correlations between parameters (d).

### Data set: BRSNW_VV_survival

Table 4: Summary of parameter estimates (parameter d is set to 1) for BRSNW_VV_survival data set

| Parameter | median | Q2.5 | Q97.5 |
|---|---|---|---|
| b | 2.352 | 1.262 | 4.288 |
| e | 239.513 | 175.276 | 443.420 |

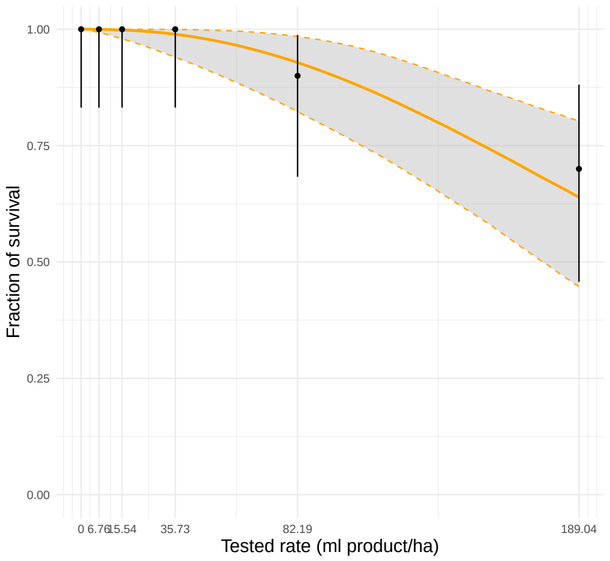

(a) Dose-response curve

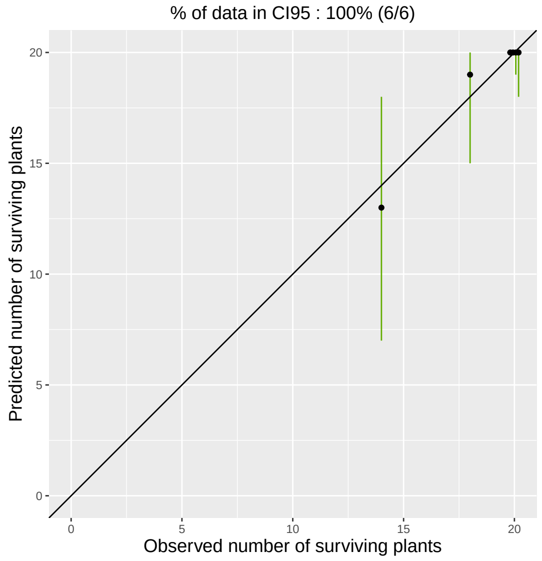

(b) Posterior predictive check (PPC)

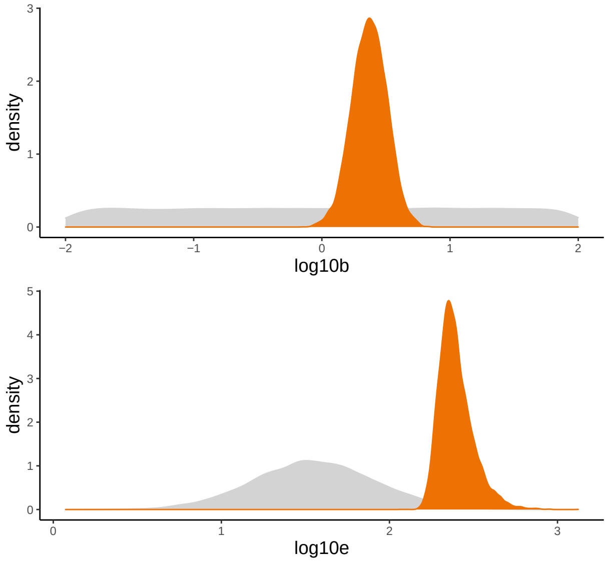

(c) Priors and posteriors

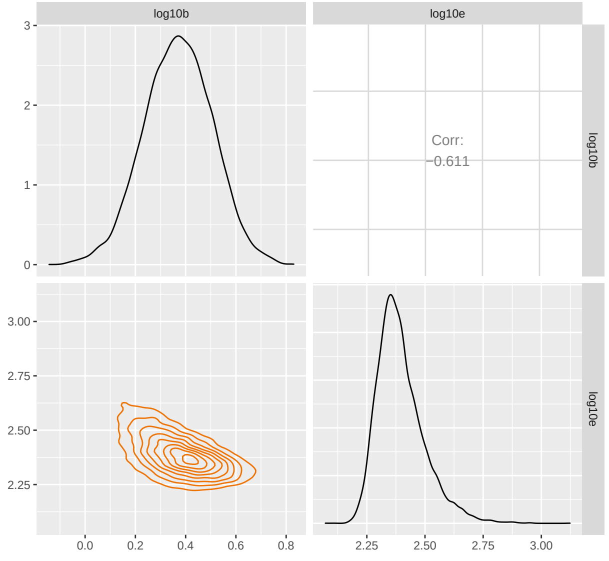

(d) Correlations between parameters

Figure 4: Dose-response curve (a), PPC (b), prior and posterior distributions (c) and correlations between parameters (d).

### Data set: CUMSA_VV_survival

Table 5: Summary of parameter estimates (parameter d is set to 1) for CUMSA_VV_survival data set

| Parameter | median | Q2.5 | Q97.5 |
|---|---:|---:|---:|
| b | 1.407 | 0.858 | 2.211 |
| e | 521.835 | 329.418 | 1139.970 |

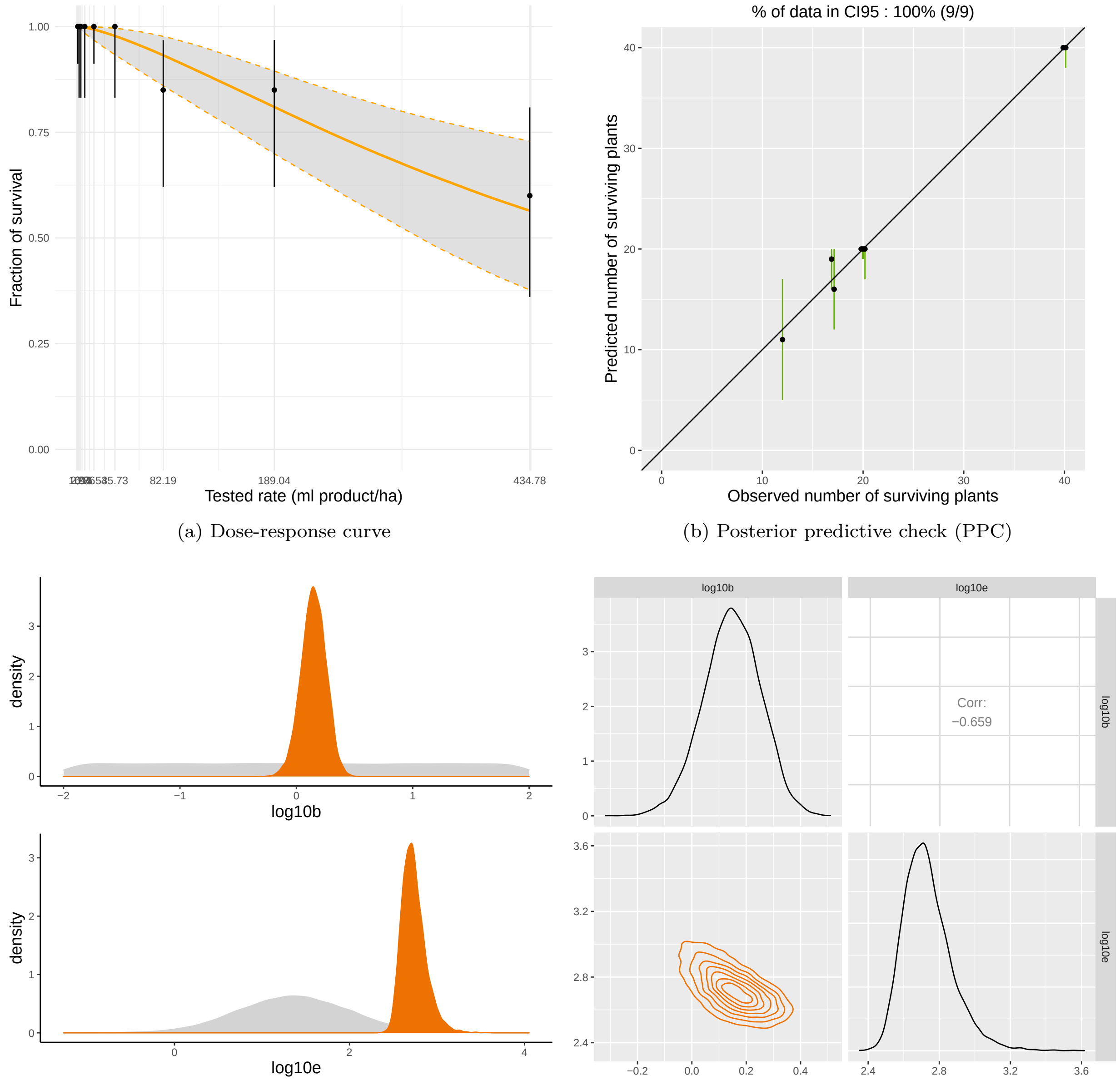

(a) Dose-response curve

(b) Posterior predictive check (PPC)

(c) Priors and posteriors

(d) Correlations between parameters

Figure 5: Dose-response curve (a), PPC (b), prior and posterior distributions (c) and correlations between parameters (d).

#### Data set: GLXMA_VV_survival

Table 6: Summary of parameter estimates (parameter d is set to 1) for GLXMA_VV_survival data set

| Parameter | median | Q2.5 | Q97.5 |
|---|---|---|---|
| b | 9.400 | 4.107 | 64.028 |
| e | 377.474 | 300.891 | 427.586 |

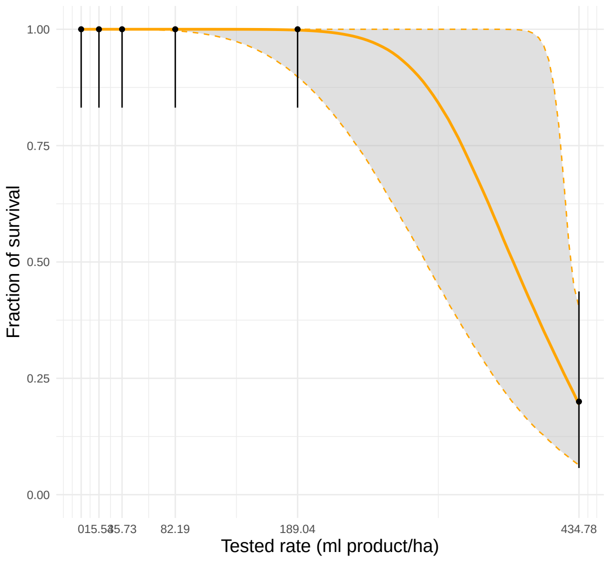

(a) Dose-response curve

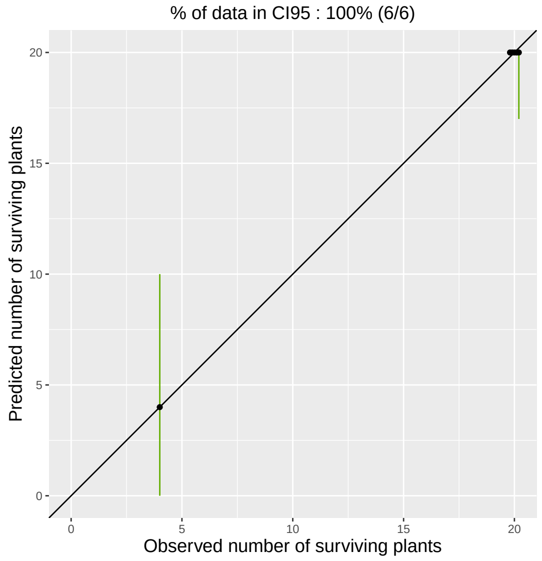

(b) Posterior predictive check (PPC)

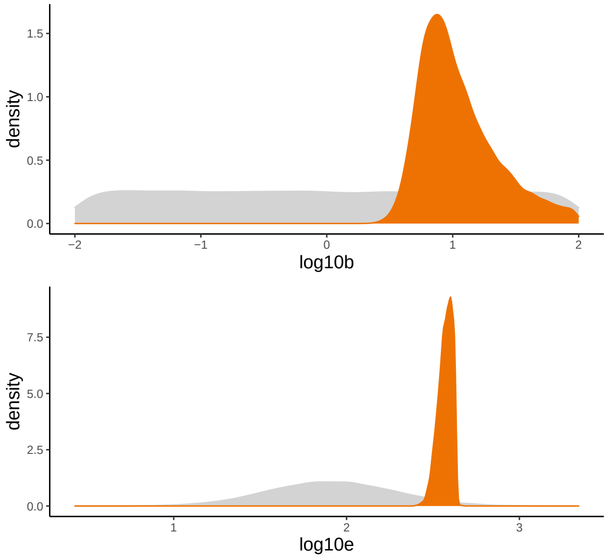

(c) Priors and posteriors

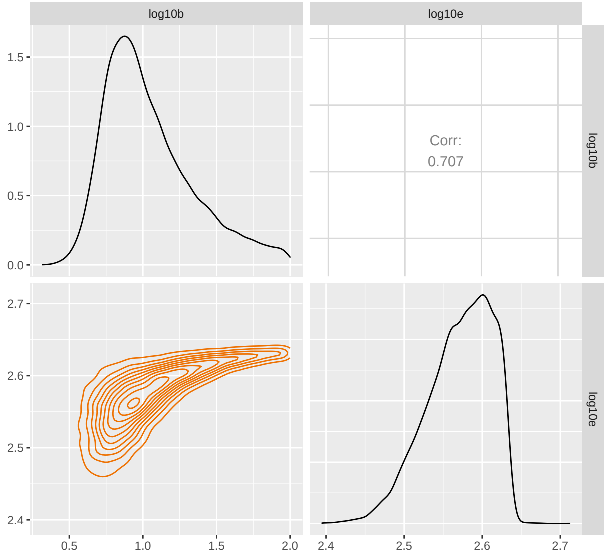

(d) Correlations between parameters

Figure 6: Dose-response curve (a), PPC (b), prior and posterior distributions (c) and correlations between parameters (d).

#### Data set: HELAN_VV_survival

Table 7: Summary of parameter estimates (parameter d is set to 1) for HELAN_VV_survival data set

| Parameter | median | Q2.5 | Q97.5 |
|---|---|---|---|
| b | 14.984 | 6.148 | 78.197 |
| e | 67.162 | 50.999 | 79.275 |

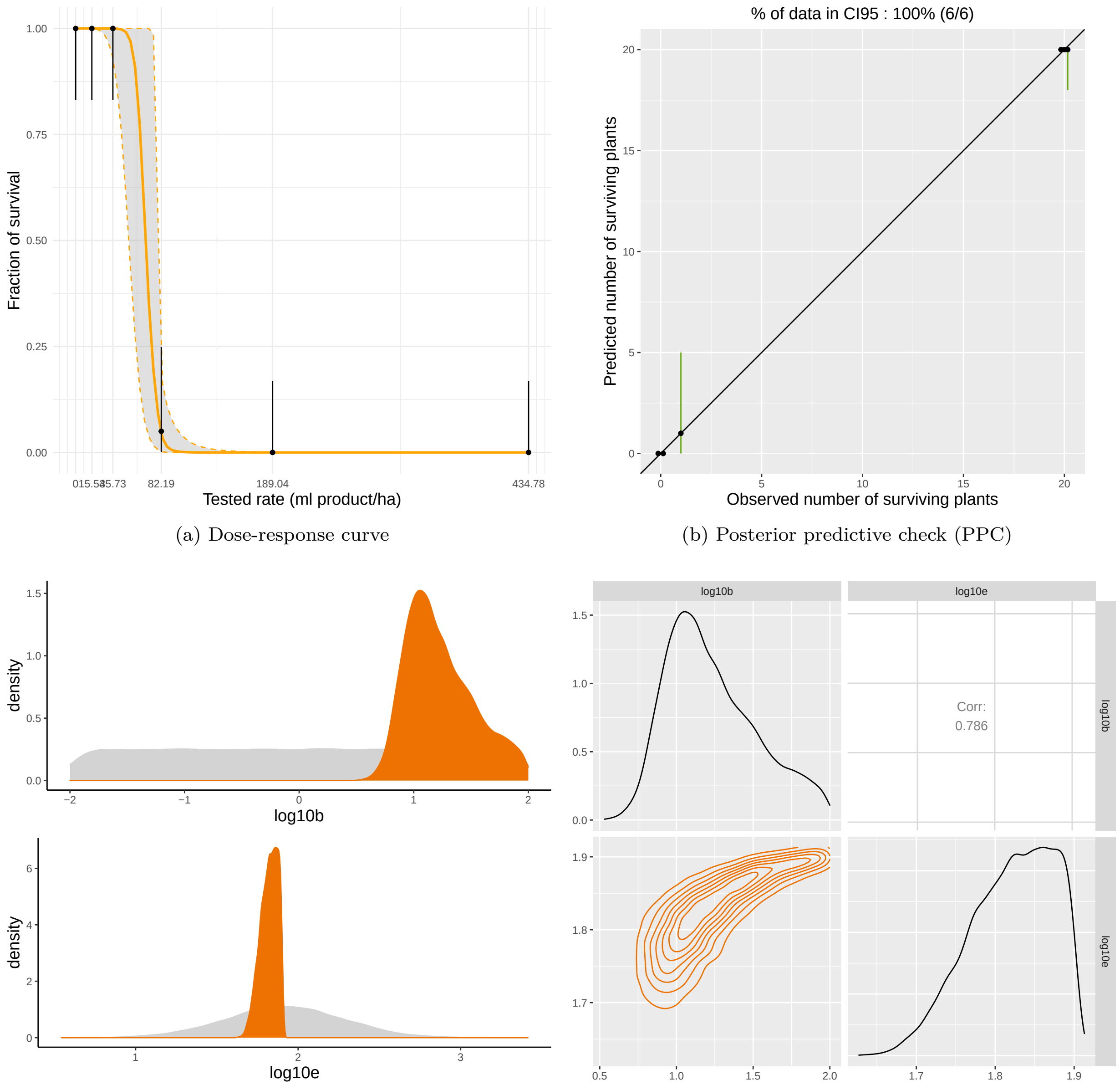

(a) Dose-response curve

(b) Posterior predictive check (PPC)

(c) Priors and posteriors

(d) Correlations between parameters

Figure 7: Dose-response curve (a), PPC (b), prior and posterior distributions (c) and correlations between parameters (d).

#### Data set: LYPES_VV_survival

Table 8: Summary of parameter estimates (parameter d is set to 1) for LYPES_VV_survival data set

| Parameter | median | Q2.5 | Q97.5 |
|---|---|---|---|
| b | 3.987 | 2.583 | 6.046 |
| e | 129.549 | 104.799 | 159.331 |

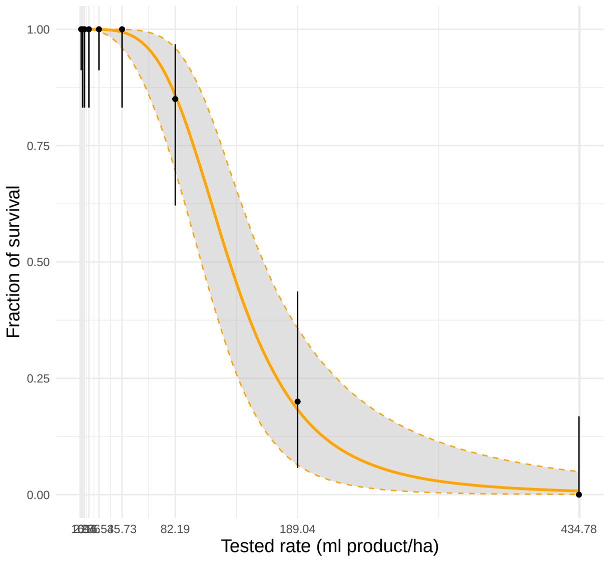

(a) Dose-response curve

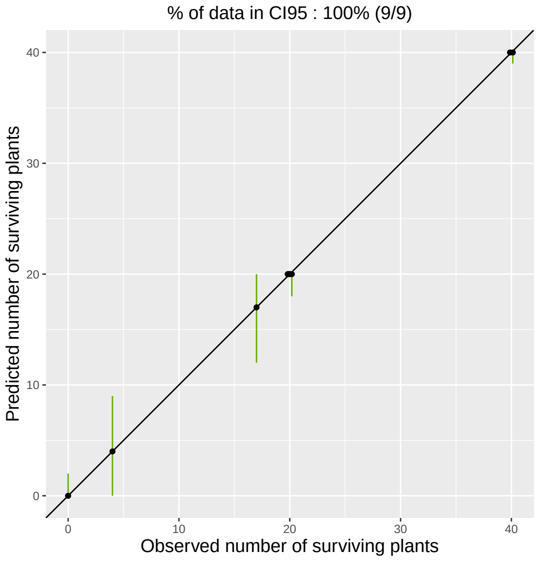

(b) Posterior predictive check (PPC)

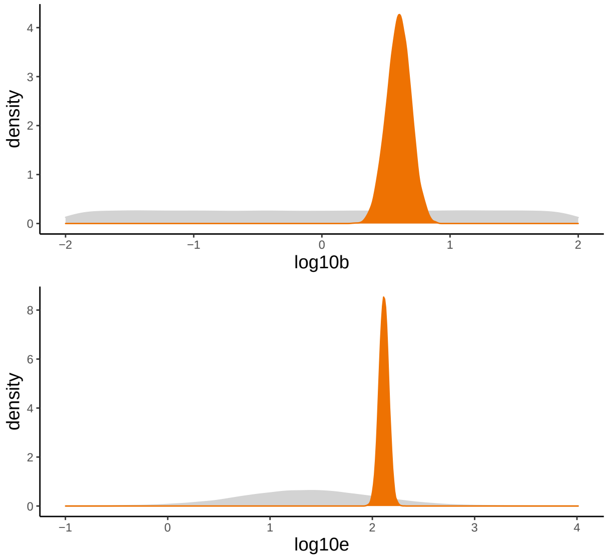

(c) Priors and posteriors

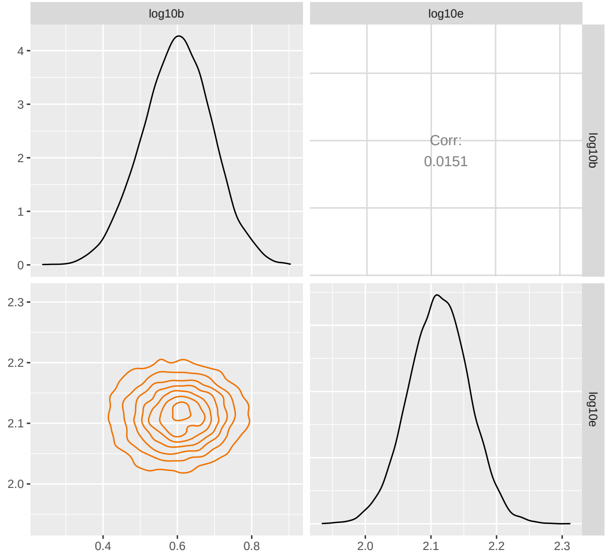

(d) Correlations between parameters

Figure 8: Dose-response curve (a), PPC (b), prior and posterior distributions (c) and correlations between parameters (d).

#### Data set: TRZAW_VV_survival

Table 9: Summary of parameter estimates (parameter d is set to 1) for TRZAW_VV_survival data set

| Parameter | median | Q2.5 | Q97.5 |
|---|---|---|---|
| b | 33.211 | 4.902 | 94.693 |
| e | 1466.795 | 1075.007 | 3369.873 |

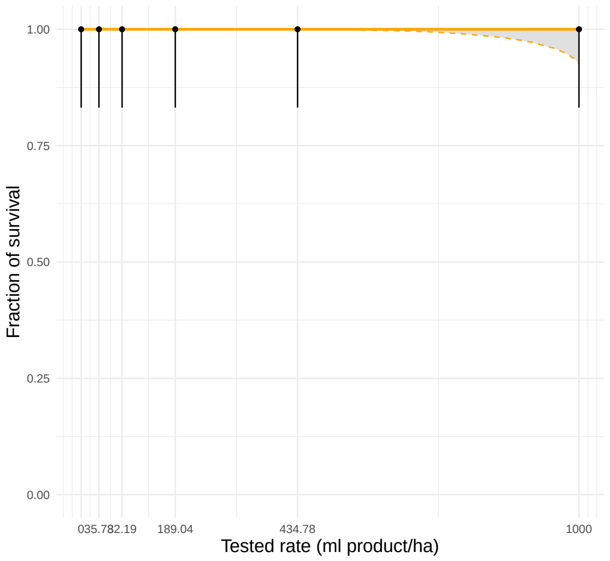

(a) Dose-response curve

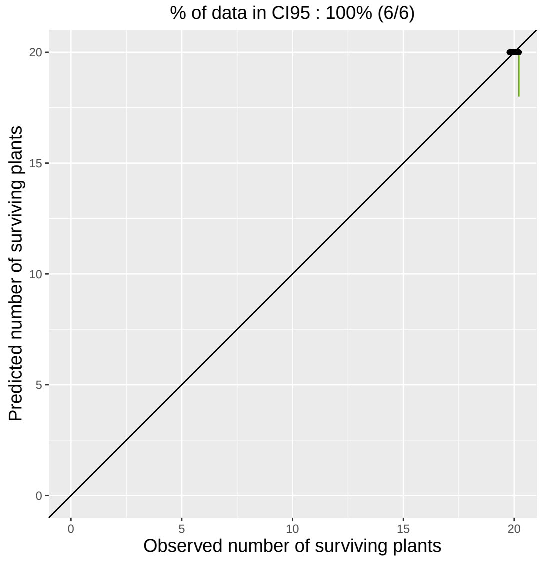

(b) Posterior predictive check (PPC)

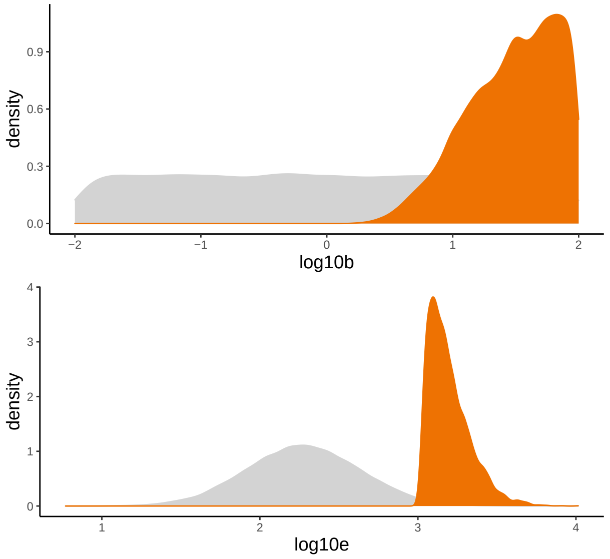

(c) Priors and posteriors

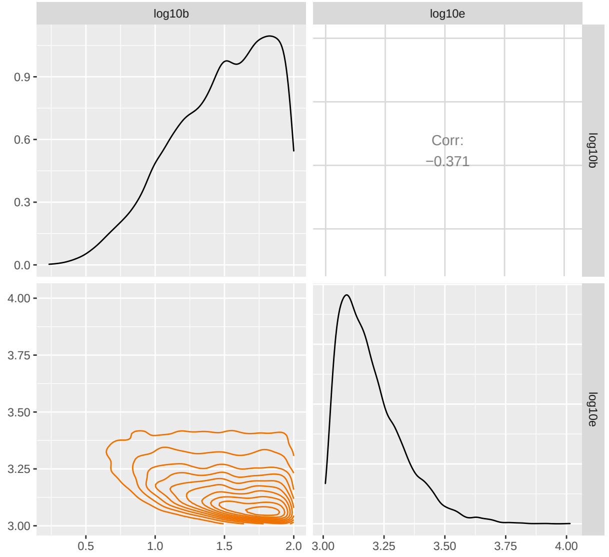

(d) Correlations between parameters

Figure 9: Dose-response curve (a), PPC (b), prior and posterior distributions (c) and correlations between parameters (d).

#### Data set: ZEAMA_VV_survival

Table 10: Summary of parameter estimates (parameter d is set to 1) for ZEAMA_VV_survival data set

| Parameter | median | Q2.5 | Q97.5 |
|---|---|---|---|
| b | 8.018 | 2.196 | 67.317 |
| e | 1219.009 | 1022.244 | 2299.026 |

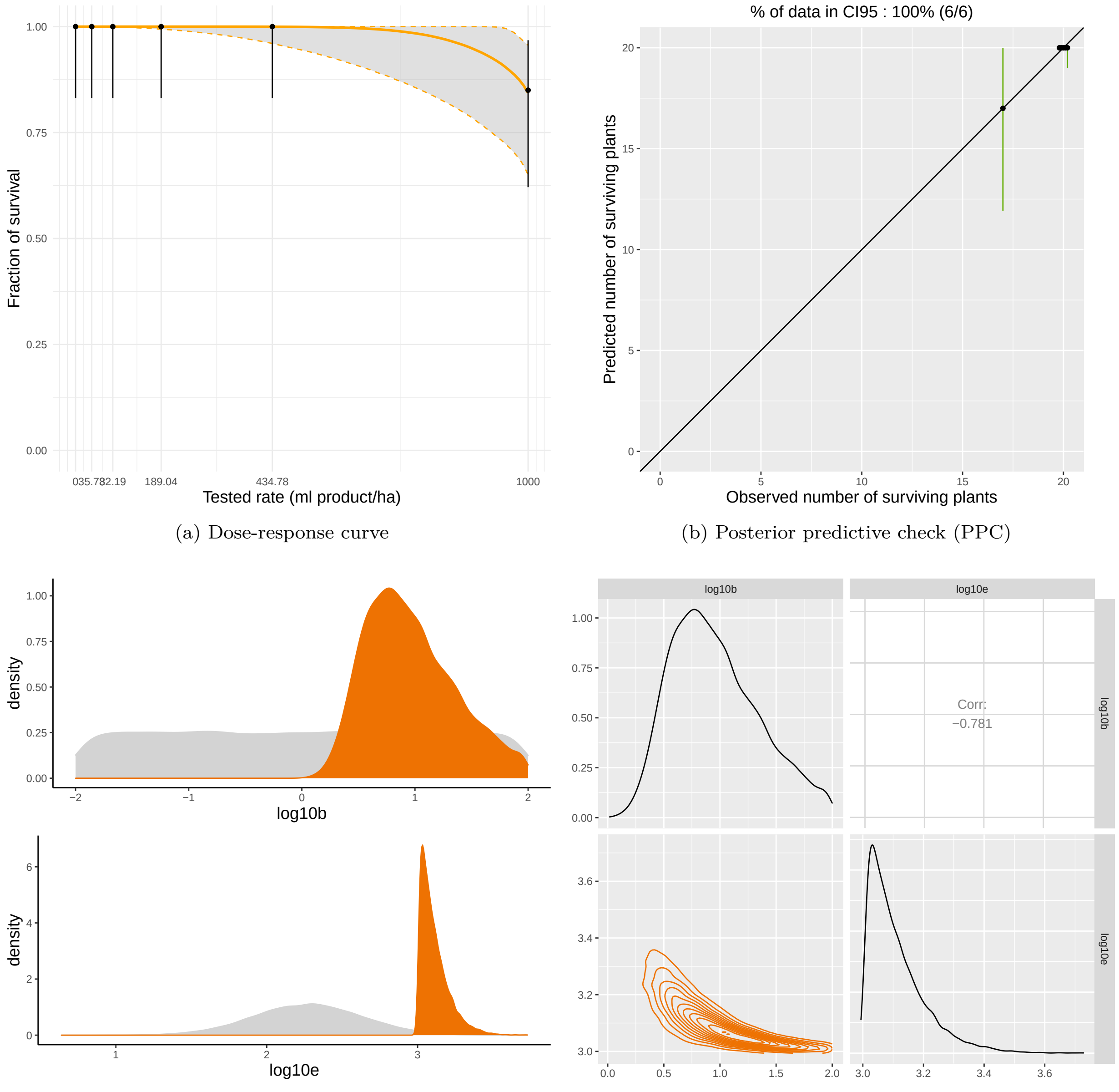

(a) Dose-response curve

(b) Posterior predictive check (PPC)

(c) Priors and posteriors

(d) Correlations between parameters

Figure 10: Dose-response curve (a), PPC (b), prior and posterior distributions (c) and correlations between parameters (d).
