## Supplementary material for "How to account for the uncertainty from standard toxicity tests in species sensitivity distributions: an example in non-target plants": Archive with reports on studies: report_VV_weight.pdf

### Dose-response analysis

Study 4

Vegetative Vigour test - shoot dry VV_weight endpoint

25 June 2020

Contact:

---

This is a report which provides results on all performed dose-response analyses for the shoot dry VV_weight endpoint of the Vegetative Vigour test for study 4.

---

### Data set: ALLCE_VV_weight

Table 1: Summary of parameter estimates for ALLCE_VV_weight data set

| Parameter | median | Q2.5 | Q97.5 |
|---|---|---|---|
| b | 0.752 | 0.153 | 3.744 |
| d | 0.209 | 0.159 | 0.262 |
| e | 150.609 | 52.123 | 370.245 |
| sigma | 0.056 | 0.043 | 0.076 |

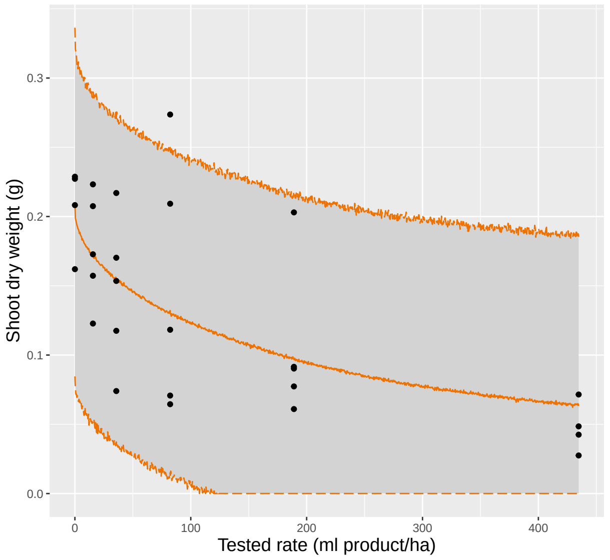

(a) Dose-response curve

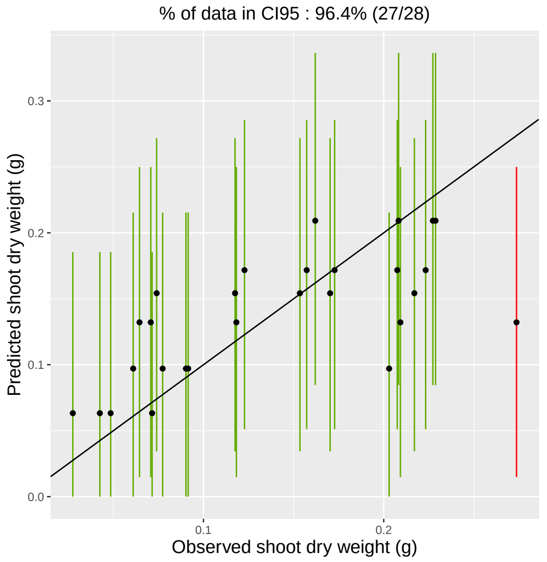

(b) Posterior predictive check (PPC)

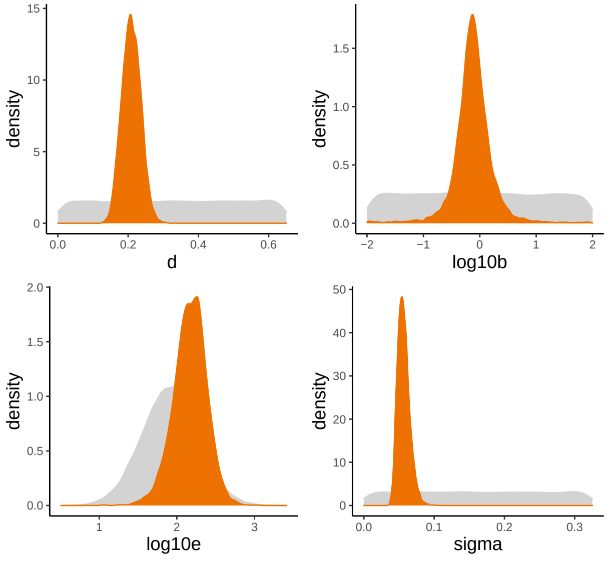

(c) Priors and posteriors

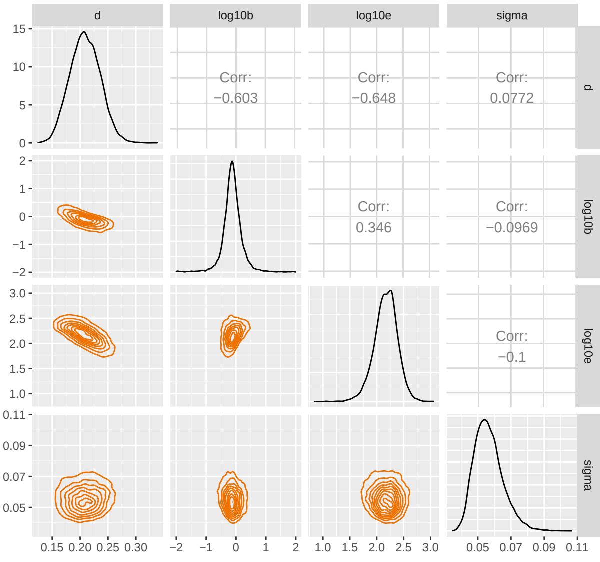

(d) Correlations between parameters

Figure 1: Dose-response curve (a), PPC (b), prior and posterior distributions (c) and correlations between parameters (d).

#### Data set: AVESA_VV_weight

Table 2: Summary of parameter estimates for AVESA_VV_weight data set

| Parameter | median | Q2.5 | Q97.5 |
|---|---|---|---|
| b | 2.507 | 1.923 | 3.346 |
| d | 2.335 | 2.178 | 2.502 |
| e | 115.250 | 101.208 | 131.442 |
| sigma | 0.193 | 0.145 | 0.275 |

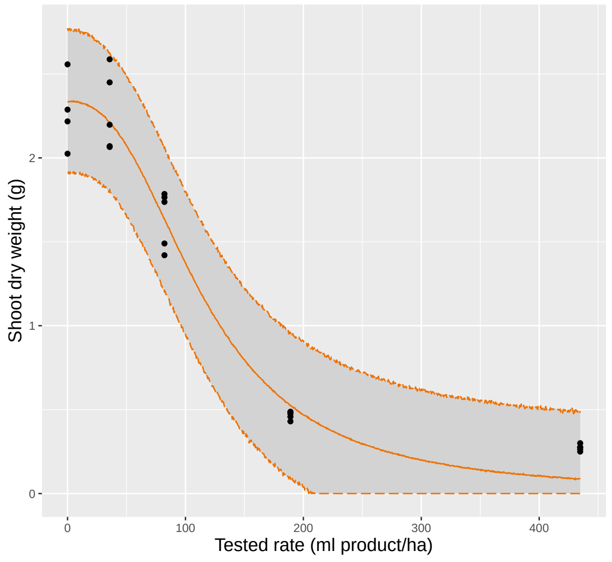

(a) Dose-response curve

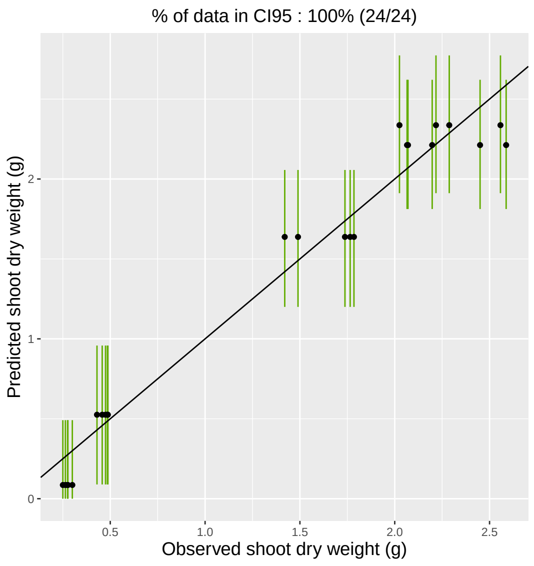

(b) Posterior predictive check (PPC)

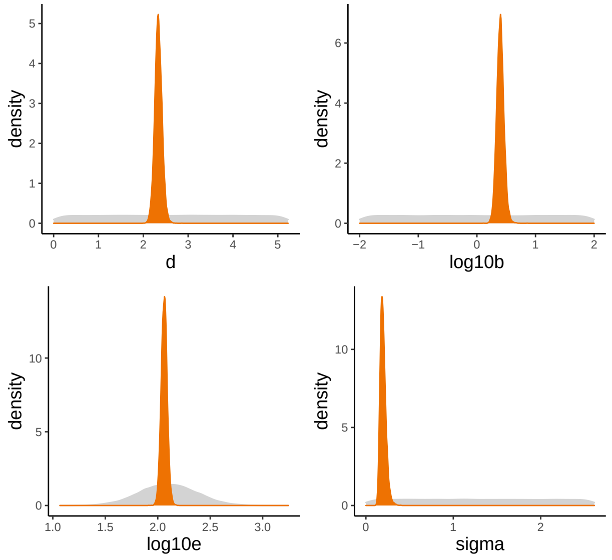

(c) Priors and posteriors

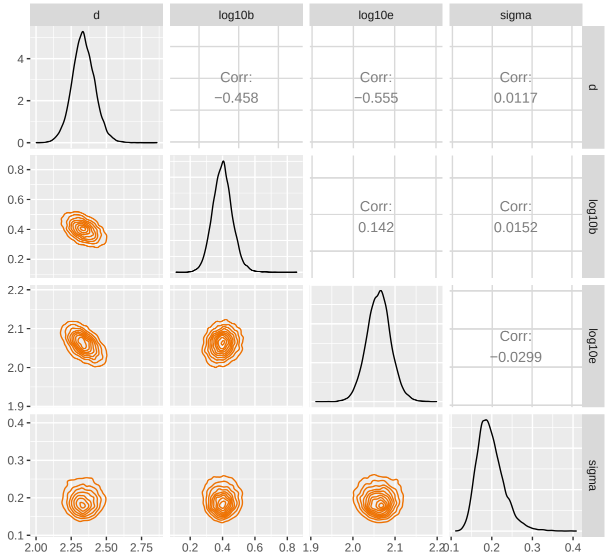

(d) Correlations between parameters

Figure 2: Dose-response curve (a), PPC (b), prior and posterior distributions (c) and correlations between parameters (d).

#### Data set: BEAVA_VV_weight

Table 3: Summary of parameter estimates for BEAVA_VV_weight data set

| Parameter | median | Q2.5 | Q97.5 |
|---|---|---|---|
| b | 1.198 | 0.888 | 1.638 |
| d | 4.245 | 3.949 | 4.536 |
| e | 8.574 | 6.797 | 10.427 |
| sigma | 0.447 | 0.371 | 0.554 |

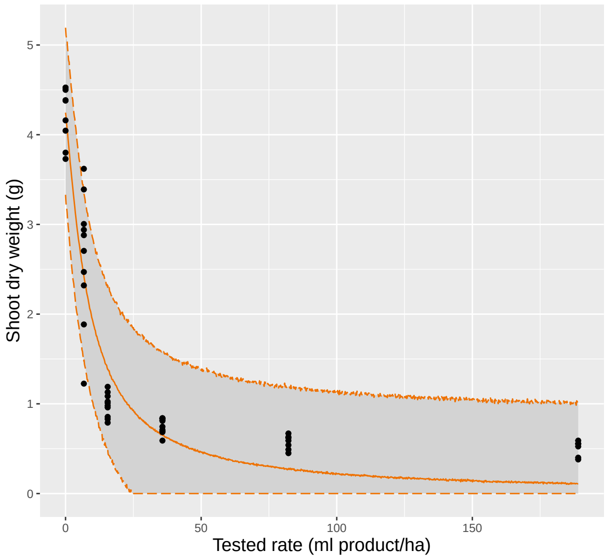

(a) Dose-response curve

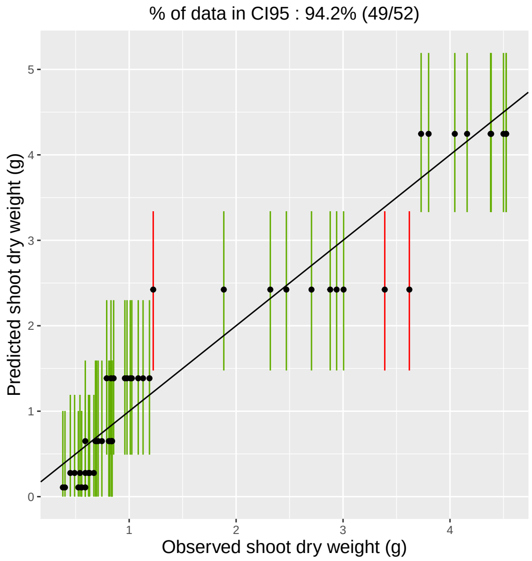

(b) Posterior predictive check (PPC)

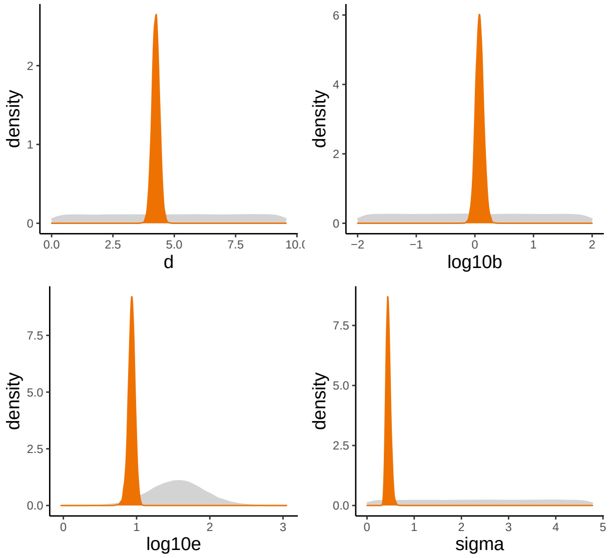

(c) Priors and posteriors

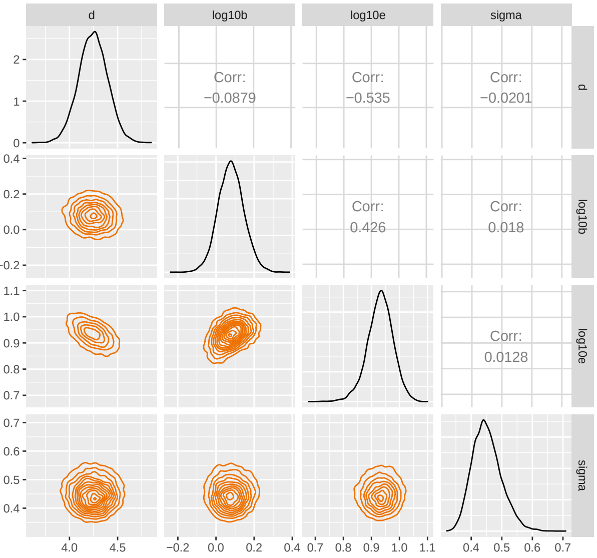

(d) Correlations between parameters

Figure 3: Dose-response curve (a), PPC (b), prior and posterior distributions (c) and correlations between parameters (d).

#### Data set: BRSNW_VV_weight

Table 4: Summary of parameter estimates for BRSNW_VV_weight data set

| Parameter | median | Q2.5 | Q97.5 |
|---|---|---|---|
| b | 1.997 | 1.606 | 2.528 |
| d | 9.193 | 8.677 | 9.732 |
| e | 19.299 | 17.089 | 21.950 |
| sigma | 0.927 | 0.774 | 1.135 |

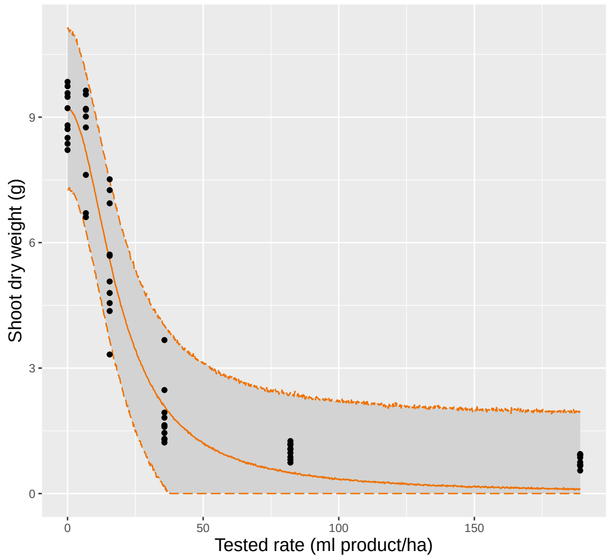

(a) Dose-response curve

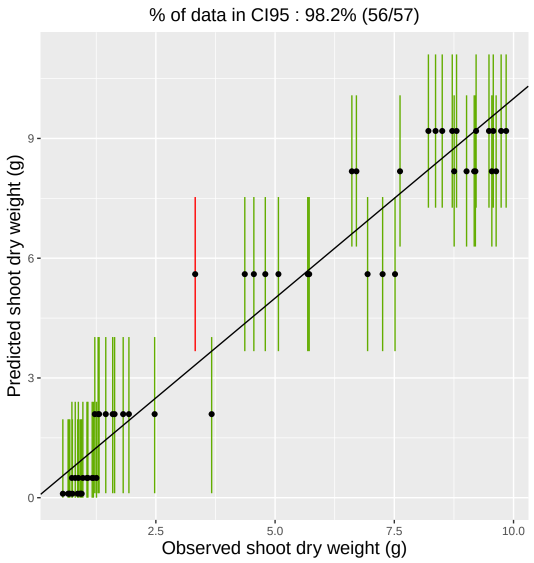

(b) Posterior predictive check (PPC)

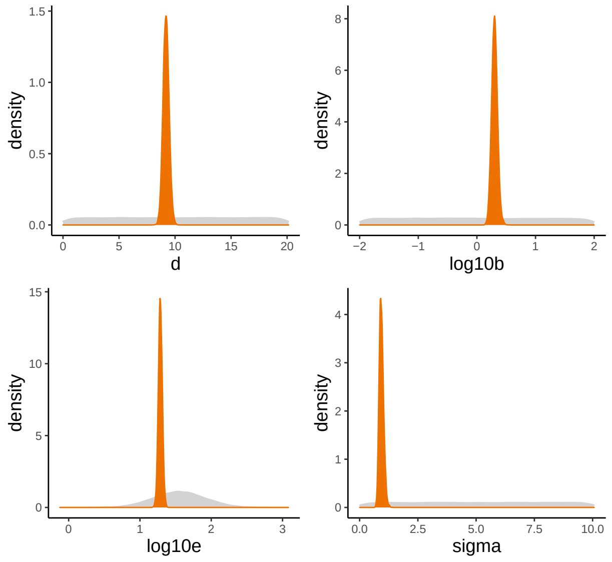

(c) Priors and posteriors

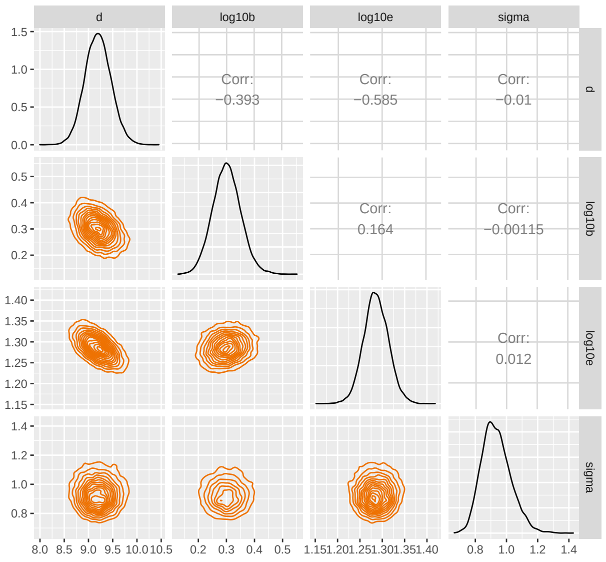

(d) Correlations between parameters

Figure 4: Dose-response curve (a), PPC (b), prior and posterior distributions (c) and correlations between parameters (d).

### Data set: CUMSA_VV_weight

Table 5: Summary of parameter estimates for CUMSA_VV_weight data set

| Parameter | median | Q2.5 | Q97.5 |
|---|---|---|---|
| b | 1.032 | 0.848 | 1.274 |
| d | 16.883 | 16.052 | 17.747 |
| e | 16.931 | 13.965 | 20.659 |
| sigma | 2.177 | 1.912 | 2.512 |

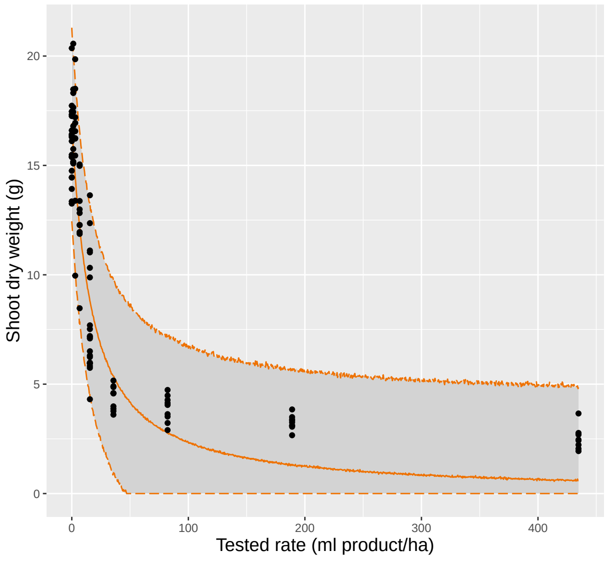

(a) Dose-response curve

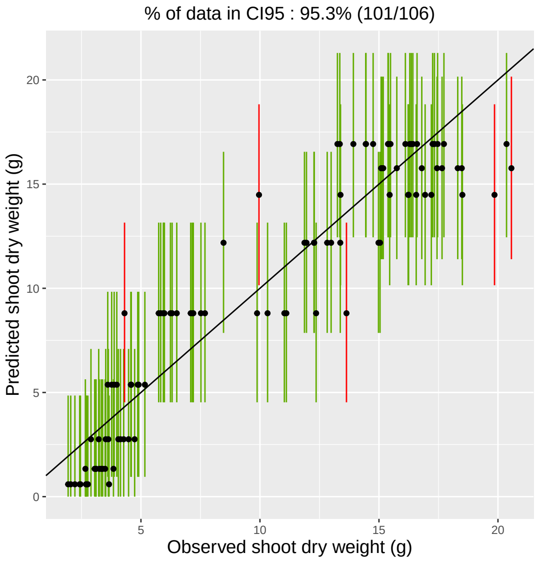

(b) Posterior predictive check (PPC)

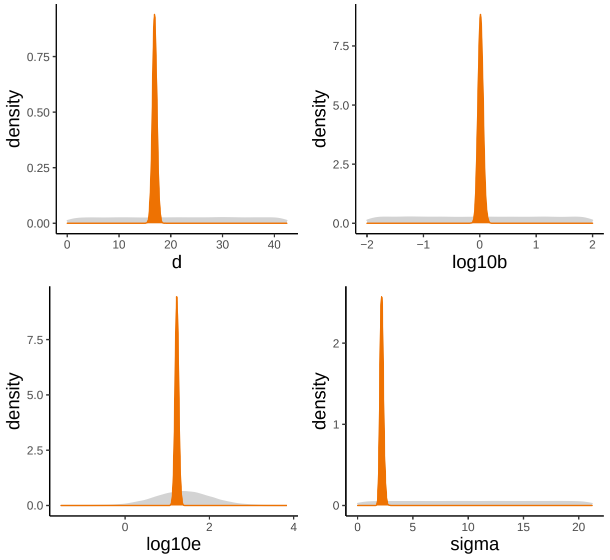

(c) Priors and posteriors

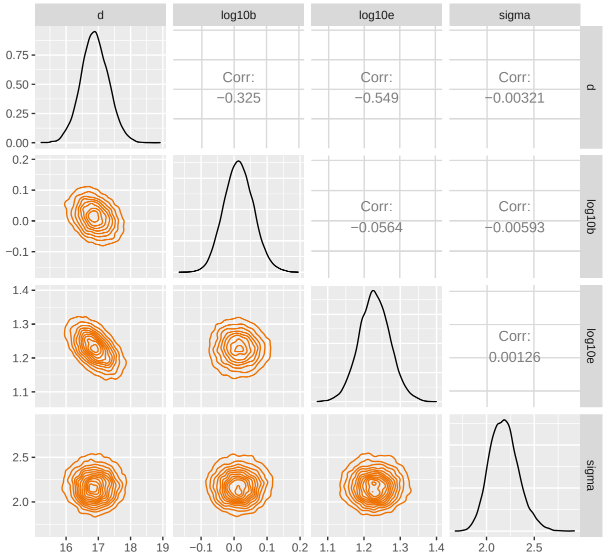

(d) Correlations between parameters

Figure 5: Dose-response curve (a), PPC (b), prior and posterior distributions (c) and correlations between parameters (d).

#### Data set: GLXMA_VV_weight

Table 6: Summary of parameter estimates for GLXMA_VV_weight data set

| Parameter | median | Q2.5 | Q97.5 |
|---|---|---|---|
| b | 1.545 | 1.279 | 1.889 |
| d | 6.491 | 6.122 | 6.851 |
| e | 33.033 | 29.031 | 37.891 |
| sigma | 0.579 | 0.482 | 0.716 |

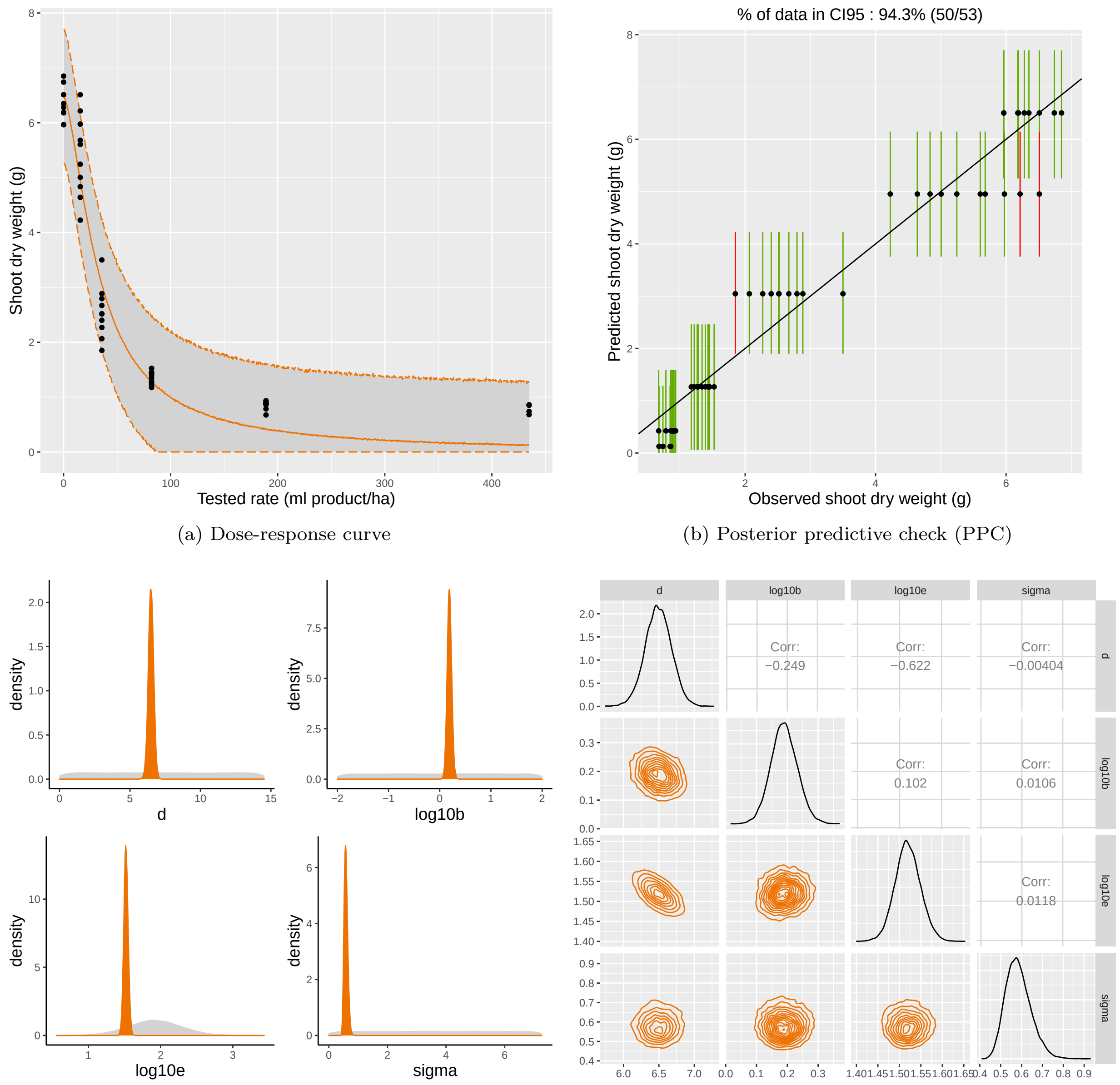

(a) Dose-response curve

(b) Posterior predictive check (PPC)

(c) Priors and posteriors

(d) Correlations between parameters

Figure 6: Dose-response curve (a), PPC (b), prior and posterior distributions (c) and correlations between parameters (d).

#### Data set: HELAN_VV_weight

Table 7: Summary of parameter estimates for HELAN_VV_weight data set

| Parameter | median | Q2.5 | Q97.5 |
|---|---|---|---|
| b | 2.308 | 1.662 | 3.130 |
| d | 3.900 | 3.591 | 4.215 |
| e | 23.907 | 20.714 | 27.804 |
| sigma | 0.461 | 0.360 | 0.626 |

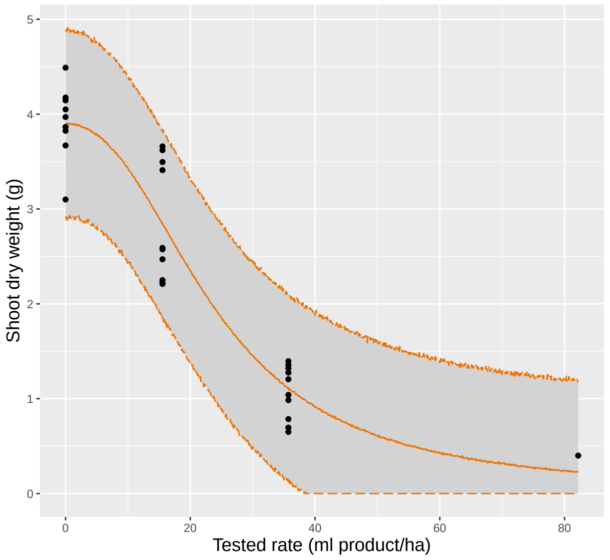

(a) Dose-response curve

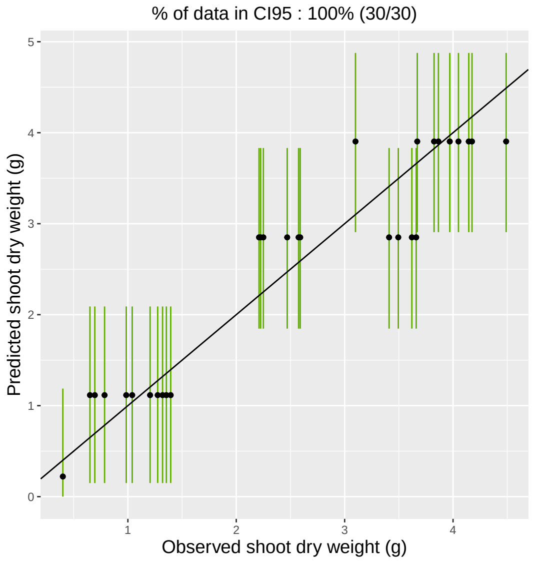

(b) Posterior predictive check (PPC)

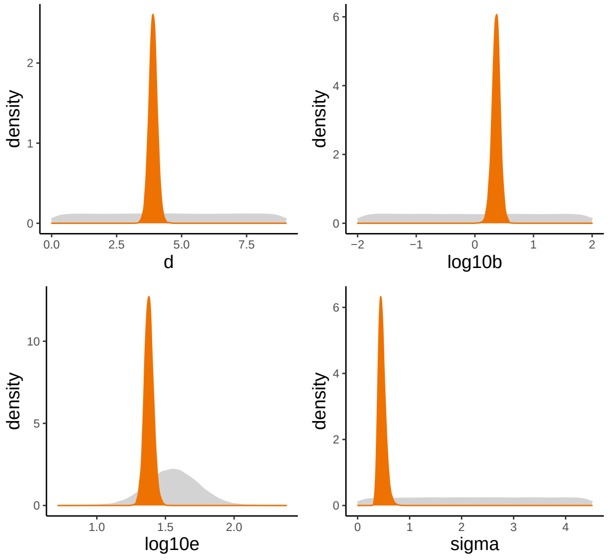

(c) Priors and posteriors

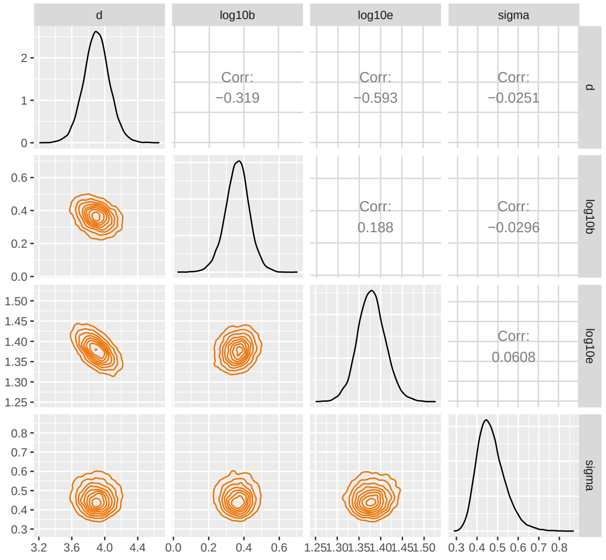

(d) Correlations between parameters

Figure 7: Dose-response curve (a), PPC (b), prior and posterior distributions (c) and correlations between parameters (d).

#### Data set: LYPES_VV_weight

Table 8: Summary of parameter estimates for LYPES_VV_weight data set

| Parameter | median | Q2.5 | Q97.5 |
|---|---|---|---|
| b | 9.088 | 3.794 | 65.630 |
| d | 6.341 | 6.044 | 6.638 |
| e | 15.172 | 14.137 | 15.608 |
| sigma | 1.032 | 0.898 | 1.211 |

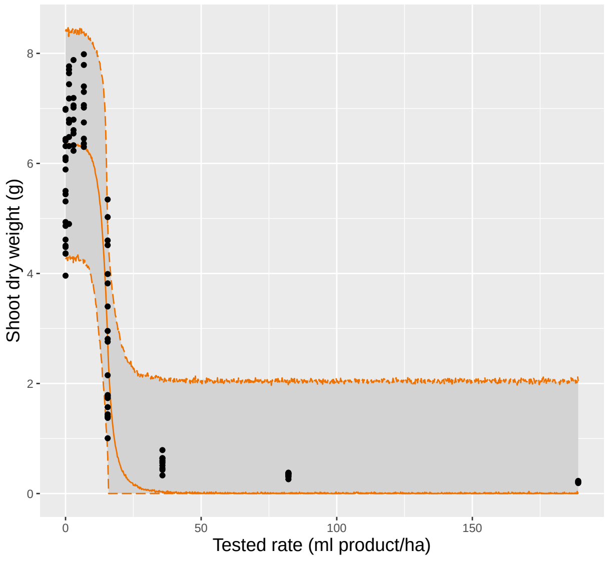

(a) Dose-response curve

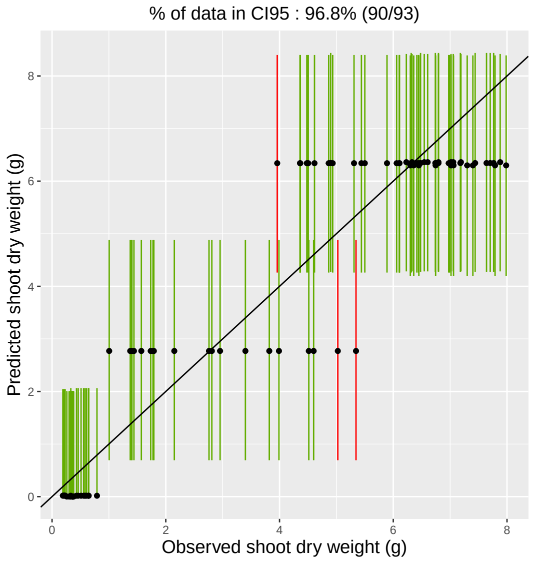

(b) Posterior predictive check (PPC)

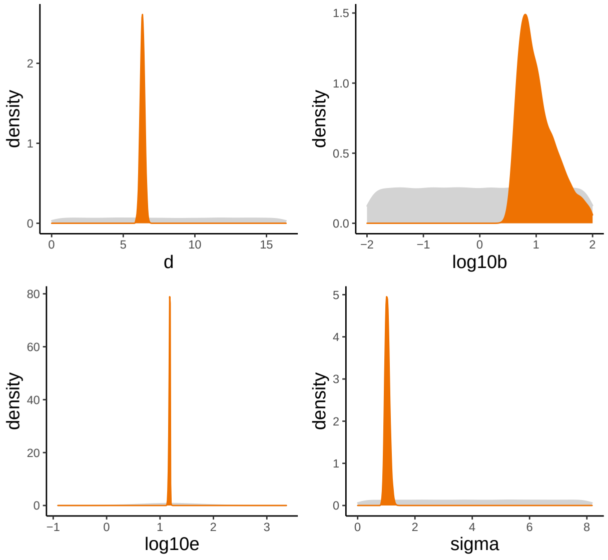

(c) Priors and posteriors

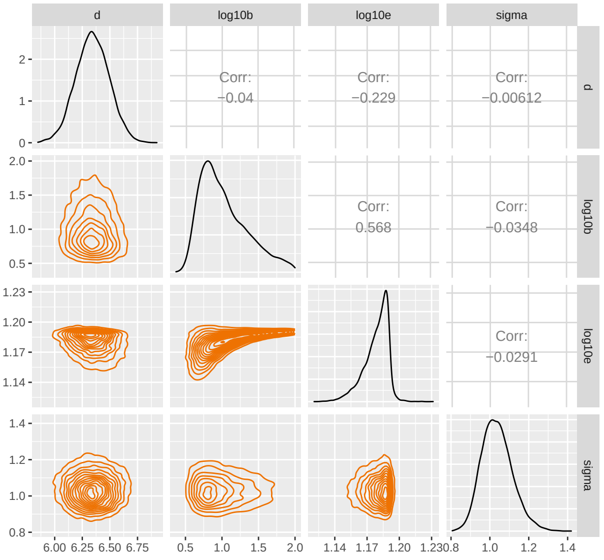

(d) Correlations between parameters

Figure 8: Dose-response curve (a), PPC (b), prior and posterior distributions (c) and correlations between parameters (d).

#### Data set: TRZAW_VV_weight

Table 9: Summary of parameter estimates for TRZAW_VV_weight data set

| Parameter | median | Q2.5 | Q97.5 |
|---|---|---|---|
| b | 2.260 | 0.927 | 49.330 |
| d | 1.757 | 1.688 | 1.841 |
| e | 2036.538 | 1042.973 | 4831.957 |
| sigma | 0.129 | 0.098 | 0.176 |

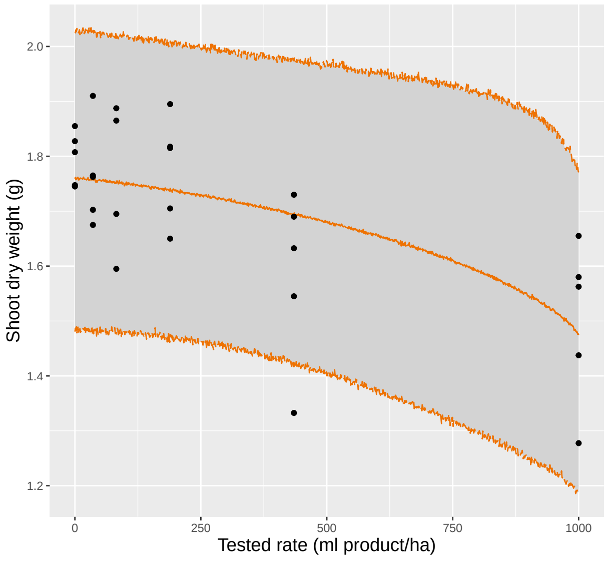

(a) Dose-response curve

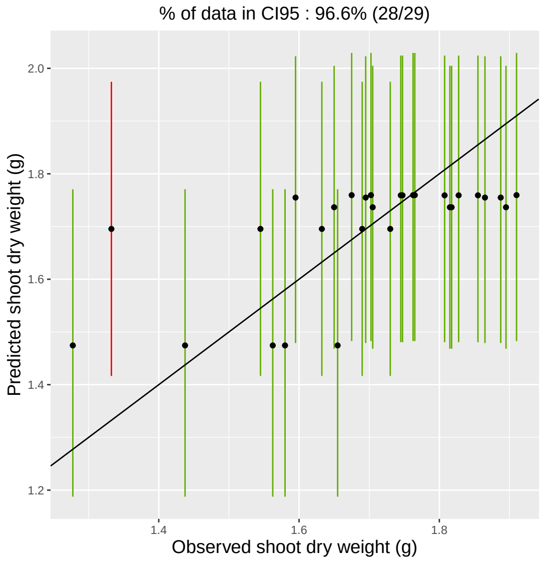

(b) Posterior predictive check (PPC)

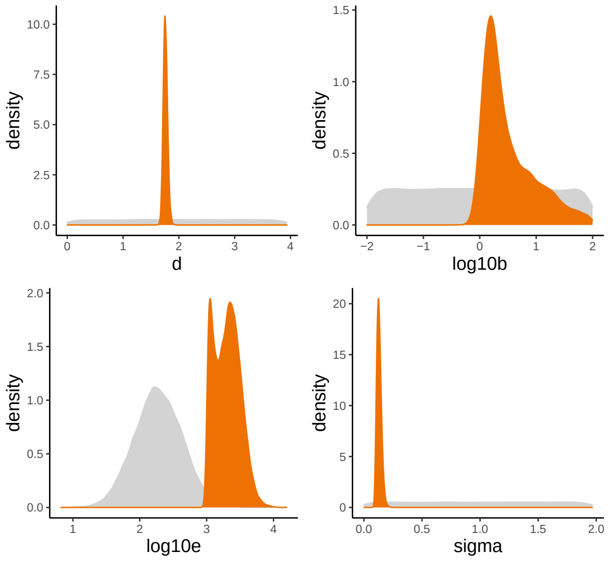

(c) Priors and posteriors

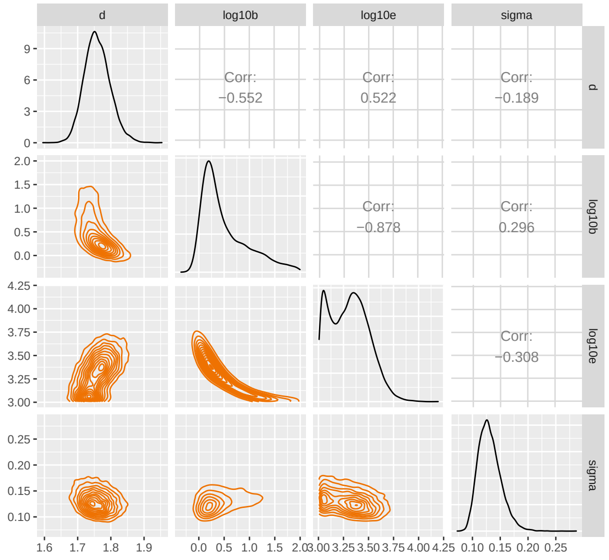

(d) Correlations between parameters

Figure 9: Dose-response curve (a), PPC (b), prior and posterior distributions (c) and correlations between parameters (d).

#### Data set: ZEAMA_VV_weight

Table 10: Summary of parameter estimates for ZEAMA_VV_weight data set

| Parameter | median | Q2.5 | Q97.5 |
|---|---|---|---|
| b | 2.950 | 2.235 | 5.011 |
| d | 10.714 | 10.269 | 11.160 |
| e | 403.884 | 367.325 | 443.155 |
| sigma | 1.188 | 0.993 | 1.462 |

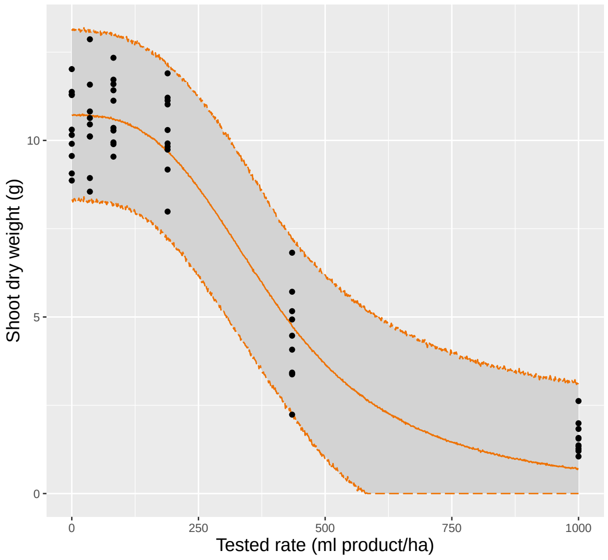

(a) Dose-response curve

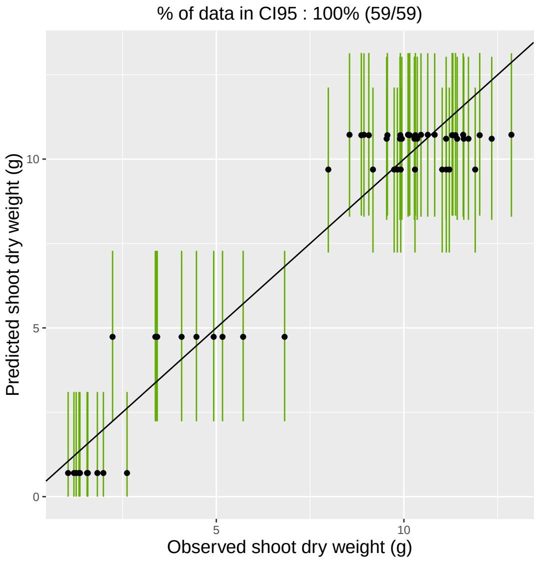

(b) Posterior predictive check (PPC)

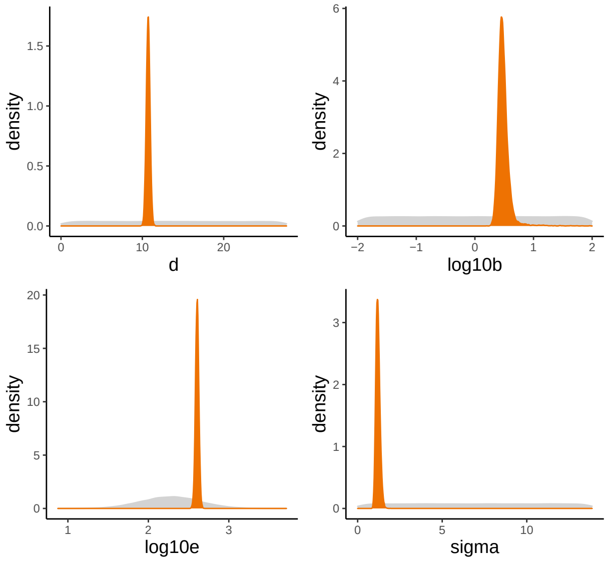

(c) Priors and posteriors

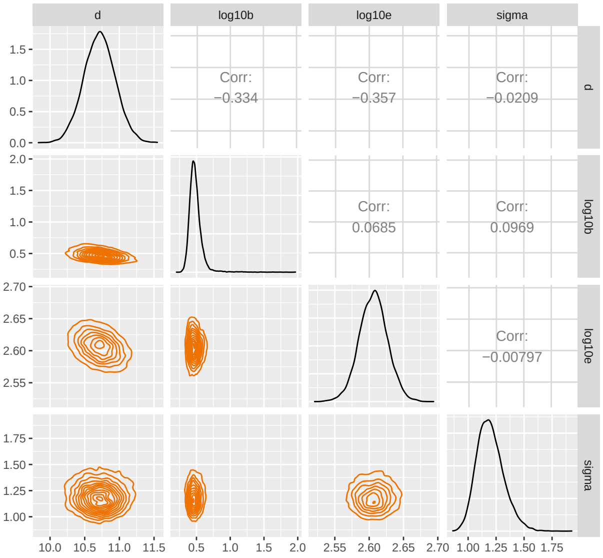

(d) Correlations between parameters

Figure 10: Dose-response curve (a), PPC (b), prior and posterior distributions (c) and correlations between parameters (d).
