## Supplementary material for "How to account for the uncertainty from standard toxicity tests in species sensitivity distributions: an example in non-target plants": Archive with reports on studies: report_SE_emergence.pdf

### Dose-response analyses

Study 5

Seedling Emergence test - emergence endpoint

25 June 2020

Contact:

---

This is a report which provides results on all performed dose-response analyses for the emergence endpoint of the Seedling Emergence test for study 5.

---

#### Data set: ALLCE_SE_emergence

Table 1: Summary of parameter estimates for ALLCE_SE_emergence data set

| Parameter | median | Q2.5 | Q97.5 |
|---|---|---|---|
| b | 5.748 | 1.380 | 66.785 |
| d | 0.786 | 0.695 | 0.862 |
| e | 1.780 | 1.472 | 3.436 |

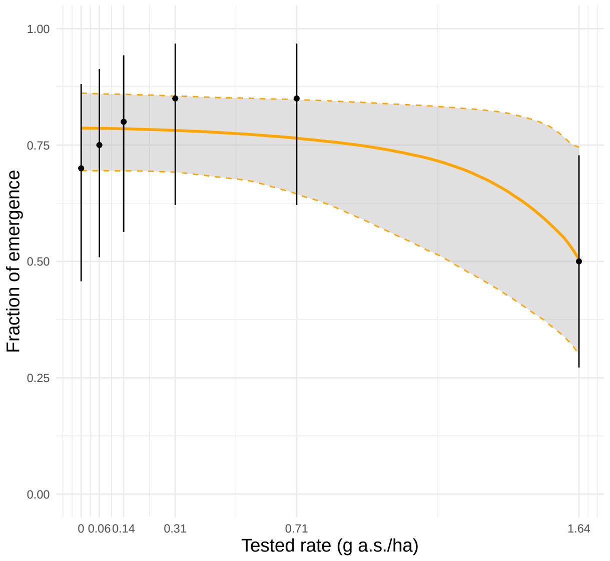

(a) Dose-response curve

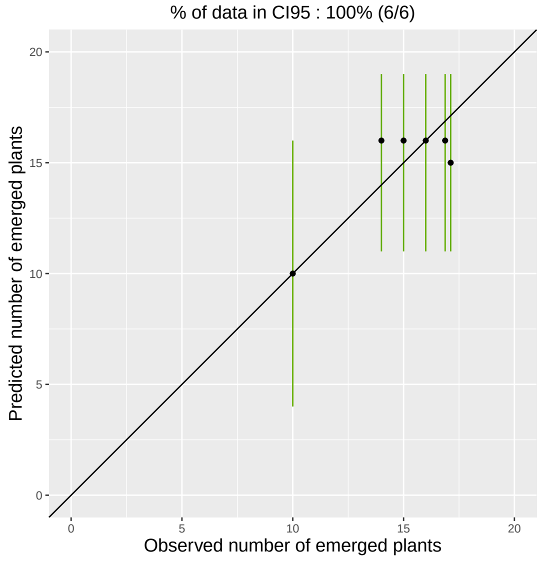

(b) Posterior predictive check (PPC)

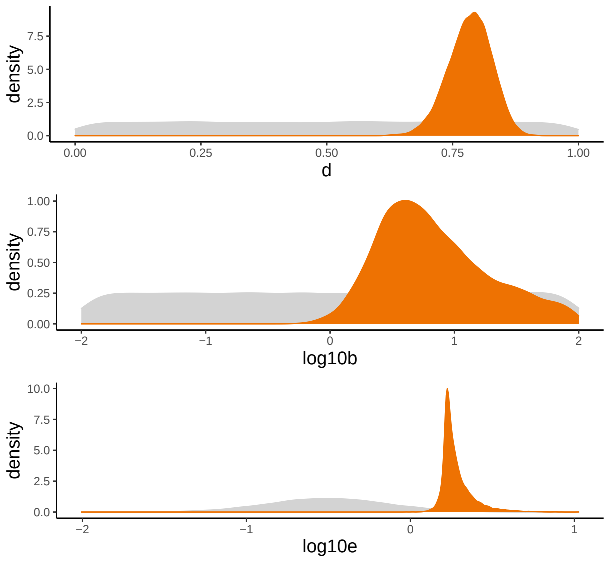

(c) Priors and posteriors

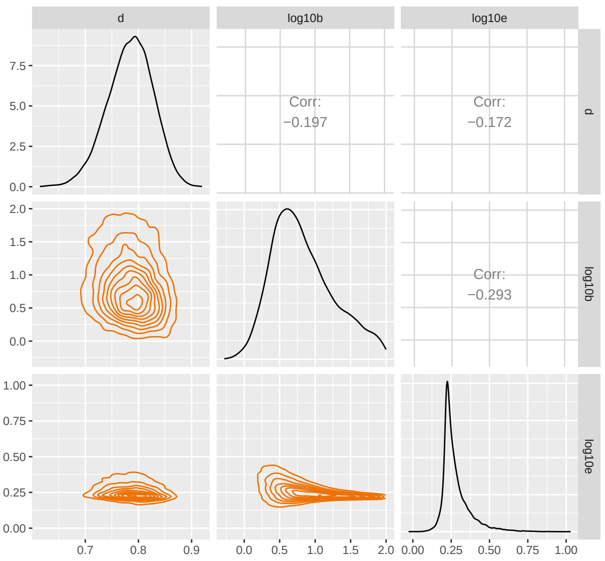

(d) Correlations between parameters

Figure 1: Dose-response curve (a), PPC (b), prior and posterior distributions (c) and correlations between parameters (d).

#### Data set: AVESA_SE_emergence

Table 2: Summary of parameter estimates for AVESA_SE_emergence data set

| Parameter | median | Q2.5 | Q97.5 |
|---|---|---|---|
| b | 24.649 | 3.344 | 93.220 |
| d | 0.981 | 0.945 | 0.997 |
| e | 12.190 | 9.144 | 27.694 |

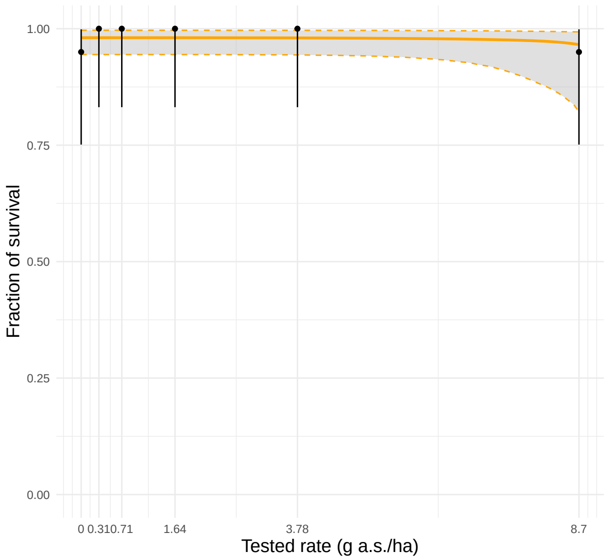

(a) Dose-response curve

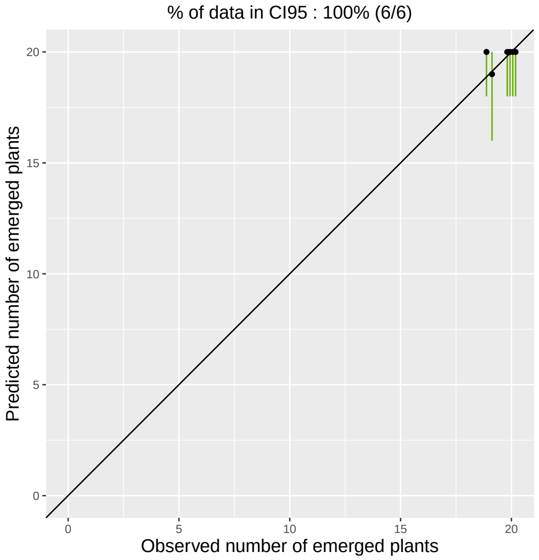

(b) Posterior predictive check (PPC)

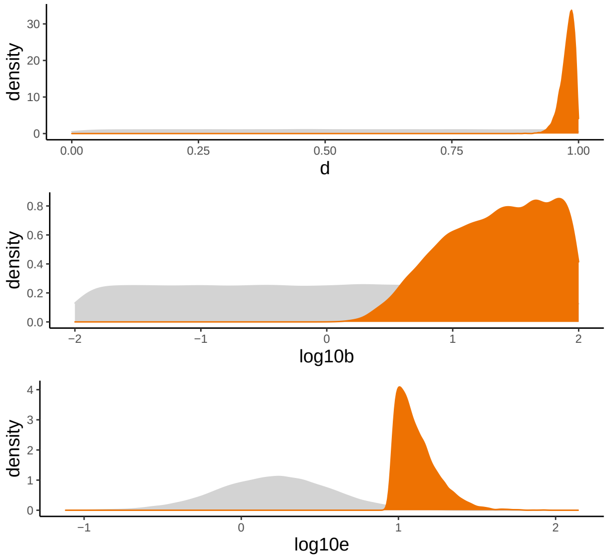

(c) Priors and posteriors

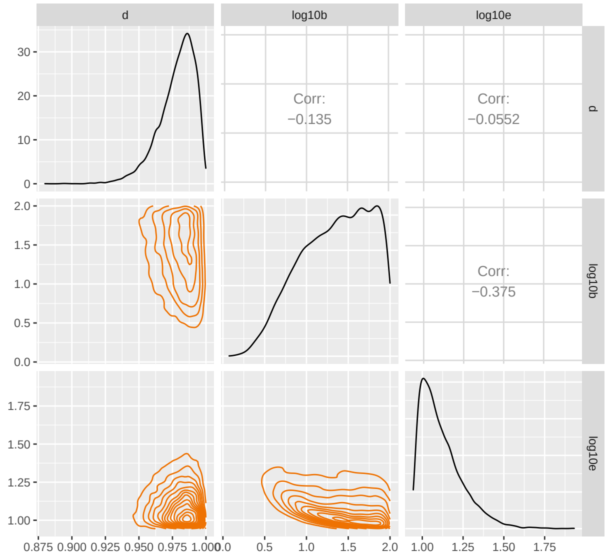

(d) Correlations between parameters

Figure 2: Dose-response curve (a), PPC (b), prior and posterior distributions (c) and correlations between parameters (d).

#### Data set: BEAVA_SE_emergence

Table 3: Summary of parameter estimates (parameter d is set to 1) for BEAVA_SE_emergence data set

| Parameter | median | Q2.5 | Q97.5 |
|---|---|---|---|
| b | 2.339 | 1.552 | 3.353 |
| e | 1.039 | 0.788 | 1.364 |

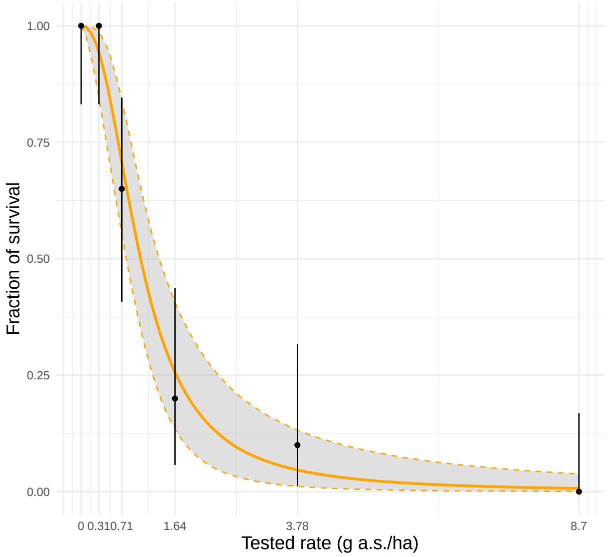

(a) Dose-response curve

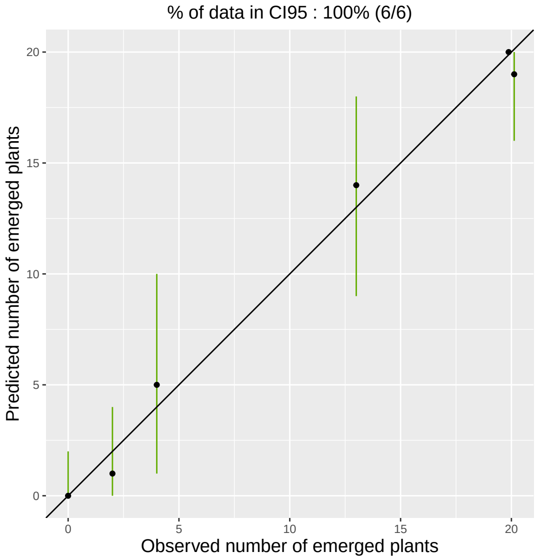

(b) Posterior predictive check (PPC)

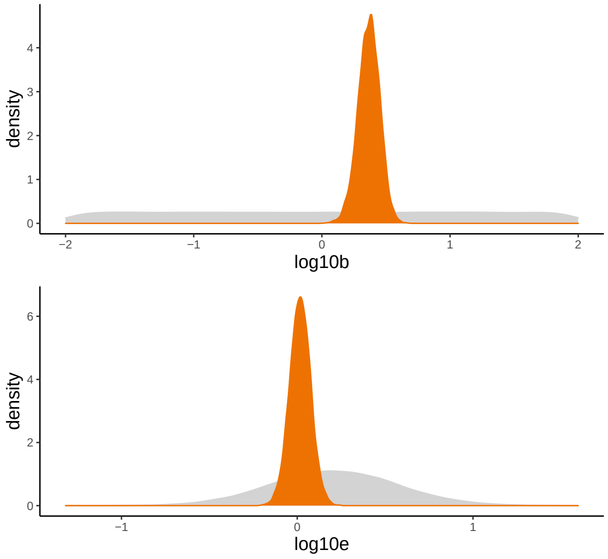

(c) Priors and posteriors

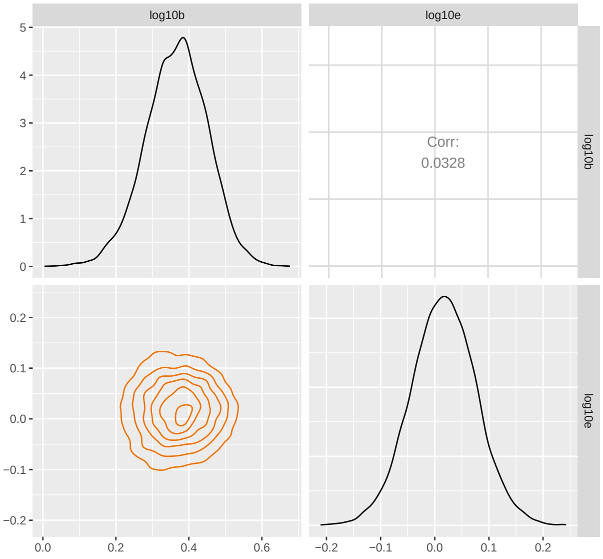

(d) Correlations between parameters

Figure 3: Dose-response curve (a), PPC (b), prior and posterior distributions (c) and correlations between parameters (d).

#### Data set: BRSNW_SE_emergence

Table 4: Summary of parameter estimates for BRSNW_SE_emergence data set

| Parameter | median | Q2.5 | Q97.5 |
|---|---|---|---|
| b | 4.407 | 2.338 | 12.679 |
| d | 0.956 | 0.892 | 0.989 |
| e | 1.012 | 0.800 | 1.417 |

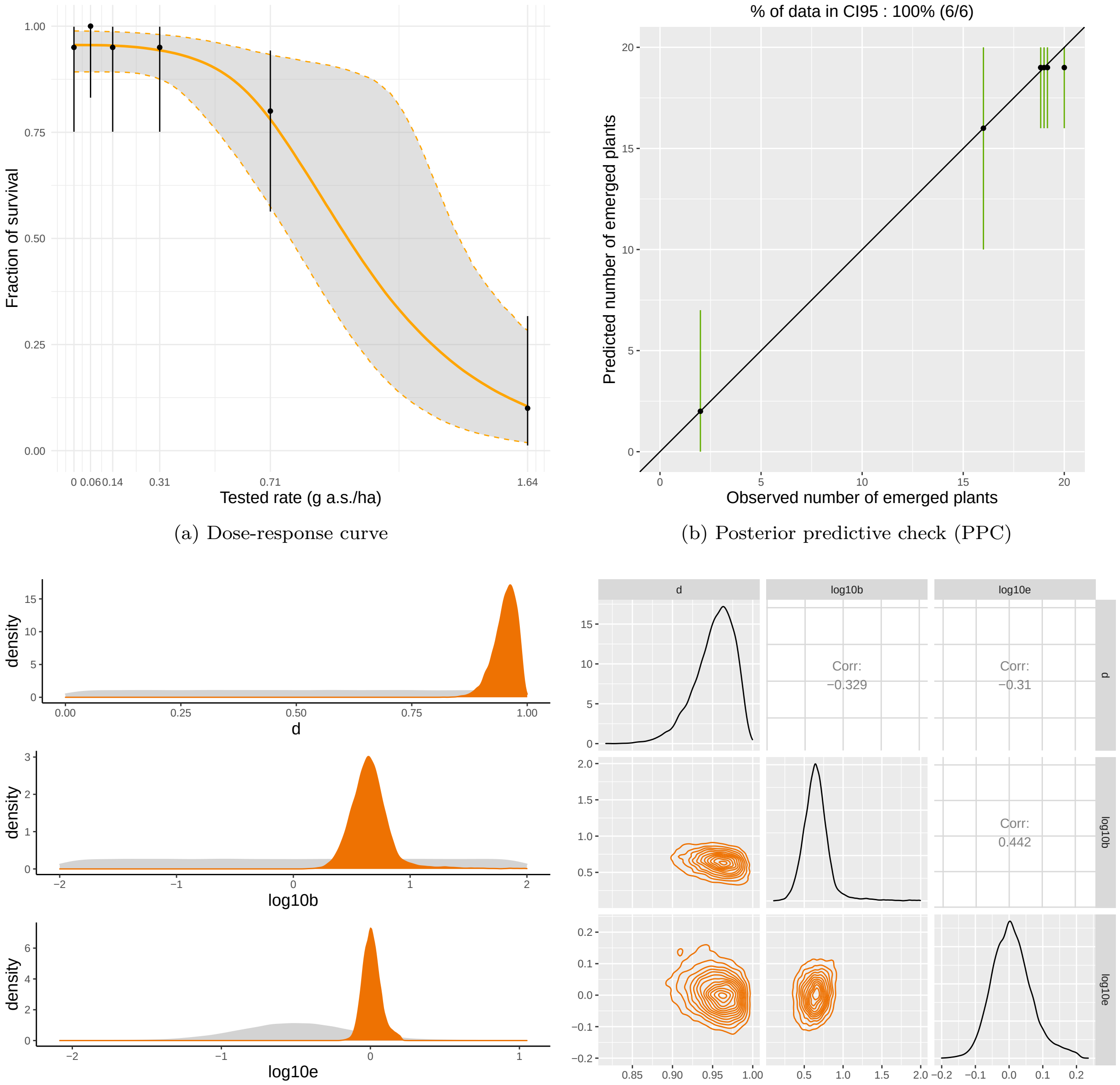

(a) Dose-response curve

(b) Posterior predictive check (PPC)

(c) Priors and posteriors

(d) Correlations between parameters

Figure 4: Dose-response curve (a), PPC (b), prior and posterior distributions (c) and correlations between parameters (d).

#### Data set: CUMSA_SE_emergence

Table 5: Summary of parameter estimates for CUMSA_SE_emergence data set

| Parameter | median | Q2.5 | Q97.5 |
|---|---|---|---|
| b | 7.652 | 0.931 | 85.180 |
| d | 0.877 | 0.799 | 0.955 |
| e | 11.905 | 8.907 | 27.043 |

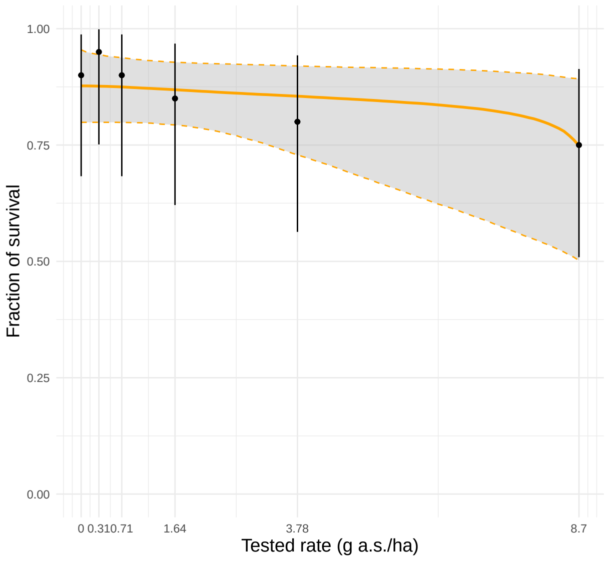

(a) Dose-response curve

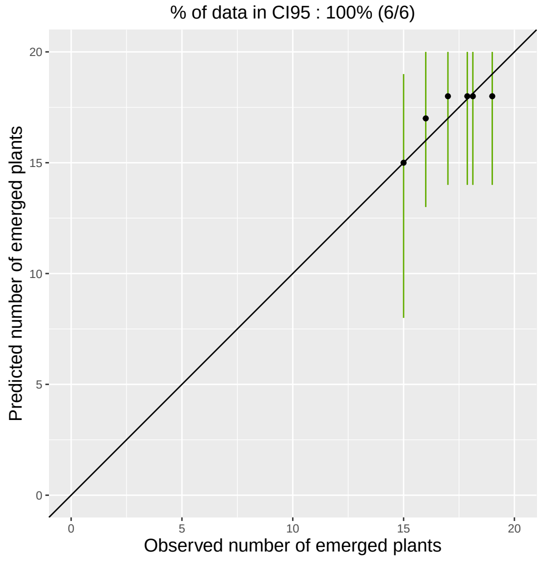

(b) Posterior predictive check (PPC)

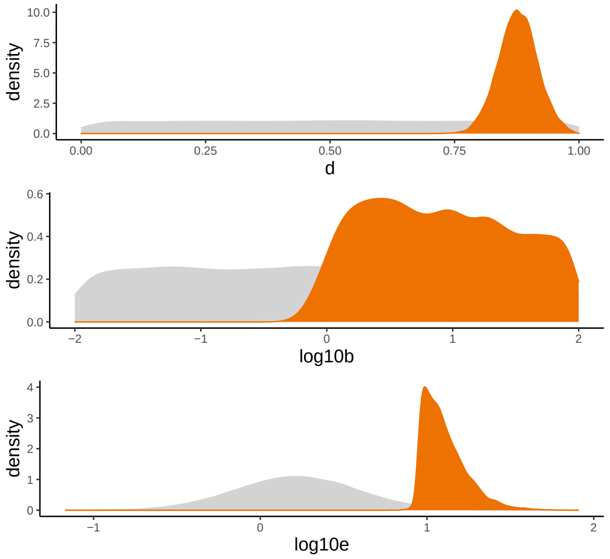

(c) Priors and posteriors

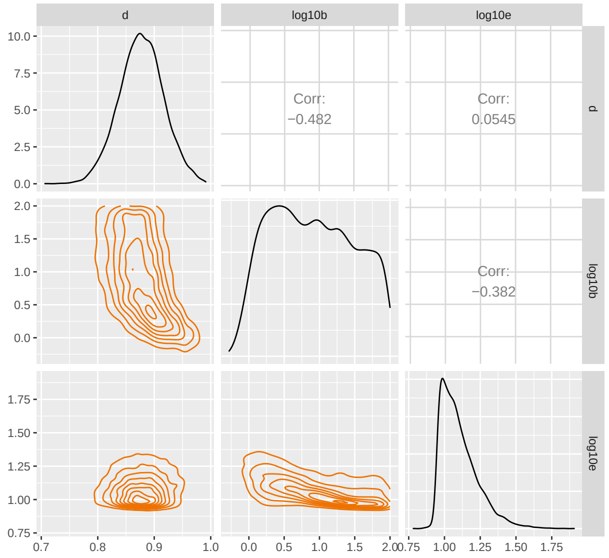

(d) Correlations between parameters

Figure 5: Dose-response curve (a), PPC (b), prior and posterior distributions (c) and correlations between parameters (d).

#### Data set: GLXMA_SE_emergence

Table 6: Summary of parameter estimates (parameter d is set to 1) for GLXMA_SE_emergence data set

| Parameter | median | Q2.5 | Q97.5 |
|---|---|---|---|
| b | 1.492 | 0.894 | 2.260 |
| e | 5.960 | 4.188 | 9.939 |

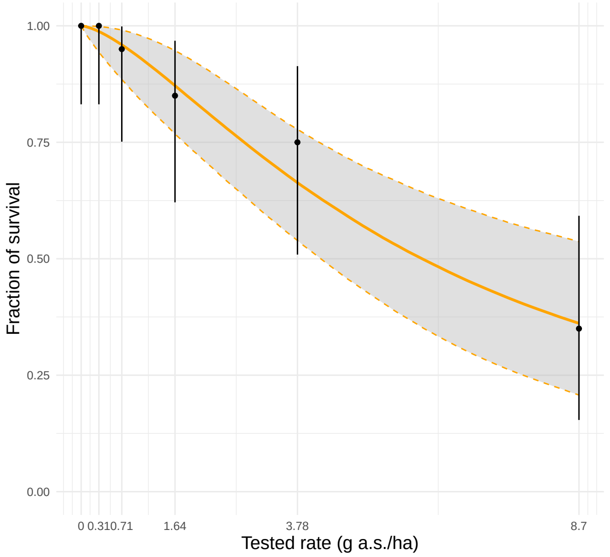

(a) Dose-response curve

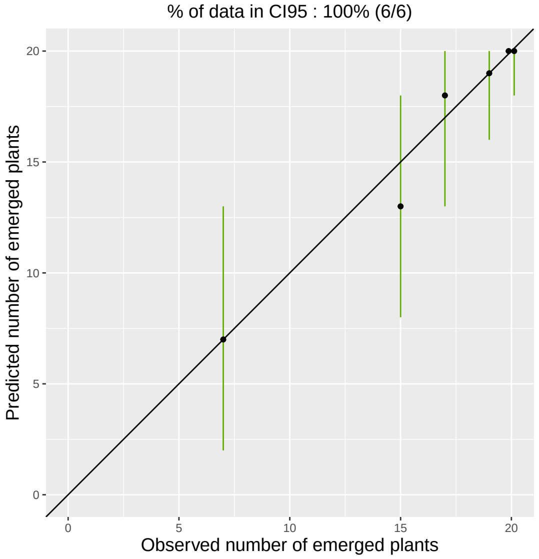

(b) Posterior predictive check (PPC)

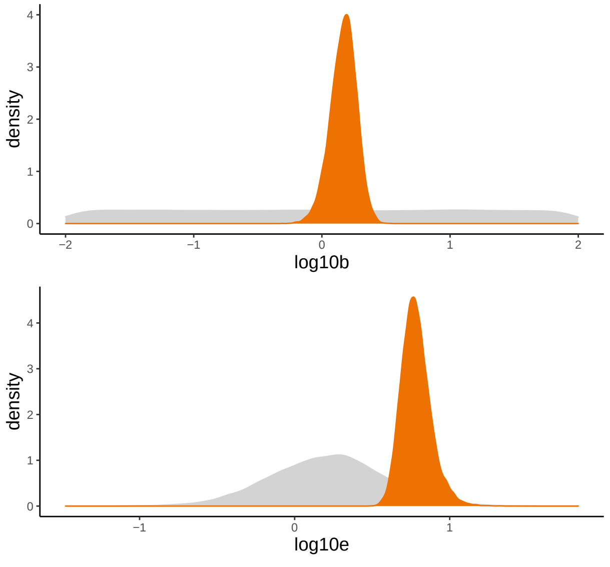

(c) Priors and posteriors

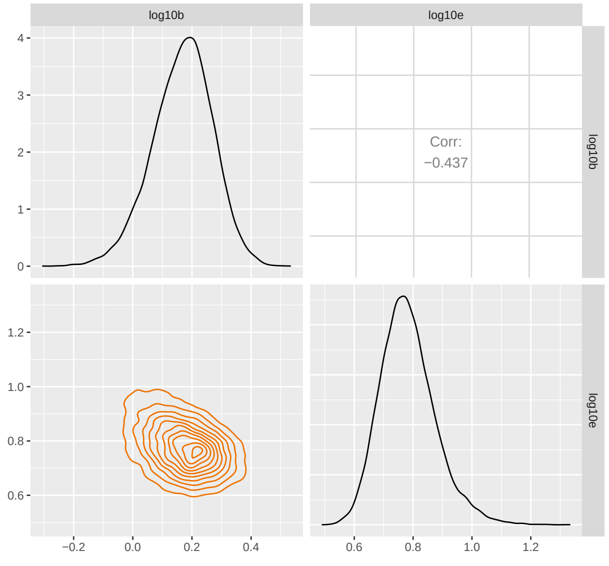

(d) Correlations between parameters

Figure 6: Dose-response curve (a), PPC (b), prior and posterior distributions (c) and correlations between parameters (d).

#### Data set: HELAN_SE_emergence

Table 7: Summary of parameter estimates for HELAN_SE_emergence data set

| Parameter | median | Q2.5 | Q97.5 |
|---|---|---|---|
| b | 1.961 | 1.214 | 2.977 |
| d | 0.960 | 0.896 | 0.990 |
| e | 11.260 | 8.234 | 16.475 |

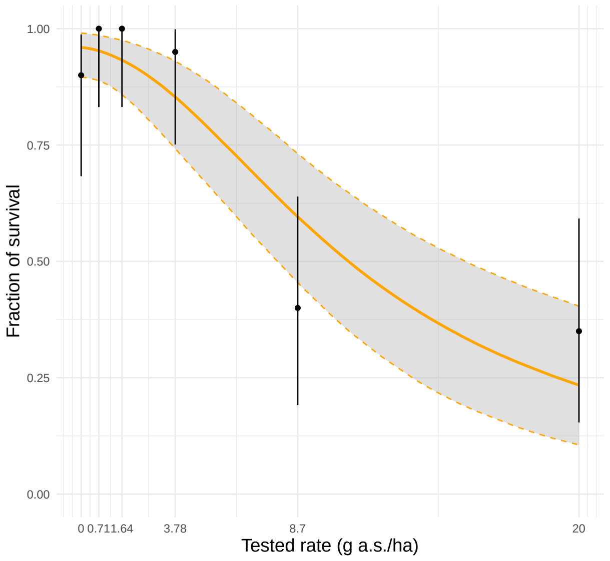

(a) Dose-response curve

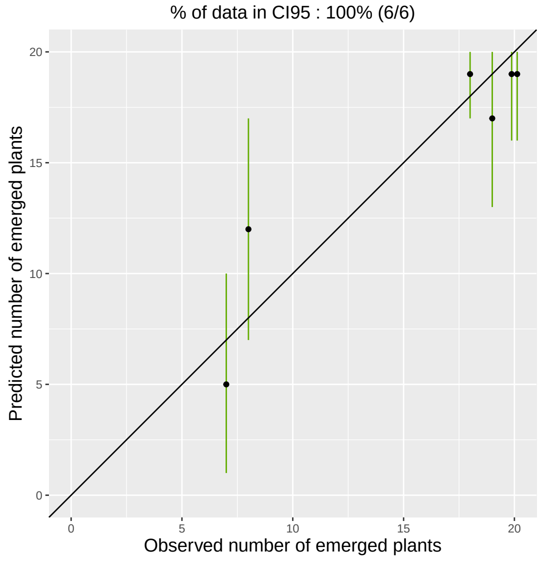

(b) Posterior predictive check (PPC)

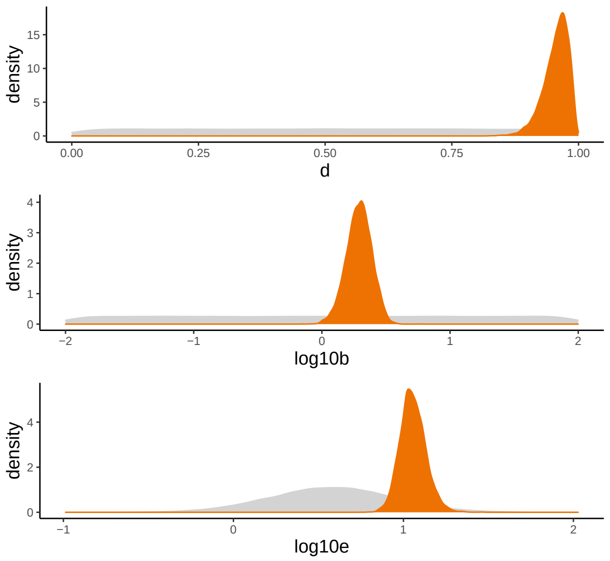

(c) Priors and posteriors

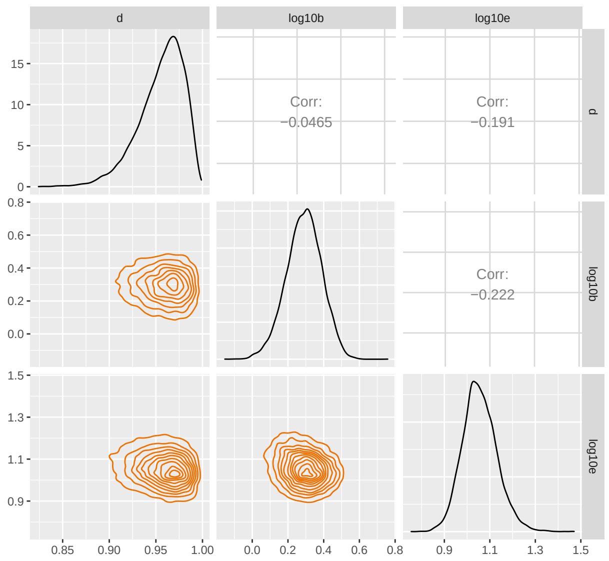

(d) Correlations between parameters

Figure 7: Dose-response curve (a), PPC (b), prior and posterior distributions (c) and correlations between parameters (d).

### Data set: LOLPE_SE_emergence

Table 8: Summary of parameter estimates for LOLPE_SE_emergence data set

| Parameter | median | Q2.5 | Q97.5 |
|---|---|---|---|
| b | 1.807 | 0.948 | 3.242 |
| d | 0.886 | 0.795 | 0.954 |
| e | 1.002 | 0.680 | 1.601 |

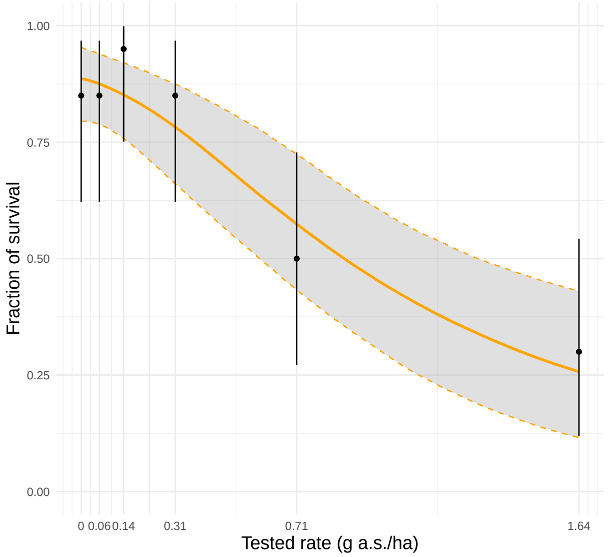

(a) Dose-response curve

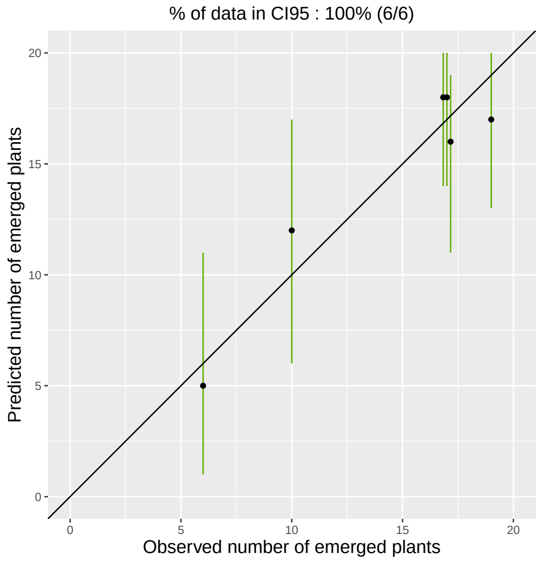

(b) Posterior predictive check (PPC)

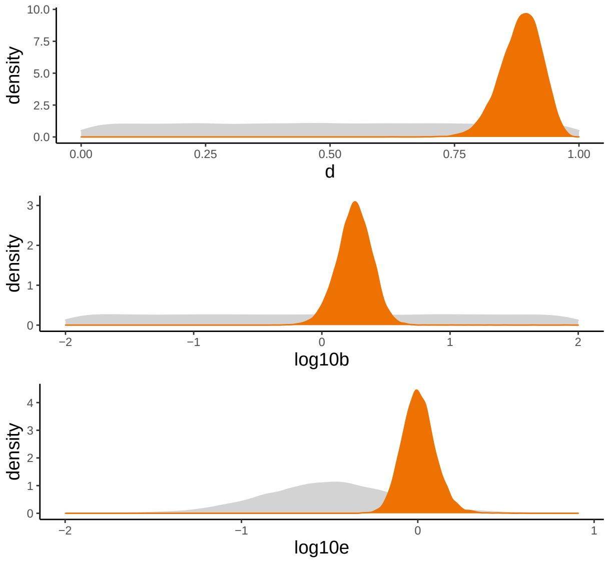

(c) Priors and posteriors

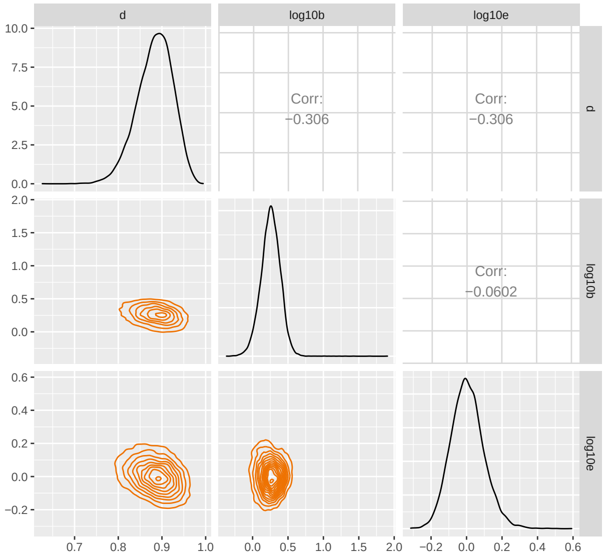

(d) Correlations between parameters

Figure 8: Dose-response curve (a), PPC (b), prior and posterior distributions (c) and correlations between parameters (d).

### Data set: LYPES_SE_emergence

Table 9: Summary of parameter estimates for LYPES_SE_emergence data set

| Parameter | median | Q2.5 | Q97.5 |
|---|---|---|---|
| b | 3.340 | 2.166 | 5.625 |
| d | 0.958 | 0.875 | 0.994 |
| e | 0.565 | 0.443 | 0.719 |

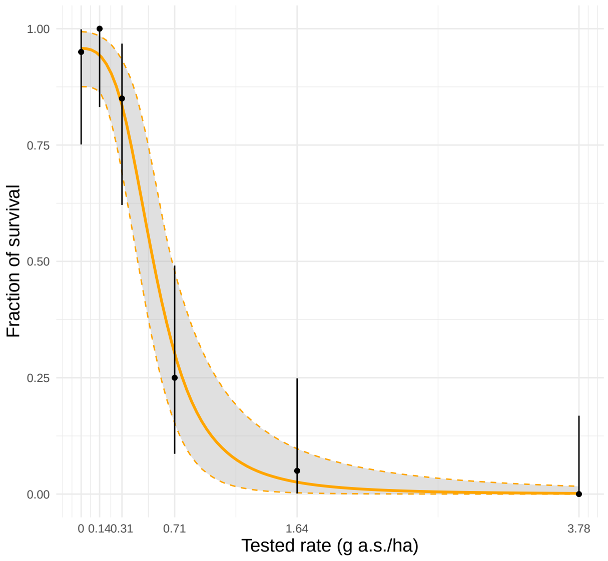

(a) Dose-response curve

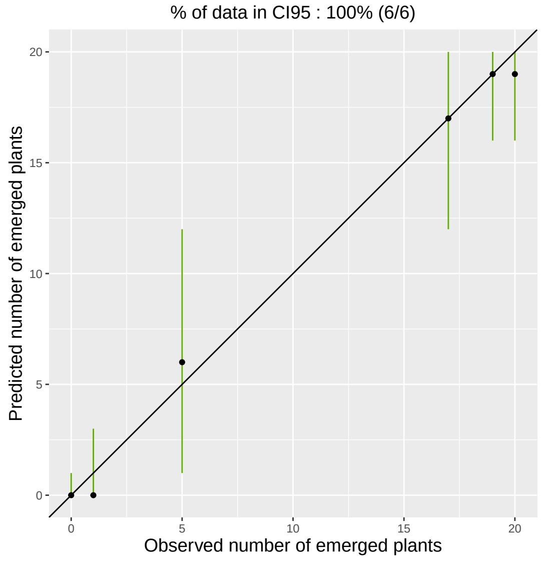

(b) Posterior predictive check (PPC)

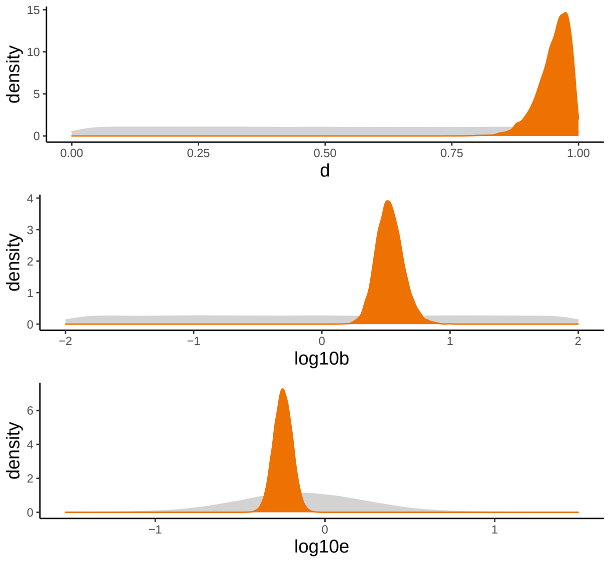

(c) Priors and posteriors

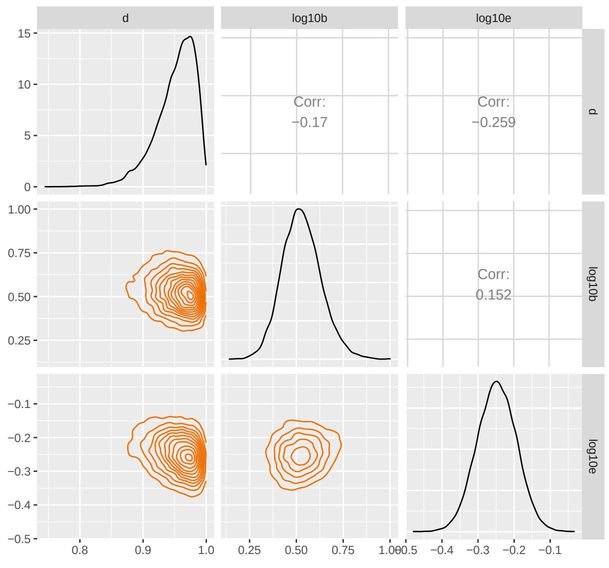

(d) Correlations between parameters

Figure 9: Dose-response curve (a), PPC (b), prior and posterior distributions (c) and correlations between parameters (d).

#### Data set: ZEAMA_SE_emergence

Table 10: Summary of parameter estimates (parameter d is set to 1) for ZEAMA_SE_emergence data set

| Parameter | median | Q2.5 | Q97.5 |
|---|---|---|---|
| b | 1.663 | 0.986 | 2.825 |
| e | 56.881 | 31.018 | 137.113 |

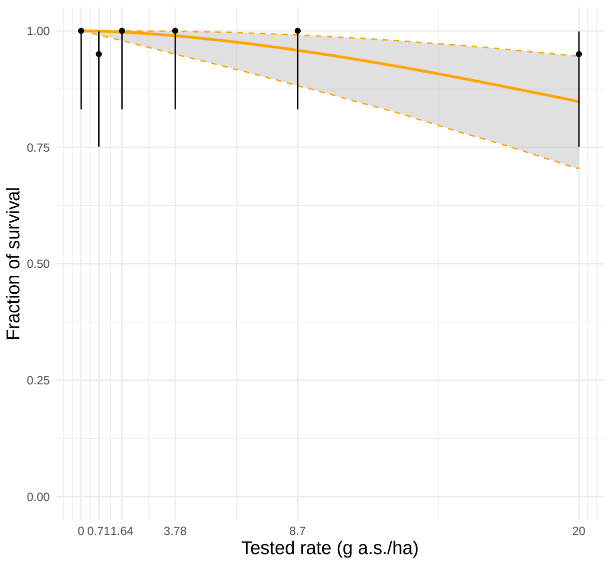

(a) Dose-response curve

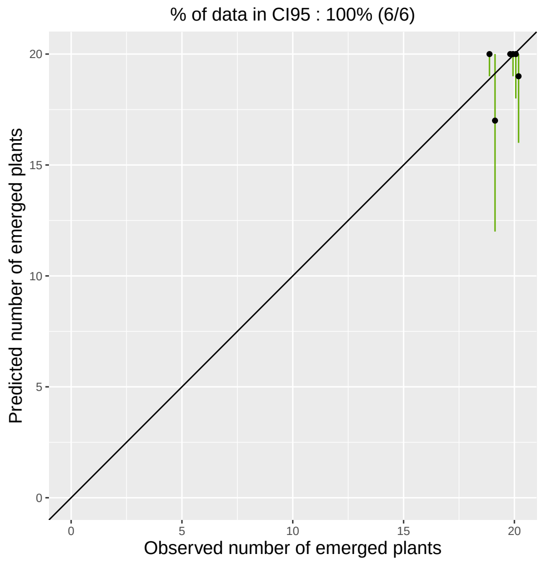

(b) Posterior predictive check (PPC)

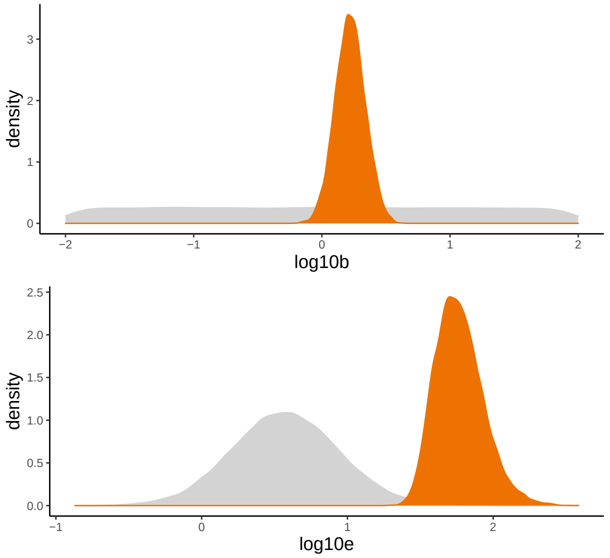

(c) Priors and posteriors

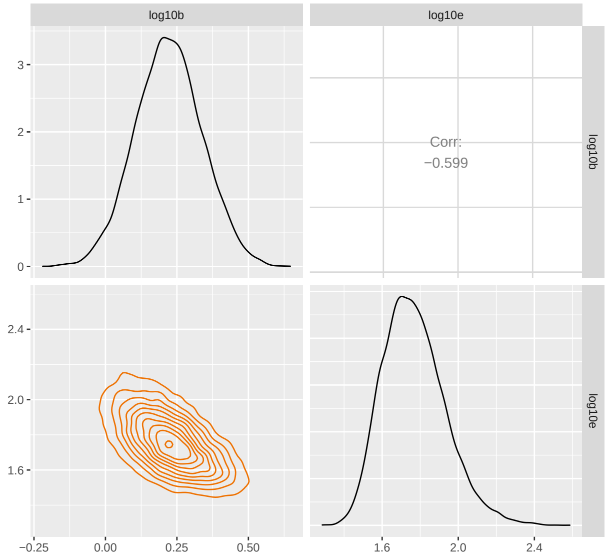

(d) Correlations between parameters

Figure 10: Dose-response curve (a), PPC (b), prior and posterior distributions (c) and correlations between parameters (d).
