## Supplementary material for "How to account for the uncertainty from standard toxicity tests in species sensitivity distributions: an example in non-target plants": Archive with reports on studies: report_SE_weight.pdf

### Dose-response analysis

#### Study 5

#### Seedling Emergence test - shoot dry SE_weight endpoint

25 June 2020

Contact:

---

This is a report which provides results on all performed dose-response analyses for the shoot dry SE_weight endpoint of the Seedling Emergence test for study 5.

---

#### Data set: ALLCE_SE_weight

Table 1: Summary of parameter estimates for ALLCE_SE_weight data set

| Parameter | median | Q2.5 | Q97.5 |
|---|---|---|---|
| b | 1.068 | 0.484 | 2.297 |
| d | 0.042 | 0.035 | 0.049 |
| e | 0.648 | 0.352 | 1.217 |
| sigma | 0.009 | 0.007 | 0.012 |

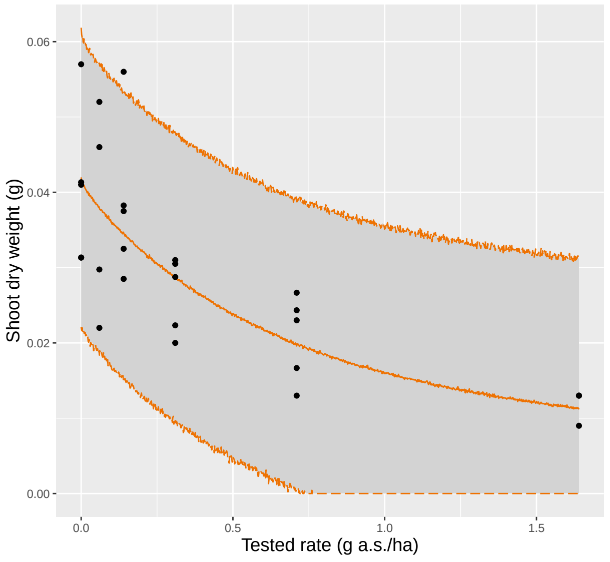

(a) Dose-response curve

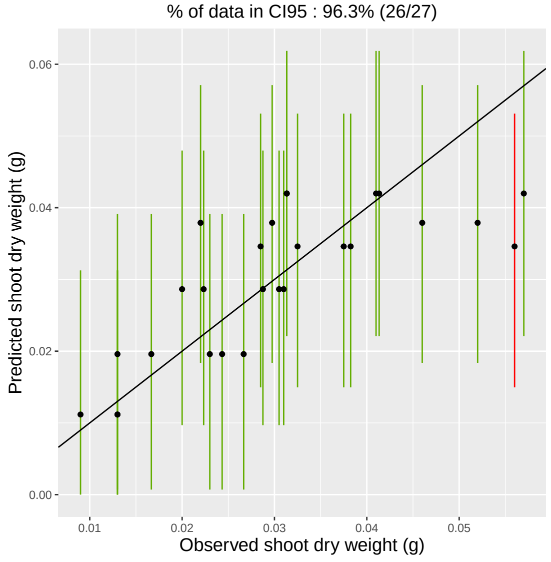

(b) Posterior predictive check (PPC)

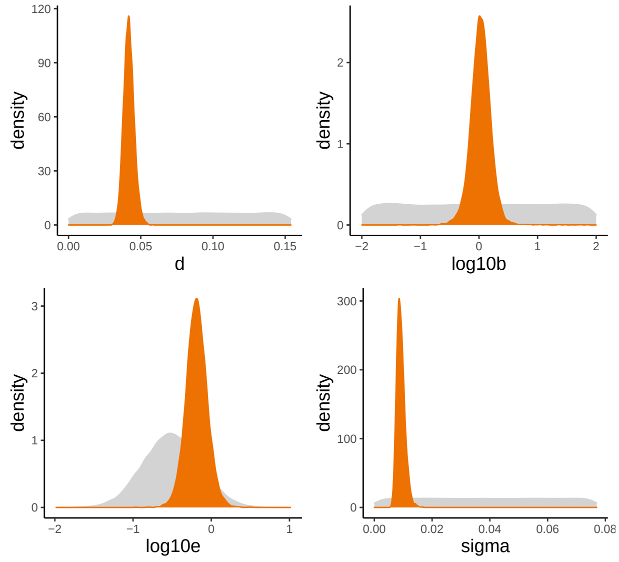

(c) Priors and posteriors

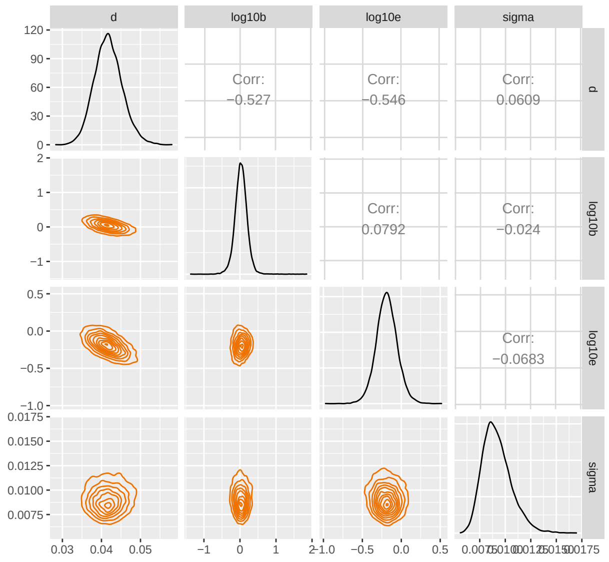

(d) Correlations between parameters

Figure 1: Dose-response curve (a), PPC (b), prior and posterior distributions (c) and correlations between parameters (d).

#### Data set: AVESA_SE_weight

Table 2: Summary of parameter estimates for AVESA_SE_weight data set

| Parameter | median | Q2.5 | Q97.5 |
|---|---|---|---|
| b | 0.358 | 0.077 | 0.771 |
| d | 0.876 | 0.740 | 1.017 |
| e | 4.728 | 1.521 | 13.596 |
| sigma | 0.133 | 0.100 | 0.187 |

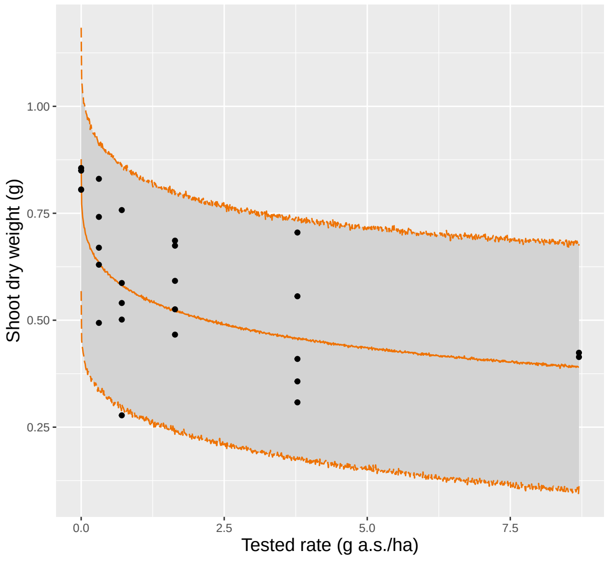

(a) Dose-response curve

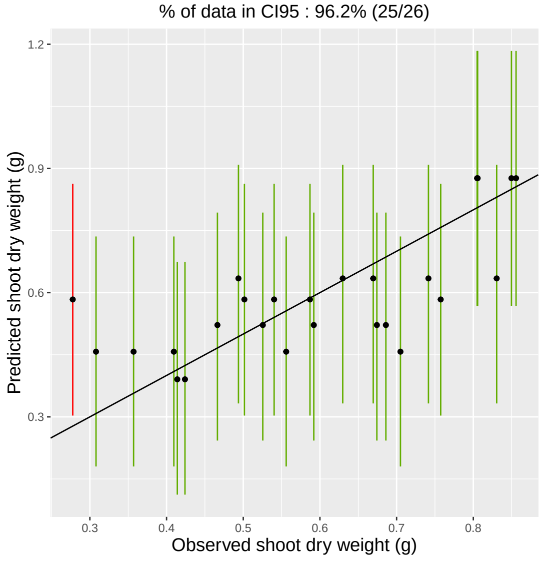

(b) Posterior predictive check (PPC)

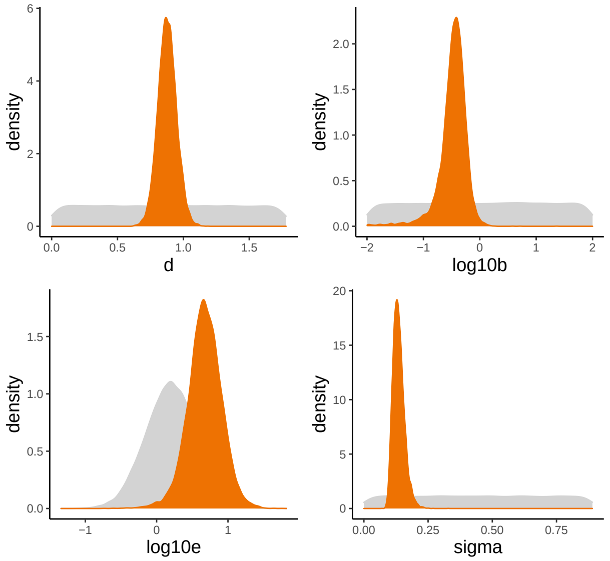

(c) Priors and posteriors

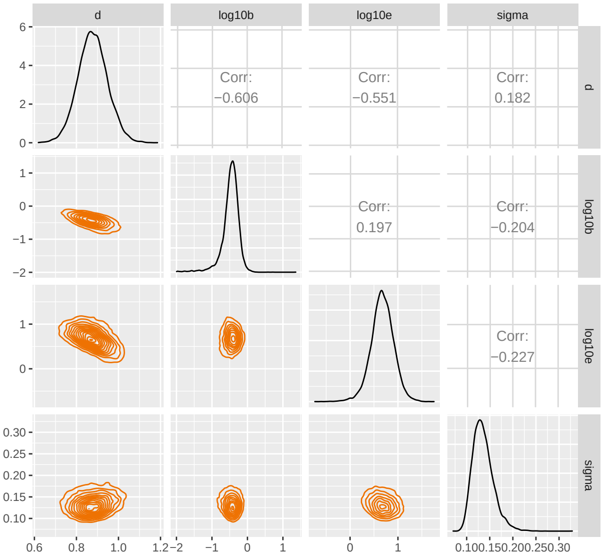

(d) Correlations between parameters

Figure 2: Dose-response curve (a), PPC (b), prior and posterior distributions (c) and correlations between parameters (d).

#### Data set: BEAVA_SE_weight

Table 3: Summary of parameter estimates for BEAVA_SE_weight data set

| Parameter | median | Q2.5 | Q97.5 |
|---|---|---|---|
| b | 6.949 | 1.175 | 82.756 |
| d | 1.911 | 1.661 | 2.206 |
| e | 1.902 | 1.330 | 3.539 |
| sigma | 0.630 | 0.498 | 0.846 |

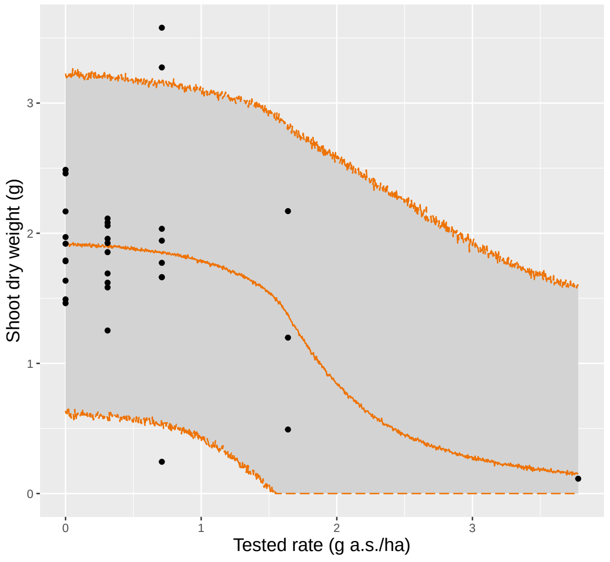

(a) Dose-response curve

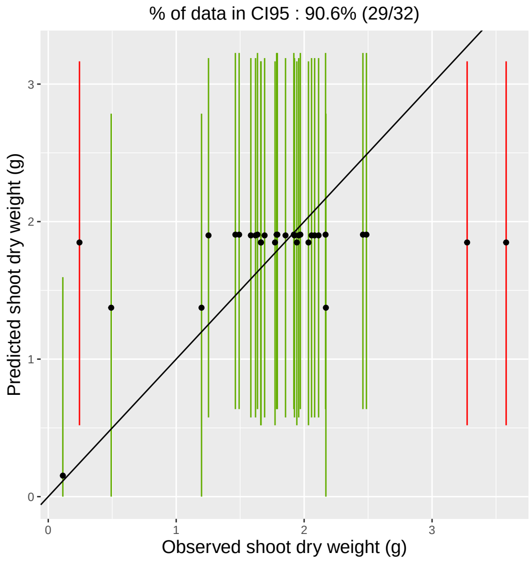

(b) Posterior predictive check (PPC)

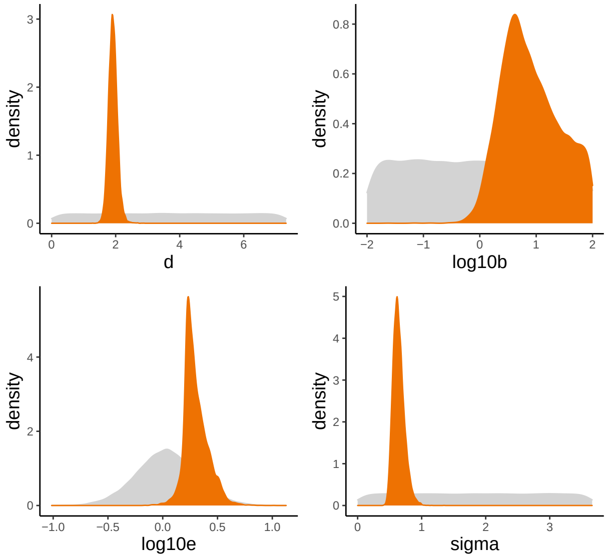

(c) Priors and posteriors

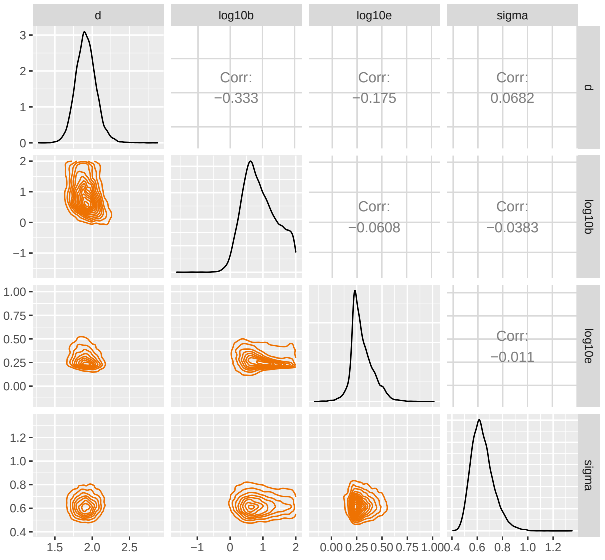

(d) Correlations between parameters

Figure 3: Dose-response curve (a), PPC (b), prior and posterior distributions (c) and correlations between parameters (d).

#### Data set: BRSNW_SE_weight

Table 4: Summary of parameter estimates for BRSNW_SE_weight data set

| Parameter | median | Q2.5 | Q97.5 |
|---|---|---|---|
| b | 2.821 | 1.431 | 10.526 |
| d | 3.095 | 2.969 | 3.239 |
| e | 1.358 | 0.995 | 1.974 |
| sigma | 0.348 | 0.285 | 0.439 |

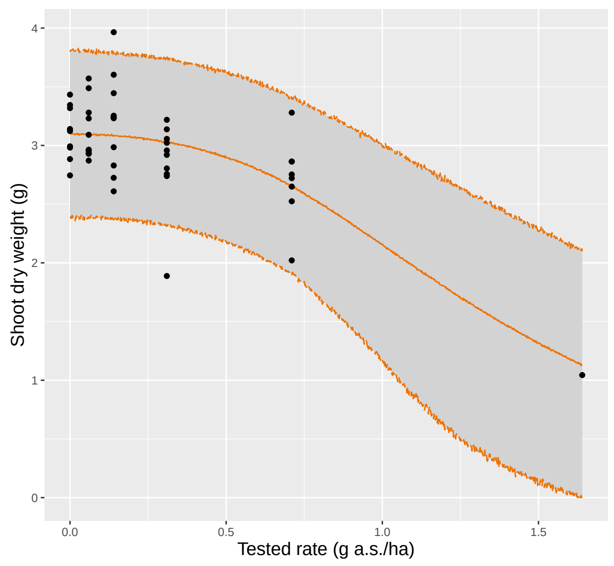

(a) Dose-response curve

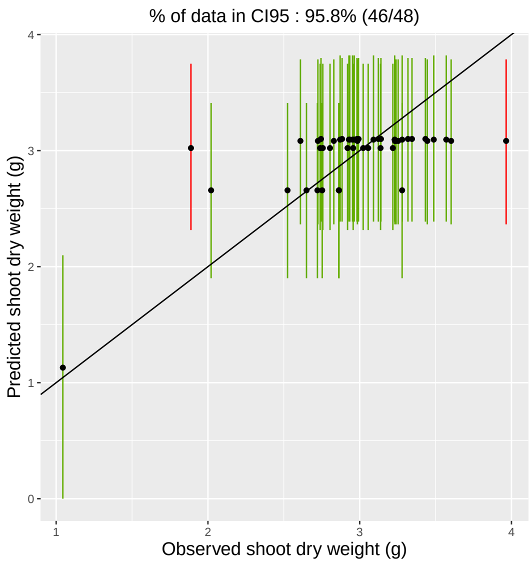

(b) Posterior predictive check (PPC)

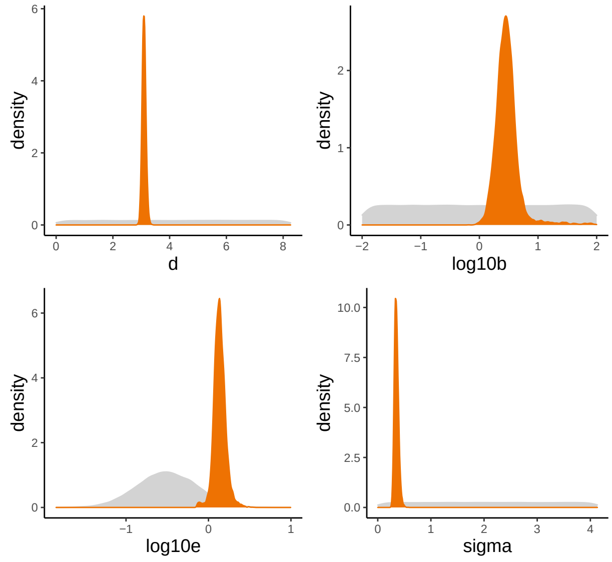

(c) Priors and posteriors

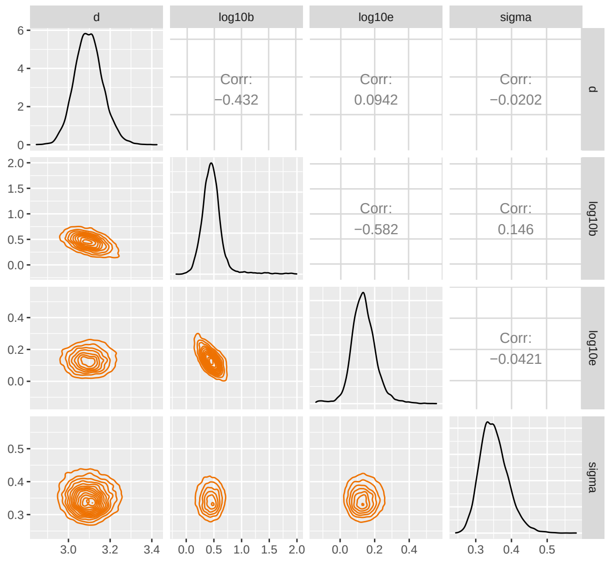

(d) Correlations between parameters

Figure 4: Dose-response curve (a), PPC (b), prior and posterior distributions (c) and correlations between parameters (d).

#### Data set: CUMSA_SE_weight

Table 5: Summary of parameter estimates for CUMSA_SE_weight data set

| Parameter | median | Q2.5 | Q97.5 |
|---|---|---|---|
| b | 5.662 | 0.738 | 85.792 |
| d | 4.654 | 4.005 | 5.643 |
| e | 3.346 | 1.745 | 7.123 |
| sigma | 1.761 | 1.417 | 2.274 |

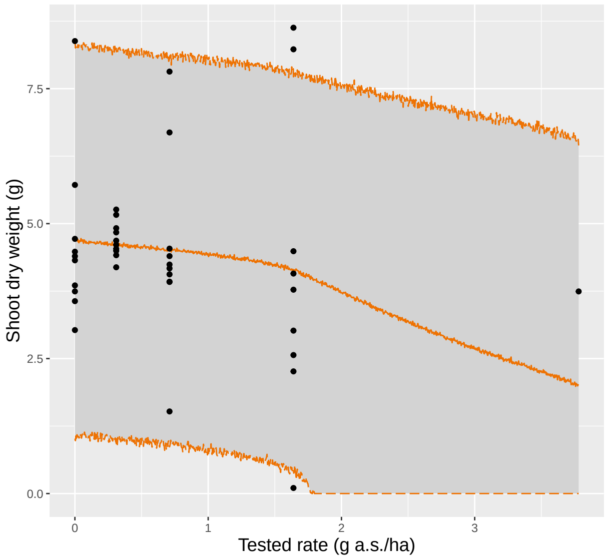

(a) Dose-response curve

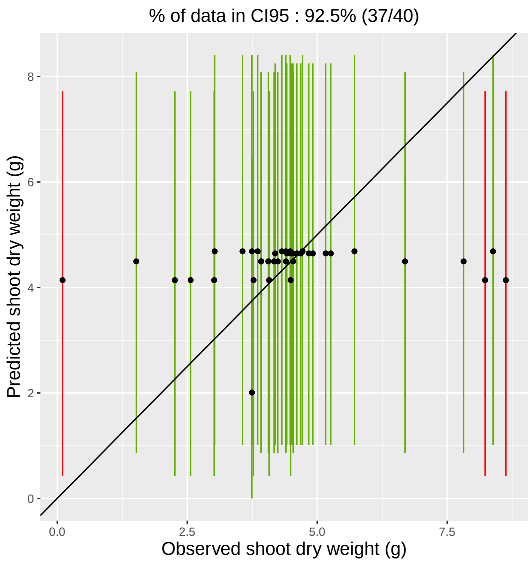

(b) Posterior predictive check (PPC)

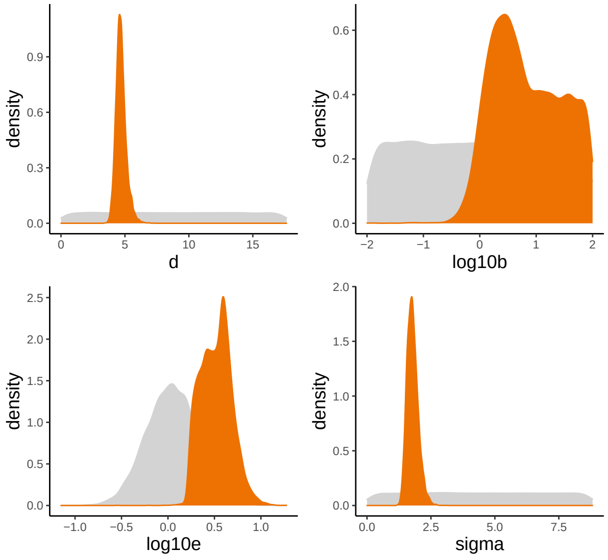

(c) Priors and posteriors

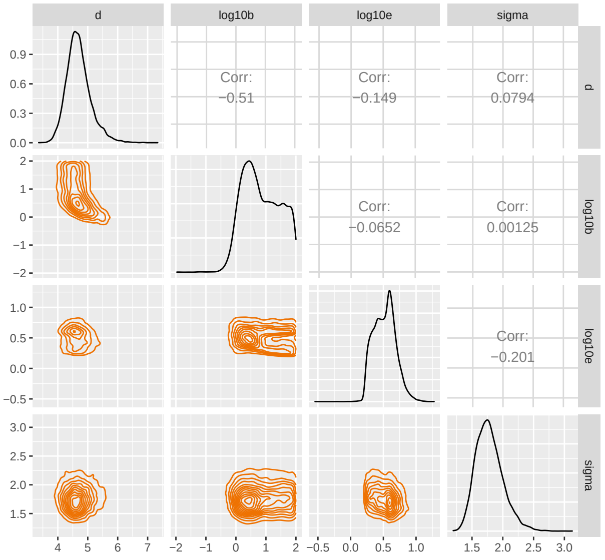

(d) Correlations between parameters

Figure 5: Dose-response curve (a), PPC (b), prior and posterior distributions (c) and correlations between parameters (d).

#### Data set: GLXMA_SE_weight

Table 6: Summary of parameter estimates for GLXMA_SE_weight data set

| Parameter | median | Q2.5 | Q97.5 |
|---|---|---|---|
| b | 0.648 | 0.346 | 1.307 |
| d | 1.553 | 1.357 | 1.763 |
| e | 7.241 | 3.714 | 17.509 |
| sigma | 0.311 | 0.257 | 0.389 |

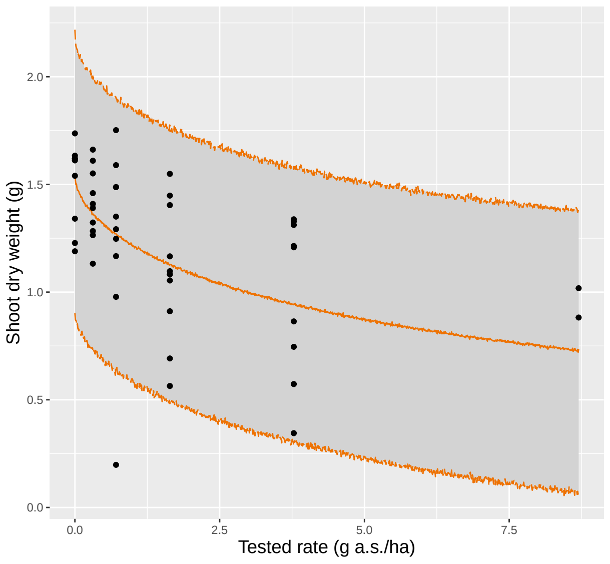

(a) Dose-response curve

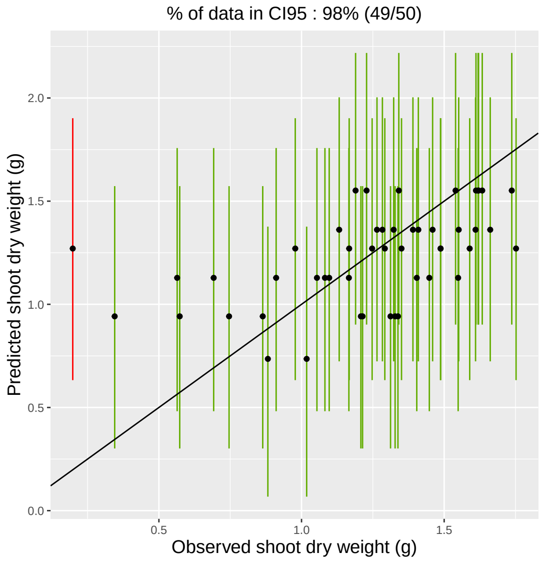

(b) Posterior predictive check (PPC)

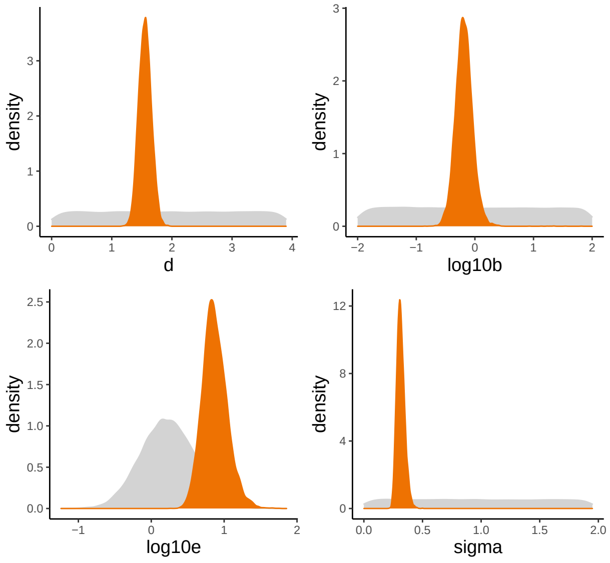

(c) Priors and posteriors

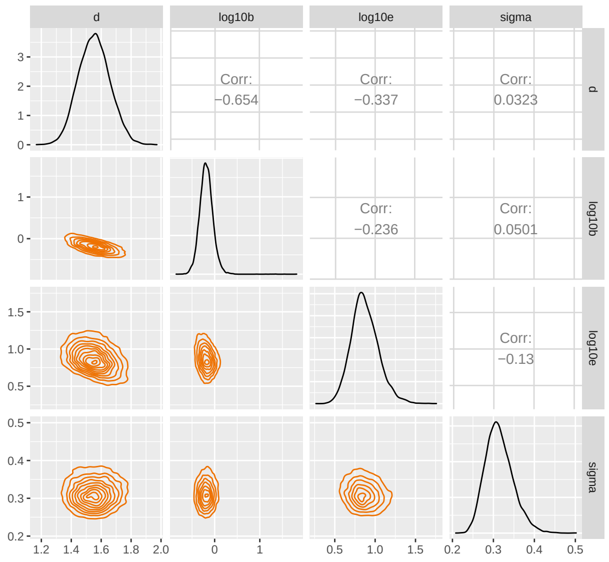

(d) Correlations between parameters

Figure 6: Dose-response curve (a), PPC (b), prior and posterior distributions (c) and correlations between parameters (d).

#### Data set: HELAN_SE_weight

Table 7: Summary of parameter estimates for HELAN_SE_weight data set

| Parameter | median | Q2.5 | Q97.5 |
|---|---|---|---|
| b | 0.901 | 0.592 | 1.408 |
| d | 0.892 | 0.820 | 0.972 |
| e | 14.277 | 9.475 | 24.435 |
| sigma | 0.135 | 0.111 | 0.169 |

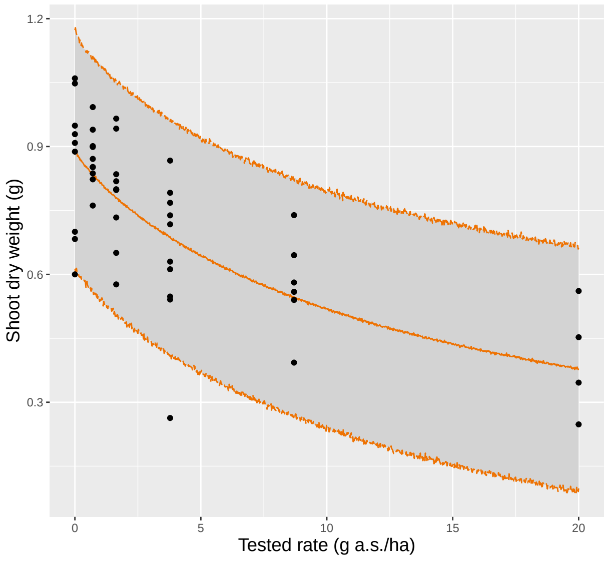

(a) Dose-response curve

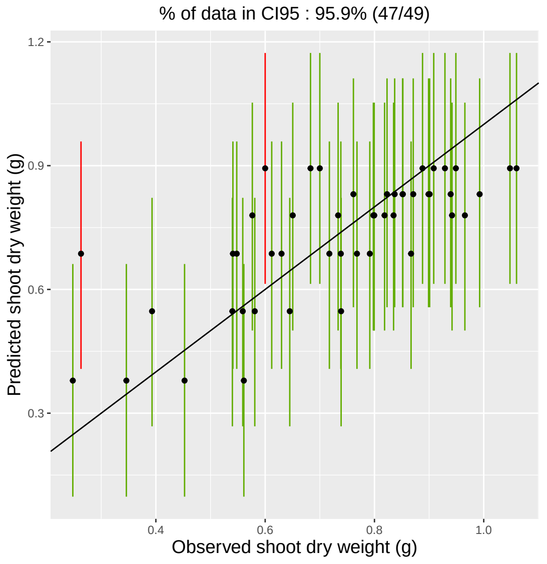

(b) Posterior predictive check (PPC)

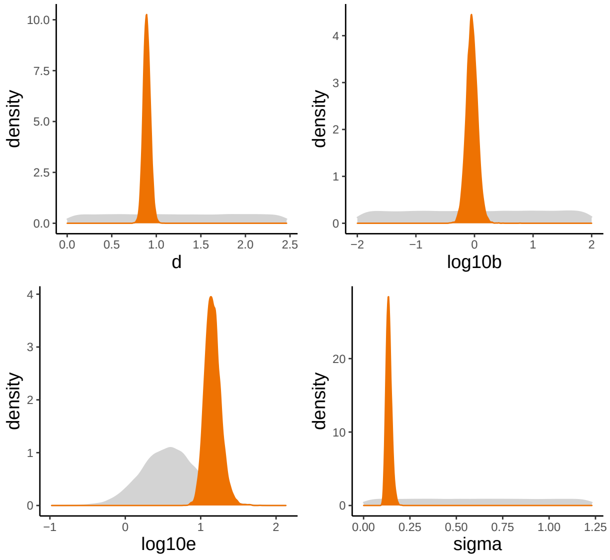

(c) Priors and posteriors

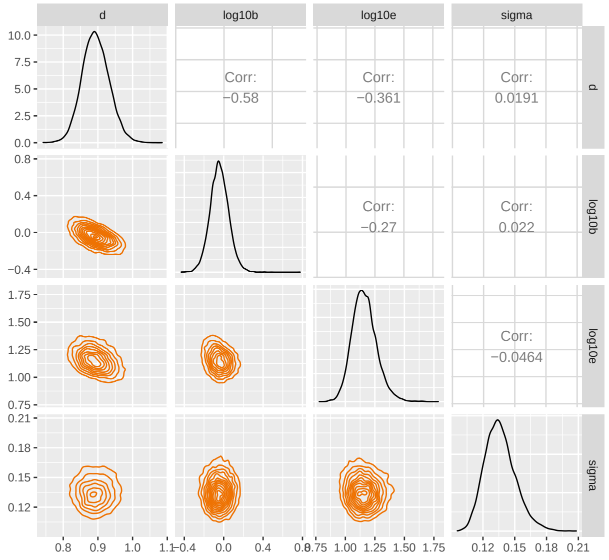

(d) Correlations between parameters

Figure 7: Dose-response curve (a), PPC (b), prior and posterior distributions (c) and correlations between parameters (d).

### Data set: LOLPE_SE_weight

Table 8: Summary of parameter estimates for LOLPE_SE_weight data set

| Parameter | median | Q2.5 | Q97.5 |
|---|---|---|---|
| b | 1.744 | 0.683 | 16.338 |
| d | 0.155 | 0.130 | 0.183 |
| e | 0.330 | 0.210 | 0.608 |
| sigma | 0.034 | 0.024 | 0.052 |

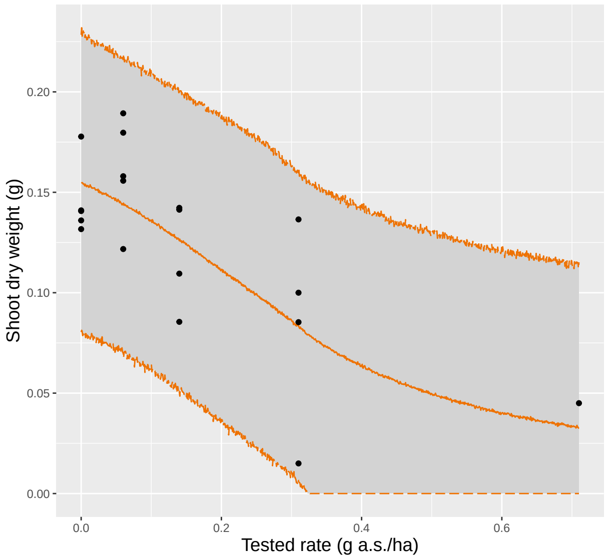

(a) Dose-response curve

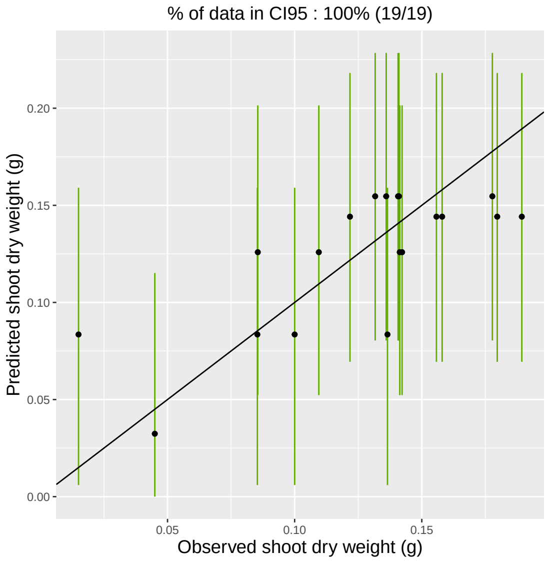

(b) Posterior predictive check (PPC)

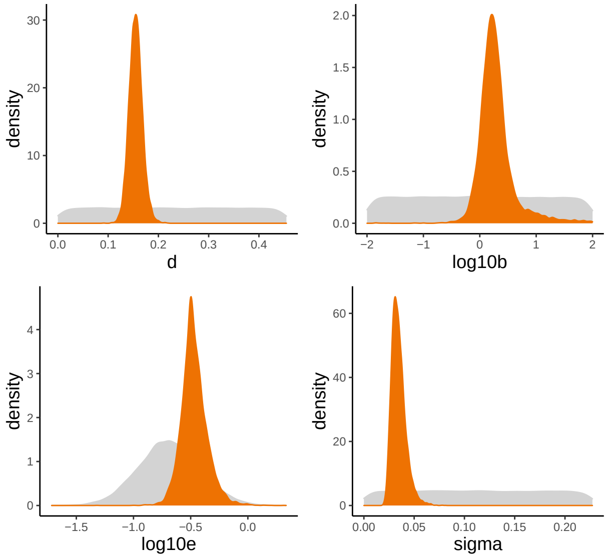

(c) Priors and posteriors

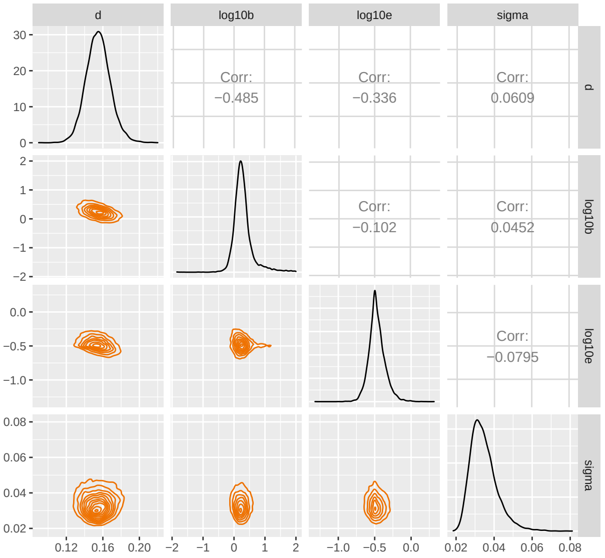

(d) Correlations between parameters

Figure 8: Dose-response curve (a), PPC (b), prior and posterior distributions (c) and correlations between parameters (d).

#### Data set: LYPES_SE_weight

Table 9: Summary of parameter estimates for LYPES_SE_weight data set

| Parameter | median | Q2.5 | Q97.5 |
|---|---|---|---|
| b | 2.810 | 0.934 | 60.826 |
| d | 1.648 | 1.499 | 1.844 |
| e | 1.525 | 0.832 | 2.727 |
| sigma | 0.345 | 0.272 | 0.460 |

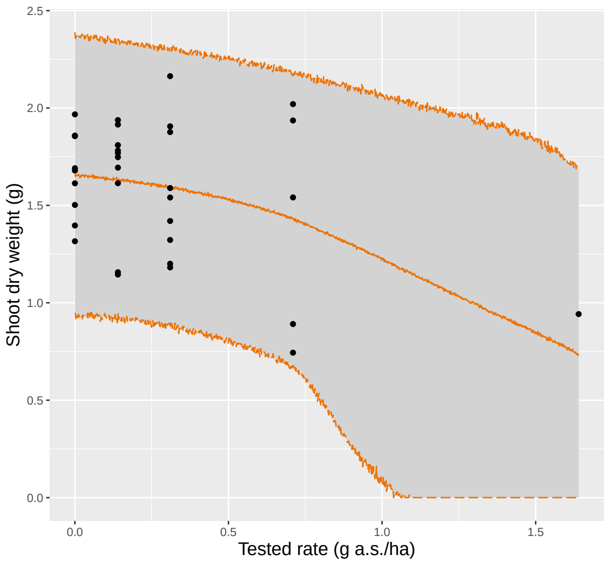

(a) Dose-response curve

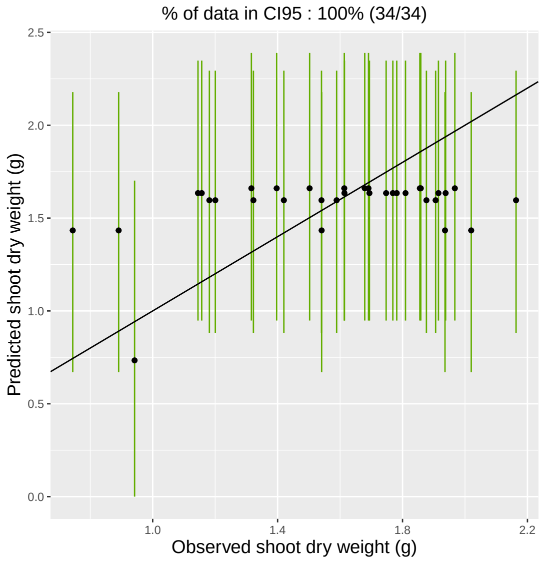

(b) Posterior predictive check (PPC)

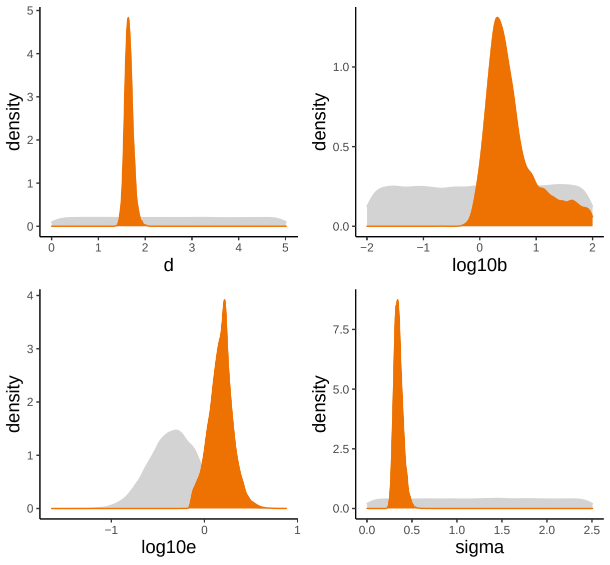

(c) Priors and posteriors

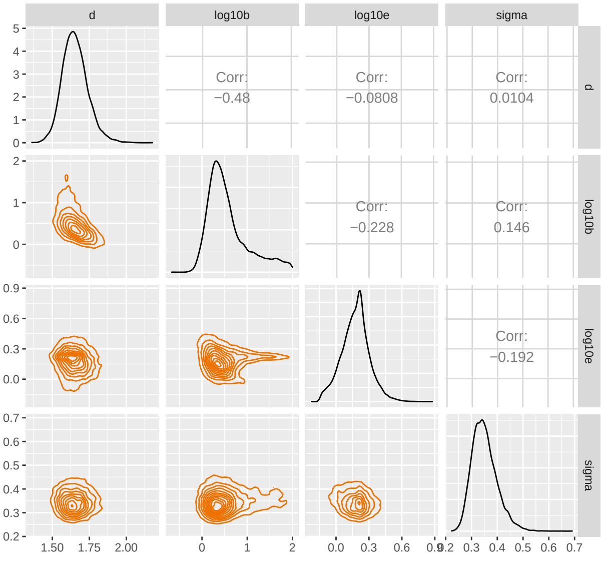

(d) Correlations between parameters

Figure 9: Dose-response curve (a), PPC (b), prior and posterior distributions (c) and correlations between parameters (d).

#### Data set: ZEAMA_SE_weight

Table 10: Summary of parameter estimates for ZEAMA_SE_weight data set

| Parameter | median | Q2.5 | Q97.5 |
|---|---|---|---|
| b | 0.850 | 0.494 | 1.807 |
| d | 4.969 | 4.531 | 5.471 |
| e | 25.897 | 18.105 | 43.959 |
| sigma | 0.728 | 0.610 | 0.888 |

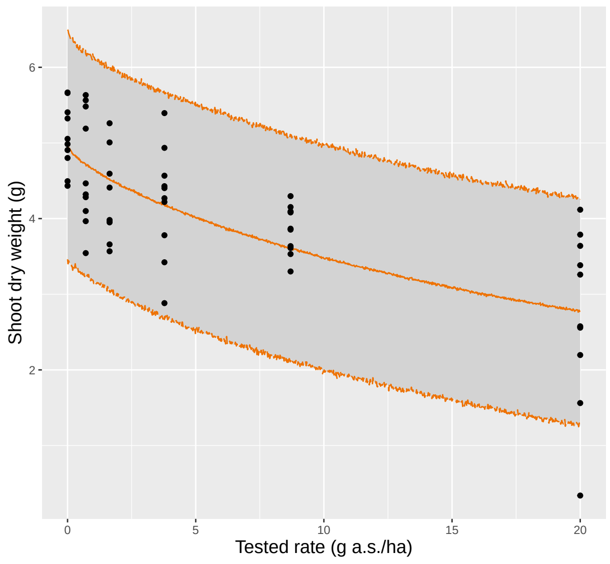

(a) Dose-response curve

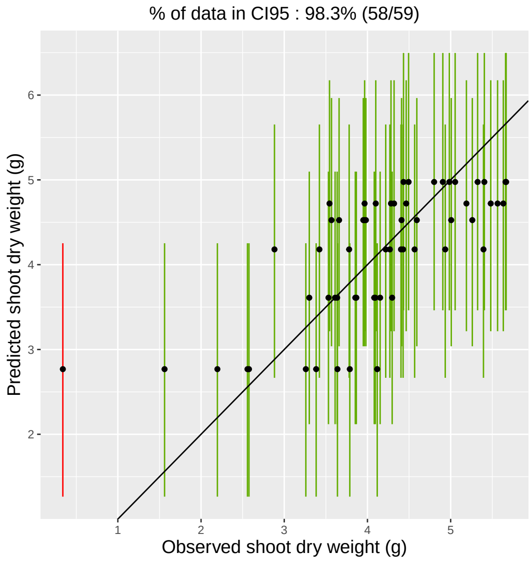

(b) Posterior predictive check (PPC)

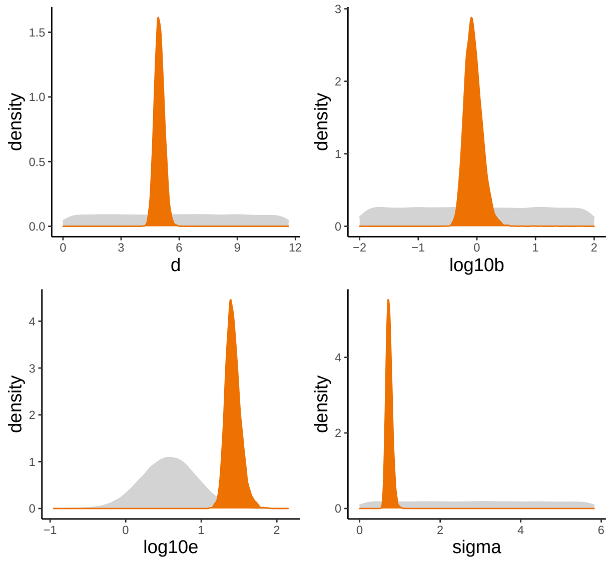

(c) Priors and posteriors

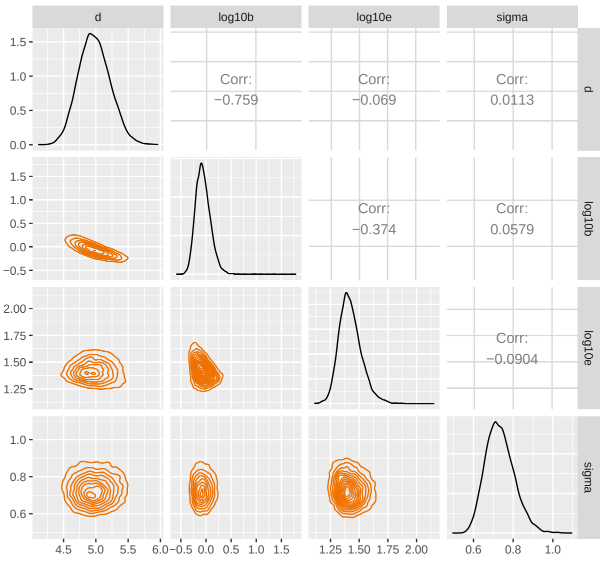

(d) Correlations between parameters

Figure 10: Dose-response curve (a), PPC (b), prior and posterior distributions (c) and correlations between parameters (d).
