## Supplementary material for "How to account for the uncertainty from standard toxicity tests in species sensitivity distributions: an example in non-target plants": Archive with reports on studies: report_VV_survival.pdf

### Dose-response analyses

#### Study 5

#### Vegetative Vigour test - survival endpoint

25 June 2020

Contact:

---

This is a report which provides results on all performed dose-response analyses for the survival endpoint of the Vegetative Vigour test for study 5.

---

### Data set: ALLCE_VV_survival

Table 1: Summary of parameter estimates (parameter d is set to 1) for ALLCE_VV_survival data set

| Parameter | median | Q2.5 | Q97.5 |
|---|---|---|---|
| b | 33.913 | 4.904 | 94.971 |
| e | 29.127 | 21.490 | 66.918 |

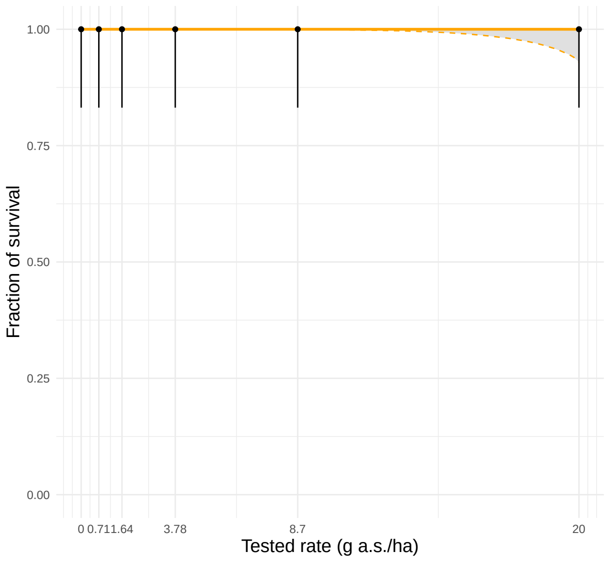

(a) Dose-response curve

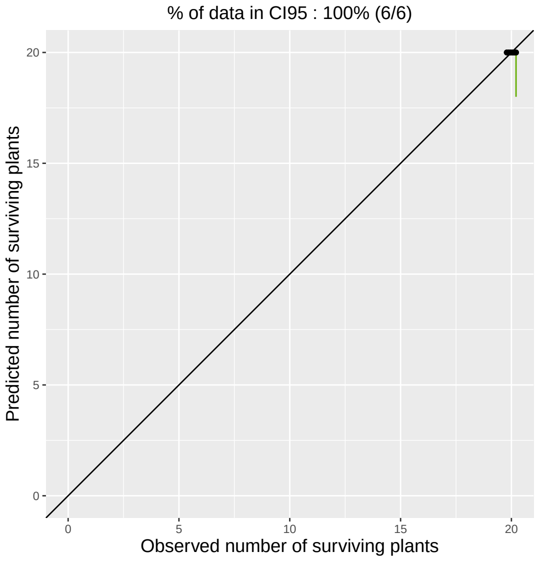

(b) Posterior predictive check (PPC)

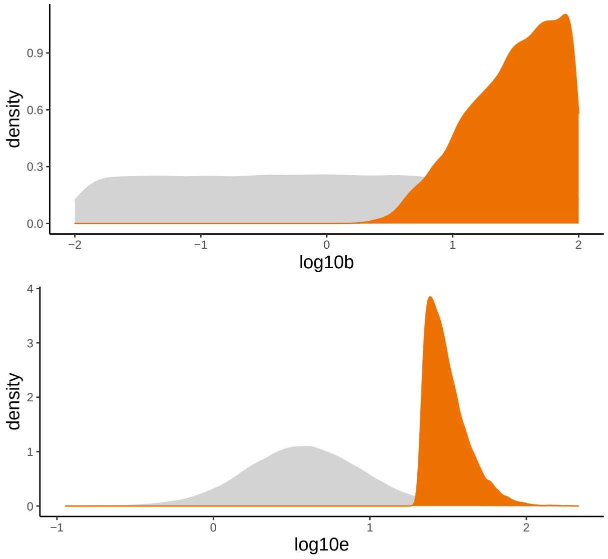

(c) Priors and posteriors

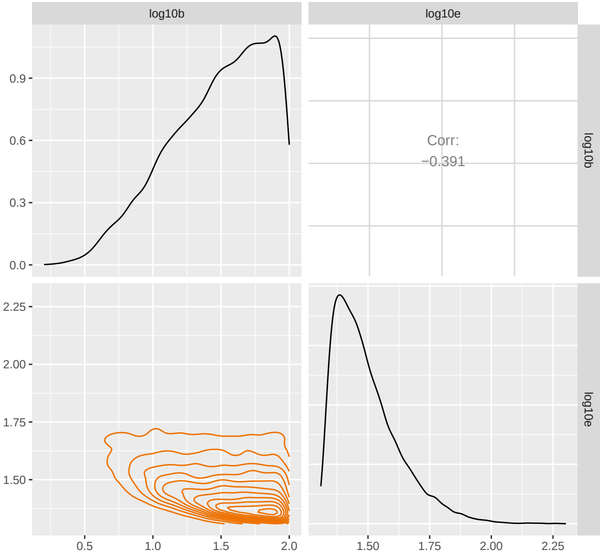

(d) Correlations between parameters

Figure 1: Dose-response curve (a), PPC (b), prior and posterior distributions (c) and correlations between parameters (d).

#### Data set: AVESA_VV_survival

Table 2: Summary of parameter estimates (parameter d is set to 1) for AVESA_VV_survival data set

| Parameter | median | Q2.5 | Q97.5 |
|---|---|---|---|
| b | 33.028 | 4.779 | 95.130 |
| e | 29.485 | 21.571 | 68.822 |

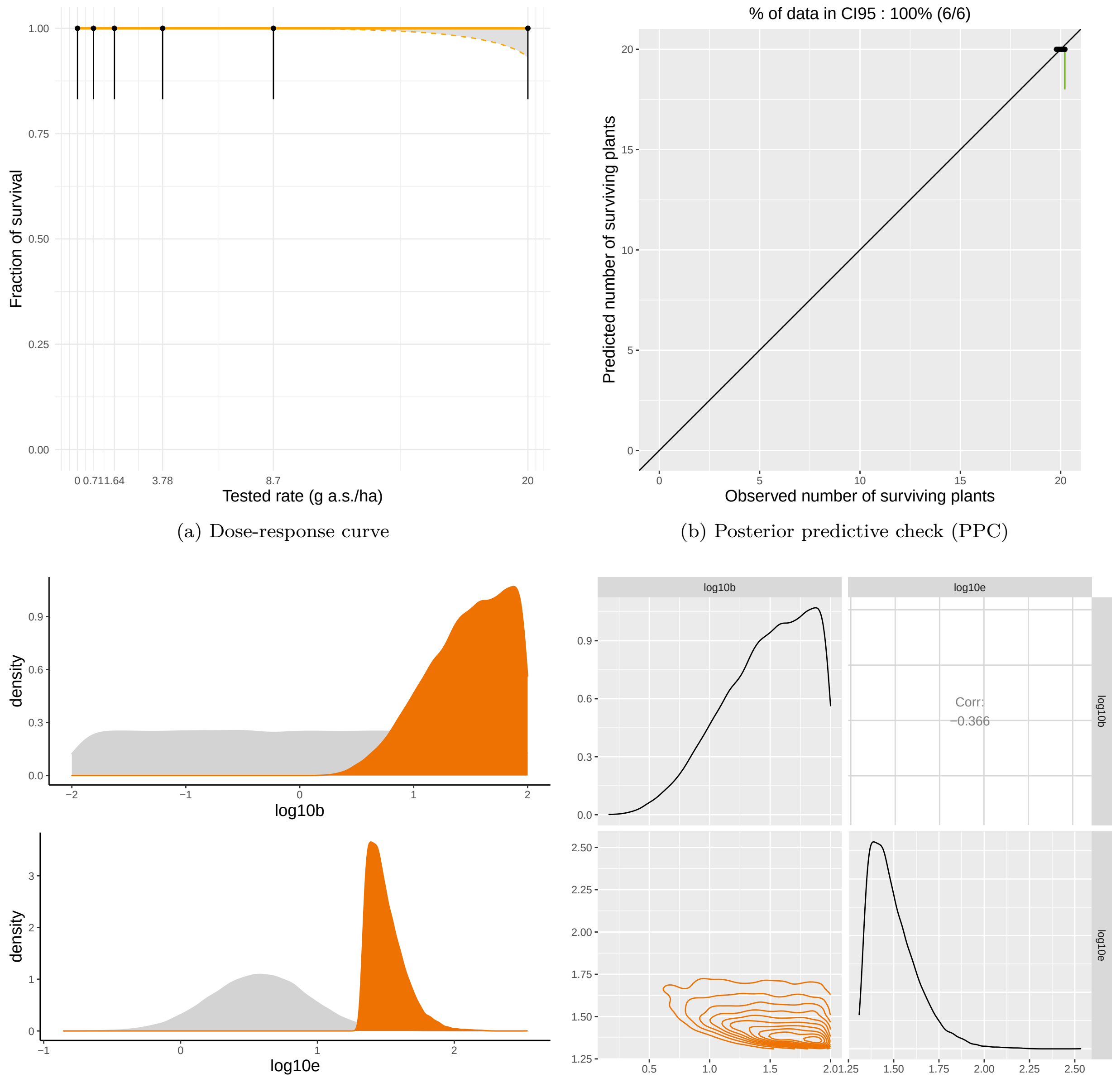

(a) Dose-response curve

(b) Posterior predictive check (PPC)

(c) Priors and posteriors

(d) Correlations between parameters

Figure 2: Dose-response curve (a), PPC (b), prior and posterior distributions (c) and correlations between parameters (d).

#### Data set: BEAVA_VV_survival

Table 3: Summary of parameter estimates (parameter d is set to 1) for BEAVA_VV_survival data set

| Parameter | median | Q2.5 | Q97.5 |
|---|---|---|---|
| b | 3.150 | 1.577 | 6.281 |
| e | 22.072 | 17.304 | 34.066 |

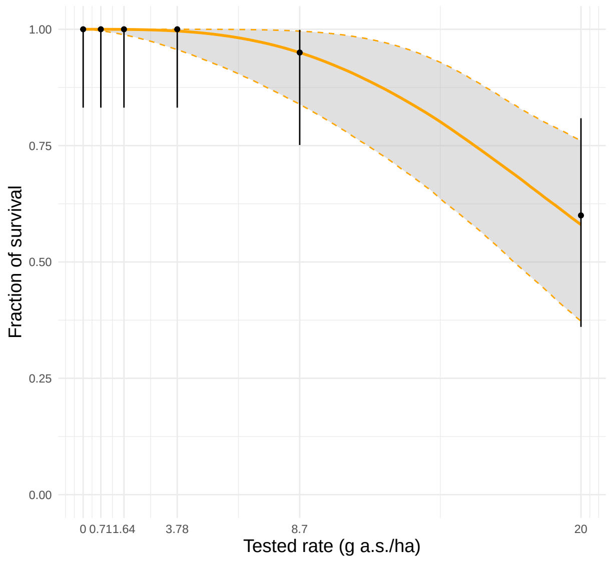

(a) Dose-response curve

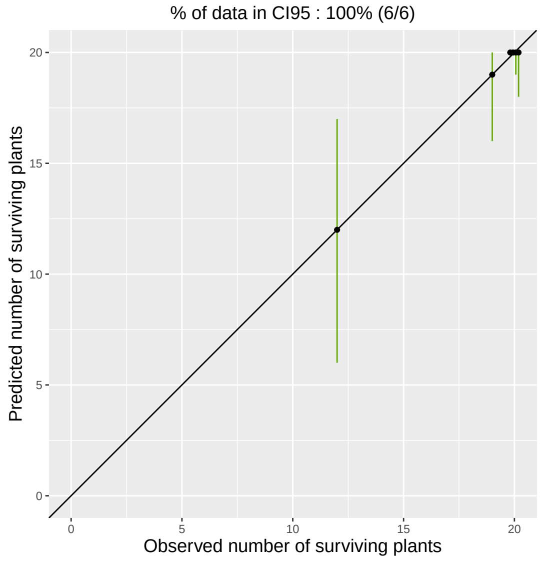

(b) Posterior predictive check (PPC)

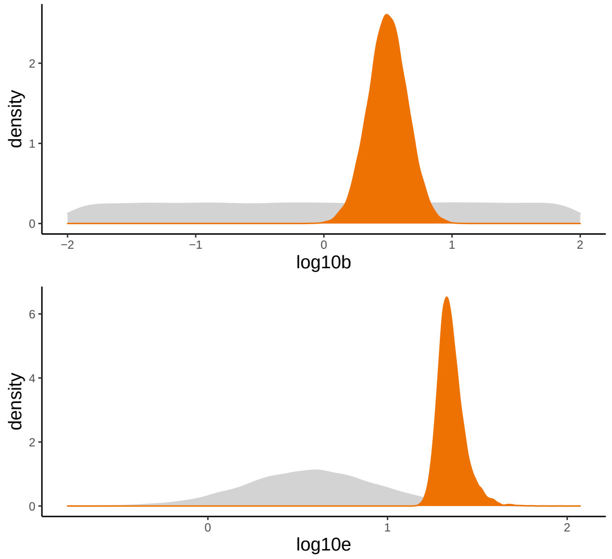

(c) Priors and posteriors

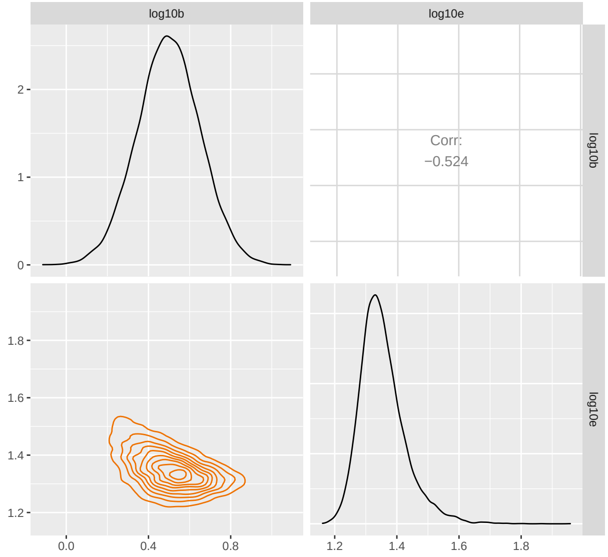

(d) Correlations between parameters

Figure 3: Dose-response curve (a), PPC (b), prior and posterior distributions (c) and correlations between parameters (d).

### Data set: BRSNW_VV_survival

Table 4: Summary of parameter estimates (parameter d is set to 1) for BRSNW_VV_survival data set

| Parameter | median | Q2.5 | Q97.5 |
|---|---|---|---|
| b | 7.556 | 2.192 | 65.037 |
| e | 22.671 | 20.165 | 36.420 |

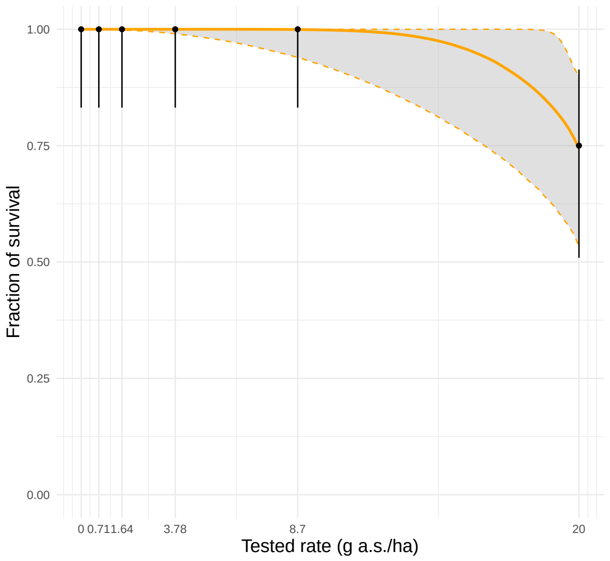

(a) Dose-response curve

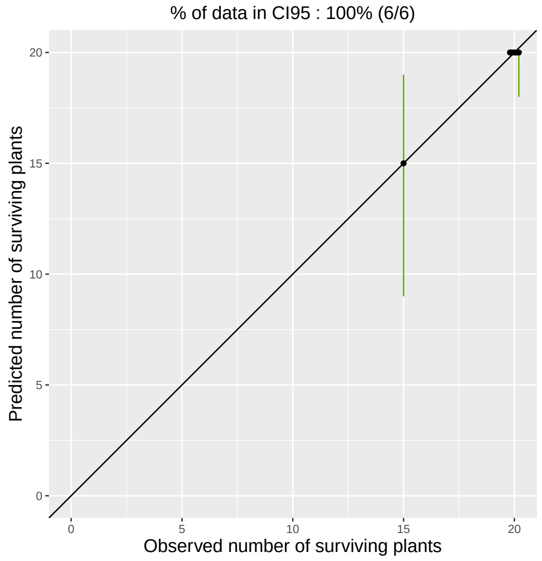

(b) Posterior predictive check (PPC)

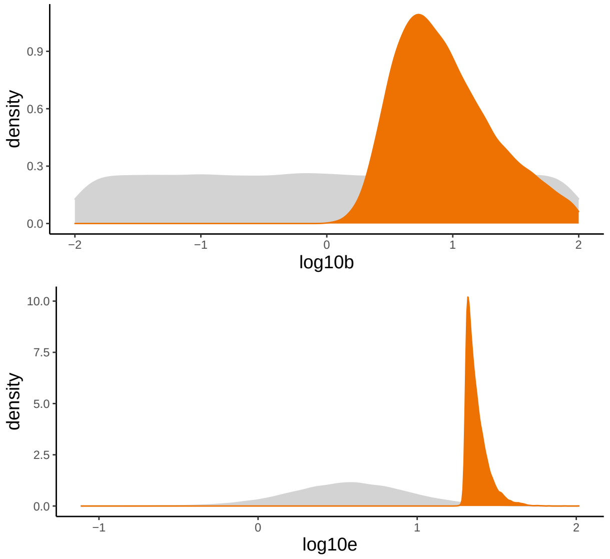

(c) Priors and posteriors

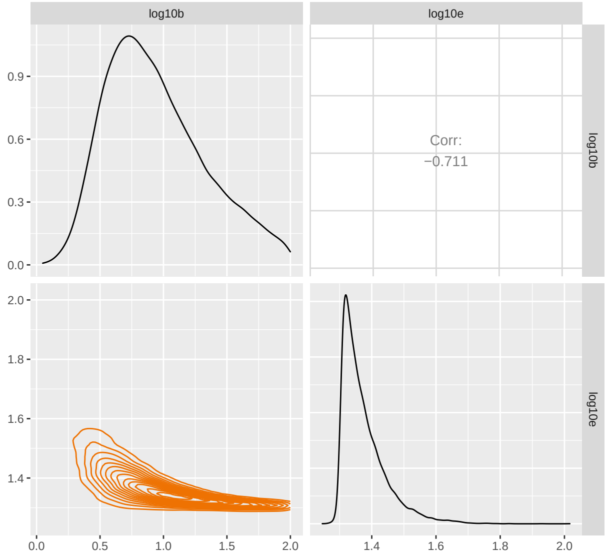

(d) Correlations between parameters

Figure 4: Dose-response curve (a), PPC (b), prior and posterior distributions (c) and correlations between parameters (d).

#### Data set: CUMSA_VV_survival

Table 5: Summary of parameter estimates (parameter d is set to 1) for CUMSA_VV_survival data set

| Parameter | median | Q2.5 | Q97.5 |
|---|---|---|---|
| b | 11.849 | 2.720 | 76.537 |
| e | 25.517 | 20.759 | 54.407 |

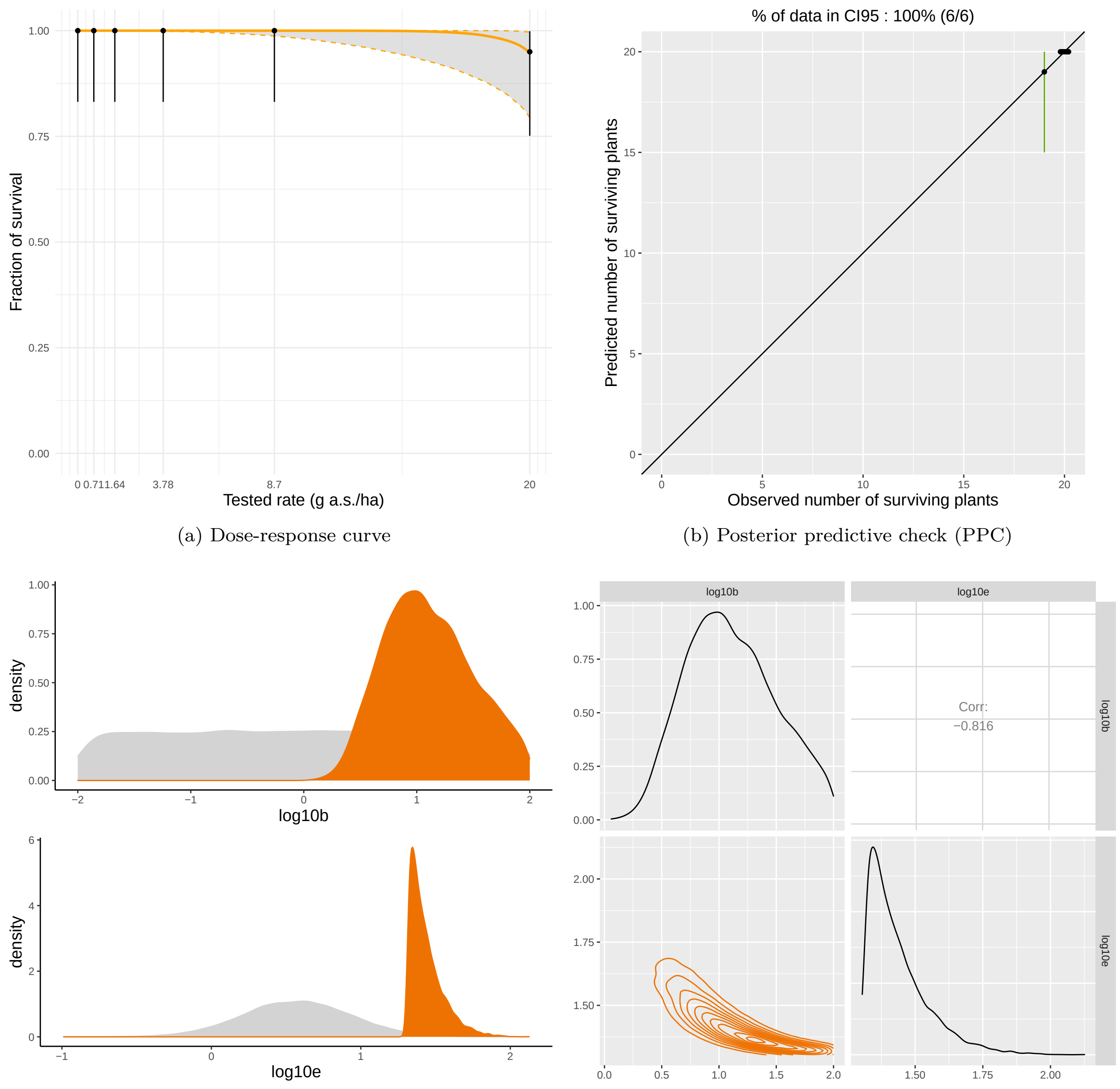

(a) Dose-response curve

(b) Posterior predictive check (PPC)

(c) Priors and posteriors

(d) Correlations between parameters

Figure 5: Dose-response curve (a), PPC (b), prior and posterior distributions (c) and correlations between parameters (d).

#### Data set: GLXMA_VV_survival

Table 6: Summary of parameter estimates (parameter d is set to 1) for GLXMA_VV_survival data set

| Parameter | median | Q2.5 | Q97.5 |
|---|---|---|---|
| b | 34.050 | 5.036 | 95.499 |
| e | 29.272 | 21.464 | 67.409 |

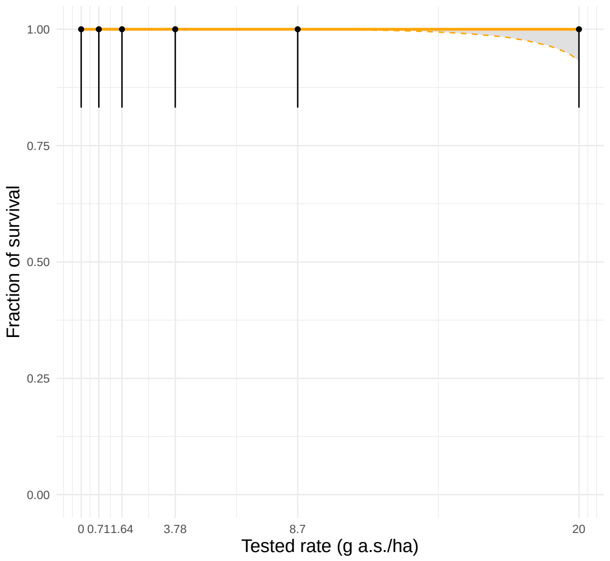

(a) Dose-response curve

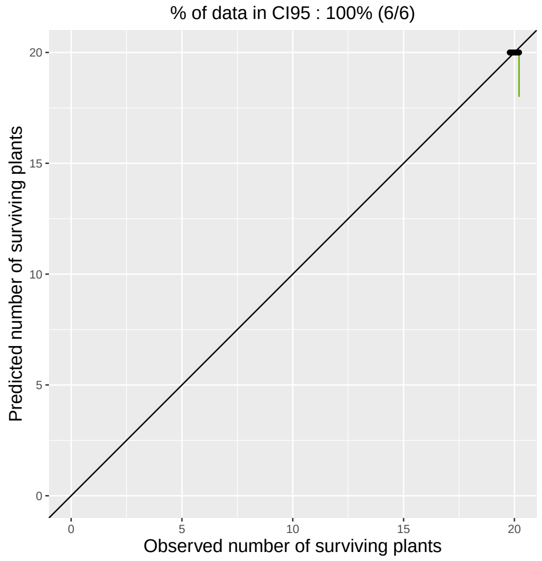

(b) Posterior predictive check (PPC)

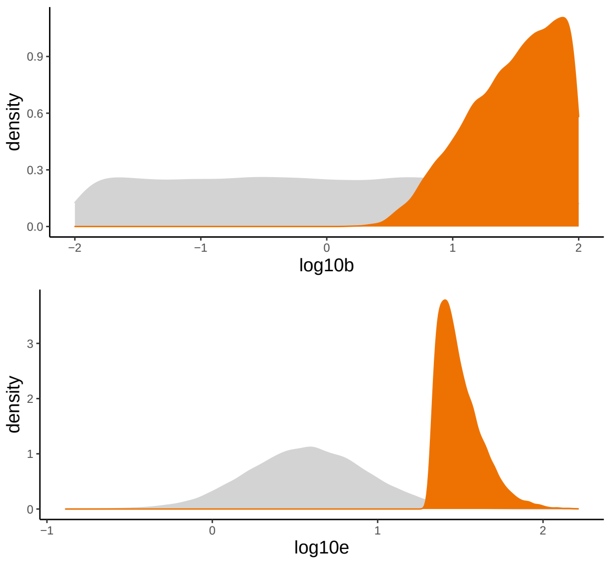

(c) Priors and posteriors

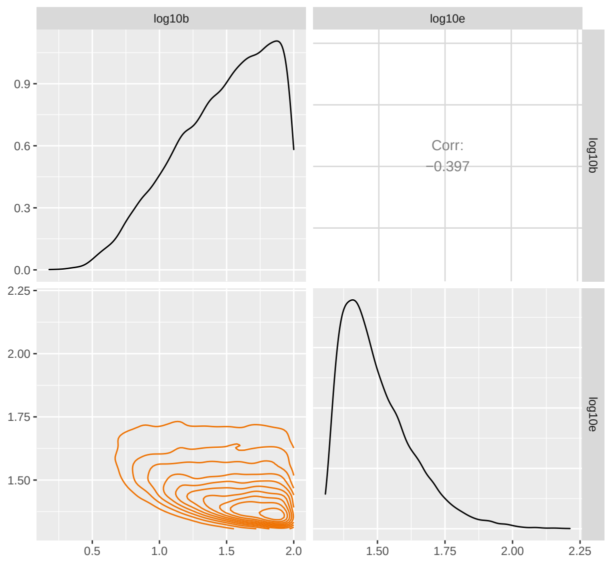

(d) Correlations between parameters

Figure 6: Dose-response curve (a), PPC (b), prior and posterior distributions (c) and correlations between parameters (d).

### Data set: HELAN_VV_survival

Table 7: Summary of parameter estimates (parameter d is set to 1) for HELAN_VV_survival data set

| Parameter | median | Q2.5 | Q97.5 |
|---|---|---|---|
| b | 34.230 | 5.033 | 95.094 |
| e | 29.006 | 21.457 | 68.555 |

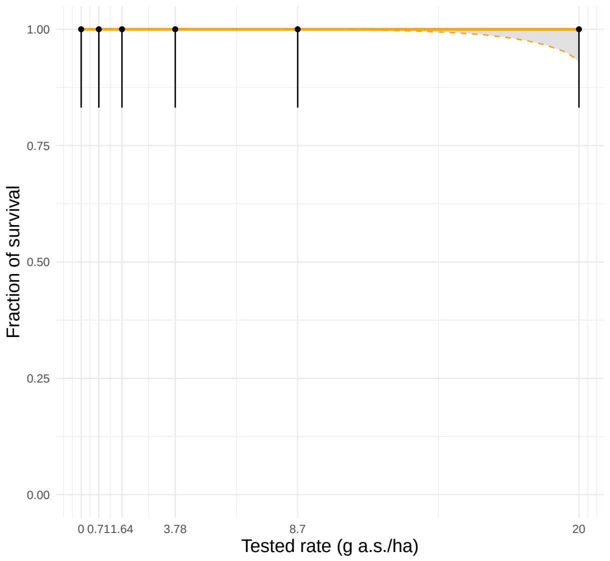

(a) Dose-response curve

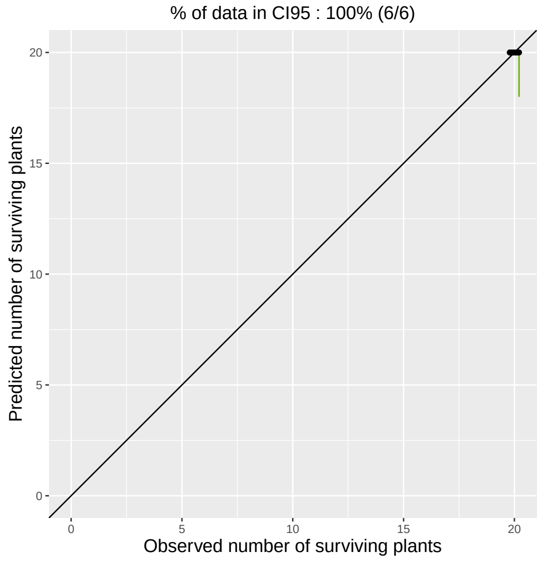

(b) Posterior predictive check (PPC)

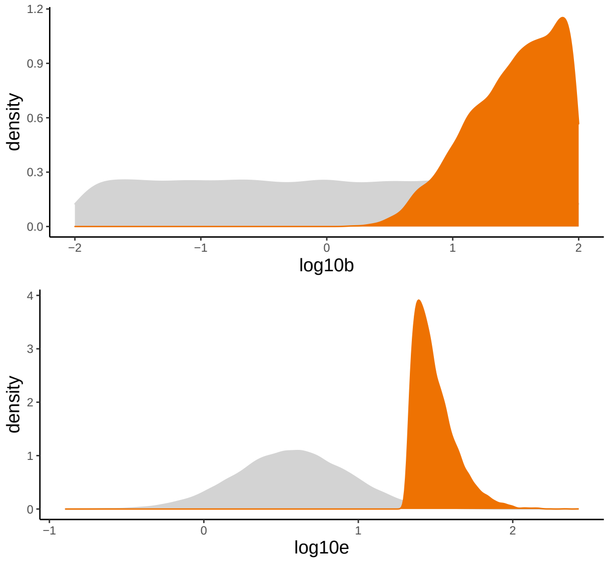

(c) Priors and posteriors

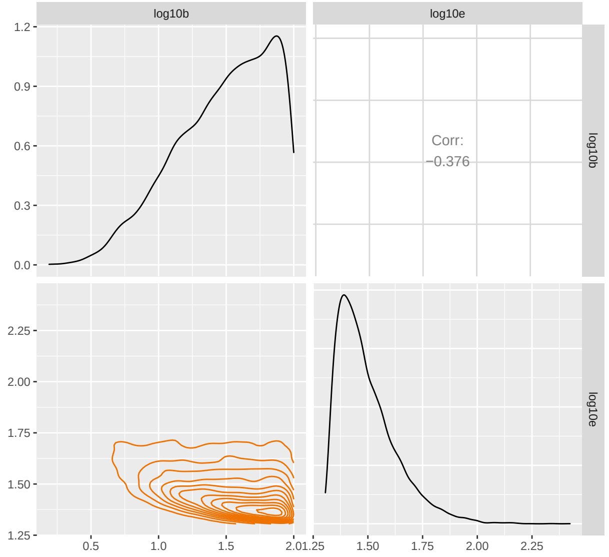

(d) Correlations between parameters

Figure 7: Dose-response curve (a), PPC (b), prior and posterior distributions (c) and correlations between parameters (d).

#### Data set: LOLPE_VV_survival

Table 8: Summary of parameter estimates (parameter d is set to 1) for LOLPE_VV_survival data set

| Parameter | median | Q2.5 | Q97.5 |
|---|---|---|---|
| b | 1.963 | 1.185 | 3.018 |
| e | 14.709 | 10.789 | 22.272 |

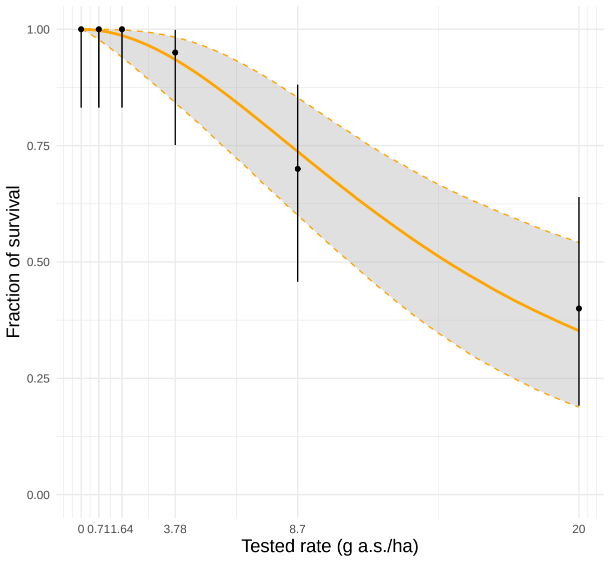

(a) Dose-response curve

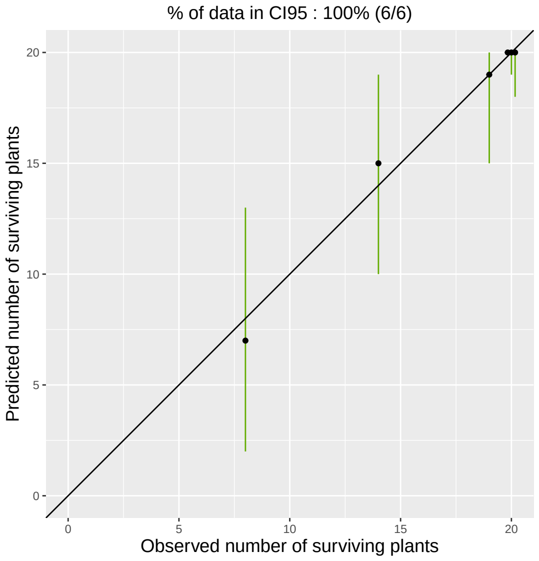

(b) Posterior predictive check (PPC)

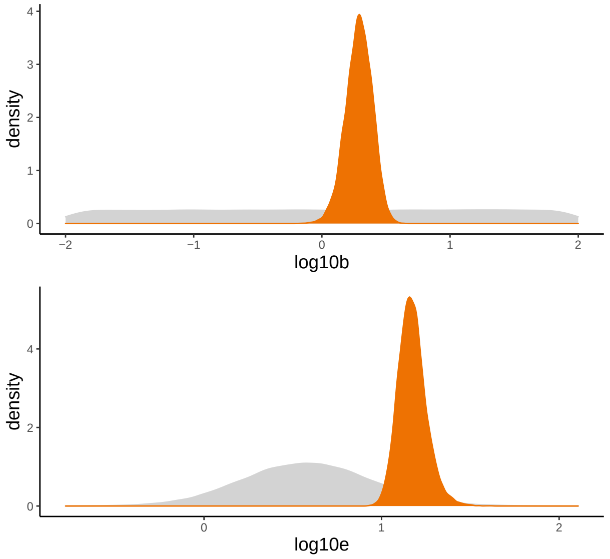

(c) Priors and posteriors

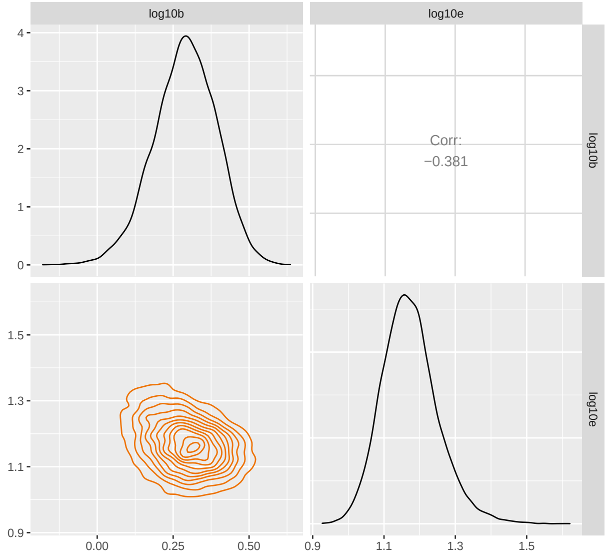

(d) Correlations between parameters

Figure 8: Dose-response curve (a), PPC (b), prior and posterior distributions (c) and correlations between parameters (d).

#### Data set: LYPES_VV_survival

Table 9: Summary of parameter estimates (parameter d is set to 1) for LYPES_VV_survival data set

| Parameter | median | Q2.5 | Q97.5 |
|---|---|---|---|
| b | 11.868 | 2.676 | 78.407 |
| e | 25.537 | 20.744 | 55.327 |

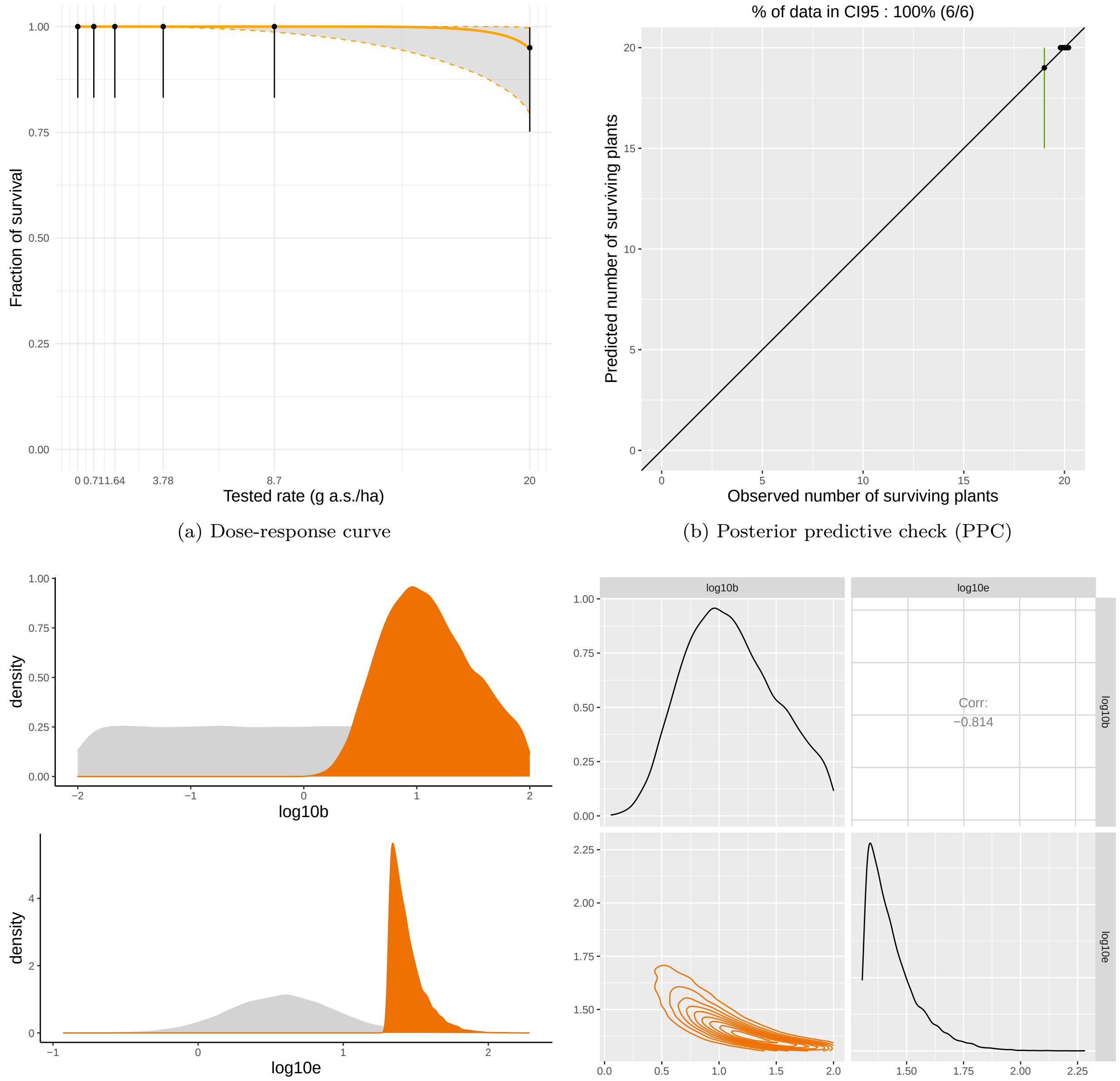

(a) Dose-response curve

(b) Posterior predictive check (PPC)

(c) Priors and posteriors

(d) Correlations between parameters

Figure 9: Dose-response curve (a), PPC (b), prior and posterior distributions (c) and correlations between parameters (d).

#### Data set: ZEAMA_VV_survival

Table 10: Summary of parameter estimates (parameter d is set to 1) for ZEAMA_VV_survival data set

| Parameter | median | Q2.5 | Q97.5 |
|---|---|---|---|
| b | 33.356 | 4.832 | 95.407 |
| e | 29.095 | 21.398 | 67.298 |

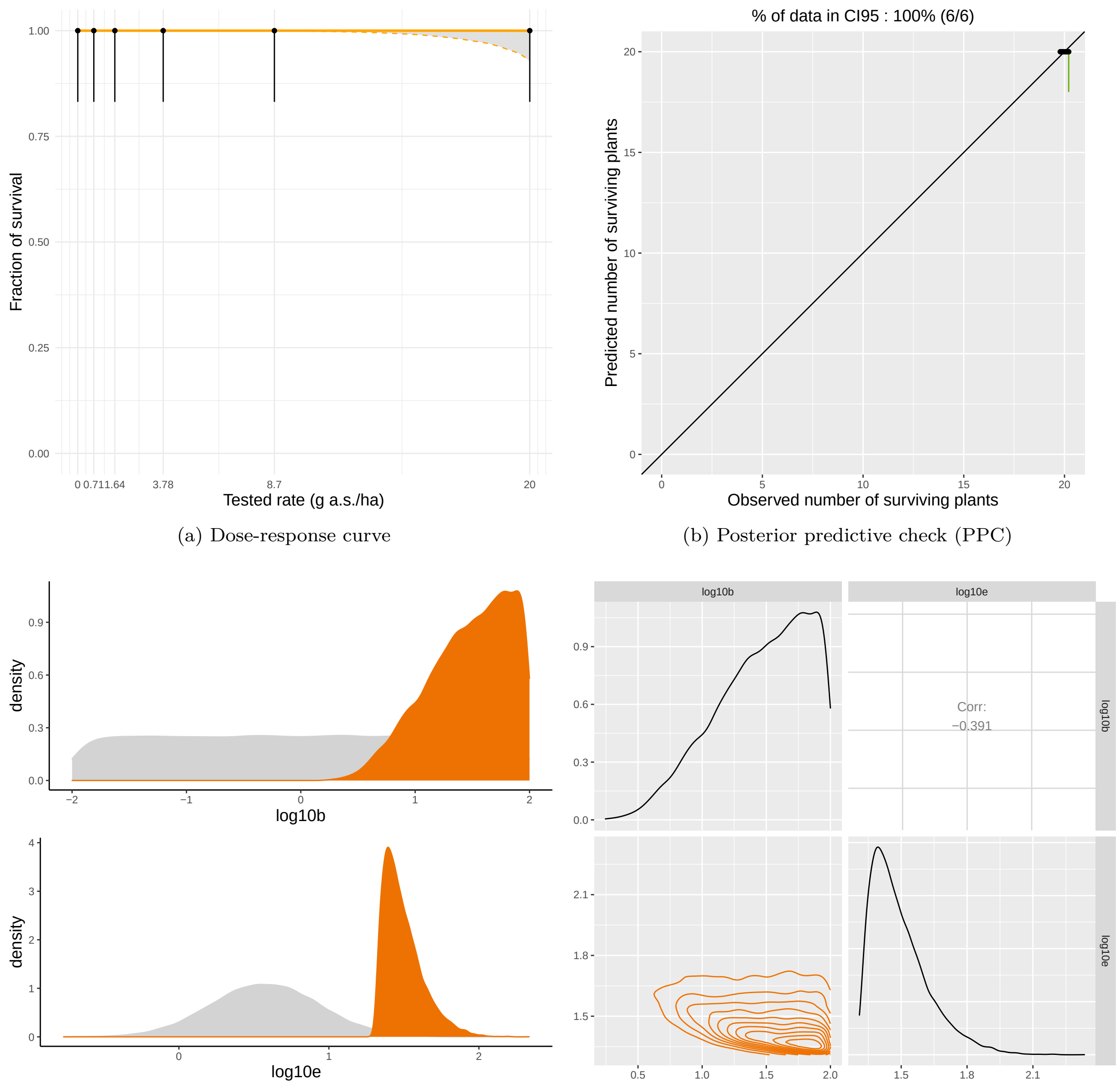

(a) Dose-response curve

(b) Posterior predictive check (PPC)

(c) Priors and posteriors

(d) Correlations between parameters

Figure 10: Dose-response curve (a), PPC (b), prior and posterior distributions (c) and correlations between parameters (d).
