## Supplementary material for "How to account for the uncertainty from standard toxicity tests in species sensitivity distributions: an example in non-target plants": Archive with reports on studies: report_SE_emergence.pdf

### Dose-response analyses

Study 6

Seedling Emergence test - emergence endpoint

25 June 2020

Contact:

---

This is a report which provides results on all performed dose-response analyses for the emergence endpoint of the Seedling Emergence test for study 6.

---

### Data set: ALLCE_SE_emergence

Table 1: Summary of parameter estimates for ALLCE_SE_emergence data set

| Parameter | median | Q2.5 | Q97.5 |
|---|---|---|---|
| b | 1.800 | 0.581 | 55.708 |
| d | 0.751 | 0.630 | 0.877 |
| e | 165.846 | 93.410 | 344.669 |

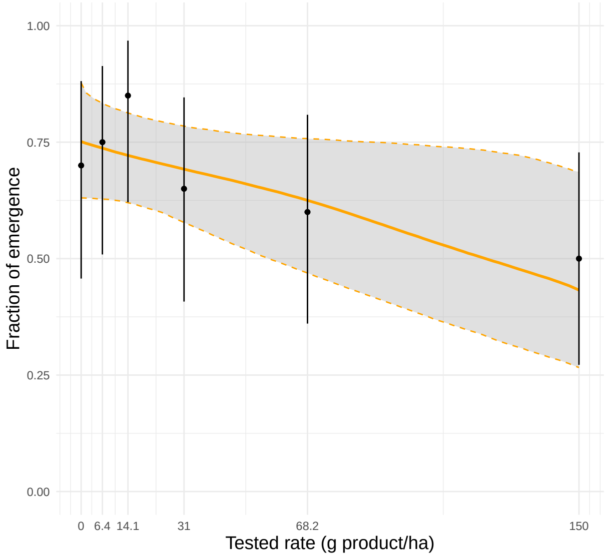

(a) Dose-response curve

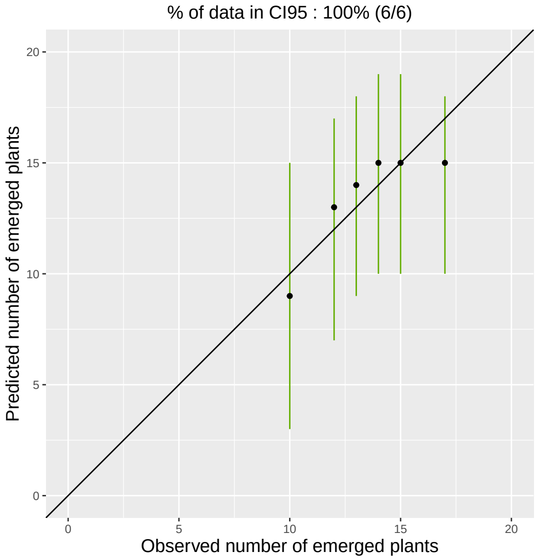

(b) Posterior predictive check (PPC)

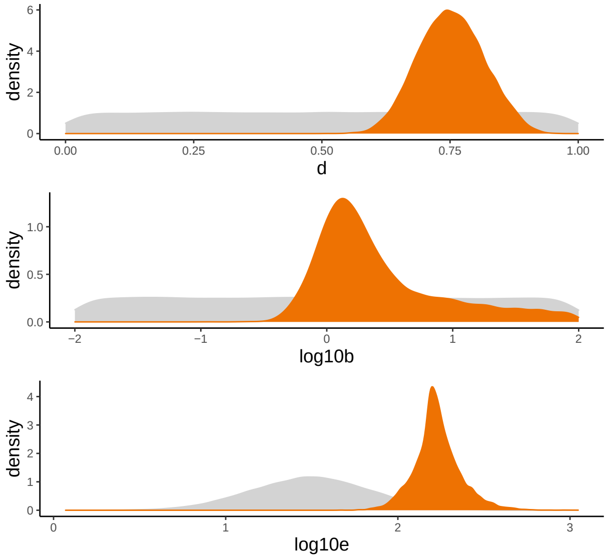

(c) Priors and posteriors

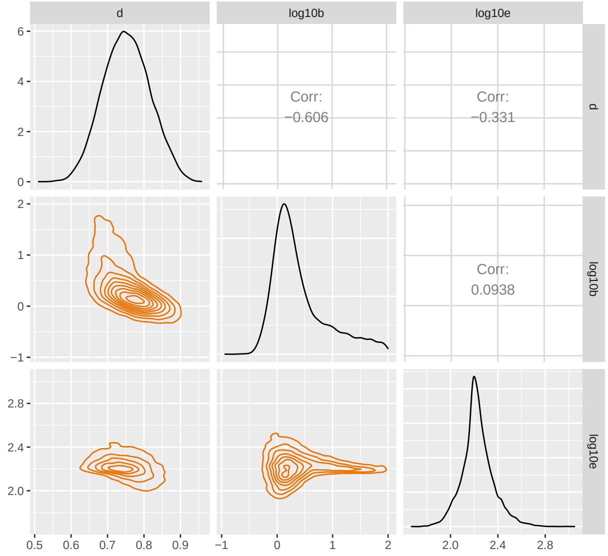

(d) Correlations between parameters

Figure 1: Dose-response curve (a), PPC (b), prior and posterior distributions (c) and correlations between parameters (d).

#### Data set: AVESA_SE_emergence

Table 2: Summary of parameter estimates (parameter d is set to 1) for AVESA_SE_emergence data set

| Parameter | median | Q2.5 | Q97.5 |
|---|---|---|---|
| b | 1.372 | 0.837 | 2.226 |
| e | 319.155 | 184.587 | 714.326 |

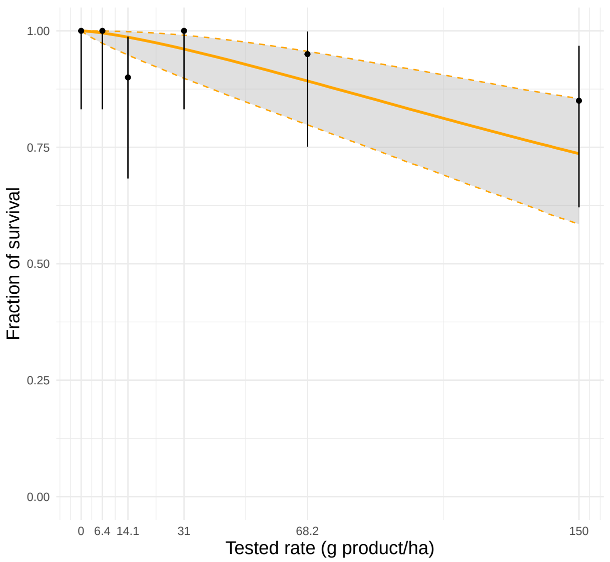

(a) Dose-response curve

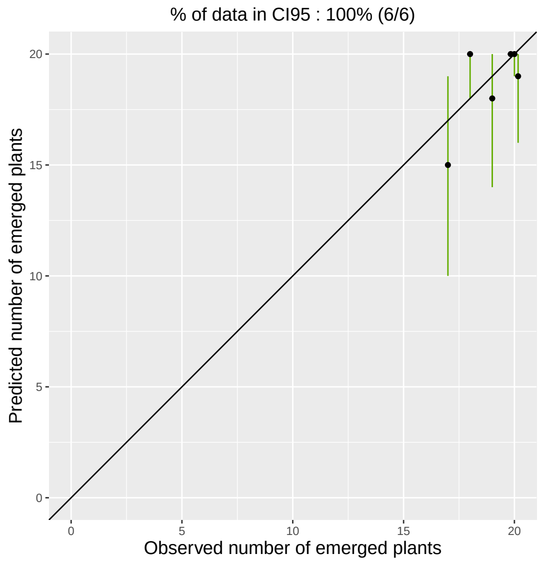

(b) Posterior predictive check (PPC)

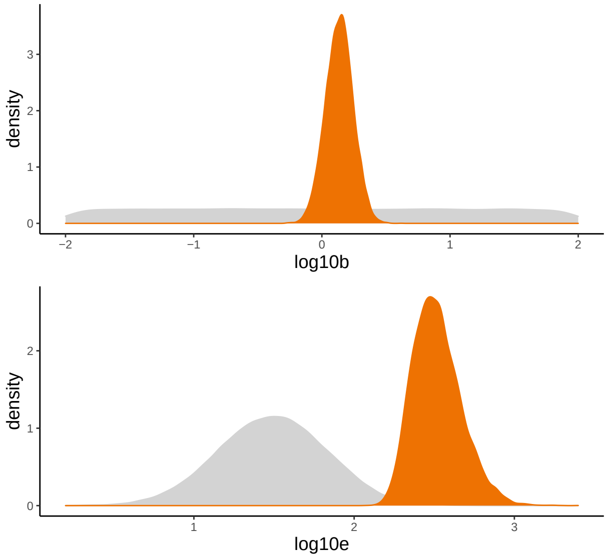

(c) Priors and posteriors

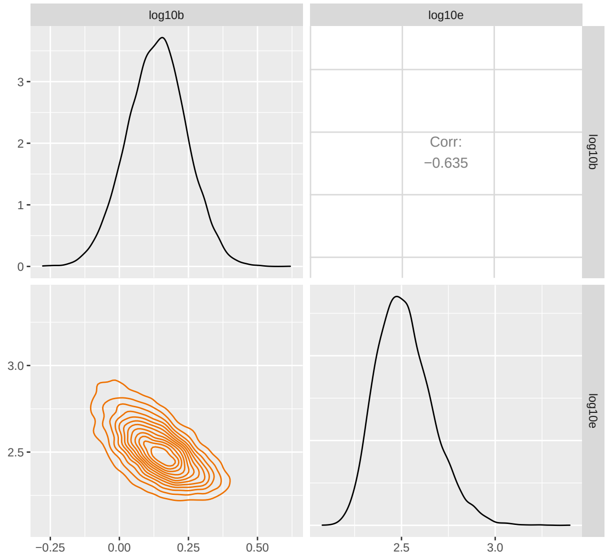

(d) Correlations between parameters

Figure 2: Dose-response curve (a), PPC (b), prior and posterior distributions (c) and correlations between parameters (d).

### Data set: BEAVA_SE_emergence

Table 3: Summary of parameter estimates (parameter d is set to 1) for BEAVA_SE_emergence data set

| Parameter | median | Q2.5 | Q97.5 |
|---|---|---|---|
| b | 2.282 | 1.257 | 4.313 |
| e | 292.419 | 188.077 | 617.988 |

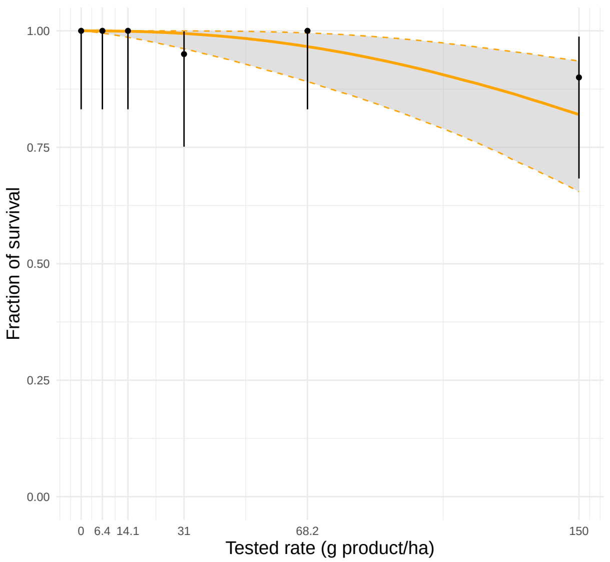

(a) Dose-response curve

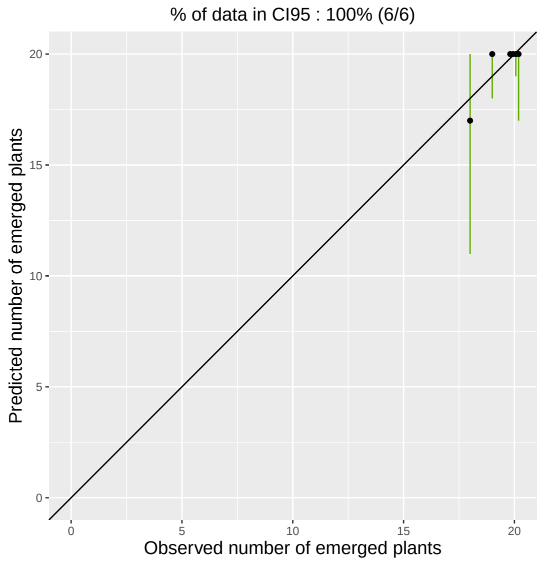

(b) Posterior predictive check (PPC)

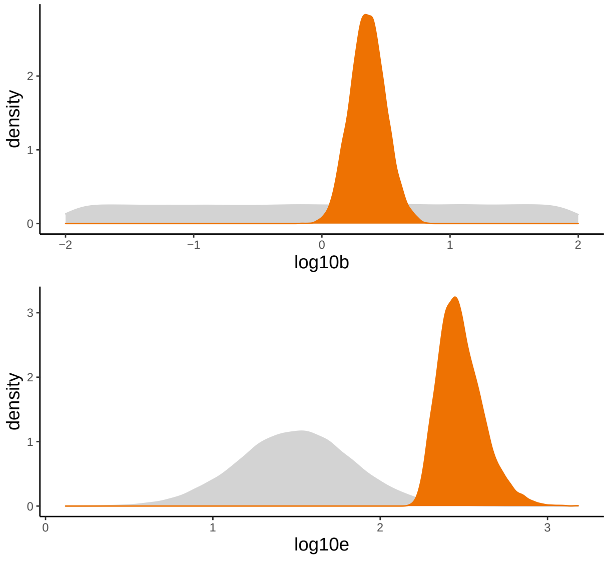

(c) Priors and posteriors

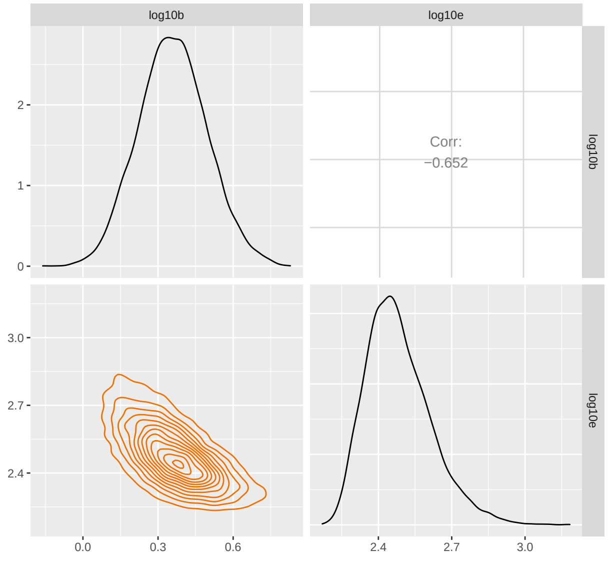

(d) Correlations between parameters

Figure 3: Dose-response curve (a), PPC (b), prior and posterior distributions (c) and correlations between parameters (d).

### Data set: BRSNW_SE_emergence

Table 4: Summary of parameter estimates (parameter d is set to 1) for BRSNW_SE_emergence data set

| Parameter | median | Q2.5 | Q97.5 |
|---|---|---|---|
| b | 2.063 | 1.325 | 2.999 |
| e | 66.484 | 50.832 | 90.237 |

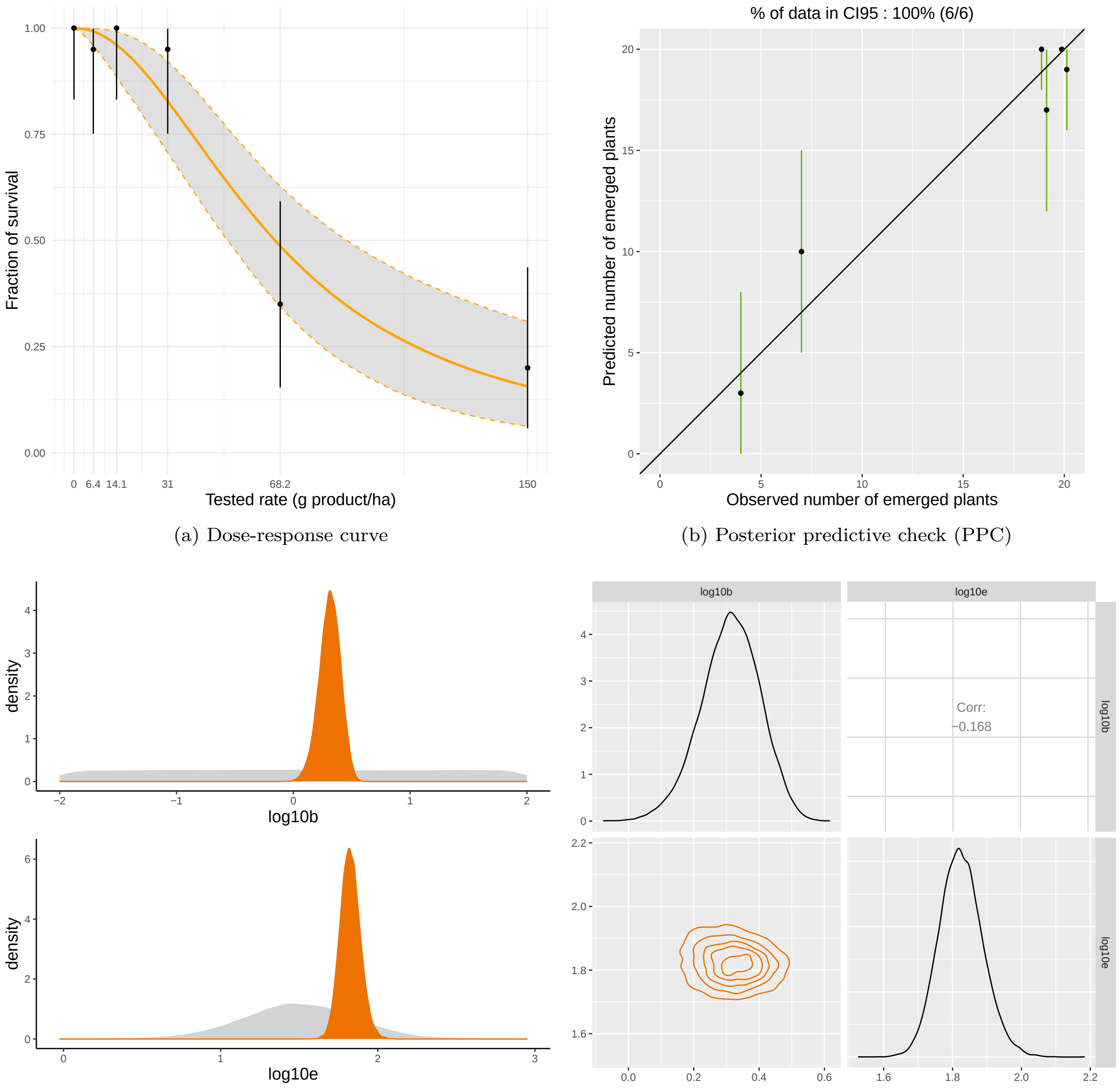

(a) Dose-response curve

(b) Posterior predictive check (PPC)

(c) Priors and posteriors

(d) Correlations between parameters

Figure 4: Dose-response curve (a), PPC (b), prior and posterior distributions (c) and correlations between parameters (d).

#### Data set: CUMSA_SE_emergence

Table 5: Summary of parameter estimates (parameter d is set to 1) for CUMSA_SE_emergence data set

| Parameter | median | Q2.5 | Q97.5 |
|---|---|---|---|
| b | 1.020 | 0.671 | 1.520 |
| e | 528.097 | 262.248 | 1274.037 |

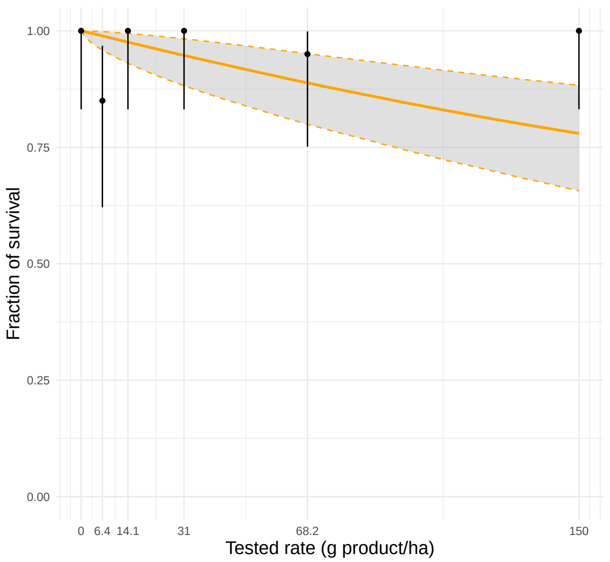

(a) Dose-response curve

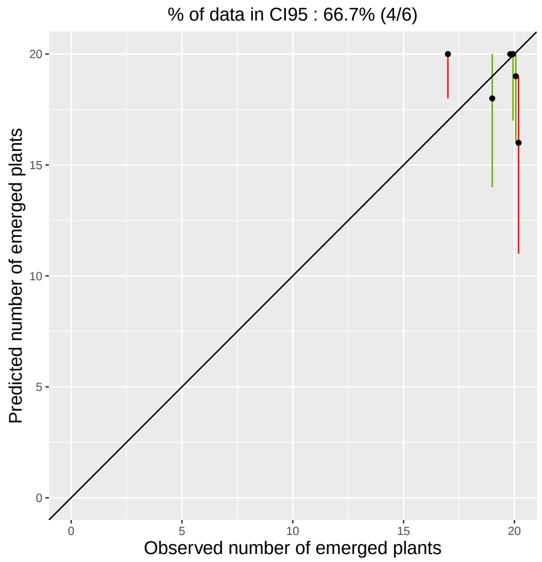

(b) Posterior predictive check (PPC)

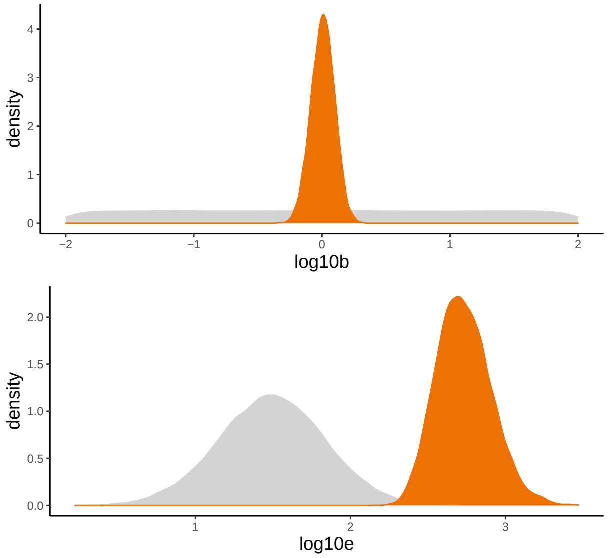

(c) Priors and posteriors

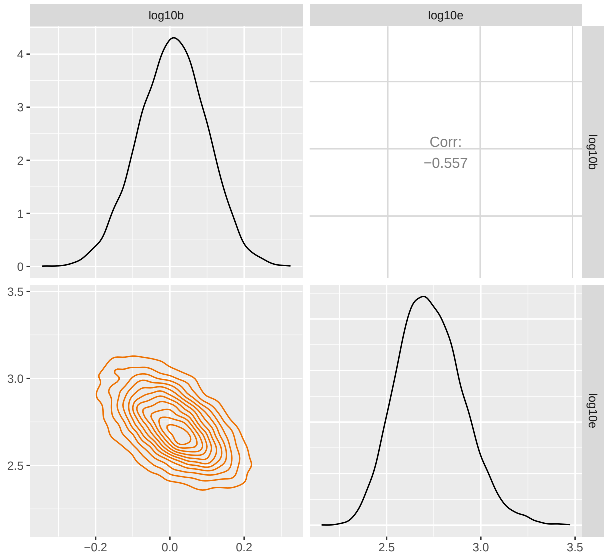

(d) Correlations between parameters

Figure 5: Dose-response curve (a), PPC (b), prior and posterior distributions (c) and correlations between parameters (d).

#### Data set: GLXMA_SE_emergence

Table 6: Summary of parameter estimates (parameter d is set to 1) for GLXMA_SE_emergence data set

| Parameter | median | Q2.5 | Q97.5 |
|---|---|---|---|
| b | 1.308 | 0.829 | 2.049 |
| e | 525.316 | 272.339 | 1279.091 |

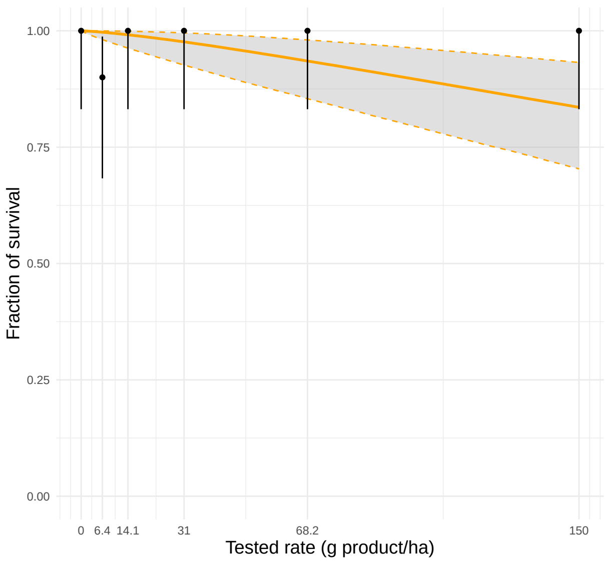

(a) Dose-response curve

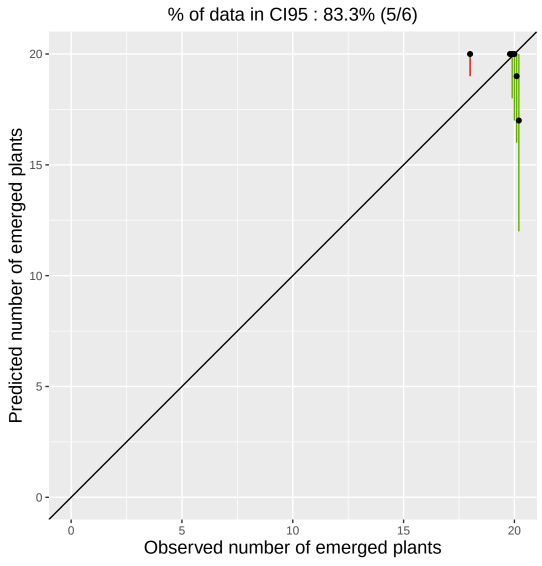

(b) Posterior predictive check (PPC)

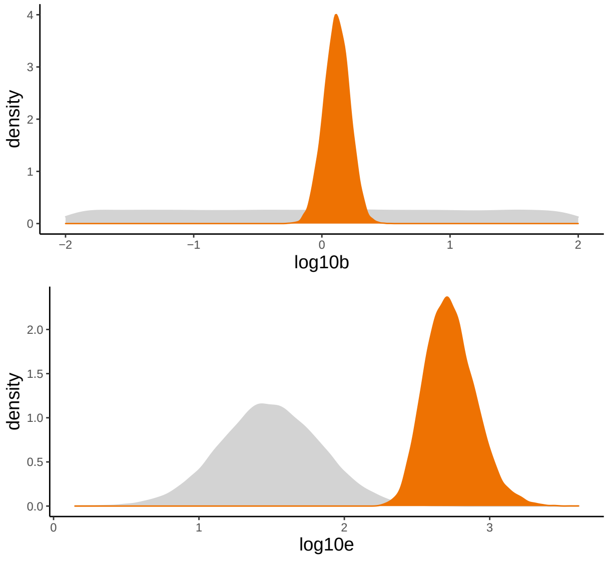

(c) Priors and posteriors

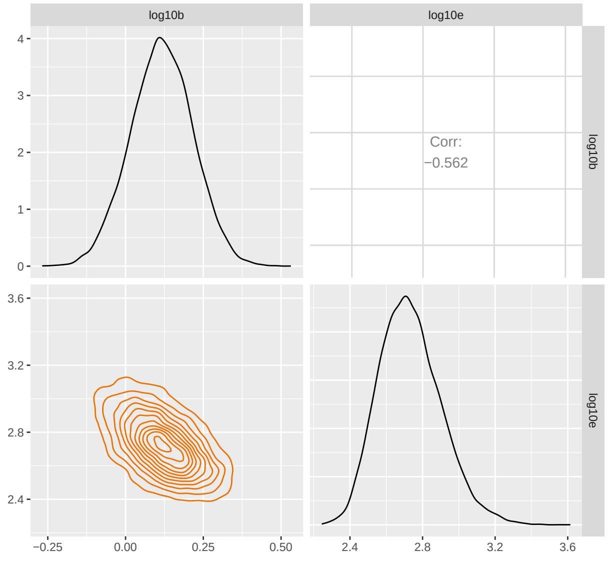

(d) Correlations between parameters

Figure 6: Dose-response curve (a), PPC (b), prior and posterior distributions (c) and correlations between parameters (d).

#### Data set: HELAN_SE_emergence

Table 7: Summary of parameter estimates (parameter d is set to 1) for HELAN_SE_emergence data set

| Parameter | median | Q2.5 | Q97.5 |
|---|---|---|---|
| b | 1.287 | 0.813 | 2.032 |
| e | 466.659 | 249.519 | 1100.256 |

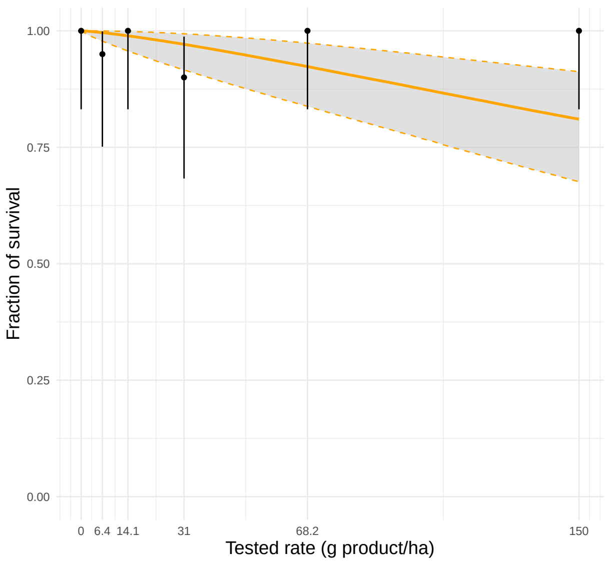

(a) Dose-response curve

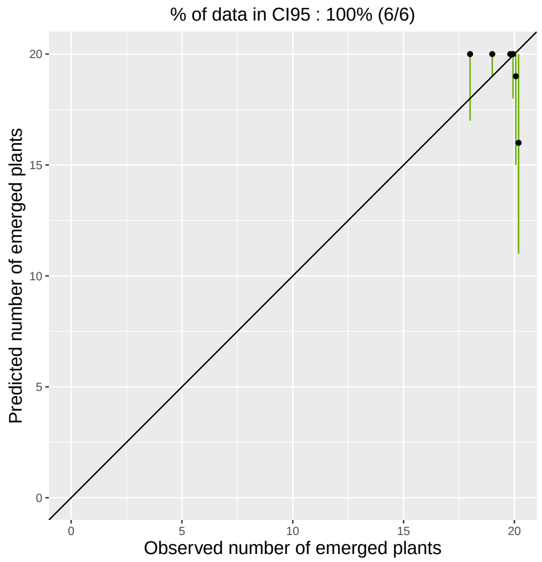

(b) Posterior predictive check (PPC)

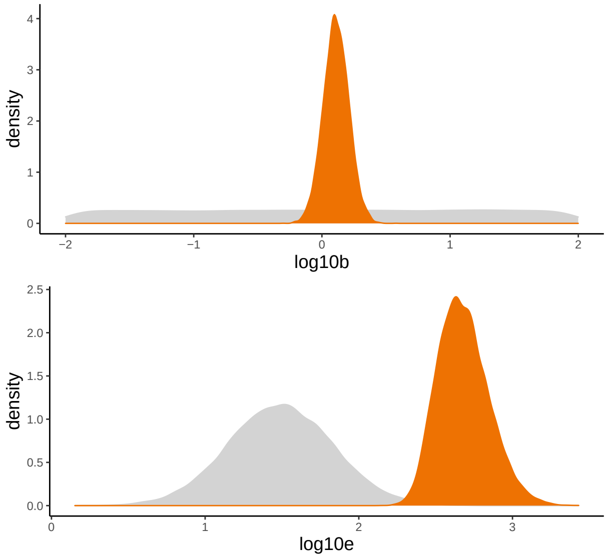

(c) Priors and posteriors

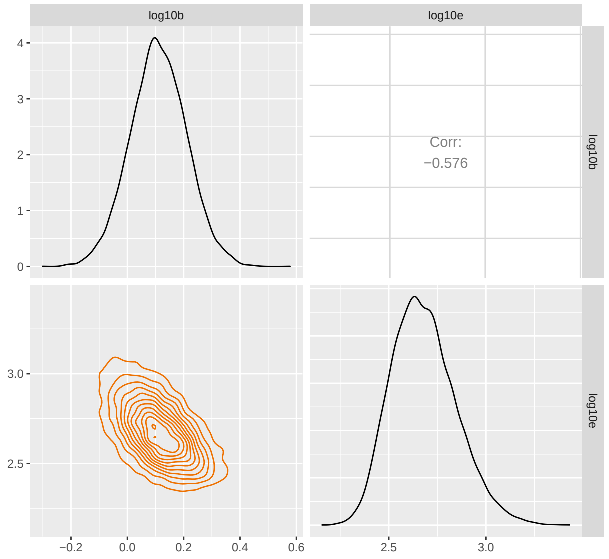

(d) Correlations between parameters

Figure 7: Dose-response curve (a), PPC (b), prior and posterior distributions (c) and correlations between parameters (d).

#### Data set: LYPES_SE_emergence

Table 8: Summary of parameter estimates (parameter d is set to 1) for LYPES_SE_emergence data set

| Parameter | median | Q2.5 | Q97.5 |
|---|---|---|---|
| b | 0.808 | 0.532 | 1.191 |
| e | 490.435 | 239.841 | 1224.281 |

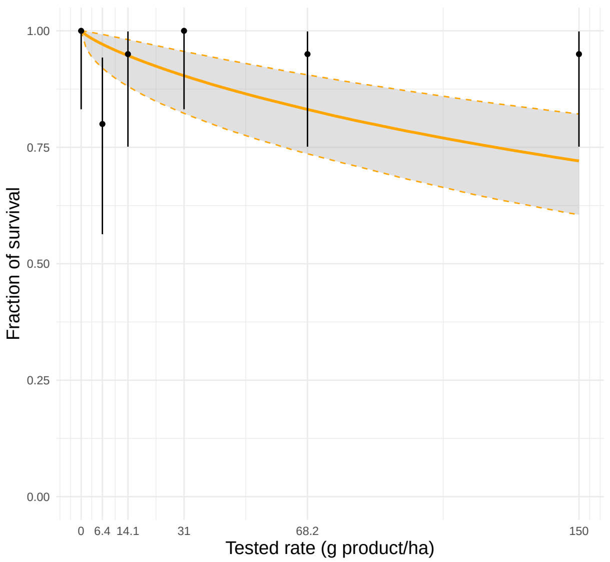

(a) Dose-response curve

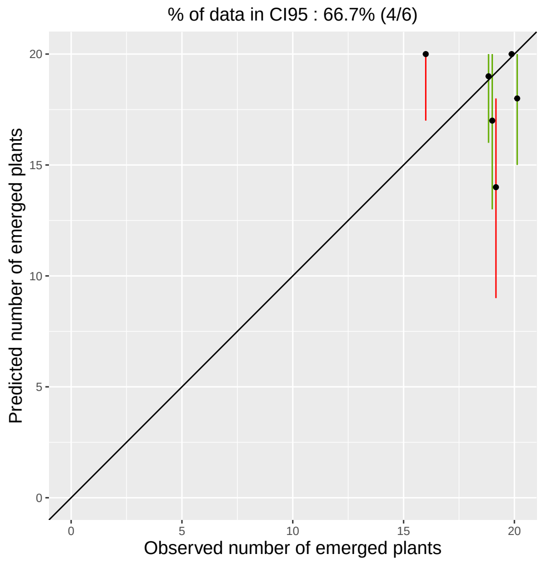

(b) Posterior predictive check (PPC)

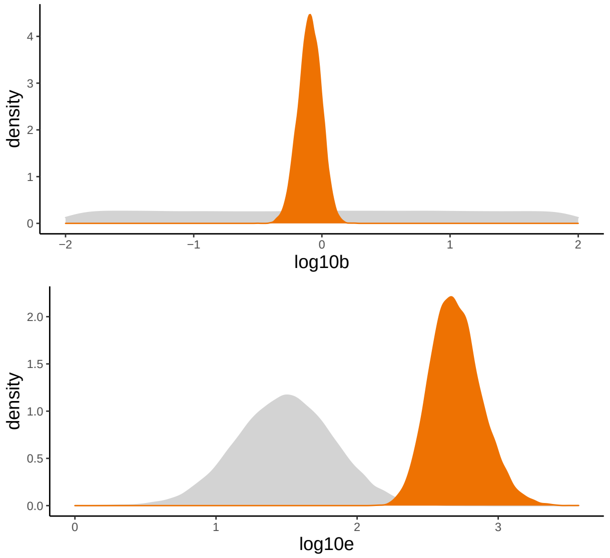

(c) Priors and posteriors

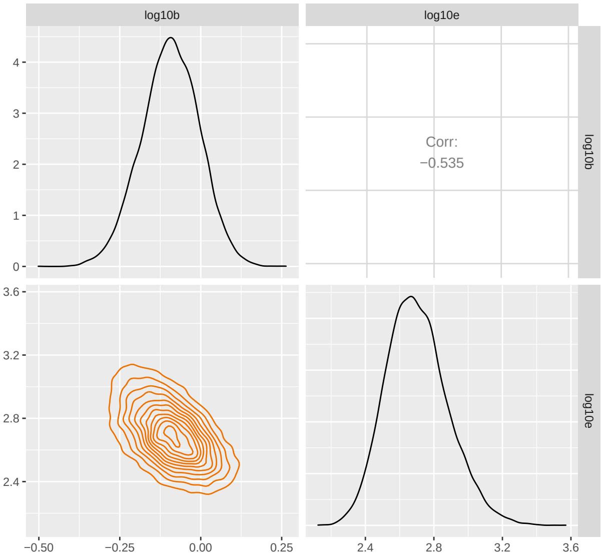

(d) Correlations between parameters

Figure 8: Dose-response curve (a), PPC (b), prior and posterior distributions (c) and correlations between parameters (d).

#### Data set: TRZAW_SE_emergence

Table 9: Summary of parameter estimates (parameter d is set to 1) for TRZAW_SE_emergence data set

| Parameter | median | Q2.5 | Q97.5 |
|---|---|---|---|
| b | 1.375 | 0.856 | 2.188 |
| e | 437.287 | 237.799 | 1049.467 |

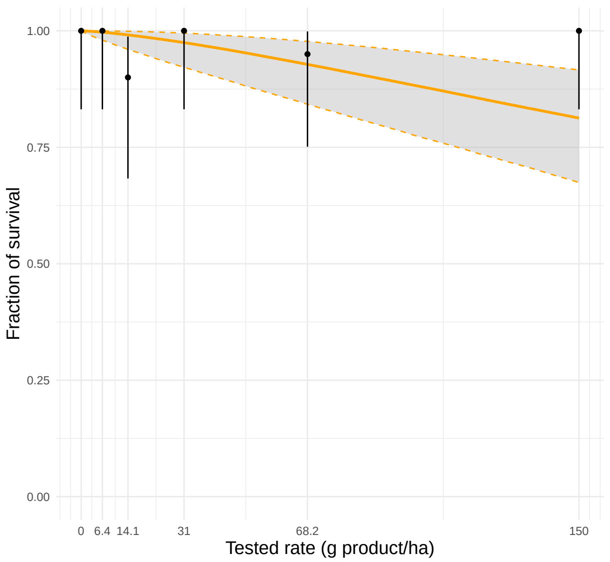

(a) Dose-response curve

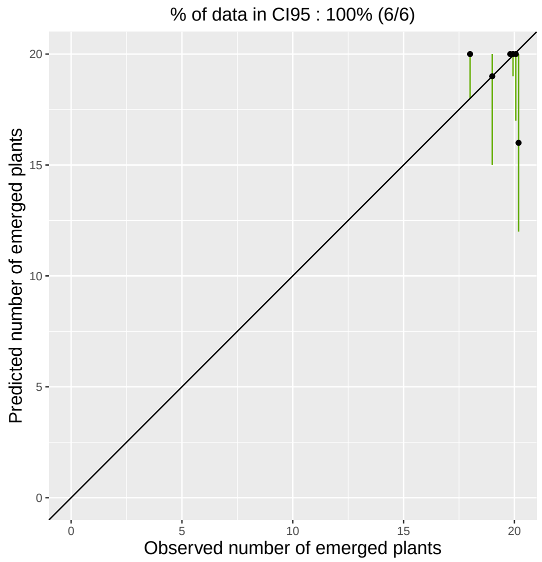

(b) Posterior predictive check (PPC)

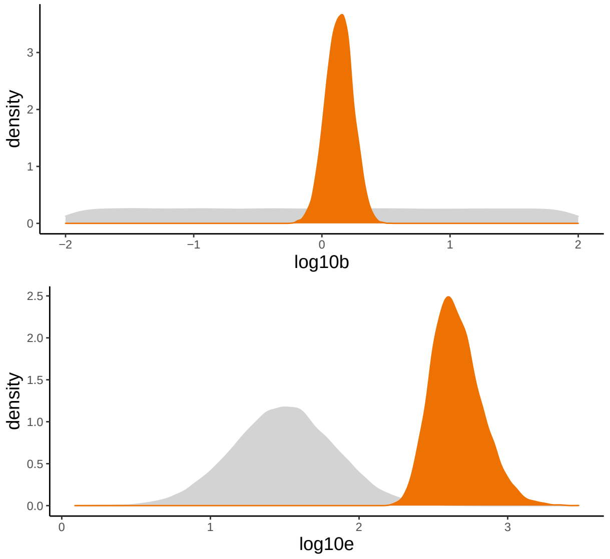

(c) Priors and posteriors

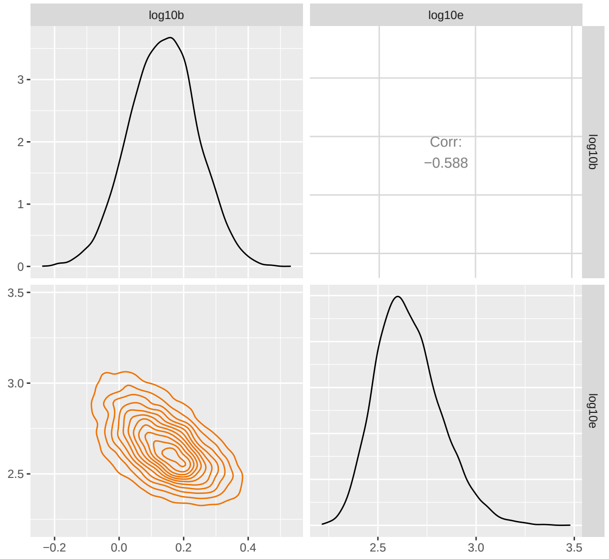

(d) Correlations between parameters

Figure 9: Dose-response curve (a), PPC (b), prior and posterior distributions (c) and correlations between parameters (d).

#### Data set: ZEAMA_SE_emergence

Table 10: Summary of parameter estimates (parameter d is set to 1) for ZEAMA_SE_emergence data set

| Parameter | median | Q2.5 | Q97.5 |
|---|---|---|---|
| b | 1.462 | 0.904 | 2.366 |
| e | 411.484 | 228.897 | 947.331 |

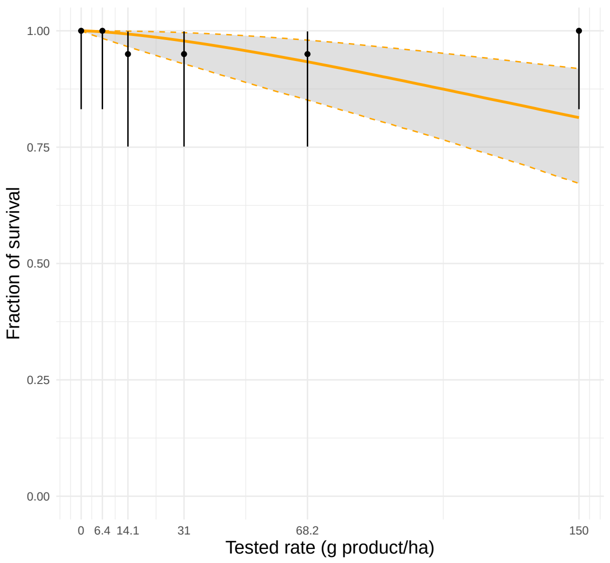

(a) Dose-response curve

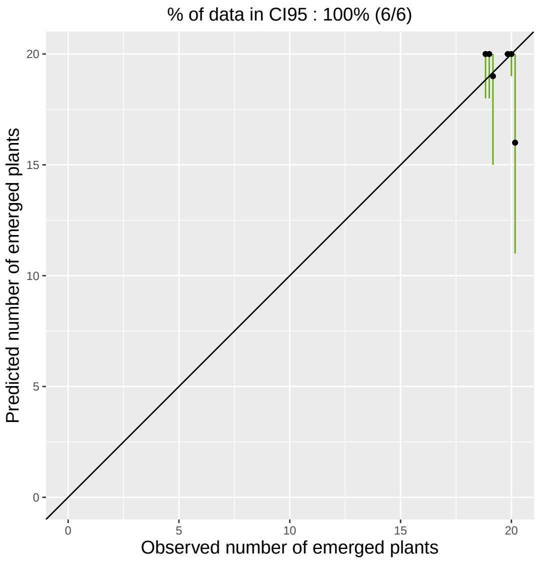

(b) Posterior predictive check (PPC)

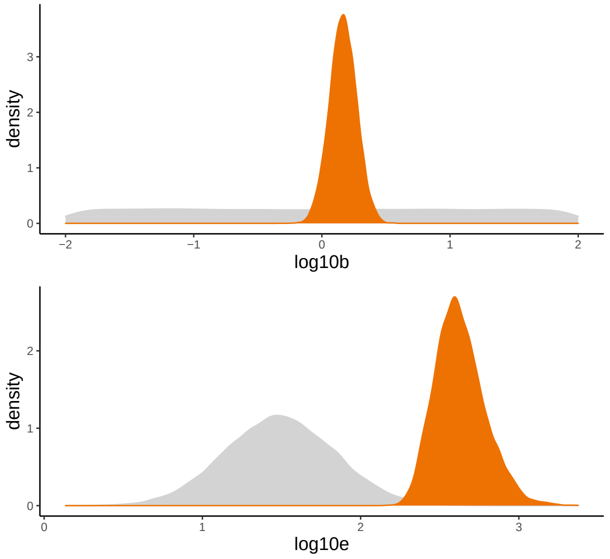

(c) Priors and posteriors

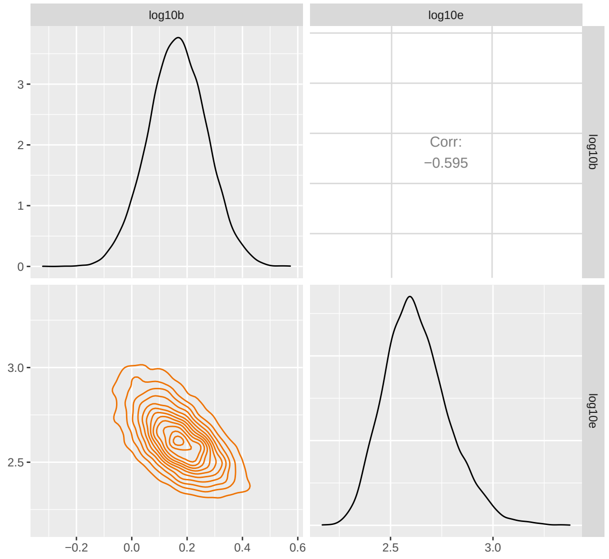

(d) Correlations between parameters

Figure 10: Dose-response curve (a), PPC (b), prior and posterior distributions (c) and correlations between parameters (d).
