## Supplementary material for "How to account for the uncertainty from standard toxicity tests in species sensitivity distributions: an example in non-target plants": Archive with reports on studies: report_SE_survival.pdf

### Dose-response analyses

Study 6

Seedling Emergence test - survival endpoint

25 June 2020

Contact:

---

This is a report which provides results on all performed dose-response analyses for the survival endpoint of the Seedling Emergence test for study 6.

---

#### Data set: ALLCE_SE_survival

Table 1: Summary of parameter estimates (parameter d is set to 1) for ALLCE_SE_survival data set

| Parameter | median | Q2.5 | Q97.5 |
|---|---|---|---|
| b | 2.468 | 1.486 | 3.859 |
| e | 20.219 | 15.065 | 27.465 |

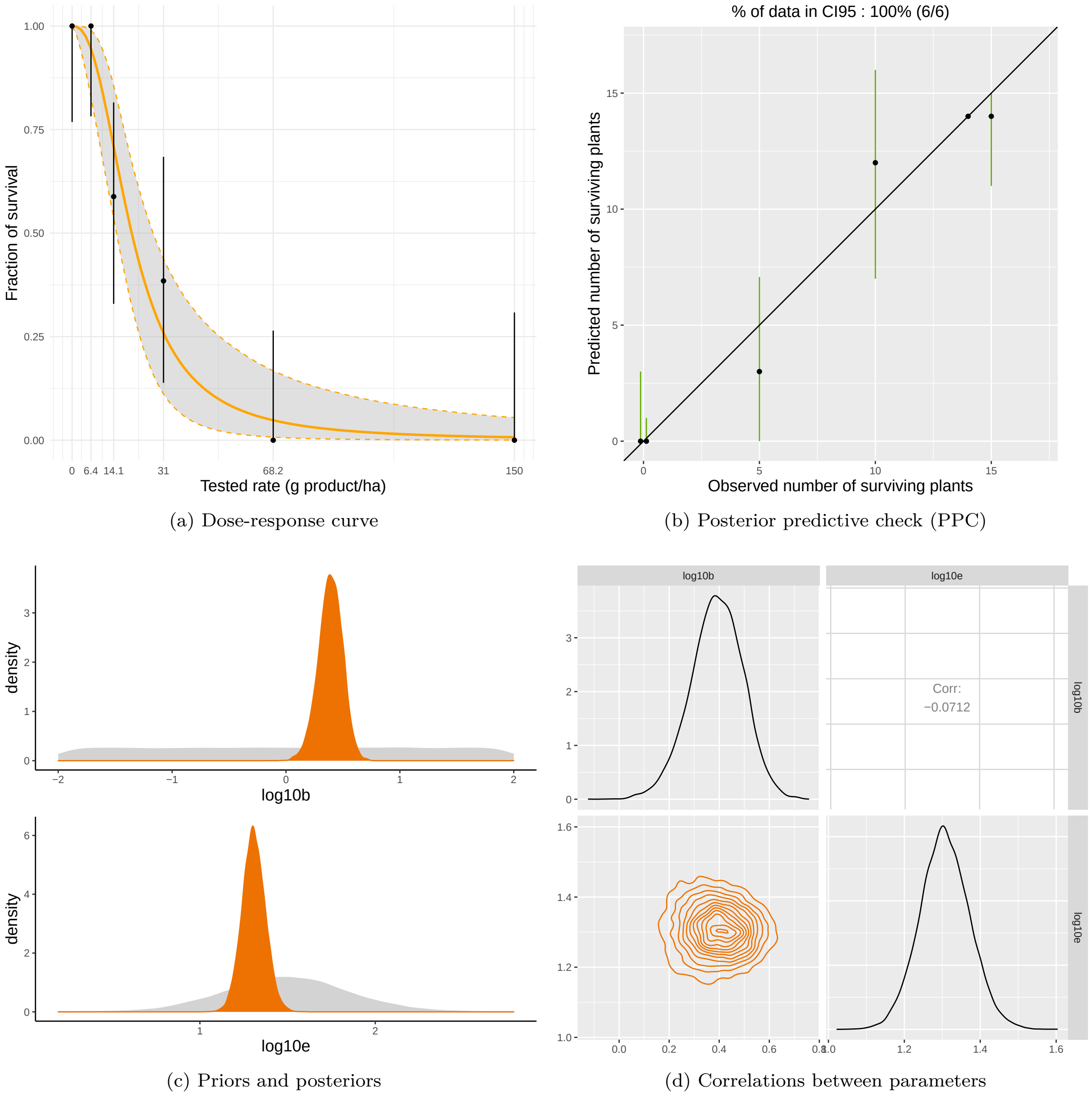

(a) Dose-response curve

(b) Posterior predictive check (PPC)

(c) Priors and posteriors

(d) Correlations between parameters

Figure 1: Dose-response curve (a), PPC (b), prior and posterior distributions (c) and correlations between parameters (d).

### Data set: AVESA_SE_survival

Table 2: Summary of parameter estimates (parameter d is set to 1) for AVESA_SE_survival data set

| Parameter | median | Q2.5 | Q97.5 |
|---|---|---|---|
| b | 2.390 | 1.318 | 4.461 |
| e | 351.254 | 211.446 | 779.351 |

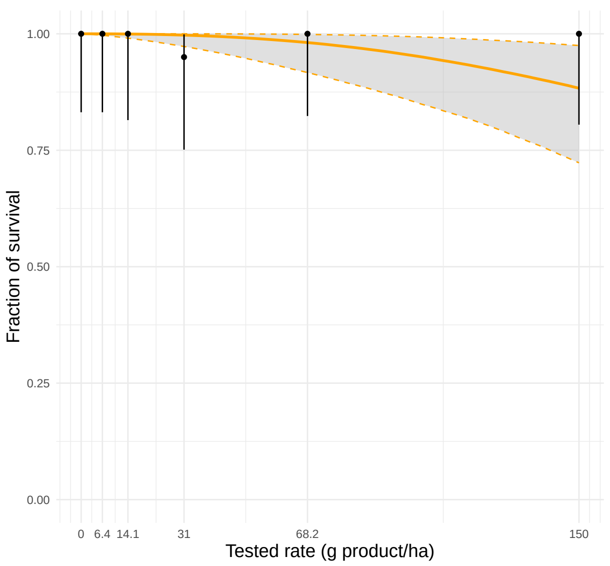

(a) Dose-response curve

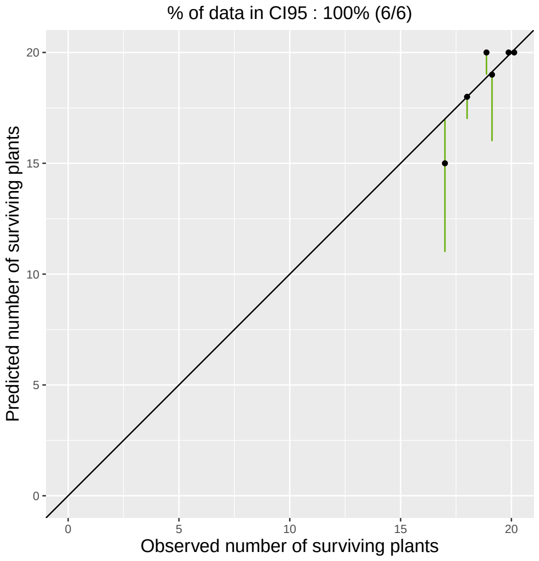

(b) Posterior predictive check (PPC)

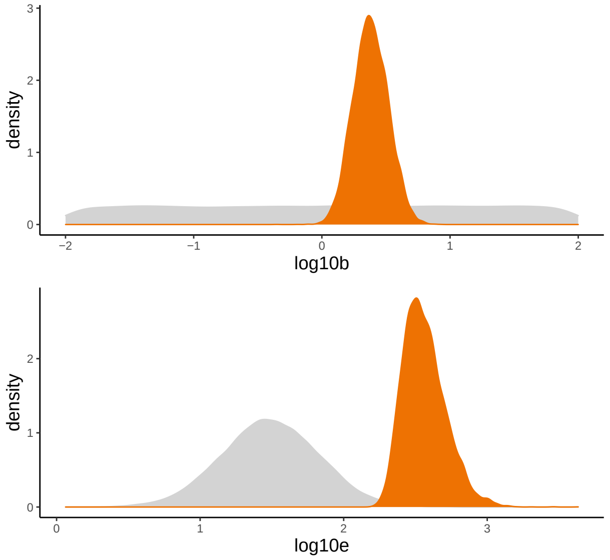

(c) Priors and posteriors

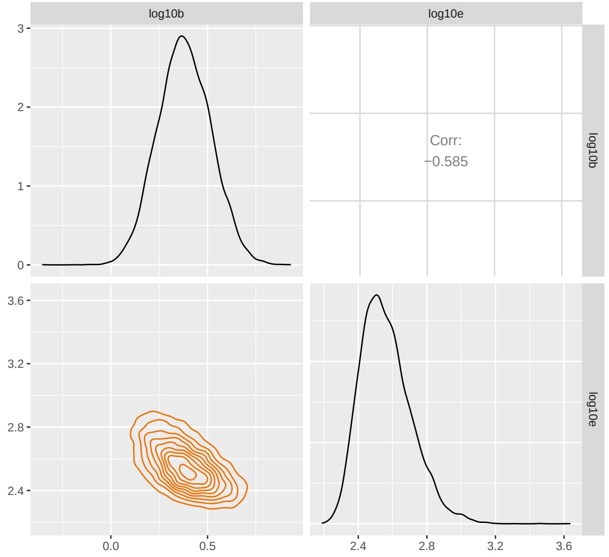

(d) Correlations between parameters

Figure 2: Dose-response curve (a), PPC (b), prior and posterior distributions (c) and correlations between parameters (d).

#### Data set: BEAVA_SE_survival

Table 3: Summary of parameter estimates (parameter d is set to 1) for BEAVA_SE_survival data set

| Parameter | median | Q2.5 | Q97.5 |
|---|---|---|---|
| b | 3.794 | 2.437 | 5.922 |
| e | 32.856 | 26.918 | 40.254 |

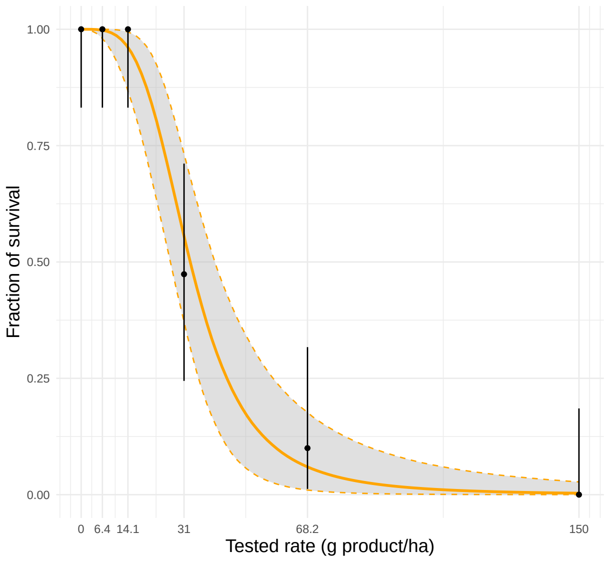

(a) Dose-response curve

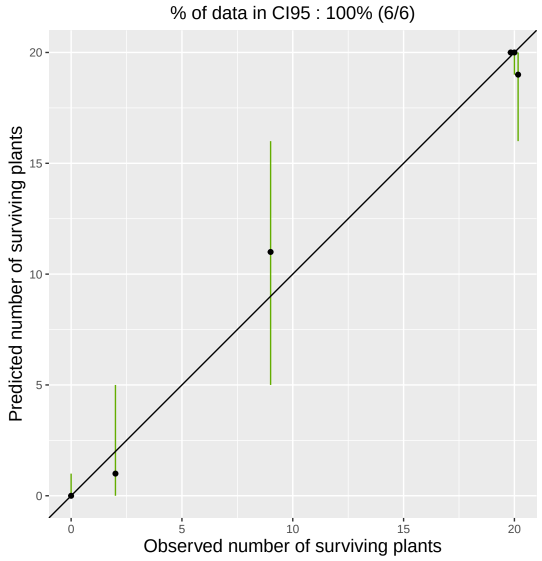

(b) Posterior predictive check (PPC)

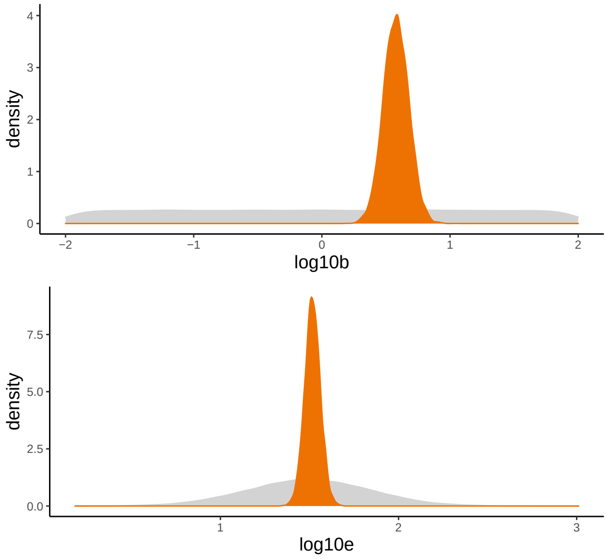

(c) Priors and posteriors

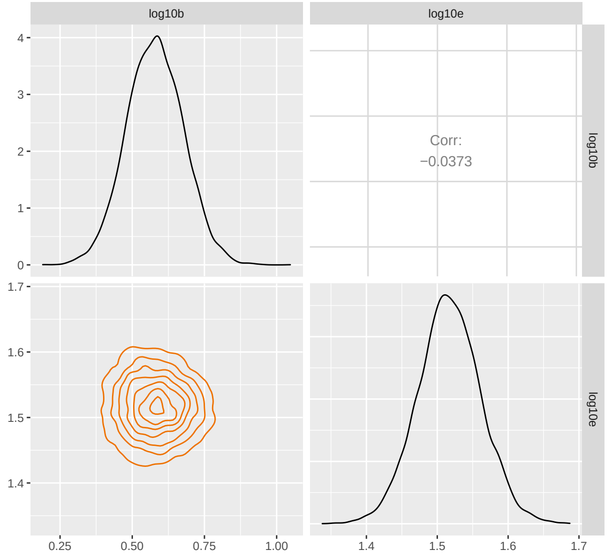

(d) Correlations between parameters

Figure 3: Dose-response curve (a), PPC (b), prior and posterior distributions (c) and correlations between parameters (d).

#### Data set: BRSNW_SE_survival

Table 4: Summary of parameter estimates (parameter d is set to 1) for BRSNW_SE_survival data set

| Parameter | median | Q2.5 | Q97.5 |
|---|---|---|---|
| b | 3.562 | 1.745 | 6.979 |
| e | 84.731 | 61.441 | 138.356 |

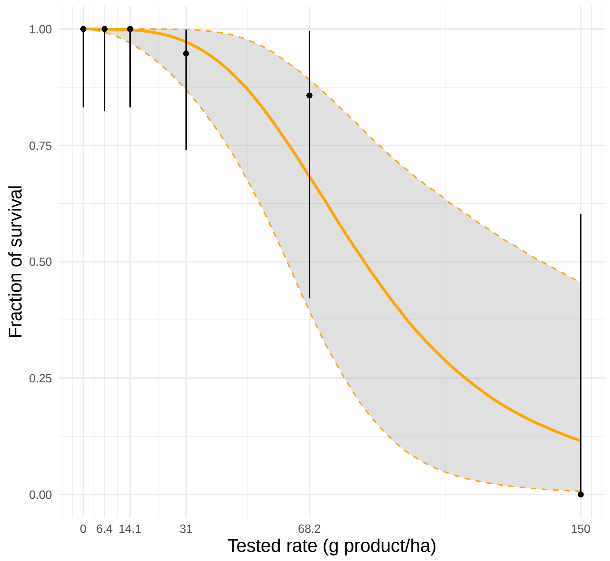

(a) Dose-response curve

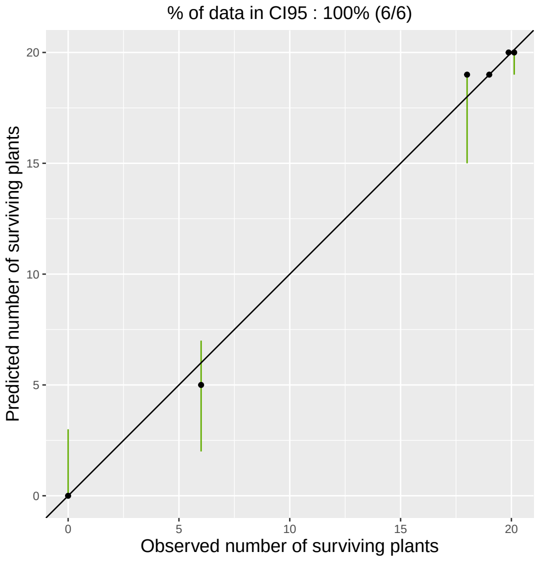

(b) Posterior predictive check (PPC)

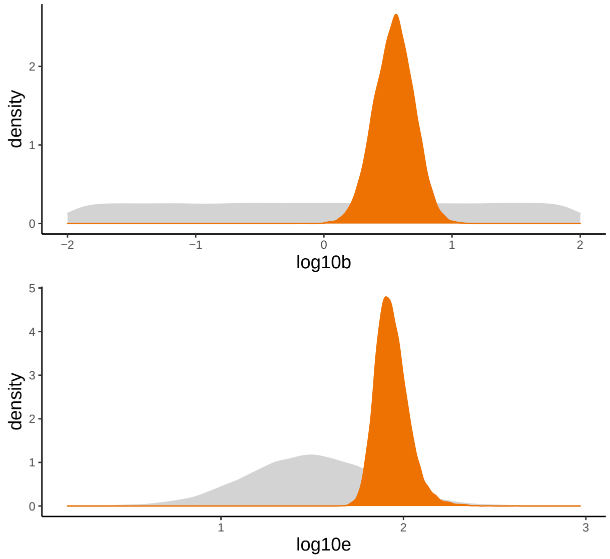

(c) Priors and posteriors

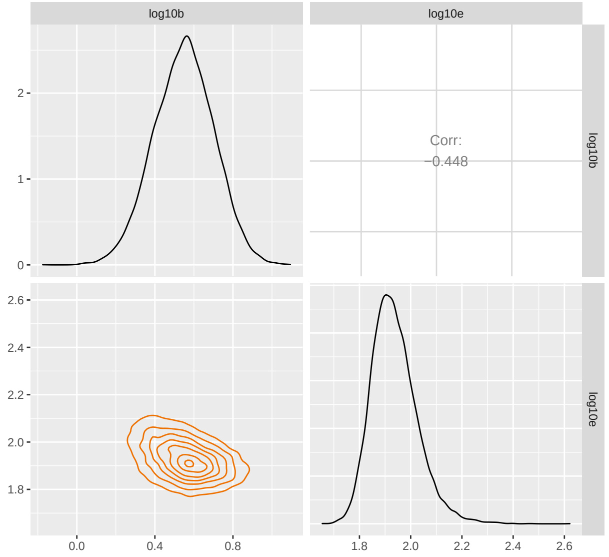

(d) Correlations between parameters

Figure 4: Dose-response curve (a), PPC (b), prior and posterior distributions (c) and correlations between parameters (d).

#### Data set: CUMSA_SE_survival

Table 5: Summary of parameter estimates (parameter d is set to 1) for CUMSA_SE_survival data set

| Parameter | median | Q2.5 | Q97.5 |
|---|---|---|---|
| b | 34.411 | 5.257 | 95.256 |
| e | 215.787 | 160.724 | 482.997 |

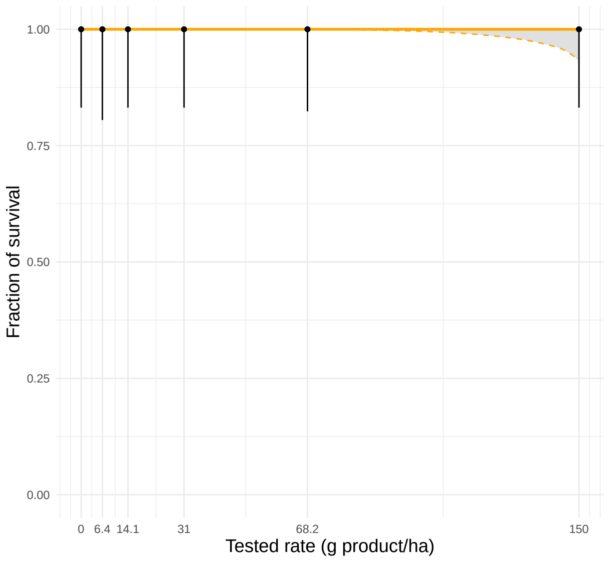

(a) Dose-response curve

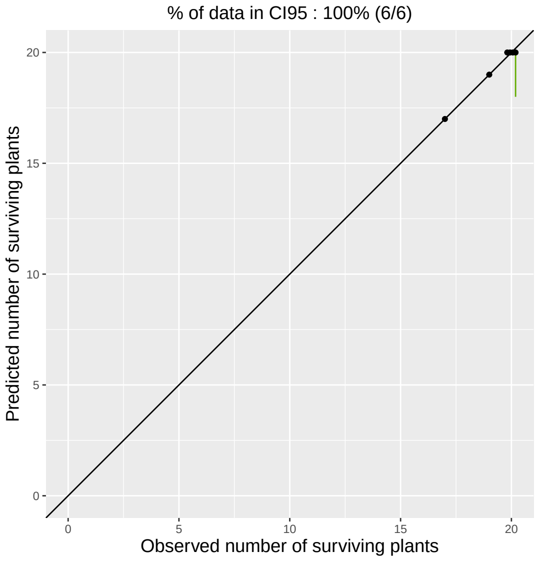

(b) Posterior predictive check (PPC)

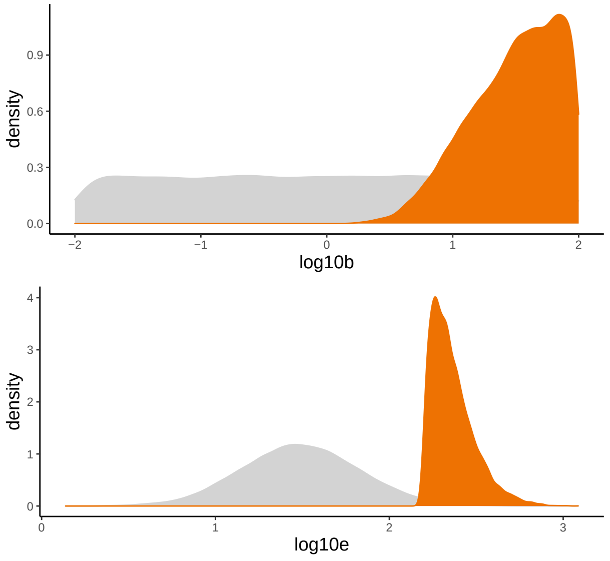

(c) Priors and posteriors

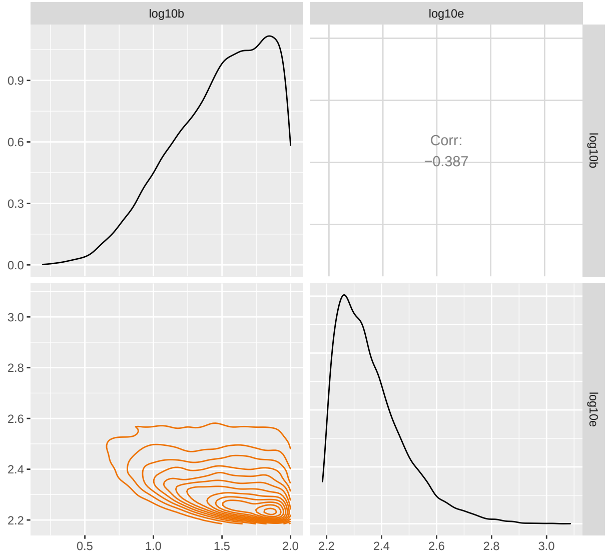

(d) Correlations between parameters

Figure 5: Dose-response curve (a), PPC (b), prior and posterior distributions (c) and correlations between parameters (d).

#### Data set: GLXMA_SE_survival

Table 6: Summary of parameter estimates (parameter d is set to 1) for GLXMA_SE_survival data set

| Parameter | median | Q2.5 | Q97.5 |
|---|---|---|---|
| b | 35.148 | 5.326 | 94.472 |
| e | 213.801 | 160.775 | 459.948 |

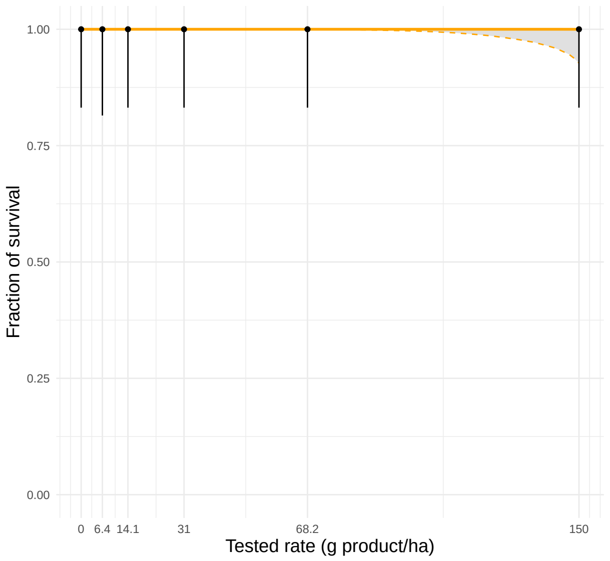

(a) Dose-response curve

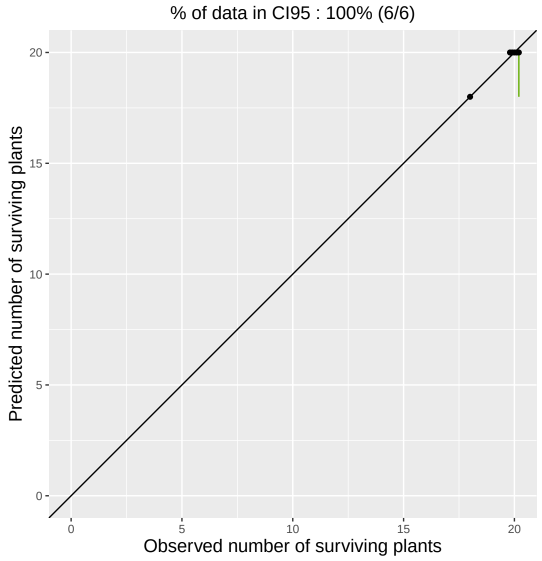

(b) Posterior predictive check (PPC)

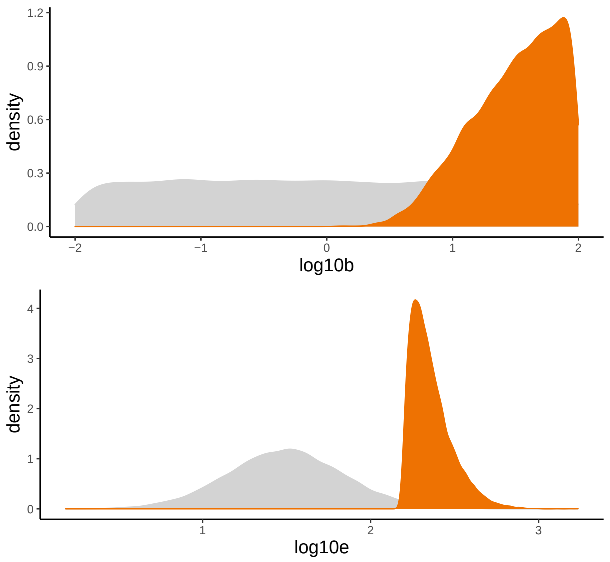

(c) Priors and posteriors

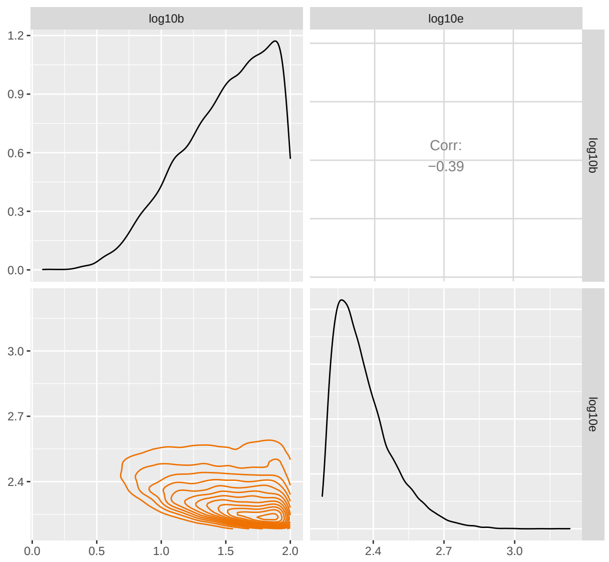

(d) Correlations between parameters

Figure 6: Dose-response curve (a), PPC (b), prior and posterior distributions (c) and correlations between parameters (d).

#### Data set: HELAN_SE_survival

Table 7: Summary of parameter estimates (parameter d is set to 1) for HELAN_SE_survival data set

| Parameter | median | Q2.5 | Q97.5 |
|---|---:|---:|---:|
| b | 3.308 | 1.842 | 5.632 |
| e | 130.690 | 104.186 | 174.427 |

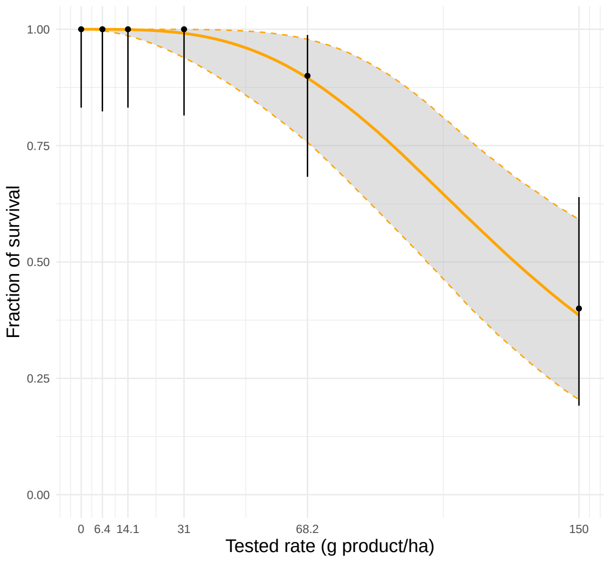

(a) Dose-response curve

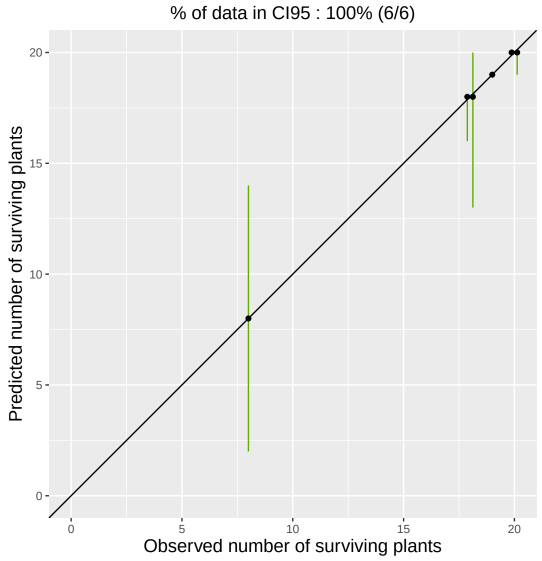

(b) Posterior predictive check (PPC)

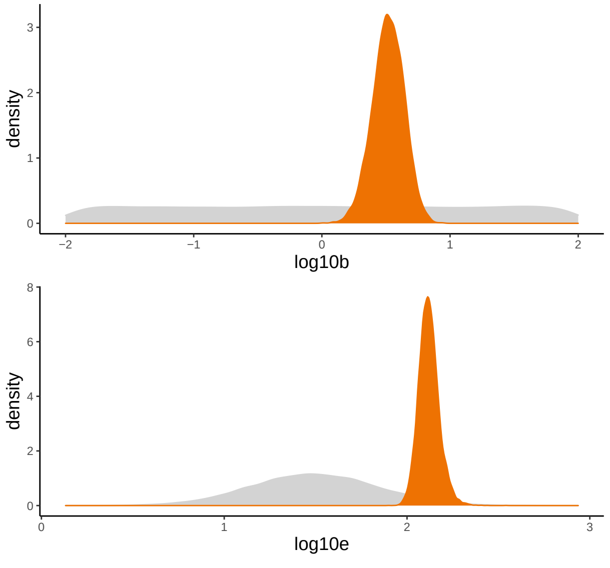

(c) Priors and posteriors

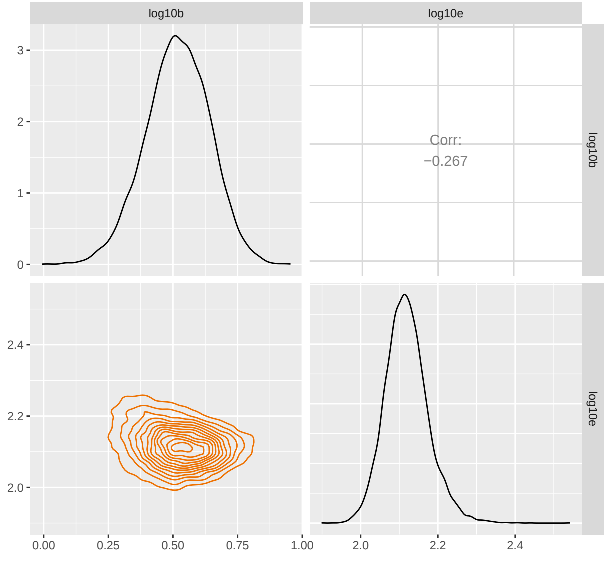

(d) Correlations between parameters

Figure 7: Dose-response curve (a), PPC (b), prior and posterior distributions (c) and correlations between parameters (d).

#### Data set: LYPES_SE_survival

Table 8: Summary of parameter estimates for LYPES_SE_survival data set

| Parameter | median | Q2.5 | Q97.5 |
|---|---|---|---|
| b | 2.628 | 1.298 | 29.481 |
| d | 0.971 | 0.913 | 0.996 |
| e | 181.078 | 136.448 | 324.500 |

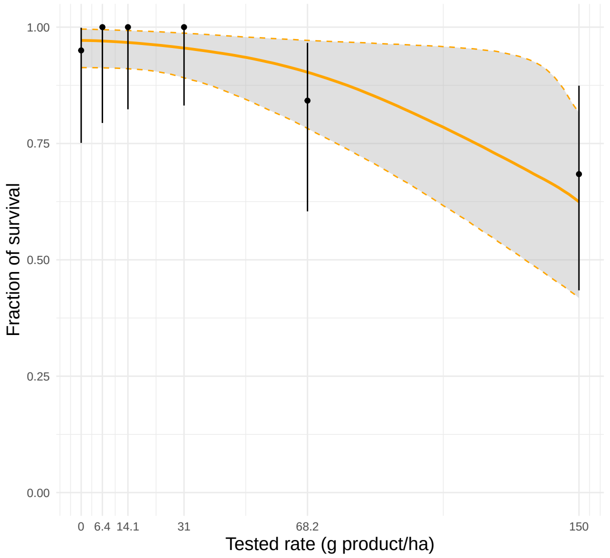

(a) Dose-response curve

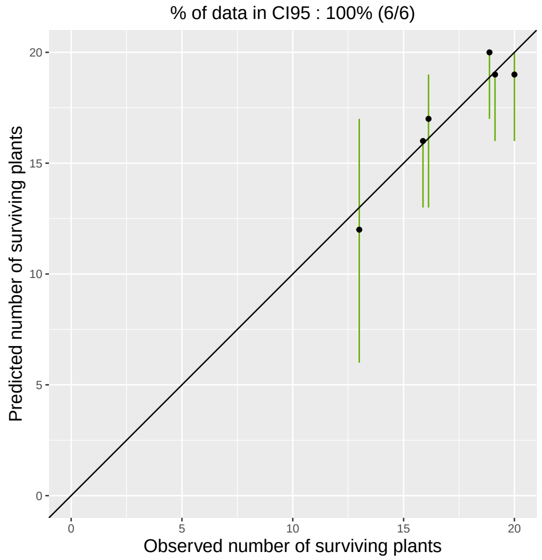

(b) Posterior predictive check (PPC)

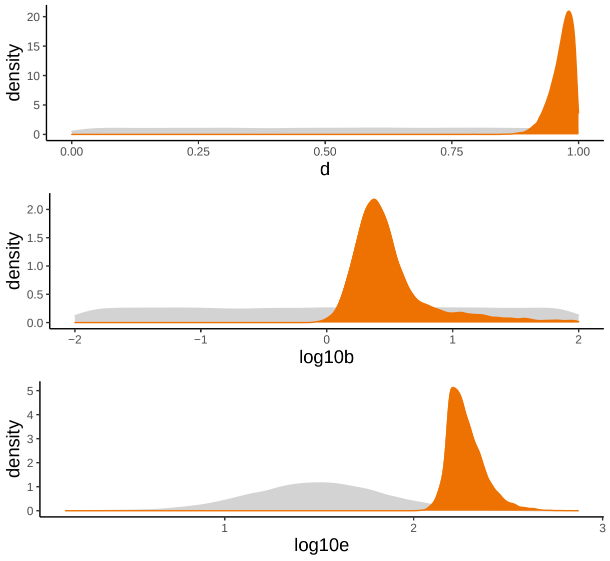

(c) Priors and posteriors

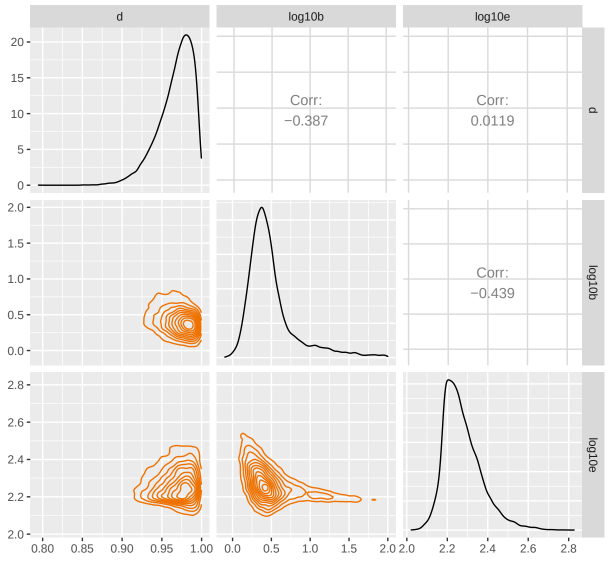

(d) Correlations between parameters

Figure 8: Dose-response curve (a), PPC (b), prior and posterior distributions (c) and correlations between parameters (d).

#### Data set: TRZAW_SE_survival

Table 9: Summary of parameter estimates (parameter d is set to 1) for TRZAW_SE_survival data set

| Parameter | median | Q2.5 | Q97.5 |
|---|---|---|---|
| b | 34.574 | 5.062 | 95.231 |
| e | 214.501 | 160.705 | 479.161 |

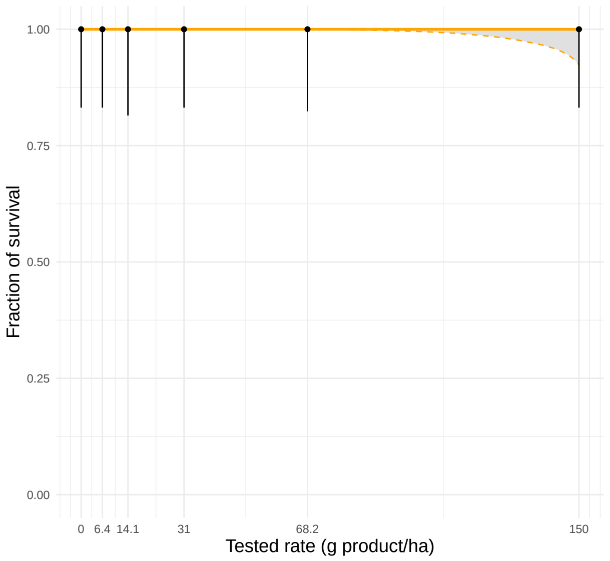

(a) Dose-response curve

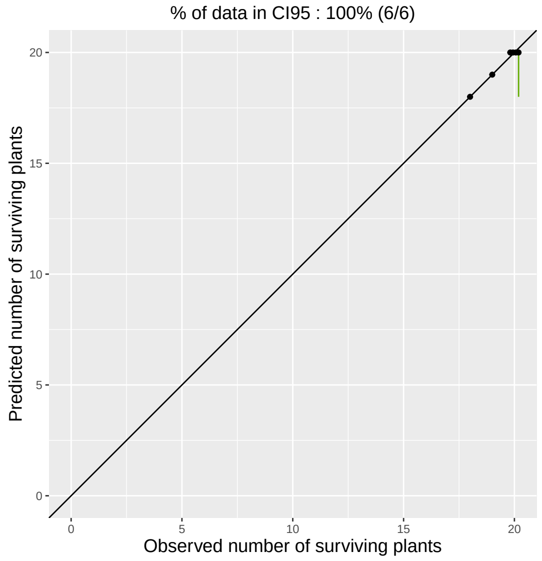

(b) Posterior predictive check (PPC)

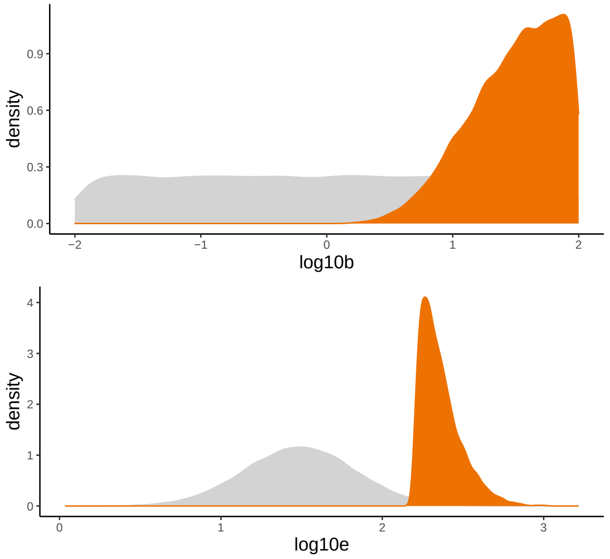

(c) Priors and posteriors

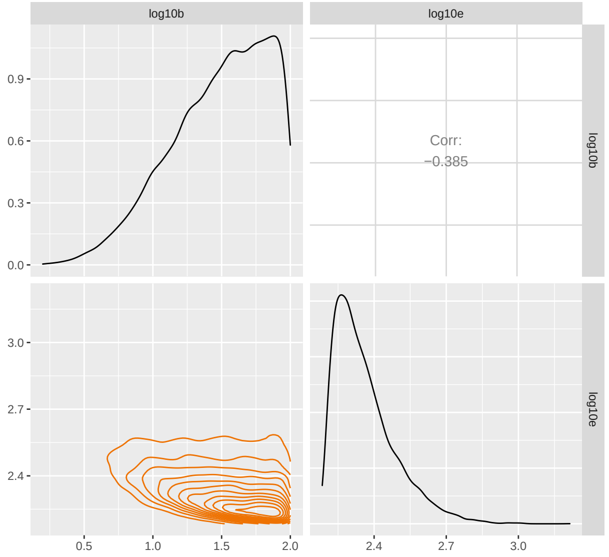

(d) Correlations between parameters

Figure 9: Dose-response curve (a), PPC (b), prior and posterior distributions (c) and correlations between parameters (d).

#### Data set: ZEAMA_SE_survival

Table 10: Summary of parameter estimates (parameter d is set to 1) for ZEAMA_SE_survival data set

| Parameter | median | Q2.5 | Q97.5 |
|---|---|---|---|
| b | 35.041 | 5.303 | 95.455 |
| e | 215.309 | 160.312 | 472.468 |

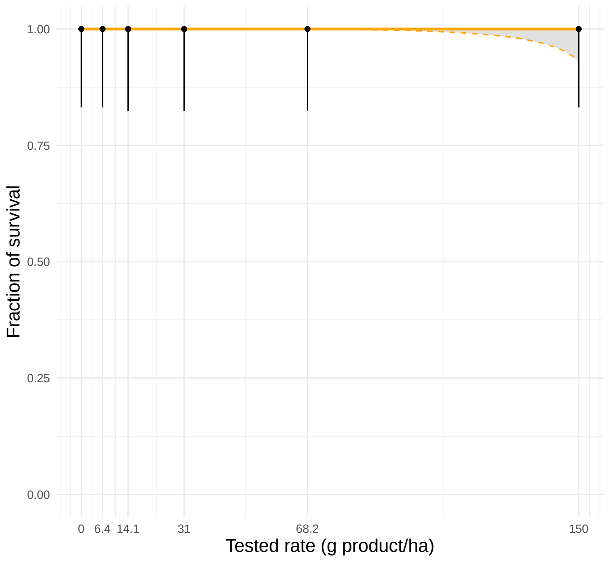

(a) Dose-response curve

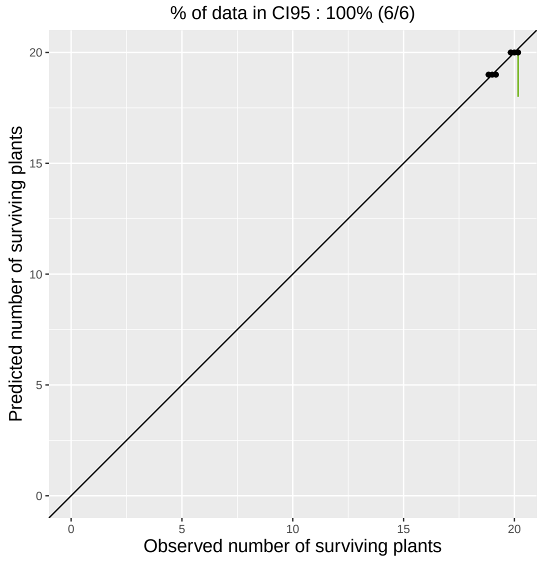

(b) Posterior predictive check (PPC)

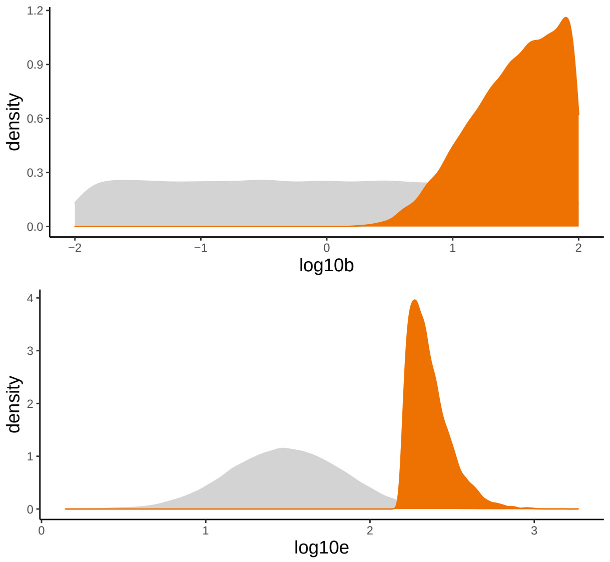

(c) Priors and posteriors

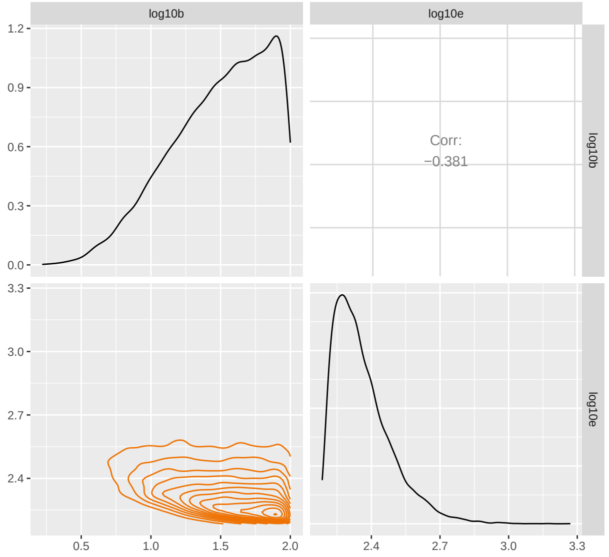

(d) Correlations between parameters

Figure 10: Dose-response curve (a), PPC (b), prior and posterior distributions (c) and correlations between parameters (d).
