## Supplementary material for "How to account for the uncertainty from standard toxicity tests in species sensitivity distributions: an example in non-target plants": Archive with reports on studies: report_SE_weight.pdf

### Dose-response analysis

Study 6

Seedling Emergence test - shoot dry SE_weight endpoint

25 June 2020

Contact:

---

This is a report which provides results on all performed dose-response analyses for the shoot dry SE_weight endpoint of the Seedling Emergence test for study 6.

---

#### Data set: ALLCE_SE_weight

Table 1: Summary of parameter estimates for ALLCE_SE_weight data set

| Parameter | median | Q2.5 | Q97.5 |
|---|---|---|---|
| b | 3.223 | 0.283 | 62.946 |
| d | 0.023 | 0.017 | 0.029 |
| e | 12.046 | 7.699 | 19.231 |
| sigma | 0.007 | 0.005 | 0.011 |

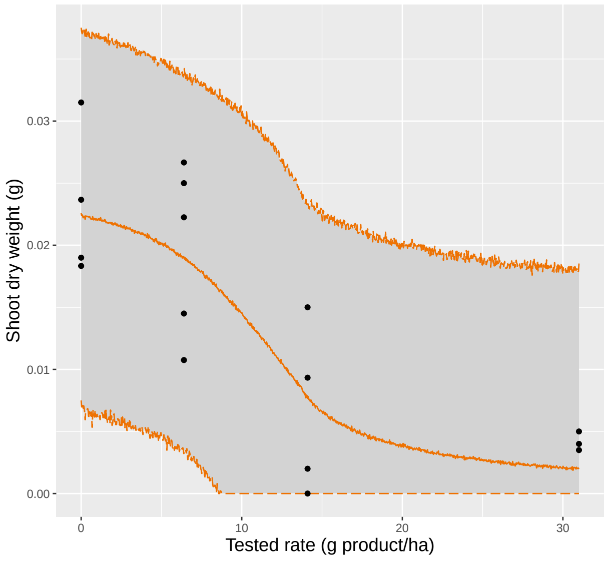

(a) Dose-response curve

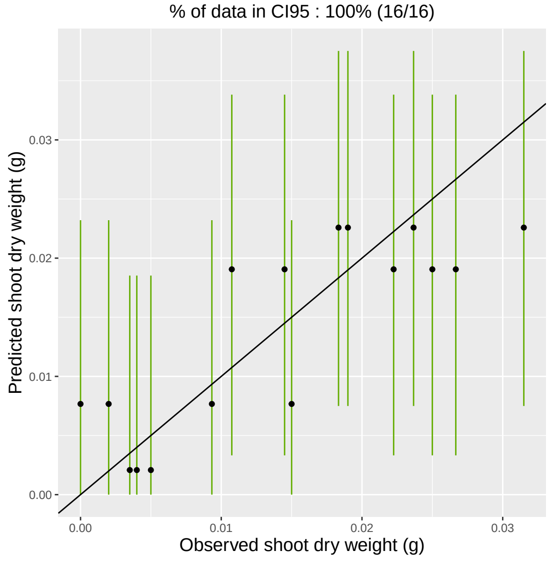

(b) Posterior predictive check (PPC)

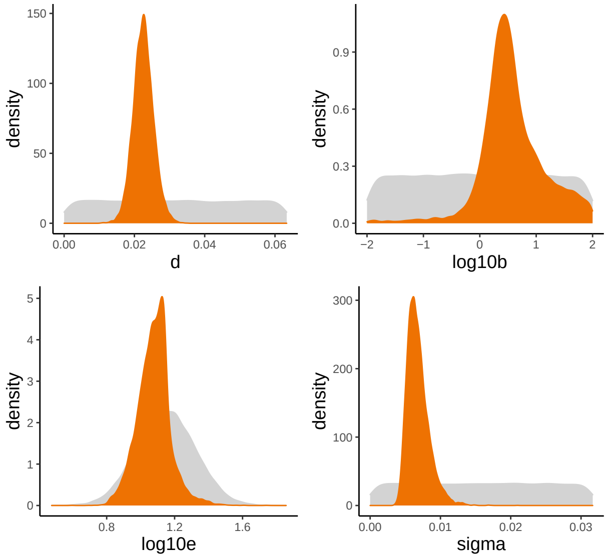

(c) Priors and posteriors

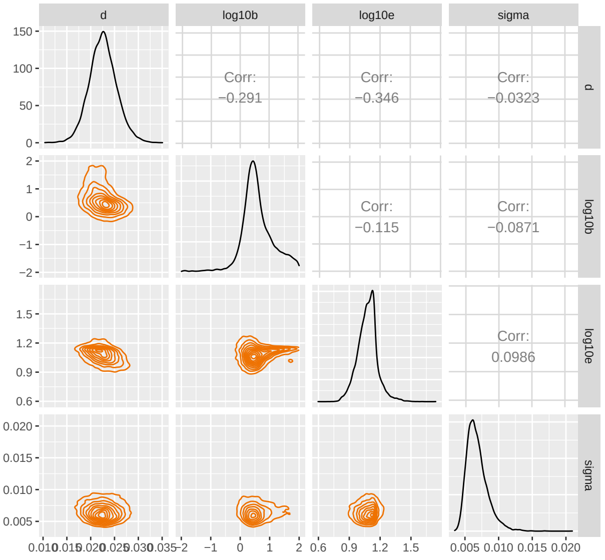

(d) Correlations between parameters

Figure 1: Dose-response curve (a), PPC (b), prior and posterior distributions (c) and correlations between parameters (d).

#### Data set: AVESA_SE_weight

Table 2: Summary of parameter estimates for AVESA_SE_weight data set

| Parameter | median | Q2.5 | Q97.5 |
|---|---|---|---|
| b | 2.114 | 1.579 | 3.046 |
| d | 0.476 | 0.438 | 0.516 |
| e | 25.207 | 21.098 | 29.880 |
| sigma | 0.048 | 0.037 | 0.065 |

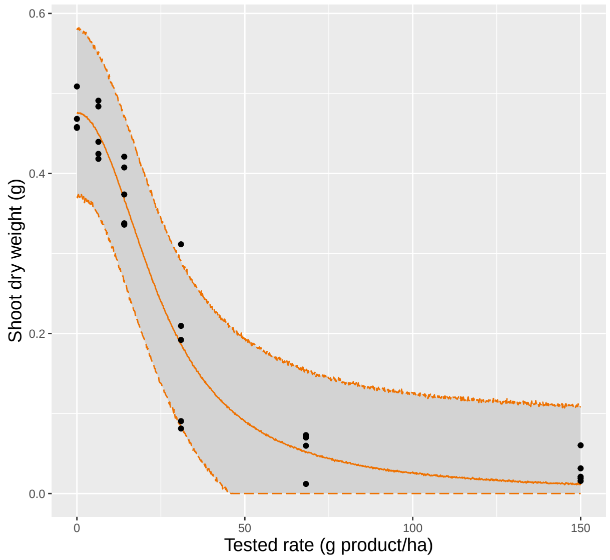

(a) Dose-response curve

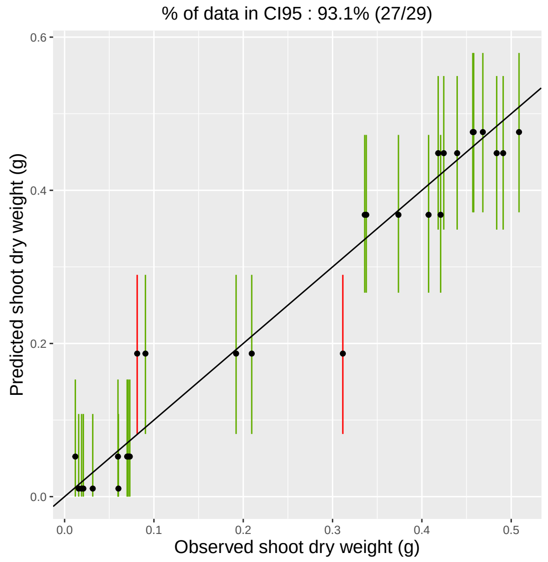

(b) Posterior predictive check (PPC)

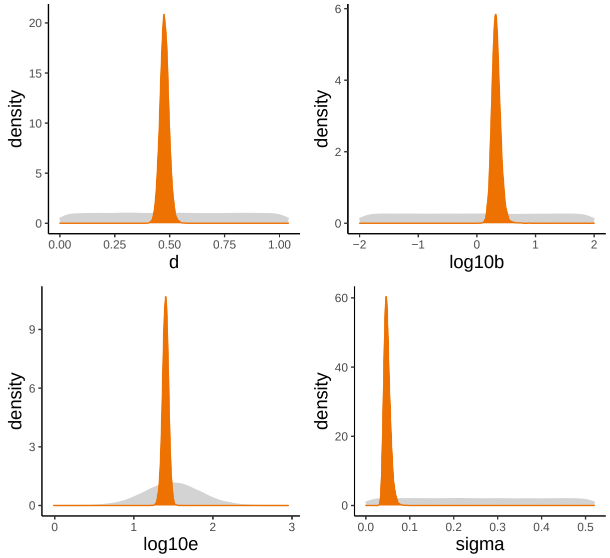

(c) Priors and posteriors

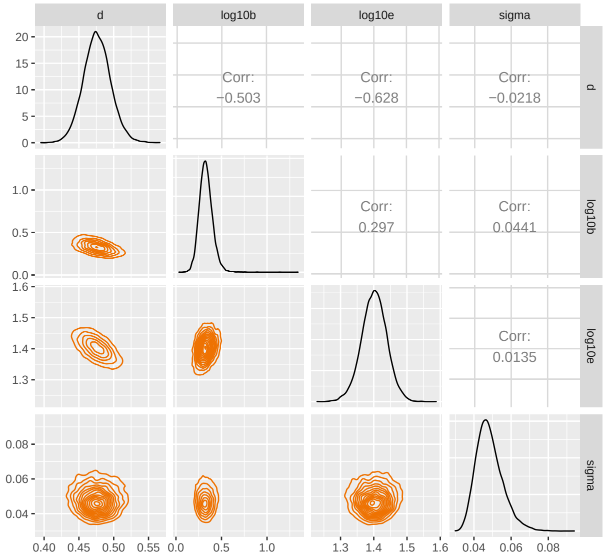

(d) Correlations between parameters

Figure 2: Dose-response curve (a), PPC (b), prior and posterior distributions (c) and correlations between parameters (d).

#### Data set: BEAVA_SE_weight

Table 3: Summary of parameter estimates for BEAVA_SE_weight data set

| Parameter | median | Q2.5 | Q97.5 |
|---|---|---|---|
| b | 2.401 | 1.807 | 3.324 |
| d | 0.574 | 0.529 | 0.619 |
| e | 11.054 | 9.637 | 12.722 |
| sigma | 0.068 | 0.055 | 0.088 |

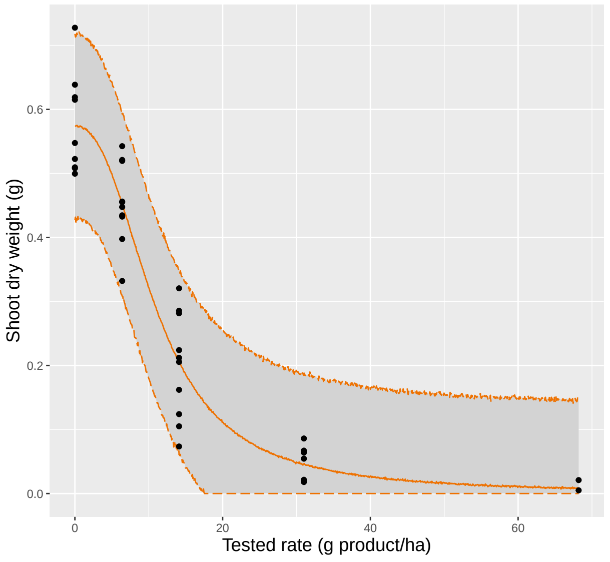

(a) Dose-response curve

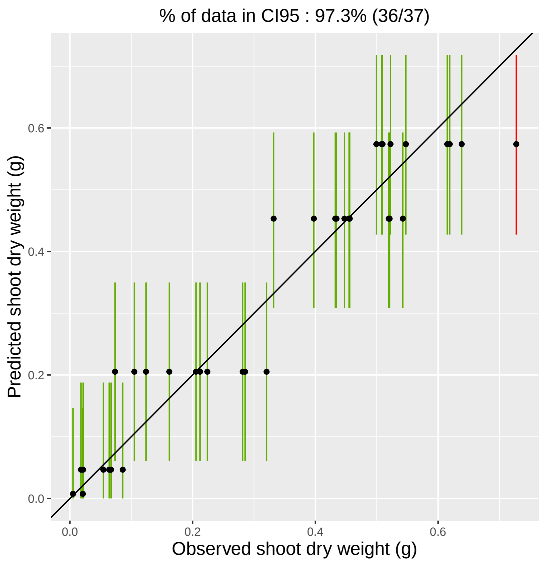

(b) Posterior predictive check (PPC)

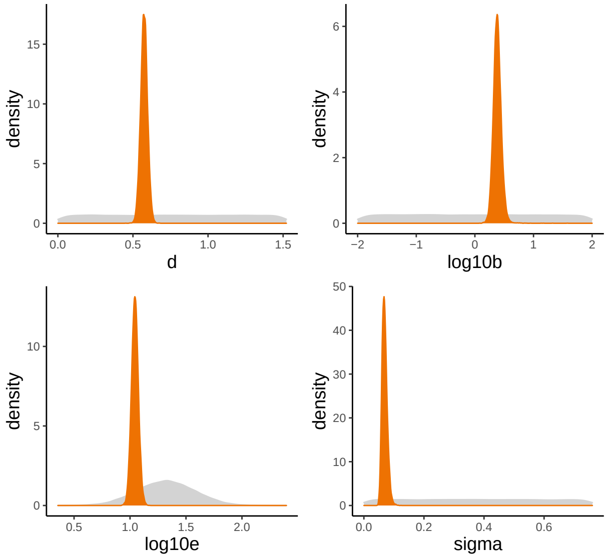

(c) Priors and posteriors

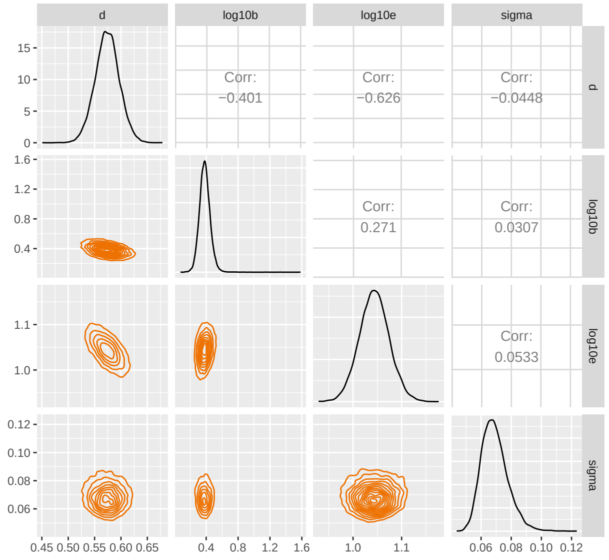

(d) Correlations between parameters

Figure 3: Dose-response curve (a), PPC (b), prior and posterior distributions (c) and correlations between parameters (d).

### Data set: BRSNW_SE_weight

Table 4: Summary of parameter estimates for BRSNW_SE_weight data set

| Parameter | median | Q2.5 | Q97.5 |
|---|---|---|---|
| b | 2.937 | 1.786 | 7.862 |
| d | 0.775 | 0.682 | 0.871 |
| e | 21.521 | 16.761 | 28.252 |
| sigma | 0.183 | 0.149 | 0.233 |

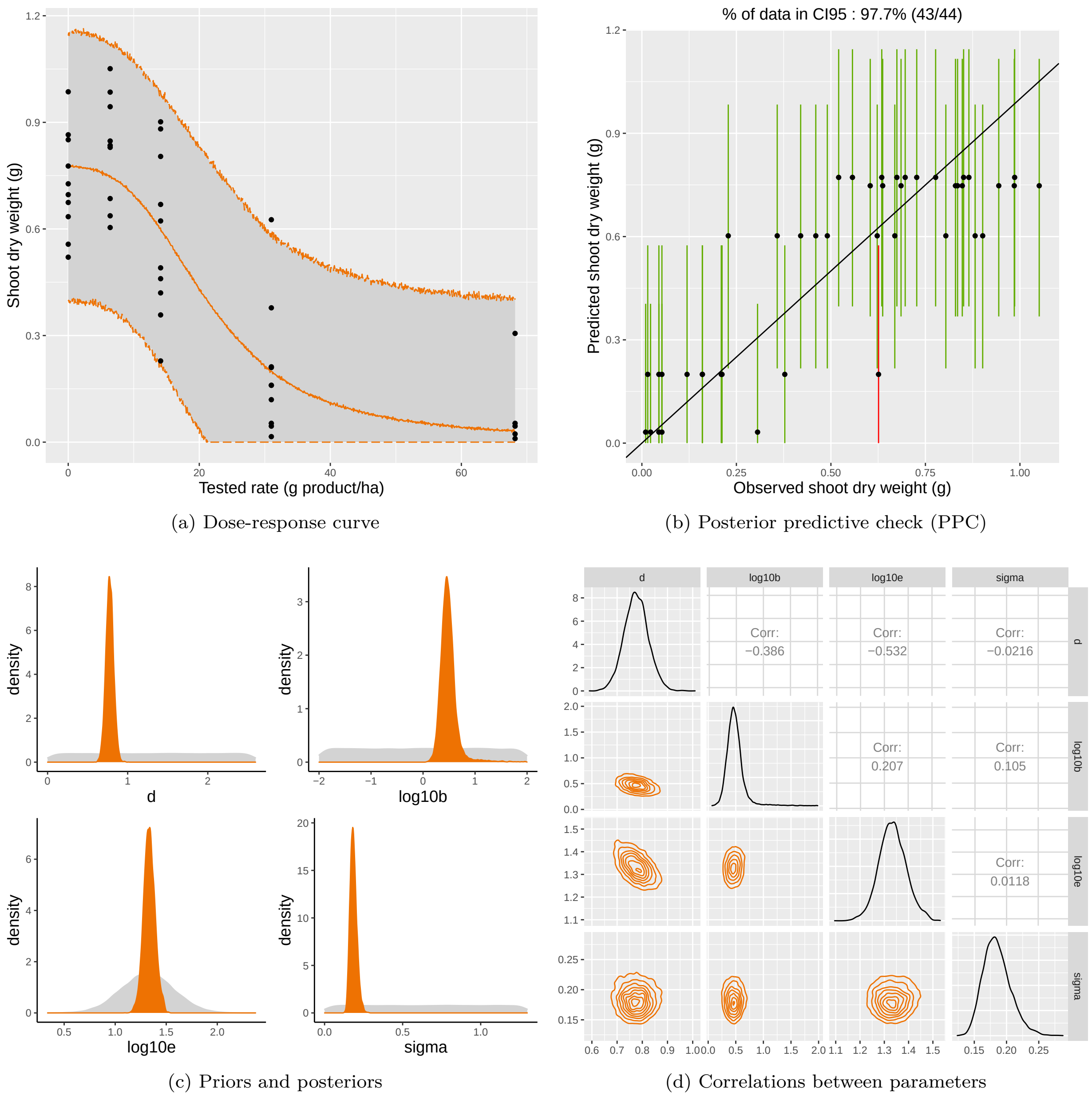

(a) Dose-response curve

(b) Posterior predictive check (PPC)

(c) Priors and posteriors

(d) Correlations between parameters

Figure 4: Dose-response curve (a), PPC (b), prior and posterior distributions (c) and correlations between parameters (d).

#### Data set: CUMSA_SE_weight

Table 5: Summary of parameter estimates for CUMSA_SE_weight data set

| Parameter | median | Q2.5 | Q97.5 |
|---|---|---|---|
| b | 1.061 | 0.780 | 1.445 |
| d | 1.897 | 1.700 | 2.097 |
| e | 28.643 | 20.359 | 40.758 |
| sigma | 0.336 | 0.281 | 0.410 |

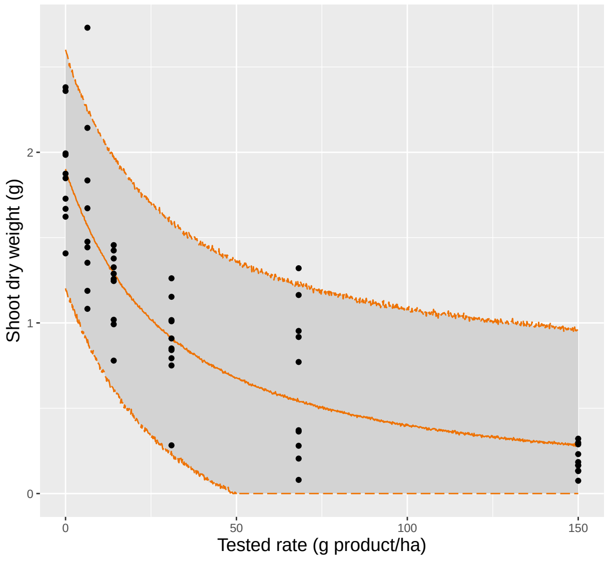

(a) Dose-response curve

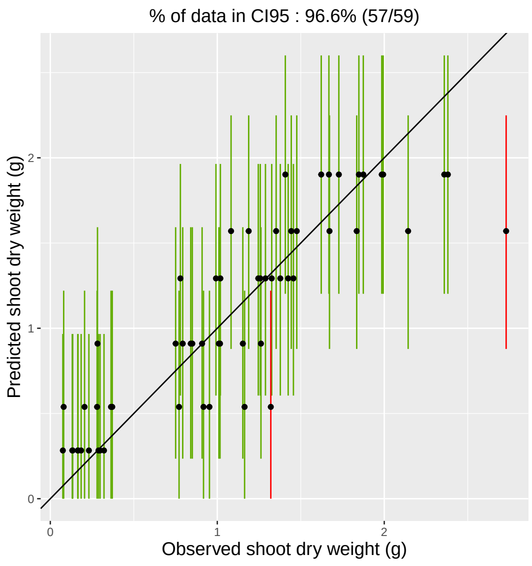

(b) Posterior predictive check (PPC)

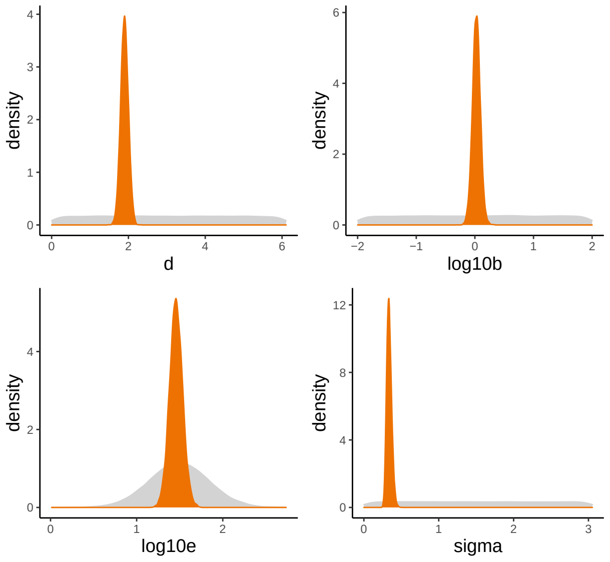

(c) Priors and posteriors

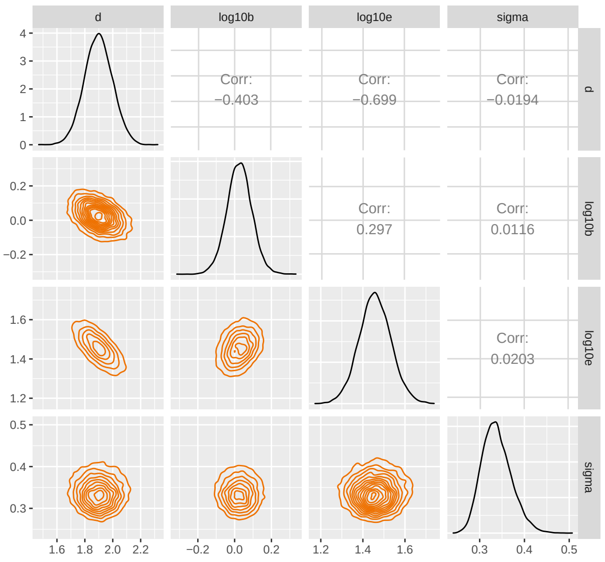

(d) Correlations between parameters

Figure 5: Dose-response curve (a), PPC (b), prior and posterior distributions (c) and correlations between parameters (d).

### Data set: GLXMA_SE_weight

Table 6: Summary of parameter estimates for GLXMA_SE_weight data set

| Parameter | median | Q2.5 | Q97.5 |
|---|---|---|---|
| b | 2.901 | 1.006 | 25.938 |
| d | 1.211 | 1.136 | 1.333 |
| e | 145.290 | 121.453 | 176.902 |
| sigma | 0.214 | 0.179 | 0.260 |

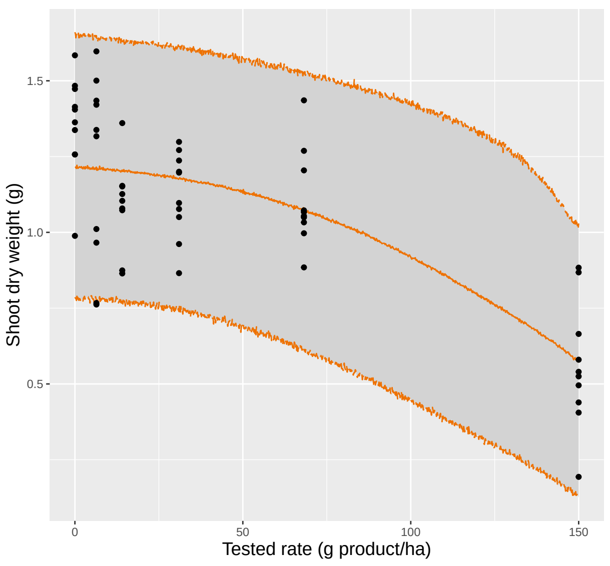

(a) Dose-response curve

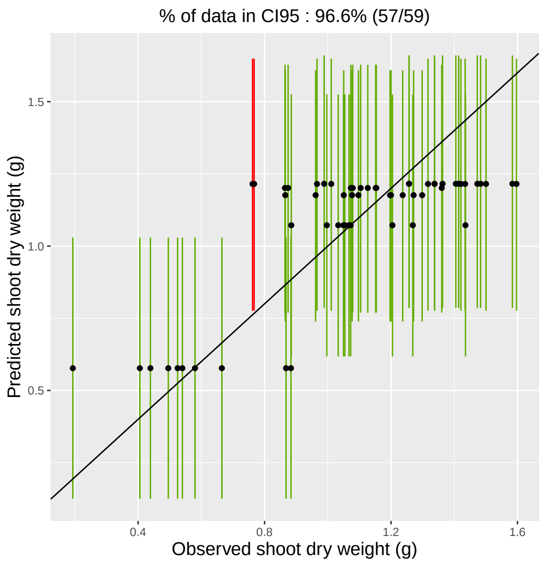

(b) Posterior predictive check (PPC)

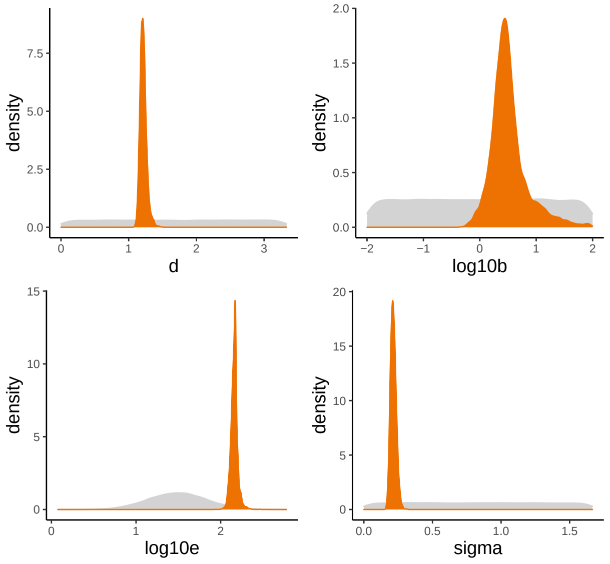

(c) Priors and posteriors

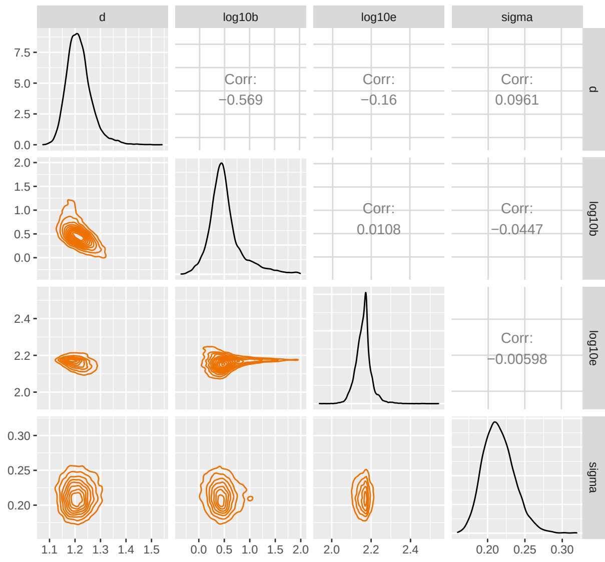

(d) Correlations between parameters

Figure 6: Dose-response curve (a), PPC (b), prior and posterior distributions (c) and correlations between parameters (d).

#### Data set: HELAN_SE_weight

Table 7: Summary of parameter estimates for HELAN_SE_weight data set

| Parameter | median | Q2.5 | Q97.5 |
|---|---|---|---|
| b | 2.423 | 1.520 | 4.328 |
| d | 0.426 | 0.389 | 0.466 |
| e | 48.604 | 38.692 | 60.364 |
| sigma | 0.085 | 0.071 | 0.105 |

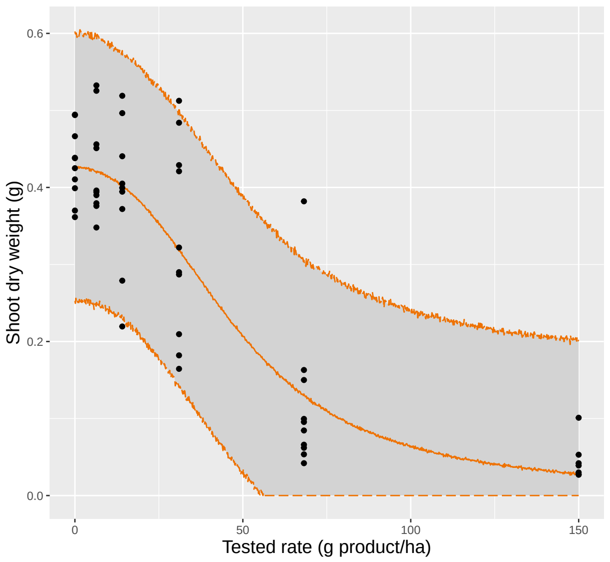

(a) Dose-response curve

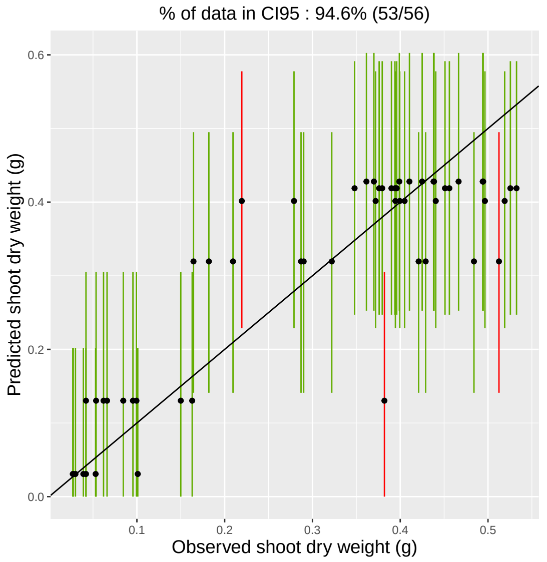

(b) Posterior predictive check (PPC)

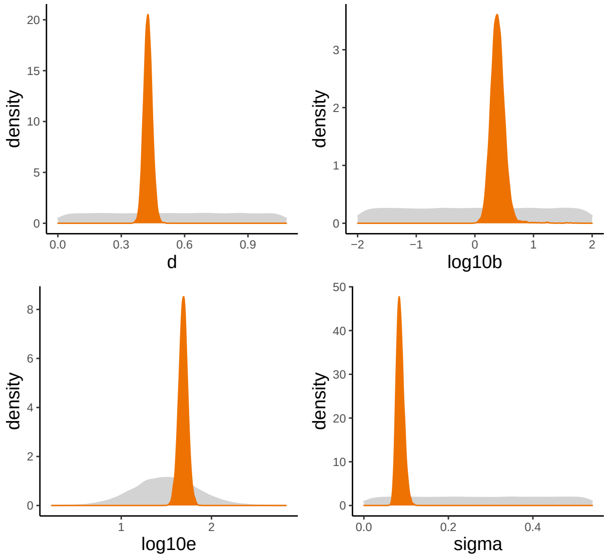

(c) Priors and posteriors

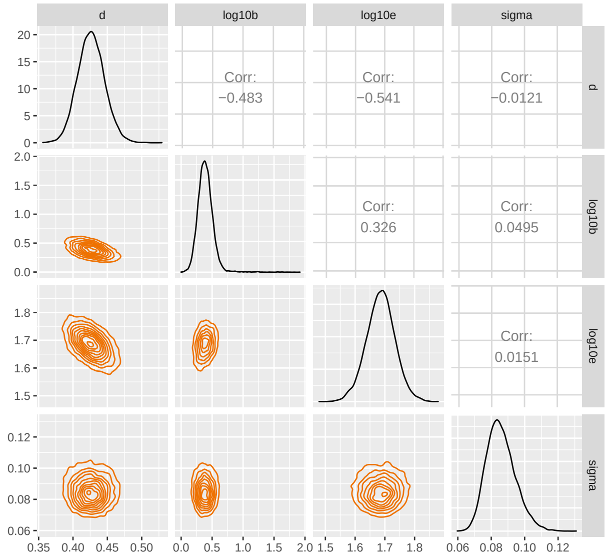

(d) Correlations between parameters

Figure 7: Dose-response curve (a), PPC (b), prior and posterior distributions (c) and correlations between parameters (d).

#### Data set: LYPES_SE_weight

Table 8: Summary of parameter estimates for LYPES_SE_weight data set

| Parameter | median | Q2.5 | Q97.5 |
|---|---|---|---|
| b | 2.819 | 2.014 | 4.190 |
| d | 0.738 | 0.678 | 0.803 |
| e | 21.317 | 18.196 | 25.120 |
| sigma | 0.114 | 0.095 | 0.141 |

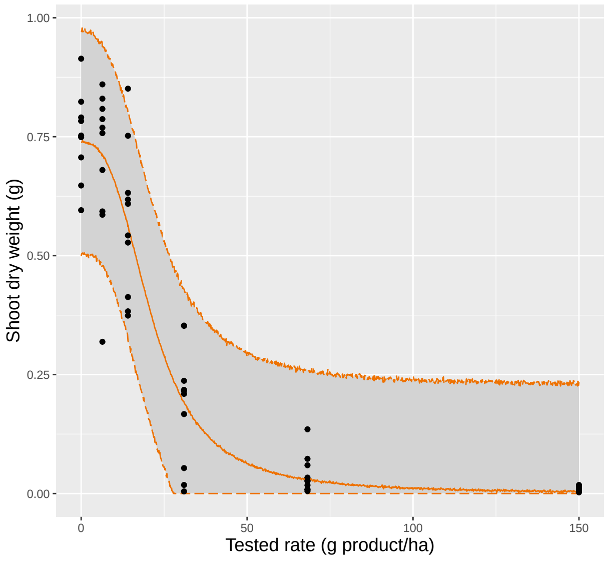

(a) Dose-response curve

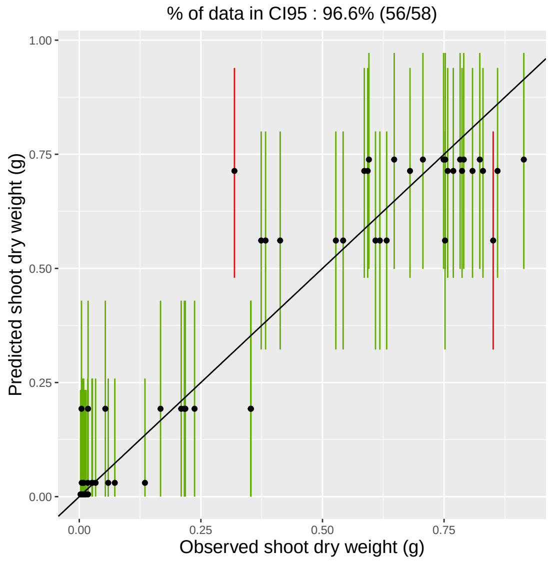

(b) Posterior predictive check (PPC)

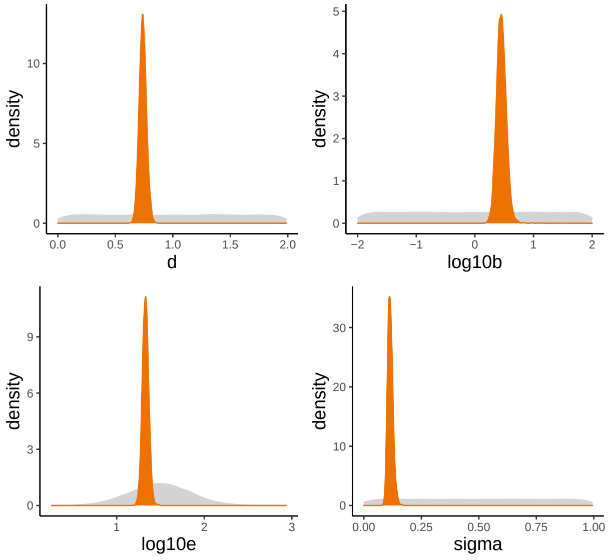

(c) Priors and posteriors

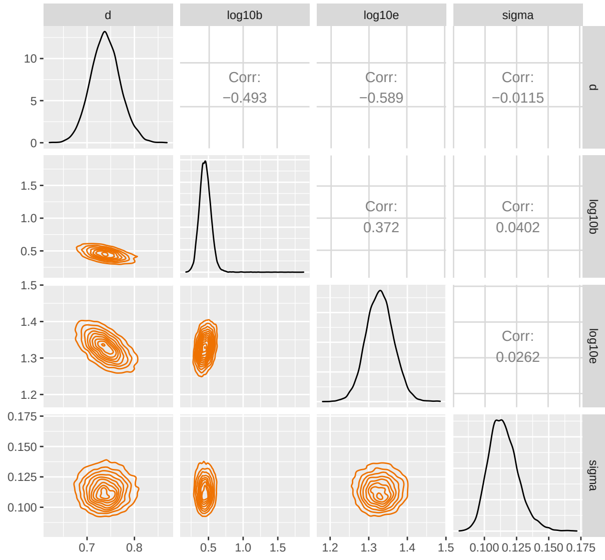

(d) Correlations between parameters

Figure 8: Dose-response curve (a), PPC (b), prior and posterior distributions (c) and correlations between parameters (d).

#### Data set: TRZAW_SE_weight

Table 9: Summary of parameter estimates for TRZAW_SE_weight data set

| Parameter | median | Q2.5 | Q97.5 |
|---|---|---|---|
| b | 1.777 | 0.992 | 4.966 |
| d | 0.331 | 0.310 | 0.356 |
| e | 167.744 | 138.635 | 229.056 |
| sigma | 0.035 | 0.027 | 0.047 |

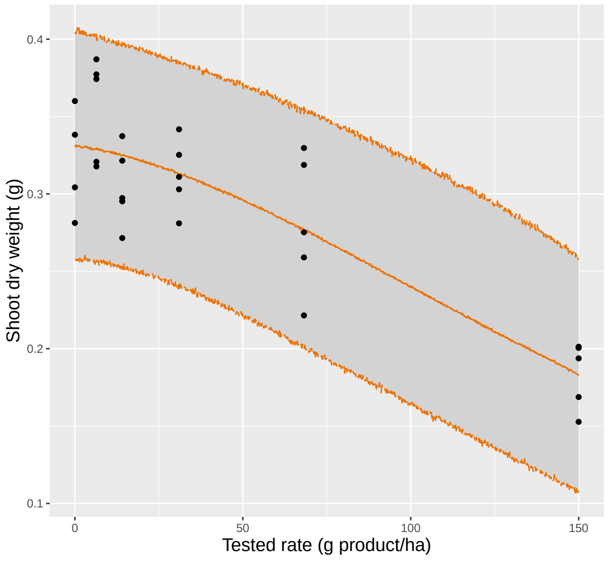

(a) Dose-response curve

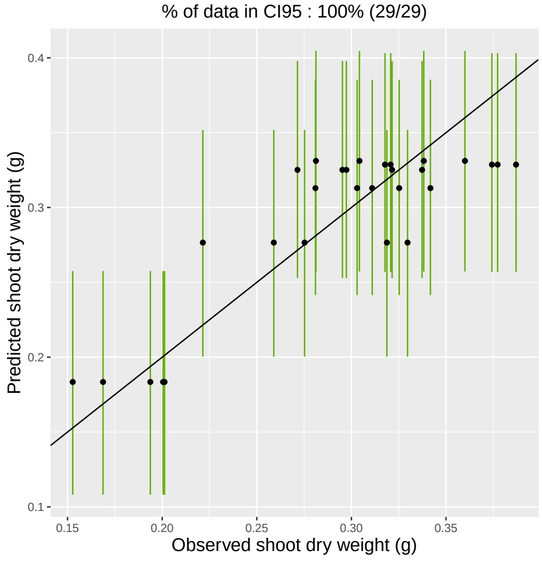

(b) Posterior predictive check (PPC)

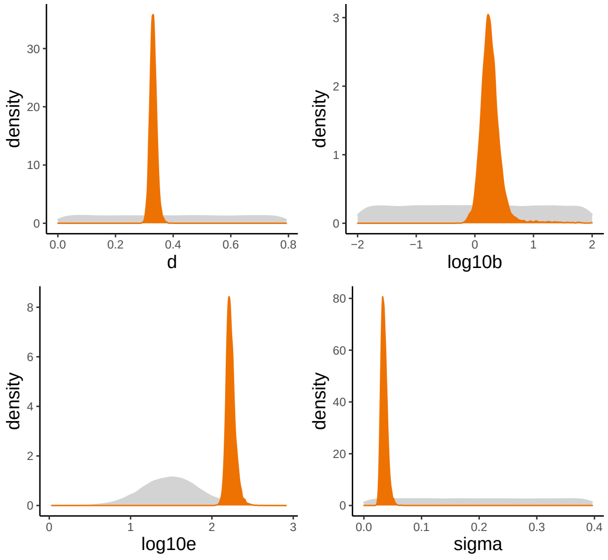

(c) Priors and posteriors

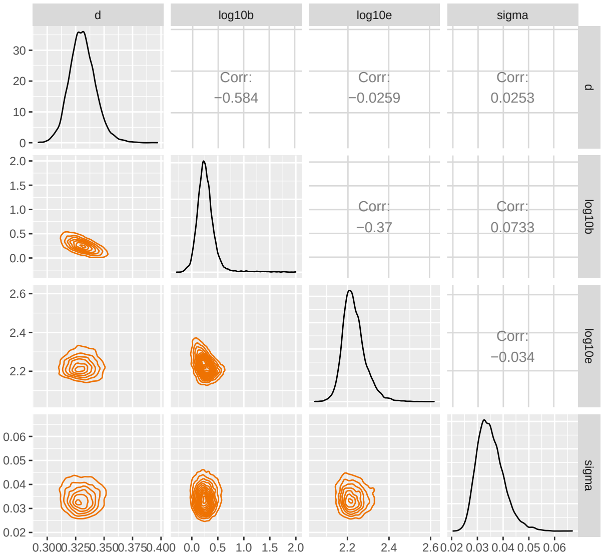

(d) Correlations between parameters

Figure 9: Dose-response curve (a), PPC (b), prior and posterior distributions (c) and correlations between parameters (d).

#### Data set: ZEAMA_SE_weight

Table 10: Summary of parameter estimates for ZEAMA_SE_weight data set

| Parameter | median | Q2.5 | Q97.5 |
|---|---|---|---|
| b | 2.072 | 1.051 | 21.372 |
| d | 1.876 | 1.762 | 2.011 |
| e | 194.498 | 153.445 | 300.683 |
| sigma | 0.316 | 0.265 | 0.387 |

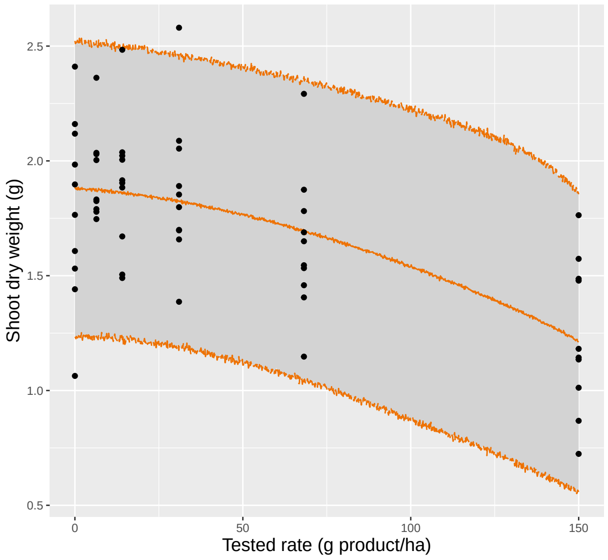

(a) Dose-response curve

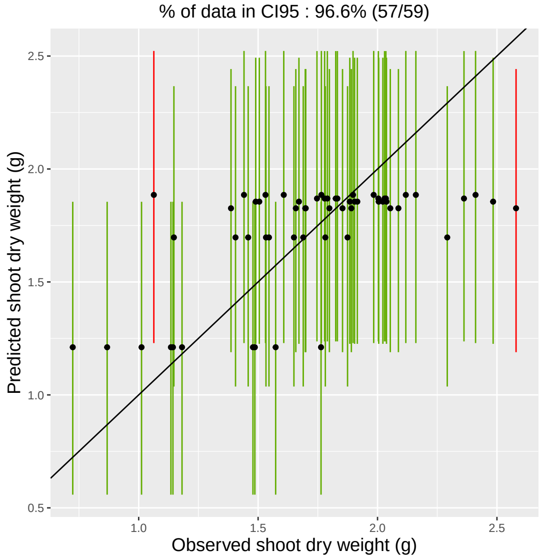

(b) Posterior predictive check (PPC)

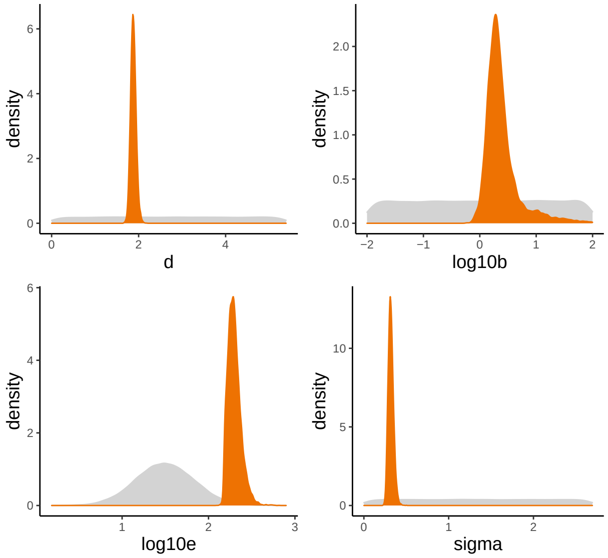

(c) Priors and posteriors

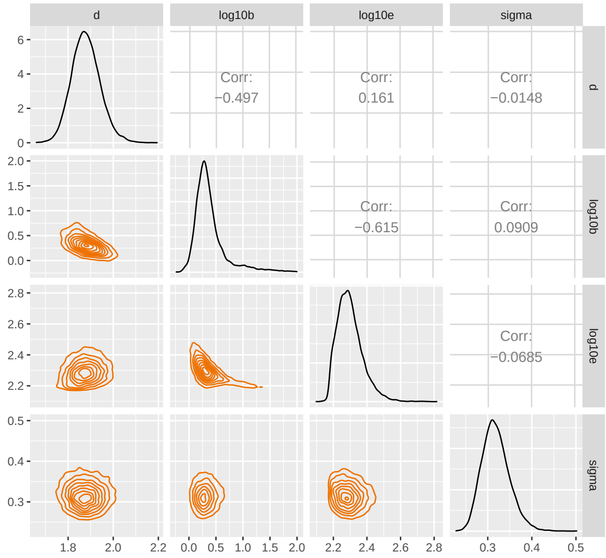

(d) Correlations between parameters

Figure 10: Dose-response curve (a), PPC (b), prior and posterior distributions (c) and correlations between parameters (d).
