## Supplementary material for "How to account for the uncertainty from standard toxicity tests in species sensitivity distributions: an example in non-target plants": Archive with reports on studies: report_VV_survival.pdf

### Dose-response analyses

Study 6

Vegetative Vigour test - survival endpoint

25 June 2020

Contact:

---

This is a report which provides results on all performed dose-response analyses for the survival endpoint of the Vegetative Vigour test for study 6.

---

#### Data set: ALLCE_VV_survival

Table 1: Summary of parameter estimates (parameter d is set to 1) for ALLCE_VV_survival data set

| Parameter | median | Q2.5 | Q97.5 |
|---|---|---|---|
| b | 8.843 | 2.337 | 68.140 |
| e | 81.658 | 69.736 | 145.767 |

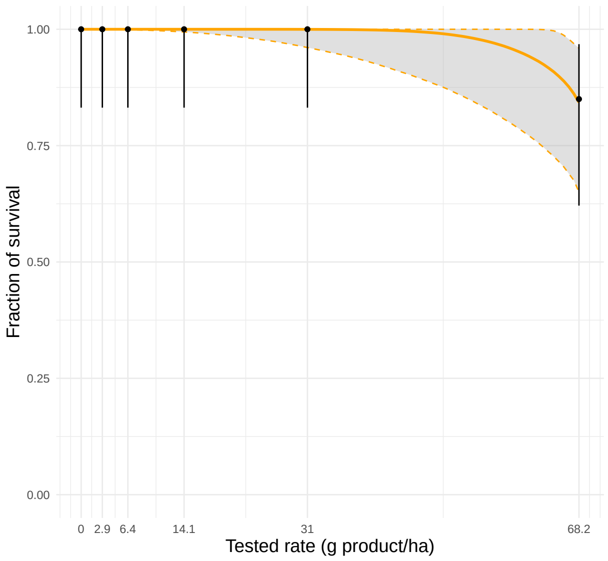

(a) Dose-response curve

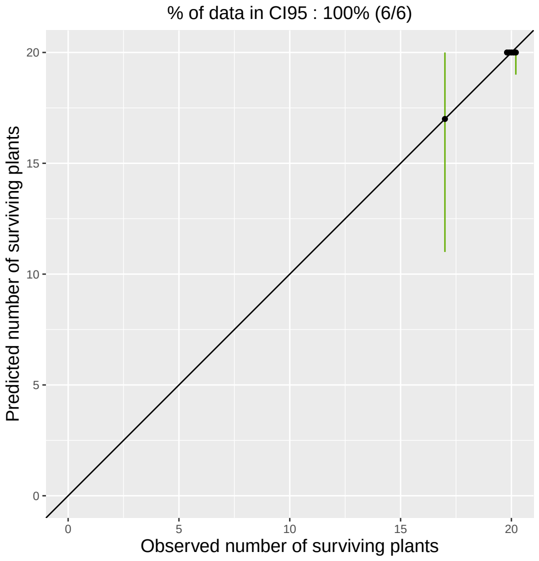

(b) Posterior predictive check (PPC)

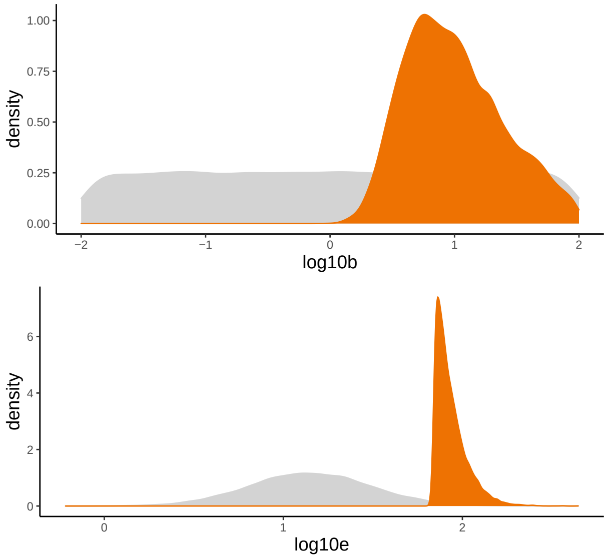

(c) Priors and posteriors

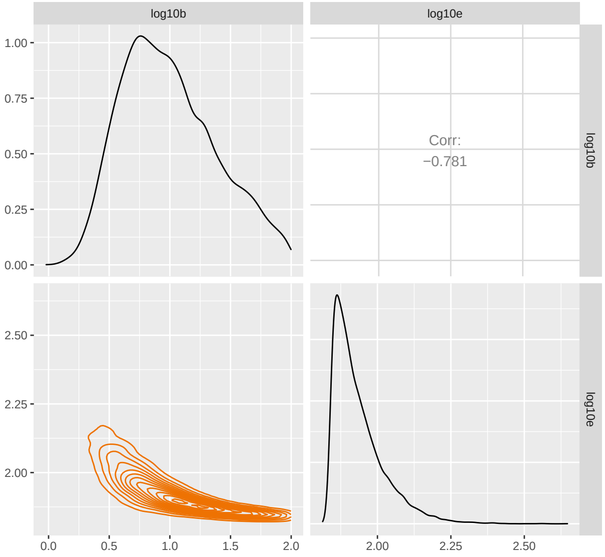

(d) Correlations between parameters

Figure 1: Dose-response curve (a), PPC (b), prior and posterior distributions (c) and correlations between parameters (d).

#### Data set: AVESA_VV_survival

Table 2: Summary of parameter estimates (parameter d is set to 1) for AVESA_VV_survival data set

| Parameter | median | Q2.5 | Q97.5 |
|---|---|---|---|
| b | 40.751 | 10.637 | 95.913 |
| e | 47.289 | 34.017 | 62.899 |

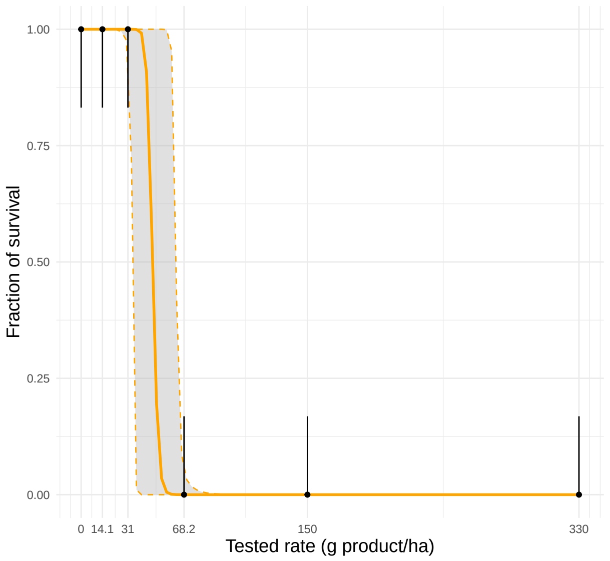

(a) Dose-response curve

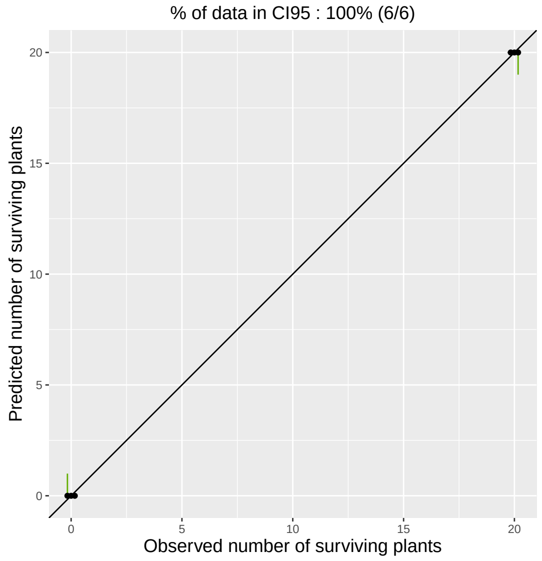

(b) Posterior predictive check (PPC)

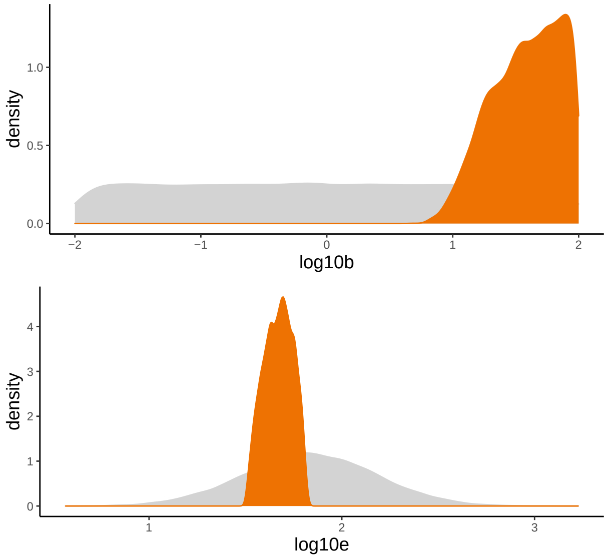

(c) Priors and posteriors

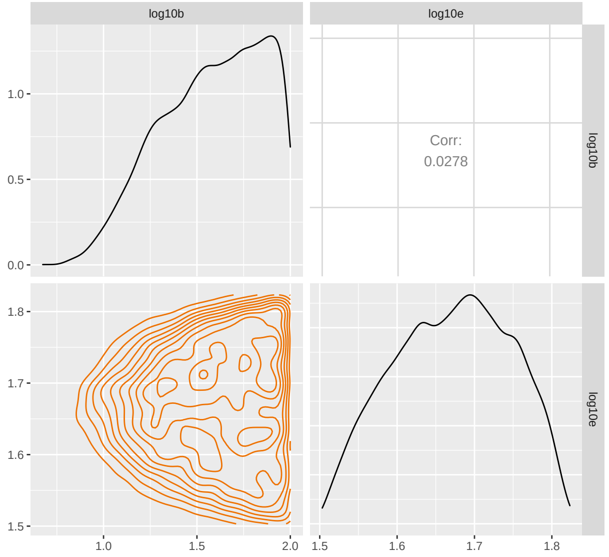

(d) Correlations between parameters

Figure 2: Dose-response curve (a), PPC (b), prior and posterior distributions (c) and correlations between parameters (d).

#### Data set: BEAVA_VV_survival

Table 3: Summary of parameter estimates (parameter d is set to 1) for BEAVA_VV_survival data set

| Parameter | median | Q2.5 | Q97.5 |
|---|---|---|---|
| b | 2.396 | 1.371 | 3.934 |
| e | 62.622 | 47.839 | 92.698 |

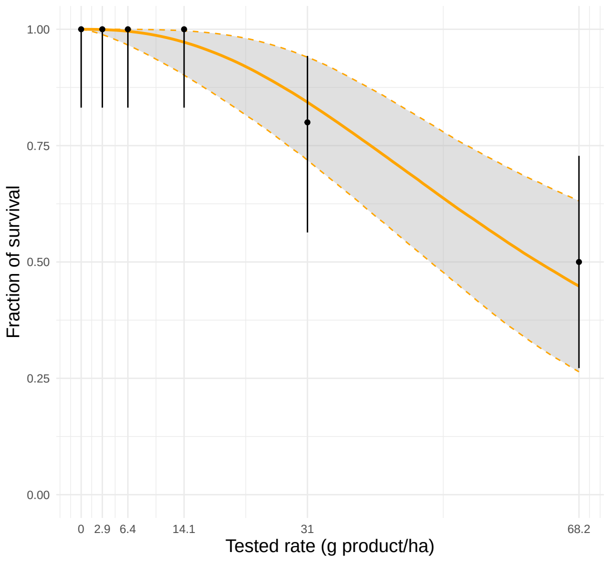

(a) Dose-response curve

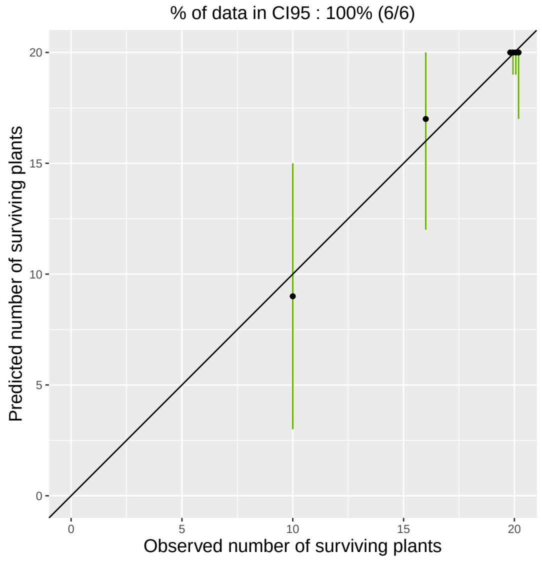

(b) Posterior predictive check (PPC)

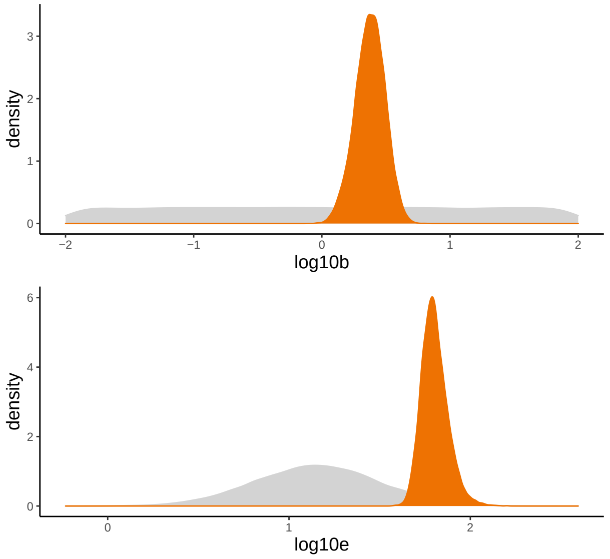

(c) Priors and posteriors

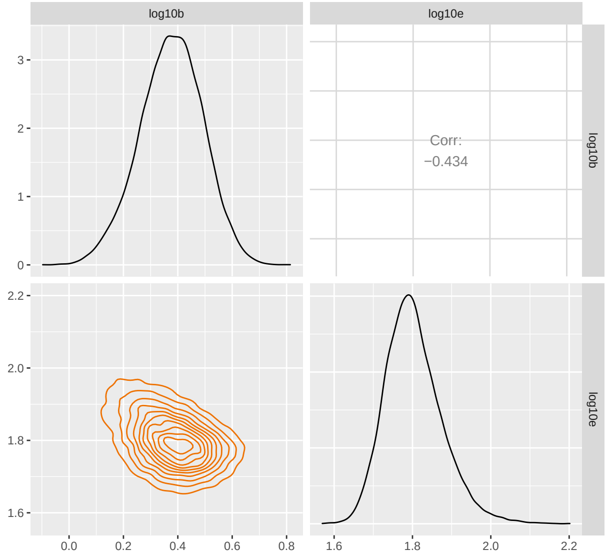

(d) Correlations between parameters

Figure 3: Dose-response curve (a), PPC (b), prior and posterior distributions (c) and correlations between parameters (d).

#### Data set: BRSNW_VV_survival

Table 4: Summary of parameter estimates (parameter d is set to 1) for BRSNW_VV_survival data set

| Parameter | median | Q2.5 | Q97.5 |
|---|---|---|---|
| b | 5.508 | 3.285 | 8.600 |
| e | 45.342 | 37.802 | 54.413 |

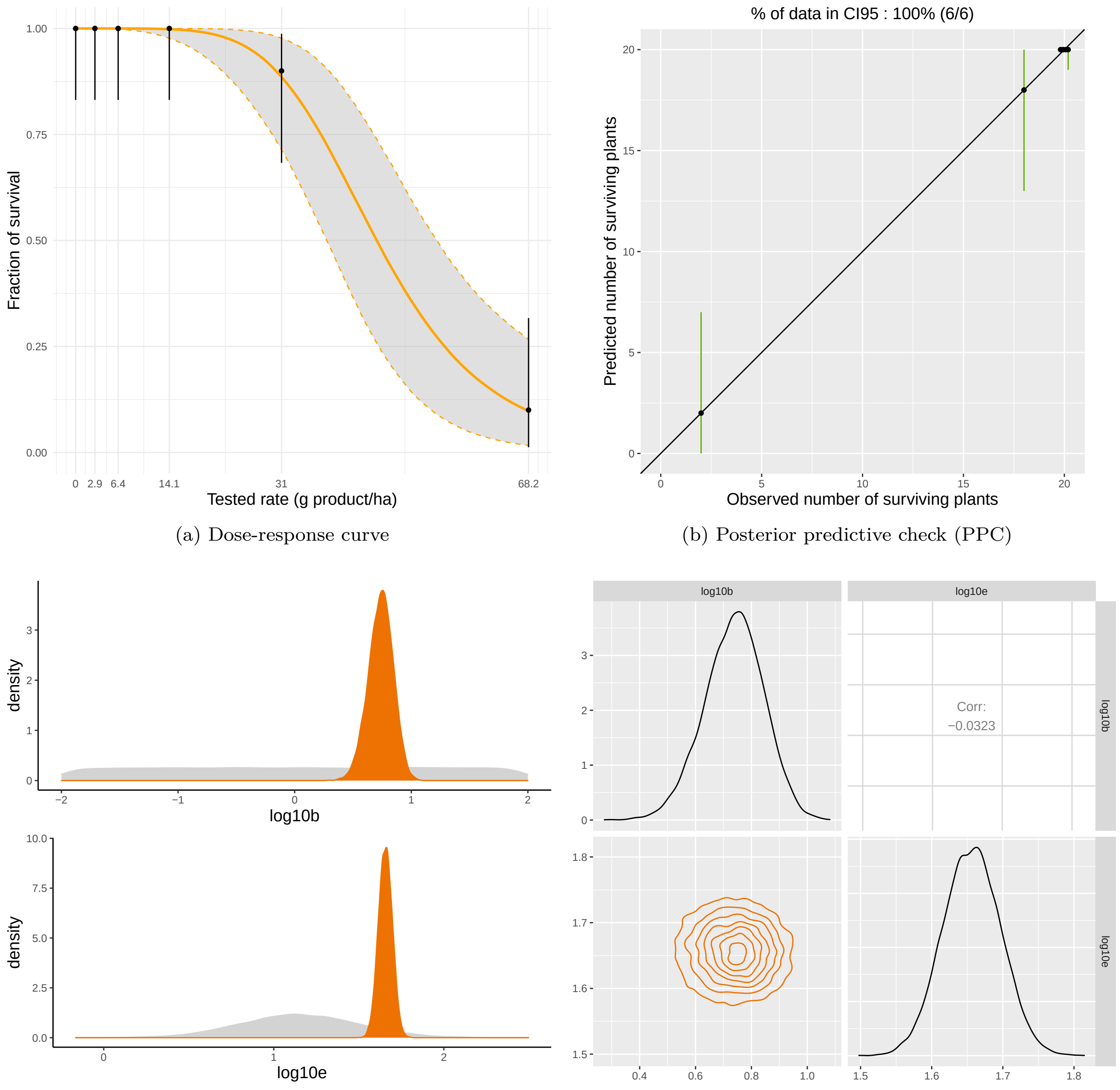

(a) Dose-response curve

(b) Posterior predictive check (PPC)

(c) Priors and posteriors

(d) Correlations between parameters

Figure 4: Dose-response curve (a), PPC (b), prior and posterior distributions (c) and correlations between parameters (d).

#### Data set: CUMSA_VV_survival

Table 5: Summary of parameter estimates (parameter d is set to 1) for CUMSA_VV_survival data set

| Parameter | median | Q2.5 | Q97.5 |
|---|---|---|---|
| b | 34.728 | 5.204 | 95.197 |
| e | 98.268 | 73.161 | 214.722 |

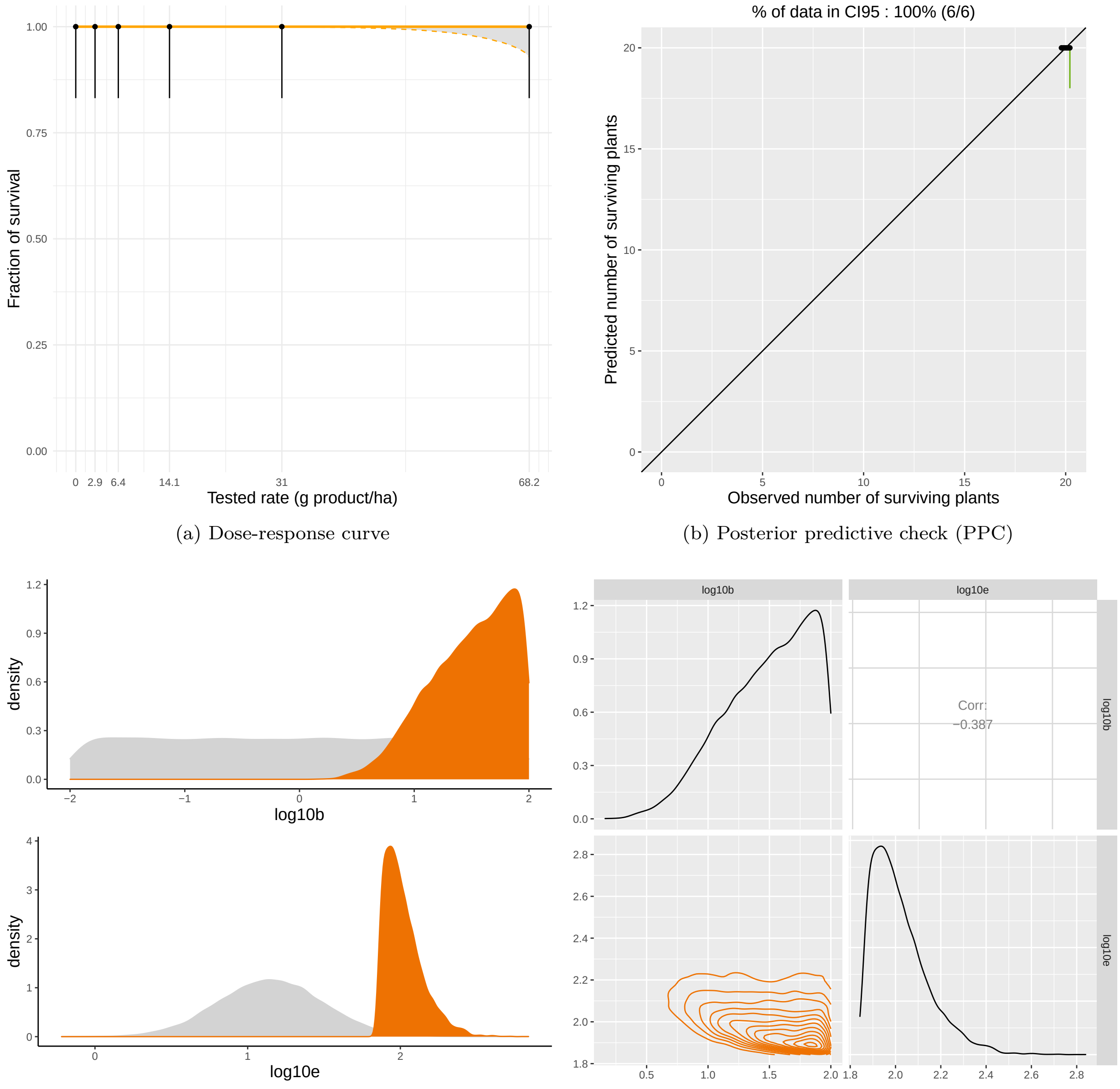

(a) Dose-response curve

(b) Posterior predictive check (PPC)

(c) Priors and posteriors

(d) Correlations between parameters

Figure 5: Dose-response curve (a), PPC (b), prior and posterior distributions (c) and correlations between parameters (d).

#### Data set: GLXMA_VV_survival

Table 6: Summary of parameter estimates (parameter d is set to 1) for GLXMA_VV_survival data set

| Parameter | median | Q2.5 | Q97.5 |
|---|---|---|---|
| b | 35.077 | 5.226 | 95.758 |
| e | 97.593 | 72.706 | 208.115 |

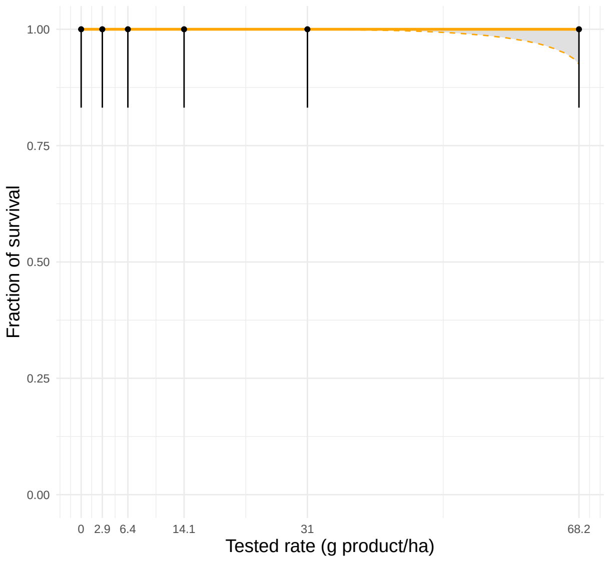

(a) Dose-response curve

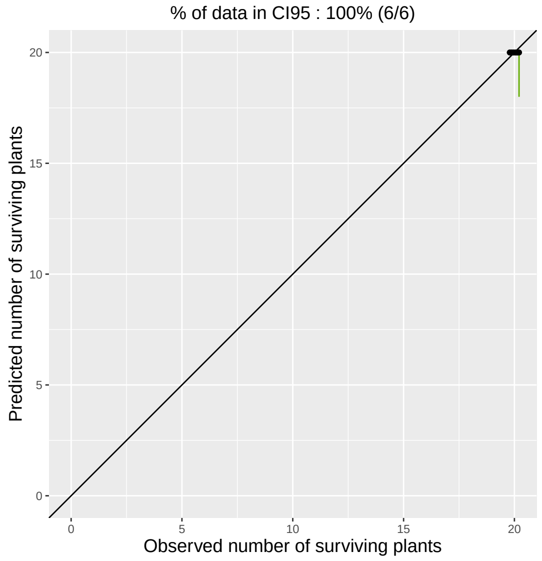

(b) Posterior predictive check (PPC)

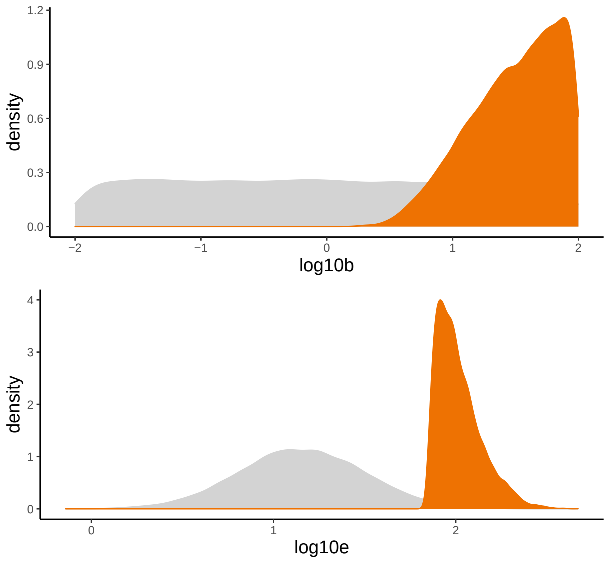

(c) Priors and posteriors

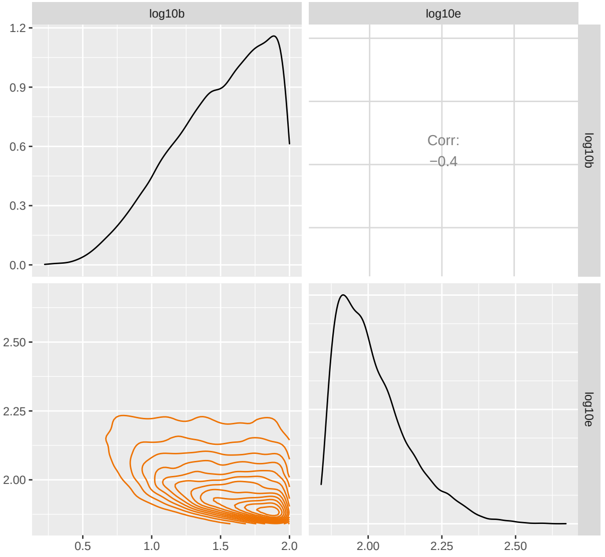

(d) Correlations between parameters

Figure 6: Dose-response curve (a), PPC (b), prior and posterior distributions (c) and correlations between parameters (d).

#### Data set: HELAN_VV_survival

Table 7: Summary of parameter estimates (parameter d is set to 1) for HELAN_VV_survival data set

| Parameter | median | Q2.5 | Q97.5 |
|---|---|---|---|
| b | 14.205 | 6.314 | 73.037 |
| e | 24.963 | 19.363 | 29.766 |

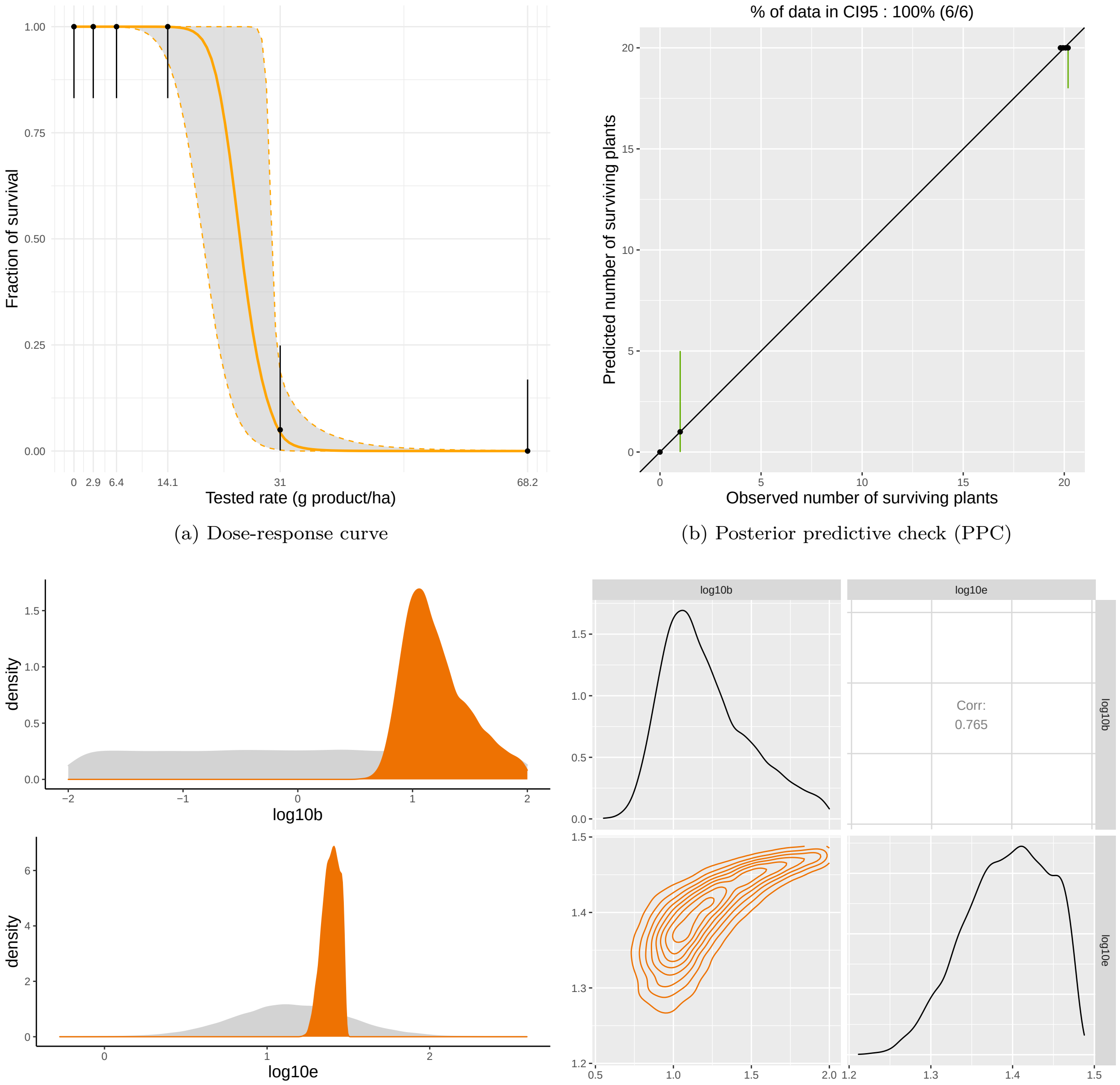

(a) Dose-response curve
(b) Posterior predictive check (PPC)
(c) Priors and posteriors
(d) Correlations between parameters

Figure 7: Dose-response curve (a), PPC (b), prior and posterior distributions (c) and correlations between parameters (d).

#### Data set: LYPES_VV_survival

Table 8: Summary of parameter estimates (parameter d is set to 1) for LYPES_VV_survival data set

| Parameter | median | Q2.5 | Q97.5 |
|---|---|---|---|
| b | 12.095 | 2.740 | 77.718 |
| e | 86.745 | 70.696 | 178.589 |

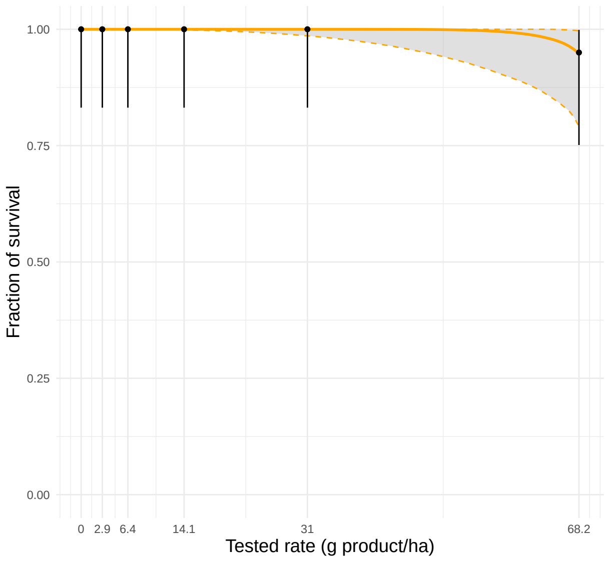

(a) Dose-response curve

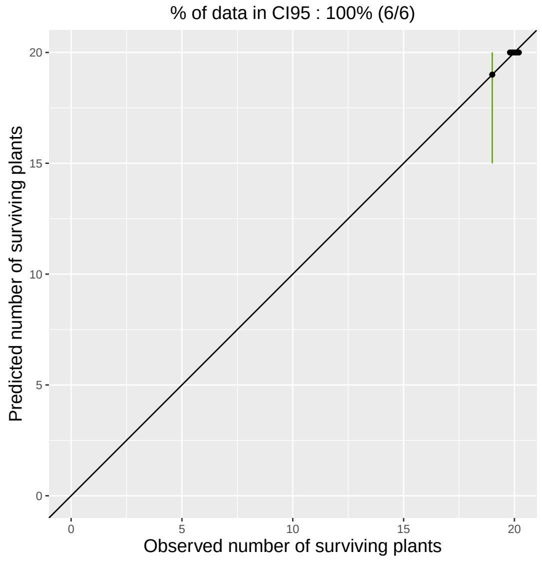

(b) Posterior predictive check (PPC)

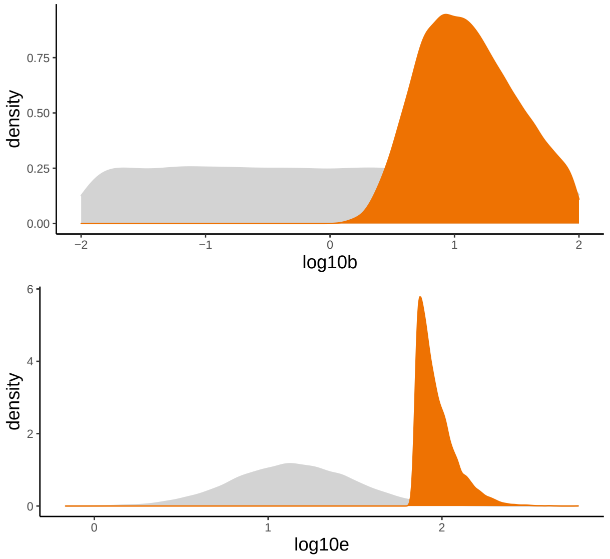

(c) Priors and posteriors

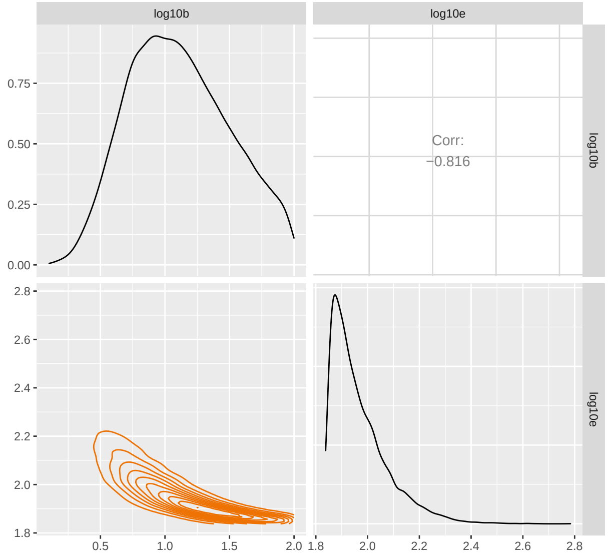

(d) Correlations between parameters

Figure 8: Dose-response curve (a), PPC (b), prior and posterior distributions (c) and correlations between parameters (d).

#### Data set: TRZAW_VV_survival

Table 9: Summary of parameter estimates (parameter d is set to 1) for TRZAW_VV_survival data set

| Parameter | median | Q2.5 | Q97.5 |
|---|---|---|---|
| b | 12.285 | 2.783 | 78.777 |
| e | 417.562 | 342.234 | 858.400 |

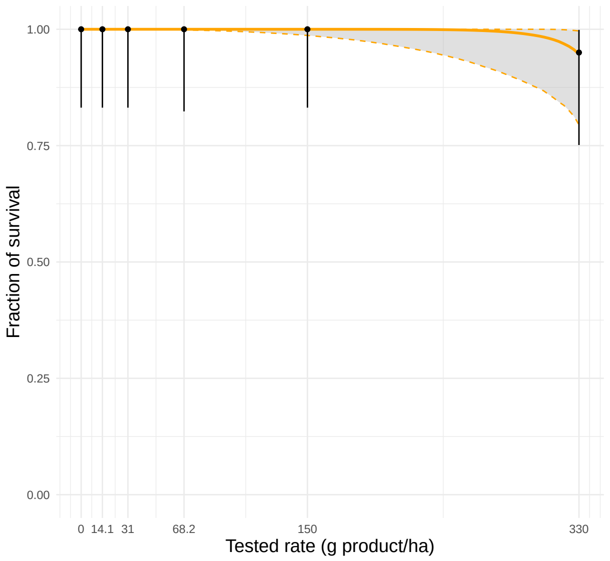

(a) Dose-response curve

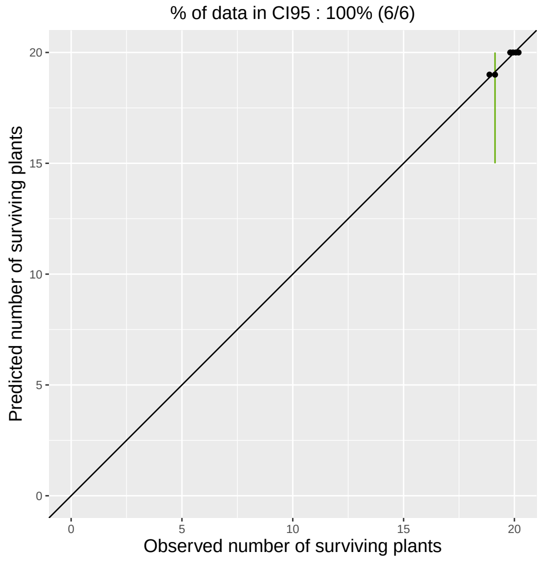

(b) Posterior predictive check (PPC)

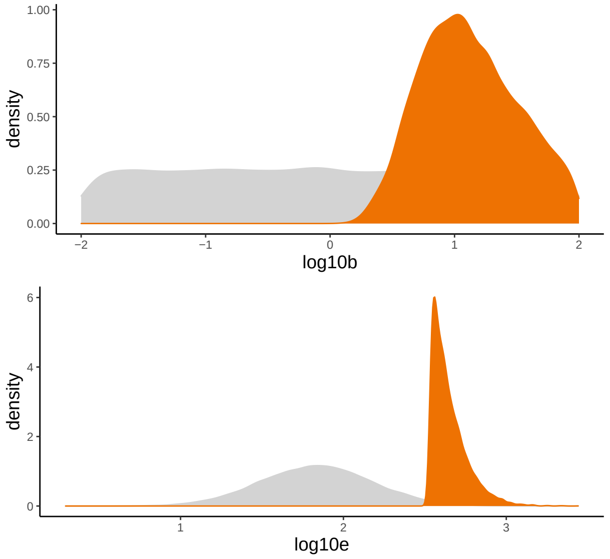

(c) Priors and posteriors

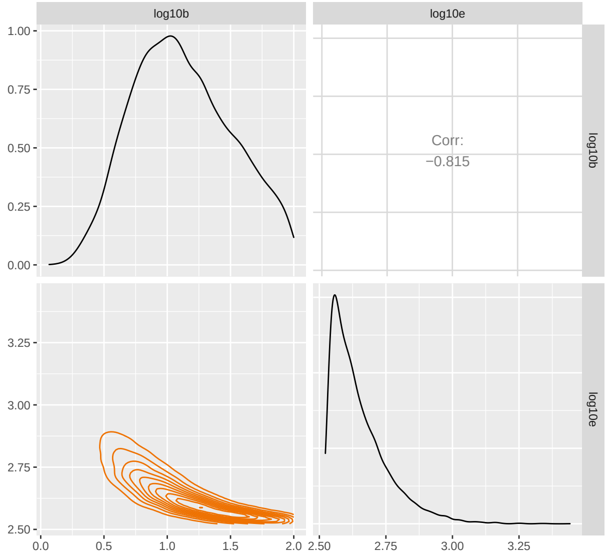

(d) Correlations between parameters

Figure 9: Dose-response curve (a), PPC (b), prior and posterior distributions (c) and correlations between parameters (d).

#### Data set: ZEAMA_VV_survival

Table 10: Summary of parameter estimates (parameter d is set to 1) for ZEAMA_VV_survival data set

| Parameter | median | Q2.5 | Q97.5 |
|---|---|---|---|
| b | 8.073 | 2.173 | 71.399 |
| e | 384.802 | 335.028 | 664.404 |

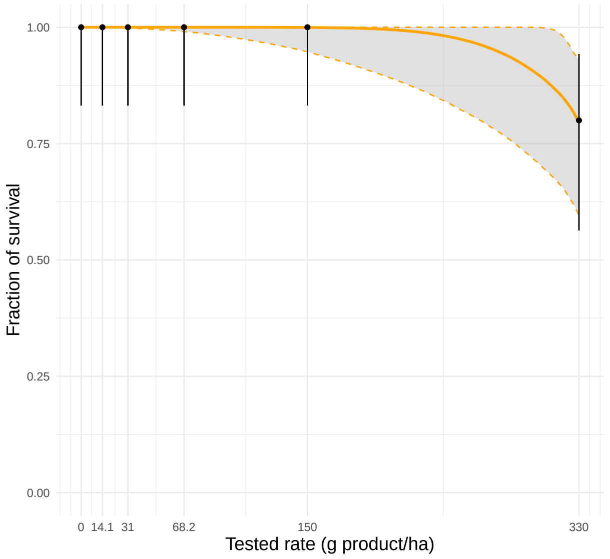

(a) Dose-response curve

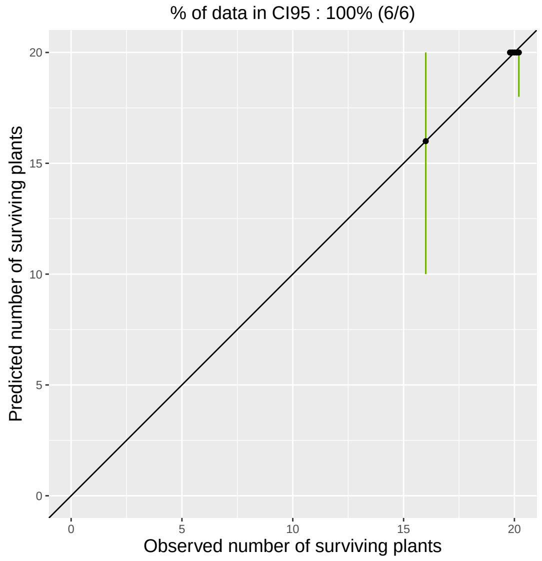

(b) Posterior predictive check (PPC)

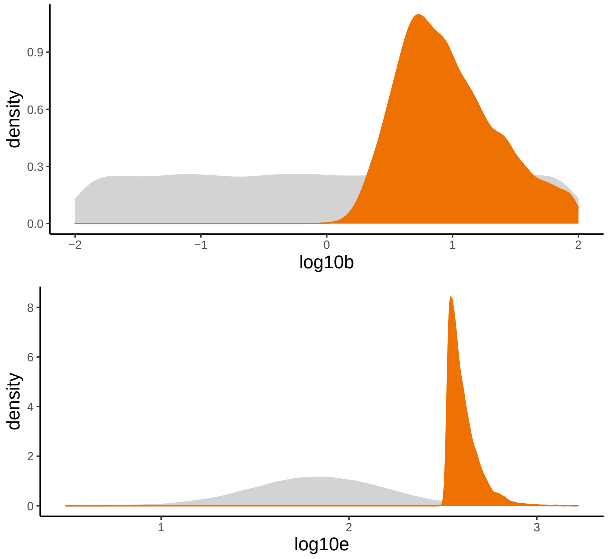

(c) Priors and posteriors

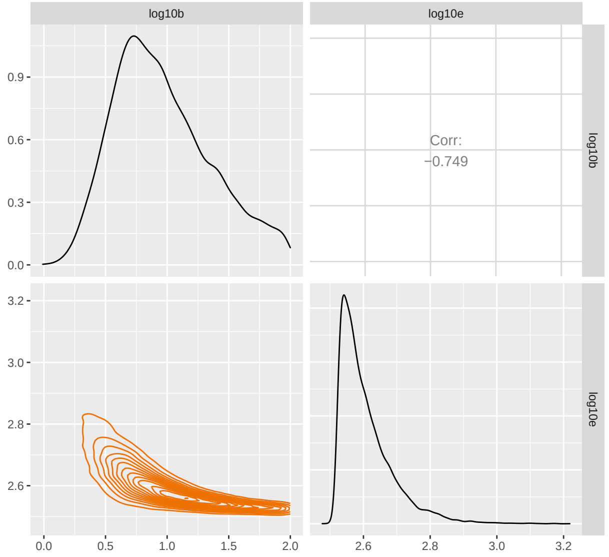

(d) Correlations between parameters

Figure 10: Dose-response curve (a), PPC (b), prior and posterior distributions (c) and correlations between parameters (d).
