## Supplementary material for "How to account for the uncertainty from standard toxicity tests in species sensitivity distributions: an example in non-target plants": Archive with reports on studies: report_VV_weight.pdf

### Dose-response analysis

#### Study 6

#### Vegetative Vigour test - shoot dry VV_weight endpoint

25 June 2020

Contact:

---

This is a report which provides results on all performed dose-response analyses for the shoot dry VV_weight endpoint of the Vegetative Vigour test for study 6.

---

#### Data set: ALLCE_VV_weight

Table 1: Summary of parameter estimates for ALLCE_VV_weight data set

| Parameter | median | Q2.5 | Q97.5 |
|---|---|---|---|
| b | 3.278 | 1.899 | 15.795 |
| d | 0.350 | 0.325 | 0.377 |
| e | 53.200 | 43.459 | 65.797 |
| sigma | 0.051 | 0.039 | 0.070 |

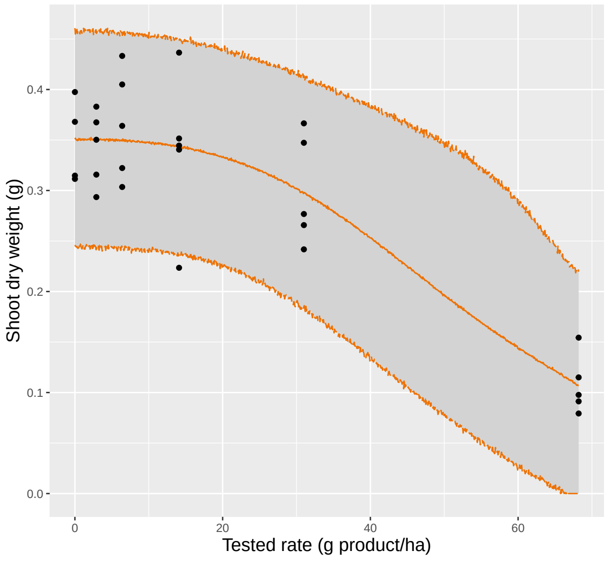

(a) Dose-response curve

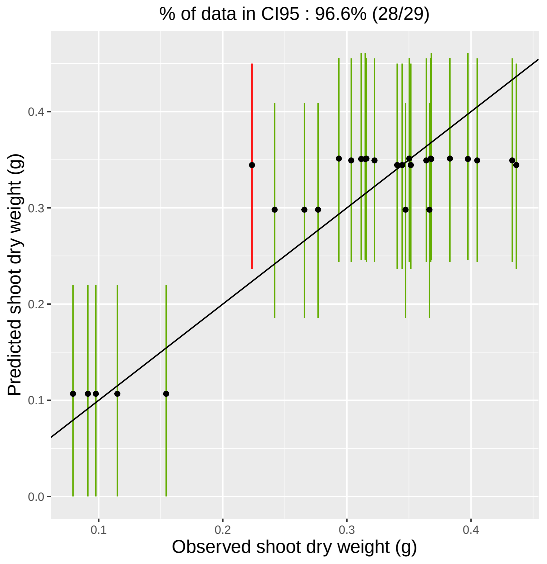

(b) Posterior predictive check (PPC)

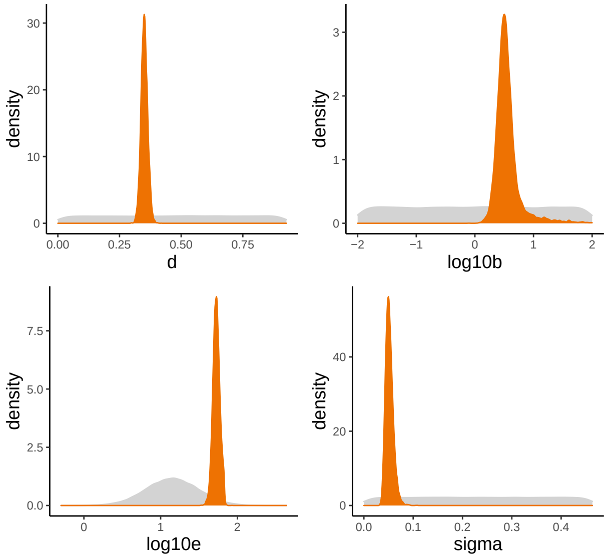

(c) Priors and posteriors

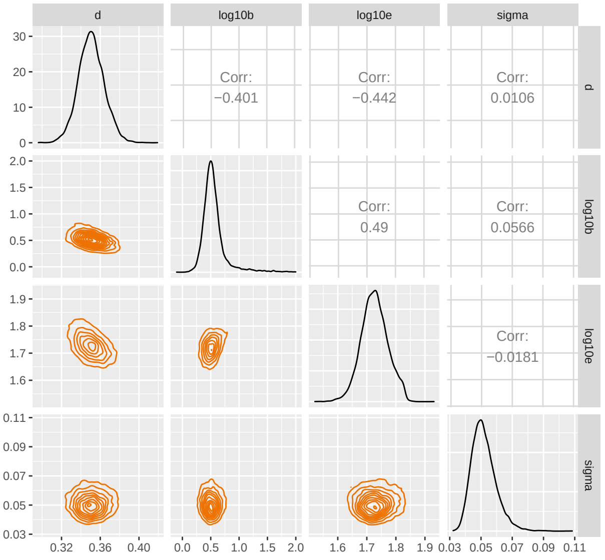

(d) Correlations between parameters

Figure 1: Dose-response curve (a), PPC (b), prior and posterior distributions (c) and correlations between parameters (d).

#### Data set: AVESA_VV_weight

Table 2: Summary of parameter estimates for AVESA_VV_weight data set

| Parameter | median | Q2.5 | Q97.5 |
|---|---|---|---|
| b | 2.275 | 1.272 | 20.065 |
| d | 1.225 | 1.092 | 1.368 |
| e | 31.971 | 26.373 | 37.222 |
| sigma | 0.112 | 0.075 | 0.196 |

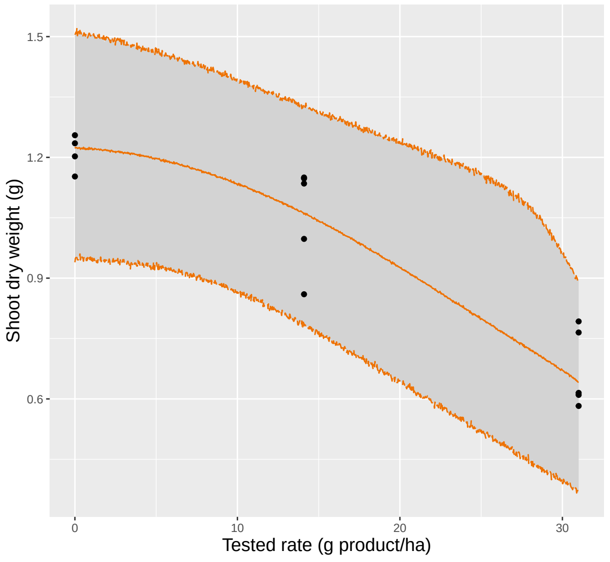

(a) Dose-response curve

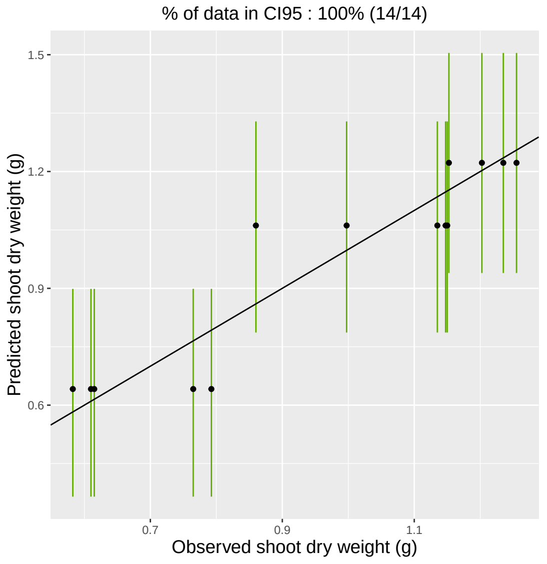

(b) Posterior predictive check (PPC)

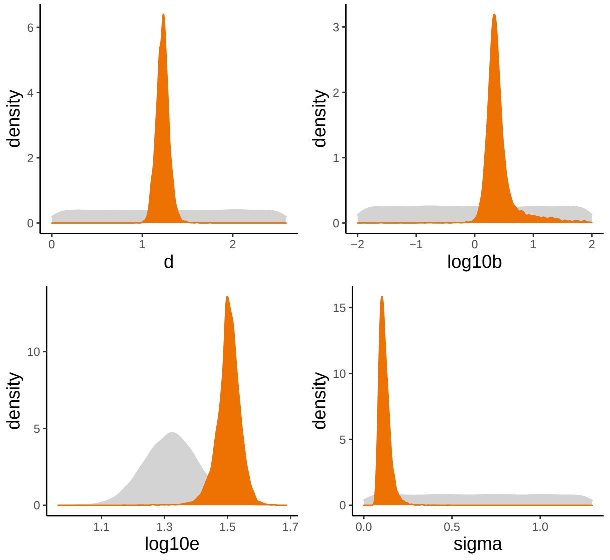

(c) Priors and posteriors

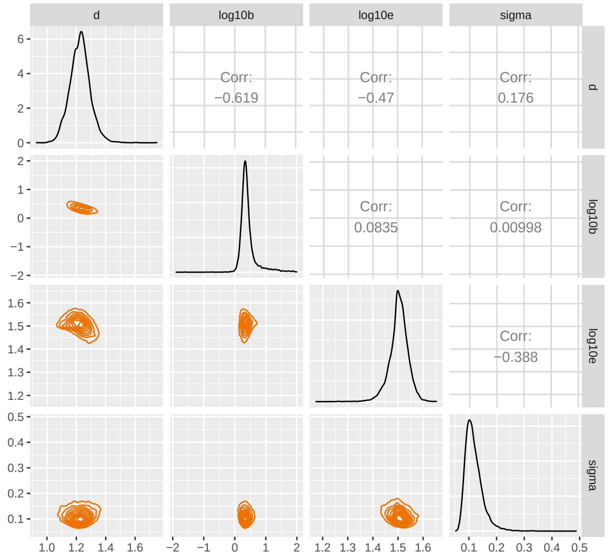

(d) Correlations between parameters

Figure 2: Dose-response curve (a), PPC (b), prior and posterior distributions (c) and correlations between parameters (d).

#### Data set: BEAVA_VV_weight

Table 3: Summary of parameter estimates for BEAVA_VV_weight data set

| Parameter | median | Q2.5 | Q97.5 |
|---|---|---|---|
| b | 1.200 | 0.932 | 1.564 |
| d | 3.545 | 3.272 | 3.826 |
| e | 7.202 | 5.817 | 9.057 |
| sigma | 0.453 | 0.378 | 0.556 |

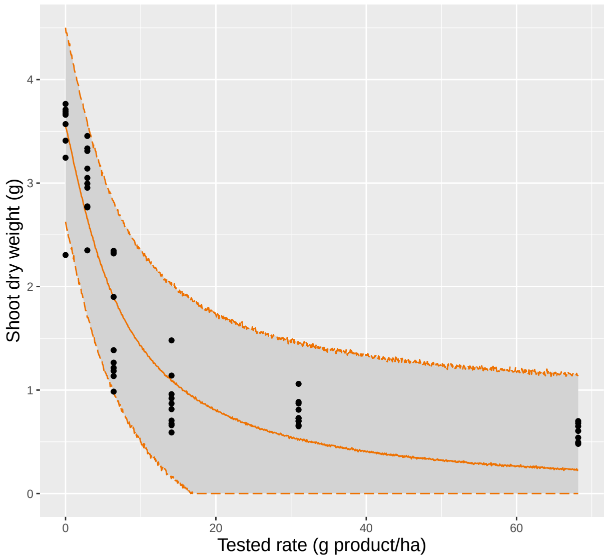

(a) Dose-response curve

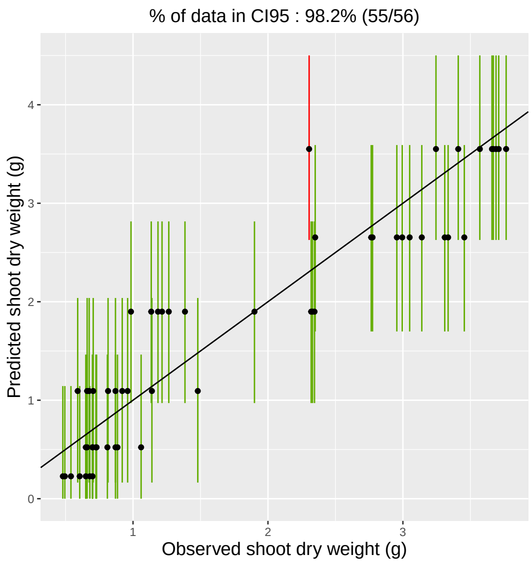

(b) Posterior predictive check (PPC)

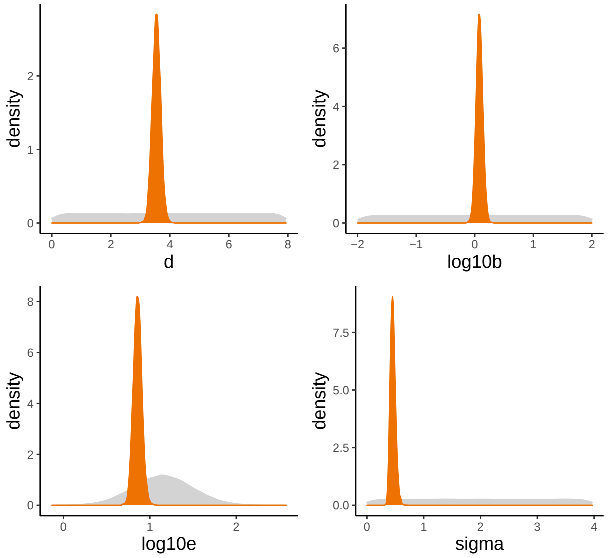

(c) Priors and posteriors

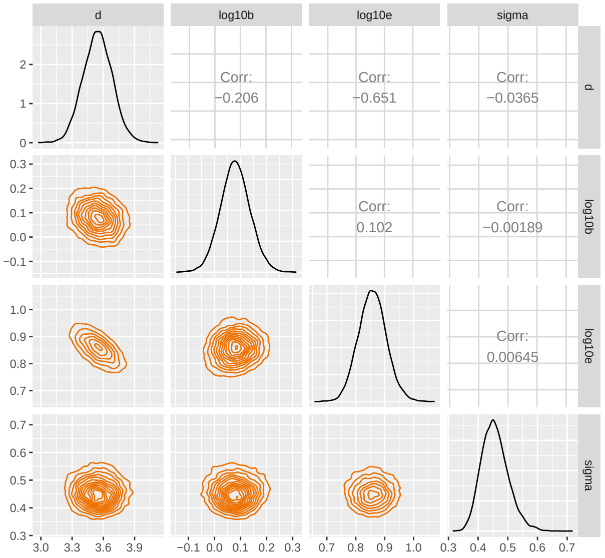

(d) Correlations between parameters

Figure 3: Dose-response curve (a), PPC (b), prior and posterior distributions (c) and correlations between parameters (d).

#### Data set: BRSNW_VV_weight

Table 4: Summary of parameter estimates for BRSNW_VV_weight data set

| Parameter | median | Q2.5 | Q97.5 |
|---|---|---|---|
| b | 1.842 | 1.306 | 2.683 |
| d | 5.065 | 4.559 | 5.593 |
| e | 8.772 | 6.973 | 11.150 |
| sigma | 0.912 | 0.754 | 1.130 |

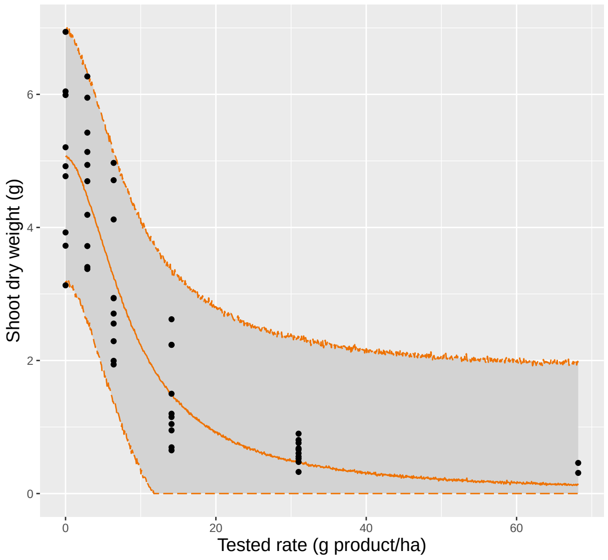

(a) Dose-response curve

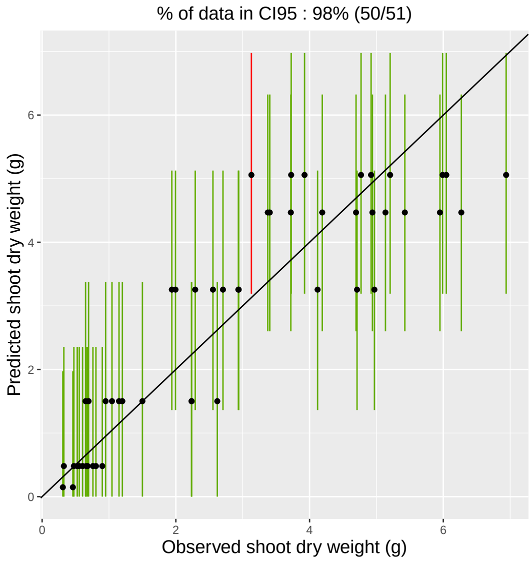

(b) Posterior predictive check (PPC)

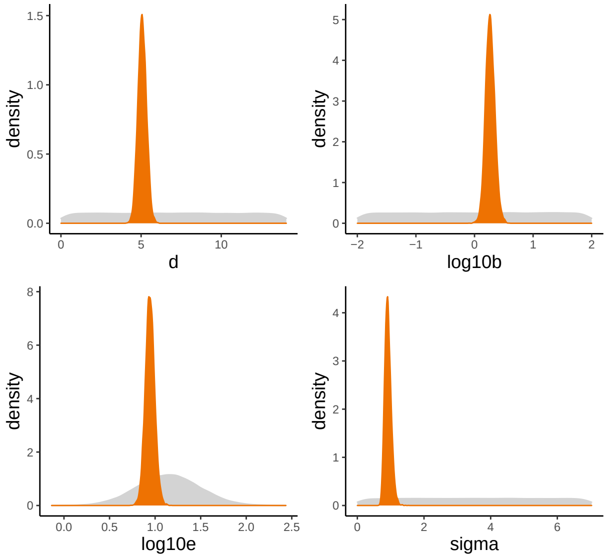

(c) Priors and posteriors

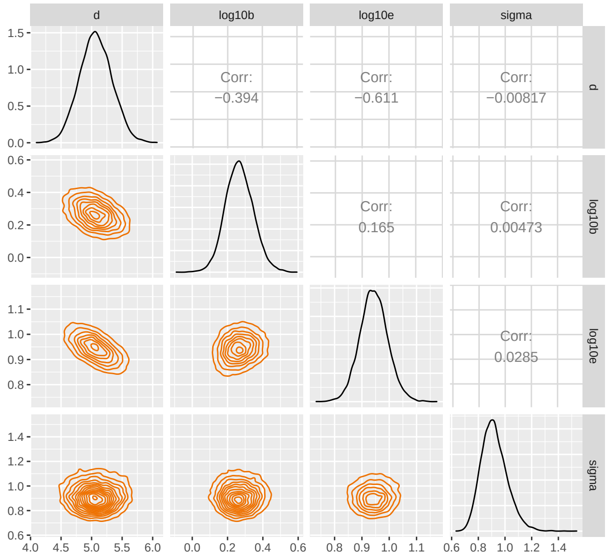

(d) Correlations between parameters

Figure 4: Dose-response curve (a), PPC (b), prior and posterior distributions (c) and correlations between parameters (d).

### Data set: CUMSA_VV_weight

Table 5: Summary of parameter estimates for CUMSA_VV_weight data set

| Parameter | median | Q2.5 | Q97.5 |
|---|---|---|---|
| b | 0.989 | 0.801 | 1.213 |
| d | 8.268 | 7.685 | 8.874 |
| e | 14.147 | 11.163 | 18.087 |
| sigma | 0.993 | 0.834 | 1.207 |

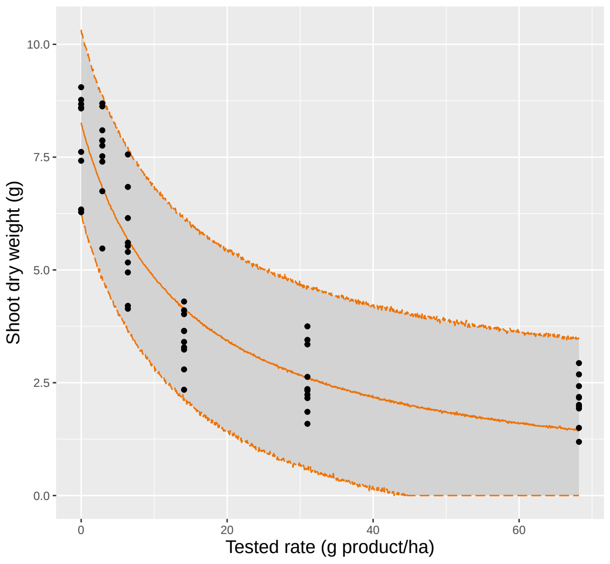

(a) Dose-response curve

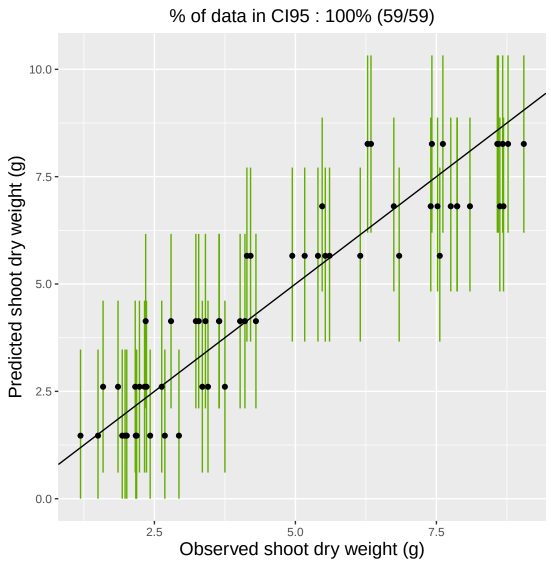

(b) Posterior predictive check (PPC)

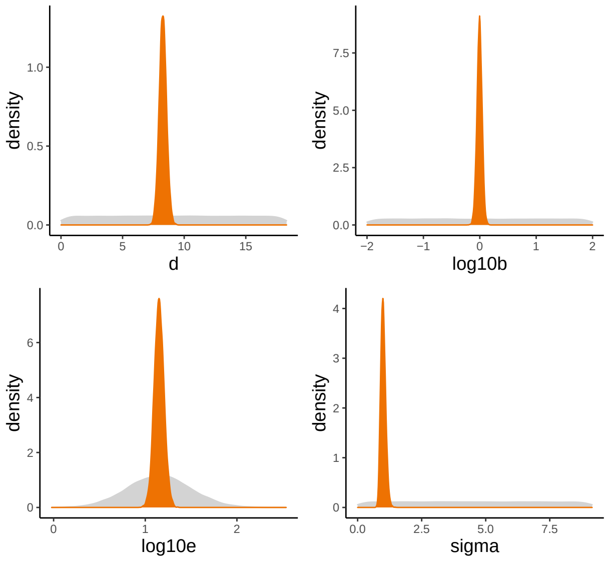

(c) Priors and posteriors

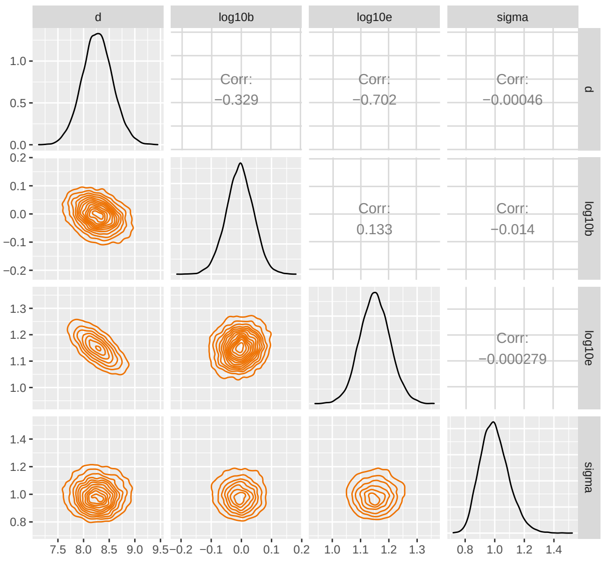

(d) Correlations between parameters

Figure 5: Dose-response curve (a), PPC (b), prior and posterior distributions (c) and correlations between parameters (d).

#### Data set: GLXMA_VV_weight

Table 6: Summary of parameter estimates for GLXMA_VV_weight data set

| Parameter | median | Q2.5 | Q97.5 |
|---|---|---|---|
| b | 1.432 | 1.219 | 1.703 |
| d | 5.129 | 4.870 | 5.400 |
| e | 14.249 | 12.358 | 16.367 |
| sigma | 0.470 | 0.395 | 0.573 |

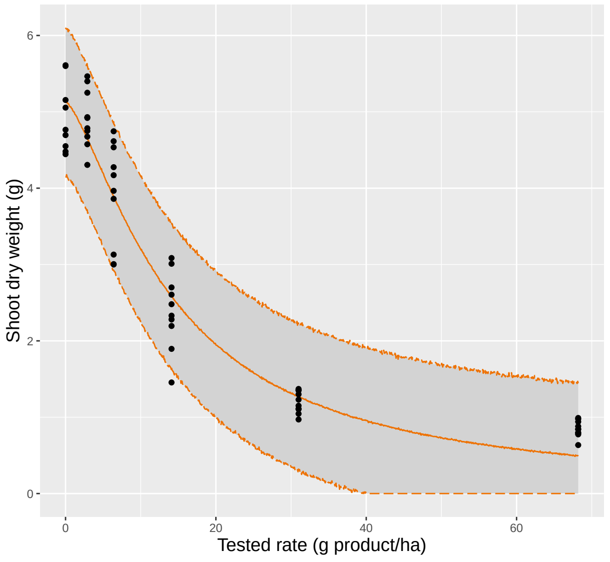

(a) Dose-response curve

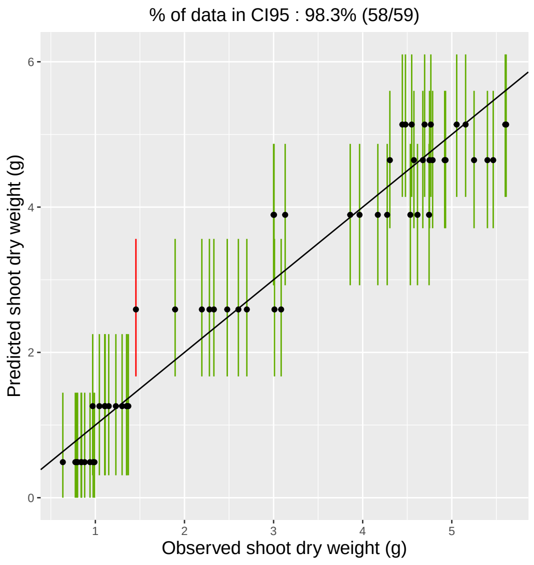

(b) Posterior predictive check (PPC)

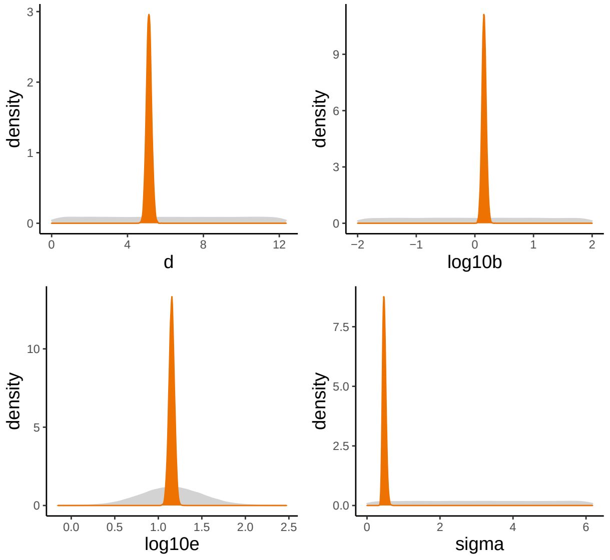

(c) Priors and posteriors

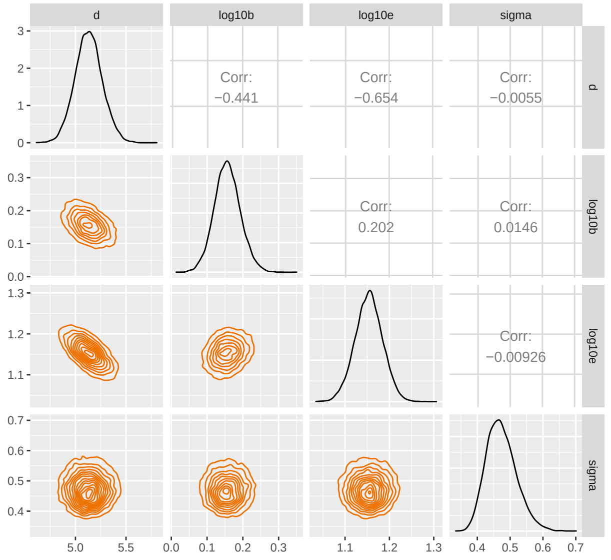

(d) Correlations between parameters

Figure 6: Dose-response curve (a), PPC (b), prior and posterior distributions (c) and correlations between parameters (d).

#### Data set: HELAN_VV_weight

Table 7: Summary of parameter estimates for HELAN_VV_weight data set

| Parameter | median | Q2.5 | Q97.5 |
|---|---|---|---|
| b | 5.860 | 3.602 | 46.854 |
| d | 2.651 | 2.562 | 2.751 |
| e | 12.793 | 11.729 | 13.931 |
| sigma | 0.217 | 0.175 | 0.278 |

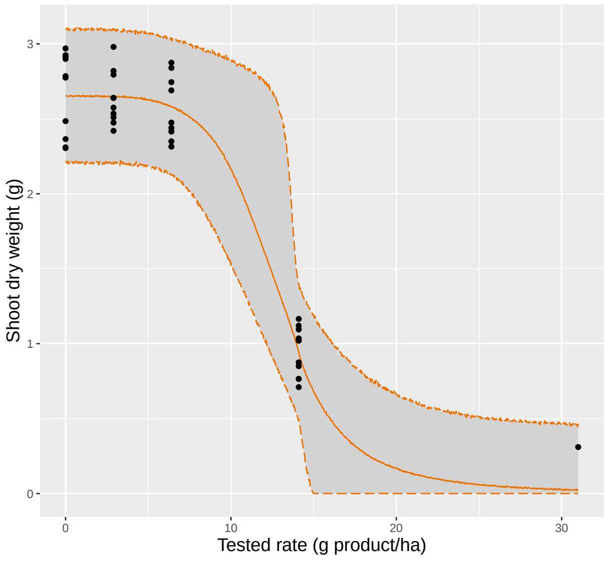

(a) Dose-response curve

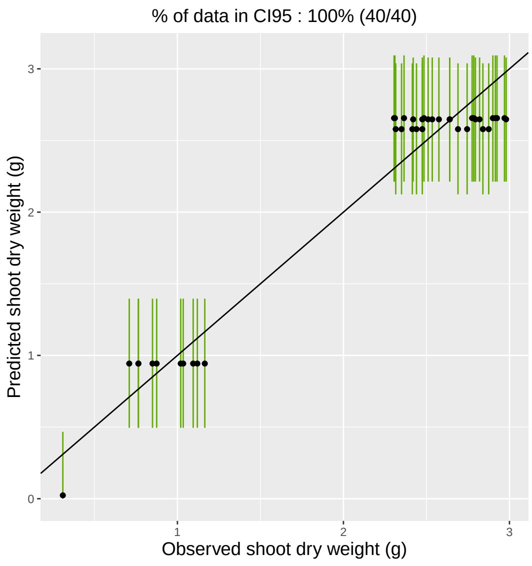

(b) Posterior predictive check (PPC)

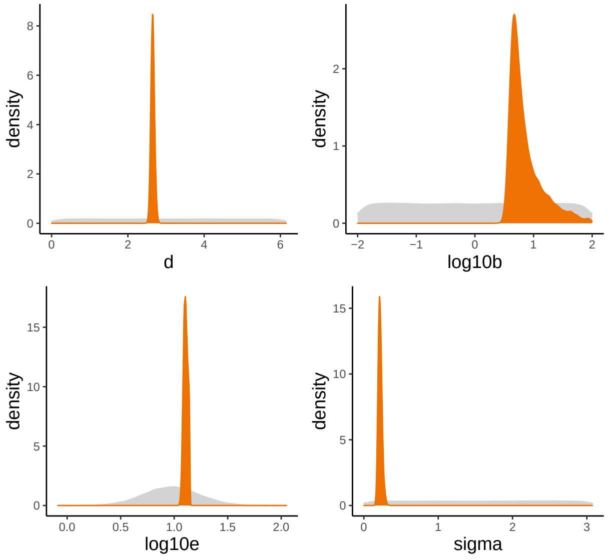

(c) Priors and posteriors

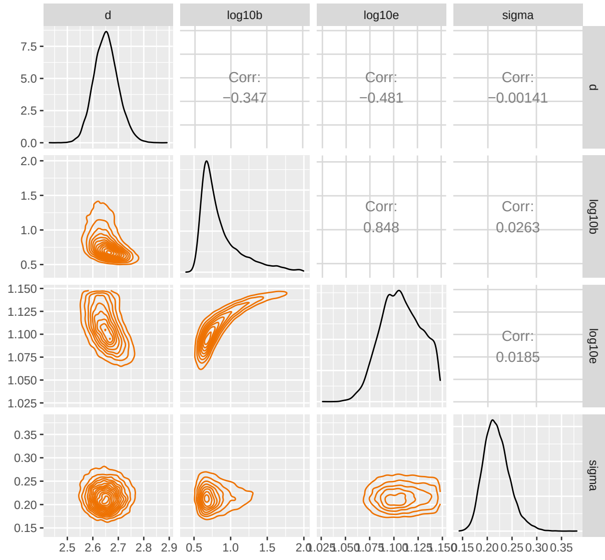

(d) Correlations between parameters

Figure 7: Dose-response curve (a), PPC (b), prior and posterior distributions (c) and correlations between parameters (d).

### Data set: LYPES_VV_weight

Table 8: Summary of parameter estimates for LYPES_VV_weight data set

| Parameter | median | Q2.5 | Q97.5 |
|---|---|---|---|
| b | 1.394 | 1.165 | 1.686 |
| d | 4.492 | 4.218 | 4.767 |
| e | 10.757 | 9.141 | 12.670 |
| sigma | 0.476 | 0.400 | 0.583 |

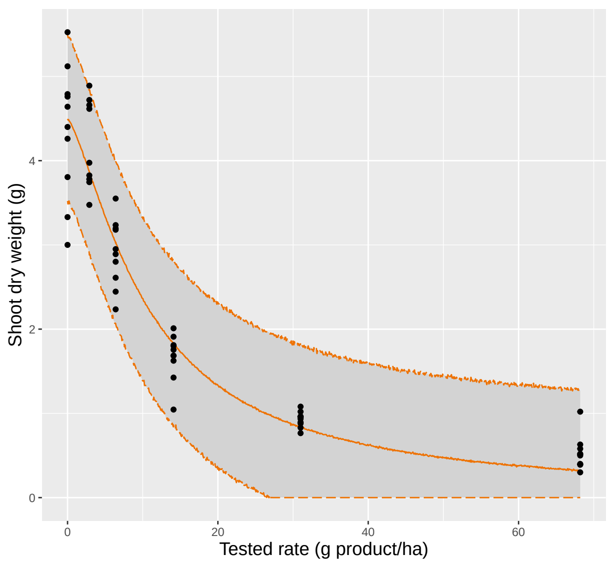

(a) Dose-response curve

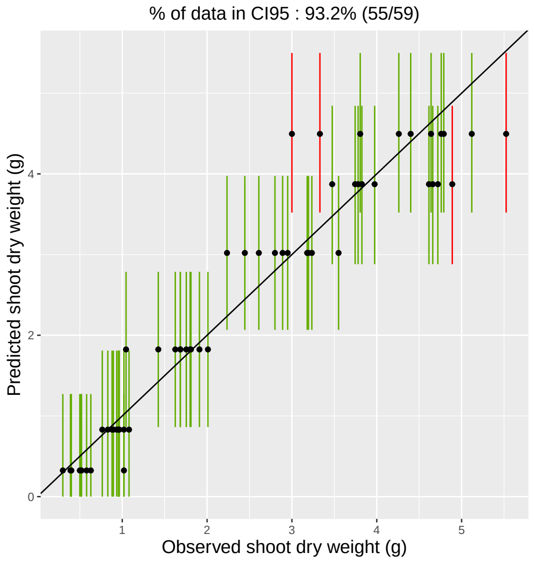

(b) Posterior predictive check (PPC)

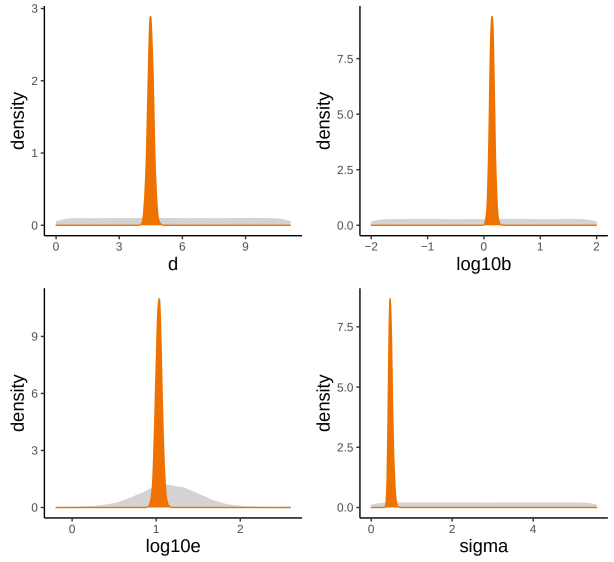

(c) Priors and posteriors

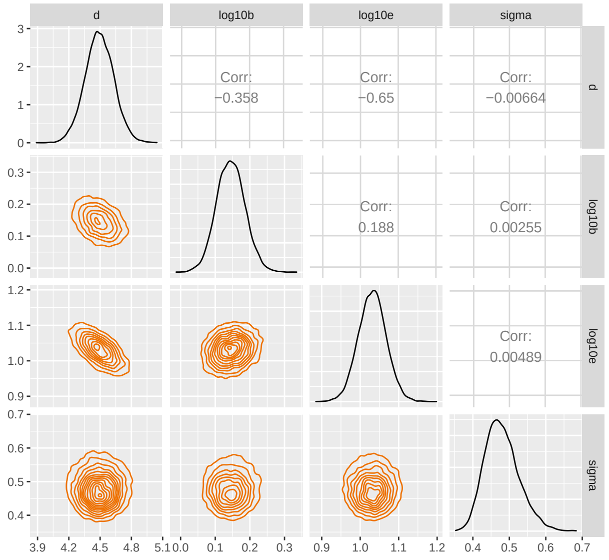

(d) Correlations between parameters

Figure 8: Dose-response curve (a), PPC (b), prior and posterior distributions (c) and correlations between parameters (d).

#### Data set: TRZAW_VV_weight

Table 9: Summary of parameter estimates for TRZAW_VV_weight data set

| Parameter | median | Q2.5 | Q97.5 |
|---|---|---|---|
| b | 2.985 | 2.185 | 4.139 |
| d | 1.293 | 1.233 | 1.353 |
| e | 248.882 | 219.795 | 281.520 |
| sigma | 0.121 | 0.094 | 0.165 |

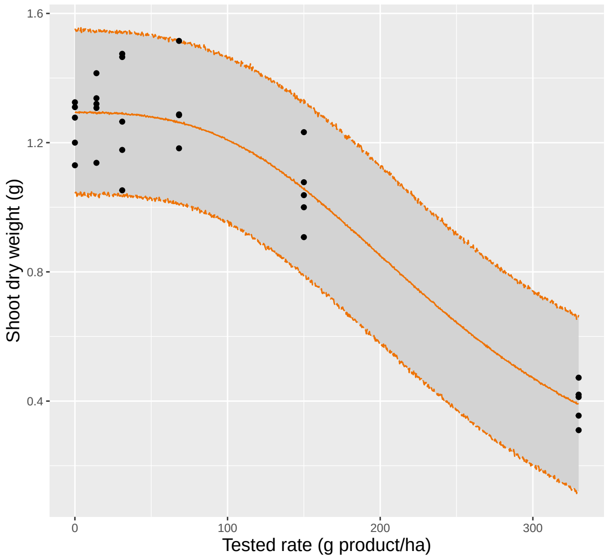

(a) Dose-response curve

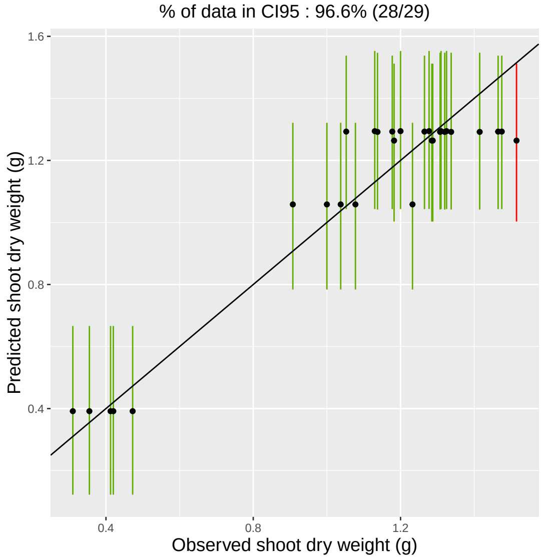

(b) Posterior predictive check (PPC)

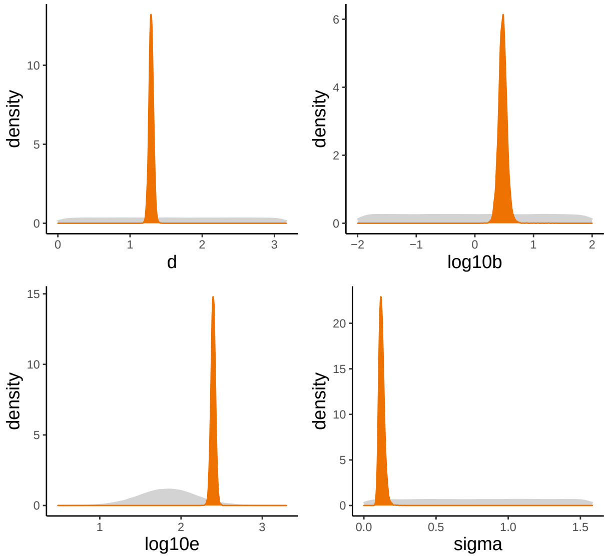

(c) Priors and posteriors

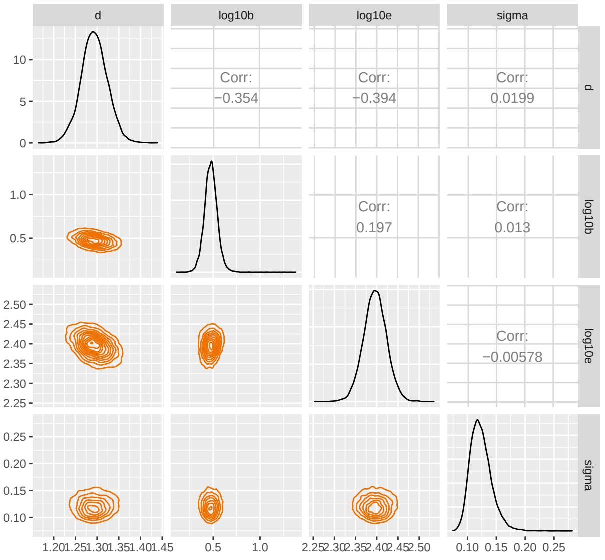

(d) Correlations between parameters

Figure 9: Dose-response curve (a), PPC (b), prior and posterior distributions (c) and correlations between parameters (d).

#### Data set: ZEAMA_VV_weight

Table 10: Summary of parameter estimates for ZEAMA_VV_weight data set

| Parameter | median | Q2.5 | Q97.5 |
|---|---|---|---|
| b | 3.503 | 2.595 | 4.869 |
| d | 7.719 | 7.367 | 8.077 |
| e | 213.231 | 189.844 | 240.396 |
| sigma | 1.040 | 0.875 | 1.269 |

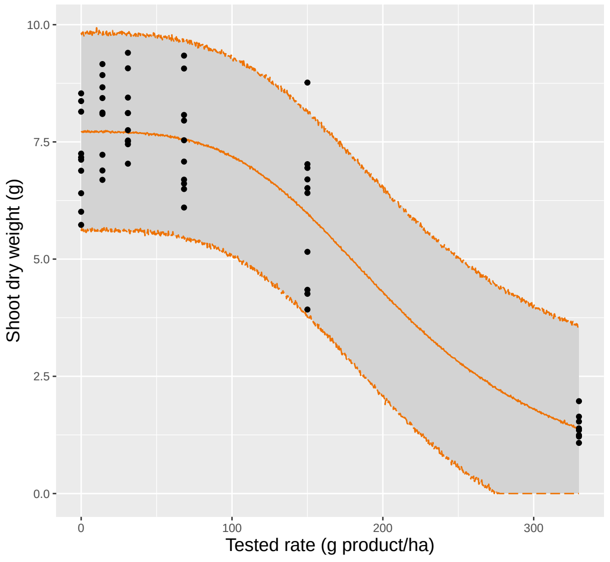

(a) Dose-response curve

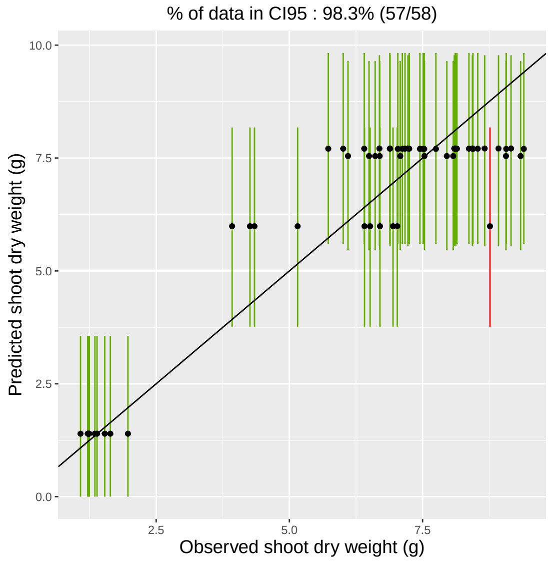

(b) Posterior predictive check (PPC)

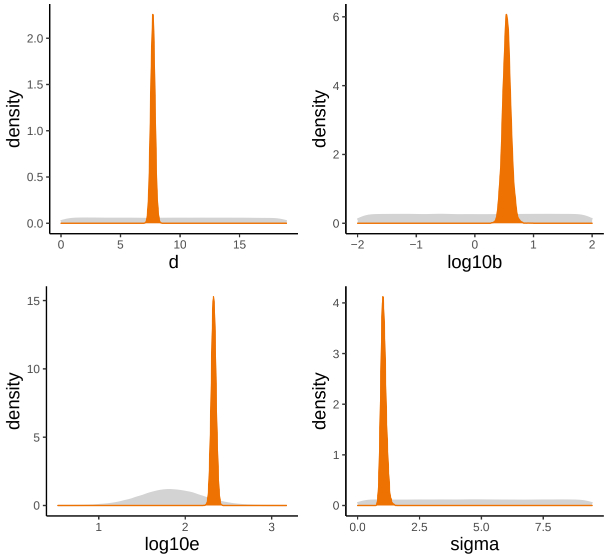

(c) Priors and posteriors

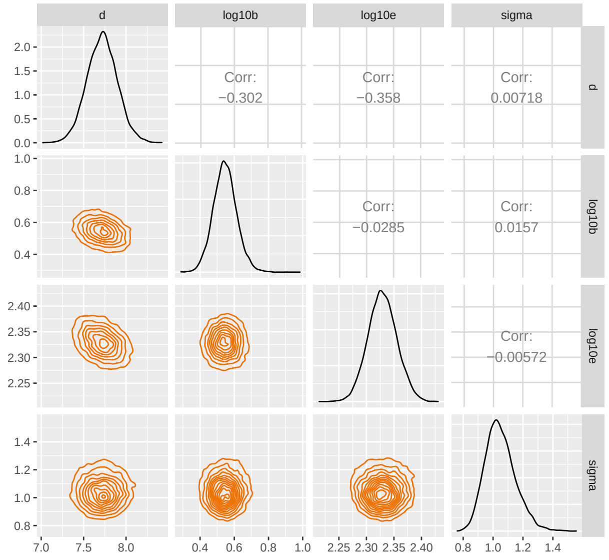

(d) Correlations between parameters

Figure 10: Dose-response curve (a), PPC (b), prior and posterior distributions (c) and correlations between parameters (d).
