## Supplementary material for "How to account for the uncertainty from standard toxicity tests in species sensitivity distributions: an example in non-target plants": Archive with reports on studies: ER50_censoring.pdf

### $ER_{50}$ censoring after dose-response analyses

Study 7

25 June 2020

Contact:

---

This document provides the $ER_{50}$ censoring after dose-response analyses for study 7, including four parts:

#### SE test - emergence endpoint

Table 2: Minimum and maximum tested rates, median, quantile 2.5% and quantile 97.5% of the $ER_{50}$ posterior distribution - emergence endpoint of the SE test of study 7

| species | min_rate | max_rate | $ER_{50}$_median | $ER_{50}$_LCI95 | $ER_{50}$_UCI95 |
|---|---|---|---|---|---|
| ALLCE | 0.166 | 90.91 | 97.05953 | 45.11447 | 334.4929 |
| AVESA | 3.880 | 90.91 | 122.90510 | 94.92559 | 261.2516 |
| BEAVA | 0.802 | 90.91 | 445.00190 | 186.97293 | 1494.5358 |
| BRSNW | 3.880 | 90.91 | 95.13338 | 62.53236 | 181.6225 |
| CUMSA | 3.880 | 90.91 | 126.03630 | 95.62384 | 275.4094 |
| GLXMA | 3.880 | 90.91 | 265.74120 | 145.00580 | 634.7157 |
| HELAN | 3.880 | 90.91 | 264.95104 | 140.47192 | 626.0006 |
| LYPES | 3.880 | 90.91 | 128.14477 | 96.39124 | 291.5196 |
| TRZAW | 3.880 | 90.91 | 169.38203 | 113.75338 | 358.2504 |
| ZEAMA | 3.880 | 90.91 | 131.39009 | 97.14600 | 294.9957 |

Table 3: Censored $ER_{50}$ according to criterion 1 and the seven decision thresholds - emergence endpoint of the SE test of study 7

| species | C1T0.0 | C1T0.2 | C1T0.4 | C1T0.5 | C1T0.6 | C1T0.8 | C1T1.0 |
|---|---|---|---|---|---|---|---|
| ALLCE | [45.11; 334.49] | [45.11; 334.49] | [45.11; 334.49] | [45.11; Inf) | [45.11; Inf) | [45.11; Inf) | [45.11; Inf) |
| AVESA | [90.91; Inf) | [90.91; Inf) | [90.91; Inf) | [90.91; Inf) | [90.91; Inf) | [90.91; Inf) | [90.91; Inf) |
| BEAVA | [90.91; Inf) | [90.91; Inf) | [90.91; Inf) | [90.91; Inf) | [90.91; Inf) | [90.91; Inf) | [90.91; Inf) |
| BRSNW | [62.53; 181.62] | [62.53; 181.62] | [62.53; 181.62] | [62.53; Inf) | [62.53; Inf) | [62.53; Inf) | [62.53; Inf) |
| CUMSA | [90.91; Inf) | [90.91; Inf) | [90.91; Inf) | [90.91; Inf) | [90.91; Inf) | [90.91; Inf) | [90.91; Inf) |
| GLXMA | [90.91; Inf) | [90.91; Inf) | [90.91; Inf) | [90.91; Inf) | [90.91; Inf) | [90.91; Inf) | [90.91; Inf) |
| HELAN | [90.91; Inf) | [90.91; Inf) | [90.91; Inf) | [90.91; Inf) | [90.91; Inf) | [90.91; Inf) | [90.91; Inf) |
| LYPES | [90.91; Inf) | [90.91; Inf) | [90.91; Inf) | [90.91; Inf) | [90.91; Inf) | [90.91; Inf) | [90.91; Inf) |
| TRZAW | [90.91; Inf) | [90.91; Inf) | [90.91; Inf) | [90.91; Inf) | [90.91; Inf) | [90.91; Inf) | [90.91; Inf) |
| ZEAMA | [90.91; Inf) | [90.91; Inf) | [90.91; Inf) | [90.91; Inf) | [90.91; Inf) | [90.91; Inf) | [90.91; Inf) |

Table 4: Censored $ER_{50}$ according to criterion 2 and the seven decision thresholds - emergence endpoint of the SE test of study 7

| species | C2T0.0 | C2T0.2 | C2T0.4 | C2T0.5 | C2T0.6 | C2T0.8 | C2T1.0 |
|---|---|---|---|---|---|---|---|
| ALLCE | [45.11; 334.49] | [45.11; Inf) | [45.11; Inf) | [45.11; Inf) | [45.11; Inf) | [45.11; Inf) | [45.11; Inf) |
| AVESA | [90.91; Inf) | [90.91; Inf) | [90.91; Inf) | [90.91; Inf) | [90.91; Inf) | [90.91; Inf) | [90.91; Inf) |
| BEAVA | [90.91; Inf) | [90.91; Inf) | [90.91; Inf) | [90.91; Inf) | [90.91; Inf) | [90.91; Inf) | [90.91; Inf) |
| BRSNW | [62.53; 181.62] | [62.53; 181.62] | [62.53; Inf) | [62.53; Inf) | [62.53; Inf) | [62.53; Inf) | [62.53; Inf) |
| CUMSA | [90.91; Inf) | [90.91; Inf) | [90.91; Inf) | [90.91; Inf) | [90.91; Inf) | [90.91; Inf) | [90.91; Inf) |
| GLXMA | [90.91; Inf) | [90.91; Inf) | [90.91; Inf) | [90.91; Inf) | [90.91; Inf) | [90.91; Inf) | [90.91; Inf) |
| HELAN | [90.91; Inf) | [90.91; Inf) | [90.91; Inf) | [90.91; Inf) | [90.91; Inf) | [90.91; Inf) | [90.91; Inf) |
| LYPES | [90.91; Inf) | [90.91; Inf) | [90.91; Inf) | [90.91; Inf) | [90.91; Inf) | [90.91; Inf) | [90.91; Inf) |
| TRZAW | [90.91; Inf) | [90.91; Inf) | [90.91; Inf) | [90.91; Inf) | [90.91; Inf) | [90.91; Inf) | [90.91; Inf) |
| ZEAMA | [90.91; Inf) | [90.91; Inf) | [90.91; Inf) | [90.91; Inf) | [90.91; Inf) | [90.91; Inf) | [90.91; Inf) |

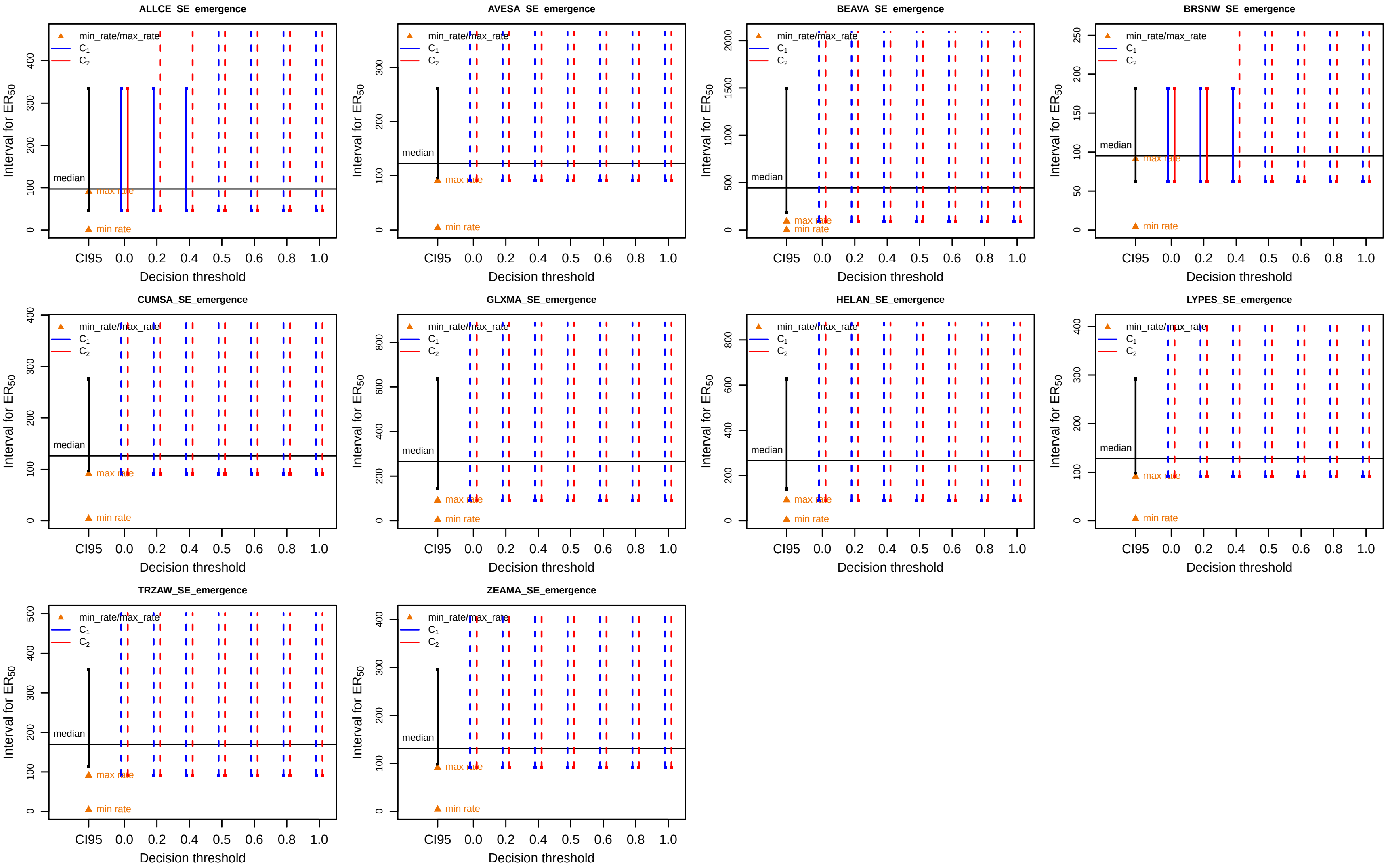


Figure 1: CI95 of $ER_{50}$ and censored $ER_{50}$ according to both criteria and the seven decision thresholds for all species - emergence endpoint of the SE test of study 7. The two orange triangles stand for the lowest and the highest tested rates. Vertical segments of three different colors (black, blue, red) represent the CI95 of $ER_{50}$, the censored $ER_{50}$ according to $C_1$ and the censored $ER_{50}$ according to $C_2$, respectively; solid vertical segments are for bounded intervals while dotted vertical segments stand for right-unbounded intervals; the black horizontal line represents the median of the $ER_{50}$.

#### SE test - survival endpoint

Table 5: Minimum and maximum tested rates, median, quantile 2.5% and quantile 97.5% of the $ER_{50}$ posterior distribution - survival endpoint of the SE test of study 7

| species | min_rate | max_rate | $ER_{50}$_median | $ER_{50}$_LCI95 | $ER_{50}$_UCI95 |
|---|---|---|---|---|---|
| ALLCE | 0.166 | 90.91 | 2.689684 | 2.058483 | 3.46178 |
| AVESA | 3.880 | 90.91 | 97.441835 | 68.021285 | 172.49433 |
| BEAVA | 0.802 | 90.91 | 13.936858 | 11.068070 | 17.63860 |
| BRSNW | 3.880 | 90.91 | 56.653232 | 41.742047 | 83.63910 |
| CUMSA | 3.880 | 90.91 | 178.796740 | 107.109086 | 395.70705 |
| GLXMA | 3.880 | 90.91 | 187.844440 | 120.473427 | 419.63462 |
| HELAN | 3.880 | 90.91 | 78.272547 | 53.723540 | 130.92953 |
| LYPES | 3.880 | 90.91 | 88.004095 | 66.529442 | 133.26106 |
| TRZAW | 3.880 | 90.91 | 129.696489 | 97.384962 | 293.64758 |
| ZEAMA | 3.880 | 90.91 | 130.078055 | 97.205533 | 285.80160 |

Table 6: Censored $ER_{50}$ according to criterion 1 and the seven decision thresholds - survival endpoint of the SE test of study 7

| species | C1T0.0 | C1T0.2 | C1T0.4 | C1T0.5 | C1T0.6 | C1T0.8 | C1T1.0 |
|---|---|---|---|---|---|---|---|
| ALLCE | [2.06; 3.46] | [2.06; 3.46] | [2.06; 3.46] | [2.06; 3.46] | [2.06; 3.46] | [2.06; 3.46] | [2.06; 3.46] |
| AVESA | [68.02; 172.49] | [68.02; 172.49] | [68.02; Inf) | [68.02; Inf) | [68.02; Inf) | [68.02; Inf) | [68.02; Inf) |
| BEAVA | [11.07; 17.64] | [11.07; 17.64] | [11.07; 17.64] | [11.07; 17.64] | [11.07; 17.64] | [11.07; 17.64] | [11.07; 17.64] |
| BRSNW | [41.74; 83.64] | [41.74; 83.64] | [41.74; 83.64] | [41.74; 83.64] | [41.74; 83.64] | [41.74; 83.64] | [41.74; 83.64] |
| CUMSA | [90.91; Inf) | [90.91; Inf) | [90.91; Inf) | [90.91; Inf) | [90.91; Inf) | [90.91; Inf) | [90.91; Inf) |
| GLXMA | [90.91; Inf) | [90.91; Inf) | [90.91; Inf) | [90.91; Inf) | [90.91; Inf) | [90.91; Inf) | [90.91; Inf) |
| HELAN | [53.72; 130.93] | [53.72; 130.93] | [53.72; 130.93] | [53.72; 130.93] | [53.72; 130.93] | [53.72; Inf) | [53.72; Inf) |
| LYPES | [66.53; 133.26] | [66.53; 133.26] | [66.53; 133.26] | [66.53; 133.26] | [66.53; Inf) | [66.53; Inf) | [66.53; Inf) |
| TRZAW | [90.91; Inf) | [90.91; Inf) | [90.91; Inf) | [90.91; Inf) | [90.91; Inf) | [90.91; Inf) | [90.91; Inf) |
| ZEAMA | [90.91; Inf) | [90.91; Inf) | [90.91; Inf) | [90.91; Inf) | [90.91; Inf) | [90.91; Inf) | [90.91; Inf) |

Table 7: Censored $ER_{50}$ according to criterion 2 and the seven decision thresholds - survival endpoint of the SE test of study 7

| species | C2T0.0 | C2T0.2 | C2T0.4 | C2T0.5 | C2T0.6 | C2T0.8 | C2T1.0 |
|---|---|---|---|---|---|---|---|
| ALLCE | [2.06; 3.46] | [2.06; 3.46] | [2.06; 3.46] | [2.06; 3.46] | [2.06; 3.46] | [2.06; 3.46] | [2.06; 3.46] |
| AVESA | [68.02; 172.49] | [68.02; 172.49] | [68.02; Inf) | [68.02; Inf) | [68.02; Inf) | [68.02; Inf) | [68.02; Inf) |
| BEAVA | [11.07; 17.64] | [11.07; 17.64] | [11.07; 17.64] | [11.07; 17.64] | [11.07; 17.64] | [11.07; 17.64] | [11.07; 17.64] |
| BRSNW | [41.74; 83.64] | [41.74; 83.64] | [41.74; 83.64] | [41.74; 83.64] | [41.74; 83.64] | [41.74; 83.64] | [41.74; 83.64] |
| CUMSA | [90.91; Inf) | [90.91; Inf) | [90.91; Inf) | [90.91; Inf) | [90.91; Inf) | [90.91; Inf) | [90.91; Inf) |
| GLXMA | [90.91; Inf) | [90.91; Inf) | [90.91; Inf) | [90.91; Inf) | [90.91; Inf) | [90.91; Inf) | [90.91; Inf) |
| HELAN | [53.72; 130.93] | [53.72; 130.93] | [53.72; 130.93] | [53.72; Inf) | [53.72; Inf) | [53.72; Inf) | [53.72; Inf) |
| LYPES | [66.53; 133.26] | [66.53; 133.26] | [66.53; Inf) | [66.53; Inf) | [66.53; Inf) | [66.53; Inf) | [66.53; Inf) |
| TRZAW | [90.91; Inf) | [90.91; Inf) | [90.91; Inf) | [90.91; Inf) | [90.91; Inf) | [90.91; Inf) | [90.91; Inf) |
| ZEAMA | [90.91; Inf) | [90.91; Inf) | [90.91; Inf) | [90.91; Inf) | [90.91; Inf) | [90.91; Inf) | [90.91; Inf) |

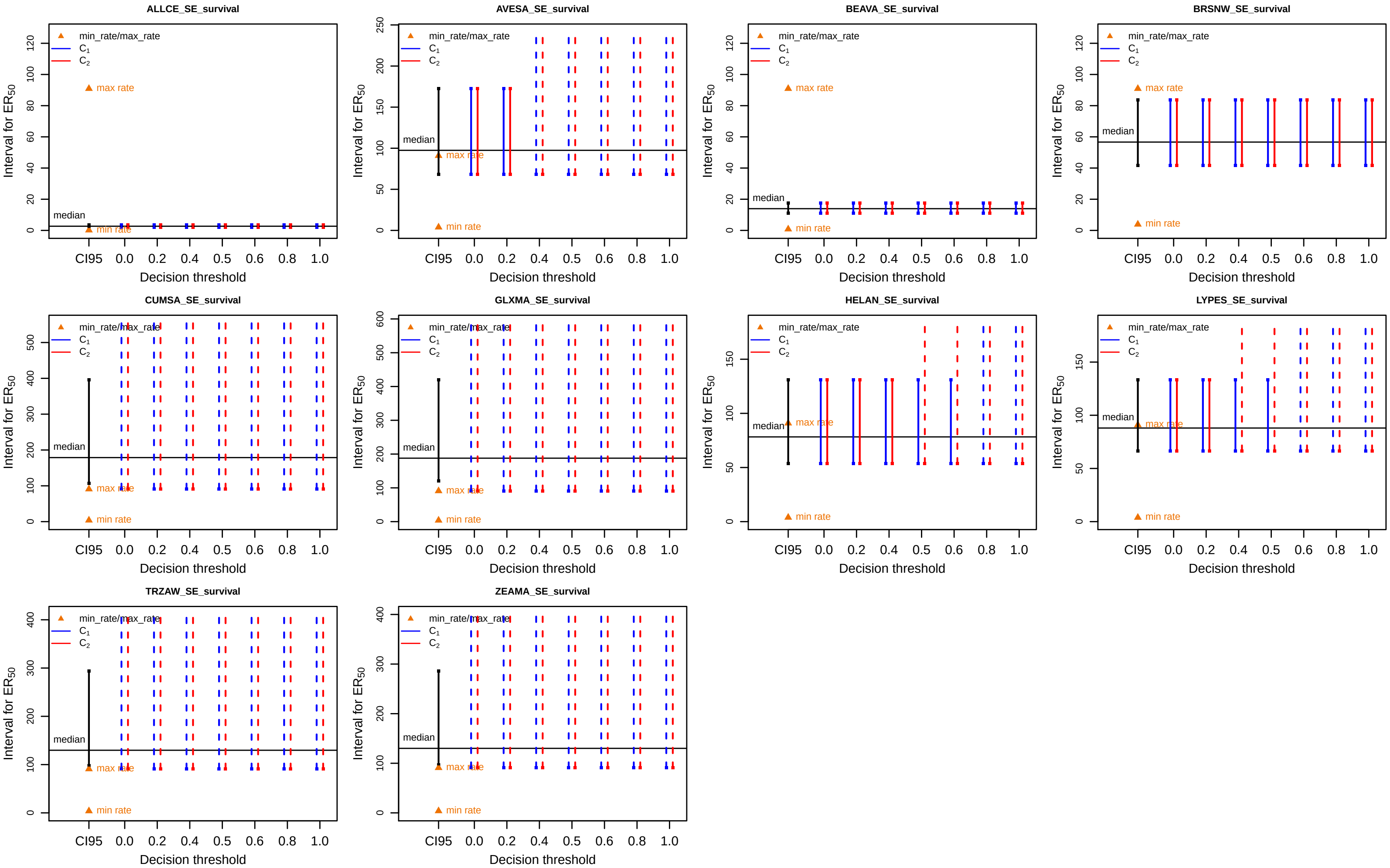


Figure 2: CI95 of $ER_{50}$ and censored $ER_{50}$ according to both criteria and the seven decision thresholds for all species - survival endpoint of the SE test of study 7. The two orange triangles stand for the lowest and the highest tested rates. Vertical segments of three different colors (black, blue, red) represent the CI95 of $ER_{50}$, the censored $ER_{50}$ according to $C_1$ and the censored $ER_{50}$ according to $C_2$, respectively; solid vertical segments are for bounded intervals while dotted vertical segments stand for right-unbounded intervals; the black horizontal line represents the median of the $ER_{50}$.

#### SE test - shoot dry weight endpoint

Table 8: Minimum and maximum tested rates, median, quantile 2.5% and quantile 97.5% of the $ER_{50}$ posterior distribution - shoot dry weight endpoint of the SE test of study 7

| species | min_rate | max_rate | $ER_{50}$_median | $ER_{50}$_LCI95 | $ER_{50}$_UCI95 |
|---|---|---|---|---|---|
| ALLCE | 0.166 | 18.78 | 0.3620842 | 0.0997492 | 1.435263 |
| AVESA | 3.880 | 90.91 | 13.6036784 | 11.4052582 | 16.097591 |
| BEAVA | 0.802 | 41.32 | 3.9756174 | 2.5908858 | 5.886766 |
| BRSNW | 3.880 | 90.91 | 11.2371002 | 9.2720797 | 13.747939 |
| CUMSA | 3.880 | 90.91 | 9.2826393 | 7.0020399 | 12.336570 |
| GLXMA | 3.880 | 90.91 | 52.5459370 | 43.1427806 | 64.219082 |
| HELAN | 3.880 | 90.91 | 10.5857543 | 7.1429281 | 15.550129 |
| LYPES | 3.880 | 90.91 | 15.8588676 | 13.6473161 | 18.115954 |
| TRZAW | 3.880 | 90.91 | 110.8278310 | 81.1229476 | 164.913404 |
| ZEAMA | 3.880 | 90.91 | 37.5100755 | 29.2653647 | 48.304069 |

Table 9: Censored $ER_{50}$ according to criterion 1 and the seven decision thresholds - shoot dry weight endpoint of the SE test of study 7

| species | C1T0.0 | C1T0.2 | C1T0.4 | C1T0.5 | C1T0.6 | C1T0.8 | C1T1.0 |
|---|---|---|---|---|---|---|---|
| ALLCE | [0.1; 1.44] | [0.1; 1.44] | [0.1; 1.44] | [0.1; 1.44] | [0.1; 1.44] | [0.1; 1.44] | [0.1; 1.44] |
| AVESA | [11.41; 16.1] | [11.41; 16.1] | [11.41; 16.1] | [11.41; 16.1] | [11.41; 16.1] | [11.41; 16.1] | [11.41; 16.1] |
| BEAVA | [2.59; 5.89] | [2.59; 5.89] | [2.59; 5.89] | [2.59; 5.89] | [2.59; 5.89] | [2.59; 5.89] | [2.59; 5.89] |
| BRSNW | [9.27; 13.75] | [9.27; 13.75] | [9.27; 13.75] | [9.27; 13.75] | [9.27; 13.75] | [9.27; 13.75] | [9.27; 13.75] |
| CUMSA | [7; 12.34] | [7; 12.34] | [7; 12.34] | [7; 12.34] | [7; 12.34] | [7; 12.34] | [7; 12.34] |
| GLXMA | [43.14; 64.22] | [43.14; 64.22] | [43.14; 64.22] | [43.14; 64.22] | [43.14; 64.22] | [43.14; 64.22] | [43.14; 64.22] |
| HELAN | [7.14; 15.55] | [7.14; 15.55] | [7.14; 15.55] | [7.14; 15.55] | [7.14; 15.55] | [7.14; 15.55] | [7.14; 15.55] |
| LYPES | [13.65; 18.12] | [13.65; 18.12] | [13.65; 18.12] | [13.65; 18.12] | [13.65; 18.12] | [13.65; 18.12] | [13.65; 18.12] |
| TRZAW | [81.12; 164.91] | [81.12; Inf) | [81.12; Inf) | [81.12; Inf) | [81.12; Inf) | [81.12; Inf) | [81.12; Inf) |
| ZEAMA | [29.27; 48.3] | [29.27; 48.3] | [29.27; 48.3] | [29.27; 48.3] | [29.27; 48.3] | [29.27; 48.3] | [29.27; 48.3] |

Table 10: Censored $ER_{50}$ according to criterion 2 and the seven decision thresholds - shoot dry weight endpoint of the SE test of study 7

| species | C2T0.0 | C2T0.2 | C2T0.4 | C2T0.5 | C2T0.6 | C2T0.8 | C2T1.0 |
|---|---|---|---|---|---|---|---|
| ALLCE | [0.1; 1.44] | [0.1; 1.44] | [0.1; 1.44] | [0.1; 1.44] | [0.1; 1.44] | [0.1; 1.44] | [0.1; 1.44] |
| AVESA | [11.41; 16.1] | [11.41; 16.1] | [11.41; 16.1] | [11.41; 16.1] | [11.41; 16.1] | [11.41; 16.1] | [11.41; 16.1] |
| BEAVA | [2.59; 5.89] | [2.59; 5.89] | [2.59; 5.89] | [2.59; 5.89] | [2.59; 5.89] | [2.59; 5.89] | [2.59; 5.89] |
| BRSNW | [9.27; 13.75] | [9.27; 13.75] | [9.27; 13.75] | [9.27; 13.75] | [9.27; 13.75] | [9.27; 13.75] | [9.27; 13.75] |
| CUMSA | [7; 12.34] | [7; 12.34] | [7; 12.34] | [7; 12.34] | [7; 12.34] | [7; 12.34] | [7; 12.34] |
| GLXMA | [43.14; 64.22] | [43.14; 64.22] | [43.14; 64.22] | [43.14; 64.22] | [43.14; 64.22] | [43.14; 64.22] | [43.14; 64.22] |
| HELAN | [7.14; 15.55] | [7.14; 15.55] | [7.14; 15.55] | [7.14; 15.55] | [7.14; 15.55] | [7.14; 15.55] | [7.14; 15.55] |
| LYPES | [13.65; 18.12] | [13.65; 18.12] | [13.65; 18.12] | [13.65; 18.12] | [13.65; 18.12] | [13.65; 18.12] | [13.65; 18.12] |
| TRZAW | [81.12; 164.91] | [81.12; Inf) | [81.12; Inf) | [81.12; Inf) | [81.12; Inf) | [81.12; Inf) | [81.12; Inf) |
| ZEAMA | [29.27; 48.3] | [29.27; 48.3] | [29.27; 48.3] | [29.27; 48.3] | [29.27; 48.3] | [29.27; 48.3] | [29.27; 48.3] |

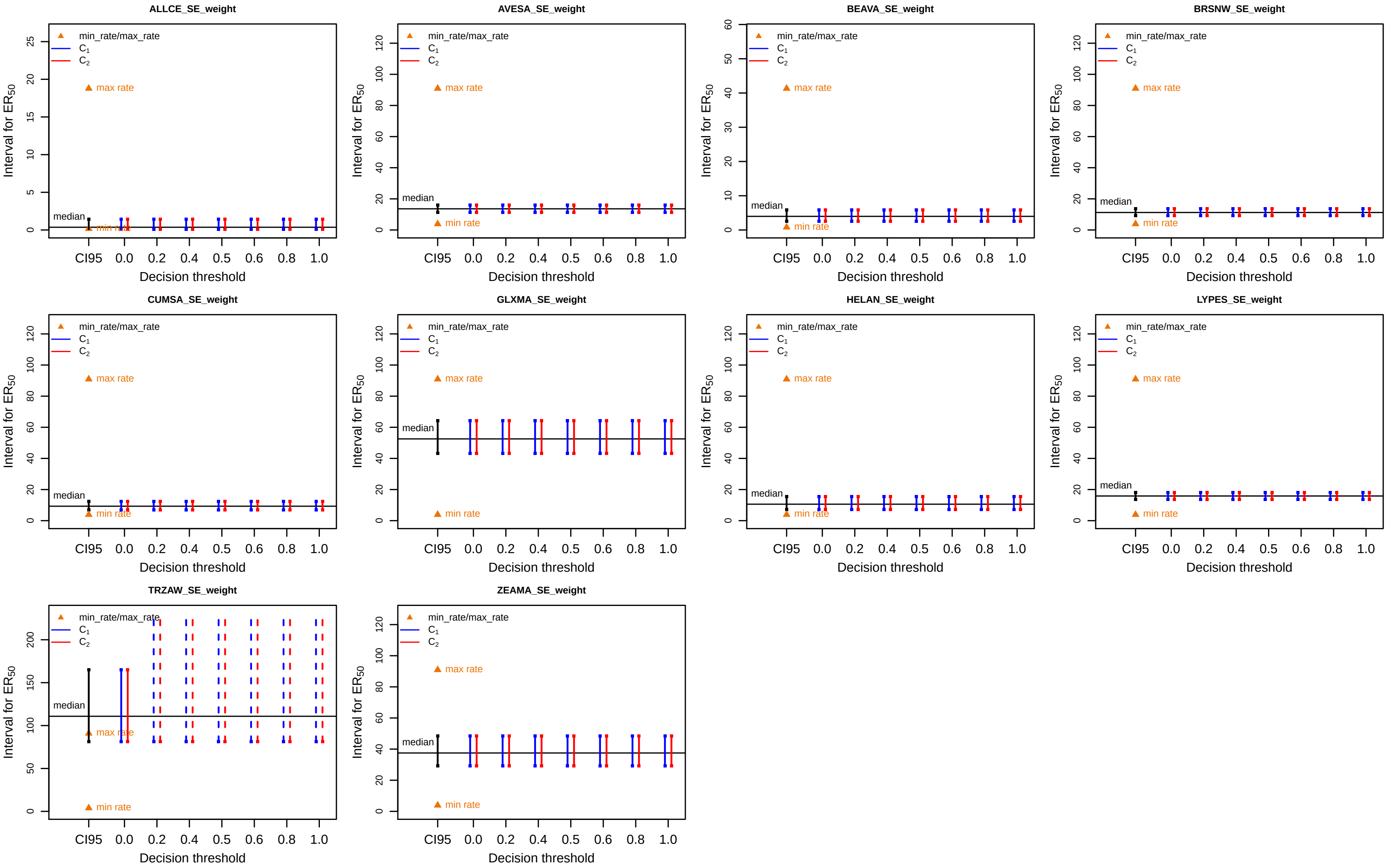


Figure 3: CI95 of $ER_{50}$ and censored $ER_{50}$ according to both criteria and the seven decision thresholds for all species - shoot dry weight endpoint of the SE test of study 7. The two orange triangles stand for the lowest and the highest tested rates. Vertical segments of three different colors (black, blue, red) represent the CI95 of $ER_{50}$, the censored $ER_{50}$ according to $C_1$ and the censored $ER_{50}$ according to $C_2$, respectively; solid vertical segments are for bounded intervals while dotted vertical segments stand for right-unbounded intervals; the black horizontal line represents the median of the $ER_{50}$.

### Vegatative vigor (VV) test of study 7

##### Raw data description (test item rates) for VV test

```
Number of species:  10
EPPO code:  ALLCE AVESA BEAVA BRSNW CUMSA GLXMA HELAN LYPES TRZAW ZEAMA
```

Table 11: Corresponding species and common names to the EPPO code

| EPPO code | Species name | Common name |
|---|---|---|
| ALLCE | *Allium cepa* | Onion |
| AVESA | *Avena sativa* | Oat |
| BEAVA | *Beta vulgaris* | Sugar beet |
| BRSNW | *Brassica napus winter* | Oilseed rape |
| CUMSA | *Cucumis sativus* | Cucumber |
| GLXMA | *Glycine max* | Soybean |
| HELAN | *Helianthus annuus* | Sunflower |
| LYPES | *Solanum lycopersicum* | Tomato |
| TRZAW | *Triticum aestivum winter* | Winter wheat |
| ZEAMA | *Zea mays subs. Amylacea* | Corn |

Table 12: Tested rates per species (g product/ha) for the VV test of study 7

| ALLCE | AVESA | BEAVA | BRSNW | CUMSA | GLXMA | HELAN | LYPES | TRZAW | ZEAMA |
|---|---|---|---|---|---|---|---|---|---|
| 0.00 | 0.00 | 0.000 | 0.000 | 0.00 | 0.00 | 0.00 | 0.00 | 0.00 | 0.00 |
| 1.76 | 1.76 | 0.364 | 0.364 | 1.76 | 3.88 | 1.76 | 1.76 | 3.88 | 3.88 |
| 3.88 | 3.88 | 0.802 | 0.802 | 3.88 | 8.54 | 3.88 | 3.88 | 8.54 | 8.54 |
| 8.54 | 8.54 | 1.760 | 1.760 | 8.54 | 18.78 | 8.54 | 8.54 | 18.78 | 18.78 |
| 18.78 | 18.78 | 3.880 | 3.880 | 18.78 | 41.32 | 18.78 | 18.78 | 41.32 | 41.32 |
| 41.32 | 41.32 | 8.540 | 8.540 | 41.32 | 90.91 | 41.32 | 41.32 | 90.91 | 90.91 |

#### VV test - survival endpoint

Table 13: Minimum and maximum tested rates, median, quantile 2.5% and quantile 97.5% of the $ER_{50}$ posterior distribution - survival endpoint of the VV test of study 7

| species | min_rate | max_rate | $ER_{50}$_median | $ER_{50}$_LCI95 | $ER_{50}$_UCI95 |
|---|---|---|---|---|---|
| ALLCE | 1.760 | 41.32 | 46.641646 | 41.601541 | 75.843785 |
| AVESA | 1.760 | 41.32 | 49.473009 | 42.230466 | 88.991649 |
| BEAVA | 0.364 | 8.54 | 7.085982 | 5.837452 | 8.589002 |
| BRSNW | 0.364 | 8.54 | 9.932272 | 8.673981 | 16.546308 |
| CUMSA | 1.760 | 41.32 | 38.936929 | 32.401060 | 42.706934 |
| GLXMA | 3.880 | 90.91 | 129.998101 | 97.333796 | 294.161230 |
| HELAN | 1.760 | 41.32 | 8.807486 | 8.055439 | 10.279506 |
| LYPES | 1.760 | 41.32 | 36.152426 | 28.797941 | 47.938910 |
| TRZAW | 3.880 | 90.91 | 129.113217 | 97.423971 | 283.965460 |
| ZEAMA | 3.880 | 90.91 | 130.044482 | 97.498618 | 281.766364 |

Table 14: Censored $ER_{50}$ according to criterion 1 and the seven decision thresholds - survival endpoint of the VV test of study 7

| species | C1T0.0 | C1T0.2 | C1T0.4 | C1T0.5 | C1T0.6 | C1T0.8 | C1T1.0 |
|---|---|---|---|---|---|---|---|
| ALLCE | [41.32; Inf) | [41.32; Inf) | [41.32; Inf) | [41.32; Inf) | [41.32; Inf) | [41.32; Inf) | [41.32; Inf) |
| AVESA | [41.32; Inf) | [41.32; Inf) | [41.32; Inf) | [41.32; Inf) | [41.32; Inf) | [41.32; Inf) | [41.32; Inf) |
| BEAVA | [5.84; 8.59] | [5.84; 8.59] | [5.84; 8.59] | [5.84; 8.59] | [5.84; 8.59] | [5.84; 8.59] | [5.84; Inf) |
| BRSNW | [8.54; Inf) | [8.54; Inf) | [8.54; Inf) | [8.54; Inf) | [8.54; Inf) | [8.54; Inf) | [8.54; Inf) |
| CUMSA | [32.4; 42.71] | [32.4; 42.71] | [32.4; 42.71] | [32.4; 42.71] | [32.4; 42.71] | [32.4; 42.71] | [32.4; Inf) |
| GLXMA | [90.91; Inf) | [90.91; Inf) | [90.91; Inf) | [90.91; Inf) | [90.91; Inf) | [90.91; Inf) | [90.91; Inf) |
| HELAN | [8.06; 10.28] | [8.06; 10.28] | [8.06; 10.28] | [8.06; 10.28] | [8.06; 10.28] | [8.06; 10.28] | [8.06; 10.28] |
| LYPES | [28.8; 47.94] | [28.8; 47.94] | [28.8; 47.94] | [28.8; 47.94] | [28.8; 47.94] | [28.8; 47.94] | [28.8; Inf) |
| TRZAW | [90.91; Inf) | [90.91; Inf) | [90.91; Inf) | [90.91; Inf) | [90.91; Inf) | [90.91; Inf) | [90.91; Inf) |
| ZEAMA | [90.91; Inf) | [90.91; Inf) | [90.91; Inf) | [90.91; Inf) | [90.91; Inf) | [90.91; Inf) | [90.91; Inf) |

Table 15: Censored $ER_{50}$ according to criterion 2 and the seven decision thresholds - survival endpoint of the VV test of study 7

| species | C2T0.0 | C2T0.2 | C2T0.4 | C2T0.5 | C2T0.6 | C2T0.8 | C2T1.0 |
|---|---|---|---|---|---|---|---|
| ALLCE | [41.32; Inf) | [41.32; Inf) | [41.32; Inf) | [41.32; Inf) | [41.32; Inf) | [41.32; Inf) | [41.32; Inf) |
| AVESA | [41.32; Inf) | [41.32; Inf) | [41.32; Inf) | [41.32; Inf) | [41.32; Inf) | [41.32; Inf) | [41.32; Inf) |
| BEAVA | [5.84; 8.59] | [5.84; 8.59] | [5.84; 8.59] | [5.84; 8.59] | [5.84; 8.59] | [5.84; 8.59] | [5.84; Inf) |
| BRSNW | [8.54; Inf) | [8.54; Inf) | [8.54; Inf) | [8.54; Inf) | [8.54; Inf) | [8.54; Inf) | [8.54; Inf) |
| CUMSA | [32.4; 42.71] | [32.4; 42.71] | [32.4; 42.71] | [32.4; 42.71] | [32.4; 42.71] | [32.4; 42.71] | [32.4; Inf) |
| GLXMA | [90.91; Inf) | [90.91; Inf) | [90.91; Inf) | [90.91; Inf) | [90.91; Inf) | [90.91; Inf) | [90.91; Inf) |
| HELAN | [8.06; 10.28] | [8.06; 10.28] | [8.06; 10.28] | [8.06; 10.28] | [8.06; 10.28] | [8.06; 10.28] | [8.06; 10.28] |
| LYPES | [28.8; 47.94] | [28.8; 47.94] | [28.8; 47.94] | [28.8; 47.94] | [28.8; 47.94] | [28.8; Inf) | [28.8; Inf) |
| TRZAW | [90.91; Inf) | [90.91; Inf) | [90.91; Inf) | [90.91; Inf) | [90.91; Inf) | [90.91; Inf) | [90.91; Inf) |
| ZEAMA | [90.91; Inf) | [90.91; Inf) | [90.91; Inf) | [90.91; Inf) | [90.91; Inf) | [90.91; Inf) | [90.91; Inf) |

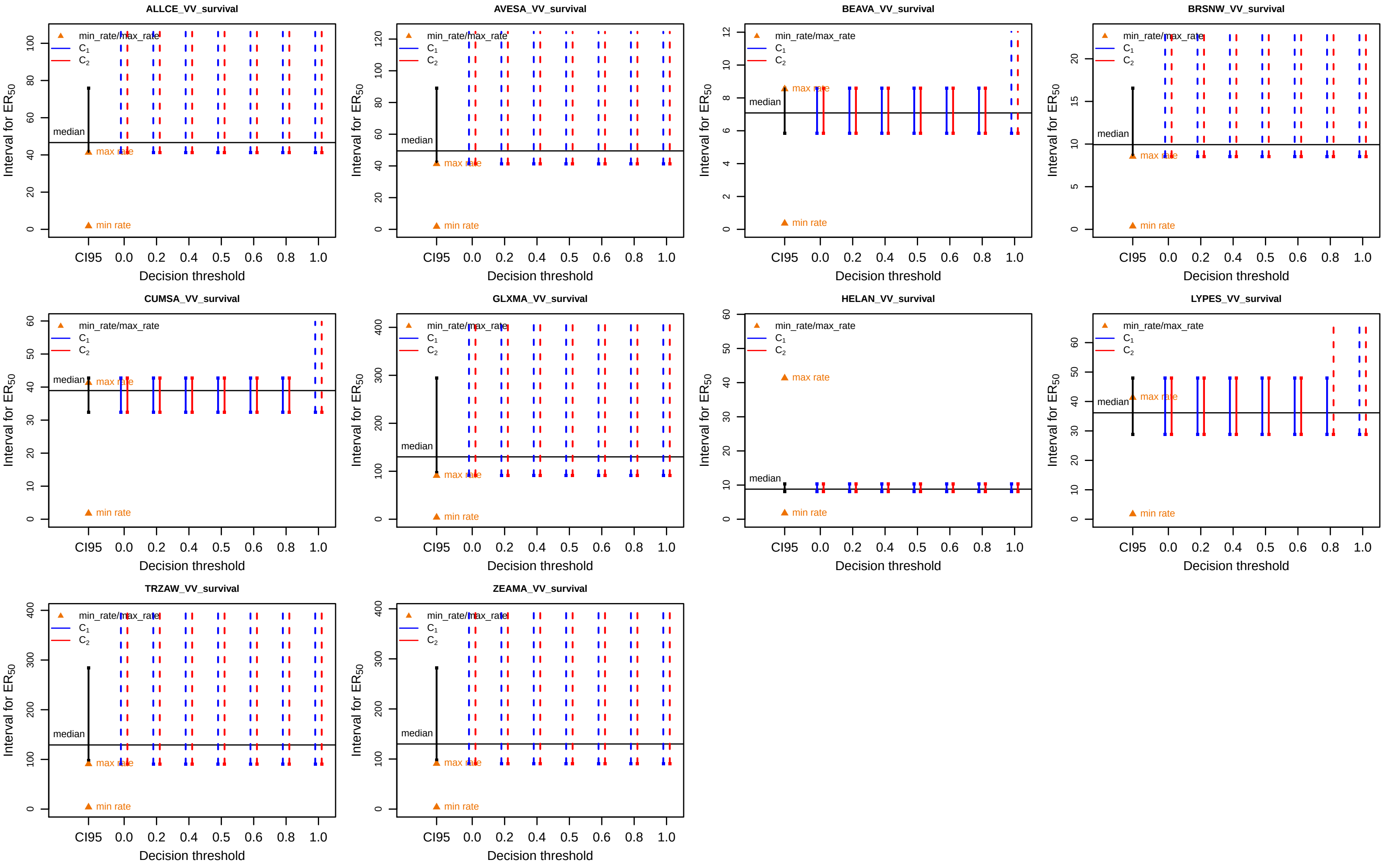


Figure 4: CI95 of $ER_{50}$ and censored $ER_{50}$ according to both criteria and the seven decision thresholds for all species - survival endpoint of the VV test of study 7. The two orange triangles stand for the lowest and the highest tested rates. Vertical segments of three different colors (black, blue, red) represent the CI95 of $ER_{50}$, the censored $ER_{50}$ according to $C_1$ and the censored $ER_{50}$ according to $C_2$, respectively; solid vertical segments are for bounded intervals while dotted vertical segments stand for right-unbounded intervals; the black horizontal line represents the median of the $ER_{50}$.

#### VV test - shoot dry weight endpoint

Table 16: Minimum and maximum tested rates, median, quantile 2.5% and quantile 97.5% of the $ER_{50}$ posterior distribution - shoot dry weight endpoint of the VV test of study 7

| species | min_rate | max_rate | $ER_{50}$_median | $ER_{50}$_LCI95 | $ER_{50}$_UCI95 |
|---|---|---|---|---|---|
| ALLCE | 1.760 | 41.32 | 37.965213 | 27.928862 | 50.457253 |
| AVESA | 1.760 | 41.32 | 29.872184 | 23.640898 | 38.034025 |
| BEAVA | 0.364 | 8.54 | 1.756003 | 1.406141 | 2.215841 |
| BRSNW | 0.364 | 8.54 | 4.065580 | 3.600104 | 4.593755 |
| CUMSA | 1.760 | 41.32 | 9.378367 | 6.943601 | 12.905670 |
| GLXMA | 3.880 | 90.91 | 8.119111 | 6.372771 | 10.193543 |
| HELAN | 1.760 | 8.54 | 5.172458 | 4.318611 | 6.279537 |
| LYPES | 1.760 | 41.32 | 4.988694 | 4.388423 | 5.633699 |
| TRZAW | 3.880 | 90.91 | 123.306567 | 94.552554 | 290.279282 |
| ZEAMA | 3.880 | 90.91 | 94.766356 | 83.961649 | 112.980031 |

Table 17: Censored $ER_{50}$ according to criterion 1 and the seven decision thresholds - shoot dry weight endpoint of the VV test of study 7

| species | C1T0.0 | C1T0.2 | C1T0.4 | C1T0.5 | C1T0.6 | C1T0.8 | C1T1.0 |
|---|---|---|---|---|---|---|---|
| ALLCE | [27.93; 50.46] | [27.93; 50.46] | [27.93; 50.46] | [27.93; 50.46] | [27.93; 50.46] | [27.93; Inf) | [27.93; Inf) |
| AVESA | [23.64; 38.03] | [23.64; 38.03] | [23.64; 38.03] | [23.64; 38.03] | [23.64; 38.03] | [23.64; 38.03] | [23.64; 38.03] |
| BEAVA | [1.41; 2.22] | [1.41; 2.22] | [1.41; 2.22] | [1.41; 2.22] | [1.41; 2.22] | [1.41; 2.22] | [1.41; 2.22] |
| BRSNW | [3.6; 4.59] | [3.6; 4.59] | [3.6; 4.59] | [3.6; 4.59] | [3.6; 4.59] | [3.6; 4.59] | [3.6; 4.59] |
| CUMSA | [6.94; 12.91] | [6.94; 12.91] | [6.94; 12.91] | [6.94; 12.91] | [6.94; 12.91] | [6.94; 12.91] | [6.94; 12.91] |
| GLXMA | [6.37; 10.19] | [6.37; 10.19] | [6.37; 10.19] | [6.37; 10.19] | [6.37; 10.19] | [6.37; 10.19] | [6.37; 10.19] |
| HELAN | [4.32; 6.28] | [4.32; 6.28] | [4.32; 6.28] | [4.32; 6.28] | [4.32; 6.28] | [4.32; 6.28] | [4.32; 6.28] |
| LYPES | [4.39; 5.63] | [4.39; 5.63] | [4.39; 5.63] | [4.39; 5.63] | [4.39; 5.63] | [4.39; 5.63] | [4.39; 5.63] |
| TRZAW | [90.91; Inf) | [90.91; Inf) | [90.91; Inf) | [90.91; Inf) | [90.91; Inf) | [90.91; Inf) | [90.91; Inf) |
| ZEAMA | [83.96; 112.98] | [83.96; 112.98] | [83.96; Inf) | [83.96; Inf) | [83.96; Inf) | [83.96; Inf) | [83.96; Inf) |

Table 18: Censored $ER_{50}$ according to criterion 2 and the seven decision thresholds - shoot dry weight endpoint of the VV test of study 7

| species | C2T0.0 | C2T0.2 | C2T0.4 | C2T0.5 | C2T0.6 | C2T0.8 | C2T1.0 |
|---|---|---|---|---|---|---|---|
| ALLCE | [27.93; 50.46] | [27.93; 50.46] | [27.93; 50.46] | [27.93; 50.46] | [27.93; Inf) | [27.93; Inf) | [27.93; Inf) |
| AVESA | [23.64; 38.03] | [23.64; 38.03] | [23.64; 38.03] | [23.64; 38.03] | [23.64; 38.03] | [23.64; 38.03] | [23.64; 38.03] |
| BEAVA | [1.41; 2.22] | [1.41; 2.22] | [1.41; 2.22] | [1.41; 2.22] | [1.41; 2.22] | [1.41; 2.22] | [1.41; 2.22] |
| BRSNW | [3.6; 4.59] | [3.6; 4.59] | [3.6; 4.59] | [3.6; 4.59] | [3.6; 4.59] | [3.6; 4.59] | [3.6; 4.59] |
| CUMSA | [6.94; 12.91] | [6.94; 12.91] | [6.94; 12.91] | [6.94; 12.91] | [6.94; 12.91] | [6.94; 12.91] | [6.94; 12.91] |
| GLXMA | [6.37; 10.19] | [6.37; 10.19] | [6.37; 10.19] | [6.37; 10.19] | [6.37; 10.19] | [6.37; 10.19] | [6.37; 10.19] |
| HELAN | [4.32; 6.28] | [4.32; 6.28] | [4.32; 6.28] | [4.32; 6.28] | [4.32; 6.28] | [4.32; 6.28] | [4.32; 6.28] |
| LYPES | [4.39; 5.63] | [4.39; 5.63] | [4.39; 5.63] | [4.39; 5.63] | [4.39; 5.63] | [4.39; 5.63] | [4.39; 5.63] |
| TRZAW | [90.91; Inf) | [90.91; Inf) | [90.91; Inf) | [90.91; Inf) | [90.91; Inf) | [90.91; Inf) | [90.91; Inf) |
| ZEAMA | [83.96; 112.98] | [83.96; 112.98] | [83.96; Inf) | [83.96; Inf) | [83.96; Inf) | [83.96; Inf) | [83.96; Inf) |

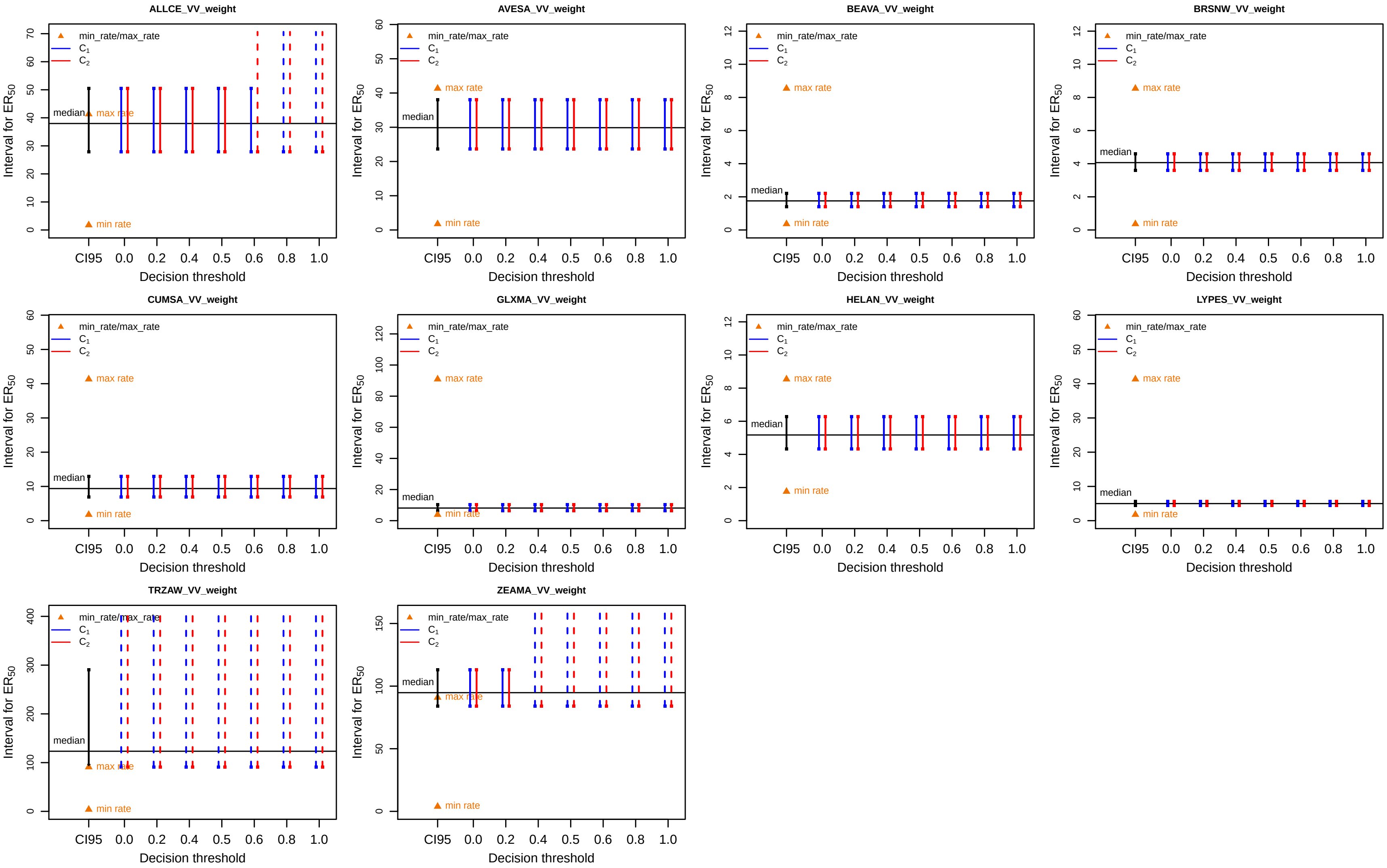


Figure 5: CI95 of $ER_{50}$ and censored $ER_{50}$ according to both criteria and the seven decision thresholds for all species - shoot dry weight endpoint of the VV test of study 7. The two orange triangles stand for the lowest and the highest tested rates. Vertical segments of three different colors (black, blue, red) represent the CI95 of $ER_{50}$, the censored $ER_{50}$ according to $C_1$ and the censored $ER_{50}$ according to $C_2$, respectively; solid vertical segments are for bounded intervals while dotted vertical segments stand for right-unbounded intervals; the black horizontal line represents the median of the $ER_{50}$.
