## Supplementary material for "How to account for the uncertainty from standard toxicity tests in species sensitivity distributions: an example in non-target plants": Archive with reports on studies: report_SE_emergence.pdf

### Dose-response analyses

Study 7

Seedling Emergence test - emergence endpoint

25 June 2020

Contact:

---

This is a report which provides results on all performed dose-response analyses for the emergence endpoint of the Seedling Emergence test for study 7.

---

#### Data set: ALLCE_SE_emergence

Table 1: Summary of parameter estimates for ALLCE_SE_emergence data set

| Parameter | median | Q2.5 | Q97.5 |
|---|---|---|---|
| b | 0.856 | 0.425 | 2.530 |
| d | 0.794 | 0.714 | 0.870 |
| e | 97.060 | 45.114 | 334.493 |

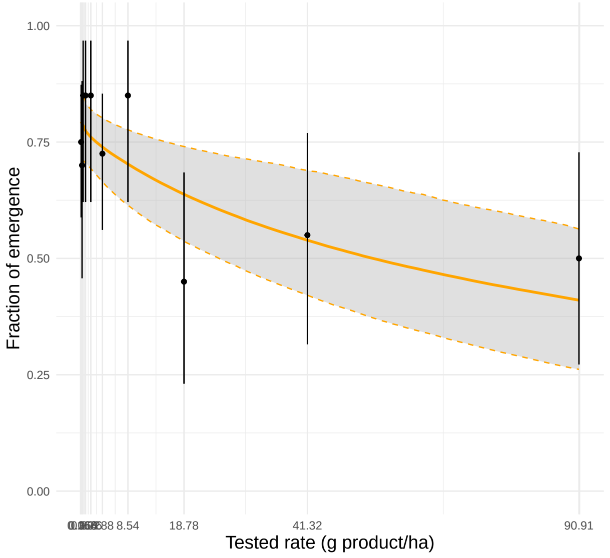

(a) Dose-response curve

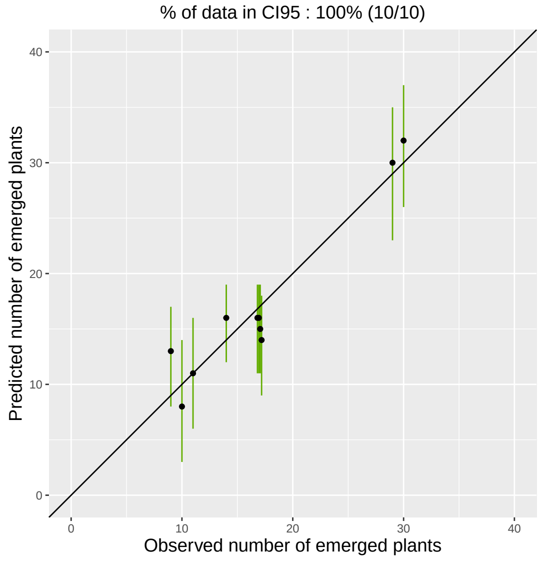

(b) Posterior predictive check (PPC)

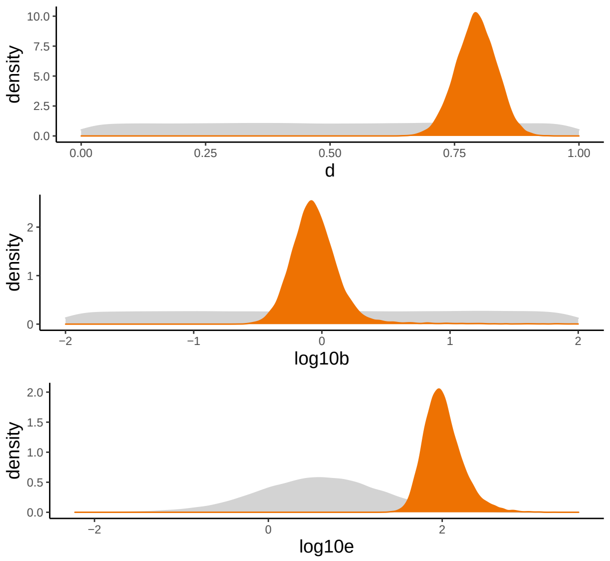

(c) Priors and posteriors

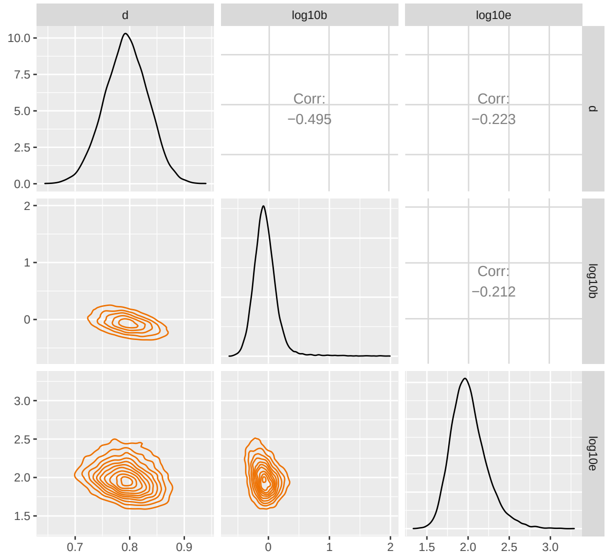

(d) Correlations between parameters

Figure 1: Dose-response curve (a), PPC (b), prior and posterior distributions (c) and correlations between parameters (d).

#### Data set: AVESA_SE_emergence

Table 2: Summary of parameter estimates for AVESA_SE_emergence data set

| Parameter | median | Q2.5 | Q97.5 |
|---|---|---|---|
| b | 22.370 | 3.199 | 93.202 |
| d | 0.950 | 0.899 | 0.981 |
| e | 122.905 | 94.926 | 261.252 |

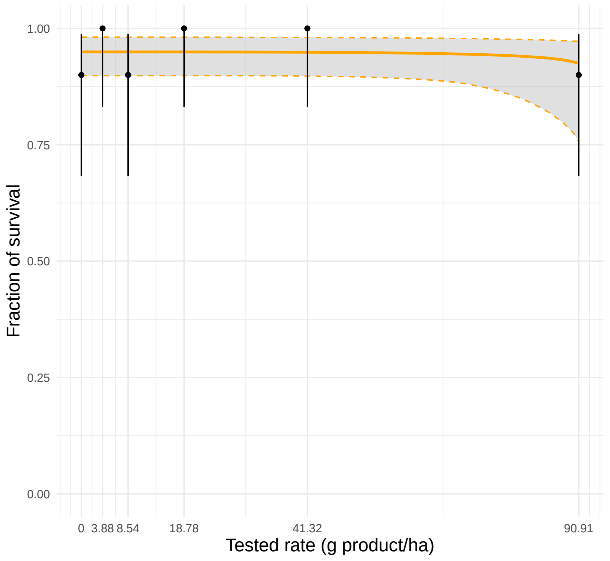

(a) Dose-response curve

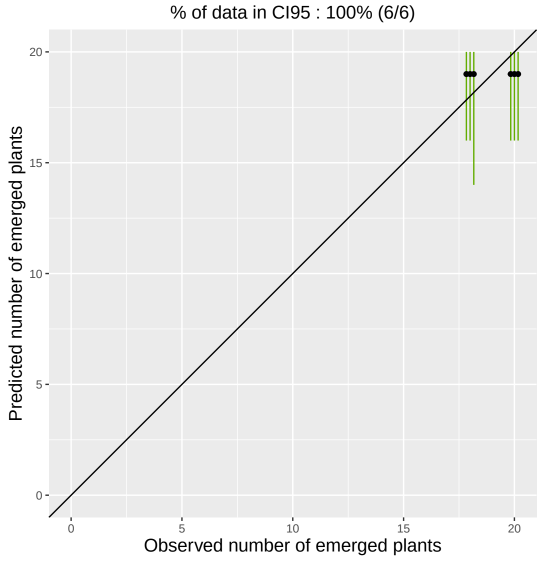

(b) Posterior predictive check (PPC)

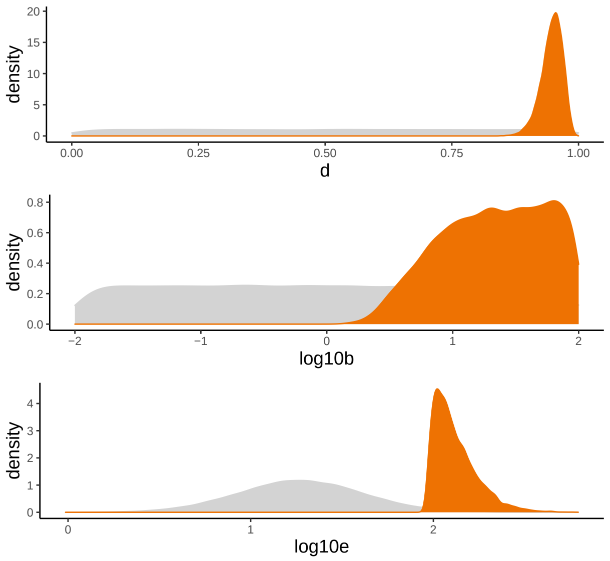

(c) Priors and posteriors

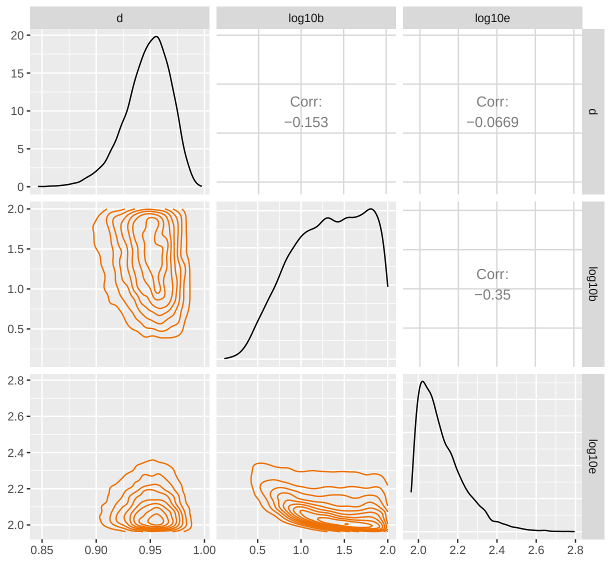

(d) Correlations between parameters

Figure 2: Dose-response curve (a), PPC (b), prior and posterior distributions (c) and correlations between parameters (d).

### Data set: BEAVA_SE_emergence

Table 3: Summary of parameter estimates (parameter d is set to 1) for BEAVA_SE_emergence data set

| Parameter | median | Q2.5 | Q97.5 |
|---|---|---|---|
| b | 1.016 | 0.668 | 1.552 |
| e | 445.002 | 186.973 | 1494.536 |

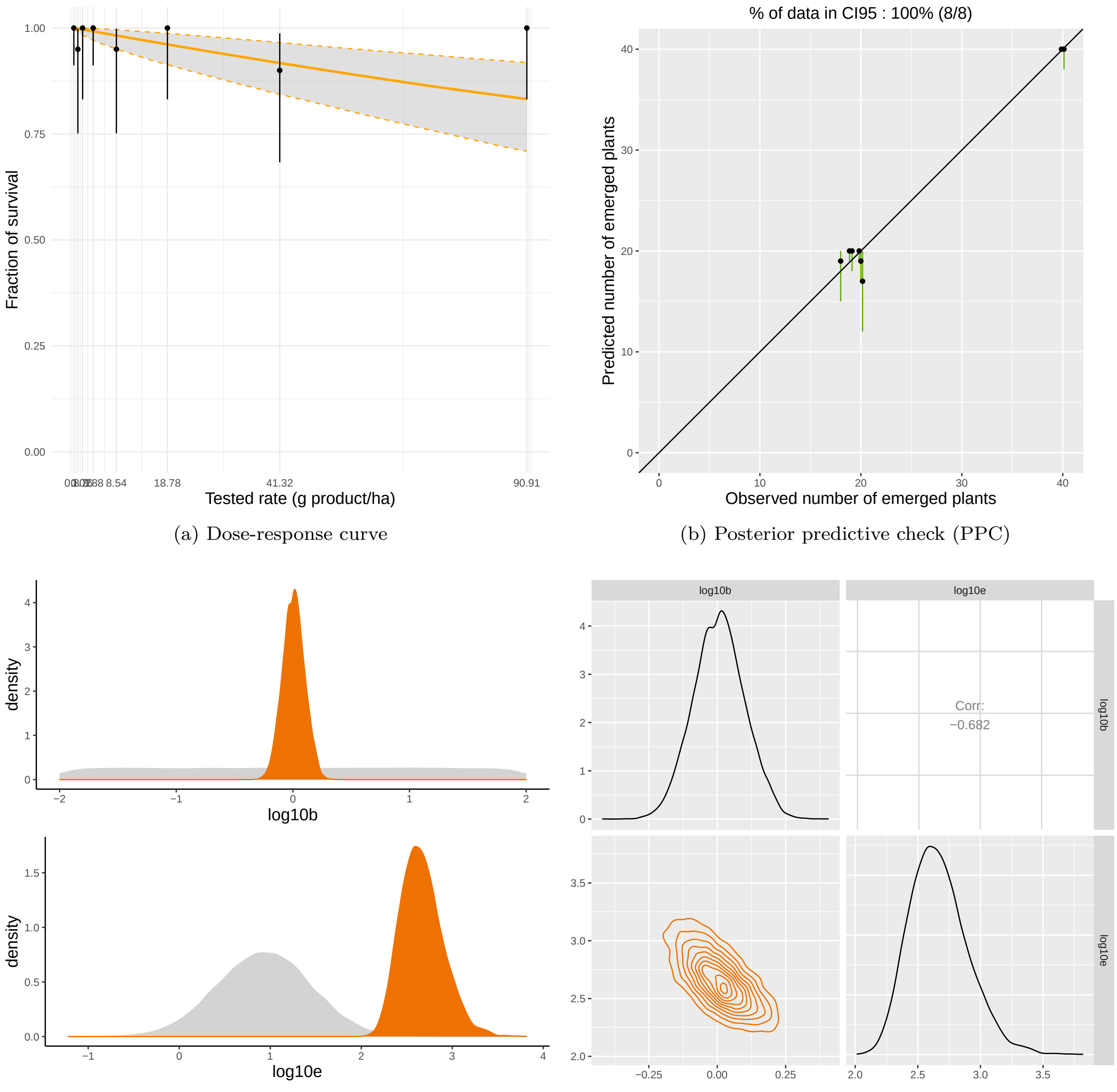

(a) Dose-response curve

(b) Posterior predictive check (PPC)

(c) Priors and posteriors

(d) Correlations between parameters

Figure 3: Dose-response curve (a), PPC (b), prior and posterior distributions (c) and correlations between parameters (d).

#### Data set: BRSNW_SE_emergence

Table 4: Summary of parameter estimates (parameter d is set to 1) for BRSNW_SE_emergence data set

| Parameter | median | Q2.5 | Q97.5 |
|---|---|---|---|
| b | 1.324 | 0.793 | 2.046 |
| e | 95.133 | 62.532 | 181.623 |

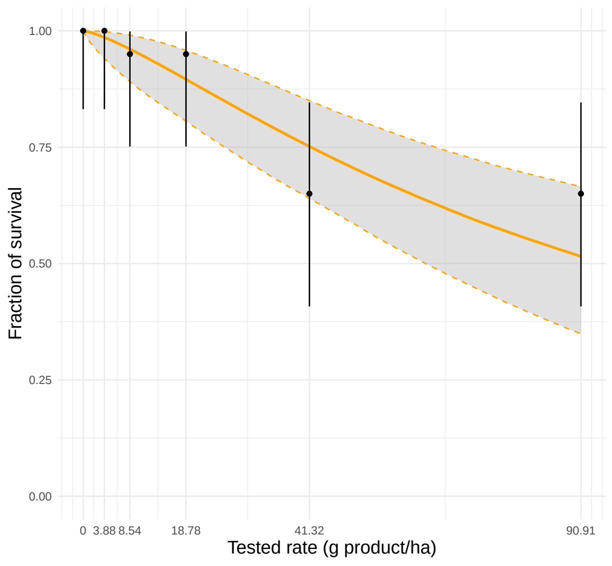

(a) Dose-response curve

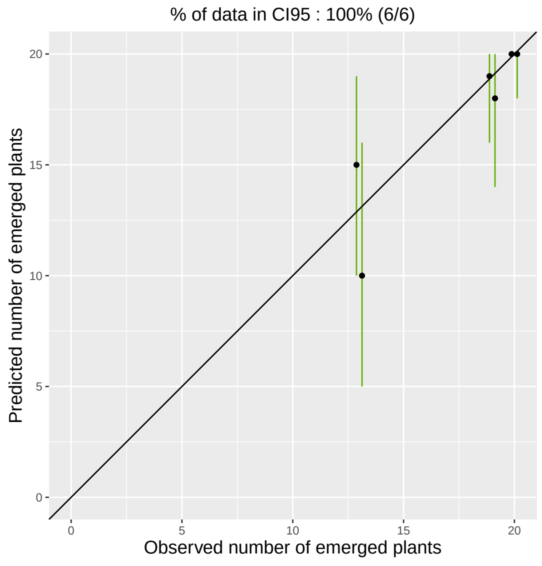

(b) Posterior predictive check (PPC)

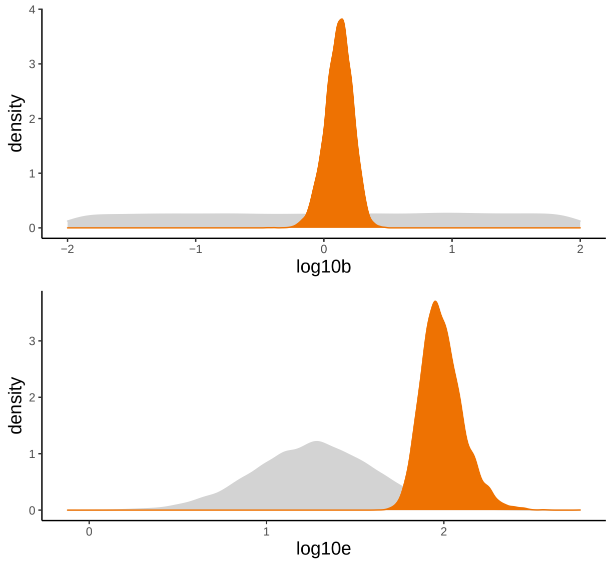

(c) Priors and posteriors

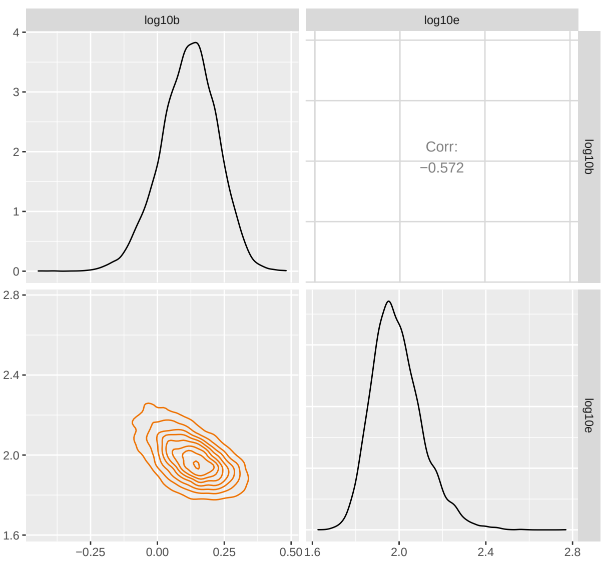

(d) Correlations between parameters

Figure 4: Dose-response curve (a), PPC (b), prior and posterior distributions (c) and correlations between parameters (d).

#### Data set: CUMSA_SE_emergence

Table 5: Summary of parameter estimates for CUMSA_SE_emergence data set

| Parameter | median | Q2.5 | Q97.5 |
|---|---|---|---|
| b | 26.439 | 3.214 | 93.735 |
| d | 0.923 | 0.865 | 0.963 |
| e | 126.036 | 95.624 | 275.409 |

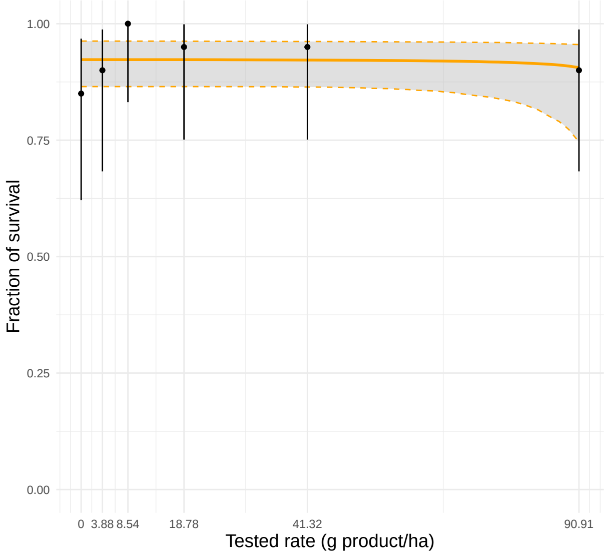

(a) Dose-response curve

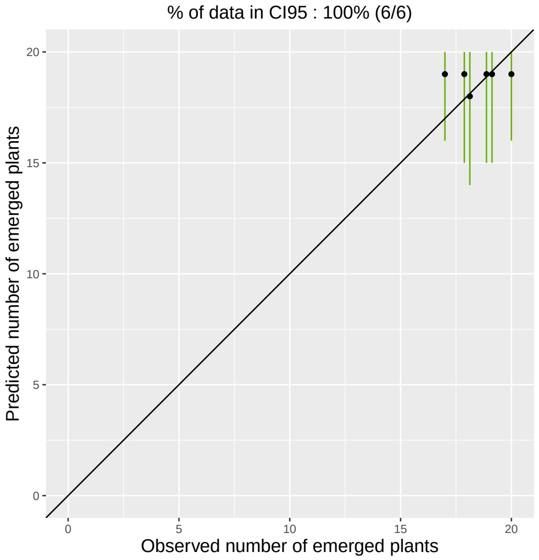

(b) Posterior predictive check (PPC)

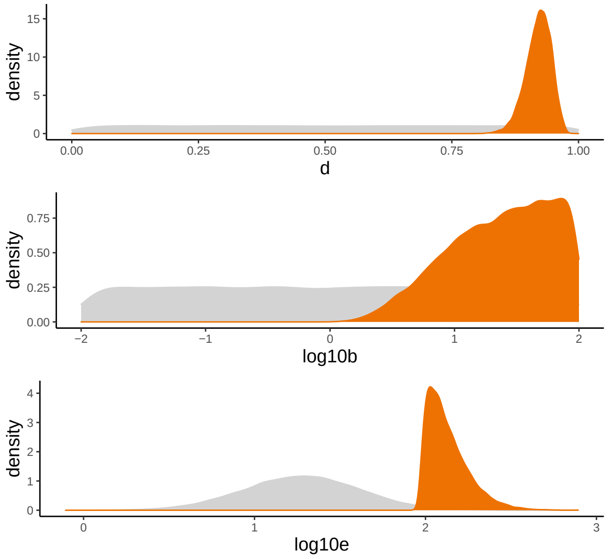

(c) Priors and posteriors

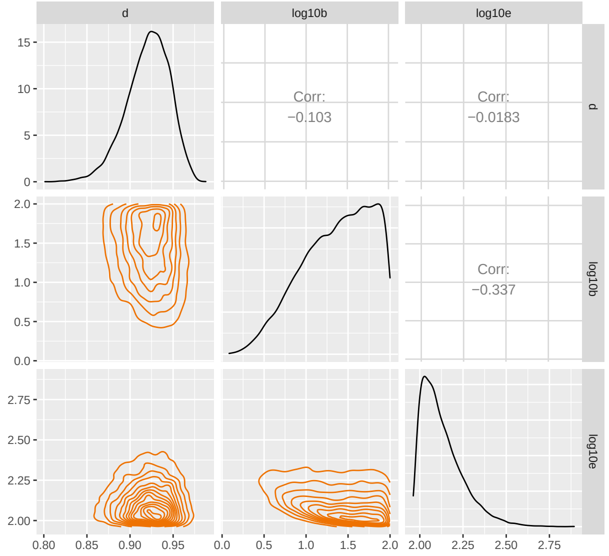

(d) Correlations between parameters

Figure 5: Dose-response curve (a), PPC (b), prior and posterior distributions (c) and correlations between parameters (d).

#### Data set: GLXMA_SE_emergence

Table 6: Summary of parameter estimates (parameter d is set to 1) for GLXMA_SE_emergence data set

| Parameter | median | Q2.5 | Q97.5 |
|---|---|---|---|
| b | 1.370 | 0.863 | 2.175 |
| e | 265.741 | 145.006 | 634.716 |

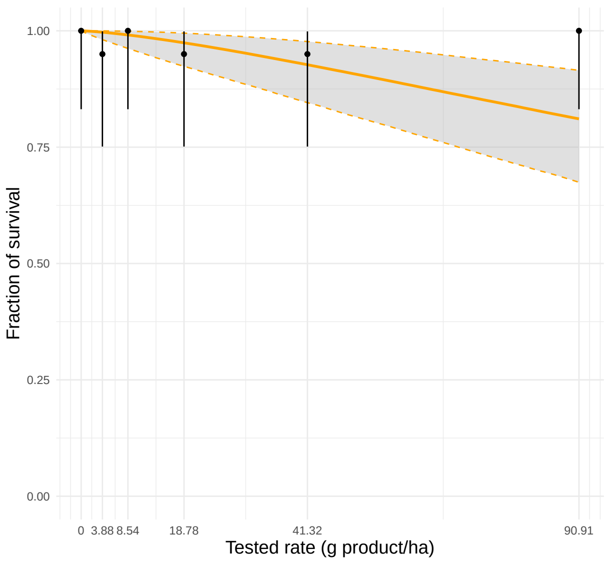

(a) Dose-response curve

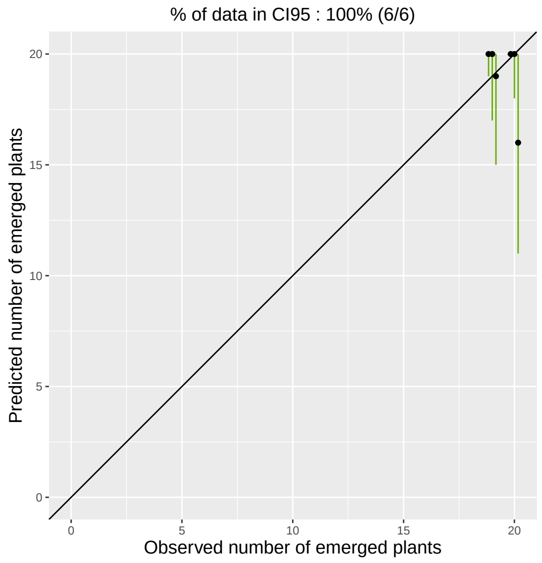

(b) Posterior predictive check (PPC)

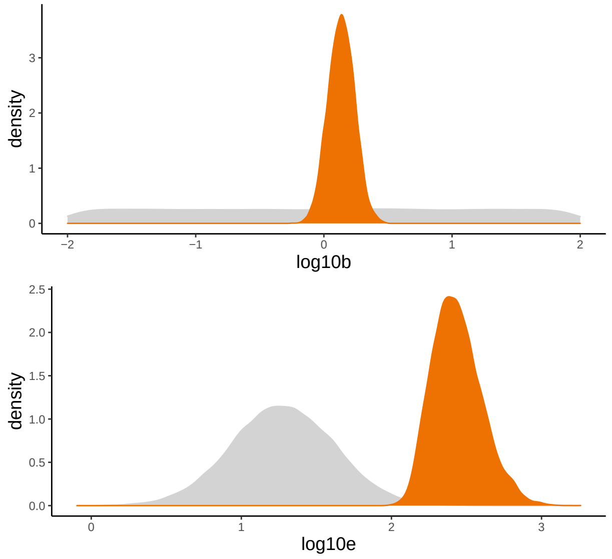

(c) Priors and posteriors

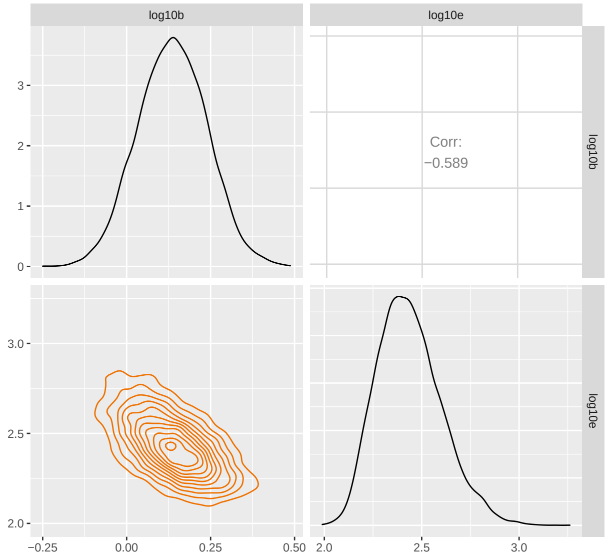

(d) Correlations between parameters

Figure 6: Dose-response curve (a), PPC (b), prior and posterior distributions (c) and correlations between parameters (d).

#### Data set: HELAN_SE_emergence

Table 7: Summary of parameter estimates (parameter d is set to 1) for HELAN_SE_emergence data set

| Parameter | median | Q2.5 | Q97.5 |
|---|---|---|---|
| b | 1.215 | 0.773 | 1.896 |
| e | 264.951 | 140.472 | 626.001 |

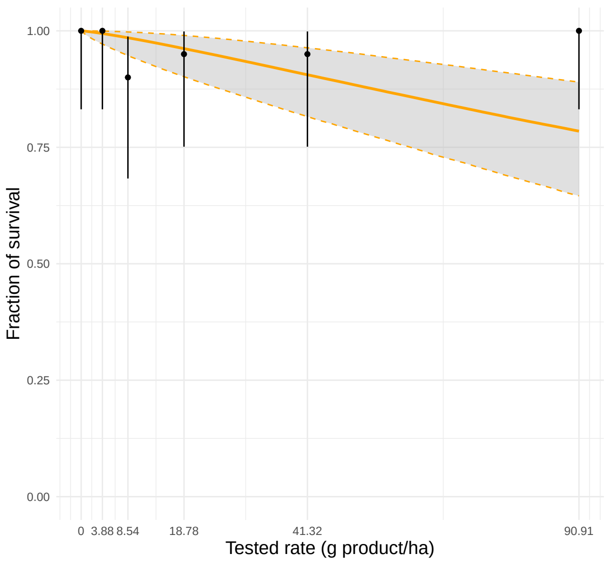

(a) Dose-response curve

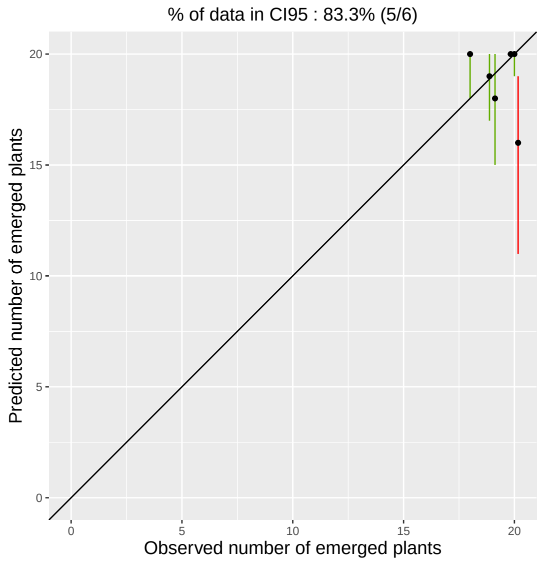

(b) Posterior predictive check (PPC)

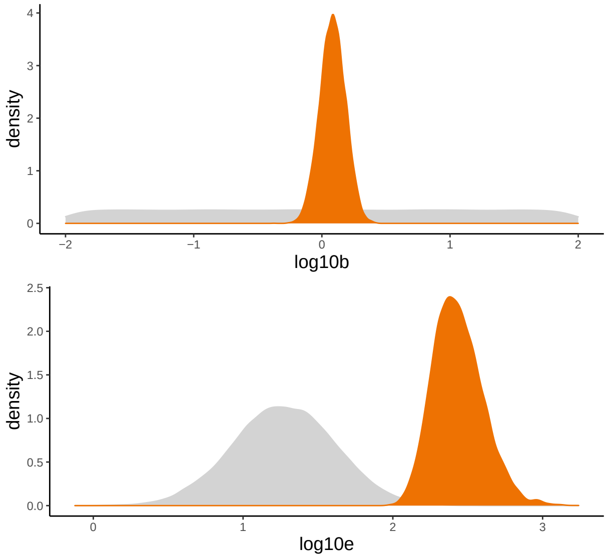

(c) Priors and posteriors

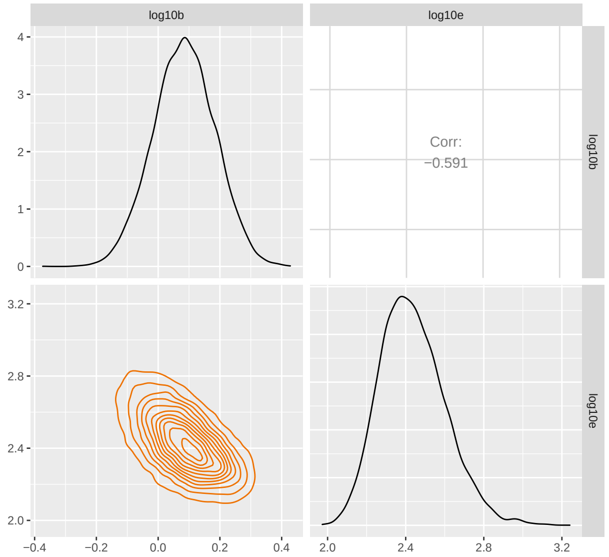

(d) Correlations between parameters

Figure 7: Dose-response curve (a), PPC (b), prior and posterior distributions (c) and correlations between parameters (d).

#### Data set: LYPES_SE_emergence

Table 8: Summary of parameter estimates for LYPES_SE_emergence data set

| Parameter | median | Q2.5 | Q97.5 |
|---|---|---|---|
| b | 29.699 | 3.190 | 94.289 |
| d | 0.946 | 0.896 | 0.978 |
| e | 128.145 | 96.391 | 291.520 |

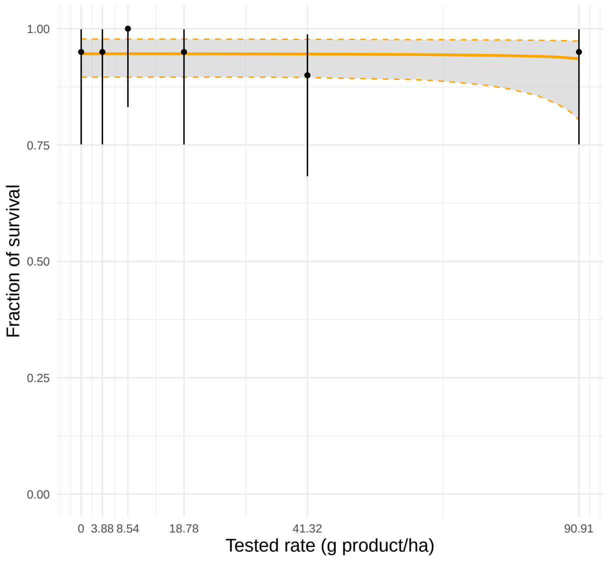

(a) Dose-response curve

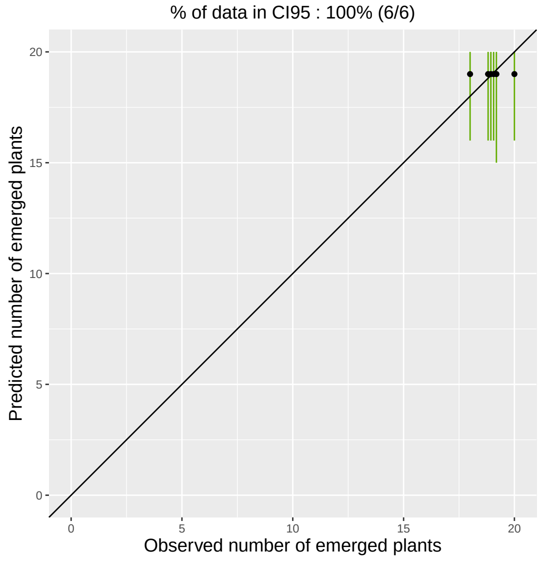

(b) Posterior predictive check (PPC)

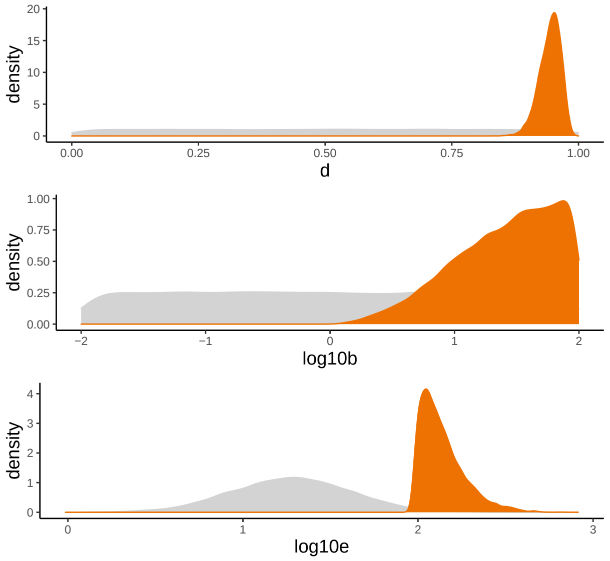

(c) Priors and posteriors

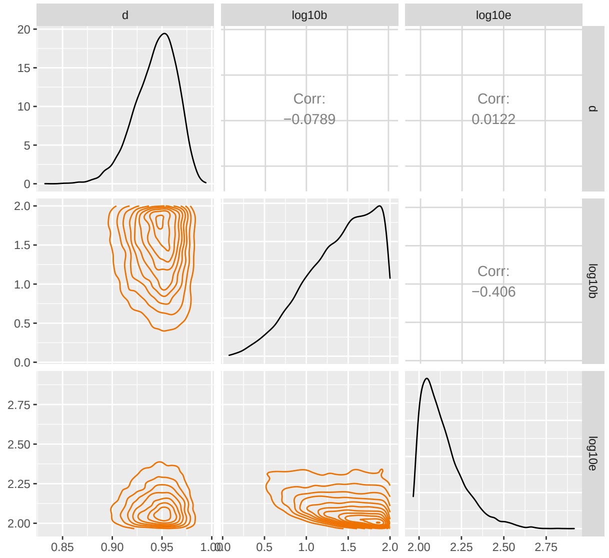

(d) Correlations between parameters

Figure 8: Dose-response curve (a), PPC (b), prior and posterior distributions (c) and correlations between parameters (d).

#### Data set: TRZAW_SE_emergence

Table 9: Summary of parameter estimates (parameter d is set to 1) for TRZAW_SE_emergence data set

| Parameter | median | Q2.5 | Q97.5 |
|---|---|---|---|
| b | 3.009 | 1.510 | 6.399 |
| e | 169.382 | 113.753 | 358.250 |

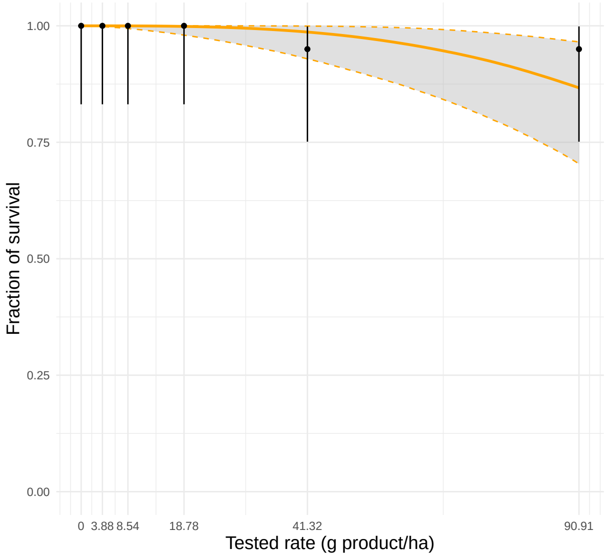

(a) Dose-response curve

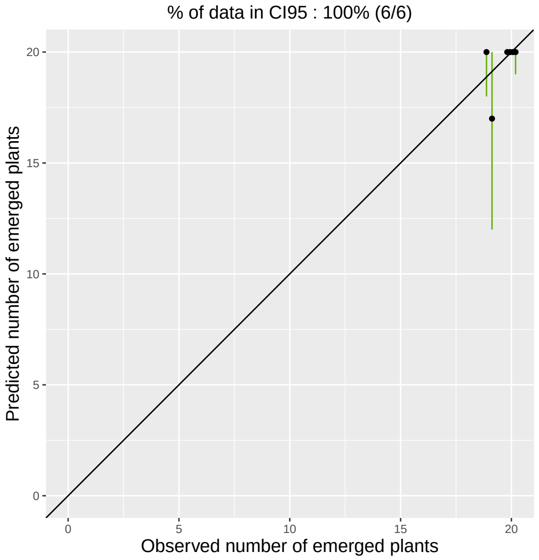

(b) Posterior predictive check (PPC)

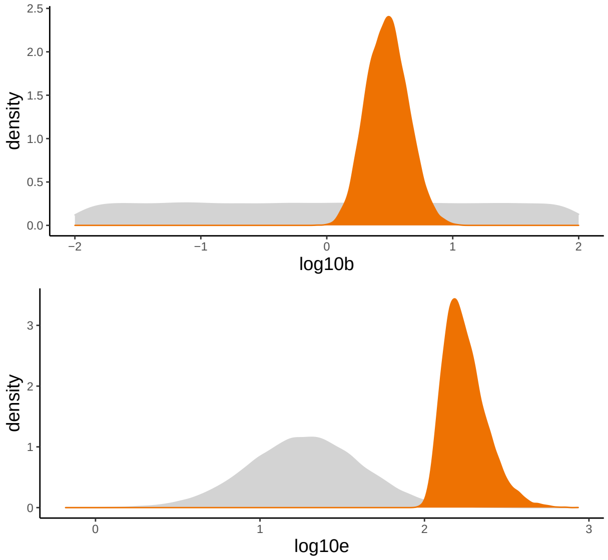

(c) Priors and posteriors

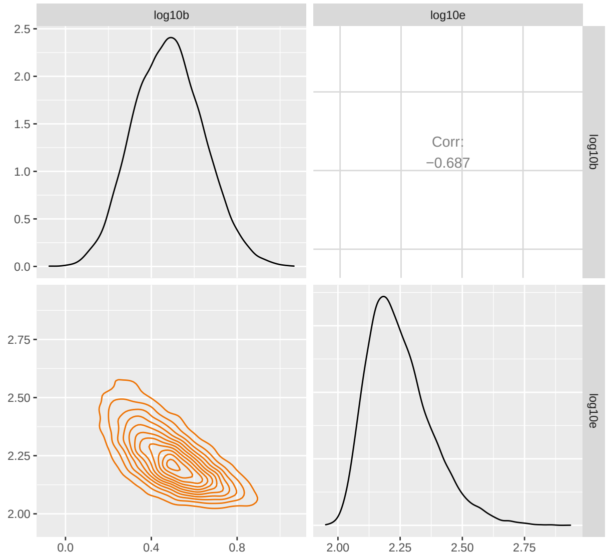

(d) Correlations between parameters

Figure 9: Dose-response curve (a), PPC (b), prior and posterior distributions (c) and correlations between parameters (d).

#### Data set: ZEAMA_SE_emergence

Table 10: Summary of parameter estimates for ZEAMA_SE_emergence data set

| Parameter | median | Q2.5 | Q97.5 |
|---|---|---|---|
| b | 34.013 | 4.981 | 95.444 |
| d | 0.970 | 0.929 | 0.991 |
| e | 131.390 | 97.146 | 294.996 |

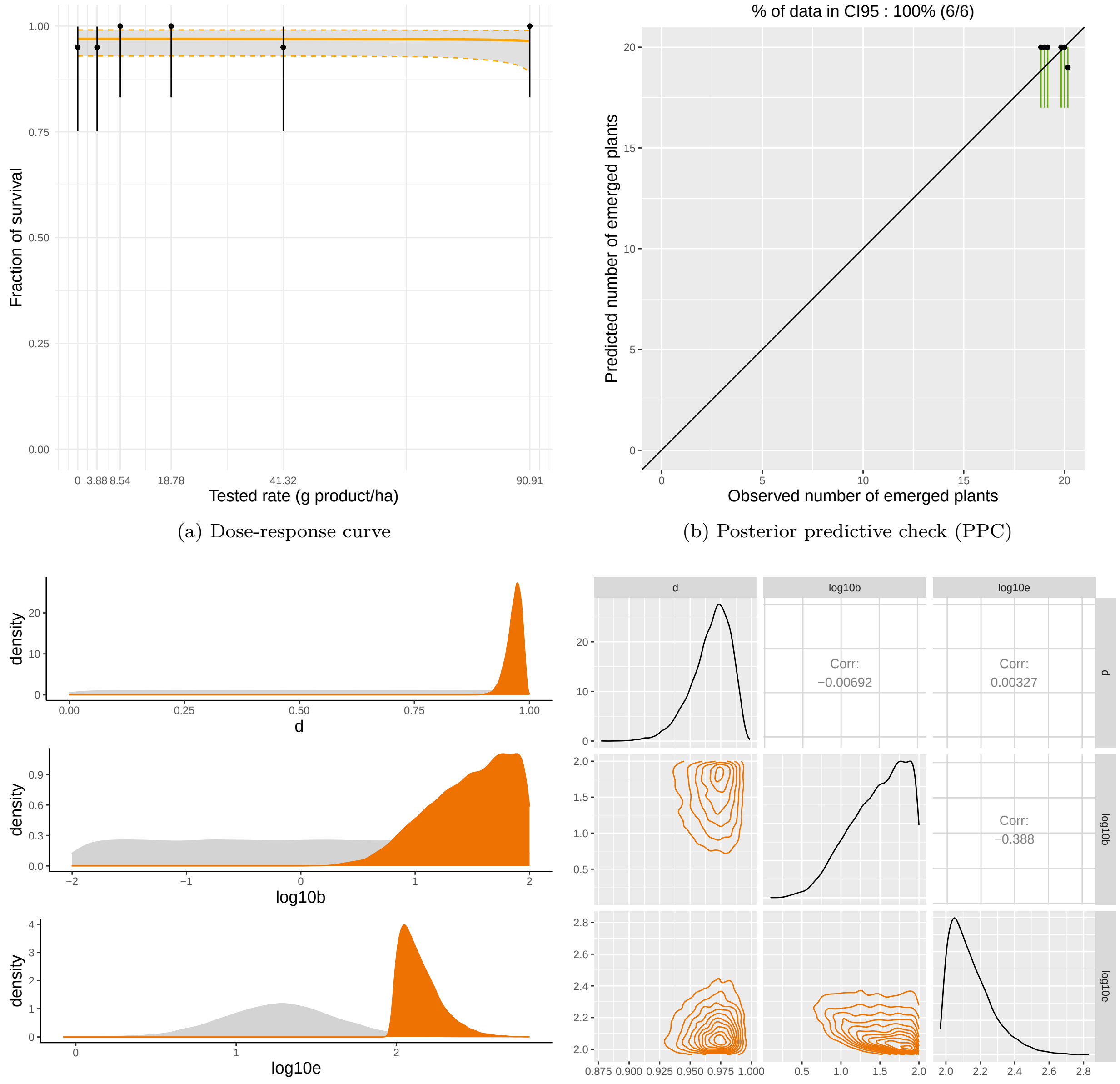

(a) Dose-response curve

(b) Posterior predictive check (PPC)

(c) Priors and posteriors

(d) Correlations between parameters

Figure 10: Dose-response curve (a), PPC (b), prior and posterior distributions (c) and correlations between parameters (d).
