## Supplementary material for "How to account for the uncertainty from standard toxicity tests in species sensitivity distributions: an example in non-target plants": Archive with reports on studies: report_SE_survival.pdf

### Dose-response analyses

Study 7

Seedling Emergence test - survival endpoint

25 June 2020

Contact:

---

This is a report which provides results on all performed dose-response analyses for the survival endpoint of the Seedling Emergence test for study 7.

---

### Data set: ALLCE_SE_survival

Table 1: Summary of parameter estimates (parameter d is set to 1) for ALLCE_SE_survival data set

| Parameter | median | Q2.5 | Q97.5 |
|---|---|---|---|
| b | 2.206 | 1.514 | 3.101 |
| e | 2.690 | 2.058 | 3.462 |

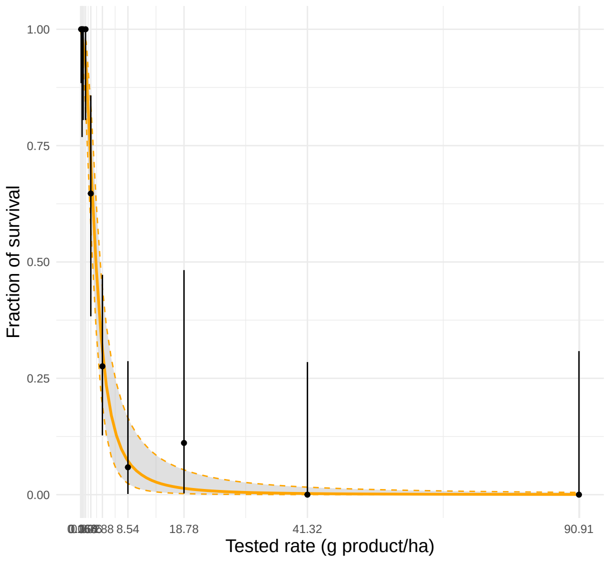

(a) Dose-response curve

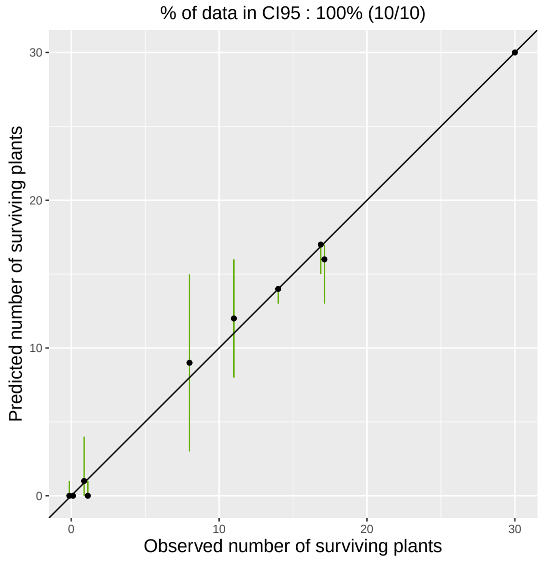

(b) Posterior predictive check (PPC)

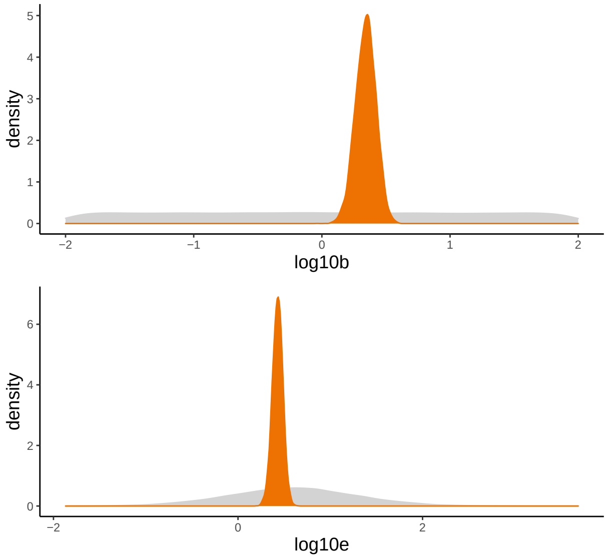

(c) Priors and posteriors

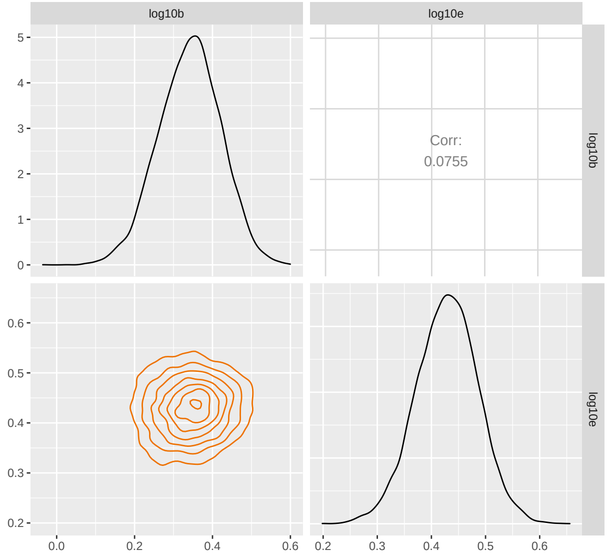

(d) Correlations between parameters

Figure 1: Dose-response curve (a), PPC (b), prior and posterior distributions (c) and correlations between parameters (d).

#### Data set: AVESA_SE_survival

Table 2: Summary of parameter estimates (parameter d is set to 1) for AVESA_SE_survival data set

| Parameter | median | Q2.5 | Q97.5 |
|---|---|---|---|
| b | 1.792 | 1.042 | 2.937 |
| e | 97.442 | 68.021 | 172.494 |

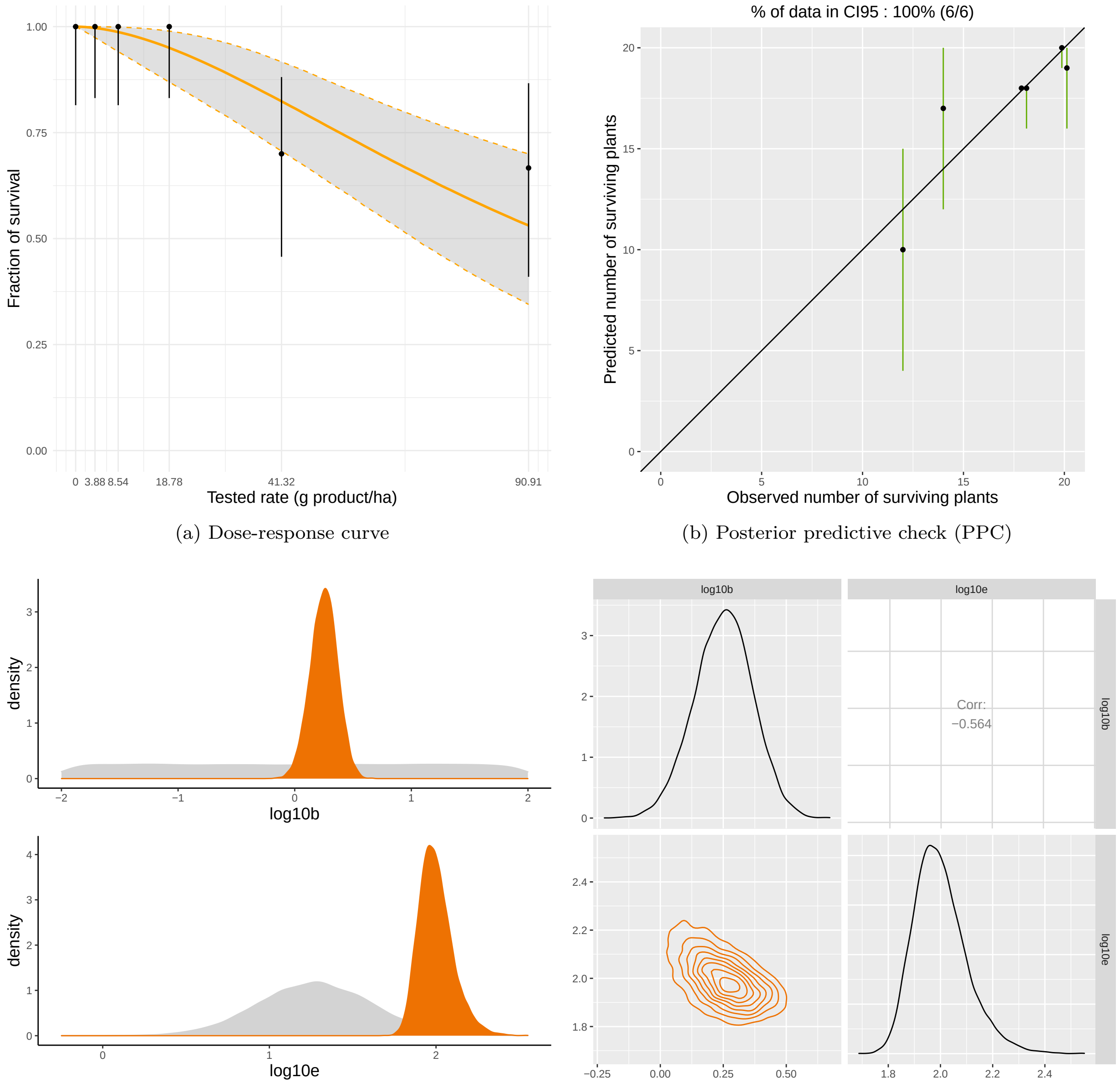

(a) Dose-response curve (b) Posterior predictive check (PPC)

(c) Priors and posteriors (d) Correlations between parameters

Figure 2: Dose-response curve (a), PPC (b), prior and posterior distributions (c) and correlations between parameters (d).

#### Data set: BEAVA_SE_survival

Table 3: Summary of parameter estimates (parameter d is set to 1) for BEAVA_SE_survival data set

| Parameter | median | Q2.5 | Q97.5 |
|---|---|---|---|
| b | 2.809 | 2.004 | 3.924 |
| e | 13.937 | 11.068 | 17.639 |

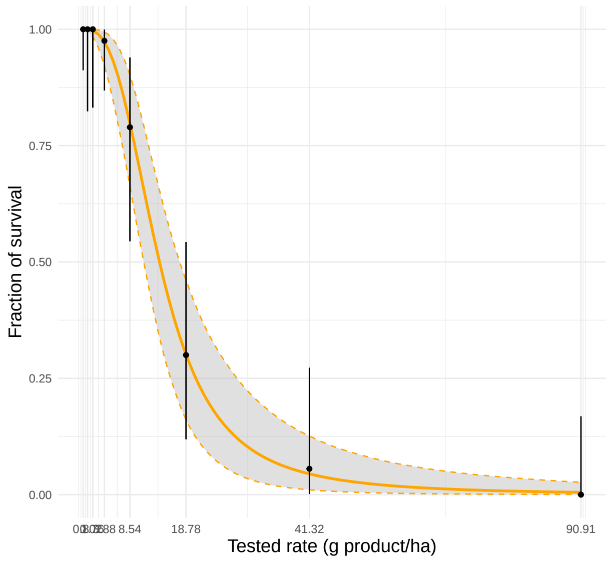

(a) Dose-response curve

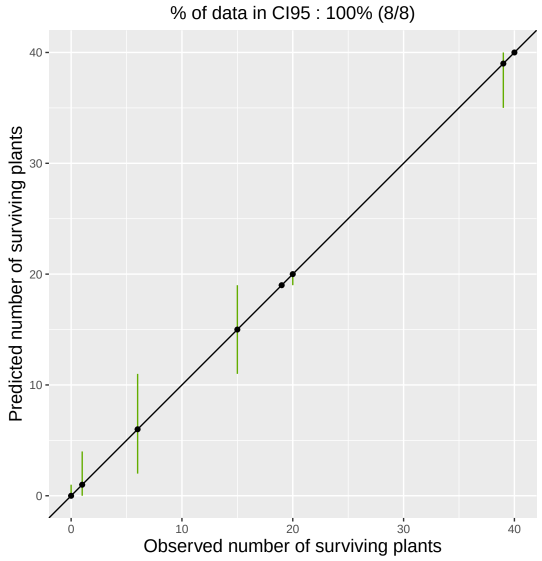

(b) Posterior predictive check (PPC)

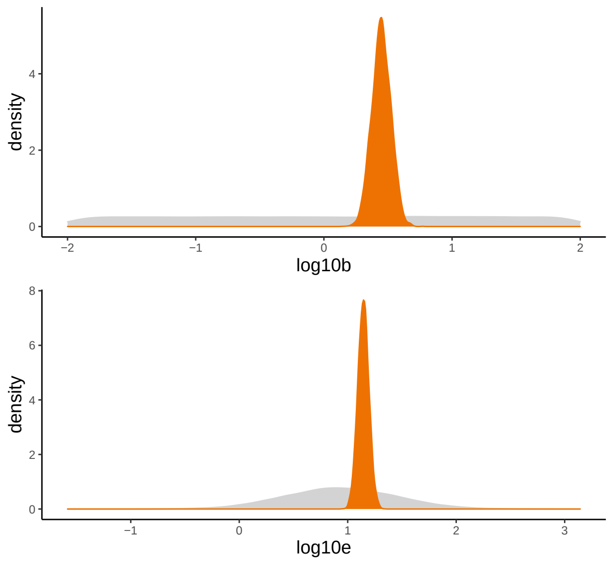

(c) Priors and posteriors

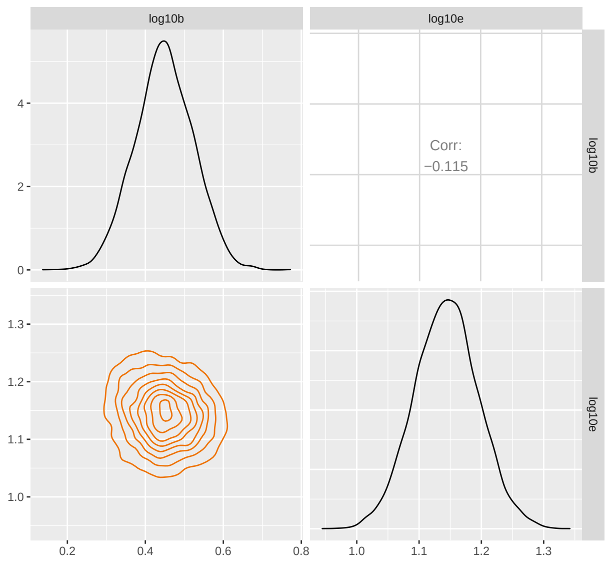

(d) Correlations between parameters

Figure 3: Dose-response curve (a), PPC (b), prior and posterior distributions (c) and correlations between parameters (d).

#### Data set: BRSNW_SE_survival

Table 4: Summary of parameter estimates (parameter d is set to 1) for BRSNW_SE_survival data set

| Parameter | median | Q2.5 | Q97.5 |
|---|---|---|---|
| b | 2.321 | 1.382 | 3.714 |
| e | 56.653 | 41.742 | 83.639 |

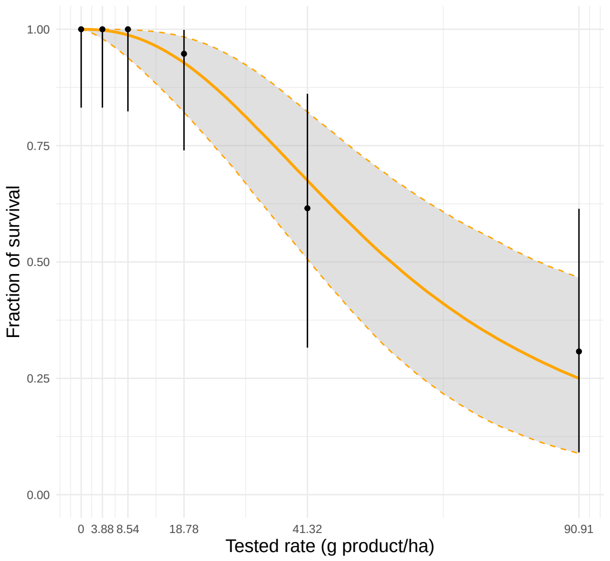

(a) Dose-response curve

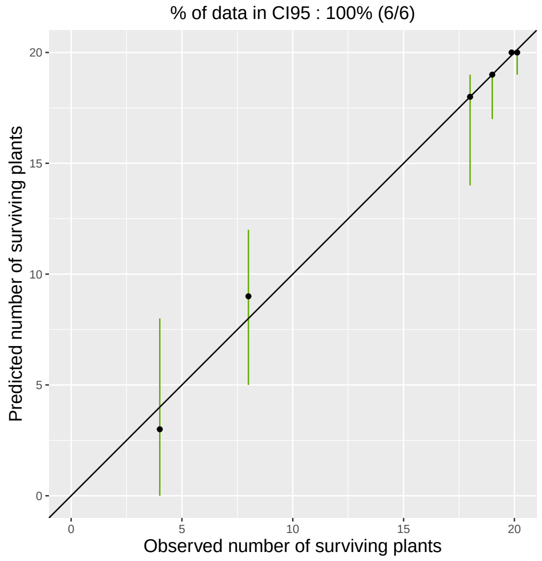

(b) Posterior predictive check (PPC)

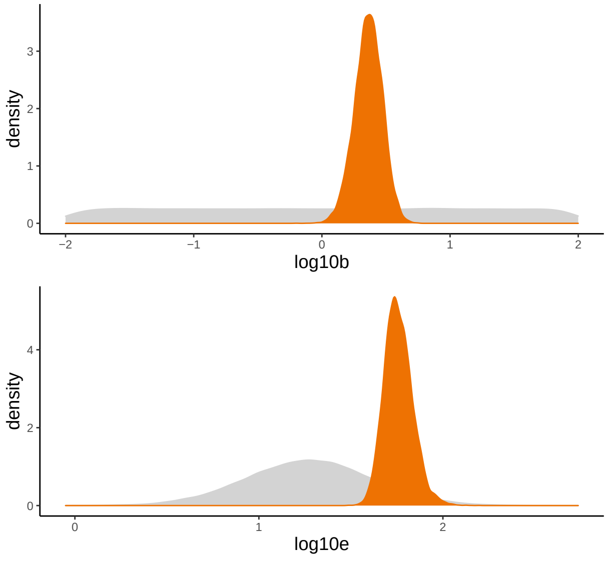

(c) Priors and posteriors

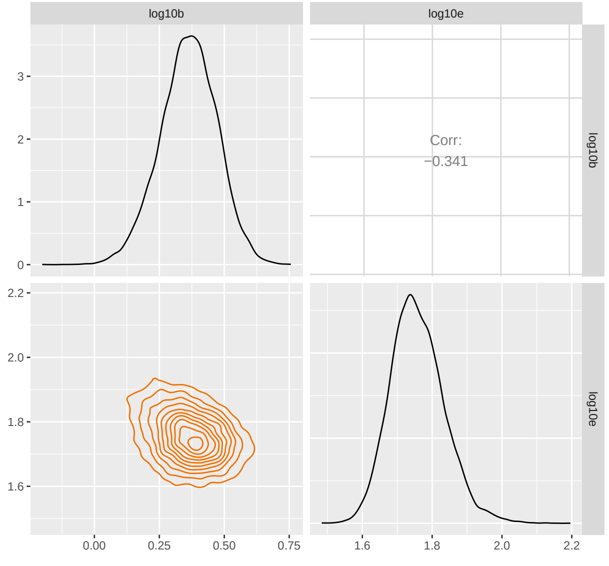

(d) Correlations between parameters

Figure 4: Dose-response curve (a), PPC (b), prior and posterior distributions (c) and correlations between parameters (d).

#### Data set: CUMSA_SE_survival

Table 5: Summary of parameter estimates (parameter d is set to 1) for CUMSA_SE_survival data set

| Parameter | median | Q2.5 | Q97.5 |
|---|---|---|---|
| b | 1.559 | 0.924 | 2.623 |
| e | 178.797 | 107.109 | 395.707 |

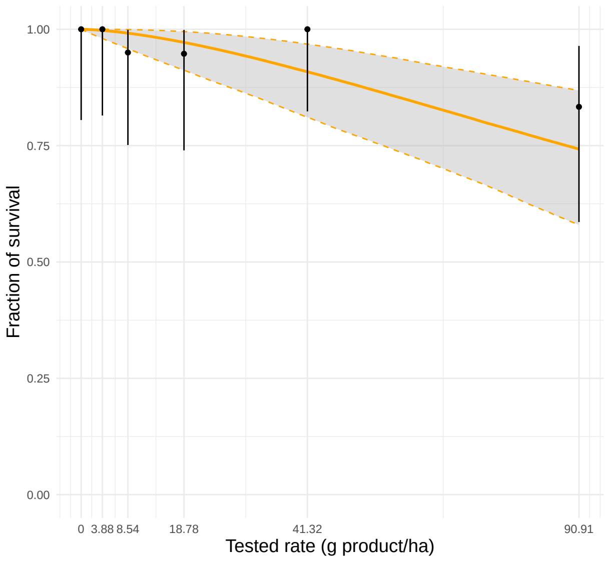

(a) Dose-response curve

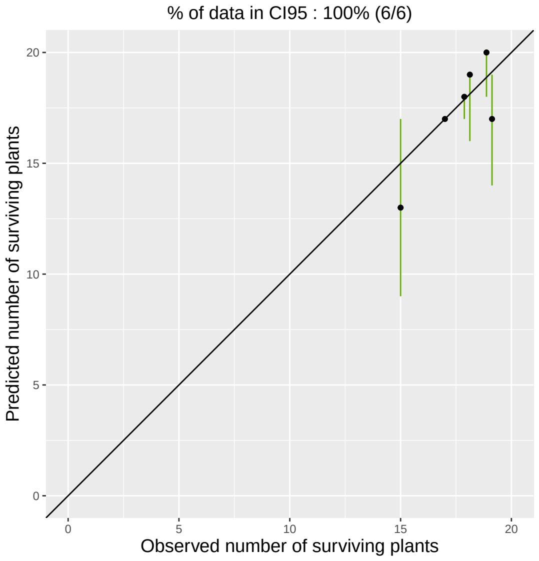

(b) Posterior predictive check (PPC)

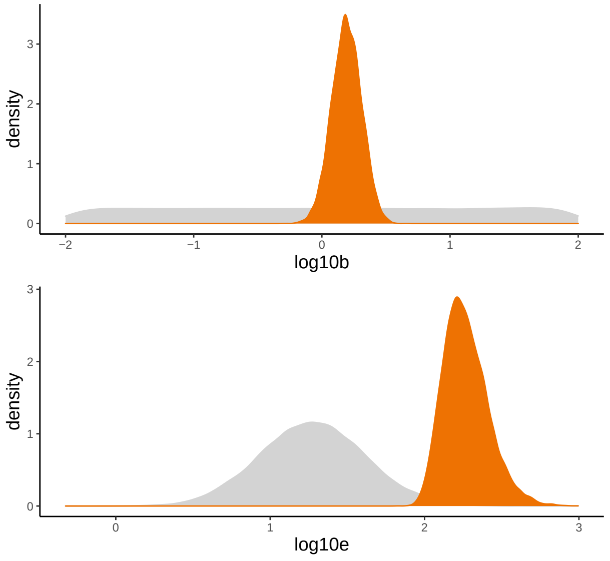

(c) Priors and posteriors

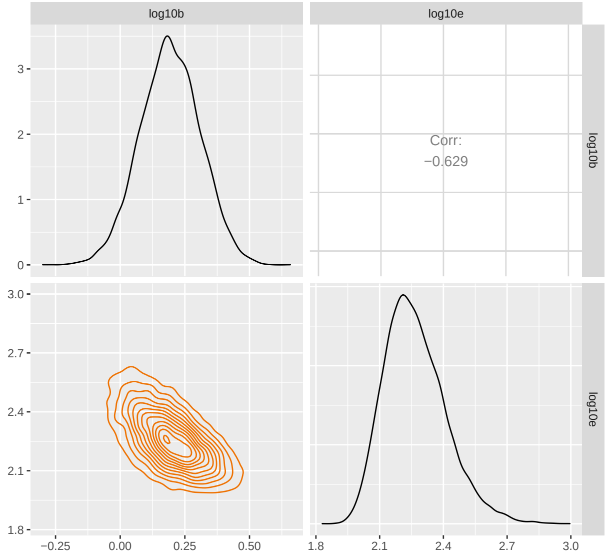

(d) Correlations between parameters

Figure 5: Dose-response curve (a), PPC (b), prior and posterior distributions (c) and correlations between parameters (d).

### Data set: GLXMA_SE_survival

Table 6: Summary of parameter estimates (parameter d is set to 1) for GLXMA_SE_survival data set

| Parameter | median | Q2.5 | Q97.5 |
|---|---|---|---|
| b | 3.201 | 1.567 | 6.961 |
| e | 187.844 | 120.473 | 419.635 |

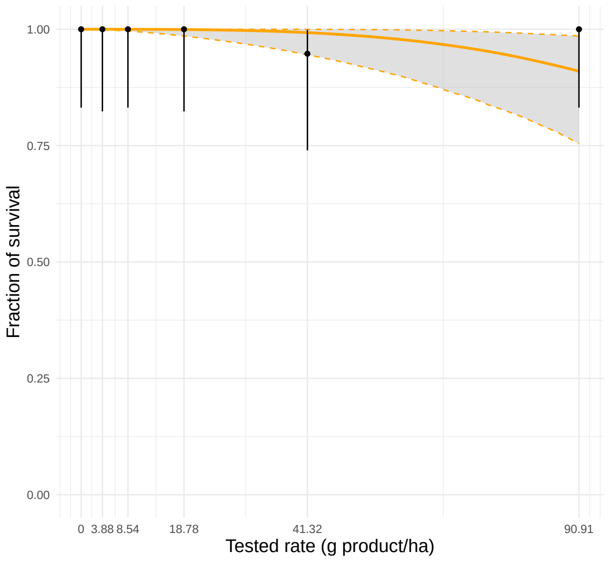

(a) Dose-response curve

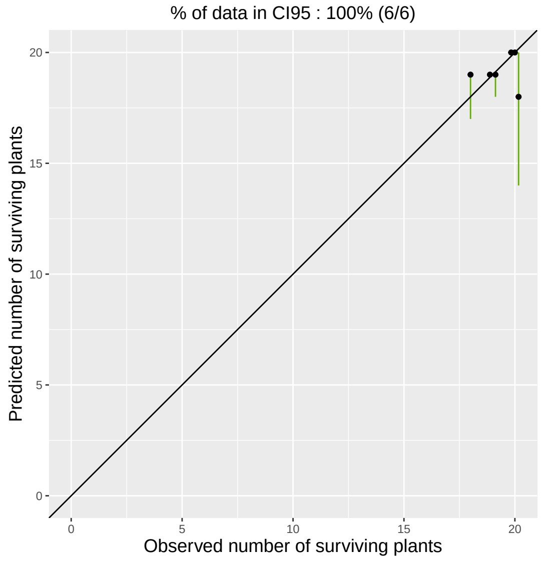

(b) Posterior predictive check (PPC)

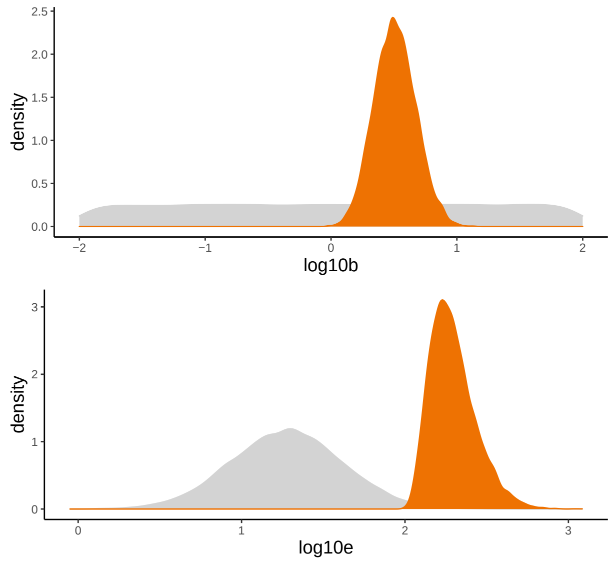

(c) Priors and posteriors

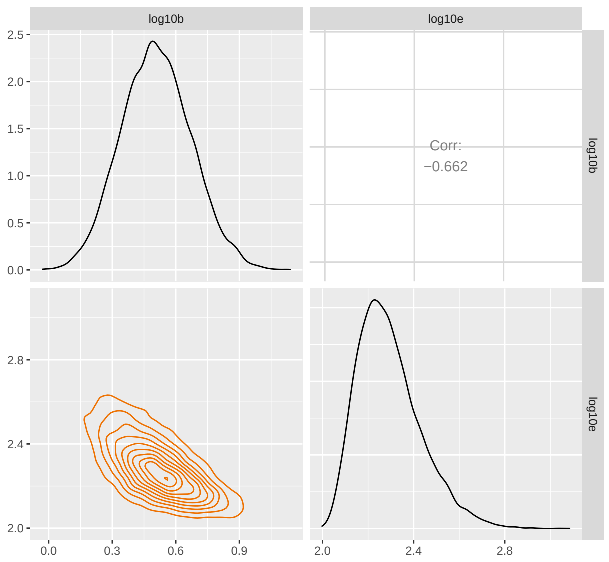

(d) Correlations between parameters

Figure 6: Dose-response curve (a), PPC (b), prior and posterior distributions (c) and correlations between parameters (d).

#### Data set: HELAN_SE_survival

Table 7: Summary of parameter estimates for HELAN_SE_survival data set

| Parameter | median | Q2.5 | Q97.5 |
|---|---|---|---|
| b | 1.829 | 0.915 | 4.218 |
| d | 0.942 | 0.862 | 0.987 |
| e | 78.273 | 53.724 | 130.930 |

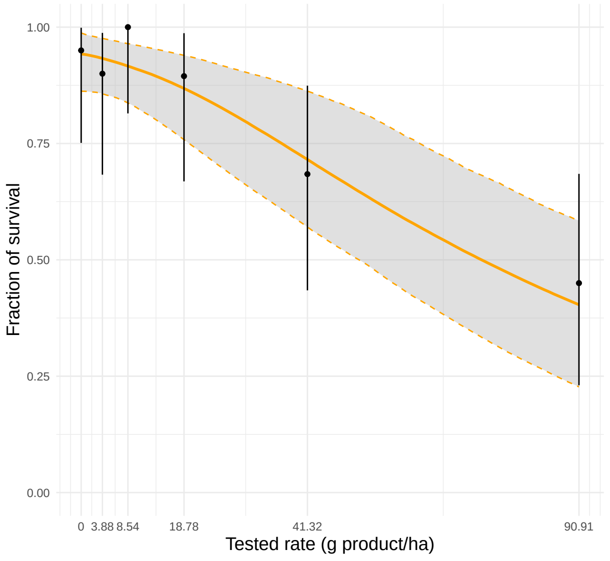

(a) Dose-response curve

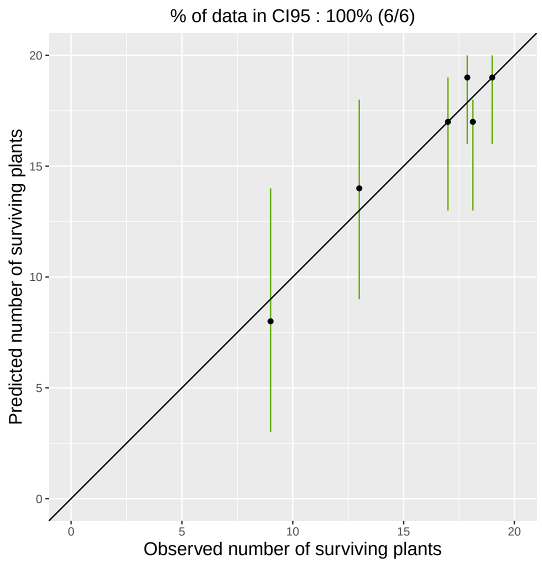

(b) Posterior predictive check (PPC)

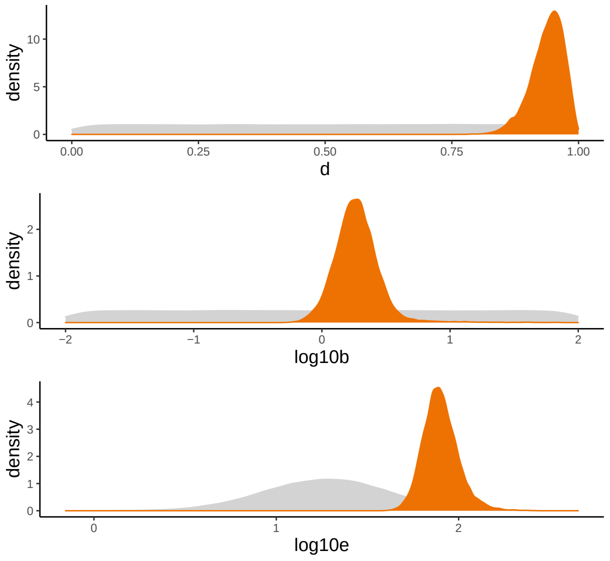

(c) Priors and posteriors

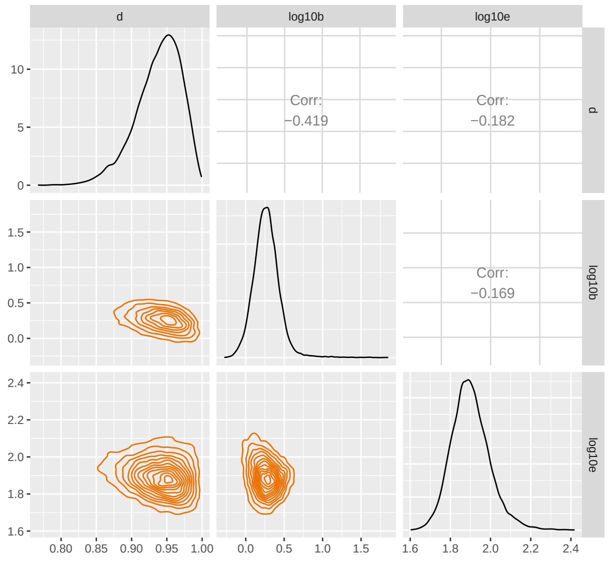

(d) Correlations between parameters

Figure 7: Dose-response curve (a), PPC (b), prior and posterior distributions (c) and correlations between parameters (d).

#### Data set: LYPES_SE_survival

Table 8: Summary of parameter estimates (parameter d is set to 1) for LYPES_SE_survival data set

| Parameter | median | Q2.5 | Q97.5 |
|---|---|---|---|
| b | 2.480 | 1.394 | 4.199 |
| e | 88.004 | 66.529 | 133.261 |

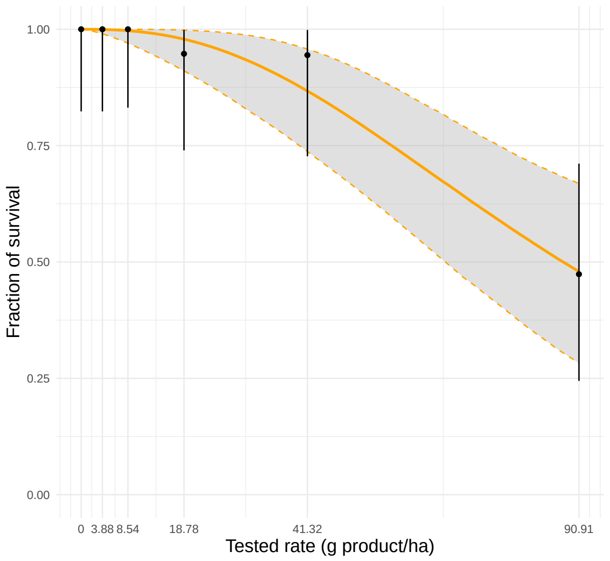

(a) Dose-response curve

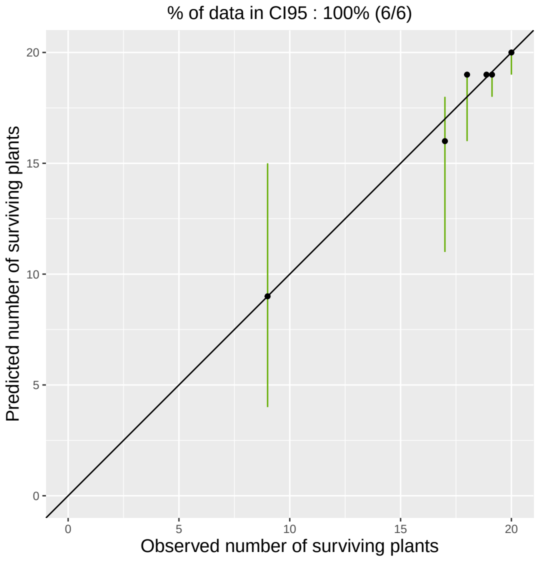

(b) Posterior predictive check (PPC)

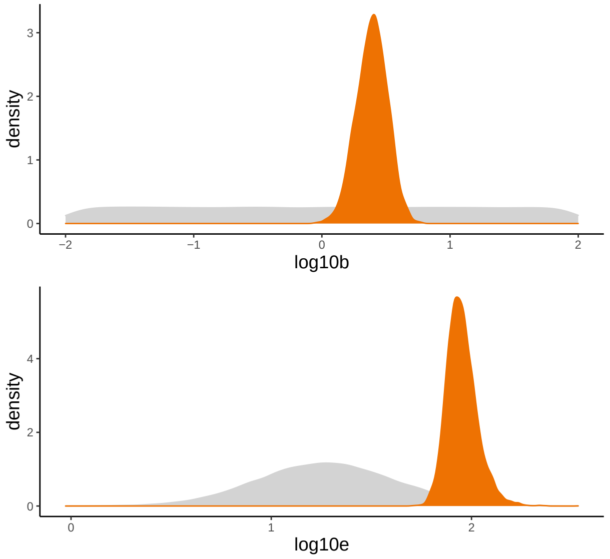

(c) Priors and posteriors

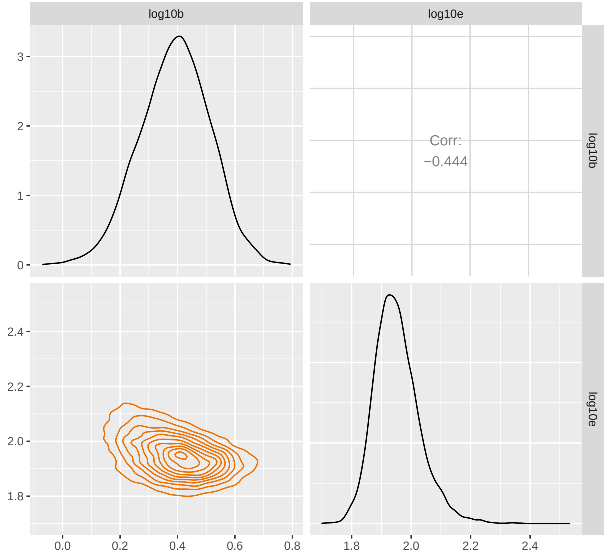

(d) Correlations between parameters

Figure 8: Dose-response curve (a), PPC (b), prior and posterior distributions (c) and correlations between parameters (d).

#### Data set: TRZAW_SE_survival

Table 9: Summary of parameter estimates (parameter d is set to 1) for TRZAW_SE_survival data set

| Parameter | median | Q2.5 | Q97.5 |
|---|---|---|---|
| b | 34.909 | 5.052 | 95.465 |
| e | 129.696 | 97.385 | 293.648 |

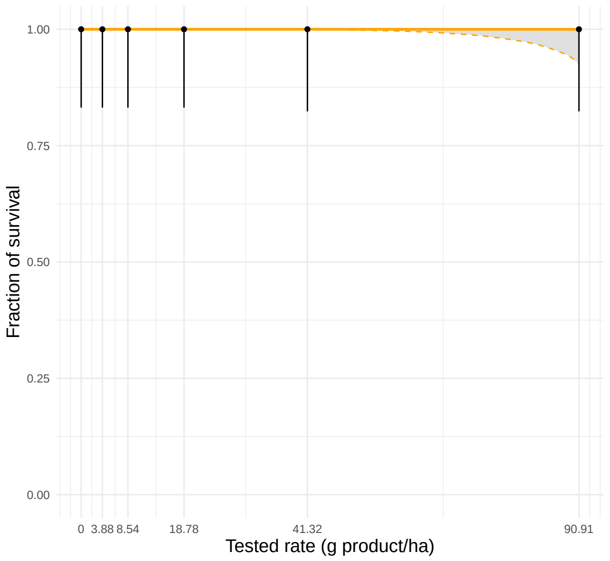

(a) Dose-response curve

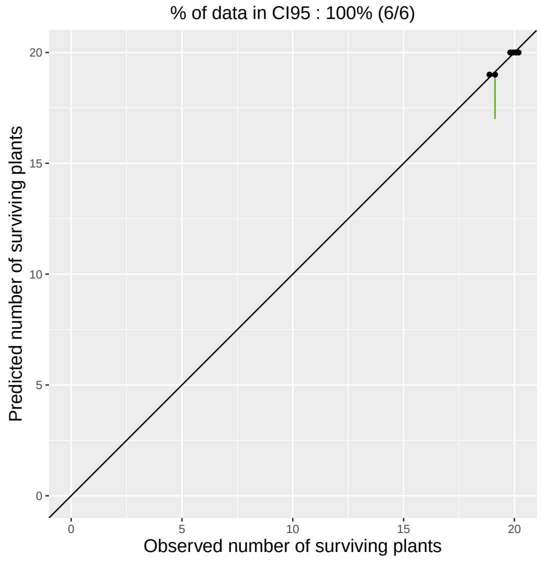

(b) Posterior predictive check (PPC)

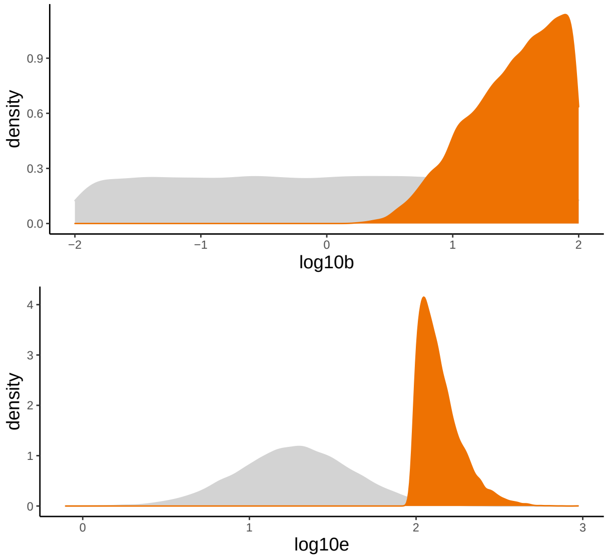

(c) Priors and posteriors

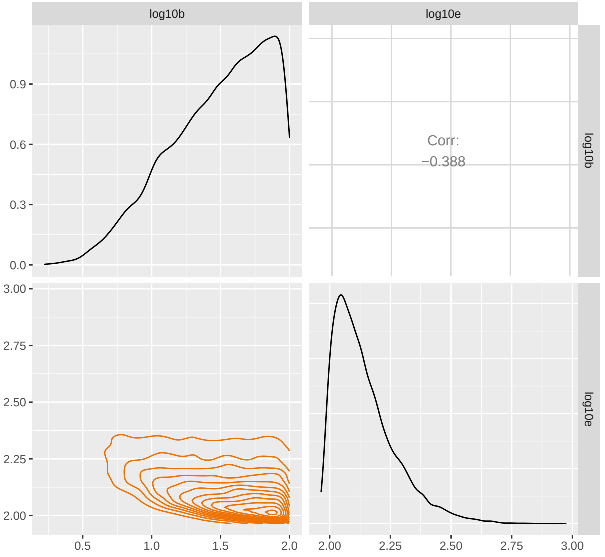

(d) Correlations between parameters

Figure 9: Dose-response curve (a), PPC (b), prior and posterior distributions (c) and correlations between parameters (d).

#### Data set: ZEAMA_SE_survival

Table 10: Summary of parameter estimates (parameter d is set to 1) for ZEAMA_SE_survival data set

| Parameter | median | Q2.5 | Q97.5 |
|---|---|---|---|
| b | 35.156 | 5.330 | 95.731 |
| e | 130.078 | 97.206 | 285.802 |

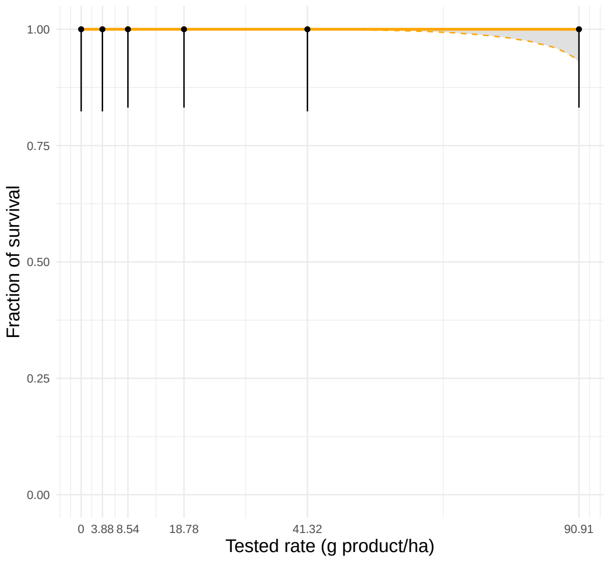

(a) Dose-response curve

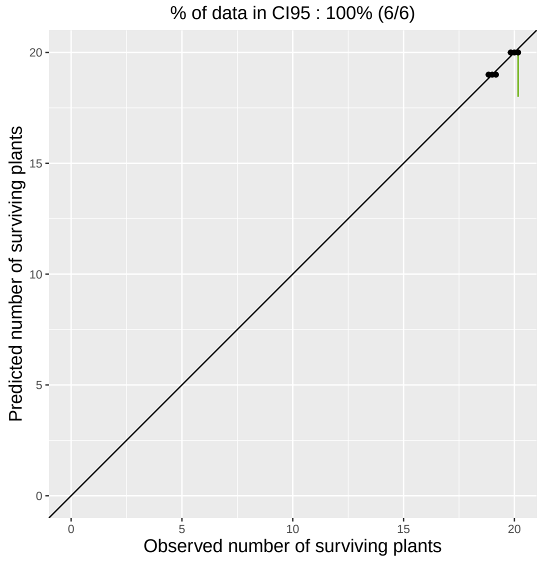

(b) Posterior predictive check (PPC)

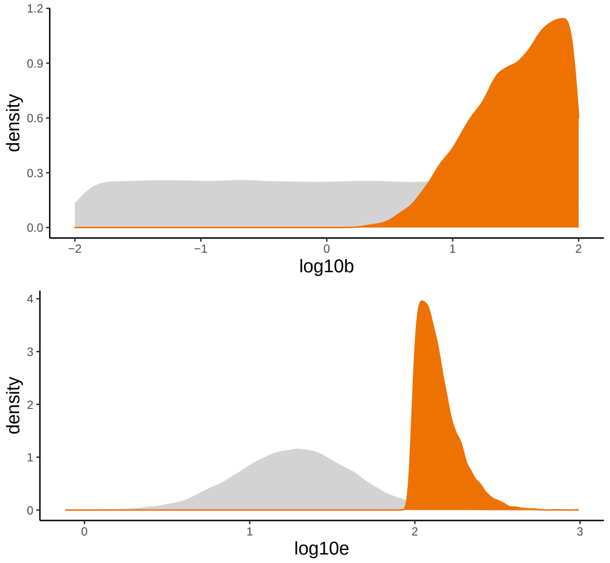

(c) Priors and posteriors

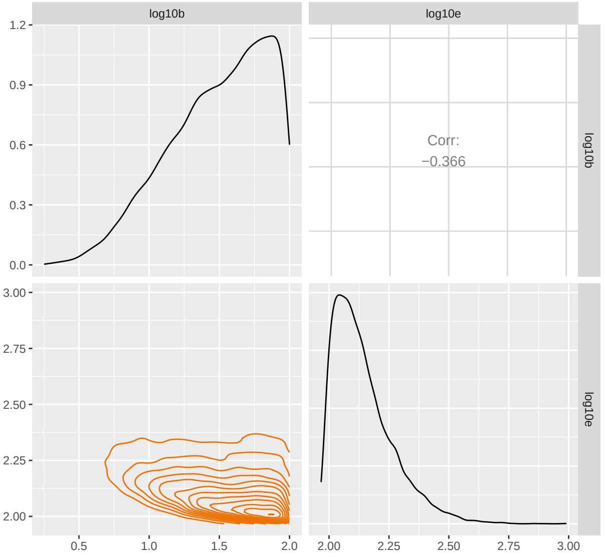

(d) Correlations between parameters

Figure 10: Dose-response curve (a), PPC (b), prior and posterior distributions (c) and correlations between parameters (d).
