## Supplementary material for "How to account for the uncertainty from standard toxicity tests in species sensitivity distributions: an example in non-target plants": Archive with reports on studies: report_SE_weight.pdf

### Dose-response analysis

Study 7

Seedling Emergence test - shoot dry SE_weight endpoint

25 June 2020

Contact:

---

This is a report which provides results on all performed dose-response analyses for the shoot dry SE_weight endpoint of the Seedling Emergence test for study 7.

---

#### Data set: ALLCE_SE_weight

Table 1: Summary of parameter estimates for ALLCE_SE_weight data set

| Parameter | median | Q2.5 | Q97.5 |
|---|---|---|---|
| b | 0.446 | 0.100 | 0.849 |
| d | 0.027 | 0.022 | 0.031 |
| e | 0.362 | 0.100 | 1.435 |
| sigma | 0.007 | 0.005 | 0.009 |

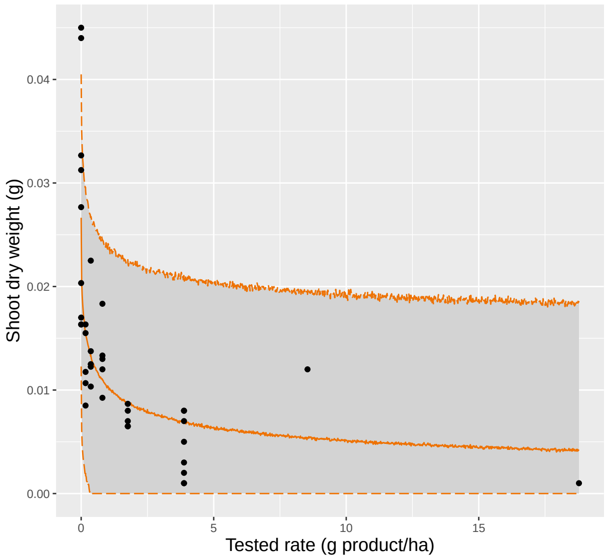

(a) Dose-response curve

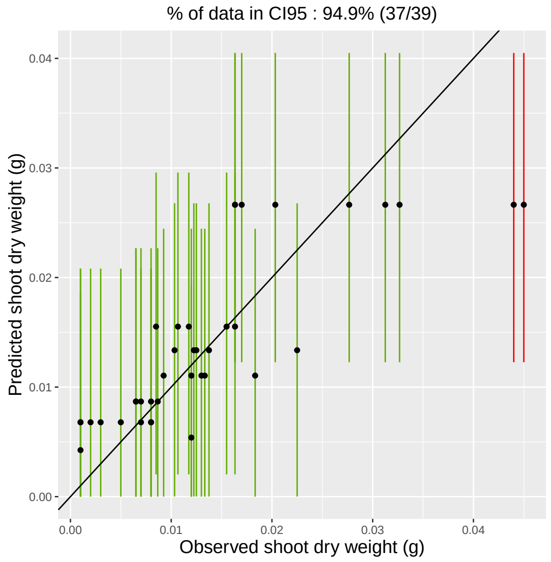

(b) Posterior predictive check (PPC)

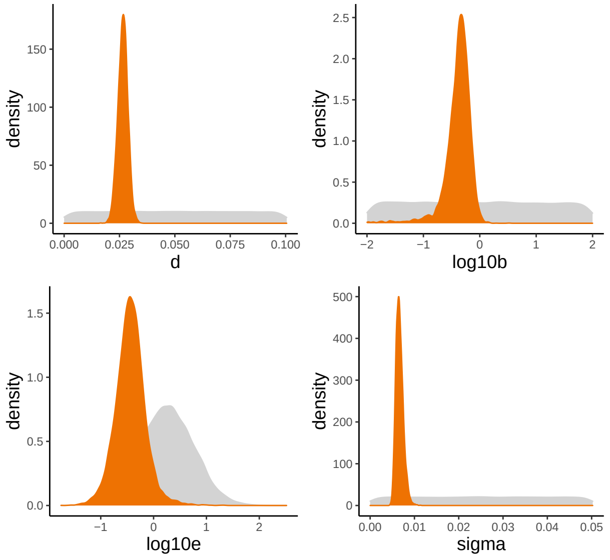

(c) Priors and posteriors

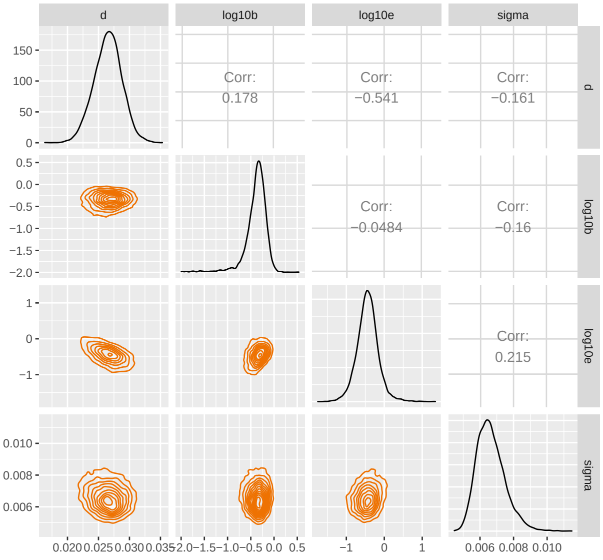

(d) Correlations between parameters

Figure 1: Dose-response curve (a), PPC (b), prior and posterior distributions (c) and correlations between parameters (d).

#### Data set: AVESA_SE_weight

Table 2: Summary of parameter estimates for AVESA_SE_weight data set

| Parameter | median | Q2.5 | Q97.5 |
|---|---|---|---|
| b | 2.026 | 1.525 | 2.845 |
| d | 0.491 | 0.453 | 0.534 |
| e | 13.604 | 11.405 | 16.098 |
| sigma | 0.046 | 0.036 | 0.063 |

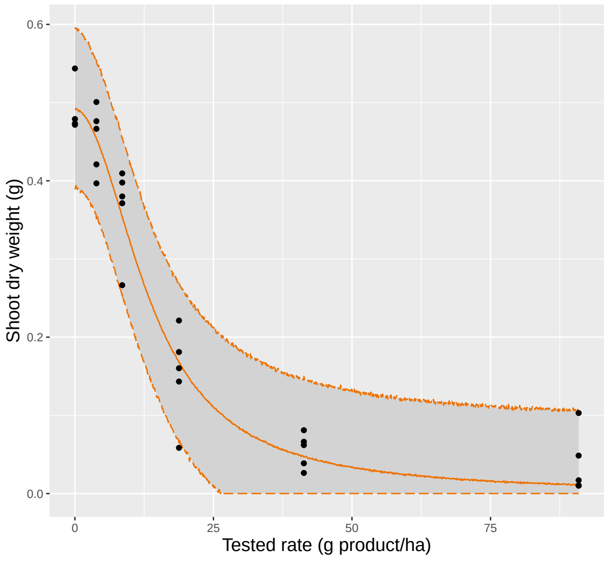

(a) Dose-response curve

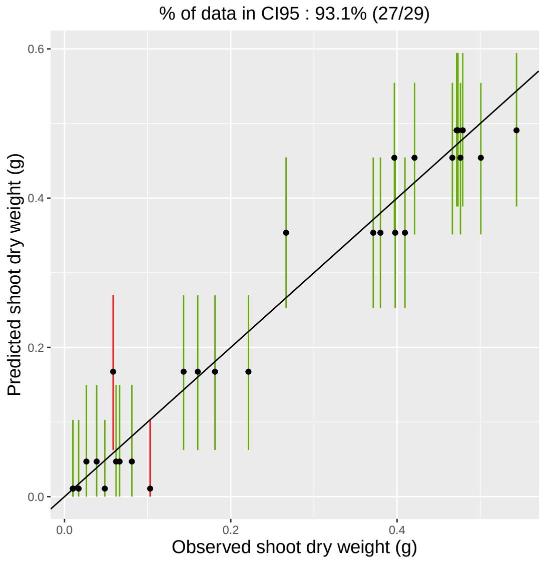

(b) Posterior predictive check (PPC)

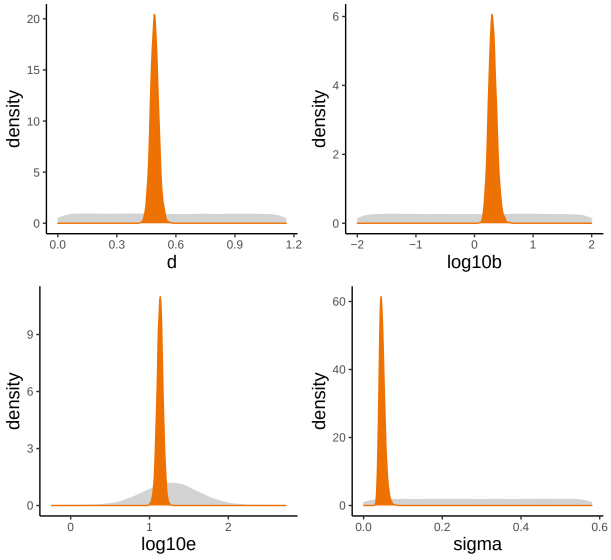

(c) Priors and posteriors

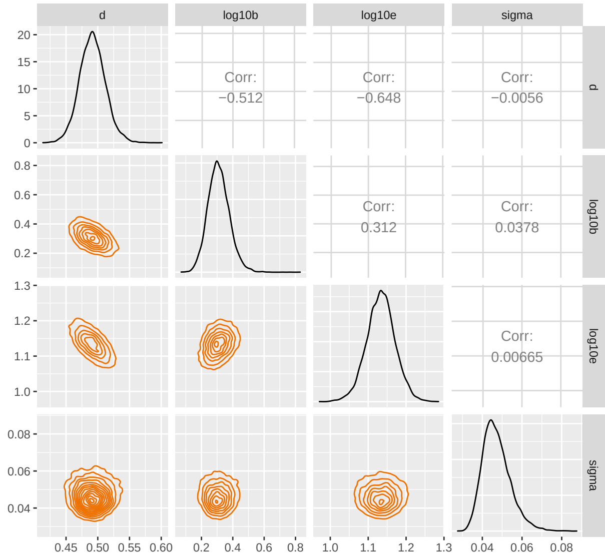

(d) Correlations between parameters

Figure 2: Dose-response curve (a), PPC (b), prior and posterior distributions (c) and correlations between parameters (d).

#### Data set: BEAVA_SE_weight

Table 3: Summary of parameter estimates for BEAVA_SE_weight data set

| Parameter | median | Q2.5 | Q97.5 |
|---|---|---|---|
| b | 1.013 | 0.589 | 1.767 |
| d | 0.419 | 0.372 | 0.467 |
| e | 3.976 | 2.591 | 5.887 |
| sigma | 0.100 | 0.085 | 0.119 |

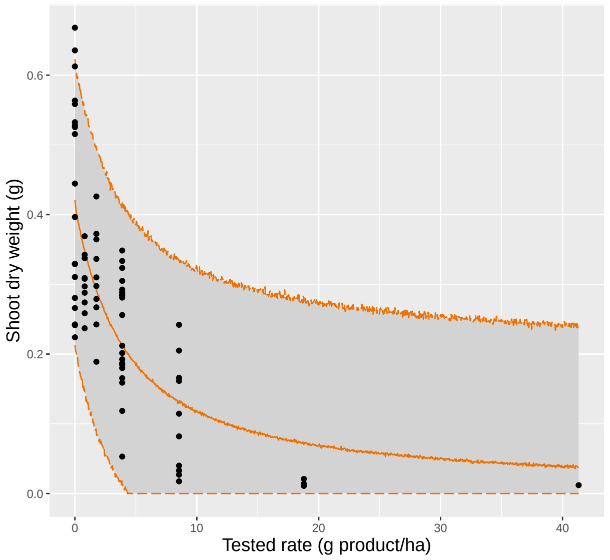

(a) Dose-response curve

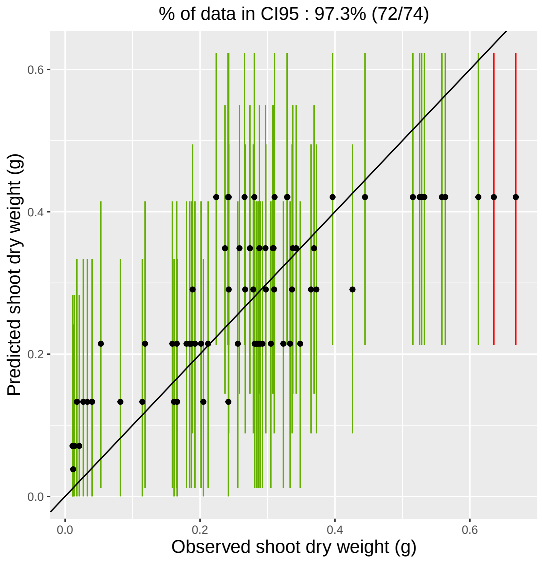

(b) Posterior predictive check (PPC)

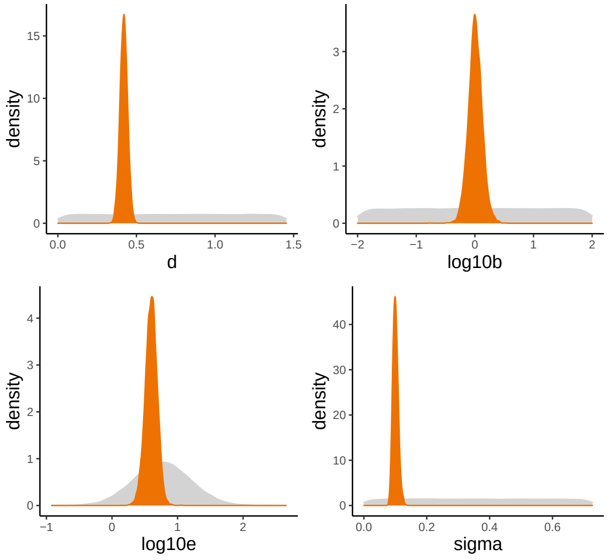

(c) Priors and posteriors

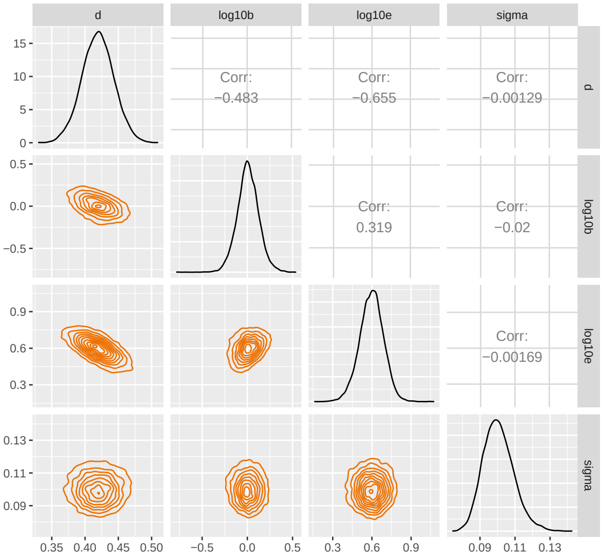

(d) Correlations between parameters

Figure 3: Dose-response curve (a), PPC (b), prior and posterior distributions (c) and correlations between parameters (d).

#### Data set: BRSNW_SE_weight

Table 4: Summary of parameter estimates for BRSNW_SE_weight data set

| Parameter | median | Q2.5 | Q97.5 |
|---|---|---|---|
| b | 3.293 | 2.110 | 8.143 |
| d | 0.974 | 0.878 | 1.077 |
| e | 11.237 | 9.272 | 13.748 |
| sigma | 0.186 | 0.153 | 0.234 |

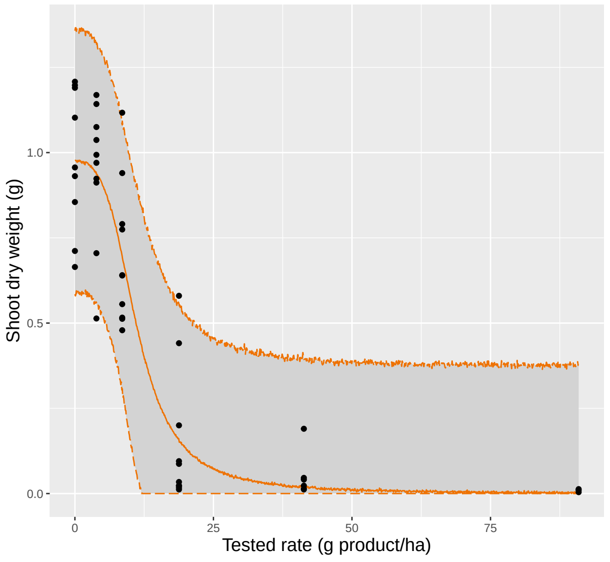

(a) Dose-response curve

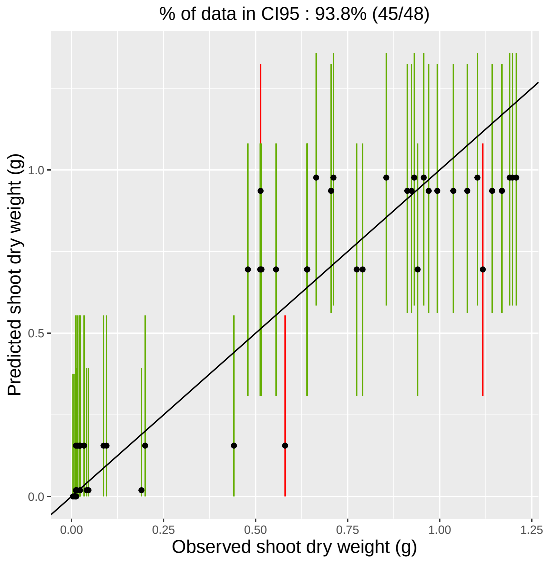

(b) Posterior predictive check (PPC)

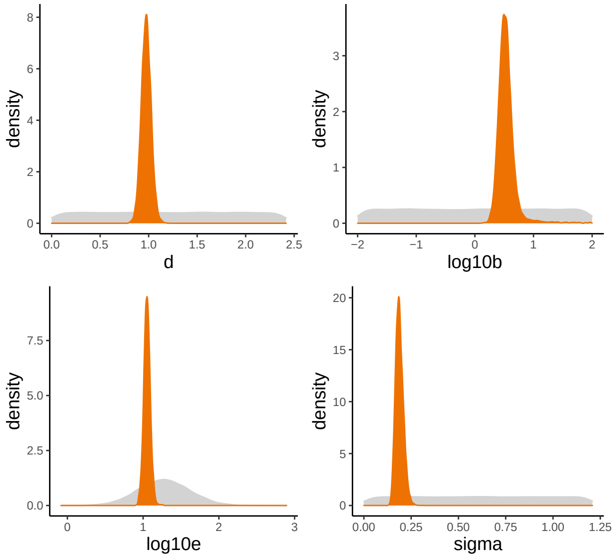

(c) Priors and posteriors

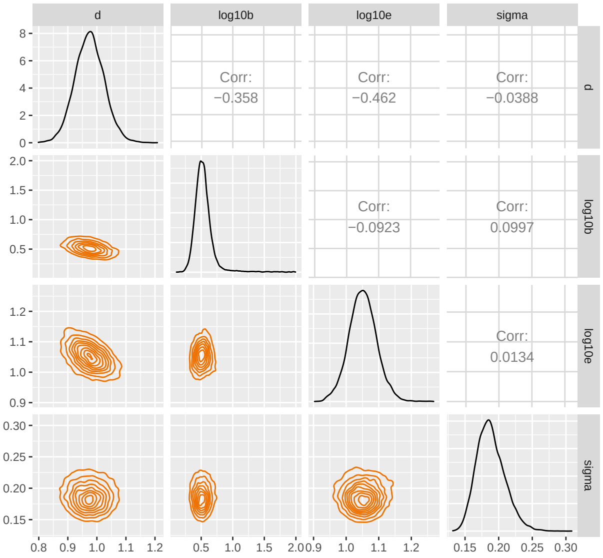

(d) Correlations between parameters

Figure 4: Dose-response curve (a), PPC (b), prior and posterior distributions (c) and correlations between parameters (d).

#### Data set: CUMSA_SE_weight

Table 5: Summary of parameter estimates for CUMSA_SE_weight data set

| Parameter | median | Q2.5 | Q97.5 |
|---|---|---|---|
| b | 1.244 | 0.932 | 1.674 |
| d | 1.489 | 1.332 | 1.647 |
| e | 9.283 | 7.002 | 12.337 |
| sigma | 0.245 | 0.205 | 0.300 |

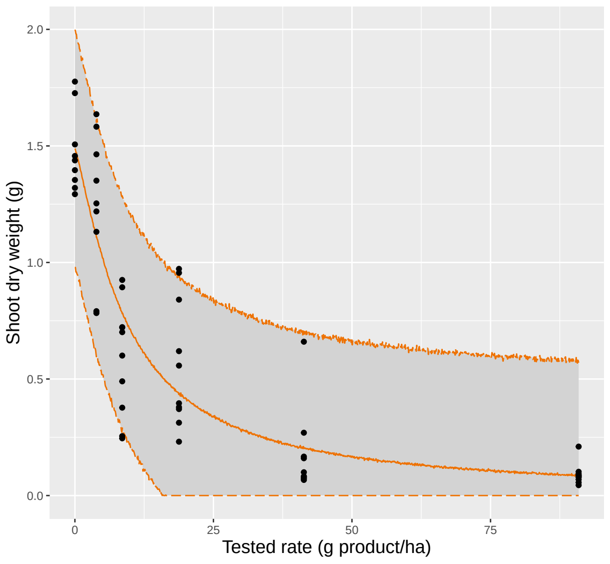

(a) Dose-response curve

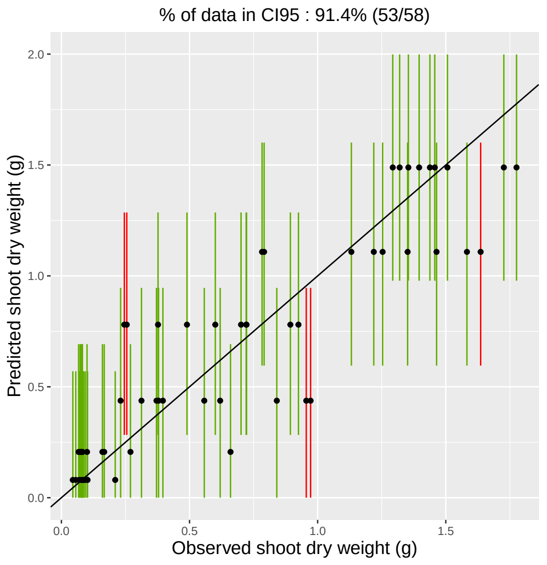

(b) Posterior predictive check (PPC)

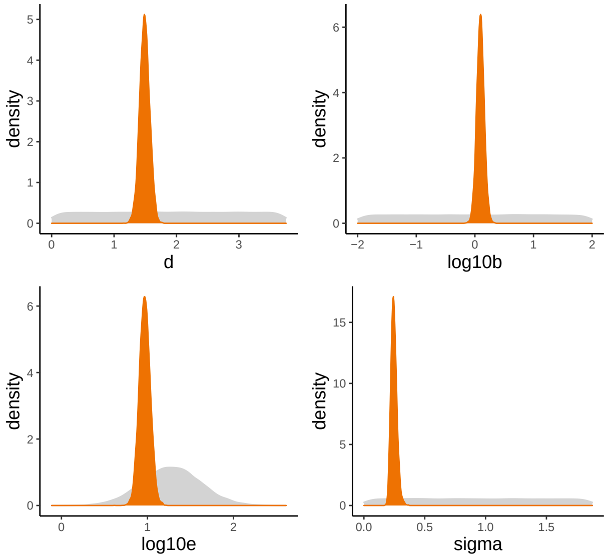

(c) Priors and posteriors

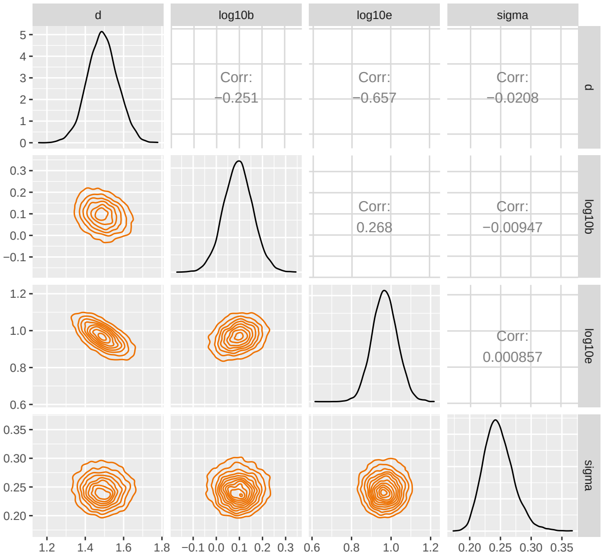

(d) Correlations between parameters

Figure 5: Dose-response curve (a), PPC (b), prior and posterior distributions (c) and correlations between parameters (d).

#### Data set: GLXMA_SE_weight

Table 6: Summary of parameter estimates for GLXMA_SE_weight data set

| Parameter | median | Q2.5 | Q97.5 |
|---|---|---|---|
| b | 2.016 | 1.391 | 2.895 |
| d | 1.134 | 1.060 | 1.213 |
| e | 52.546 | 43.143 | 64.219 |
| sigma | 0.198 | 0.166 | 0.242 |

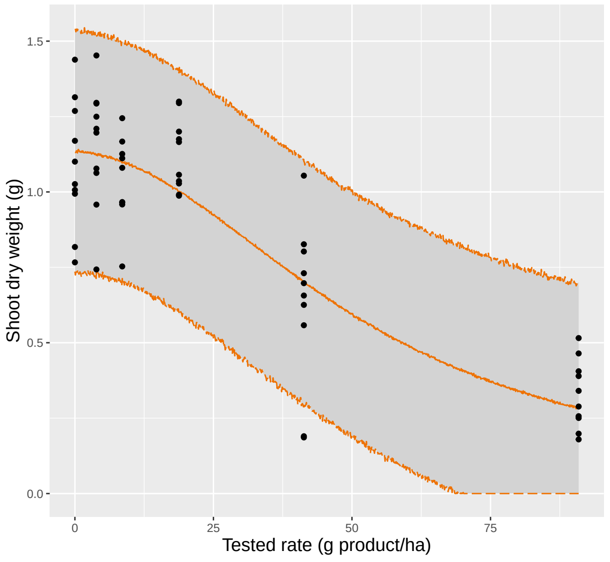

(a) Dose-response curve

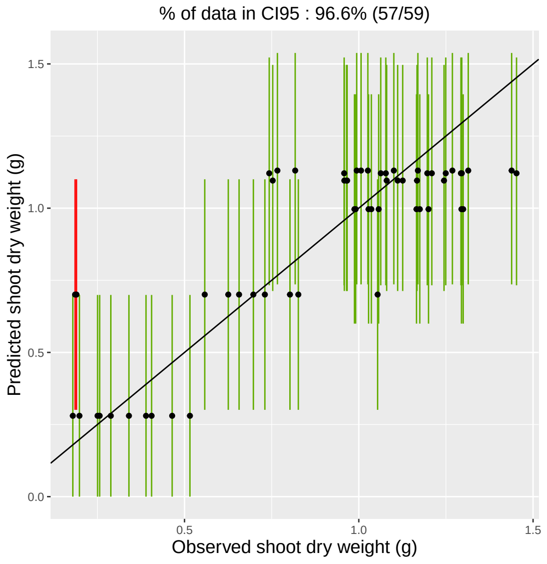

(b) Posterior predictive check (PPC)

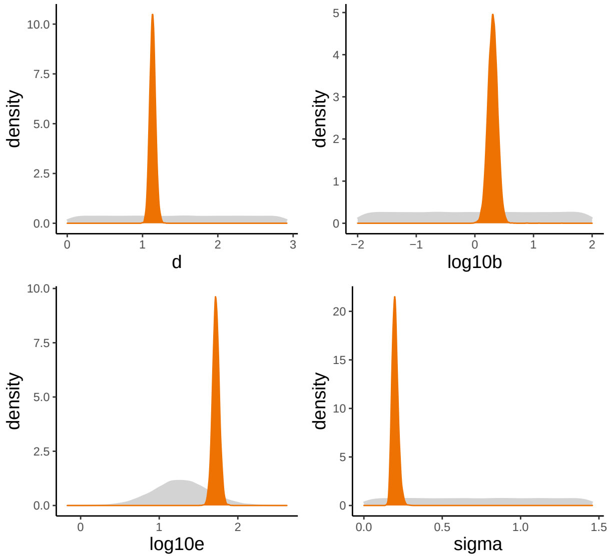

(c) Priors and posteriors

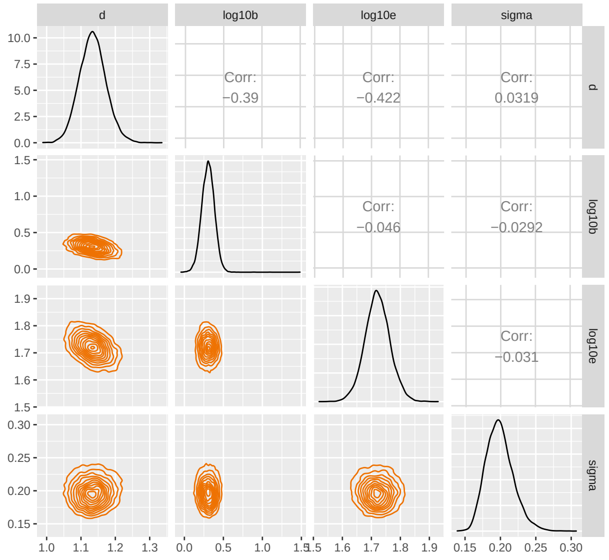

(d) Correlations between parameters

Figure 6: Dose-response curve (a), PPC (b), prior and posterior distributions (c) and correlations between parameters (d).

#### Data set: HELAN_SE_weight

Table 7: Summary of parameter estimates for HELAN_SE_weight data set

| Parameter | median | Q2.5 | Q97.5 |
|---|---|---|---|
| b | 1.319 | 0.828 | 2.299 |
| d | 0.501 | 0.431 | 0.573 |
| e | 10.586 | 7.143 | 15.550 |
| sigma | 0.114 | 0.095 | 0.141 |

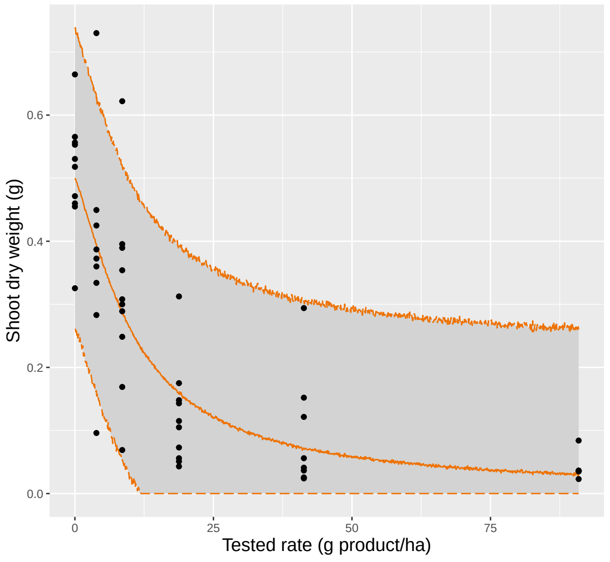

(a) Dose-response curve

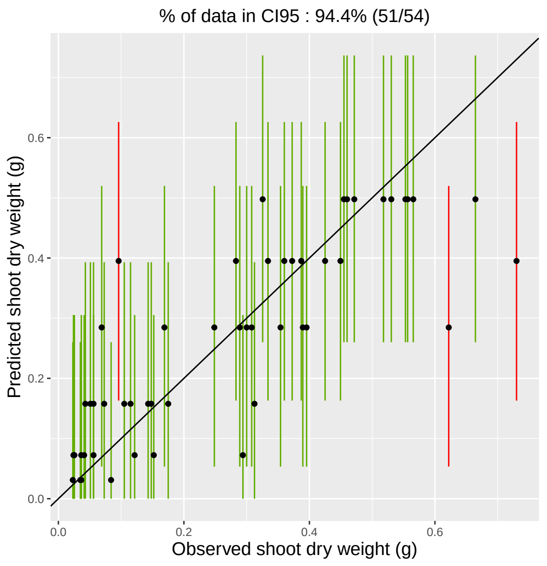

(b) Posterior predictive check (PPC)

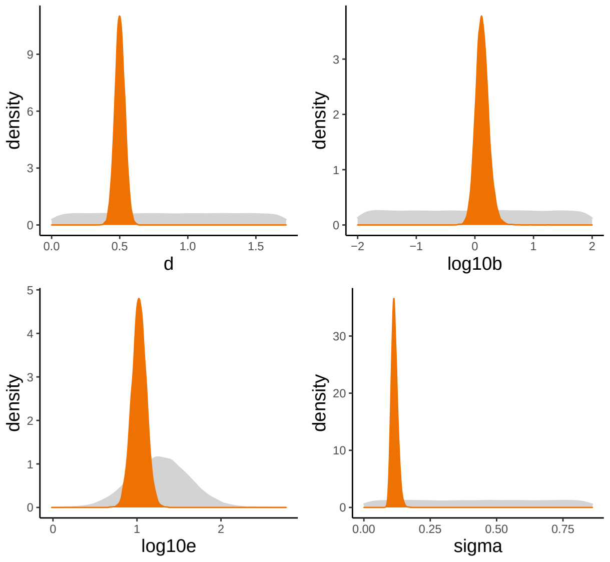

(c) Priors and posteriors

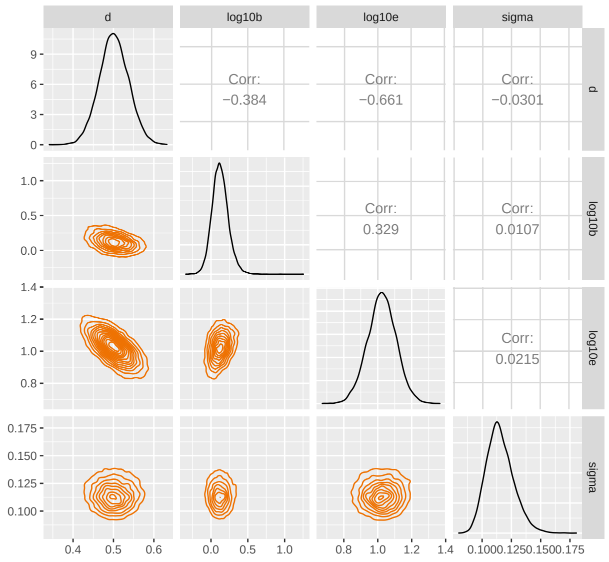

(d) Correlations between parameters

Figure 7: Dose-response curve (a), PPC (b), prior and posterior distributions (c) and correlations between parameters (d).

#### Data set: LYPES_SE_weight

Table 8: Summary of parameter estimates for LYPES_SE_weight data set

| Parameter | median | Q2.5 | Q97.5 |
|---|---|---|---|
| b | 2.270 | 1.731 | 3.195 |
| d | 1.101 | 1.029 | 1.179 |
| e | 15.859 | 13.647 | 18.116 |
| sigma | 0.133 | 0.111 | 0.163 |

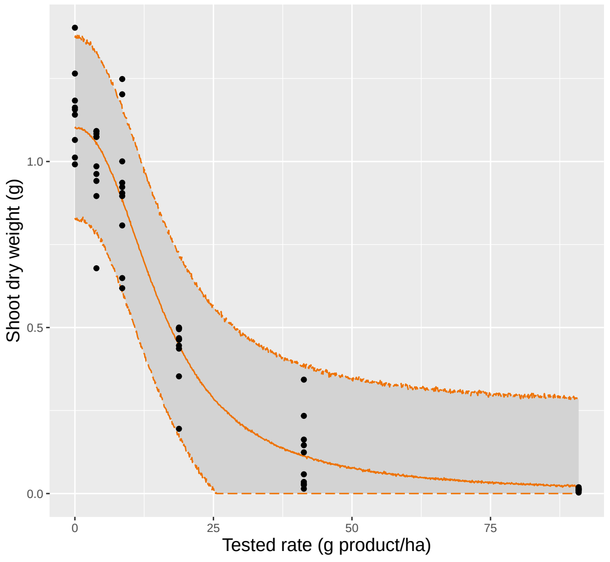

(a) Dose-response curve

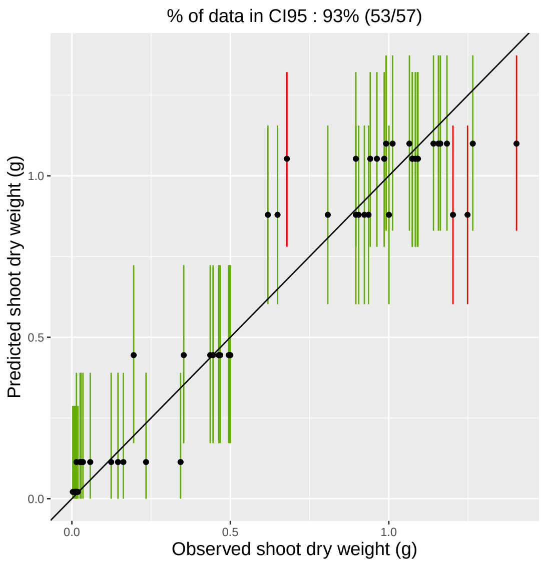

(b) Posterior predictive check (PPC)

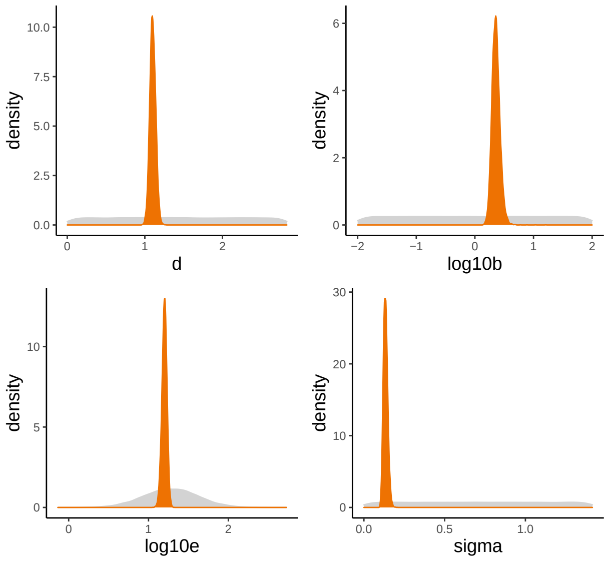

(c) Priors and posteriors

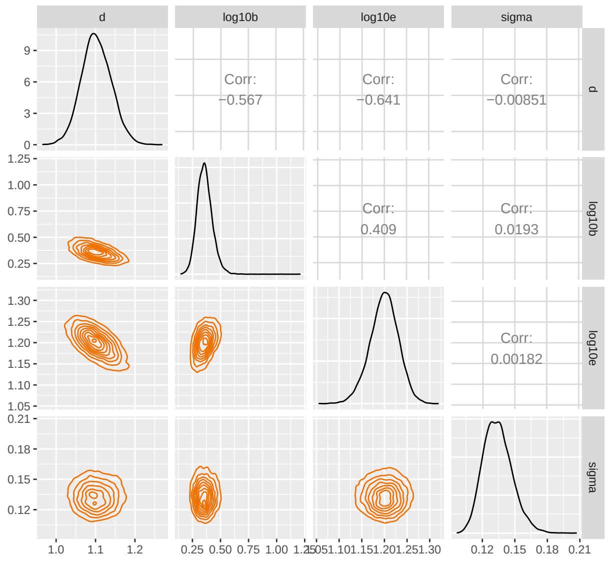

(d) Correlations between parameters

Figure 8: Dose-response curve (a), PPC (b), prior and posterior distributions (c) and correlations between parameters (d).

#### Data set: TRZAW_SE_weight

Table 9: Summary of parameter estimates for TRZAW_SE_weight data set

| Parameter | median | Q2.5 | Q97.5 |
|---|---|---|---|
| b | 0.921 | 0.609 | 1.450 |
| d | 0.397 | 0.369 | 0.431 |
| e | 110.828 | 81.123 | 164.913 |
| sigma | 0.034 | 0.027 | 0.047 |

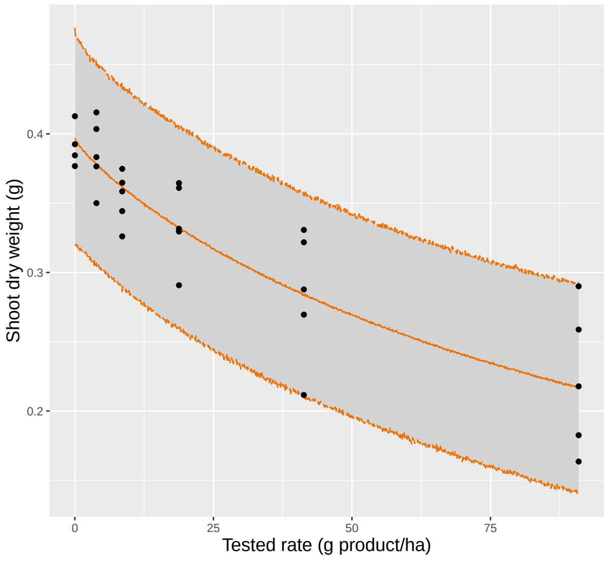

(a) Dose-response curve

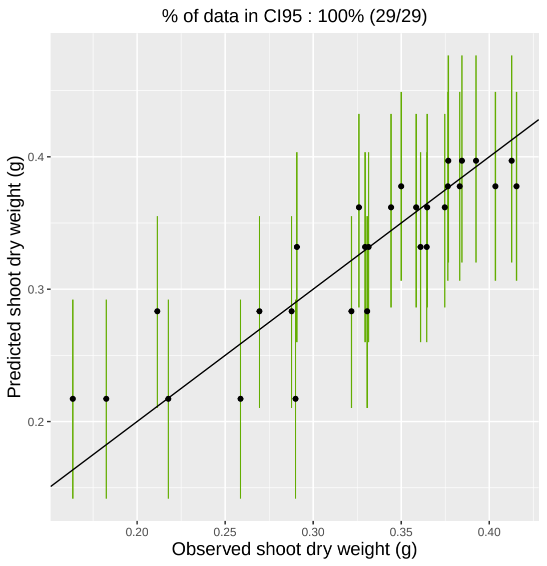

(b) Posterior predictive check (PPC)

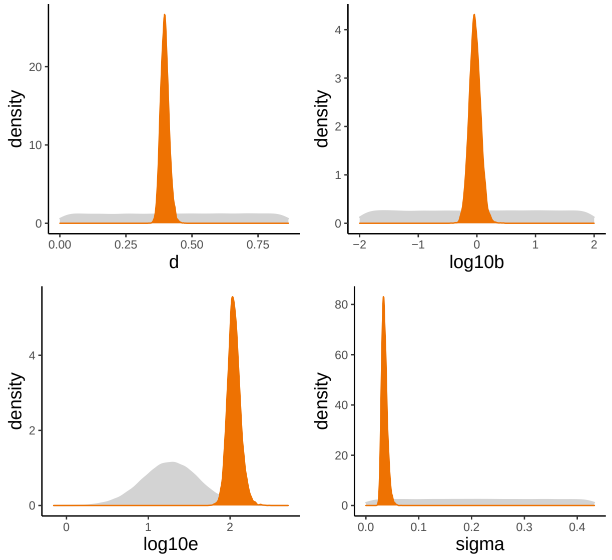

(c) Priors and posteriors

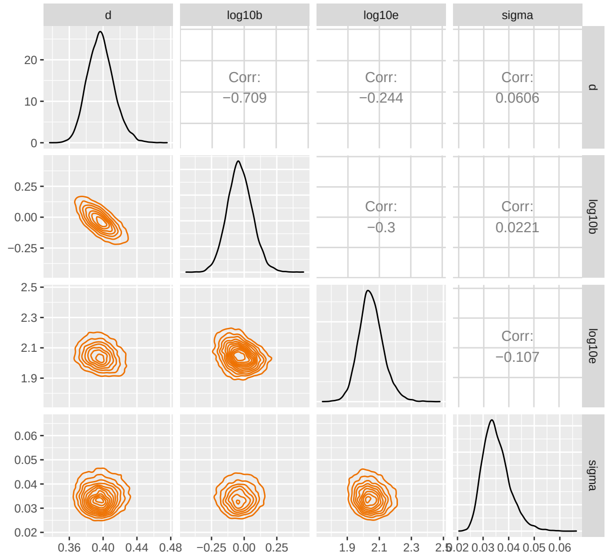

(d) Correlations between parameters

Figure 9: Dose-response curve (a), PPC (b), prior and posterior distributions (c) and correlations between parameters (d).

#### Data set: ZEAMA_SE_weight

Table 10: Summary of parameter estimates for ZEAMA_SE_weight data set

| Parameter | median | Q2.5 | Q97.5 |
|---|---|---|---|
| b | 1.454 | 1.054 | 2.039 |
| d | 2.509 | 2.312 | 2.725 |
| e | 37.510 | 29.265 | 48.304 |
| sigma | 0.425 | 0.357 | 0.520 |

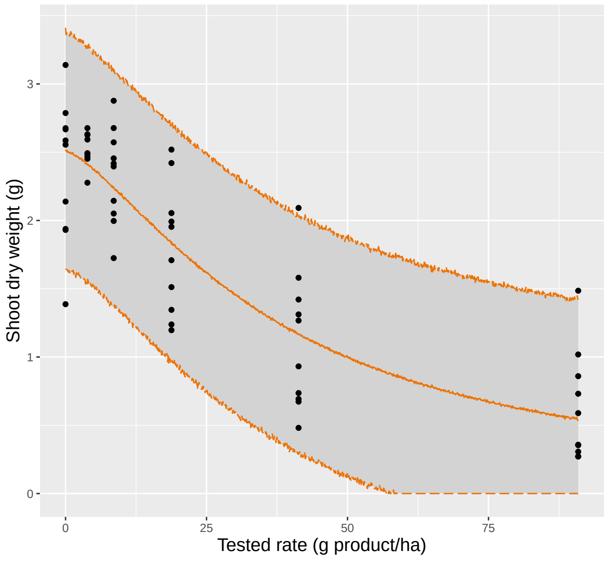

(a) Dose-response curve

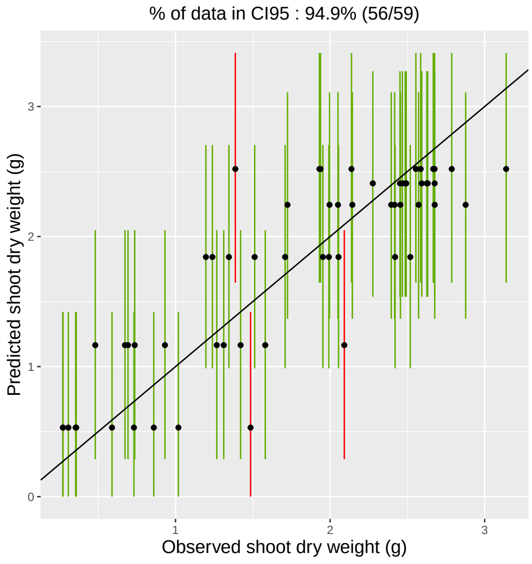

(b) Posterior predictive check (PPC)

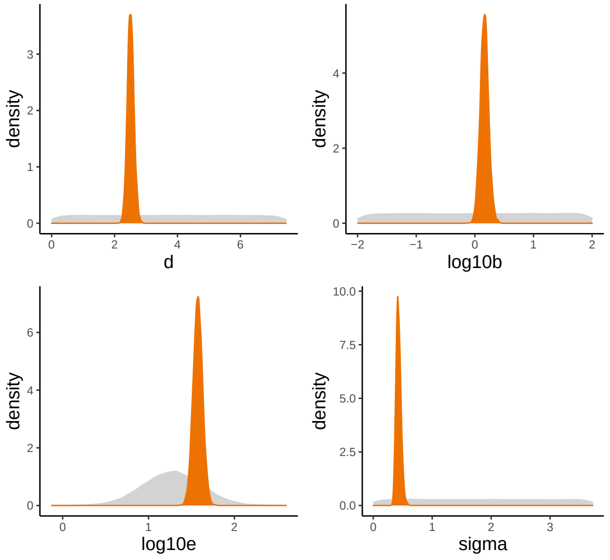

(c) Priors and posteriors

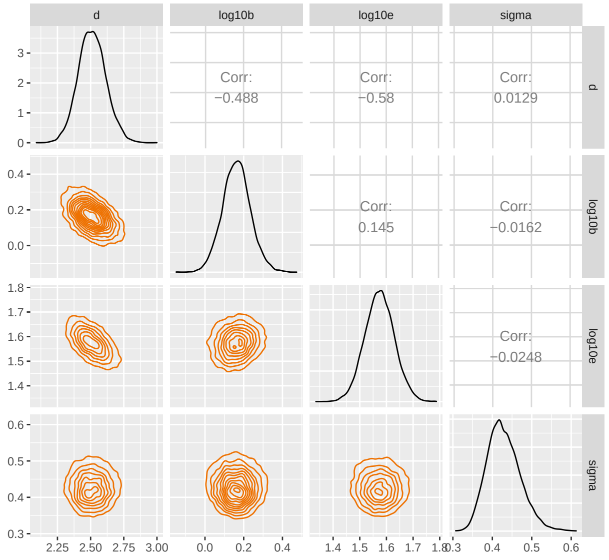

(d) Correlations between parameters

Figure 10: Dose-response curve (a), PPC (b), prior and posterior distributions (c) and correlations between parameters (d).
