## Supplementary material for "How to account for the uncertainty from standard toxicity tests in species sensitivity distributions: an example in non-target plants": Archive with reports on studies: report_VV_survival.pdf

### Dose-response analyses

Study 7

Vegetative Vigour test - survival endpoint

25 June 2020

Contact:

---

This is a report which provides results on all performed dose-response analyses for the survival endpoint of the Vegetative Vigour test for study 7.

---

#### Data set: ALLCE_VV_survival

Table 1: Summary of parameter estimates (parameter d is set to 1) for ALLCE_VV_survival data set

| Parameter | median | Q2.5 | Q97.5 |
|---|---|---|---|
| b | 7.778 | 2.234 | 66.578 |
| e | 46.642 | 41.602 | 75.844 |

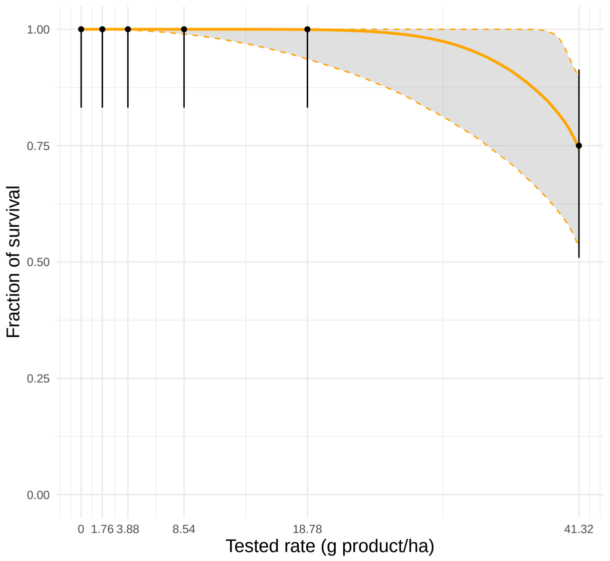

(a) Dose-response curve

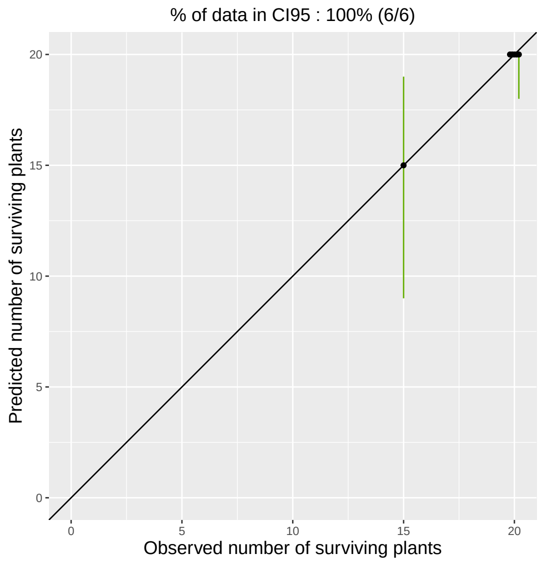

(b) Posterior predictive check (PPC)

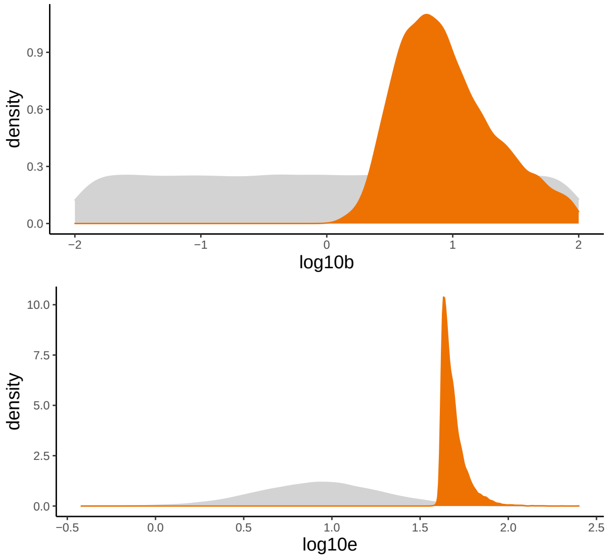

(c) Priors and posteriors

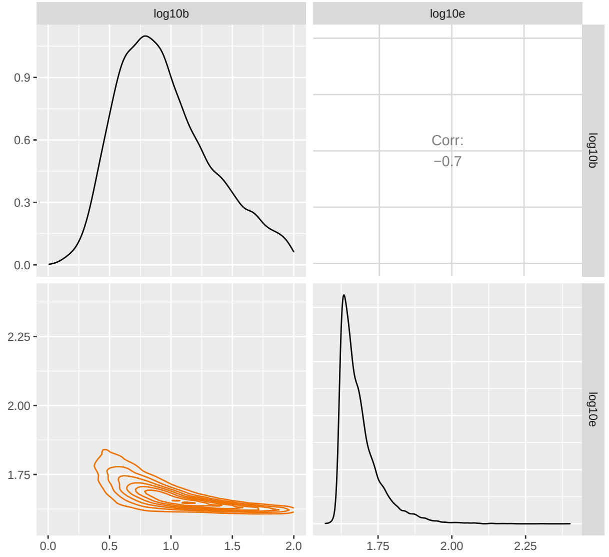

(d) Correlations between parameters

Figure 1: Dose-response curve (a), PPC (b), prior and posterior distributions (c) and correlations between parameters (d).

#### Data set: AVESA_VV_survival

Table 2: Summary of parameter estimates (parameter d is set to 1) for AVESA_VV_survival data set

| Parameter | median | Q2.5 | Q97.5 |
|---|---|---|---|
| b | 8.682 | 2.355 | 71.351 |
| e | 49.473 | 42.230 | 88.992 |

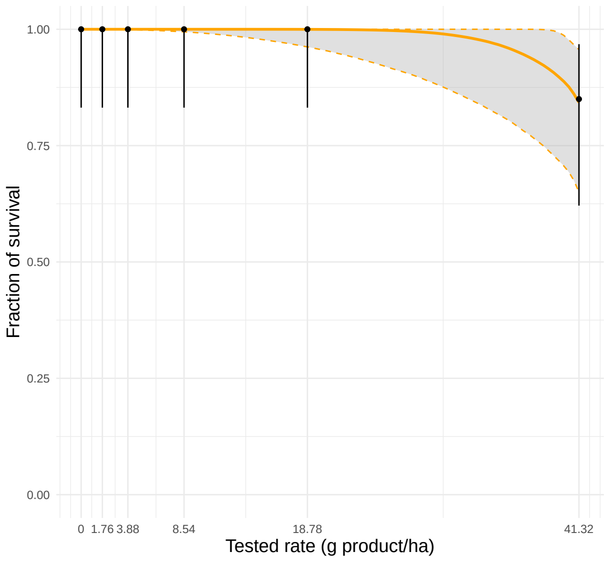

(a) Dose-response curve

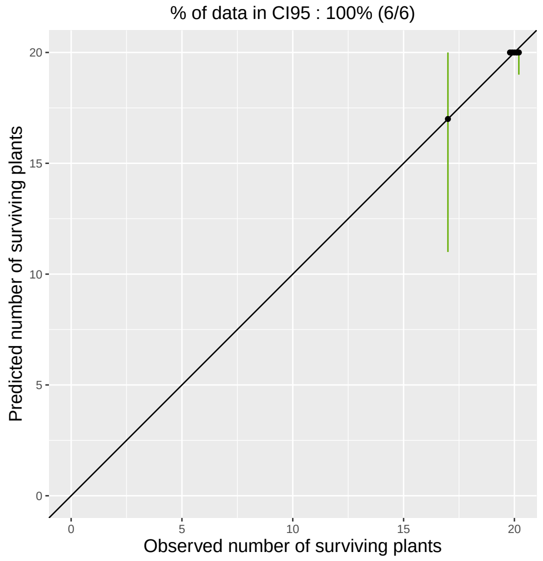

(b) Posterior predictive check (PPC)

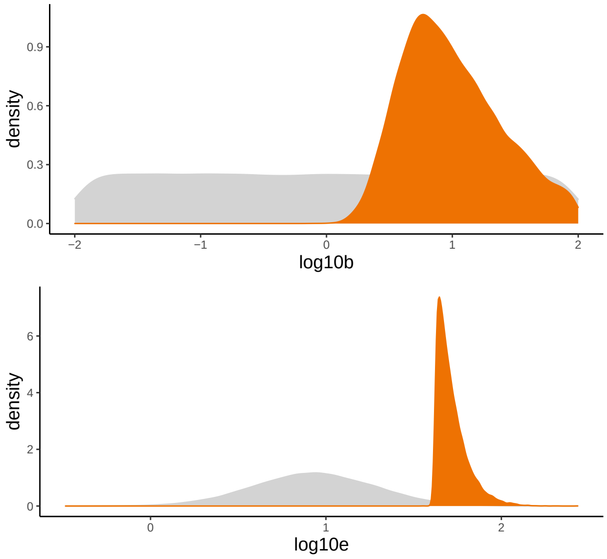

(c) Priors and posteriors

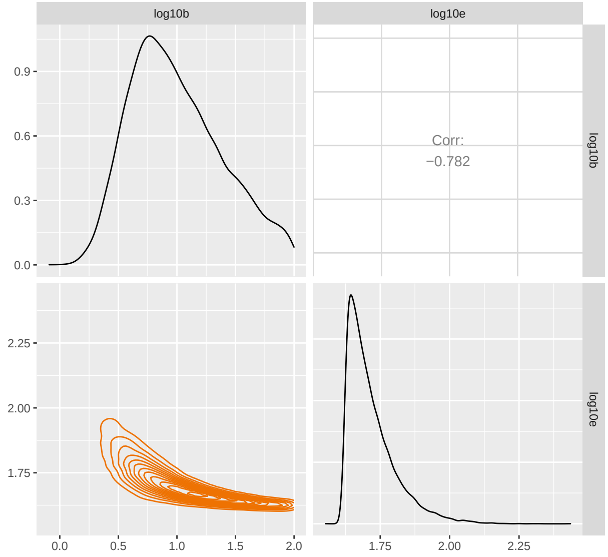

(d) Correlations between parameters

Figure 2: Dose-response curve (a), PPC (b), prior and posterior distributions (c) and correlations between parameters (d).

### Data set: BEAVA_VV_survival

Table 3: Summary of parameter estimates (parameter d is set to 1) for BEAVA_VV_survival data set

| Parameter | median | Q2.5 | Q97.5 |
|---|---|---|---|
| b | 4.600 | 2.593 | 8.054 |
| e | 7.086 | 5.837 | 8.589 |

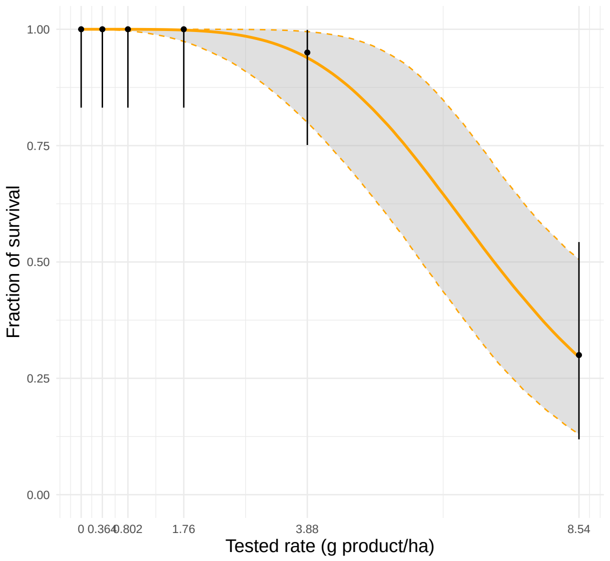

(a) Dose-response curve

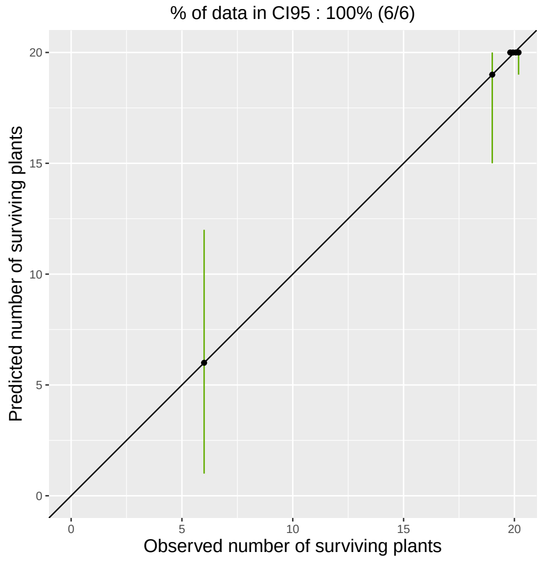

(b) Posterior predictive check (PPC)

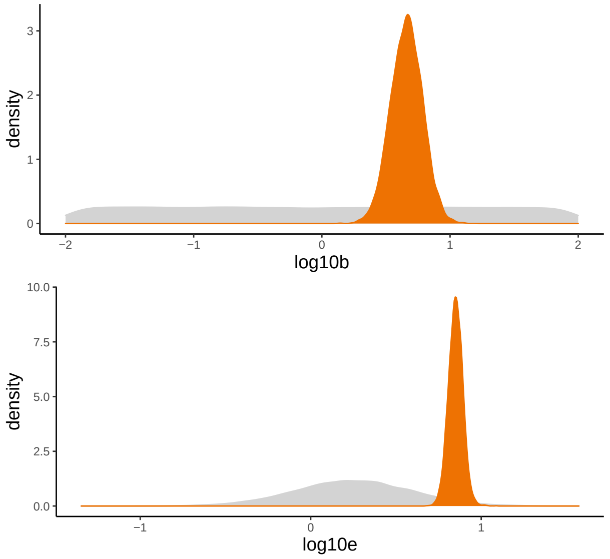

(c) Priors and posteriors

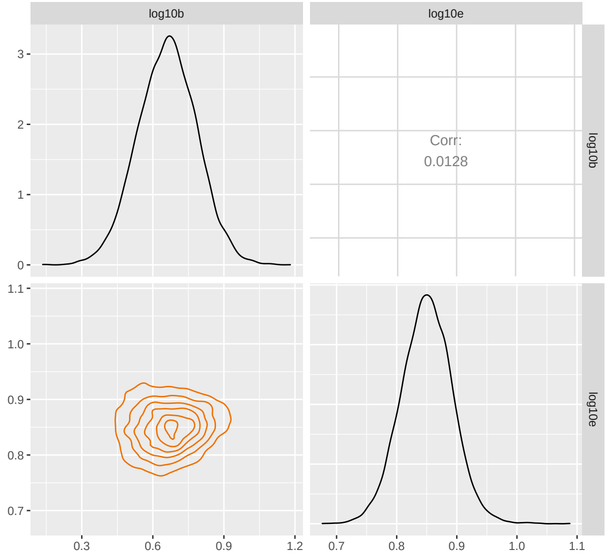

(d) Correlations between parameters

Figure 3: Dose-response curve (a), PPC (b), prior and posterior distributions (c) and correlations between parameters (d).

#### Data set: BRSNW_VV_survival

Table 4: Summary of parameter estimates (parameter d is set to 1) for BRSNW_VV_survival data set

| Parameter | median | Q2.5 | Q97.5 |
|---|---|---|---|
| b | 8.138 | 2.236 | 69.153 |
| e | 9.932 | 8.674 | 16.546 |

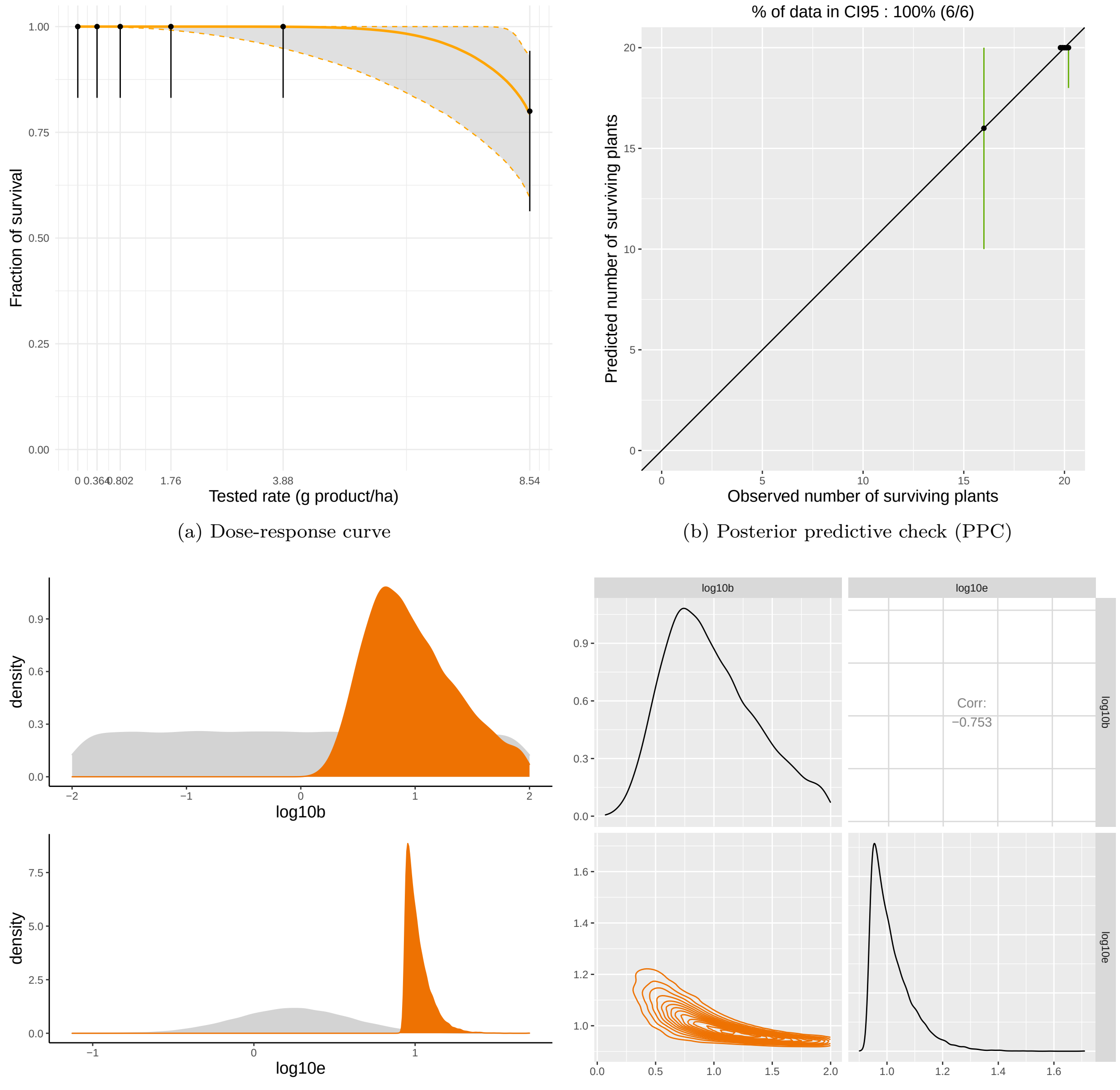

(a) Dose-response curve

(b) Posterior predictive check (PPC)

(c) Priors and posteriors

(d) Correlations between parameters

Figure 4: Dose-response curve (a), PPC (b), prior and posterior distributions (c) and correlations between parameters (d).

### Data set: CUMSA_VV_survival

Table 5: Summary of parameter estimates (parameter d is set to 1) for CUMSA_VV_survival data set

| Parameter | median | Q2.5 | Q97.5 |
|---|---|---|---|
| b | 8.805 | 3.385 | 64.355 |
| e | 38.937 | 32.401 | 42.707 |

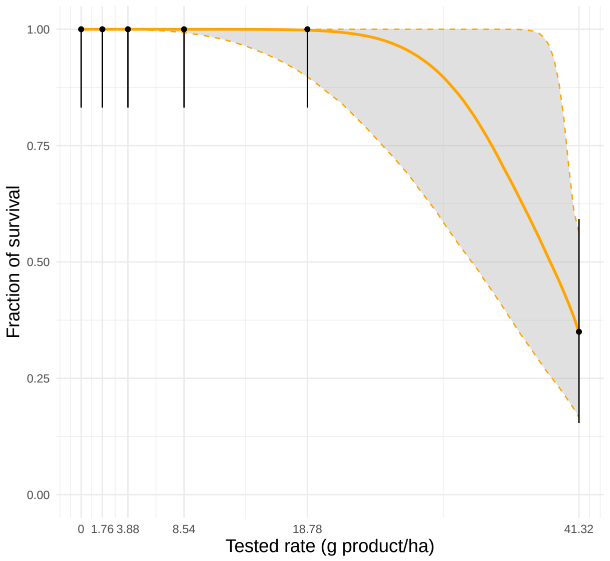

(a) Dose-response curve

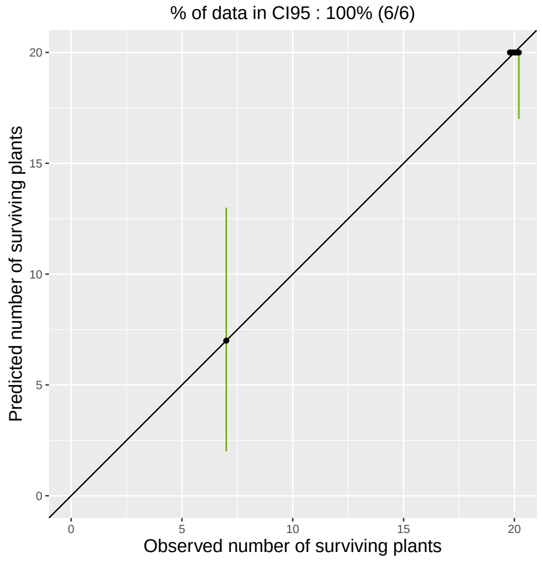

(b) Posterior predictive check (PPC)

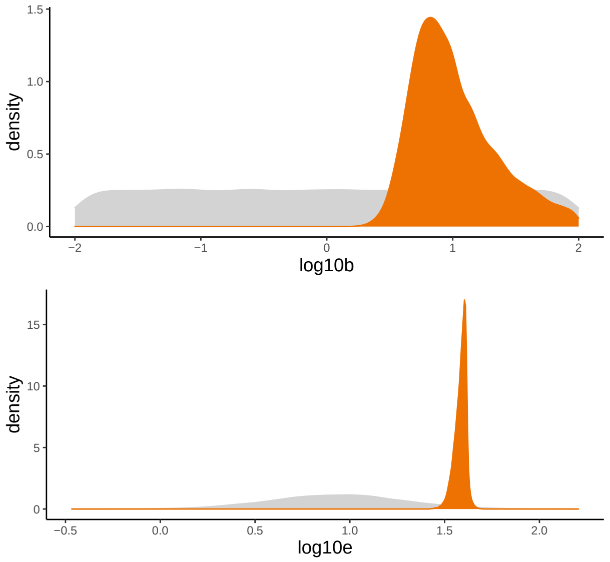

(c) Priors and posteriors

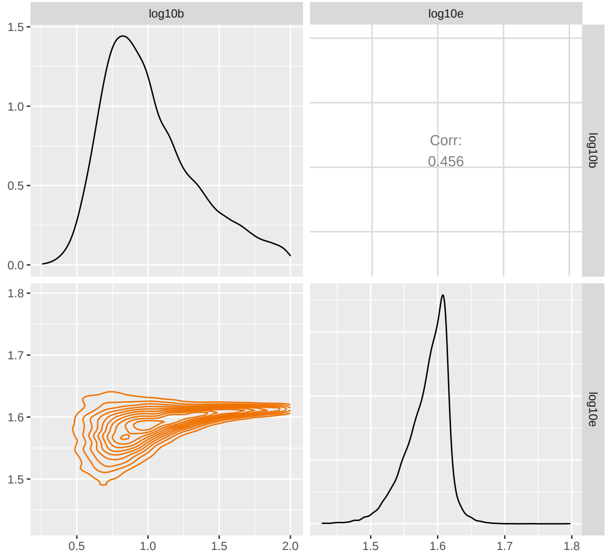

(d) Correlations between parameters

Figure 5: Dose-response curve (a), PPC (b), prior and posterior distributions (c) and correlations between parameters (d).

#### Data set: GLXMA_VV_survival

Table 6: Summary of parameter estimates (parameter d is set to 1) for GLXMA_VV_survival data set

| Parameter | median | Q2.5 | Q97.5 |
|---|---|---|---|
| b | 35.315 | 5.129 | 95.033 |
| e | 129.998 | 97.334 | 294.161 |

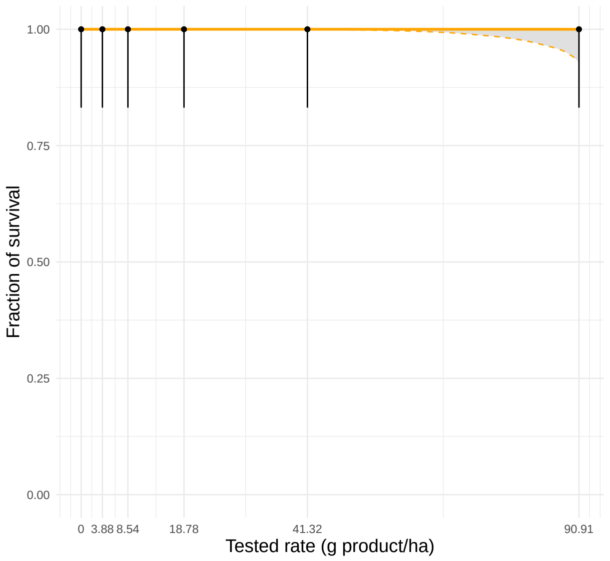

(a) Dose-response curve

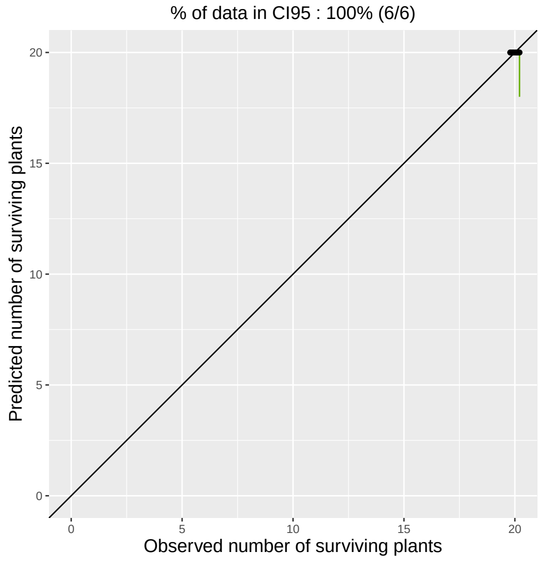

(b) Posterior predictive check (PPC)

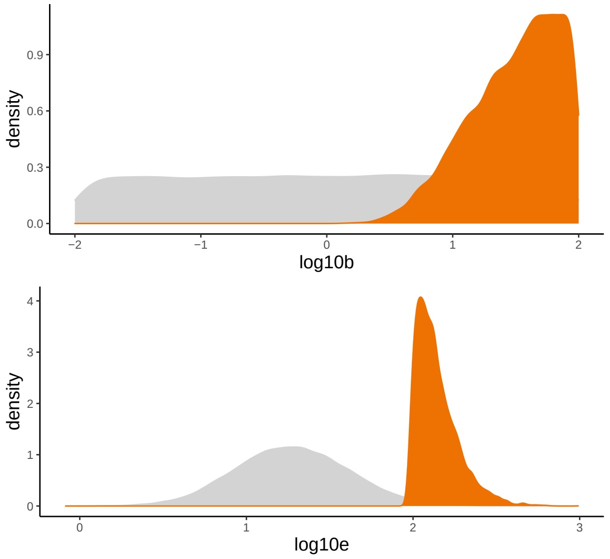

(c) Priors and posteriors

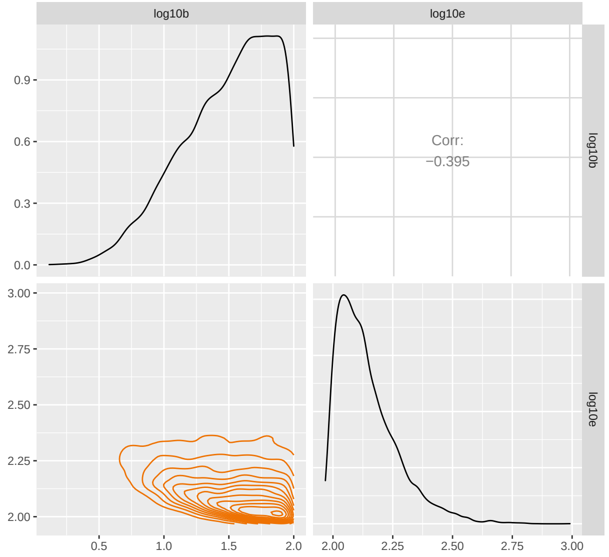

(d) Correlations between parameters

Figure 6: Dose-response curve (a), PPC (b), prior and posterior distributions (c) and correlations between parameters (d).

#### Data set: HELAN_VV_survival

Table 7: Summary of parameter estimates (parameter d is set to 1) for HELAN_VV_survival data set

| Parameter | median | Q2.5 | Q97.5 |
|---|---|---|---|
| b | 9.688 | 3.981 | 68.507 |
| e | 8.807 | 8.055 | 10.280 |

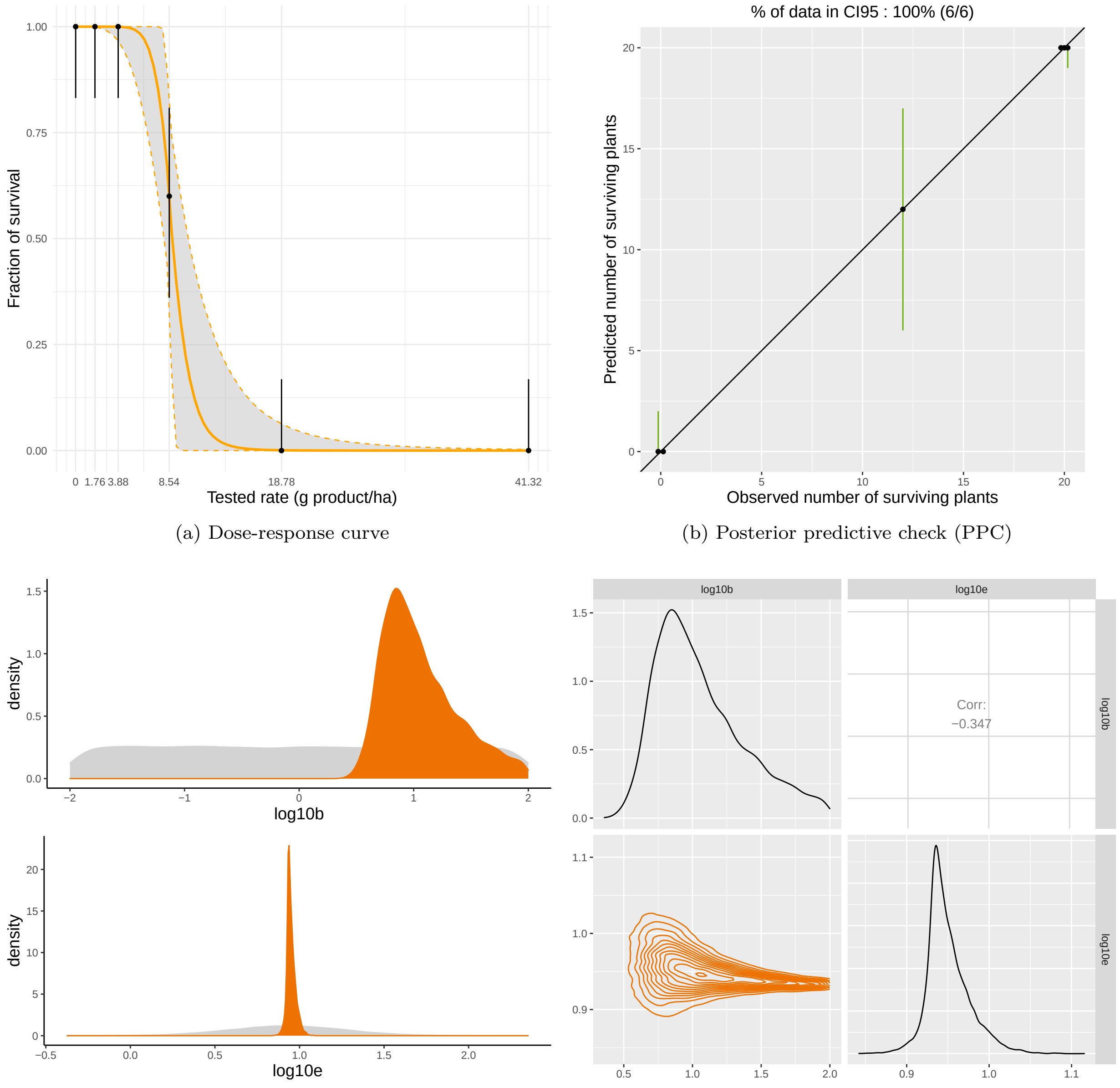

(a) Dose-response curve

(b) Posterior predictive check (PPC)

(c) Priors and posteriors

(d) Correlations between parameters

Figure 7: Dose-response curve (a), PPC (b), prior and posterior distributions (c) and correlations between parameters (d).

#### Data set: LYPES_VV_survival

Table 8: Summary of parameter estimates (parameter d is set to 1) for LYPES_VV_survival data set

| Parameter | median | Q2.5 | Q97.5 |
|---|---|---|---|
| b | 3.353 | 1.895 | 5.649 |
| e | 36.152 | 28.798 | 47.939 |

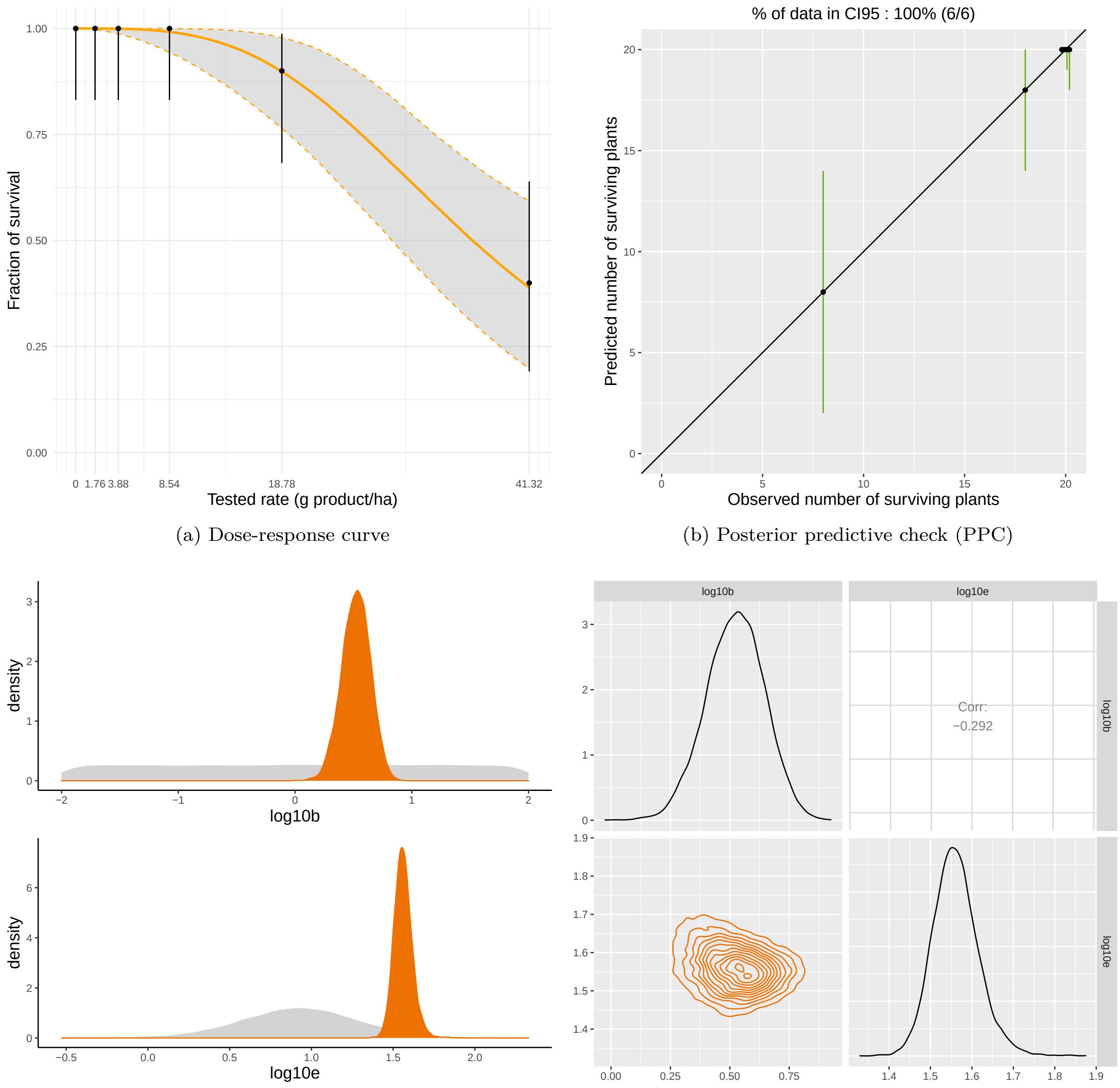

(a) Dose-response curve

(b) Posterior predictive check (PPC)

(c) Priors and posteriors

(d) Correlations between parameters

Figure 8: Dose-response curve (a), PPC (b), prior and posterior distributions (c) and correlations between parameters (d).

#### Data set: TRZAW_VV_survival

Table 9: Summary of parameter estimates (parameter d is set to 1) for TRZAW_VV_survival data set

| Parameter | median | Q2.5 | Q97.5 |
|---|---|---|---|
| b | 35.802 | 5.390 | 95.218 |
| e | 129.113 | 97.424 | 283.965 |

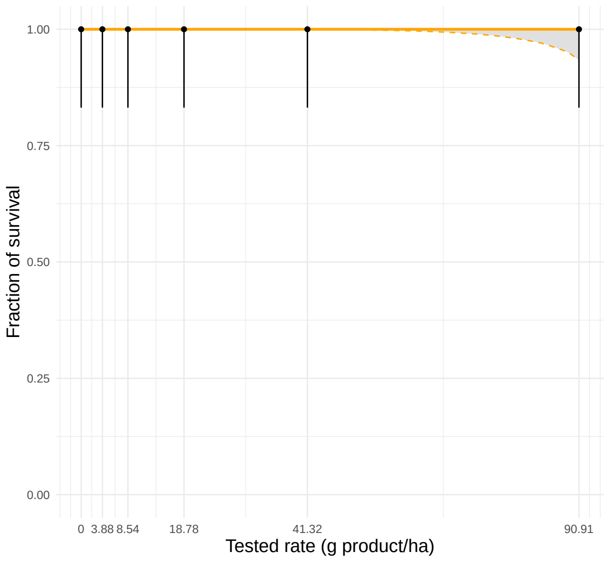

(a) Dose-response curve

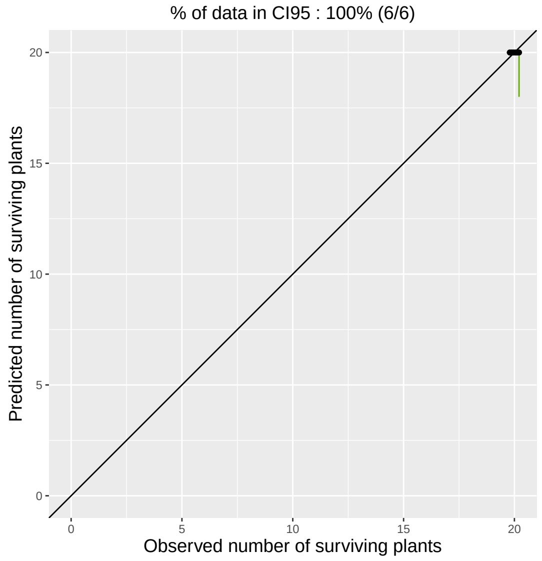

(b) Posterior predictive check (PPC)

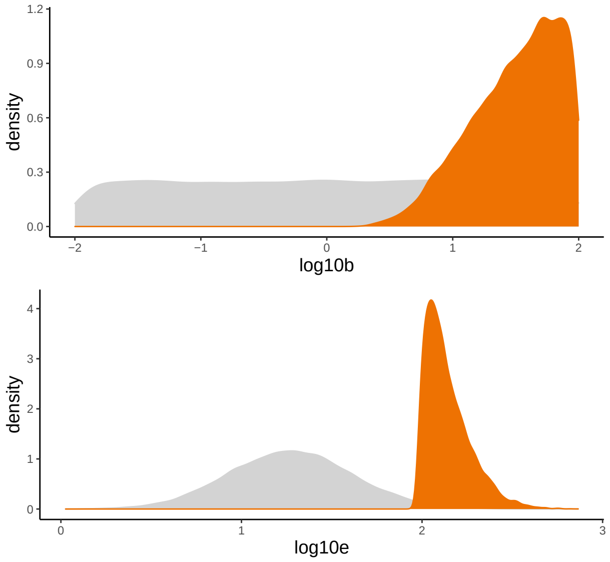

(c) Priors and posteriors

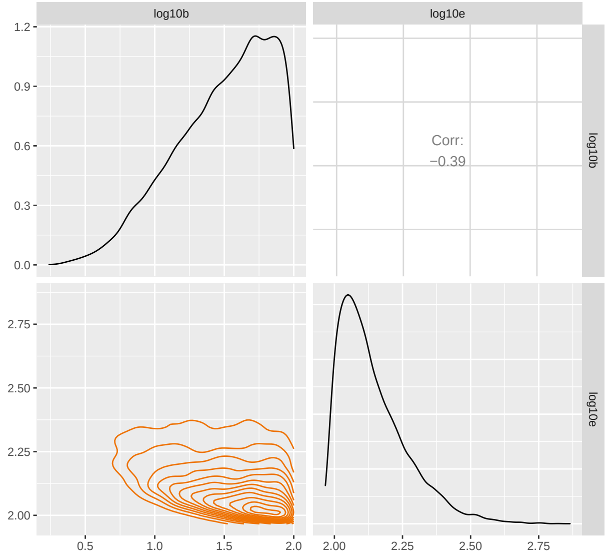

(d) Correlations between parameters

Figure 9: Dose-response curve (a), PPC (b), prior and posterior distributions (c) and correlations between parameters (d).

#### Data set: ZEAMA_VV_survival

Table 10: Summary of parameter estimates (parameter d is set to 1) for ZEAMA_VV_survival data set

| Parameter | median | Q2.5 | Q97.5 |
|---|---|---|---|
| b | 34.830 | 5.240 | 94.849 |
| e | 130.044 | 97.499 | 281.766 |

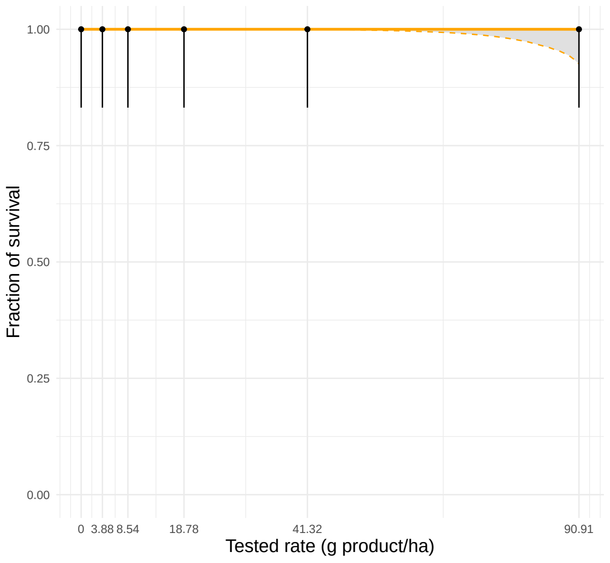

(a) Dose-response curve

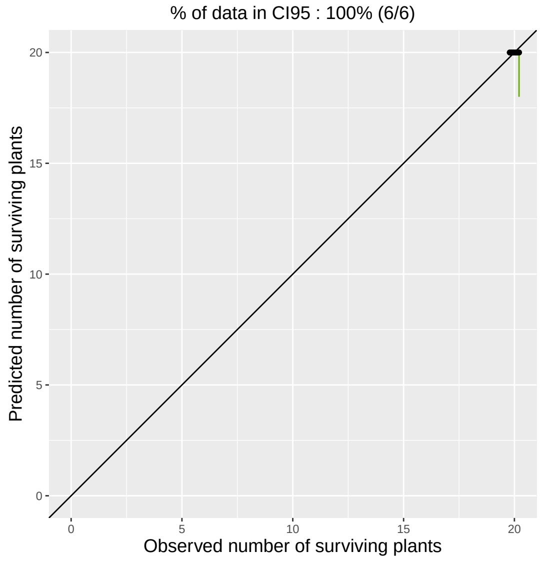

(b) Posterior predictive check (PPC)

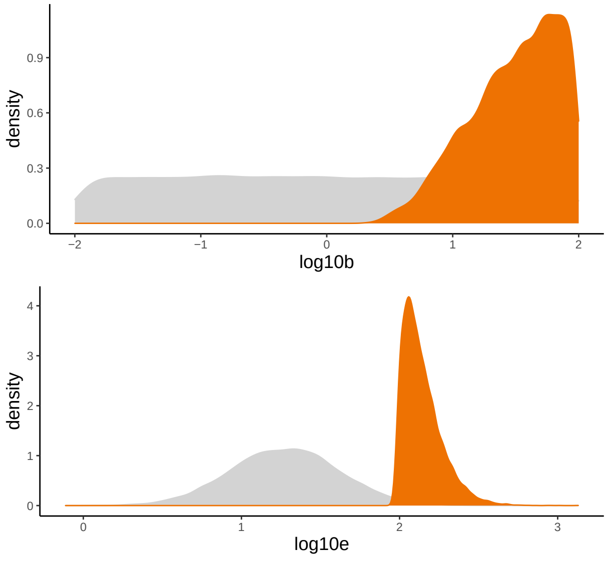

(c) Priors and posteriors

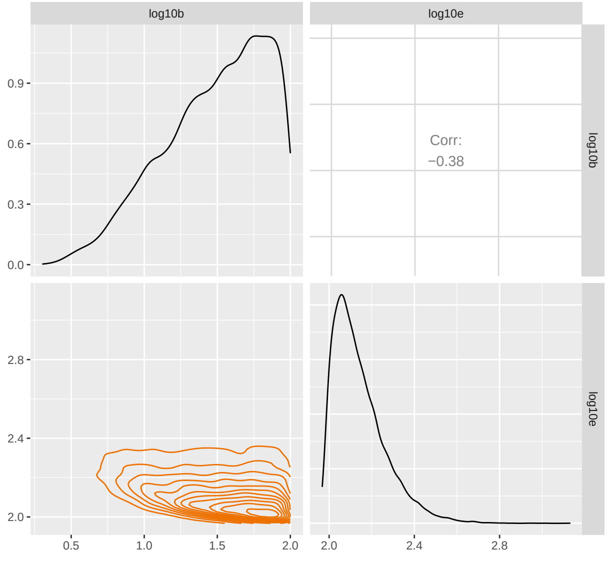

(d) Correlations between parameters

Figure 10: Dose-response curve (a), PPC (b), prior and posterior distributions (c) and correlations between parameters (d).
