## Supplementary material for "How to account for the uncertainty from standard toxicity tests in species sensitivity distributions: an example in non-target plants": Archive with reports on studies: report_VV_weight.pdf

### Dose-response analysis

Study 7

Vegetative Vigour test - shoot dry VV_weight endpoint

25 June 2020

Contact:

---

This is a report which provides results on all performed dose-response analyses for the shoot dry VV_weight endpoint of the Vegetative Vigour test for study 7.

---

#### Data set: ALLCE_VV_weight

Table 1: Summary of parameter estimates for ALLCE_VV_weight data set

| Parameter | median | Q2.5 | Q97.5 |
|---|---|---|---|
| b | 2.650 | 1.198 | 21.399 |
| d | 0.216 | 0.195 | 0.241 |
| e | 37.965 | 27.929 | 50.457 |
| sigma | 0.042 | 0.033 | 0.057 |

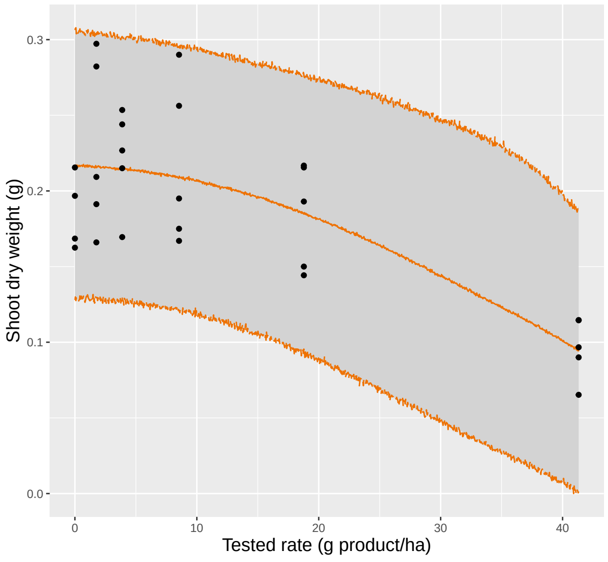

(a) Dose-response curve

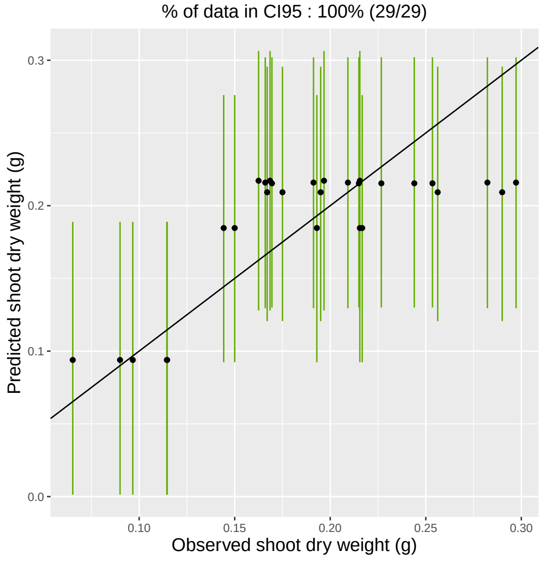

(b) Posterior predictive check (PPC)

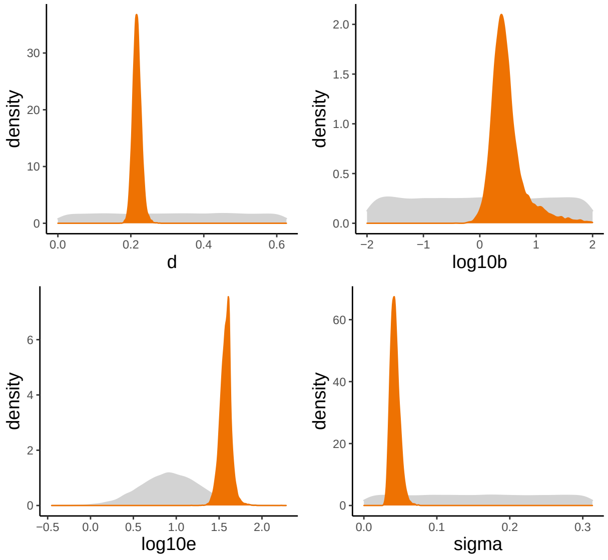

(c) Priors and posteriors

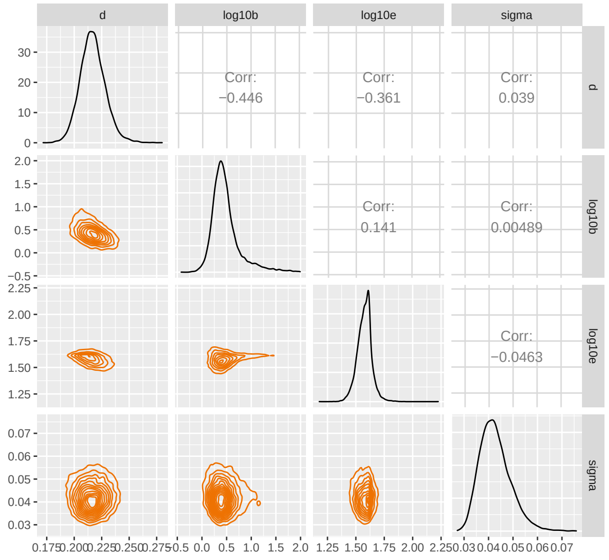

(d) Correlations between parameters

Figure 1: Dose-response curve (a), PPC (b), prior and posterior distributions (c) and correlations between parameters (d).

#### Data set: AVESA_VV_weight

Table 2: Summary of parameter estimates for AVESA_VV_weight data set

| Parameter | median | Q2.5 | Q97.5 |
|---|---|---|---|
| b | 3.202 | 1.922 | 9.033 |
| d | 1.706 | 1.573 | 1.842 |
| e | 29.872 | 23.641 | 38.034 |
| sigma | 0.274 | 0.211 | 0.376 |

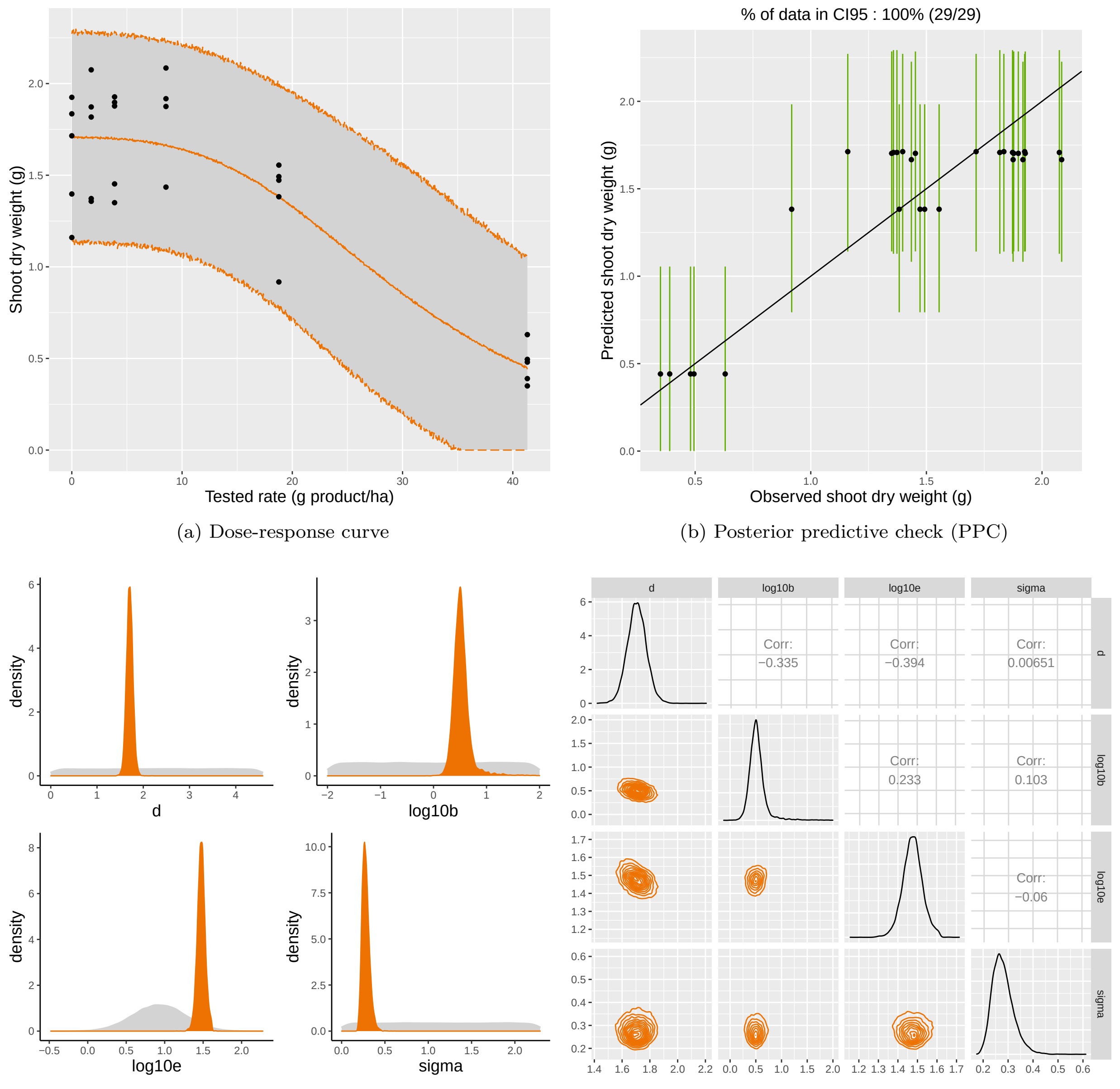

(a) Dose-response curve

(b) Posterior predictive check (PPC)

(c) Priors and posteriors

(d) Correlations between parameters

Figure 2: Dose-response curve (a), PPC (b), prior and posterior distributions (c) and correlations between parameters (d).

#### Data set: BEAVA_VV_weight

Table 3: Summary of parameter estimates for BEAVA_VV_weight data set

| Parameter | median | Q2.5 | Q97.5 |
|---|---|---|---|
| b | 1.865 | 1.316 | 2.827 |
| d | 2.255 | 2.052 | 2.472 |
| e | 1.756 | 1.406 | 2.216 |
| sigma | 0.412 | 0.344 | 0.510 |

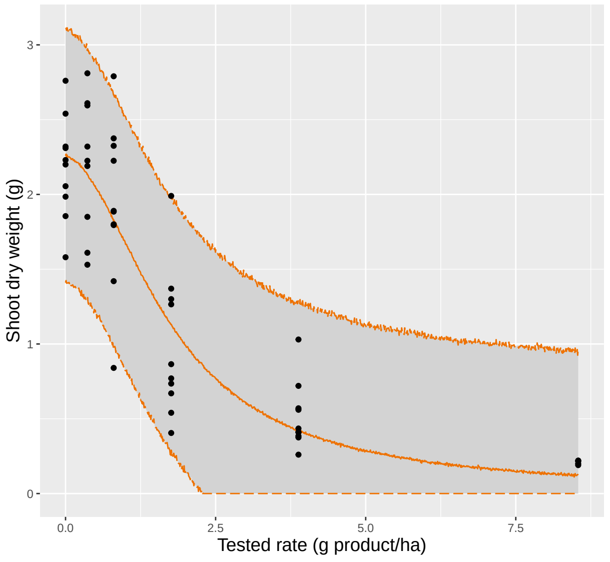

(a) Dose-response curve

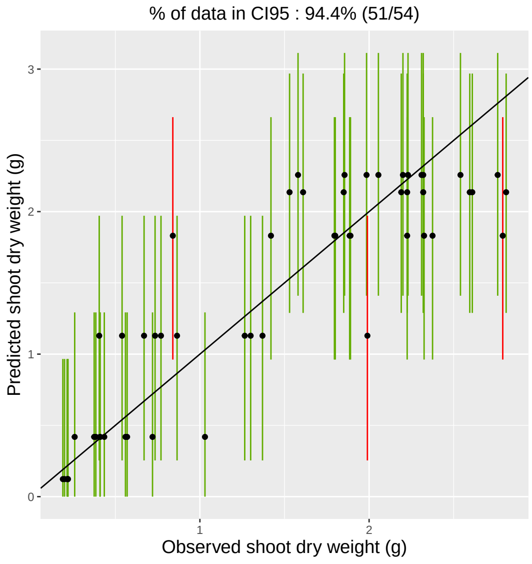

(b) Posterior predictive check (PPC)

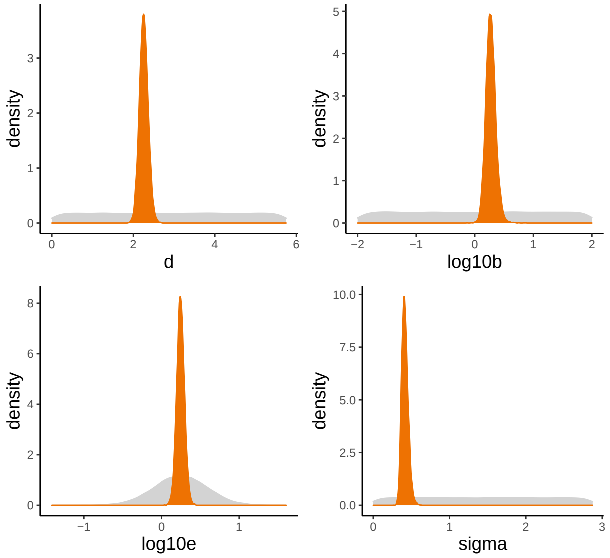

(c) Priors and posteriors

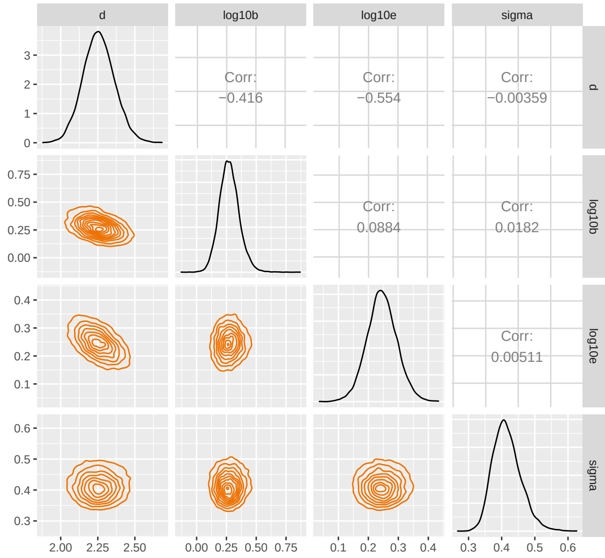

(d) Correlations between parameters

Figure 3: Dose-response curve (a), PPC (b), prior and posterior distributions (c) and correlations between parameters (d).

#### Data set: BRSNW_VV_weight

Table 4: Summary of parameter estimates for BRSNW_VV_weight data set

| Parameter | median | Q2.5 | Q97.5 |
|---|---|---|---|
| b | 3.021 | 2.167 | 7.829 |
| d | 4.810 | 4.557 | 5.070 |
| e | 4.066 | 3.600 | 4.594 |
| sigma | 0.692 | 0.582 | 0.846 |

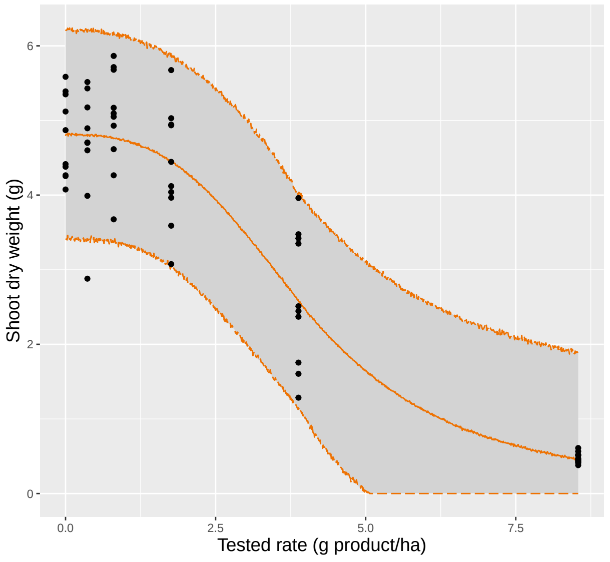

(a) Dose-response curve

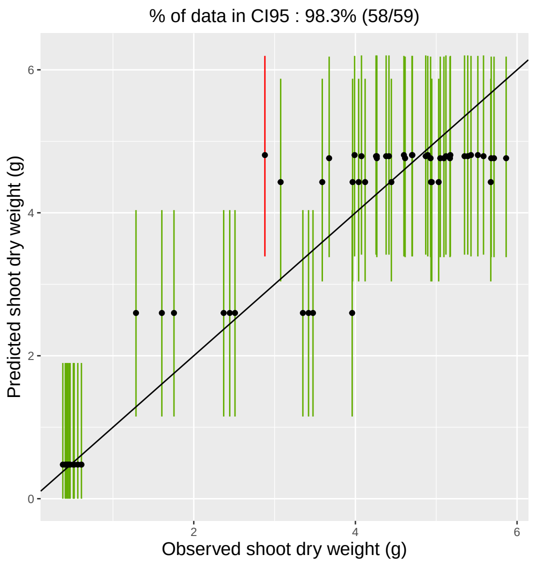

(b) Posterior predictive check (PPC)

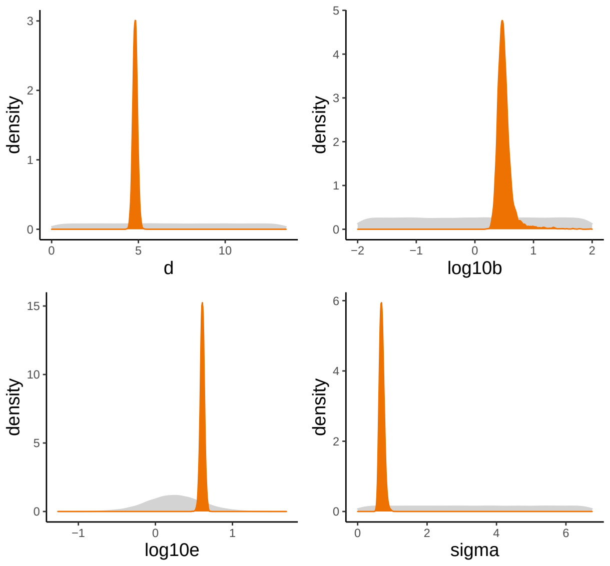

(c) Priors and posteriors

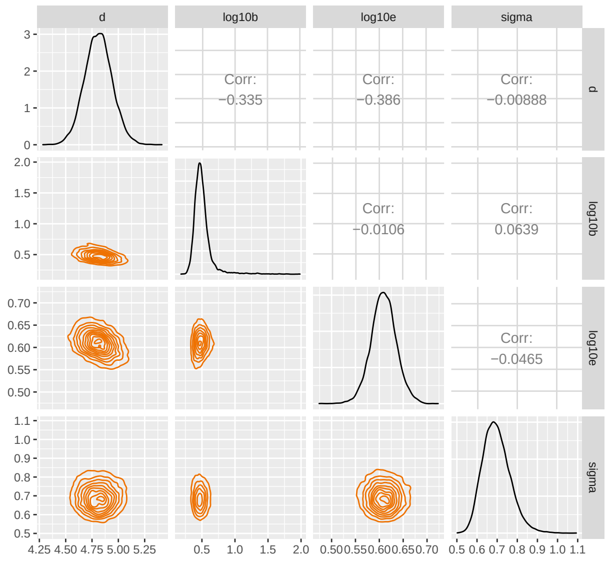

(d) Correlations between parameters

Figure 4: Dose-response curve (a), PPC (b), prior and posterior distributions (c) and correlations between parameters (d).

#### Data set: CUMSA_VV_weight

Table 5: Summary of parameter estimates for CUMSA_VV_weight data set

| Parameter | median | Q2.5 | Q97.5 |
|---|---|---|---|
| b | 0.822 | 0.632 | 1.052 |
| d | 9.891 | 9.105 | 10.687 |
| e | 9.378 | 6.944 | 12.906 |
| sigma | 1.250 | 1.047 | 1.541 |

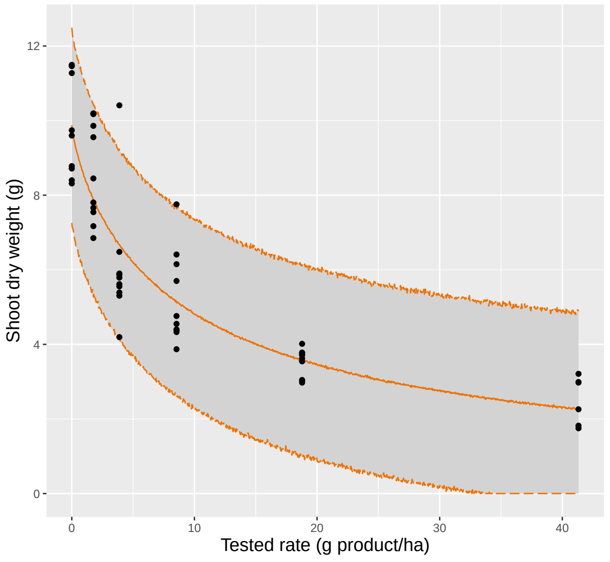

(a) Dose-response curve

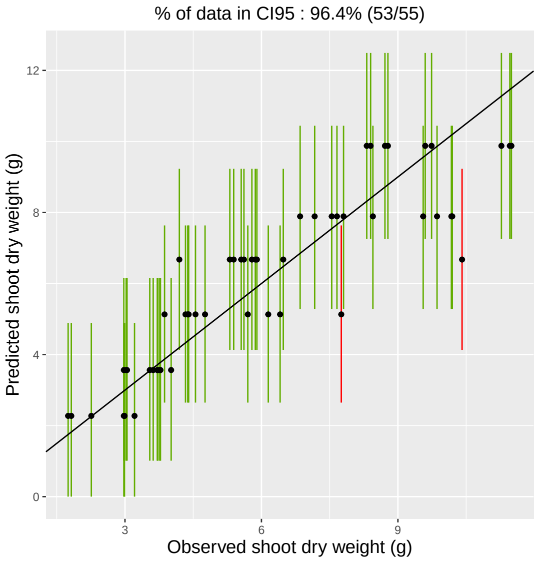

(b) Posterior predictive check (PPC)

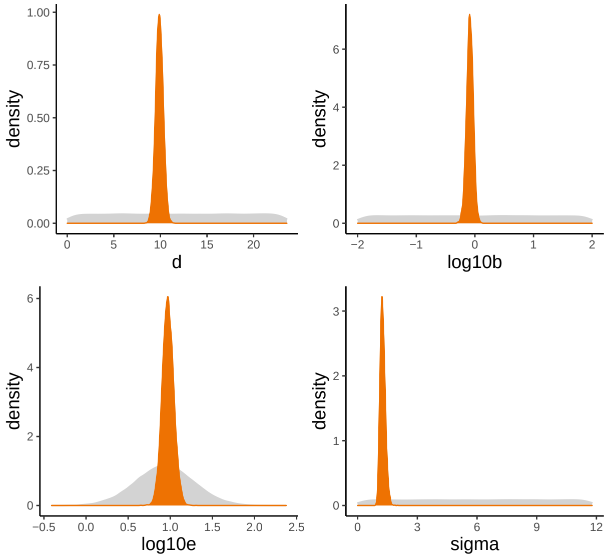

(c) Priors and posteriors

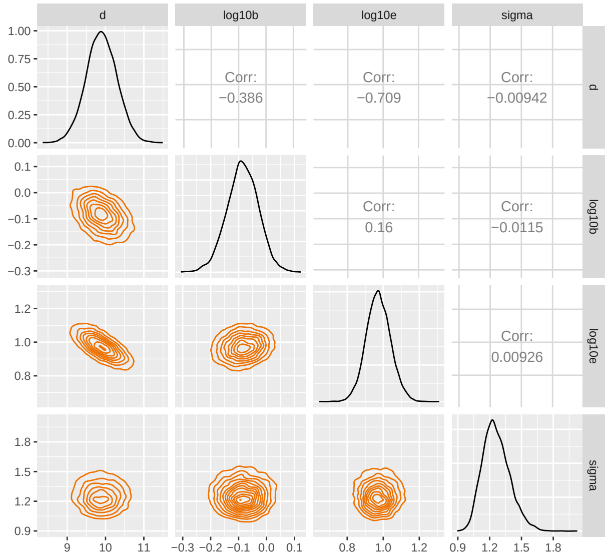

(d) Correlations between parameters

Figure 5: Dose-response curve (a), PPC (b), prior and posterior distributions (c) and correlations between parameters (d).

#### Data set: GLXMA_VV_weight

Table 6: Summary of parameter estimates for GLXMA_VV_weight data set

| Parameter | median | Q2.5 | Q97.5 |
|---|---|---|---|
| b | 0.902 | 0.730 | 1.102 |
| d | 3.071 | 2.863 | 3.280 |
| e | 8.119 | 6.373 | 10.194 |
| sigma | 0.315 | 0.264 | 0.383 |

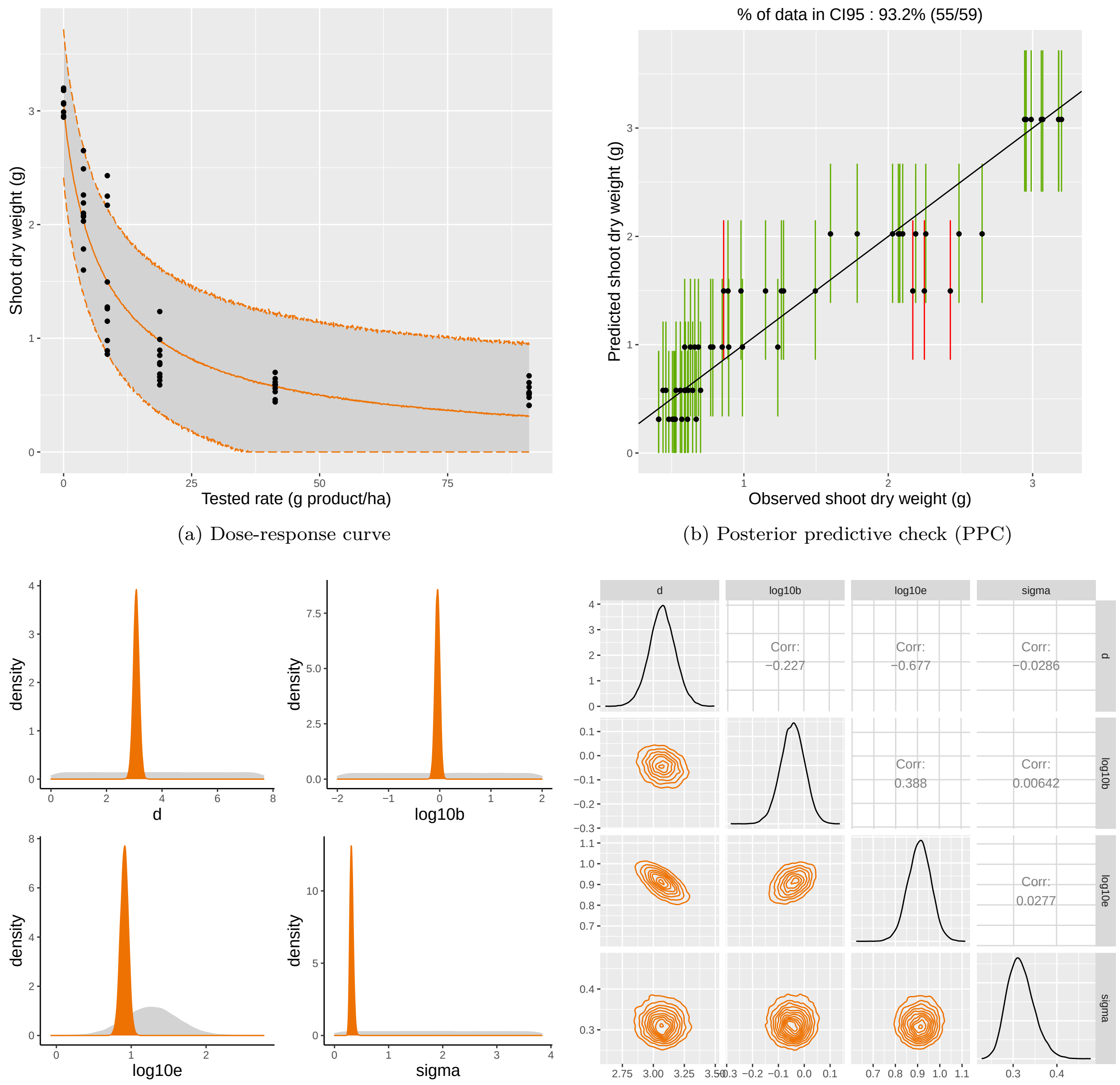

(a) Dose-response curve

(b) Posterior predictive check (PPC)

(c) Priors and posteriors

(d) Correlations between parameters

Figure 6: Dose-response curve (a), PPC (b), prior and posterior distributions (c) and correlations between parameters (d).

#### Data set: HELAN_VV_weight

Table 7: Summary of parameter estimates for HELAN_VV_weight data set

| Parameter | median | Q2.5 | Q97.5 |
|---|---|---|---|
| b | 2.634 | 1.756 | 4.070 |
| d | 1.742 | 1.585 | 1.907 |
| e | 5.172 | 4.319 | 6.280 |
| sigma | 0.299 | 0.238 | 0.389 |

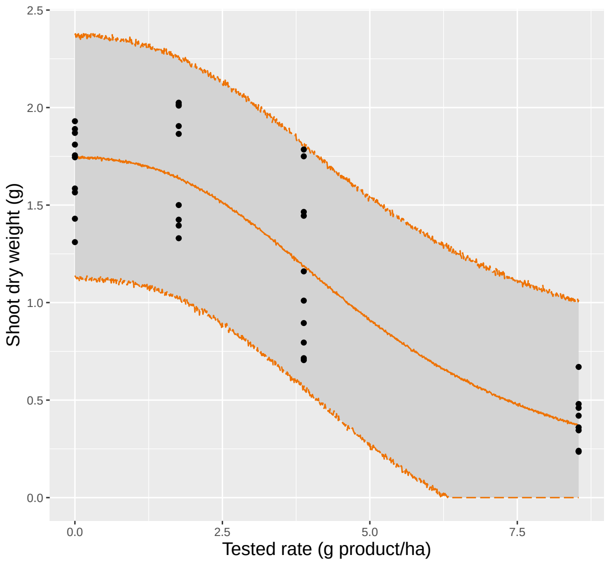

(a) Dose-response curve

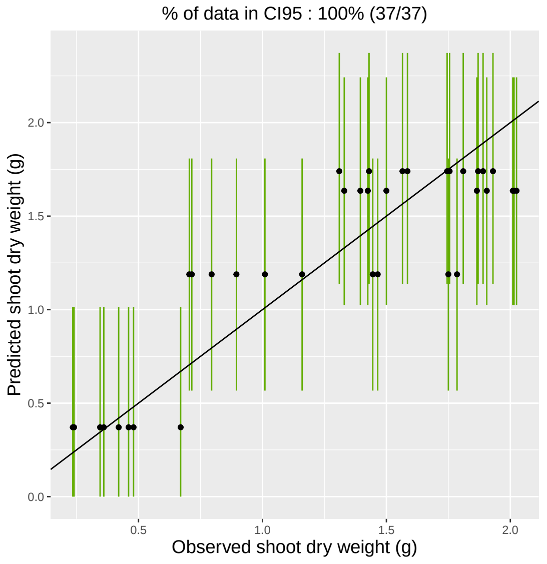

(b) Posterior predictive check (PPC)

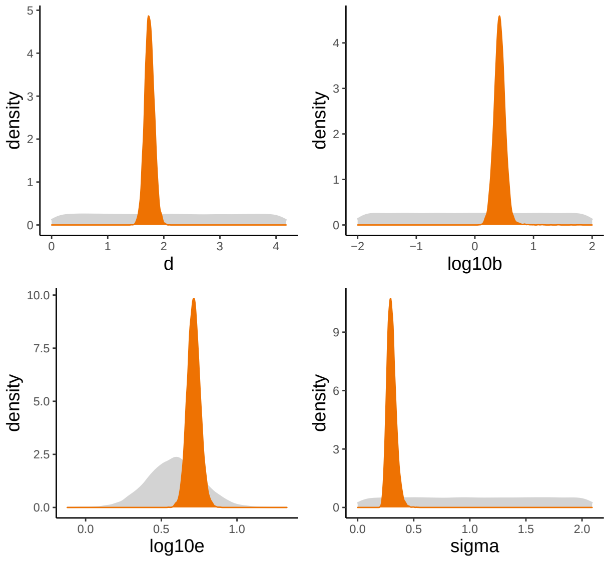

(c) Priors and posteriors

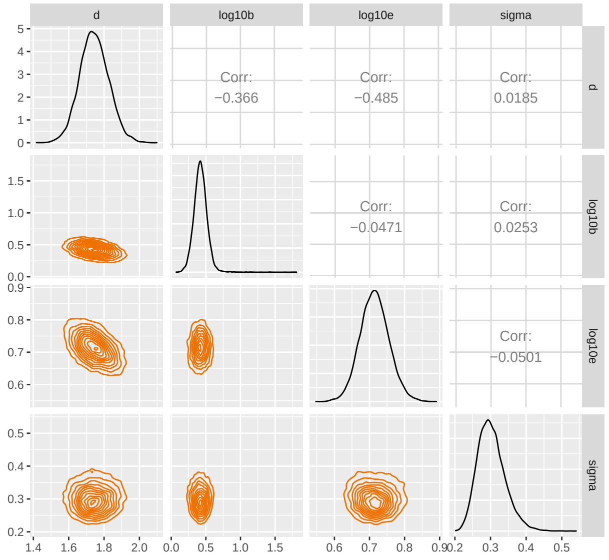

(d) Correlations between parameters

Figure 7: Dose-response curve (a), PPC (b), prior and posterior distributions (c) and correlations between parameters (d).

#### Data set: LYPES_VV_weight

Table 8: Summary of parameter estimates for LYPES_VV_weight data set

| Parameter | median | Q2.5 | Q97.5 |
|---|---|---|---|
| b | 2.078 | 1.676 | 2.642 |
| d | 4.098 | 3.850 | 4.358 |
| e | 4.989 | 4.388 | 5.634 |
| sigma | 0.412 | 0.344 | 0.508 |

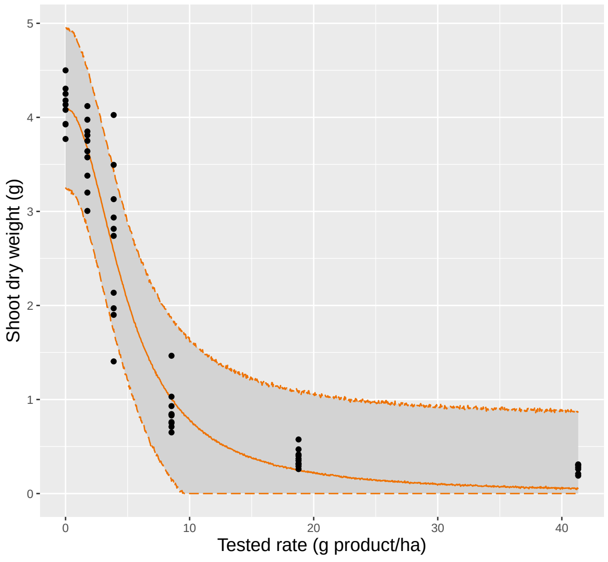

(a) Dose-response curve

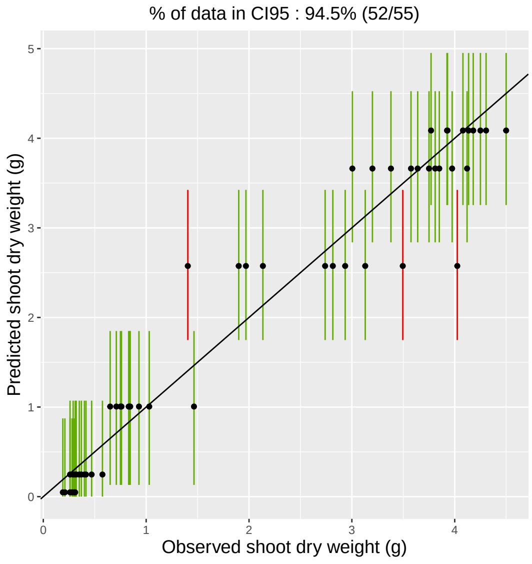

(b) Posterior predictive check (PPC)

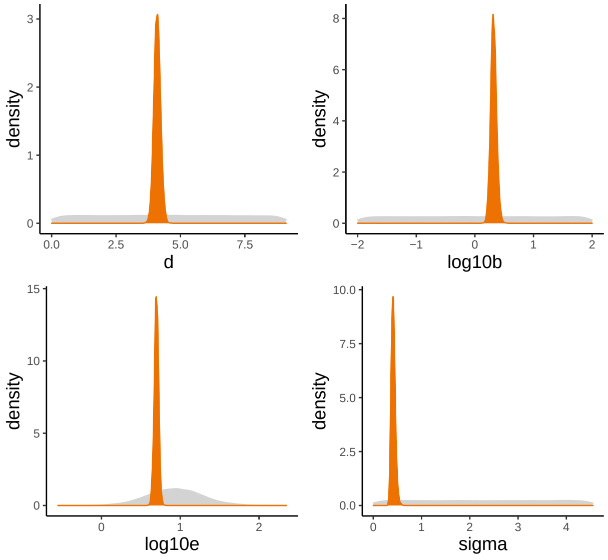

(c) Priors and posteriors

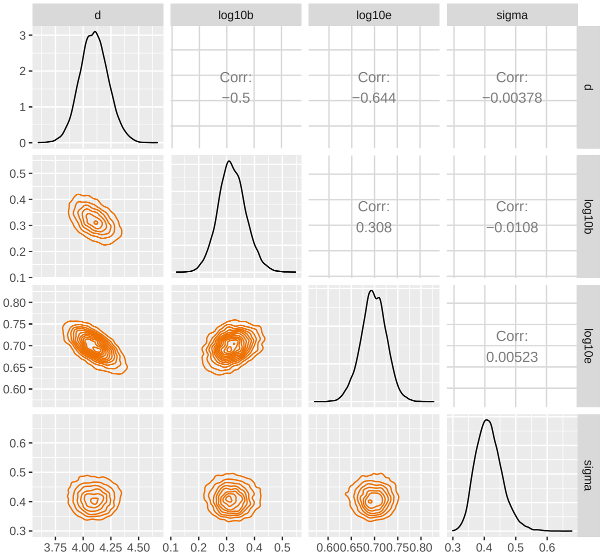

(d) Correlations between parameters

Figure 8: Dose-response curve (a), PPC (b), prior and posterior distributions (c) and correlations between parameters (d).

#### Data set: TRZAW_VV_weight

Table 9: Summary of parameter estimates for TRZAW_VV_weight data set

| Parameter | median | Q2.5 | Q97.5 |
|---|---|---|---|
| b | 14.210 | 2.099 | 89.264 |
| d | 0.624 | 0.604 | 0.645 |
| e | 123.307 | 94.553 | 290.279 |
| sigma | 0.047 | 0.036 | 0.063 |

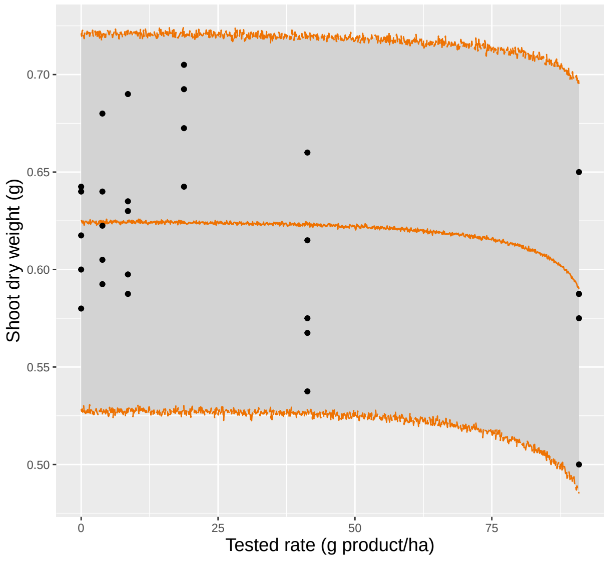

(a) Dose-response curve

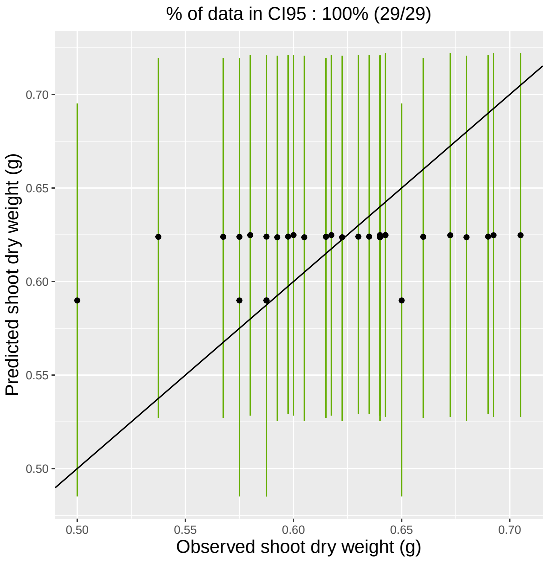

(b) Posterior predictive check (PPC)

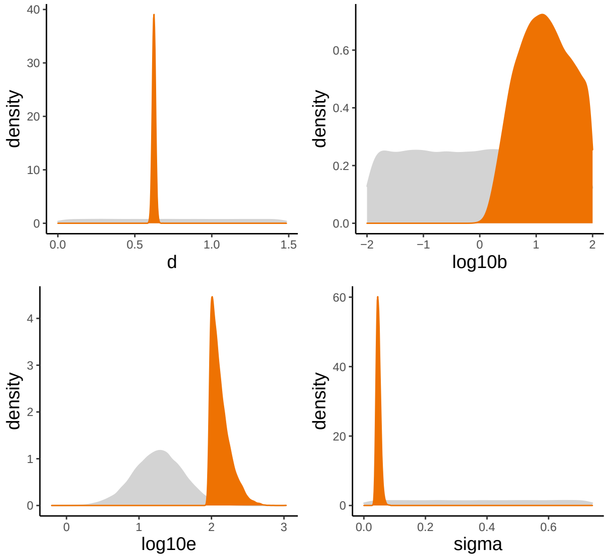

(c) Priors and posteriors

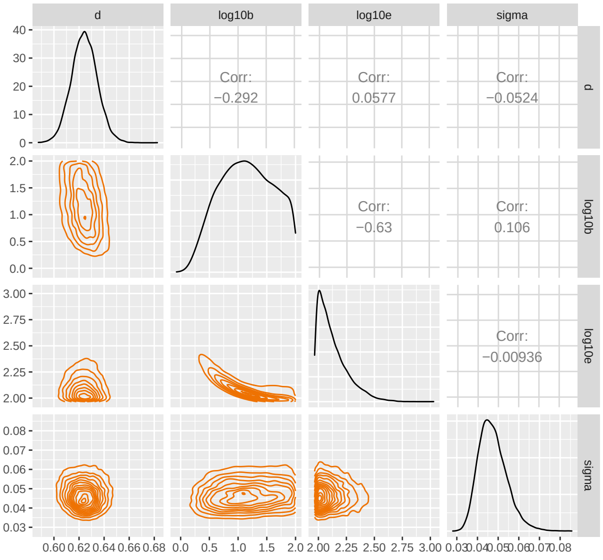

(d) Correlations between parameters

Figure 9: Dose-response curve (a), PPC (b), prior and posterior distributions (c) and correlations between parameters (d).

### Data set: ZEAMA_VV_weight

Table 10: Summary of parameter estimates for ZEAMA_VV_weight data set

| Parameter | median | Q2.5 | Q97.5 |
|---|---|---|---|
| b | 2.644 | 1.659 | 12.564 |
| d | 4.669 | 4.453 | 4.900 |
| e | 94.766 | 83.962 | 112.980 |
| sigma | 0.634 | 0.528 | 0.777 |

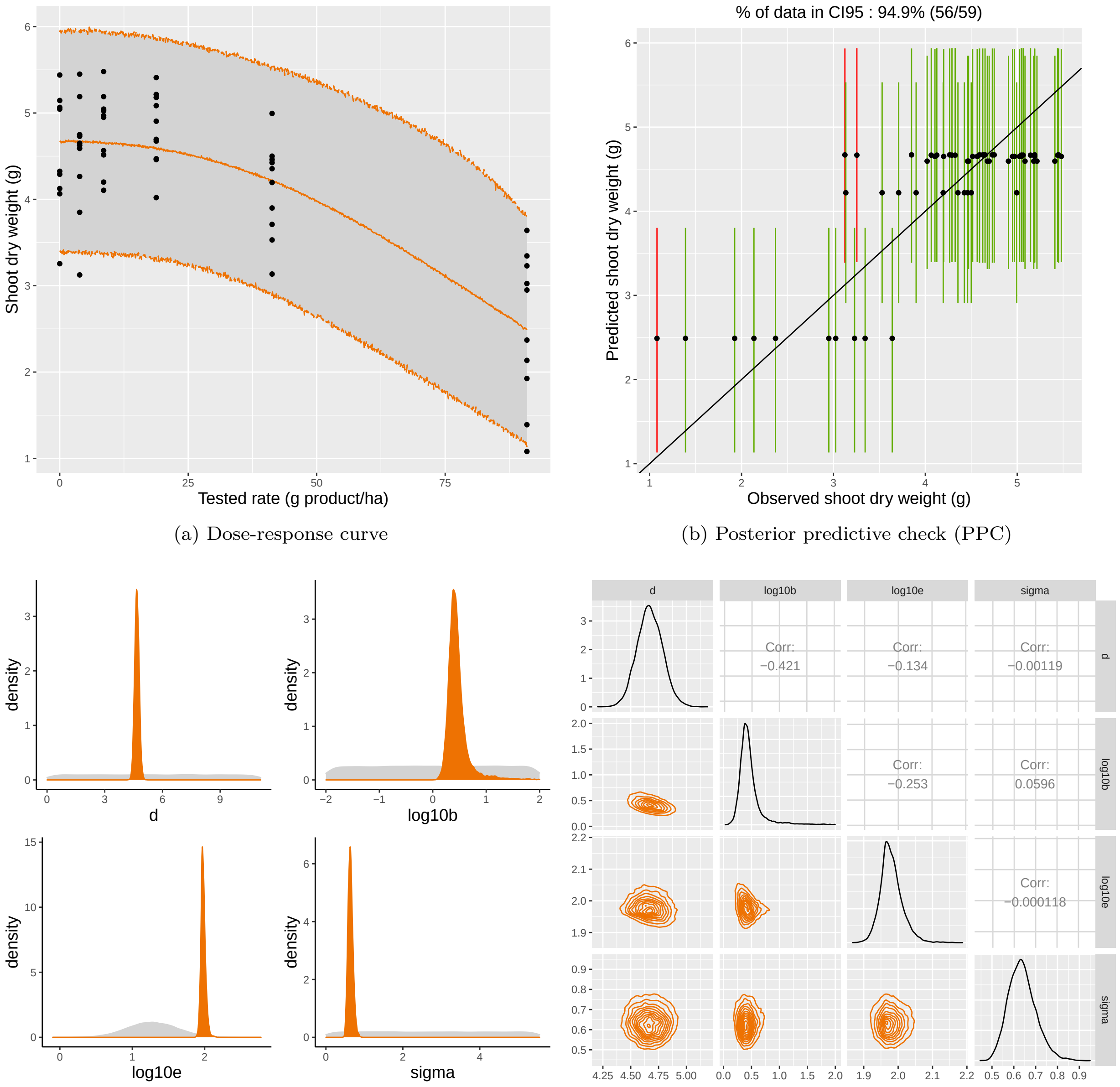

(a) Dose-response curve

(b) Posterior predictive check (PPC)

(c) Priors and posteriors

(d) Correlations between parameters

Figure 10: Dose-response curve (a), PPC (b), prior and posterior distributions (c) and correlations between parameters (d).
