## Supplemental Table 1 for "How to account for the uncertainty from standard toxicity tests in species sensitivity distributions: an example in non-target plants"

Sandrine Charles Dan Wu Virginie Ducrot

29 June 2020

Contact:

S1 Table: Corresponding species and common names to the EPPO code

| EPPO code | Species name | Common name |
|---|---|---|
| ALLCE | Allium cepa | Onion |
| AVESA | Avena sativa | Oat |
| BEAVA | Beta vulgaris | Sugar beet |
| BRSNW | Brassica napus winter | Oilseed rape |
| BRSOL | Brassica oleracea | Cabbage |
| CUMSA | Cucumis sativus | Cucumber |
| FAGES | Fagopyrum esculentum | Buckwheat |
| GLXMA | Glycine max | Soybean |
| HELAN | Helianthus annuus | Sunflower |
| LOLPE | Lolium perenne | Perennial ryegrass |
| LYPES | Solanum lycopersicum | Tomato |
| RAPSR | Raphanus sativus | Radish |
| TRZAW | Triticum aestivum winter | Winter wheat |
| ZEAMA | Zea mays subs. Amylacea | Corn |
