## Supplemental Table 2 for "How to account for the uncertainty from standard toxicity tests in species sensitivity distributions: an example in non-target plants"

Sandrine Charles Dan Wu Virginie Ducrot

29 June 2020

Contact:

S2 Table: Formulation of active substances in seven products

| Product number | Chemical family | Mode of action |
|---|---|---|
| 1 | Diphenylethers + Oxyacetamides + Pyridinecarboxamides | Inhibition of carotenoid synthesis + Inhibition of cell division |
| 2 | Pyridinecarboxamides | Inhibition of carotenoid synthesis |
| 3 | Benzofuran + Uracil + Carbamate | Inhibition of lipid synthesis + Inhibition of photosynthesis at photosystem II |
| 4 | Sulfonylureas + Phenoxy + Sulfonly-amino-carbonyl-triazolinones | Synthetic auxin + Inhibition of acetolactate synthase |
| 5 | Triazines | Inhibition of cellulose synthesis |
| 6 | Sulfonylureas + Sulfonly-amino-carbonyl-triazolinones | Inhibition of acetolactate synthase |
| 7 | Sulfonylureas + Sulfonly-amino-carbonyl-triazolinones | Inhibition of acetolactate synthase |

*Note:* The proportion of active substances is different between products 6 and 7.
